# Aging and Obesity Prime the Methylome and Transcriptome of Adipose Stem Cells for Disease and Dysfunction

**DOI:** 10.1101/2022.09.26.509507

**Authors:** Shaojun Xie, Sulbha Choudhari, Chia-Lung Wu, Karen Abramson, David Corcoran, Simon G. Gregory, Jyothi Thimmapurum, Farshid Guilak, Dianne Little

## Abstract

The epigenome of stem cells occupies a critical interface between genes and environment, serving to regulate expression through modification by intrinsic and extrinsic factors. We hypothesized that aging and obesity, which represent major risk factors for a variety of diseases, synergistically modify the epigenome of adult adipose stem cells (ASCs). Using integrated RNA- and targeted bisulfite-sequencing in murine ASCs from lean and obese mice at 5- and 12- months of age, we identified global DNA hypomethylation with either aging or obesity, and a synergistic effect of aging combined with obesity. The transcriptome of ASCs in lean mice was relatively stable to the effects of age, but this was not true in obese mice. Functional pathway analyses identified a subset of genes with critical roles in progenitors and in diseases of obesity and aging. Specifically, *Mapt, Nr3c2, App, and Ctnnb1* emerged as potential hypomethylated upstream regulators in both aging and obesity (AL vs YL and AO vs YO), and *App*, *Ctnnb1, Hipk2, Id2,* and *Tp53* exhibited additional effects of aging in obese animals. Further, *Foxo3* and *Ccnd1* were potential hypermethylated upstream regulators of healthy aging (AL vs YL), and of the effects of obesity in young animals (YO vs YL), suggesting that these factors could play a role in accelerated aging with obesity. Finally, we identified candidate driver genes that appeared recurrently in all analyses and comparisons undertaken. Further mechanistic studies are needed to validate the roles of these genes capable of priming ASCs for dysfunction in aging- and obesity-associated pathologies.

## Introduction

The epigenome occupies a critical interface between the response of genes to the environment, serving to regulate expression through coordinated modifications by intrinsic and extrinsic factors. In this regard, adult stem cells have the potential to acquire epigenetic memory of these environmental factors over a lifetime. Of the many factors influencing genetic modification, aging and obesity appear to have significant contributions to the risk of multiple pathologic conditions. Aging is a ubiquitous intrinsic factor influencing epigenetic changes.^1; 2^ Typically, aging in both somatic tissues and adult stem cells is associated with CpG island hypermethylation and global DNA hypomethylation.^2–6^ Age related hypomethylation at specific sites in differentiated tissues is associated with disease, but the extent to which DNA methylation change is a consequence of aging or whether it contributes to aging is unknown.^2^

Superimposed on the inevitable progression of age, over 40% of Americans are obese; therefore, diet-induced obesity is a common extrinsic factor that impacts the epigenome. Consequently, the effects of obesity, high-fat diet, and increased adiposity on DNA methylation are being explored in a variety of tissues relevant to human health. These previous studies have found maternal and offspring high-fat diet increases epigenetic ‘age’ in the mouse liver.^6^ Additionally, diabetic and obese phenotypes and clinical traits have identified characteristic differential methylation patterns in both mouse and human adipocytes, in adipose tissue, and in liver.^7; 8^ Differential methylation of periprostatic adipose tissue containing adult stem cells in patients with prostate cancer likely contributes to impaired lipid metabolism and immune dysregulation.^9^ Despite these studies, the characteristic effects of obesity on the epigenomes of progenitor cells are not well understood even though there are widespread phenotypic and molecular effects of obesity on these cells.^10; 11^ Taken together, these findings suggest that obesity may in fact represent a disease of the body’s stem cells.^12^ These effects include reduced ‘stemness’, impaired differentiation, an increase in expression of pro-inflammatory genes, impaired immunomodulation and anti-inflammatory roles, dysregulated metabolism, and altered motility.^10; 11^ Obesity also induces dysregulation of cell-cell cross-talk, both between different progenitor cell populations in different tissue locations,^13^ and between progenitor cell populations and entirely difference cell types, including solid tumors.^14^ In this regard, adipose-derived stem cells (ASCs) contribute to regulation of adipose tissue homeostasis, and thus epigenetic modifications of these cells could have significant long-term and intergenerational consequences on factors such as body weight, insulin resistance, and overall health.^15^ Furthermore, ASCs and other adult stem cells are attractive progenitor cell sources for regenerative medicine and tissue engineering purposes. However, the regenerative potential of progenitor cells identified in pre-clinical studies has not been matched by their therapeutic efficacy in humans.^16^ The reasons for this lack of therapeutic efficacy are multi-factorial. While several studies have examined the role of dietary intervention to slow epigenetic aging in somatic tissues, the role of high-fat diet and obesity in the epigenetic and transcriptomic dysfunction of progenitor cells is less well understood. In addition, the interactions between age and modifiable factors that could impact the regenerative potential of autologous ASCs and other adult progenitor cells have not been studied.

Here, we hypothesized that age-related disruption of both the methylome and transcriptome of ASCs would be exacerbated by high-fat diet induced obesity. Our aims were 1) to characterize the methylome and transcriptome of murine ASCs harvested from lean, obese, young and old mice, 2) to identify putative driver genes for epigenetic and transcriptomic disruption occurring with aging and obesity, 3) identify candidate functional pathways dysregulated in aging ASCs in diet-induced obesity, and 4) to identify candidate biomarkers for obesity and aging in murine ASCs, as a first step to identification of the ‘healthy’ ASC phenotype.

## Materials and Methods

### Animals, Tissue and Cell Isolation

All procedures were approved by the Duke University IACUC. Male C57BL6/J mice (n=12/group) were purchased at 19 weeks of age from the diet-induced obesity colony at The Jackson Laboratory. Beginning at 6 weeks of age, control normal bodyweight ‘lean (L)’ mice were fed control lean diet (10% kcal fat, D12450B, Research Diets Inc.), and ‘obese (O)’ mice were fed lard-based high-fat diet (60 %kcal fat, D12492, Research Diets, Inc.). Mice were group housed (4-5 animals/cage) under standard light cycle and laboratory animal environmental conditions. ‘Young (Y)’ mice were acclimated for 1 week before euthanasia at 20 weeks of age, whereas ‘Aged (A)’ mice were sacrificed at 52 weeks of age. The original intent of the study was to age the mice until 2-years of age, but increased mortality in the ‘Aged Obese (AO)’ mice was noted beginning at age 10 months, such that by 51 weeks of age only n=8 remained, and therefore, aged mice were sacrificed at 52 weeks of age in order to maintain statistical power of our study design. Animals were euthanized by cervical dislocation following isoflurane anesthesia. Then, the subcutaneous inguinal fat pad was isolated bilaterally and digested at 37°C for 1-1.5 hours in 0.2 % collagenase type I (Worthington). Murine ASCs were isolated from digested fat, enriched for Sca-1^+^, CD34^+^ CD31^-^, CD45^-^, Ter119^-^ markers using Fluorescent-activated cell-soring (FACS) and expanded using previously described methods.^17^ Briefly, sorted cells were cultured under hypoxic conditions (2% O_2_, 5% CO_2_) in α-Modified Eagle’s medium, 20% fetal bovine serum and 1% penicillin/streptomycin/amphotericin B at 37°C. Media was replaced every 3 days, and cells were trypsinized in 0.25% trypsin-EDTA at 90% confluence. Cells were harvested and frozen at Passage 2 in aliquots of 1-2x 10^6^ murine ASCs for independent DNA and RNA extraction. Four groups of murine ASCs were prepared for the current study: YL (young lean), YO (young obese), AL (aged lean) and AO (aged obese).

### DNA and RNA extraction and library preparation

Following DNA extraction using Qiagen’s All Prep kit (Qiagen catalogue # 80284) standard methods, libraries were prepared using the SureSelect Methyl-Seq Library Prep Kit ILM (Agilent Catalog # 5500-0128) included the indices for pooled sequencing, and SureSelectXT Mouse Methyl-Seq Reagent Kit (Agilent Catalog # 931052) . Representative library preparation quality control data are provided in (**Supplementary Methods Figure 1**). Briefly, after shearing and post-end repair, DNA fragment length was evaluated using the Agilent 2100 BioAnalyzer DNA1000chip or Agilent 4200 TapeStation High Sensitivity D1000 ScreenTape to confirm a DNA fragment size peak between 125-175bp. Post-ligation, DNA fragment size was confirmed in the same way to be 200-300bp, with adequate yield (>350ng) of adapter-ligated DNA (mean = 516±80ng). Quality assessment and quantification of indexed DNA samples was performed using the 2100 BioAnalyzer or TapeStation to verify fragment peak size of 250-300bp. Following extraction, RNA was evaluated by Nanodrop and quantified by Ribogreen. Samples were only used if RIN>7 and if total yield exceeded 1µg.

The David H. Murdock Research Institute Genomics core facility performed 100bp paired-end sequencing, throughput and quality control in accordance with the ENCODE guidelines, using an entire 8-lane flowcell dedicated to the samples, with each sample run over two separate lanes on a HiSeq2500. Data were trimmed and demultiplexed according to the standard Illumina sequencing pipeline. Briefly, samples were separated based on Illumina barcodes, and only sequencing reads that had 100% similarity match for the barcodes were used for downstream analysis.

### Data Processing and Annotation

Both Methyl-Seq and RNA-Seq data were processed using the TrimGalore (version 0.4.1) toolkit ^18^ which employs Cutadapt (version 1.8.3)^19^ to trim low quality bases and Illumina sequencing adapters from the 3’ end of the reads. Only read pairs where each read of the mate-pair was 20nt or longer after trimming were kept for further analysis. Windows were retained if they had at least 10 reads in a single sample.

### Methyl-Seq Data Analysis

Reads were mapped to the GRCm38 version of the mouse genome using the Bismark (version 0.17.0)^20^ bisulfite converted sequence read aligner. Bismark utilizes the Bowtie2 (version 2.2.4) ^21^ alignment algorithm as part of its processing pipeline. The *MarkDuplicatesWithMateCigar* function from the Picard Toolkit (version 2.4.1)^22^ was used to eliminate amplification artifacts.

### Identification of differentially methylated cytosines (DMC) and differentially methylated windows (DMW)

The MethylKit Bioconductor (version 1.2.0)^23; 24^package was used to process, normalize, and analyze differences between samples. Differential methylation was identified both on the base level for individual CpG sites (differentially methylated cytosines; DMC) as well as over 200 base pair sliding windows (differentially methylated windows; DMW) in tileMethylCounts (R version 3.3.0)^25^ with a 100 base pair step size. The input was reduced to only those regions that had reads that overlapped with the target regions in the SureSelect capture kit. CpG sites and sliding windows were annotated to their nearest gene according to the mm10 GRCm38v73 version of the mouse transcriptome.^26^ The false discovery rate was calculated to control for multiple hypothesis testing for each comparison. For calculating differential methylation, the following parameters were used: overdispersion=”MN”, test=”Chisq”. For A vs Y comparisons of age, diet (O and L) were included as covariates; for O vs L comparisons of diet, age (A and Y) were included as covariates in the model. Default parameters were used for all other packages.

### Multivariate analysis Partial Least Squares Discriminant Analysis (PLS-DA)

Partial least squares (PLS) is a versatile algorithm which can be used to predict either continuous or discrete/categorical variables. PLS-DA was applied to the data for multivariate analysis of the Methyl-seq data. Variable Importance in Projection (VIP) scores estimate the importance of each variable in the projection used in a PLS model and are often used for variable selection. Variables with VIP score above 1 were used as potential biomarkers of lean, obese, young, or aged status. Based on the number of variables with VIP score above 1, candidates were additionally filtered and analyzed based on a VIP score above 2. R package ropls^27^ was used for the analysis. Specifically, matrices of methylation levels for each sample were generated by the function *percMethylation* of methylKit and were used as X variables. Diet and/or age were provided as Y responses. One additional advantage of PLS-DA is the lack of dependence on p values; thus, VIP scores provide a means to rank the relative importance of potential biomarkers.

### Distribution of DMW and DMC within CpG islands and Genic Regions

An in-house script was used to generate the input file for *annotatr* from MethylKit results, then the R function *read_regions in annotatr* ^28^ was used to read into the input data. The R function *plot_categorical in annotatr* ^28^ was used to plot the distribution of DMW and DMC. Distributions of DMW and DMC were evaluated based on CpG annotation and genic annotation. CpG islands as the basis for all CpG annotations were determined using *AnnotationHub* ^29^ mouse package. CpG shores were defined as 2Kb upstream/downstream from the ends of the CpG islands. CpG shelves were defined as an additional 2Kb upstream/downstream of the farthest upstream/downstream limits of the CpG shores, excluding the CpG islands and CpG shores. The remaining genomic regions were annotated as inter-CpG islands (Inter-CGI) (http://bioconductor.org/packages/release/bioc/vignettes/annotatr/inst/doc/annotatr-vignette.html).

Murine gene region annotations were determined by functions from *GenomicFeatures* and data from the *TxDb.Mmusculus.UCSC.mm10* and *org.Mm.eg.db* packages in Bioconductor. Genic annotations included 1-5Kb upstream of the transcription start site (TSS), the promoter (< 1Kb upstream of the TSS), 5’UTR, coding sequence (CDS, exons, introns, and intergenic regions (http://bioconductor.org/packages/release/bioc/vignettes/annotatr/inst/doc/annotatr-vignette.html). Genic annotations were generated using GTF file for Mus_musculus.GRCm38.73 from ENSEMBL database using custom Python script.

### RNA-Seq Data Analysis

Reads were mapped to the GRCm38v73 version of the mouse genome and transcriptome ^26^using the STAR RNA-seq alignment tool.^30^ Reads were kept for subsequent analysis if they mapped to a single genomic location. Read counts for each gene feature for each animal were compiled using the HTSeq tool (Version 0.7.0)^31; 32^. The parameters used for HTSeq-Count were set for unstranded RNA-Seq. Counts from all biological replicates were merged into one file using custom Perl scripts to generate a merged read count matrix for all samples. The merged count matrix was used for downstream differential gene expression analysis, and the quality of the count matrix was verified by determining basic statistics including data range and matrix size prior to downstream evaluation. Genes that had at least 5 or more counts in at least 50% of the animals from each group were retained for subsequent analysis. Normalization, gene-wise dispersion, and differential gene expression (DGE) analysis between different groups was carried out using the DESeq2^33^ (version 1.12.4) Bioconductor ^24^ package with the R statistical programming environment (version 3.4.0).^25^ To identify the genes that affected differently by age across different diet groups, interaction analysis was conducted in DESeq2.

Non-coding RNA (ncRNA) annotations were subsetted within the differential gene expression obtained by the DESeq2 method in ENSEMBL, and Biomart (archived version of release 75) was used to identify the closest coding RNA.

### Weighted correlation network analysis (WGCNA) for construction of co-expression network

Based on the anticipated complexity of the DGE data, WGCNA^34; 35^ was used to identify clusters (modules) of highly correlated genes related to the external sample traits of age and obesity (using eigengene network methodology). As described above, the merged count matrix was used after retaining genes that had 5 or more counts in at least 50% of the samples. After normalization the log2 transformed counts were used as input to WGCNA (WGCNA, version 1.61). The function “*pickSoftThreshold*” was used to pick an approximate power value. Then “*blockwiseModules*” (networkType = “signed hybrid” and TOMtype = “signed”) was used to construct the co-expression network using: Soft thresholding power = 9, minimum module size = 30, the module detection sensitivity deepSplit=2, and cut height for merging of modules 0.25 (implying that modules whose eigengenes are correlated above 1 − 0.25 = 0.75 will be merged). Finally, *“chooseTopHubInEachModule”* returned genes in each module with the highest connectivity to all differentially expressed genes; these genes are often regulatory genes and represent candidate biomarkers.^36^

### Multivariate analysis of RNA-Seq data using PLS-DA

As described for Methyl-Seq, PLS-DA was also applied to multivariate analysis of the RNA-Seq data. Variables with VIP score >1 were used as potential biomarkers for the biological phenotype, and based on the numbers of variables with VIP score >1, further analysis was performed on potential biomarkers with VIP score >2. Specifically, normalized read counts were used as X variables, and diet and age were provided as Y responses in the R package *ropls*.

### Integration of Methyl-seq and RNA-seq data

ClosestBed in BEDTools ^37^ was used to extract the genes that were close to DMW. First the nearest genes and ncRNA that had a distance <2 kb with respect to the DMW were extracted using ClosestBed with the arguments: -D b. Then annotation information of the genes and ncRNA were added. For each comparison, only the DMW with genes <2kb were shown. The corresponding methylKit results and the DESeq2 results were also included using in-house scripts.

### Attribution of Biological Relevance

Ingenuity Pathways Analysis (IPA) (Qiagen, version 01-12) was used to examine the various comparisons, modules, and integrated data. Directionality of gene expression was considered for RNA-Seq data, but not for Methyl-Seq data, or for evaluation of genes in proximity to ncRNA identified in differential gene expression analysis. There were a large number of DMW with VIP>1; therefore, to keep datasets within IPA software limits for pathways analysis of the possible effects of hypo- and hypermethylation together, evaluation of biological function of DMW biomarker and upstream candidates was restricted to those with only VIP>2. Further, given the current limitations of building pathways in commercially available software from differential methylation data, only the directionality of the RNA species that were identified in both Methyl-Seq and RNA-Seq datasets were considered.

Analysis of differential gene expression between groups (Padj<0.05 comparisons) was performed in IPA in the context of biological knowledge relating to upstream cause of the differential gene expression and probable downstream effects, including diseases and functions used a combined enrichment score assessing overlap between observed and predicted gene sets (Fisher’s exact test P-value) and a Z-score to predict activation (positive z-score) or inhibition (negative z-score) state of the regulator based on match of observed and predicted up- or down- regulation patterns.^38^

### Generation of figures

All figures were generated using *ggplot2* and *cowplot* in R^25^, in IPA, or in JMP v 13.2.0 (SAS). For ease of identification in figures presenting both Methyl-Seq and RNA-Seq data, and because both datasets were corrected for FDR, q value was used to denote statistical significance for Methyl-seq data, and padj was used to denote statistical significance for RNA-seq data.

## Results

### Quality Control Outcomes

DNA and RNA yield and library preparation were of adequate quality to allow both Methyl-Seq and RNA-Seq data from the same animal to be used for n=7 YO, n=5 YL, n=5 AO, and n=6 AL mice. Additionally, there were n=1 YO, n=3 YL, n=1 AO, and n=1 AL datasets available for RNA-Seq only, and thus these were included in analyses of RNA-Seq data. Importantly, only animals with paired Methyl-Seq and RNA-Seq datasets were used for integration analyses. For all analyses, the focus was placed on the following biologically relevant comparisons: AO vs AL, AO vs YO, AL vs YL, and YO vs YL, A vs L and O vs L, with the AL vs YL comparison considered the healthy aging paradigm in this model.

### Differential Methylation Results

#### Single-base resolution DNA methylome using targeted bisulfite sequencing platform

DNA methylation was quantified in single-base resolution for a total of 109 Mb of coverage using targeted bisulfite sequencing. On average, 83% of the targeted regions were covered by at least one read and 59% by at least 10 reads (**Supplemental Table 1**). The medians of the coverage of the samples ranged from 8 to 20, with an average of 14. Bimodal distribution of CpG methylation levels were observed in all individual samples but with no observed difference in global methylation levels across the samples at both single-base level and window level (**Figure 1A,B**).

**Figure 1:**
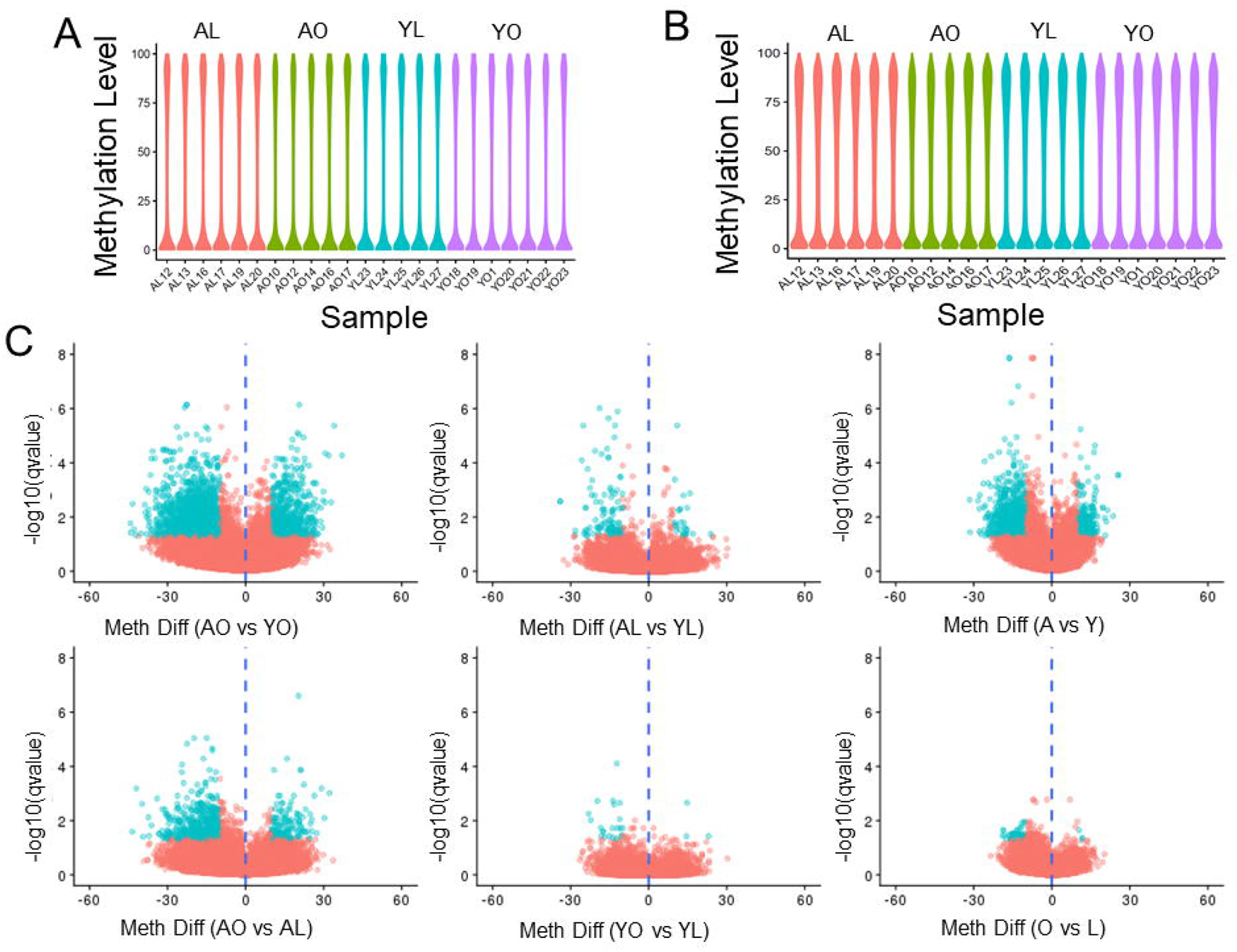
Violin plots showing distribution methylation levels for CpG (A) and differentially methylated windows (DMW) (B), and (C) volcano plots of methylation difference from MethylKit results for each group (A&B) and biologically relevant comparison (C), aged lean (AL), aged obese (AO), young lean (YL), young obese (YO), aged vs young (A vs Y), and obese vs lean (O vs L).

#### Identification of DMC and DMW demonstrated global hypomethylation in aged and/or obese mice (Supplemental Table 2 and Supplemental Table 3)

Consistent with the results observed in **Figure 1A**, when the biologically relevant comparisons (AO vs YO, AL vs YL, AO vs AL, YO vs YL, A vs Y, O vs L) were evaluated, it was observed that methylation difference were centered at zero **(Figure 1C)** The percentage of windows with methylation difference below 5% ranged from 79% to 93% across all comparisons, with an obvious bias between the number of hyper- and hypomethylated windows (**Figure 1C**). The percentage of hypomethylated windows (methylation difference < 0) ranged from 38.4% to 44% demonstrating that aged and/or obese mice had more windows with hypomethylation compared to young and/or lean mice.

This bias in favor of hypomethylation was also reflected in the DMW (windows with q ≤ 0.05 and methylation level differences >±10%), and DMC (**Supplemental Table 4**). Further, there were more total DMW in aging in obese mice compared to lean mice, and in both aged and young obese mice compared to lean aged or young mice respectively, suggesting that obesity was synergistic or at least additive to the overall effect of aging. There were a greater proportion of DMC not in DMW for obese mice, particularly with increased age, suggesting that DMC changes in obesity were more robust to statistical evaluation than methylation changes in lean mice or that there is a clustering of DMC not in DMW in obese, but not in lean mice.

We then evaluated the distribution of DMW and DMC based on the annotation of CGI in the mouse genome. Using all the analyzable windows for each comparison pair as background, DMW were less common in CG islands, CG shores and CpG shelves and more common in interCGI regions **(Figure 2A)**. For the distribution of DMW based genic annotations compared to background, we found that DMW were primarily in intron followed by intergenic regions **(Figure 2B)**. Distribution of methylation level by gene region (**Figure 2C**) suggested even distribution between hyper- and hypomethylation at 1-5kb upstream, CDS and intronic sites, whereas promoters and 5’UTR were generally hypomethylated and had lower density of methylation compared to other genic regions, while 3’UTR seemed to have a preponderance of hypermethylation and the highest density of methylation. There was no obvious difference in global gene region methylation density or level between animal groups (DMC data similar, data not shown).

**Figure 2:**
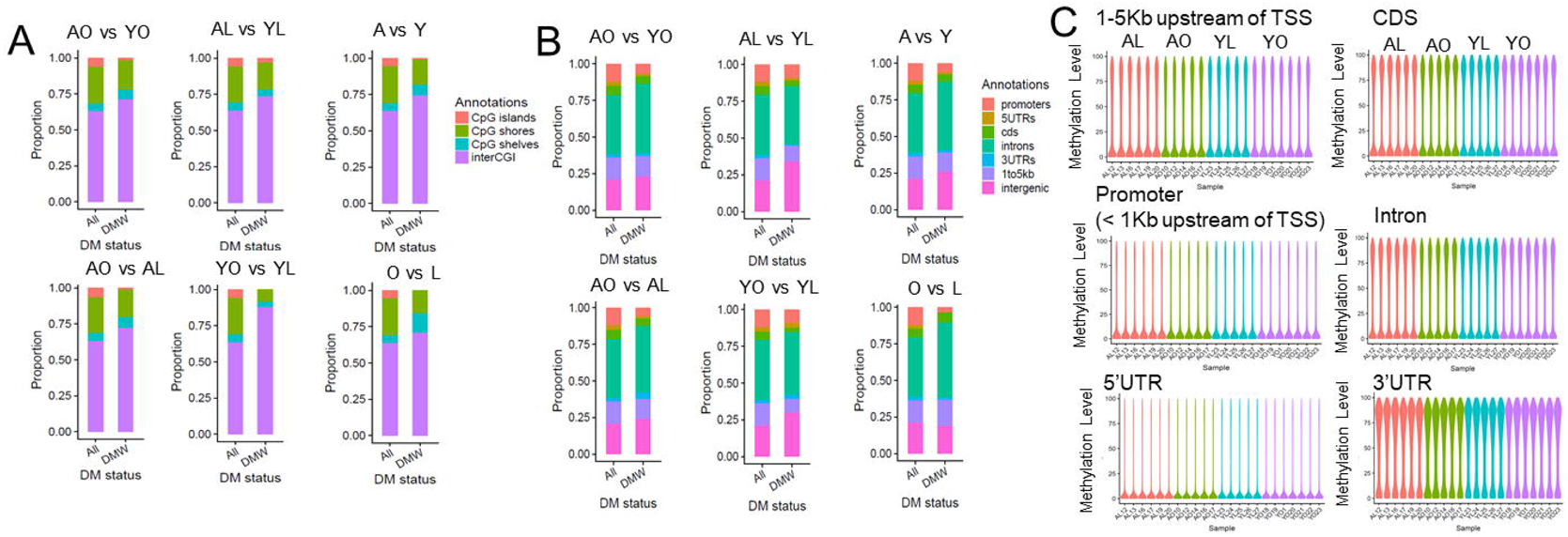
Distribution of differentially methylated windows for each biologically relevant comparison (see Figure 1 legend) based on (A) annotation to CpG islands, shores, shelves, or inter-CpG island (CGI) regions, or annotation to (B) genic region. (C) Violin plots of methylation level for the different genic regions in the different groups.

### Analysis of the DMW using PLS-DA identified several differentially methylated (q<0.05) and differentially expressed genes (padj<0.05) involved in upstream mechanistic networks and in cellular dysfunction

Score plots of PLS-DA model in the plane of the first predictive (t1) and the second predictive (t2) components for the comparisons were shown in **Figure 3A**, and when comparisons were made between the four groups of mice in **Figure 3B.** While there was good separation between the data (R2Y), the response variance explained by t(1) was modest, leading to modest predictive performance of the model (Q2Y).

**Figure 3:**
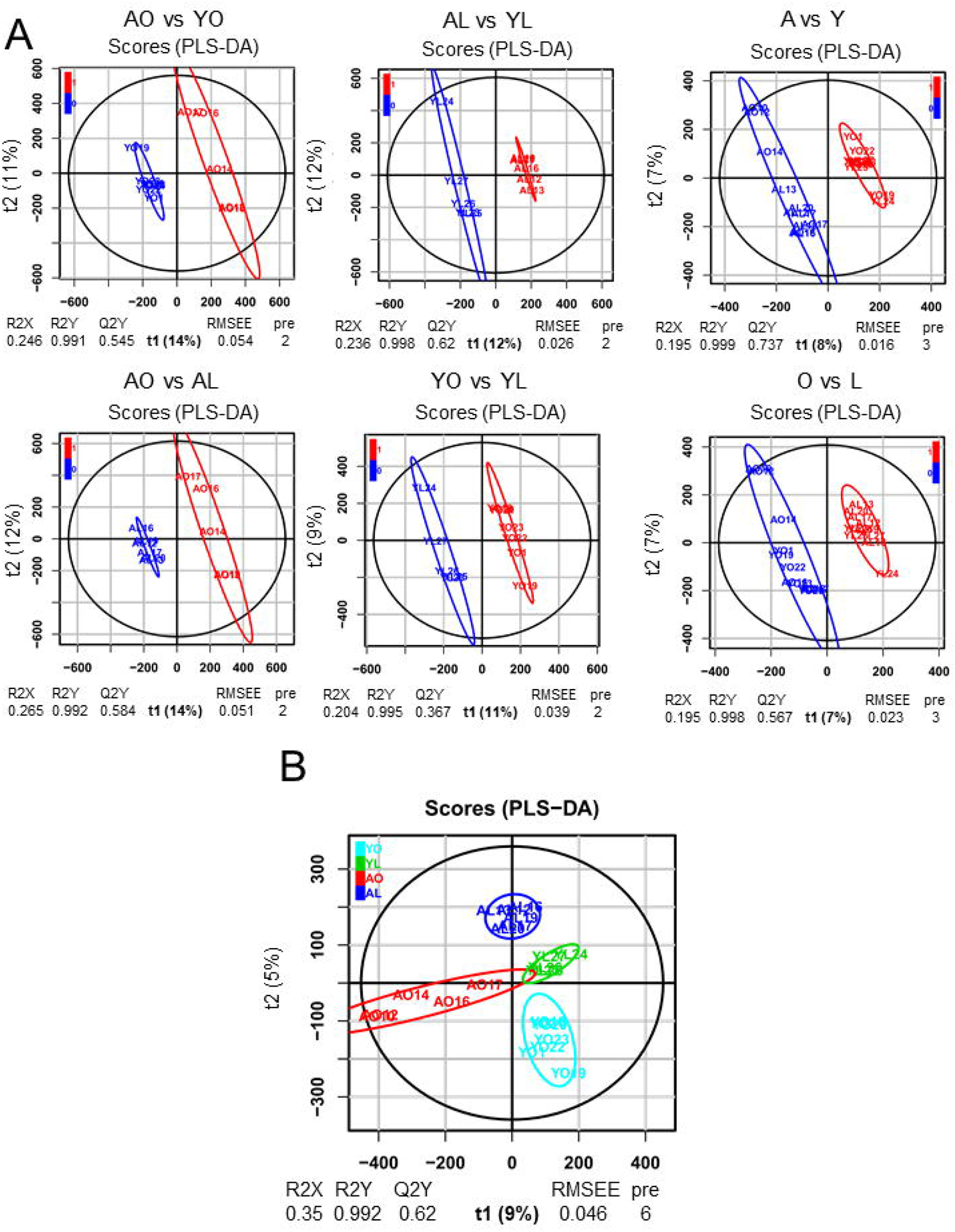
Score plots of partial least squares discriminant analysis (PLS-DA) mode in the plane of the first predictive (t1) and the second predictive (t2) components for the different comparisons (A), and between the four different groups (see Figure 1 legend). The percentage of response variance explained by the predictor component only (t1) is indicated in parentheses. R2X (respectively R2Y) means percentage of predictor (respectively response) variance explained by the full model. Q2Y means predictive performance of the model estimated by cross-validation. For the classification model, the ellipses corresponding to 95% of the multivariate normal distributions with the samples covariance for each class is shown.

Because we used the stringent cut of VIP>1 and q<0.05, the comparison between multivariate and univariate analysis **(Figure 4AB and Supplemental Table 5)** yielded only a small proportion of the total DMW. A summary of the number of DMW with VIP scores >1 or >2 in **Figure 4C** is filtered by total, qvalue<0.05 for PLS-DA and padj<0.05 for the associated comparison of gene expression (Full dataset in **Supplemental Table 6**). Then, to improve understanding of the interconnectedness of highly significant VIP scores for each comparison between groups of animals, mechanistic networks were generated in IPA for all upstream regulators for hyper- and hypomethylated DMW with VIP>2 associated with significant differences in gene expression (padj<0.05) present within each biologically relevant comparison dataset (**Figure 5A**) and examined for commonality between comparison groups (**Figure 5B**). *Mapt* was the only gene involved in both hyper- and hypomethylated upstream mechanistic networks for several biological comparisons (AO vs YO hypomethylated, AL vs YL hypomethylated, YO vs YL hypermethylated), while *Foxo3* and *Ccnd1* were involved in hypermethylated biological comparisons only (AL vs YL and YO vs YL). Several additional genes were involved in several biological comparisons of hypomethylated upstream mechanistic networks: *Nr3c2* for AO vs YO and AL vs YL comparisons, *App* and *Ctnnb1* for AL vs AL, AO vs YO and AL vs YL comparisons, and *Hipk2*, *Id2* and *Tp53* overlapped between AO vs AL and AO vs YO comparisons (**Figure 5B**).

**Figure 4:**
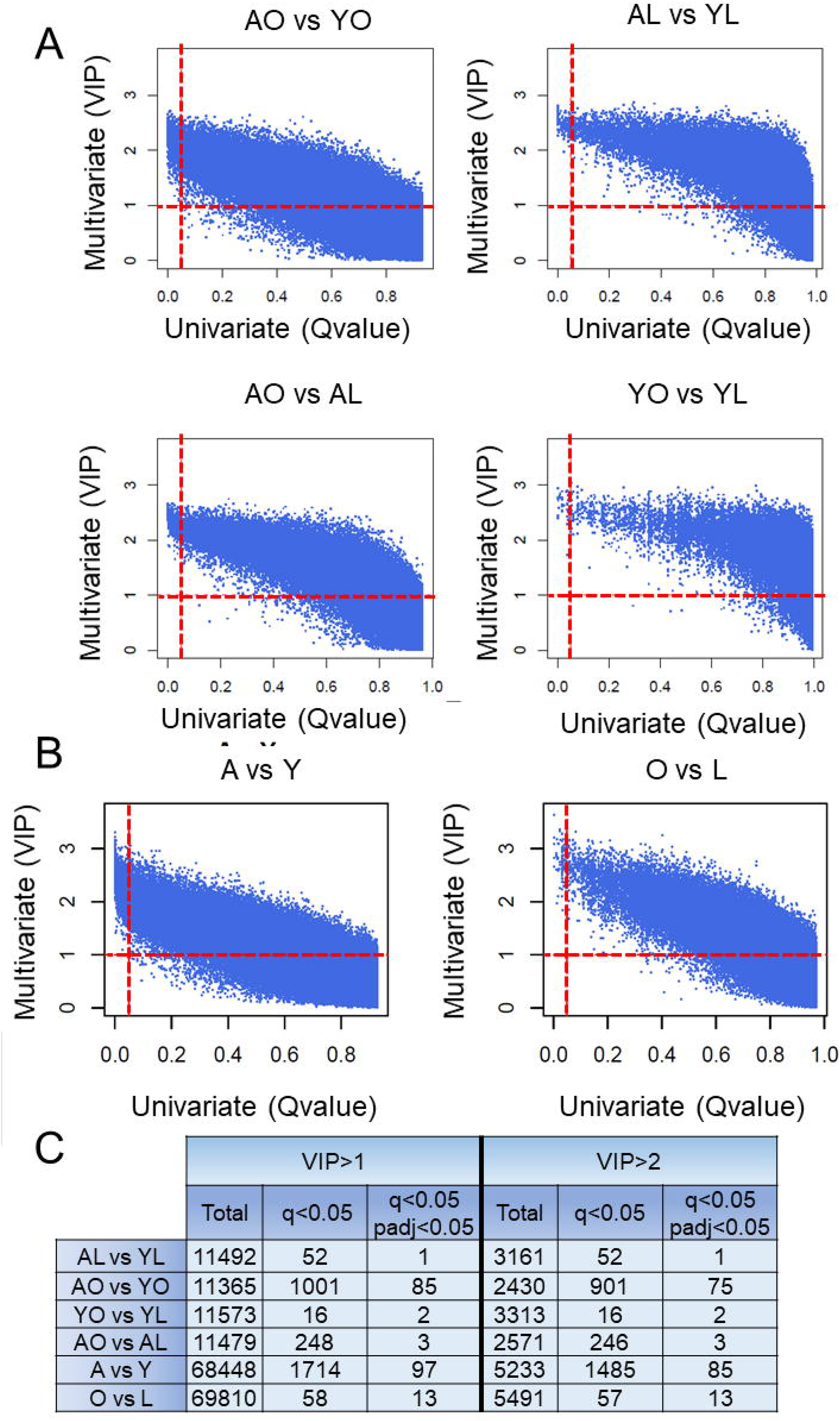
(A, B) Scatter plots of multivariate analysis by partial least squares discriminant analysis (PLS-DA) compared to univariate analysis by DESeq2 for the various biologically relevant comparisons (see Figure 1 legend). Variable Importance in Projection (VIP) scores and q-values were used together to identify potential biomarkers for the effects of age and diet. (C) Summary of the numbers of differentially methylated windows identified using these methods for the cut-offs established, including the effect of adding the associated differential gene expression at padj<0.05.

**Figure 5:**
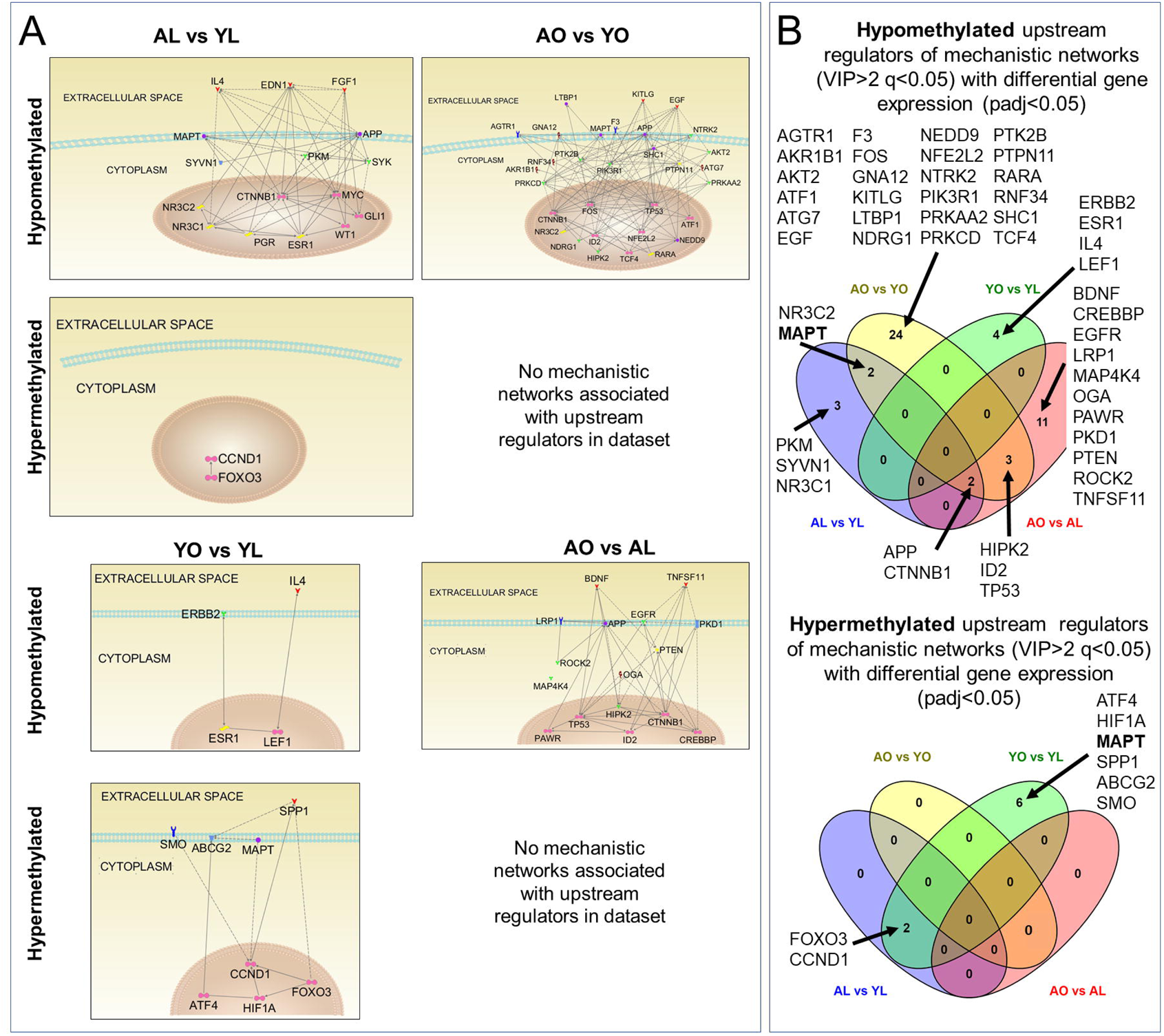
(A) Upstream regulators and mechanistic networks identified in Ingenuity Pathways Analysis (IPA) of hypo- and hypermethylated differentially methylated windows identified with Variable Importance in Projection (VIP) scores >2 and q<0.05 by partial least squares discriminant analysis (PLS-DA). (B) Overlap between comparisons for hypo- and hypermethylated upstream regulators of mechanistic networks with differential gene expression identified in IPA. https://bioinfogp.cnb.csic.es/tools/venny/index.html

Finally, diseases and functions were overlaid onto the network of upstream regulators, and the top ten most significant were summarized for each comparison in **Figure 6**. In general, for all comparisons, the disease and function terms ‘tumor’, ‘apoptosis’, ‘proliferation’, ‘development’, ‘cell cycle’ were highly represented, particularly for ‘epithelial’, ‘gland’ and ‘fibroblast’ tissues or cell lines.

**Figure 6:**
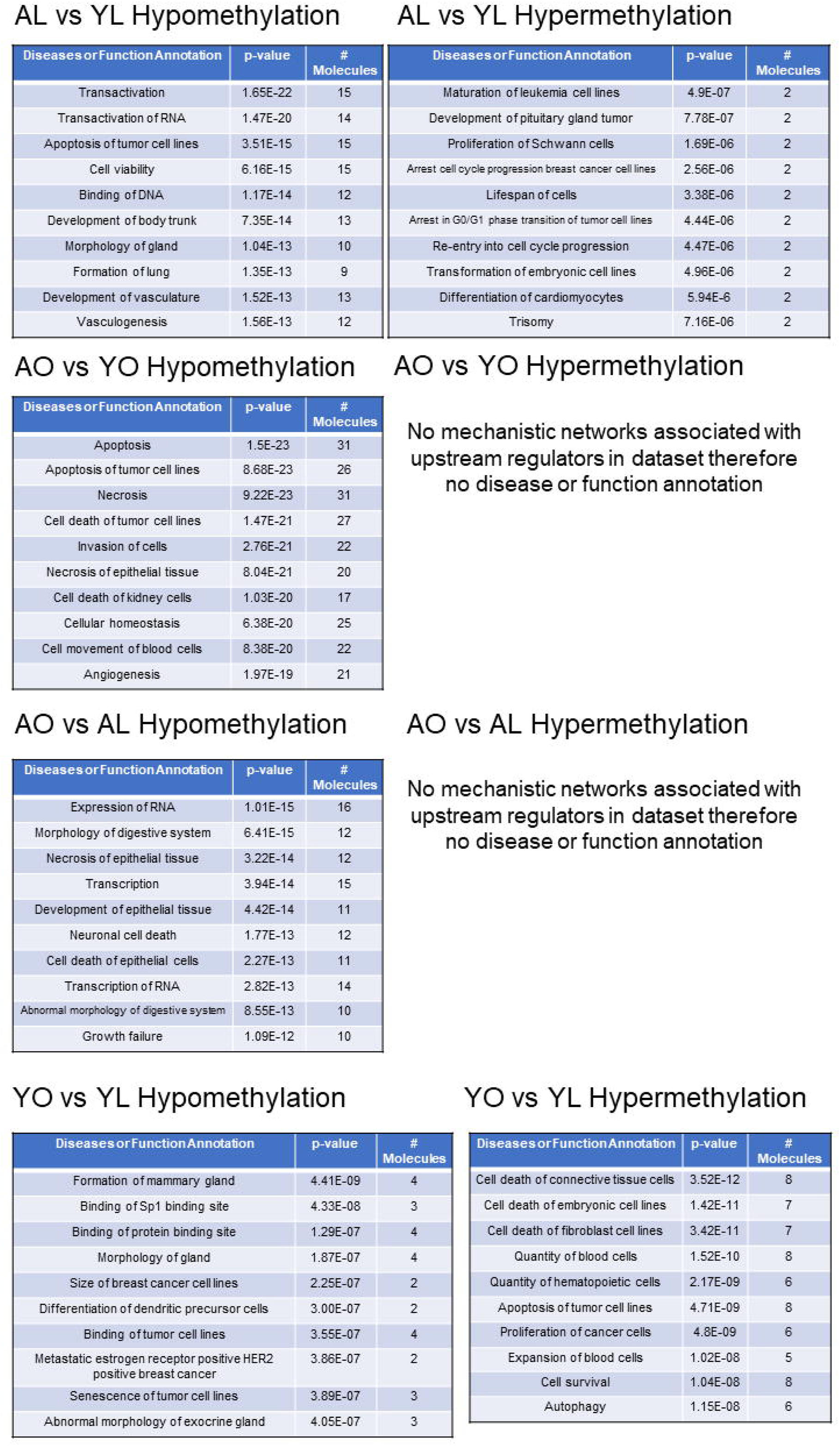
Summary of ‘top ten’ most significant diseases and functions annotations in Ingenuity Pathways Analysis (IPA) for the networks of upstream regulators and mechanistic networks identified for each comparison in Figure 5.

### Diet or Obesity Related Methylation Drift with Age was not observed in these Murine ASCs

To evaluate if age- or obesity related drift in methylation could be identified in our data from mice at 20- and 52-weeks of age, we first extracted all windows and CpG with differential methylation across the entire dataset, and a standard deviation >10 (7.41% of windows and 12.82% of CpG for AL vs YL) was selected to identify the most variable sites for each individual mouse around a mean of all analyzable windows or CpG. We expected this would separate young from old lean mice into two distinct clusters (**Supplemental Figure 2A,B**). However, this was not observed, and when the same process was repeated for the aging in obesity comparison, AO vs YO, while a greater proportion of DMW and CpG had SD>10 (12.53% DMW and 17.37% of CpG having SD>10), clustering into distinct clusters was also not observed **(Supplemental Figure 2 D,E)** suggesting that age-related methylation drift reported by others^39^ did not occur over this age range in murine ASCs. We checked our results using a less stringent cut off to identify less substantial methylation drift, a standard deviation of >5, and similarly, age-related methylation drift was not observed for either AL vs YL or AO vs YO comparisons. Analysis of 32 CpG sites that overlapped with a differentiated multi-tissue DNA methylation clock across the mouse lifespan^5^, also did not cluster the mice into phenotypic groups for either comparison (**Supplemental Figure 2C,F**) even though this clock has previously identified ‘rejuvenation’ of fibroblasts to induced pluripotent stem cells^5^.

### RNA-Seq Results

#### Differential Gene Expression

Cluster dendrograms and principal components analysis for differential expression of all RNA biotypes (**Figure 7A, B**) demonstrated considerable divergence between the groups, particularly with increasing age and obesity. Heat maps by age or diet condition (**Figure 7C**) and experimental group (**Figure 7D**) demonstrated a substantial number of RNA species impacted most notably by the combination of aging and obesity.

**Figure 7:**
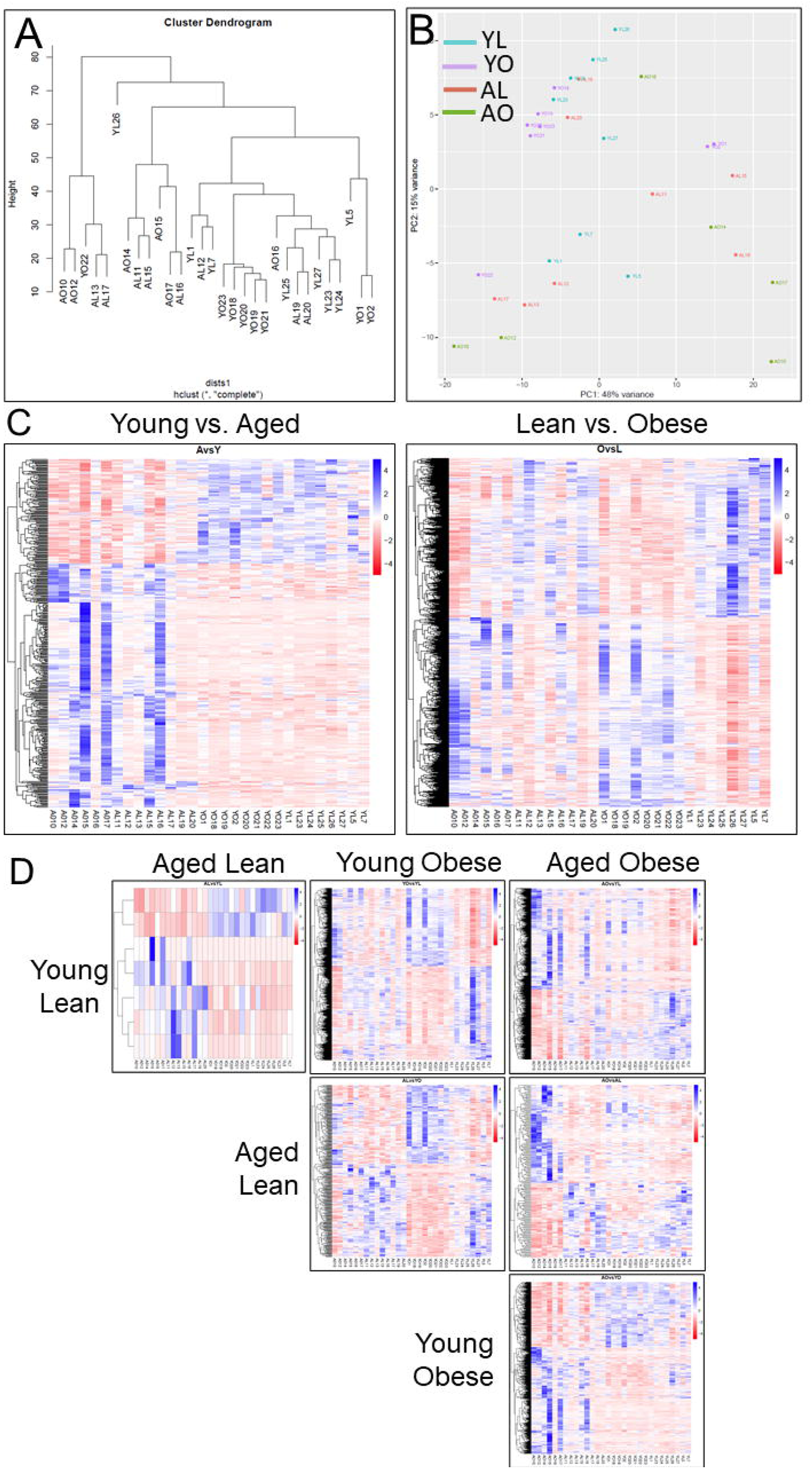
(A) Cluster dendrogram of differential expression for all genes from RNA-Seq analysis, and (B) Principal component analysis for mRNA biotypes. Heat maps by diet or age condition (C) and experimental group comparisons (D) for all RNA biotypes.

When evaluating differential RNA biotype expression (**Supplemental Table 7**) and summarized in **Table 1**, obesity (O vs L) influenced expression of almost 5-fold more RNA biotypes (Padj<0.05) than aging (A vs Y), and this effect was most profound when obesity was superimposed on aging (AO vs YO compared to AL vs YL). In this regard, the ASCs from lean mice demonstrated a remarkably stable transcriptome with aging (AL vs YL), with differential expression of only 7 RNA biotypes; whereas there was >120-fold increase in number of RNA biotypes with significantly different expression in AO vs YO mice. With respect to directionality of RNA biotype expression, aging generally increased expression, whereas the influence of obesity was to moderate the increase in gene expression. Similarly, for the magnitude of expression changes (>2-fold change), obesity reduced the number of RNA biotypes with large changes in expression induced by aging, regardless of whether this was for a >2-fold increase or decrease in expression (**Table 1; Supplemental Table 7**). Overall mRNA transcripts demonstrated similar patterns as the patterns of ‘ALL’ RNA biotypes. While our RNA extraction technique was not designed to capture all small RNAs and non-coding RNAs, we identified 1752 of these species following annotation of the data in ENSEMBL (**Supplemental Table 7)**. While many of these other biotypes demonstrated no significant differences (padj>0.05) in expression of specific RNA species with aging or obesity, many of these RNA biotypes followed the same general patterns as for as for mRNA transcripts.

**Table 1:** Summary of numbers and proportions of differentially expressed RNA biotypes (increased and decreased expression), and of numbers and proportions of differentially expressed RNA biotypes with >2-fold increase or decrease in expression.

| Comparison | Gene and Transcript Biotype | Total Number of Biotypes | Padj <0.05 | %Padj <0.05 | Of Padj<0.05 Biotypes |  |  |  |  |  |  |  |
| --- | --- | --- | --- | --- | --- | --- | --- | --- | --- | --- | --- | --- |
|  |  |  |  |  | Total Increase | % Increase | Total Decrease | % Decrease | >2-fold Increase | %>2-fold Increase | >2-fold Decrease | % >2-fold Decrease |
| O vs L | ALL | 14987 | 2075 | 13.85 | 1124 | 54.17 | 951 | 45.83 | 76 | 6.76 | 60 | 6.31 |
| A vs Y | ALL | 14987 | 424 | 2.83 | 297 | 70.05 | 127 | 29.95 | 122 | 41.08 | 14 | 11.02 |
| YO vs YL | ALL | 14987 | 1774 | 11.84 | 807 | 45.49 | 967 | 54.51 | 65 | 8.05 | 123 | 12.72 |
| AO vs YL | ALL | 14987 | 2052 | 13.69 | 1213 | 59.11 | 839 | 40.89 | 348 | 28.69 | 152 | 18.12 |
| AO vs AL | ALL | 14987 | 189 | 1.26 | 108 | 57.14 | 81 | 42.86 | 60 | 55.56 | 39 | 48.15 |
| AL vs YO | ALL | 14987 | 367 | 2.45 | 198 | 53.95 | 169 | 46.05 | 62 | 31.31 | 37 | 21.89 |
| AL vs YL | ALL | 14987 | 7 | 0.05 | 5 | 71.43 | 2 | 28.57 | 5 | 100.00 | 1 | 50.00 |
| AO vs YO | ALL | 14987 | 842 | 5.62 | 521 | 61.88 | 321 | 38.12 | 256 | 49.14 | 67 | 20.87 |

#### Biological Relevance of Differential Gene Expression

In lean mice, we identified no significant effect of aging on IPA designated ontological functions, or upstream regulators. In ‘healthy aging’ mice (AL vs YL), the only altered and scored networks identified involved ‘cellular development, cellular growth and proliferation’ and ‘hematological system development and function’, and diseases and functions involved ‘Cell Death and Survival’, ‘Gene Expression’, and ‘Tissue Morphology’ (**Supplemental Table 8**). However, all other comparisons examined involving obesity identified dozens of putative upstream regulators, networks, functions, and diseases and functions impacted, many of which have been extensively associated with diseases of obesity (**Supplemental Table 8**). DESeq2 was used to examine the interaction between age and diet, and 7 differentially expressed (padj<0.05) genes were identified: *Ccl9, Gucy1b1, Ltbp4, Kif1a, Cmtm4, Itpr3,*and *Piezo2*. (**Figure 8A,B, Supplemental Table 9**)

**Figure 8:**
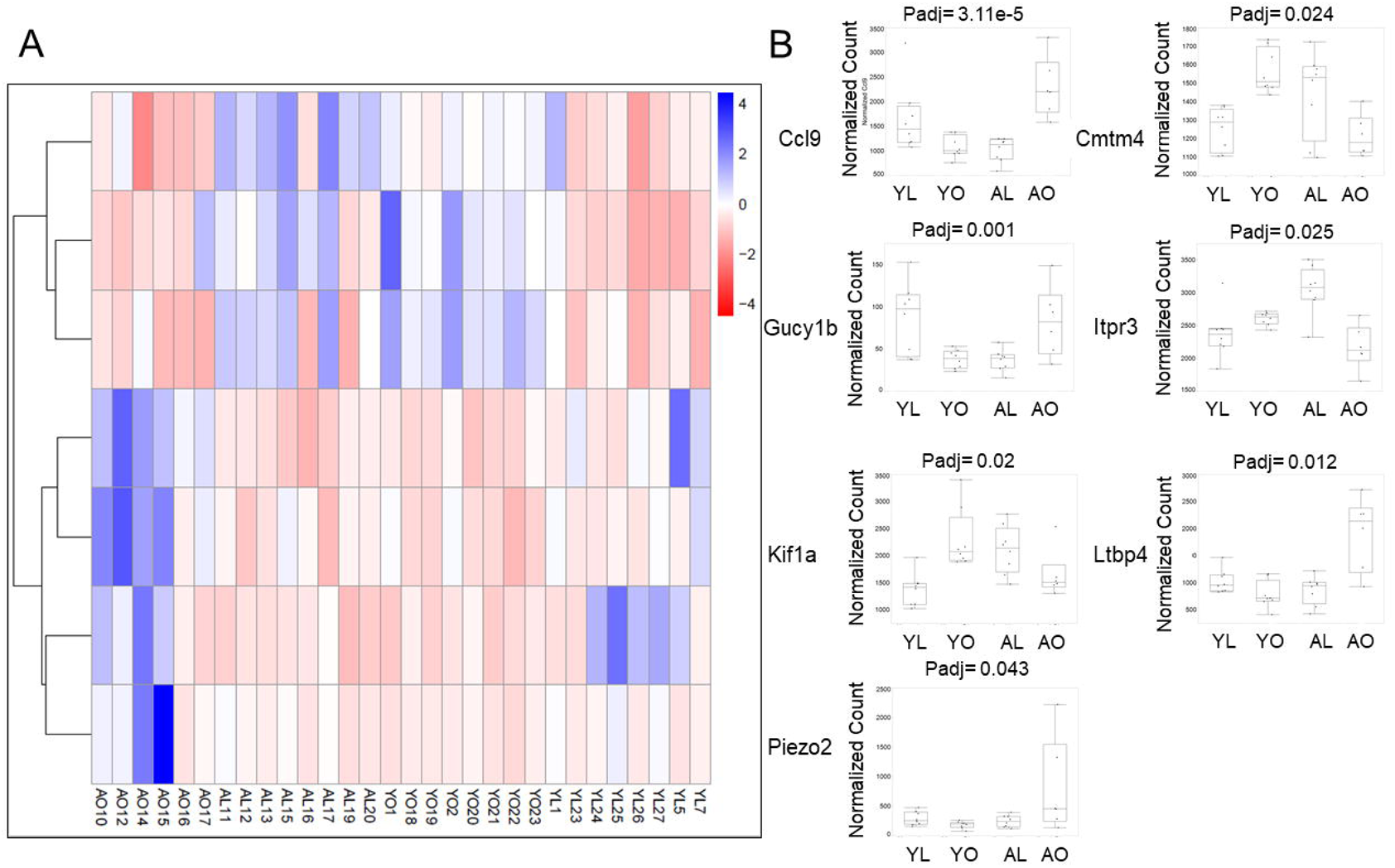
Heat map (A) and normalized counts (B) for seven genes differentially expressed (padj<0.05) with diet and age identified by DESeq2 analysis.

#### WGCNA Results

Based on the complexity of the differential expression data, eigengene network methodology WGCNA was used to identify unique (no overlapping RNA biotypes) modules of highly correlated genes related to the external sample traits of age and diet in all mice, and bodyweight at euthanasia for aged mice. Each module was a cluster of highly interconnected genes related (positive or negative correlation) to diet or age, and for aged mice for the external trait of bodyweight at euthanasia. First, considering all groups of mice, aged and obese condition were assigned trait values =1, and young or lean condition were assigned trait values of 0. Relationships between the consensus module eigengenes (the first principal component of a given module) and sample traits were generated (**Figure 9A, B**), and biotypes present in each module are found in (**Supplemental Table 10)**. Module eigengenes were considered as representative of the gene expression profiles in a module. For age, there were 5 modules negatively correlated with age, with p<0.05, and 7 modules positively correlated with age. For diet, there were 2 modules negatively correlated with diet, but no modules positively correlated with diet. To better assess the gene modules specifically associated with bodyweight at sacrifice in aged mice rather than diet itself, trait value of bodyweight was assigned and the WGCNA analysis was repeated (**Figure 9C-E**) and biotypes present in each module are found in (**Supplemental Table 11)**. This time, there were 5 modules negatively correlated with bodyweight, and 4 modules positively correlated with bodyweight, at p<0.05.

**Figure 9:**
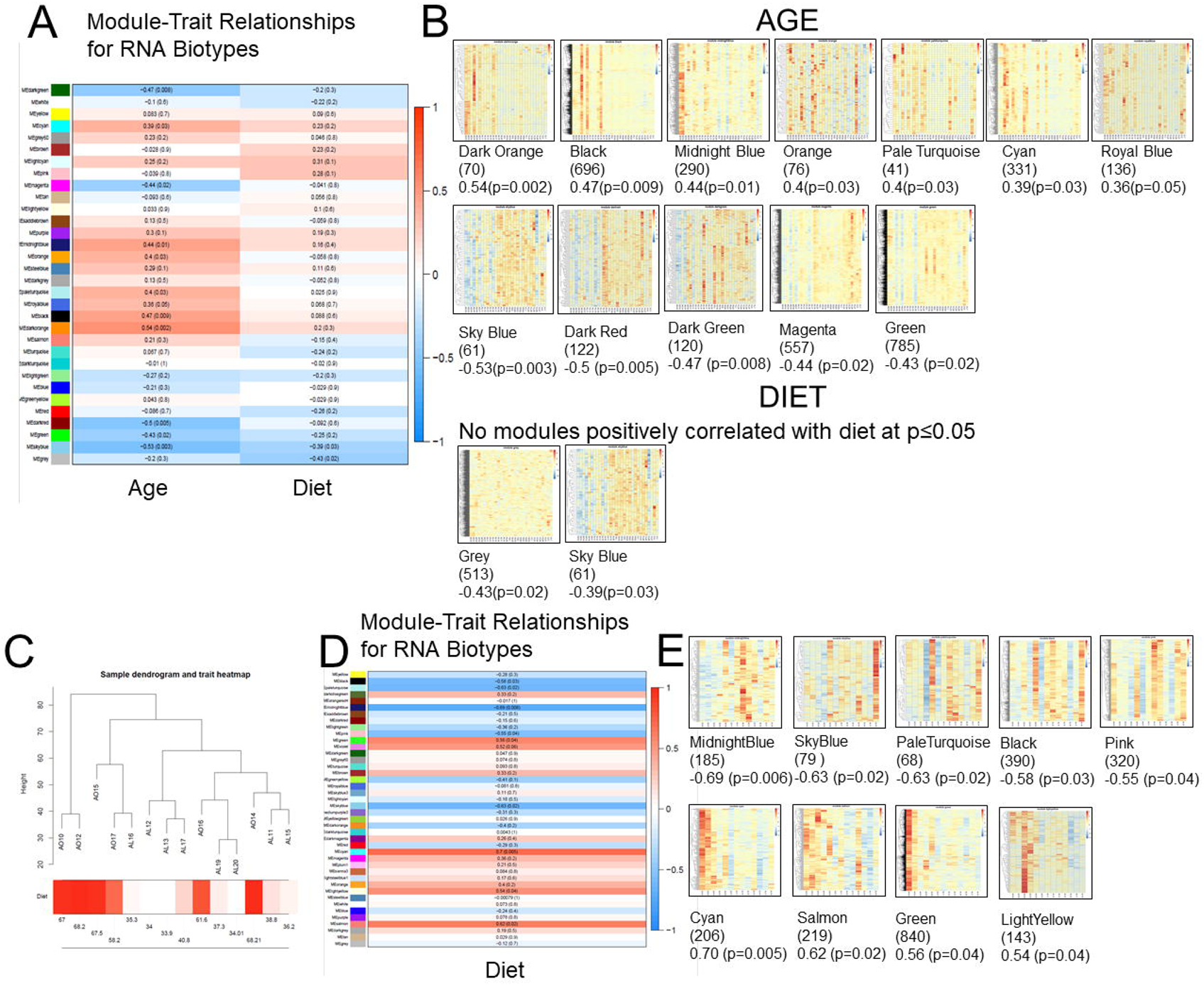
Module-trait relationships (A) reporting the relationships of consensus eigengenes (the first principal component of a given module) and sample traits. Each row in the figure corresponds to a consensus module, and each column to a trait. Numbers in the table report the correlations of the corresponding module eigengenes and traits, with the p-values printed in the correlations in parentheses. The table is color coded by correlation according to the color legend. Heatmaps (B) for significant (p<0.05; parentheses) modules for age and diet, with number of biotypes and the r-value highlighted. (C) Sample dendrogram and trait heatmap, (D) module-trait relationships, and (E) heatmaps of significant modules for bodyweight of aged mice at sacrifice.

#### ncRNA in WGCNA modules

Some modules contained large numbers of ncRNA biotypes (Age and Diet, range 3.9-61.8% ncRNA of total RNA biotypes (**Supplemental Table 12**), bodyweight at euthanasia in aged mice, range 2.8-26.6% ncRNA of total RNA biotypes (**Supplemental Table 13**). While ncRNAs that were annotated were included within the baseline IPA analysis for each module, we attempted to gain further additional insight into the possible function of the ncRNAs within the context of the coding RNA also present within the module. Each ncRNA species within distinct WGCNA modules significantly associated with age, diet, or bodyweight at euthanasia was examined for the presence of coding transcripts within ±20 kb of their locus. 43% modules contained ncRNA that was within ±20kb of a coding RNA species in the same module, and all modules had at least one in-module ncRNA species (range 1-72 coding species) within ±20kb of an mRNA species not in the original module. Further, several of these mRNA species were within 20kb of more than one ncRNA species within a module, suggesting the possibility of multiple additional regulatory pathways, both within modules, between modules, and to mRNA species not associated with a module. Further, in pathways analysis of the mRNA within 20kb of ncRNA species, aged obese mice had different nodes to those identified in aged lean mice, and the connectedness of mRNA within 20kb of ncRNA within networks was reduced in obesity, suggesting the combination of aging and obesity could disrupt normal communication between mRNA species regulated by ncRNA.

#### Biological Relevance of Significant WGCNA Modules

IPA analysis was performed on the modules with the associated fold-change expression data for the most relevant biological comparisons (YO vs YL, AO vs AL, AO vs YO, and AL vs YL), and separately on the nearest protein-coding data from the evaluation of ncRNA within the modules. The later was performed without regard for directionality of potential expression since the effect of change in expression of ncRNA on the expression of the nearest mRNA was unknown. The “*ChooseTopHubInEachModule*” function was performed using WGCNA (version 1.61) R package and then network analysis was performed in IPA to examine interconnectedness between the genes identified for each module across different regions of the cell (**Figure 10**), with top hub genes *Tgfb1, Mcl1* and *Mapk14* noted as especially inter-connected with other module top hub genes for all conditions (**Figure 10A**). Significant canonical pathways for these top hub genes were ‘Type I Diabetes Mellitus Signaling’, ‘Inhibition of Angiogenesis by TSP1’, ‘Antigen Presentation Pathway’, ‘Role of Osteoclasts, Osteoblasts, and Chondrocytes in Rheumatoid Arthritis’, and ‘T Helper Cell Differentiation’. When examining the interactions of diet with bodyweight in aged mice (**Figure 10B**), *HLA-A, Gas6*, and *Nedd9* were noted as especially interconnected with other module top hub genes, and significant canonical pathways for these top hub genes were ‘Neuroprotective Role of THOP1 in Alzheimer’s Disease’, ‘Protein Ubiquitination Pathway’, ‘Sucrose Degradation V (Mammalian)’, ‘Ketogenesis’, and ‘Mevalonate Pathway I’. IPA analysis of the RNA biotypes in the modules revealed wide ranging canonical pathways, upstream, causal, networks, and diseases and functions involved. In this regard, top diseases and functions across all modules were related to oncogenic or neoplastic processes, or to tumors (Diseases and Functions presented as Upset Plots in **Supplemental Figure 3**).

**Figure 10:**
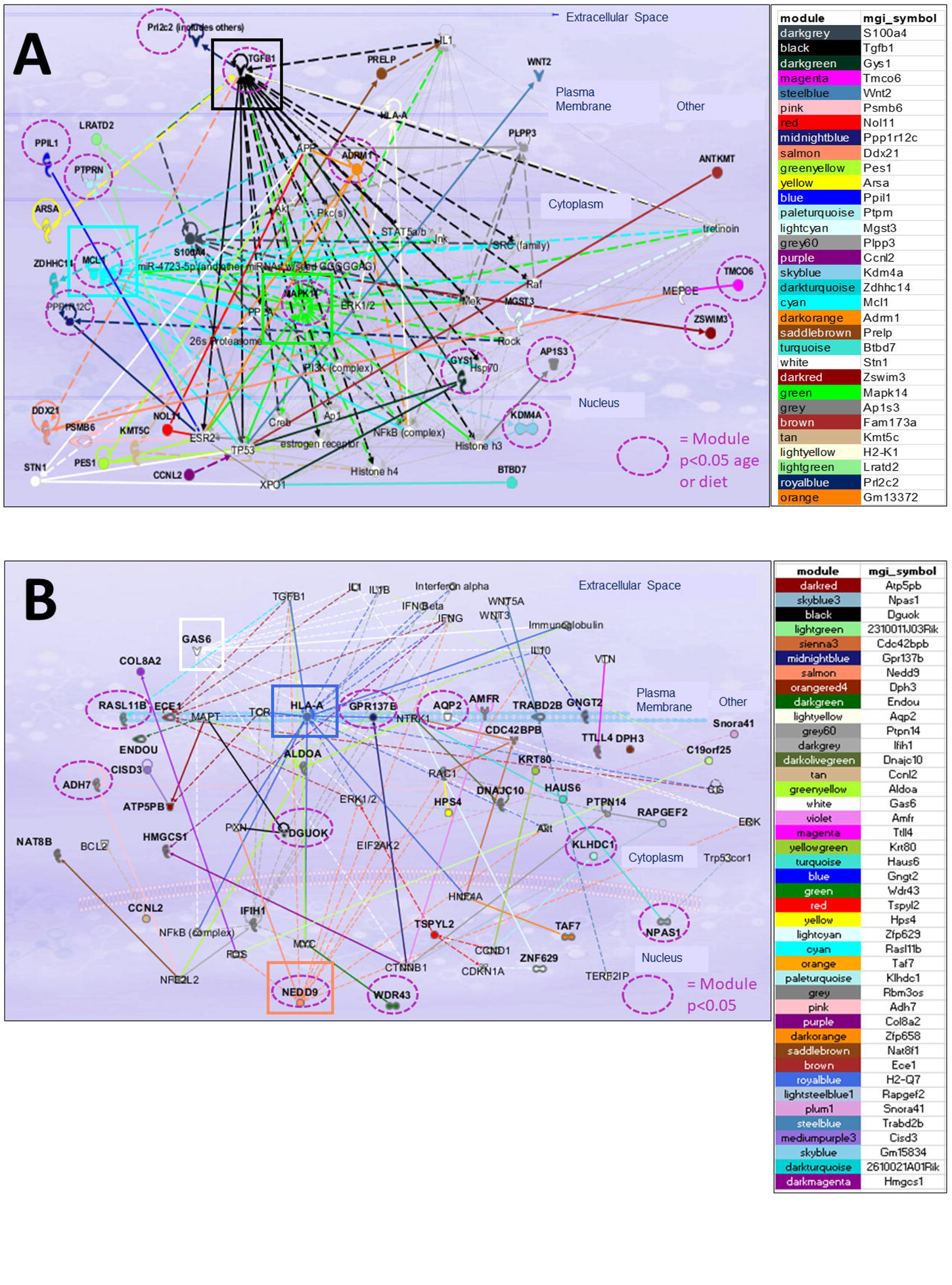
Networks generated in Ingenuity Pathways Analysis (IPA) and organized by cellular localization to examine interconnectedness between top hub genes in each significant modules identified by weighted correlation network analysis (WGCNA) for all conditions (A) and for the interactions of diet with bodyweight in aged mice at euthanasia (B). For clarity in presentation, non-hub genes have been removed, if they could be removed without impacting the interconnectivity between hub genes.

#### PLS-DA

As expected from the preceding analyses, score plots of PLS-DA model in the plane of the first predictive (t1) and the second predictive (t2) components for age (**Figure 11A**) and diet (**Figure 11B**) for Ensembl genes demonstrated a greater effect of diet than age, with less separation between age components than diet (**Supplemental Table 14**). However, PLS-DA analysis of genes from Ensembl database for the interaction of age and diet did not have a good performance based on Q2Y values (0.193). Scatter plots of univariate analysis in DESeq2 (**Figure 11C,D**) and multivariate analysis in PLS-DA demonstrated several genes for both age (**Figure 11C**) and diet (**Figure 11D**) with VIP scores greater than 1 and Padj <0.05.

**Figure 11:**
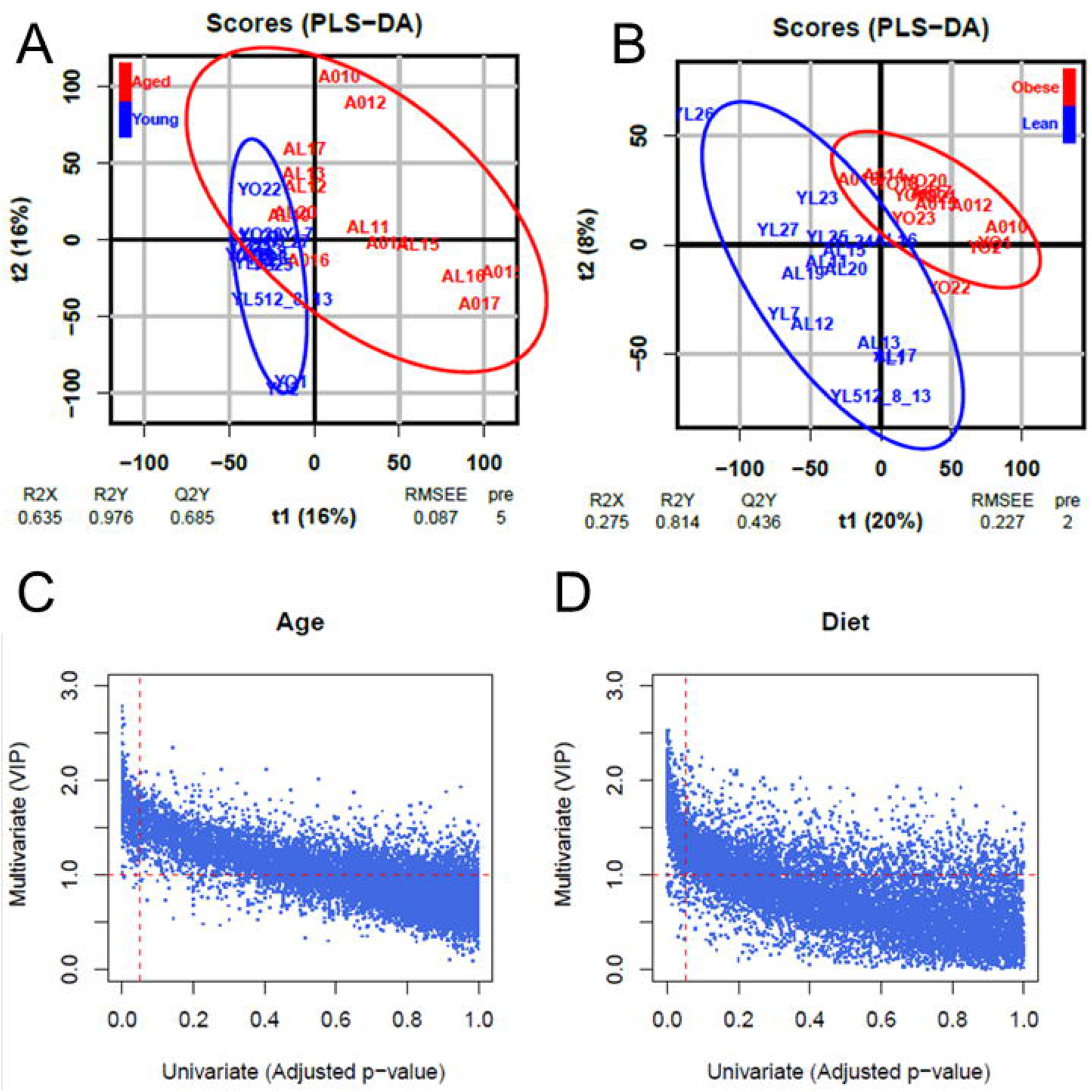
Score plots of partial least squares discriminant analysis (PLS-DA) mode in the plane of the first predictive (t1) and the second predictive (t2) components for RNA biotype responses to age (A) and diet (B). The percentage of response variance explained by the predictor component only (t1) is indicated in parentheses. R2X (respectively R2Y) means percentage of predictor (respectively response) variance explained by the full model. Q2Y means predictive performance of the model estimated by cross-validation. For the classification model, the ellipses corresponding to 95% of the multivariate normal distributions with the samples covariance for each class is shown. Scatter plots of multivariate analysis by partial least squares discriminant analysis (PLS-DA) compared to univariate analysis by DESeq2 for age (C) and diet (D). Variable Importance in Projection (VIP) scores and padj-values were used together to identify potential RNA biomarkers for the effects of age and diet.

There were 420 genes with VIP≥1 and p<0.05 related to age, of which 385 were also differentially expressed, and 2013 genes related to diet of which 1610 were also differentially expressed, with 9 genes VIP≥1 and p<0.05 for both age and diet (*Capn6, Dusp6, Gzme, Il11, Itih2, Klra4, Rap1gap2, Relt, Tnfrsf12a*), all of which were also differentially expressed (**Figure 12**). Given the versatility of PLS-DA to predict discrete variables, in this case obesity or aging, the PLS-DA transcripts with VIP ≥2 and p<0.05 (45 genes for age (**Supplemental Figure 4**), 105 for diet (**Supplemental Figure 5**), including 1 gene (*Capn6*) differentially expressed and VIP≥2 for both age and diet) were considered as potential ASC biomarkers for obesity and/or age. Given that identification of putative driver genes for ASC dysfunction in aging and obesity was an aim of this work transcriptional regulators with VIP≥2 were considered for additional evaluation. Of the 105 differentially expressed genes with VIP≥2 for diet, 6 were transcriptional regulators (*Cdkn2a, FoxG1, Ncoa3, Pax3, Plagl1* and *Plag1*). Of the 45 differentially expressed genes with VIP≥2 for age, 9 were transcriptional regulators (*En2, Gzf1, Shox2, Sox12, Tfec, Vhl, Zbtb18, Zfp78,* and *Zfp992*).

**Figure 12:**
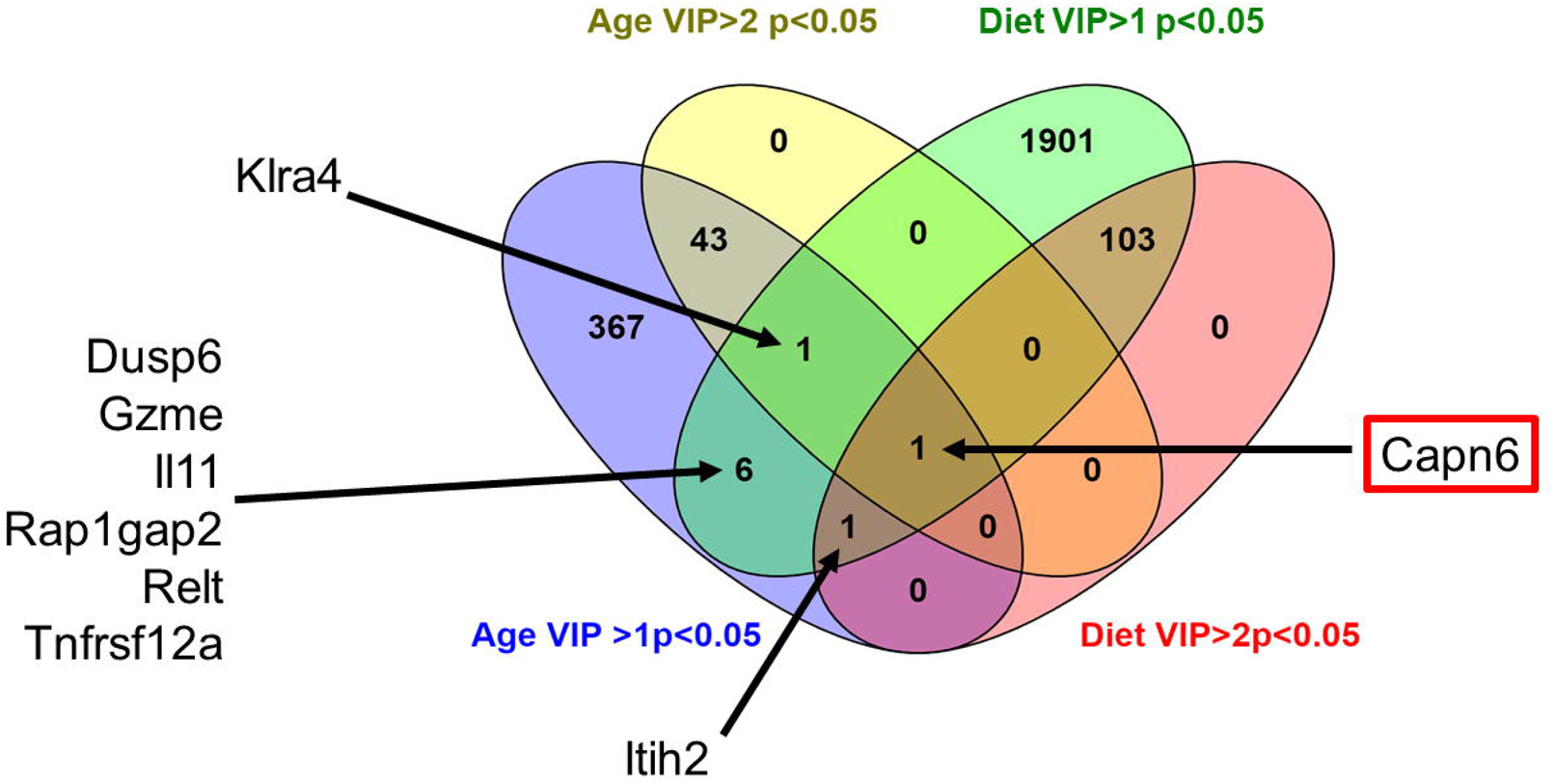
Overlap between RNA biotypes with Variable Importance in Projection (VIP) scores >1, or >2 and padj<0.05 related to age or diet. https://bioinfogp.cnb.csic.es/tools/venny/index.html

Finally, comparison of PLS-DA age dataset to genes previously associated with methylation drift in blood during aging in mice^39^, 15 genes of the original 32 validated in that study overlapped with the current data, (**Supplemental Figure 6**), but none were significantly associated with AL vs YL comparison, only *Tapbp* was significantly increased in AO vs YO (padj = 0.02). *Sox11* (p=0.03), and *Trhde* (p=0.05) were increased and *Cdh13* (p=0.015) decreased (*Cdh13*) in YO vs YL comparison, and only *Pax3* (p=0.004) was decreased in AO vs AL comparison.

#### Integration of Methyl-seq and RNA-seq data

To further investigate the correlation between variation of DNA methylation and gene expression, genes **(Supplemental Table 15)** or significantly differentially expressed genes (padj<0.05) **(Supplemental Table 16**) that had a distance less than 2 kb from the DMW were extracted (DMC data not shown). AO vs YO had the greatest number of significant DMW (159) and DMC (29 genes) (q<0.05) in genic regions with significant differential expression (padj<0.05) compared to all other biologically relevant comparisons (AL vs YL: 1 DMC, 2 DMW, YO vs YL: 1 DMC, 3 DMW, AO vs AL: 1 DMC, 9 DMW) (**Table 2**). The majority of differentially expressed genes contained only one DMW or DMC, and as expected from the methylome wide analysis of genic location, the majority of DMC or DMW were contained within the introns of differentially expressed genes. Also as expected more DMC or DMW sites in differentially expressed genes were hypomethylated than hypermethylated, but this difference was generally greater compared to the differential methylation when assessed across all available loci. Interestingly, however, hypermethylation of DMC or DMW was not associated with decreased gene expression except for DMW in AO vs YO, while as expected hypomethylation of DMC or DMW was more frequently associated with increased gene expression in aging, but in obesity when hypomethylation was more frequently associated with decreased gene expression (**Table 2**). To visualize these methylation and gene expression data, boxplots were generated for both the methylation data and gene expression data for the DMW that showed significant methylation difference and had a significantly differentially expressed gene nearby (distance < 2 kb) (**Supplemental Figure 7**).

**Table 2:** (A) Numbers of differentially methylated cytosines (DMC) within differentially expressed RNA biotypes for biologically relevant comparisons. (B) Numbers of differentially methylated windows (DMW) within differentially expressed RNA biotypes for biologically relevant comparisons. Genic location from Annotatr of DMC (C) and DMV (D) within differentially expressed biotypes for biologically relevant comparisons. Association of hyper-or hypomethylated DMC (E) or DMW (F) with increased or decreased RNA biotype expression.

|  | #DMC/Significantly Differentially Expressed Gene |  |  |  |  |  |  |  |
| --- | --- | --- | --- | --- | --- | --- | --- | --- |
|  | #DMC | 1 | 2 | 3 | 4 | 5 | 6 | 12 |
| AL vs YL | 1 | 1 |  |  |  |  |  |  |
| AO vs YO | 29 | 20 | 6 | 2 | 1 |  |  |  |
| YO vs YL | 1 | 1 |  |  |  |  |  |  |
| AO vs AL | 1 | 1 |  |  |  |  |  |  |

**B**
|  | #DMW/Significantly Differentially Expressed Gene |  |  |  |  |  |  |  |
| --- | --- | --- | --- | --- | --- | --- | --- | --- |
|  | #DMW | 1 | 2 | 3 | 4 | 5 | 6 | 12 |
| AL vs YL | 2 |  | 1 |  |  |  |  |  |
| AO vs YO | 159 | 45 | 36 | 6 | 3 | 0 | 0 | 1 |
| YO vs YL | 3 | 3 |  |  |  |  |  |  |
| AO vs AL | 9 | 1 | 1 | 2 |  |  |  |  |

**C**
| Genic Location (Annotatr) of Differential Methylation in Significantly Differentially Expressed Gene |  |  |  |  |  |  |  |
| --- | --- | --- | --- | --- | --- | --- | --- |
| #DMC | 1-5kB Upstream | Promotor (<1kB) | 5'UTR | CDS | Intron | Exon | 3'UTR |
| AL vs YL |  |  |  |  | 1 |  |  |
| AO vs YO | 9 | 4 | 1 | 2 | 29 | 1 | 4 |
| YO vs YL |  |  |  |  | 1 |  |  |
| AO vs AL |  |  |  | 1 |  |  |  |

**D**
| #DMW | 1-5kB Upstream | Promotor (<1kB) | 5'UTR | CDS | Intron | Exon | 3'UTR |
| --- | --- | --- | --- | --- | --- | --- | --- |
| AL vs YL |  |  |  |  | 2 |  |  |
| AO vs YO | 35 | 27 | 6 | 15 | 136 | 19 | 5 |
| YO vs YL | 1 |  |  |  | 4 |  |  |
| AO vs AL |  | 2 | 2 | 2 | 9 | 5 |  |

**E**
| #DMC | Number Hypo-methylated | Number Hyper-methylated | %Hypo-methylated | % Hypo-methylated and increased expression | % Hypo-methylated and decreased expression | %Hyper-methylated and increased expression | %Hypermethylated and decreased expression |
| --- | --- | --- | --- | --- | --- | --- | --- |
| AL vs YL | 1 | 0 | 100 | 100 | 0 | 0 | 0 |
| AO vs YO | 35 | 9 | 80 | 71 | 29 | 100 | 0 |
| YO vs YL | 1 | 0 | 100 | 0 | 100 | 0 | 0 |
| AO vs AL | 0 | 1 | 0 | 0 | 0 | 100 | 0 |

F
| #DMW | Number Hypo-methylated | Number Hyper-methylated | %Hypo-methylated | % Hypo-methylated and increased expression | % Hypo-methylated and decreased expression | %Hyper-methylated and increased expression | %Hyper-methylated and decreased expression |
| --- | --- | --- | --- | --- | --- | --- | --- |
| AL vs YL | 2 | 0 | 100 | 100 | 0 | 0 | 0 |
| AO vs YO | 123 | 56 | 69 | 76 | 24 | 64 | 36 |
| YO vs YL | 3 | 1 | 75 | 50 | 50 | 100 | 0 |
| AO vs AL | 7 | 8 | 47 | 0 | 100 | 100 | 0 |

To identify putative biomarkers for changes associated with age and obesity in ASCs, using PLS-DA analyses from both differentially methylated (DMeth) (**Supplemental Table 6**) and differentially expressed (DEx) (**Supplemental Table 14**), common elements were identified for each comparison. As previously stated, *Capn6* was the only common element between DEx Diet and DEx Age comparisons with VIP>2 and padj<0.05; however, this was not present in any of the DMeth Comparisons with VIP>2. Eight additional biotypes (*Il11, Itih2, Relt, Tnfrsf12a, Klra4, Rap1gap2, Dusp6, Gzme*) were identified in common between DEx Diet and DEx Age comparisons with VIP>1 and padj<0.05. For DMeth comparisons of AL vs YL with VIP>1 and q<0.05, there were 3 common elements with DEx Age (*Nfatc2, Tbk1, Fnip2*), and 10 with DEx Diet *(Pvt1, Parm1, Atf6, Gdap2, 2210408F21Rik, Gpr85, Mboat1, Hipk2, Aff1, Sfi1*). For DMeth comparisons of AO vs YO with VIP>1 and q<0.05, there were 160 common elements with DEx Diet comparisons with VIP>1 and padj<0.05, and 56 common elements with DEx Age comparisons with VIP>1 and padj<0.05, with an additional 2 elements (*Itih2, Dusp6*) in common with both DEx Diet and DEx Age comparisons. For DMeth YO vs YL comparison against DEx Diet, there were 2 common elements (*Actn1, Tnpo1*), and no elements in common with Age. For DMeth AO vs AL comparison, there were 45 common elements with DEx Diet with VIP>1 and padj<0.05 and 11 common elements with DEx Age with VIP>1 and padj<0.05, with an additional element in common with both DEx Diet and DEx Age (*Dusp6*). For DMeth A vs Y comparison, there were 149 common elements with DEx Diet with VIP>1 and padj<0.05 and 42 common elements with DEx Age with VIP>1 and padj<0.05, with and additional an additional 2 elements in common with both DEx Diet and DEx Age (*Itih2, Dusp6*). For DMeth O vs L comparison, there were 10 common elements with DEx Diet with VIP>1 and padj<0.05 (*Adam19, Tiam1, Mboat1, Sorcs2, Ablim1, Intu, Col12a1, Cyb5d1, Ppp2r3a, Nop56*) and 2 common elements with DEx Age with VIP>1 and padj<0.05 (*Sptb, Ifngr2*).

Finally, the list of elements identified as significant in each analysis, comparison, or module were combined into a single table, and the number of times each element appeared in ≥4 analyses and comparisons was recorded and rank ordered (**Figure 13, Supplemental Table 17, and Supplemental Table 18**). Consistent with other analyses, for the AL vs YL comparison, few elements appeared multiple times, with *Nfatc2* appearing a total of 8 times (**Figure 13A**). In contrast, for the AO vs YO comparison (**Figure 13B**), a number of genes appeared in 9 analyses (*Akap13, Baiap2, Cd44, Gm15663, Jup, Msx1, Noct, Rarg, Rhobtb1,* and *Slc35e4*). As suggested from the previous analyses described here, there were fewer elements identified recurrently for the YO vs YL comparison (**Figure 13C**) (*Actn1* and *Tnpo1* present in 7 analyses each) than for the AO vs AL comparison (*Gbx2* (7), *Fam83f* (7), *Adh7.* (6) *Tnfsf11* (6)) (**Figure 13D**). For the A vs Y comparison (**Figure 13E**), the number of elements appearing in multiple analyses increased, with *Dusp6* appearing 11 times, followed by *Itih2, Jup, Noct, Slc35e4, Stard13, and Stat5b* each appearing in 10 analyses. For the O vs L comparison (**Figure 13F**), the number of elements appearing in multiple analyses was more diffuse, with *Ablim1* (9), *Cyb5d1* (9), *Adam19* (8) *Col12a1* (8), *Dhrs9* (7), *Intu* (7), *Nop56* (7), *Pax3* (7), *Slc2a9* (7), *Slc38a4* (7), *Sorcs2* (7), and *Tspan5* (7) the most common recurring elements. Finally, when all comparisons were considered together, (**Figure 13G**) *Dusp6* was identified in 35 analyses, and *Aff1, Nfatc2, Itih2, Start13, Baiap2, Noct, Pvt11, Stat5b, Cd44,* and *Tbx3* were all identified in at least 25 analyses.

**Figure 13:**
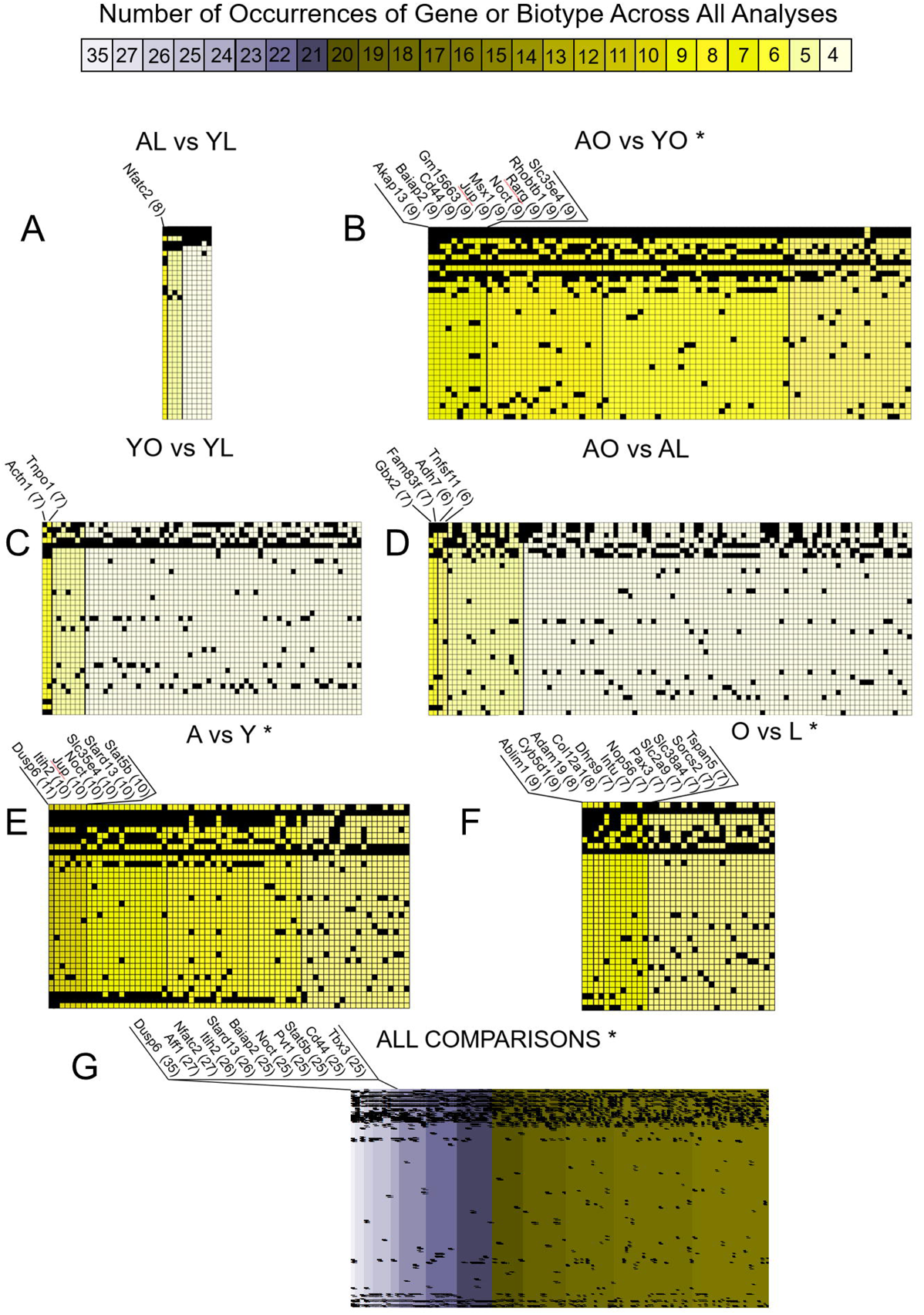
‘Top Hit’ summary of the rank order frequency of genes and biotypes occurring across all analyses performed in the current study for A) AL vs YL, B) AO vs YO, C) YO vs YL, D) AO vs AL, E) A vs Y, F) O vs L, and G) All Comparisons. Columns represent rank order of frequency then alphabetized biotype names, rows represent individual analyses and comparisons. Number in parentheses after individual biotype abbreviation is number of times that specific biotype appears across all analyses. * Represents truncation of table for elements appearing in fewer analyses for clarity. Full dataset available for all elements appearing >=4 analyses in **Supplemental Table 18.**

## Discussion

We identified substantial global DNA hypomethylation with aging or obesity using RNA-Seq, Methyl-Seq and integrated RNA-Seq and Methyl-Seq analyses in murine ASCs from lean and obese mice at 5- and 12-months of age, and identify an apparent additive or synergistic effect of aging combined with obesity. In contrast, the transcriptome of aging lean mice (AL vs YL), was remarkably stable with only seven differentially expressed genes. PLS-DA and WGCNA were helpful in ranking DMW and genes according to possible importance as biomarkers, and in identifying modules of related genes, respectively. Functional analysis of pathways identified by PLS-DA and WGCNA provided substantial insight into potential upstream mechanistic networks and potential diseases and functions associated with these changes. When these and other analyses were evaluated across all biologic comparisons, relatively few recurrent genes were identified in multiple comparisons; many of these were genes already known for their critical roles in progenitors and in diseases of obesity and aging. Therefore, these genes are considered to be potential drivers capable of priming ASCs for dysfunction with aging and obesity.

ASCs have important roles in tissue homeostasis and immunomodulation and pathophysiologic roles in enhancing proliferation and cancer stem cell-like properties[reviewed in ^40^],^41; 42^ in sensing the metabolic environment, and in acting upstream of inflammatory responses.^43^ The substantial interest in their use in regenerative medicine applications demands a better understanding of the factors that drive ASCs from their physiologic role to a pathophysiologic role *in situ*, and in understanding mechanisms that diminish their evolving therapeutic importance in regenerative medicine. Notably, the regenerative capacity of stem cells is attenuated if the cells are harvested from aged patients with chronic disease.^44^ As in obesity, age-related changes to ASCs and other mesenchymal stem cells (MSCs) reduces their regenerative potential, and ASCs and MSCs are sensitive to environmental obesogens.^45–48^ Further, MSCs from obese human donors have impaired expression of adhesion markers (CD29, CD44, CD166), MSC markers (CD73, CD90, CD105), increased expression of epithelial marker CD31, and have impaired proliferation, decreased osteogenic and adipogenic differentiation, increased senescence and evidence of endoplasmic reticulum stress.^49^ In contrast, our previous data demonstrate enhanced adipogenesis and osteogenesis, but impaired chondrogenic differentiation in murine ASCs from high-fat diet-induced C57BL6/J obese mice.^17^ Overfeeding of neonatal mice before weaning induces metabolic re-programming of ASCs towards increased adipogenic differentiation, and decreased osteogenic differentiation,^50^ demonstrating that dysregulation in obesity can begin early in life, which implicates a role for epigenetic regulation.

DNA methylation at the 5’ site of cytosine is a critical component of the heritable and modifiable epigenetic landscape. It contributes to precise fine-tuning and precision of gene expression patterns that are required for stem cell viability, proliferation, pluripotency, differentiation, transdifferentiation, and reprogramming.^51^ The global hypomethylation we identified in aged or obese mice, with an apparent synergistic or additive effect in both aged and obese mice, was primarily in inter-CGI regions and with regard to genic location, primarily intronic followed by intergenic regions. The greater number of differentially expressed biotypes in AO vs YO compared with AL vs YL suggests greater dysregulation of epigenetic co-factors and machinery in obesity compared to aging in ASCs from lean animals. CpG islands represent a minority of the genome; the majority of CpG are found in repetitive DNA elements, located in intergenic regions.^52^ While much about the function of these repetitive elements is unknown, they are critical for chromosomal organization and function, for example forming centromeres and telomeres.^52^ Repetitive elements are notoriously unstable, and are frequently a source of mutations but epigenetic mechanisms have evolved to effectively improve the stability of these regions.^52^ Thus, widespread intergenic and inter-CGI hypomethylation is expected to reduce the efficacy of these epigenetic mechanisms, and reduce stability of repetitive elements, thus predisposing to mutations and dysfunction. This instability induced by widespread hypomethylation is a key factor in a rapidly expanding range of devastating diseases, including the chronic effects of environmental factors.^52^ Combined with the abnormalities of genic region methylation and RNA biotype identified, this intergenic, inter-CGI hypomethylation could be devastating for normal ASC function. In support of our findings, human ASCs from obese (BMI 32.6LJ±LJ2.2LJkg/m^2^, age 34.3LJ±LJ7.4 years) compared to lean (BMI 22.4LJ±LJ12LJkg/m^2^, age 44.3LJ±LJ9.2 years) donors demonstrated global hypomethylation,^53^ even though the obese population in this study was slightly younger overall than the general obese population. Similarly, whole blood samples^54^ and bone marrow-derived MSCs [reviewed^55^] demonstrate global demethylation with age, and this demethylation was predominantly distributed away from CGI. Also in agreement with our findings, most of the differential methylation was also overwhelmingly in non-CGI regions in obese ASCs within transcribed regions of gene bodies followed by intergenic regions,^53^ whereas we have identified enrichment more specifically in intronic regions followed by intergenic regions. While the precise epigenetic mechanism of this global hypomethylation remains to be determined, obese porcine ASCs had reduced global DNA hydroxymethylation and H3K4m3 marks, whereas H3K9me3 and H3K27me3 marks were unchanged compared to lean porcine ASCs,^56^ but the age of the pigs was not considered as a factor in these studies. These hydroxymethylation and H3K4m3 findings were associated with genomic transcriptional repression.^56^ In the current study, DNA hypomethylation was generally associated with increased gene expression with aging, as expected. However, in obesity, DNA hypomethylation was associated with *decreased* gene expression and no association between DNA hypermethylation and decreased gene expression was observed. These intriguing findings are also consistent with our observation that obesity moderated the number of RNA biotypes with large changes in gene expression, particularly for the increase in expression of RNA biotypes we observed with aging. Our finding that despite the extent of DNA methylation changes observed, ‘healthy’ aging (AL vs YL) maintained a remarkably stable transcriptome, with only seven differentially expressed genes is in line with the findings of metanalyses of gene microarray studies across a range of tissues.^57^ Also in line with previous studies,^57^ is the preponderance of over- rather than under-expression in ‘healthy’ aging (AL vs YL). As expected therefore, for this comparison, changes in gene expression were not associated with significant effects on function, or on upstream regulators of any pathways – networks involved included those expected for aging, cellular growth, proliferation, death, and survival as seen previously in ASCs derived from old donors.^58^ We were surprised to discover however, that all comparisons involving obesity identified dozens of impacted potential upstream regulators, networks, functions and diseases, many of which have been extensively associated with diseases for which obesity is a risk factor.

The lack of age-related epigenetic drift using existing methylation markers of aging was surprising, even when evaluating similar markers in multiple other species or strains of mice.^39; 59^ However, our results were consistent with data from muscle stem cells from young (1.5-2.1 months) and old (23.3-27 months) mice, a wider age range than we evaluated.^60^ The slight increase in DNA methylation age identified in the muscle stem cells lagged far behind the chronological age of the mice. Current DNA methylation aging ‘clocks’ are developed with data from bulk tissues, rather than stem cell populations, and since stem cells generally represent a very small proportion of the overall cellularity of a tissue, it is possible that differentiated cell populations drive the characteristics of the epigenetic aging ‘clock’ at a tissue level. If similar findings are true for stem cells from a variety of sources over the full lifespan, accurate matching of stem cell and chronological age may entail development of an entirely different set of methylation marks to account for their apparently slower rate of epigenetic aging. Given the findings of this study and the diseases and functions identified as modified by obesity and aging in ASCs, this could be useful in a similar way to screen individual donor ASCs derived from subcutaneous adipose tissue for risk of disease or dysfunction, similar to how markers of epigenetic age are being used.^61^

Non-codingRNA play a critical role in biological regulation, obesity,^62^ in the beneficial effect of caloric restriction,^3^ in identifying disease sub-phenotypes,^63–65^ and in the modification of the epigenetic landscape.^66^ Reducing the complexity of differential expression data using WGCNA extracted clusters of highly co-expressed genes and ncRNA biotypes related to age, diet, or bodyweight at sacrifice. Many of the loci of ncRNA biotypes were close to those of in-module mRNA biotypes or were associated with another mRNA biotype contained within another module. Thus, ncRNA biotypes could provide a level of connectedness between different clusters or functions of modules. This connectedness reduced with obesity providing a potential mechanism of ncRNA and regulatory pathway dysregulation in obesity whereby there could be an overall reduction in ncRNA regulation. The modules significantly associated with diet and age had a wide range of diseases and functions, but the top hit for every module was related to tumors, neoplasia, or to oncogenesis. Interconnectedness of modules associated with age and obesity examined identified *Tgfb1, Mcl1*, (anti-apoptotic gene) and *Mapk14* as highly interconnected regulatory factors related to the transcriptional changes, whereas the same analysis for body weight at sacrifice associated with high-fat diet or control diet found *HLA-A, Gas6,* (regulation of cell proliferation) and *Nedd9* (focal adhesion protein) to be the most highly interconnected. Together with the differences in top canonical signaling pathways identified for the two comparisons, this provides a useful starting point for future studies evaluating how bodyweight gain in response to a specific diet influences disease risk. Together, our data support previous findings of WGCNA in subcutaneous fat suggesting obesity could accelerate the aging process,^67^ even though functions associated with lipid and carbohydrate metabolism and the immune system and inflammatory response were the dominant pathways involved with aging in subcutaneous adipose tissue. Interestingly, a study of sex-related differences in subcutaneous adipose tissue in obesity that specifically included ncRNA identified a large number of differentially expressed ncRNA biotypes. NcRNAs were over-represented in epigenetic and transcriptional regulatory pathways, and dysregulation of ncRNA species was identified as potential link with cancer and neurodegenerative diseases in sex-related differences in obesity, similar to the interactions of age and obesity in the current study, and similarly suggesting a role for ncRNA in changes in expression of obesity-related mRNA biotypes.^68^ Therefore, improved understanding of the emerging functions of ncRNA in the modification of the epigenetic landscape^66^ merits further study. With additional more comprehensive study of the role of ncRNA in aging and obesity of ASCs, and in sex-differences between ASCs, WGCNA could identify potential targets and interactions for modulation.

In contrast to the WGCNA approach, VIP scores generated from PLS-DA allowed us to identify individual variables. These variables can separate the experimental groups as potential biomarkers for the effects of diet and age in both the Methyl-Seq and RNA-Seq data individually, in combination, and to compare between the variables that best distinguished the groups in Methyl-Seq to differential gene expression. *Mapt, Nr3c2, App, and Ctnnb1* emerged as potential hypomethylated upstream regulators in both aging and obesity (AL vs YL and AO vs YO), and *App*, *Ctnnb1, Hipk2, Id2,* and *Tp53* with the additional effects of aging in obese animals, with *Foxo3* and *Ccnd1* as potential hypermethylated upstream regulators of healthy aging (AL vs YL), but also for the effects of obesity in young animals (YO vs YL), suggesting that these factors could play a role in accelerated aging with obesity. As expected, based on global hypomethylation data, there appeared to be loss of variables with differential expression and hypermethylation in obesity combined with aging, with only *Foxo3* and *Ccnd1* associated with both healthy aging and with obesity in young animals. All of the functions of these upstream regulators were consistent with disease and function terms associated with cancer, neurodegenerative diseases, and abnormalities in the cell cycle.

PLS-DA analysis of RNA-Seq data identified nine genes with VIP>1 for both age and diet that were also differentially expressed emerged: *Capn6, Dusp6, Gzme, Il11, Itih2, Klra4, Rap1gap2, Relt*, and *Tnfrsf12a.* Interestingly, none of these potential biomarkers for both age and diet with differential gene expression were identified as transcriptional regulators, but when considering differentially expressed biomarker candidates with VIP≥2 for diet, transcriptional regulators were identified (i.e., *Cdkn2a, FoxG1, Ncoa3, Pax3, Plagl1,* and *Plag1*). For age by similar filters, 9 transcriptional regulators were identified, *En2, Gzf1, Shox2, Sox12, Tfec, Vhl, Zbtb18, Zfp78,* and *Zfp992*. Once again, several of these genes have been previously associated with ASC dysfunction, or with many of the diseases and functions identified in pathways analysis.

From all analyses used to evaluate the data, no RNA biotypes consistently emerged recurrently across datasets or comparisons between the groups. However, the disease and functional effects of obesity and age on ASCs were remarkably consistent. This, together with the high level of connectivity between hub genes, lead us to ask if potential driver genes could be identified based on the degree of their recurrent appearance in different analyses and between group comparisons, rather than necessarily appearing as a ‘top hit’ appearance in a specific module or single analysis. The current definition of driver gene relates mostly to cancer – a gene whose mutations increase cell growth under specific intracellular microenvironmental conditions.^69^ Here we did not sequence individual ASCs for clonal mutations already associated with malignancy or with stem cell dysfunction, nor would the genetic mutation rate in these baseline conditions be expected to be elevated above stochastic low levels. Therefore, tumor datasets used to identify cancer genes are not likely to be relevant in the current study and many of the driver gene prediction tools developed for cancer are unlikely to be useful; individual analyses revealed differential methylation and differential expression of many RNA biotypes already associated with cancer. There are currently no established frameworks for identifying epigenetic drivers in cancer or any other discipline,^70^ or their effect on downstream pathways.^71^ However, given that epigenetic aberrations can themselves impact the mutation rate and genetic instability in carcinogenesis,^71^ we attempted to determine RNA biotypes that could ultimately be responsible for driving ASC dysfunction down many of the identified pathways. We attempted to identify a pre-primed driver ‘switch’ for ASC dysfunction on the premise that epigenetic ‘priming’ can lead to cancer development.^72^ For example, could we identify a critical switch if the local ASC niche became permissive of such dysfunction *in situ*, if the ASCs were implanted into a dysfunctional niche, if the ASCs were exposed to exogenous signaling or differentiation cues, or if the ASC dysfunction attained a level capable of influencing the niche itself, and of function of surrounding cells? To begin to answer this question, we used the general approach applied to identification of cancer driver genes, to evaluate the data using multiple robust methods and identify genes predicted by more than one method,^69^ by looking for RNA biotypes that appeared as significant or important recurrently in multiple analyses and in multiple comparisons. When the data from all analyses and comparisons were manually curated, and rank ordered by frequency of occurrence (**Figure 13, Supplemental Table 18)**, the number of individual RNA recurrent biotypes was low for healthy comparisons, highlighting *Nfatc2* in healthy aging of ASCs. *Nfatc2* is a calcium-regulated member of the Nuclear Factor of Activated T-cells transcription factor family that regulates a number of key cellular functions including differentiation and adaptation,^73^ cytokine gene expression in immune responses,^74^ sarcomas and round cell sarcomas,^75; 76^ and adipogenesis.^77^ Moreover, *Nfatc2* is a key regulator of chondrogenesis including in MSCs,^78^ contributes to normal cartilage homeostasis, and is regulated through epigenetic mechanisms.^79^ Additionally *Nfatc2* demonstrates age-dependent increases in expression in other cell types.^80^ *Nfatc2* therefore makes ‘biological sense’ as a potential driver of aging in ASCs.

As suggested from our individual analyses, the addition of obesity to aging increased the number of RNA biotypes occurring recurrently throughout the entire dataset. When rank-ordered by frequency of appearance for all group comparisons, *Dusp6* was identified in 35 analyses, and *Aff1, Nfatc2, Itih2, Stard13, Baiap2, Noct, Pvt1, Stat5b, Cd44,* and *Tbx3* were all identified at least 25 times, of which *Dusp6, Nfatc2, Itih2, Stard 13, Baiap2, Noct, Stat5b,* and *Tbx3* were all also differentially methylated and differentially expressed in at least one biologic comparison. All of these mRNA biotypes had known functions consistent with putative roles as drivers of ASC priming to dysfunction and disease with aging and obesity: Depletion of *AFF1* improves osteogenic differentiation of MSCs via inhibition of DKK1, an inhibitor of Wnt/β-catenin signaling, whereas overexpression impairs osteoblast differentiation,^81^ and *AFF1* depletion also enhances adipogenesis.^82^ In combination with *KMT2A* overexpression, *AFF1* is associated with poor prognosis in acute lymphoblastic leukemia.^83^ *ITIH2* was recently identified as a core gene for colorectal cancer liver metastasis, with overexpression associated with poor survival,^84^ and as a hub gene in metastatic uveal melanoma,^85^ whereas in multiple other primary solid tumors, downregulation of *ITIH2* is identified.^86^ Elevations of serum ITIH2 are associated with metabolic dysfunction in obese dogs compared to those without metabolic dysfunction.^87^ STARD13 is a GTPase activating protein for Rho GTPases, and is a tumor suppressor that regulates cell survival and apoptosis. Its role in normal Rho GTPase function also means dysregulation impacts tumor invasiveness and metastasis^88^, while downregulation of *Stard13* in combination with *Stat3* and *Casp2* contributes to the anti-aging effects of caloric restriction on the liver.^89^ BAIAP2 (IRSp53) is part of a ubiquitin-dependent switch that stabilizes cortical actin at points of cell-cell contact to align membranes and facilitate cell fusion at key points in development, a process that is key for formation of osteoclasts, myofibers (including cardiac), giant cells, and placental development,^90^ and deletion is embryonic lethal, while deletion in adult mice demonstrate failure to regulate synaptic plasticity and demonstrate learning deficiencies associated with the hippocampus.^91^ Overexpression of *Noct*, a gene essential for regulating circadian rhythm output in enterocytes and bone, is expressed in antiphase to IGF-1 but is nonrhythmic in white adipose tissue under normal feeding states, despite its interactions with PPARLJ. However, in conditions of restricted feeding, *Noct* becomes rhythmic. *Noct* deletion confers resistance to high-fat diet induced-obesity, and conversely, *Noct* overexpression reduces fat mass in male mice under normal diet conditions and reduces adipocyte size, through its de- adenylase activity and mitochondrial actions.^92^ *Pvt1* is a long non-coding RNA regulated by p53 that has been identified as pro-fibrotic^93^, and an oncogene, with over-expression or high copy number associated with many types of cancers, and with negative prognosis. *PVT1* has multiple variants in human, some of which are linear, and some of which are circular, but both of these transcript-types control cell proliferation via effects on MYC and CDKN1A targets, facilitate tumor cell invasion and metastasis by promoting epithelial-mesenchymal transition, and both have anti-apoptotic functions.^94^ On the other hand, circular *PVT1* expression is reduced in senescent human tendon progenitors,^95^ but inhibition of other isoforms of *PVT1* is associated with reduced senescence and increased proliferation.^96^ *PVT1* isoforms also interact with Notch signaling, pathways with wide-ranging effects on cell proliferation, development, differentiation and homeostasis.^97^ Constitutive activation of *Stat5b* promotes adipogenesis along with *Stat5a* via *PPAR*_γ_,^98^ and is a growth hormone signaling intermediate in the regulation of postnatal growth and adiposity.^99^ CD44, the well-known MSC marker^100^ and receptor for hyaluronan, is also a cancer stem cell marker associated with drug resistance, epithelial-mesenchymal transition, metastasis, and survival.^101^ *Tbx3* is dynamically regulated during maintenance and induction of pluripotency in embryonic stem cells,^102^ and is a critical regulator in development. In the adult, *TBX3* is frequently overexpressed in epithelial and mesenchymal cancers and is a regulator of cancer stemness,^103^ and is a critical regulator of pluripotency and in induced pluripotency.^103^

The most frequently recurrent gene*, Dusp*6 is associated with transition between naïve and primed pluripotency states in mouse embryonic stem cells, is part of core pluripotency circuitry, acts upstream of *Oct4,* and *Sox2,* and as such has been described as a ‘guardian’ of pluripotency.^104; 105^ *Dusp6* is in the MAP kinase phosphatase family and reduces the high levels of basal ERK signaling and pulses typically present in stem cells to the low levels of ERK signaling in differentiated cells,^106^ thus *Dusp6* promotes early differentiation at the time of commitment,^107^ acting as a switch that transitions stem cells to differentiated cells. *Dusp6* is a key regulator in many other processes and tissues, and in cellular stress responses.^108^ In primary adipocytes, *Dusp6* expression is greater in high-fat diet obese mice compared to lean controls, and *Dusp6* knock out impairs glucose tolerance, and confers resistance to obesity, depending on genetic background.^109^ *Dusp6/8* knockout in mice confers resistance to diet-induced obesity and improves metabolic parameters via increases in ERK1/2 phosphorylation suggesting a critical role for *Dusp* regulation of ERK 1/2 in metabolism.^110^ *Dusp6* plays a critical role in the methylation landscape, and is critical for completion of global demethylation in naïve embryonic stem cells, re-setting the epigenome.^111^ *Dusp6* has other epigenetic effects, for example *Dusp6* inhibition is also one effect of Hdac3 inhibitors, and *Dusp6* has partial anabolic effects on Hdac3 depleted chondrocytes.^112^ Finally, *Dusp6 i*s also important in p53 mediated cell senescence, with *Dusp6* expressed at greater levels in senescent cells; Erk1/2 being the major kinase that controls cell proliferation.^113^ Thus, *Dusp6* could be a key driver of both epigenetic, transcriptional, and functional results of obesity and aging in ASCs, and awaits further study. While it was somewhat surprising that differential expression and differential methylation was not identified for more than one comparison, the *Dusp6* protein and mRNA have very short half-lives, and stability of these species was not investigated here. ^114^

This study highlights the substantial interactions between one inevitable intrinsic factor, and one highly prevalent extrinsic factor across part of the life course. Identifying the epigenetic drivers of other life-course factors, their interactions, and the pathways impacted by these drivers could provide targets to mitigate the subsequent genetic instability or induced dysfunction. For example, given the critical role ASCs play in the tumor microenvironment in a variety of cancers and cancer metastases, [reviewed^40^], understanding and mitigating epigenetic and transcriptomic disruption in response to intrinsic and extrinsic risk factors could be critical to preventing cancer development and progression. Further, a large proportion of the ASC literature is focused on their potential for treatment or immunomodulation of several diseases. While some limitations of ASCs in this role have been identified for age, sex and adiposity, the effects of other factors remain largely unexplored. Finally, data from this study suggest epigenetic and transcriptomic risks of the life course on ASCs could explain some of the potentially devastating side effects of using unproven stem cell interventions that have recently been highlighted,^115^ together with unexpected failures in clinical trials compared to preclinical studies.^116^

## Limitations

ASCs are not a homologous population and others have begun to establish the diversity of ASC populations, using single cell RNA sequencing (scRNA-Seq).^117^ With this in mind, we do not know if the shifts identified here result from changes within the cell population homogeneously, from changes in abundance or function of individual cell populations, or universally within cell populations currently defined as ASCs but further study to perform paired scRNA-Seq with the bulk RNA-Seq data evaluated here from the same individuals could resolve this question. We were therefore unable to evaluate ASC subset changes recently identified using scRNA-Seq in visceral adipose tissue in obese mice^118^ but many of the biological pathways identified in the current study were also previously reported by scRNA-Seq analysis. Transcriptional variability generally increases with age in scRNA-Seq studies^119^ and even though we used a homogeneous cell population sorted by cell surface markers for ASCs, it is possible that the bulk DNA methylation and transcriptomic data evaluated here masked some heterogeneity within a relatively uniform cell population. Clearly, however, transcriptional heterogeneity is substantially increased in obese compared to lean mice. A potential limitation of the work described here is the use of murine ASCs, with the associated limitations of an isogenic background and homogenous laboratory animal environmental conditions and exposome. However, this model system does allow the effects of individual intrinsic and extrinsic factors and their interactions to be evaluated specifically. Our hope is that with future studies, a complex map of the exposome could be built to begin to understand how the exposome of ASC and MSC donors interacts. Indeed, the data presented here highlight an additional area of dissonance between mouse preclinical studies and human clinical outcomes in regenerative medicine that have become increasingly evident ^116^ - that intrinsic and extrinsic variables of the exposome which we shown disrupt both the methylome and the transcriptome require a multi-factorial modeling approach within in the tightly controlled laboratory animal environment to fully elaborate the effects of each individual factor. Nonetheless, evaluation of these variables in carefully controlled combinations in future multi-dimensional studies could substantially advance our understanding of how the exposome could impact regenerative therapies, through more advanced application of the computational approaches tested here, both in larger and more complex future independent mouse datasets and through mining of open source large datasets where the exposome of the human donors is likely unknown.

The effect of expansion of the mASCs to Passage 2 on the methylome and transcriptome is unknown, but is of translational relevance. While expansion represents a change from the native environment, industry sponsored allogeneic MSCs allow manufacture of up to 1 million doses per donor;^116^ thus, data from expanded ASCs are translationally relevant for treatment, transplantation, regenerative medicine and tissue engineering applications. Human bone-marrow derived stem cells undergo genome-wide demethylation with long term culture (Passage 3-12). In this study all Passage 2 cells from these isogenic animals were subjected to similar isolation and expansion protocols suggesting that the differences identified resulted from the underlying biological phenotype.

## Conclusions

Our data show that while global hypomethylation of ASCs occurs with aging, the transcriptome remains remarkably stable. This global hypomethylation is exacerbated and becomes dysregulated with obesity; simultaneously the number of dysregulated RNA biotypes in the transcriptome increases more than 120-fold in obese mice with aging compared to lean mice. Multiple analyses applied here allude to the potential importance of this dysregulation in diseases associated with obesity and aging, and for the potential of driver genes in these ASCs to cause this dysregulation. These driver genes are particularly important to validate in further studies given the widespread distribution of ASCs within their niche, their role in many diseases, and in regenerative medicine strategies.

## Acknowledgments

The authors acknowledge funding from the Duke Pepper OAIC P31 AG028716-08, a gift from Ruby E. White to Duke University School of Medicine, NIH grants AR 059784 (DL), AR065764 (DL), AR 073882 (DL), AG 046927 (FG), AG 015768 (FG), AR 075899 (C-LW), and funds from the Department of Basic Medical Sciences, Purdue University College of Veterinary Medicine. The authors would like to acknowledge invaluable technical assistance from Meredith Bostrom and Garron Wright (DHMRI) for sequencing.

## Data Availability Statement

The raw data that support the findings of this study are openly available in: (Currently in progress in Gene Expression Omnibus (GEO) link will be available shortly, embargo released upon acceptance for publication). All other data are available as supplemental tables and figures.

## Author Contributions

Little D, Guilak F, Thimmapurum J, Wu C-L, Gregory SG, and Corcoran D conceived and designed the research. Little D, Guilak F, Thimmapurum J, Wu C-L, Abramson K, Xie S, and Choudhari S performed the research and acquired the data. Little D, Thimmapurum J, Guilak F, Xie S, and Choudhari S analyzed and interpreted the data. All authors were involved in drafting and revising the manuscript.

## Supplemental Figure Legends

**Supplemental Figure 1:** Representative electropherograms from end-repaired DNA quality control using a 2100 BioAnalyzer DNA1000 Chip (**A**) and Agilent 2200 TapeStation and Tapestation HS D1000 ScreenTape (undiluted) (**B**). Representative electropherograms from adapter-ligated DNA using 2100 BioAnalyzer DNA1000 Chip (**C**), and Agilent 2200 TapeStation and Tapestation HS D1000 ScreenTape (undiluted) (**D**). Representative electropherograms final indexed sample DNA using 2100 BioAnalyzer High Sensitivity DNA Assay (**E**), and Agilent 2200 TapeStation and Tapestation HS D1000 ScreenTape (undiluted) (**F**).

**Supplemental Figure 2:** (A) Differentially methylated windows (DMW) and (B) Differentially methylated cytosines (DMC) for AL vs YL comparison with >10 standard deviation difference from mean, (C) 32 differentially methylated sites previously identified as components of methylation clock did not separate AL from YL. Similarly (D) DMW and (E) DMC for AO vs YO comparison with >10 standard deviations from mean, and (F) 32 differentially methylated sites previously identified as component of methylation clock also did not separate AO from YO.

**Supplemental Figure 3:** UpSet plots for weighted correlation network analysis (WGCNA) modules associated with diet and age ordered by r-value with ‘Top 20’ Disease and Functions from Ingenuity Pathways Analysis. Diseases and Functions are identified and ranked by significance and by predicted activation z-score. Set size represents number of in-module biotypes associated with that disease or function. Modules with a positive (A) or negative (B) association with age, modules with a (C) negative association with diet, and modules with a (D) negative and positive (E) association with bodyweight at sacrifice in aged mice.

**Supplemental Figure 4:** Differentially expressed RNA biotypes with age with variable importance in projection (VIP) scores ≥ 2 and p<0.05.

**Supplemental Figure 5:** Differentially expressed RNA biotypes with diet with variable importance in projection (VIP) scores ≥ 2 and p<0.05.

**Supplemental Figure 6:** Expression of 15 genes identified by PLS-DA previously associated with age-associated methylation drift PMID: 28912502 *p≤0.05.

**Supplemental Figure 7:** Box plots of methylation data and gene expression for differential methylated windows (DMW) for the biological comparisons that showed both significant methylation differences and significant difference in RNA biotype expression within 2kb.

## Supplemental Tables

**Supplemental Table 1:** Coverage of individual sample DNA methylome obtained using the targeted bisulfite sequencing platform.

**Supplemental Table 2:** Differential methylated cytosine (DMC) identified for the various biologically relevant comparisons.

**Supplemental Table 3:** Differential methylated windows (DMW) identified for the various biologically relevant comparisons.

**Supplemental Table 4:** Summary of number of hypomethylated DMC and DMW, and comparison of the number of DMC located in DMW for the various biologically relevant comparisons.

**Supplemental Table 5:** Variable importance in projection scores (VIP) following PLS-DA modeling of DMW for the various biologically relevant group comparisons.

**Supplemental Table 6:** Combined results of PLS-DA on DNA methylation data and comparison of RNA biotype expression for the various biologically relevant group comparisons.

**Supplemental Table 7:** Transcript biotypes by number, significance (padj<0.05), direction change in expression, and > or < 2-fold expression changes for the different biologically relevant group comparisons.

**Supplemental Table 8:** Biological relevance of differential gene expression attributed using Ingenuity Pathways Analysis for the various biologically relevant group comparisons.

**Supplemental Table 9:** Outcomes of DESeq2 used to examine the interactions between age and diet.

**Supplemental Table 10:** Results of WGCNA for RNA biotypes presented by module when evaluated for age and diet.

**Supplemental Table 11:** Results of WCNA for RNA biotypes presented by module when evaluated for bodyweight at sacrifice.

**Supplemental Table 12:** Summary of RNA biotypes present in each module significantly associated with age or diet.

**Supplemental Table 13:** Summary of RNA biotypes present in each module significantly associated with bodyweight at sacrifice.

**Supplemental Table 14:** Variable importance in projection scores for RNA biotypes following PLS-DA modeling for the effects of age and diet.

**Supplemental Table 15:** Summary of differential methylation and RNA biotype expression correlations for various comparisons of biological groups.

**Supplemental Table 16:** Summary of differential methylation and significantly differentially expressed RNA biotype expression correlations for various comparisons of biological groups.

**Supplemental Table 17:** List of elements identified as significant in each analysis, comparison, or module compiled into a single table, then rank ordered by frequency, followed by alphabetic sorting. Each element had to appear in ≥4 analysis to appear in this table.

**Supplemental Table 18:** Formatted from Supplemental Table 17, and the complete version of Figure 13.

