## Supplementary figures and images for "Aging and Obesity Prime the Methylome and Transcriptome of Adipose Stem Cells for Disease and Dysfunction"

### Supplemental Figure 1

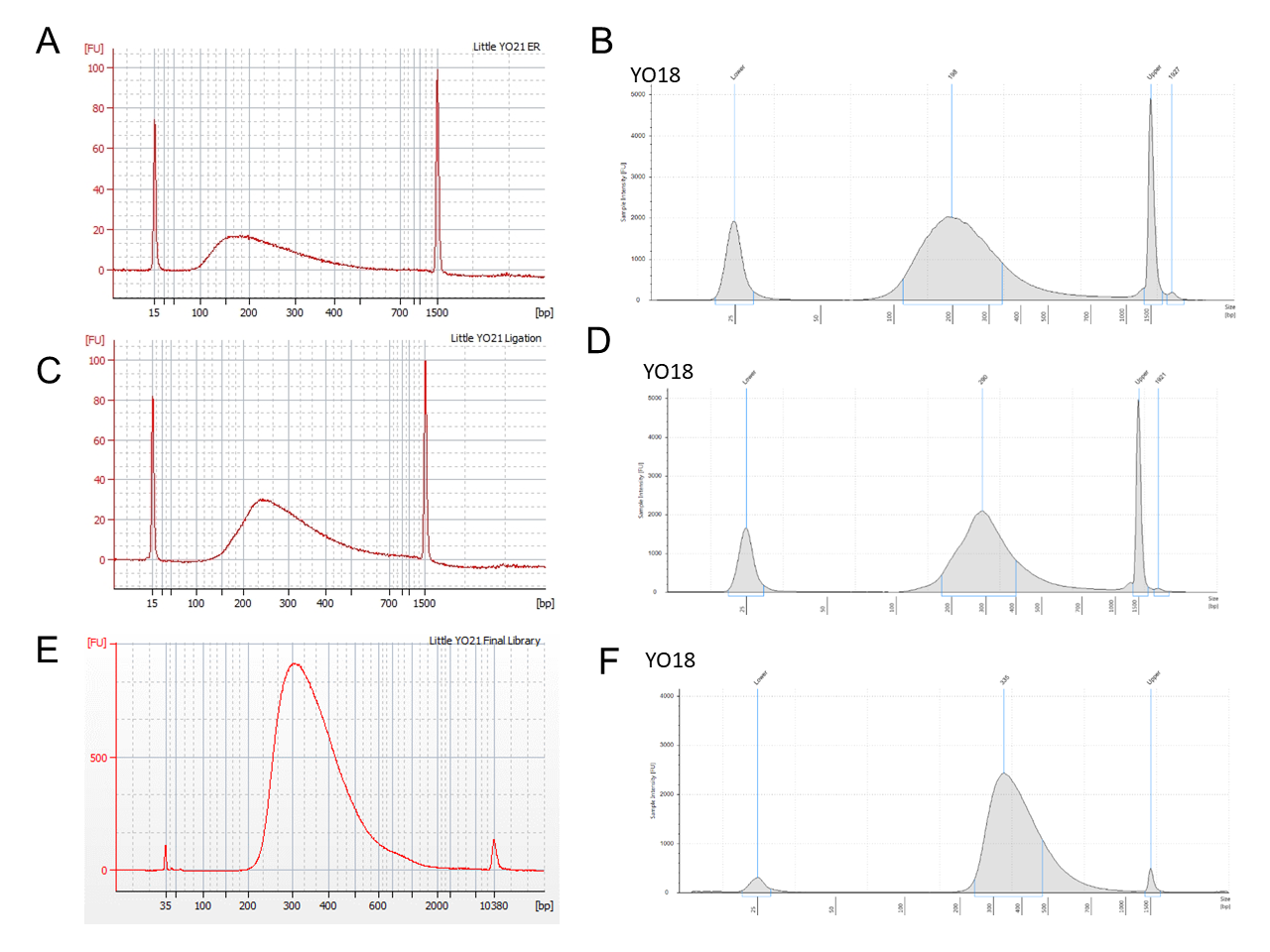

### Supplemental Figure 2

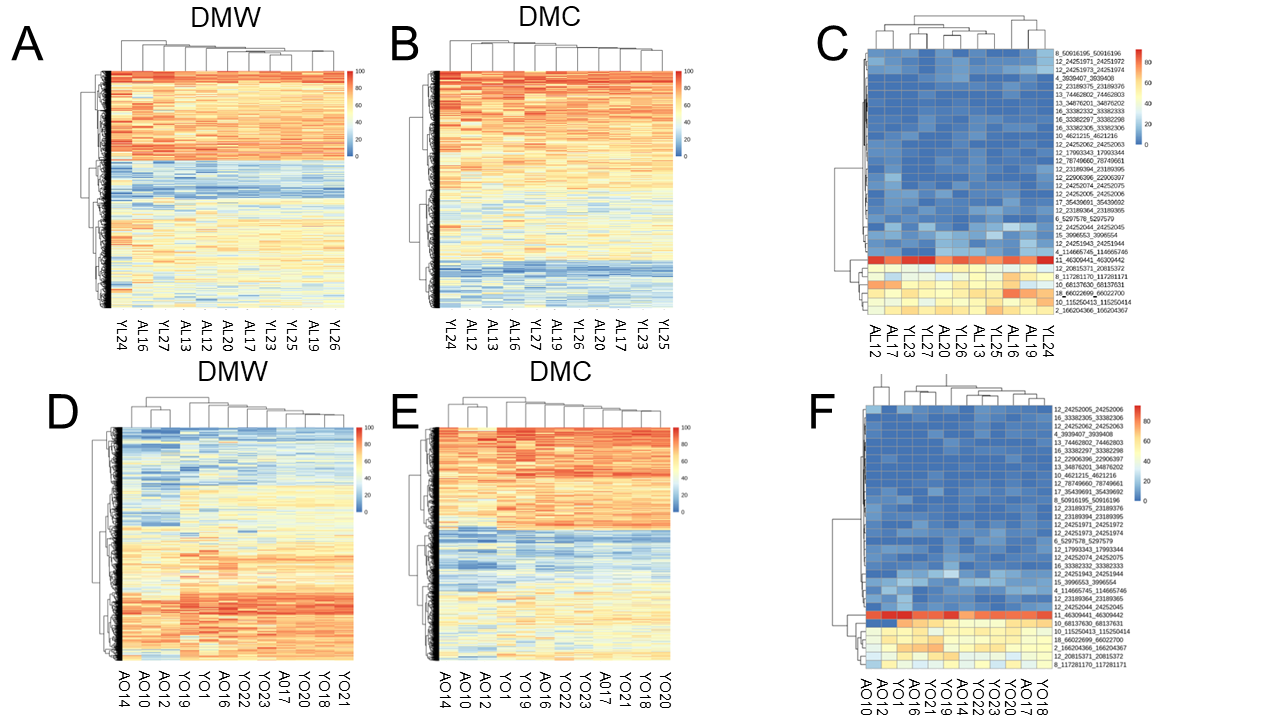

### Supplemental Figure 6

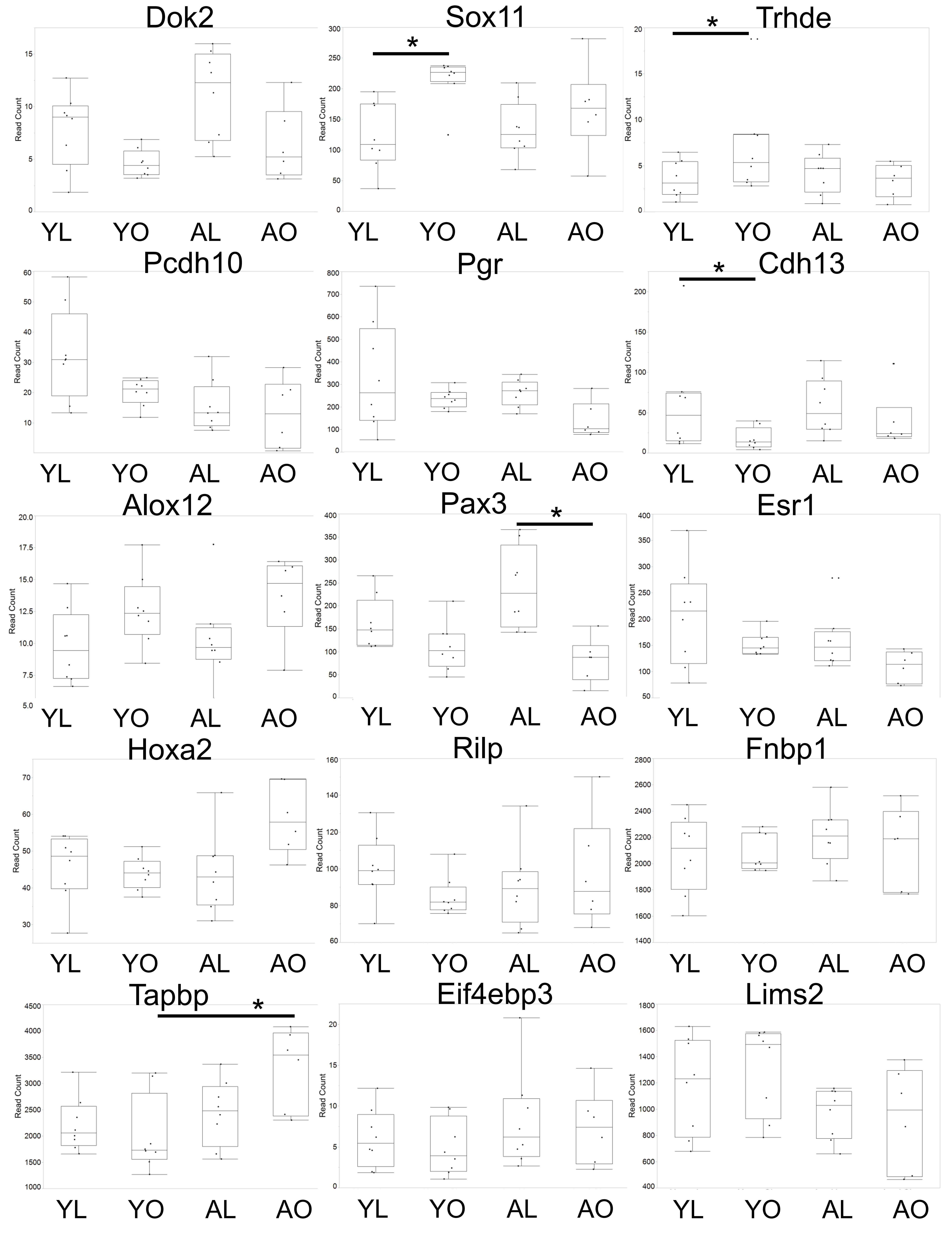
