## Supplemental Figure 3 for "Aging and Obesity Prime the Methylome and Transcriptome of Adipose Stem Cells for Disease and Dysfunction"

UpSet Plots for WGCNA modules associated with diet and age ordered by r-value with 'Top 20' Disease and Functions from Ingenuity Pathways Analysis identified and ranked by significance and by predicted activation z-score. Set size represents number of in-module biotypes associated with that disease or function.

A. Positive Association with Age

Dark Orange Module
Module Size = 70 biotypes
r-value = 0.54
p= 0.002
Positive Association with Age

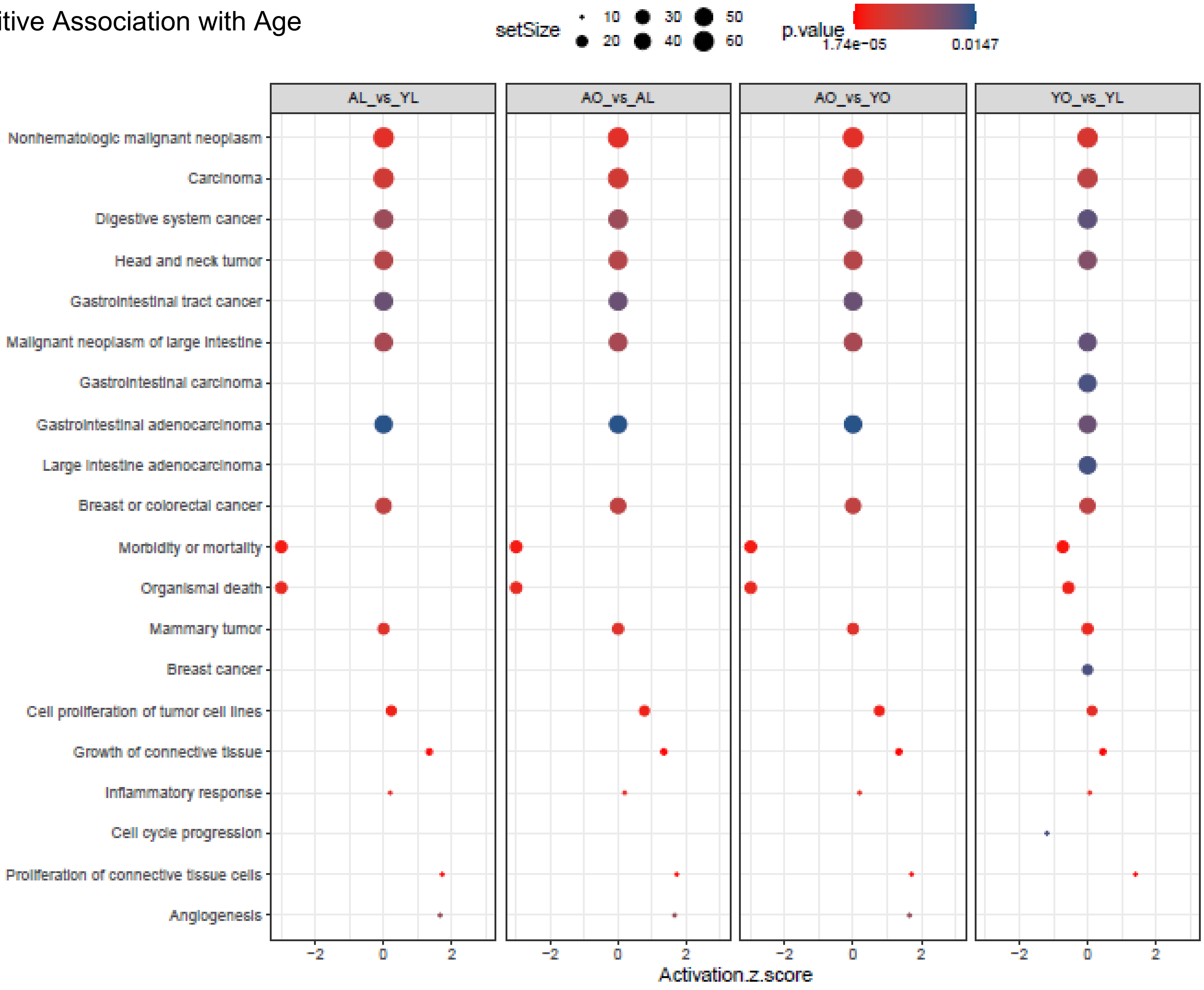

Black Module
Module Size = 696 biotypes
r-value = 0.47
p= 0.0009
Positive Association with Age

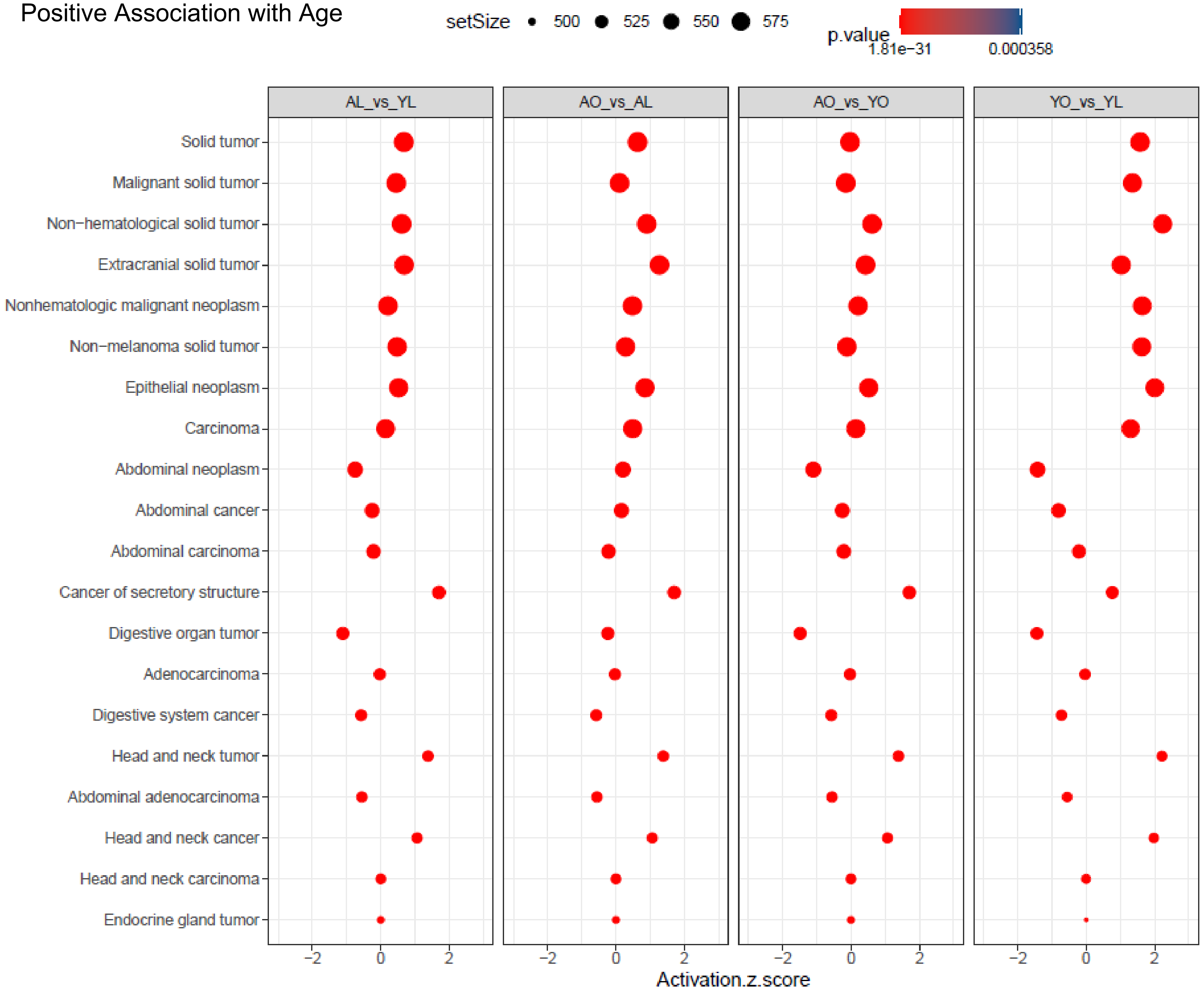

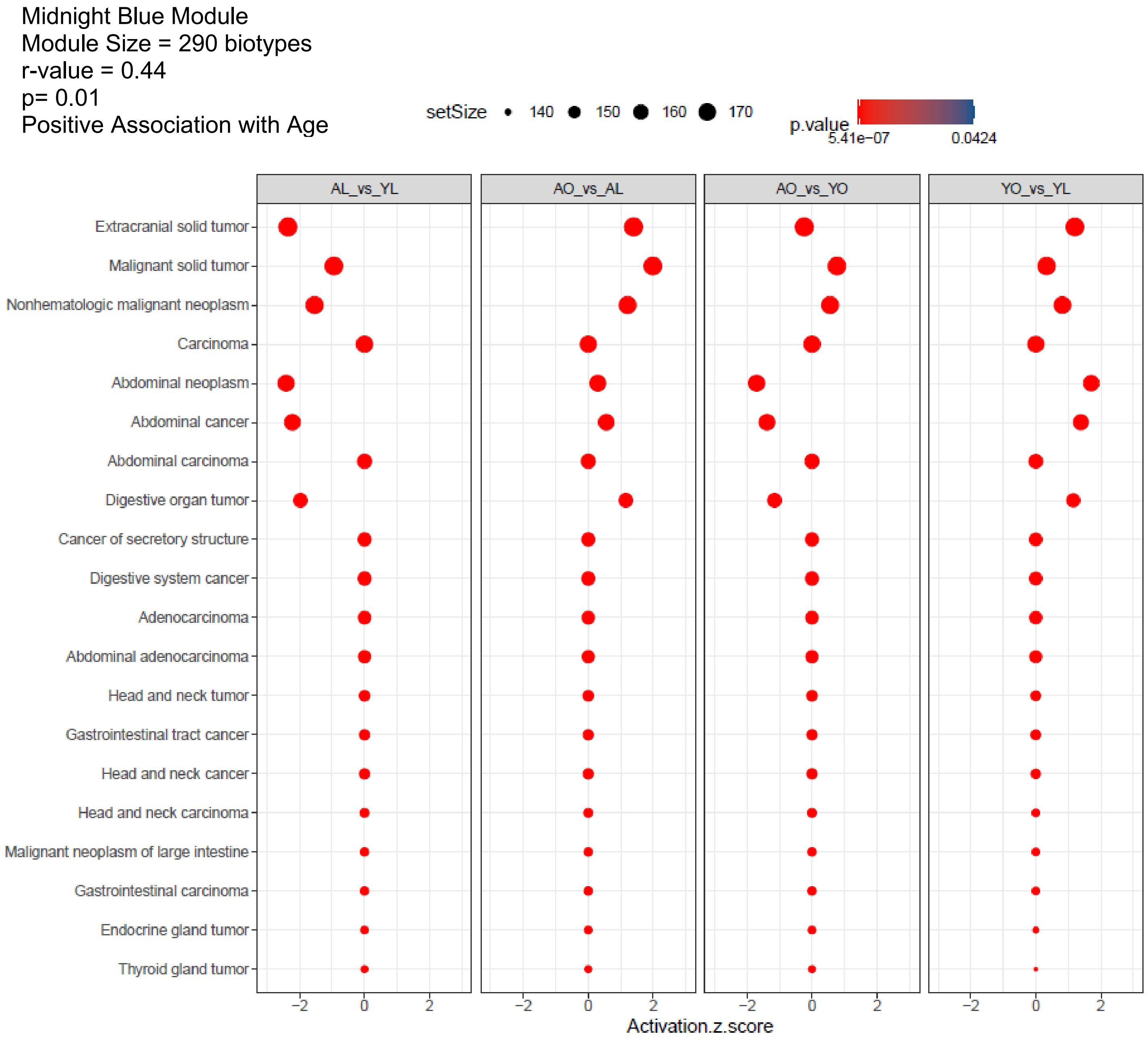

Midnight Blue Module
Module Size = 290 biotypes
r-value = 0.44
p= 0.01
Positive Association with Age
setSize
140
150
160
170
p.value
5.41e−07
0.0424
AL_vs_YL
AO_vs_AL
AO_vs_YO
YO_vs_YL
Extracranial solid tumor
Malignant solid tumor
Nonhematologic malignant neoplasm
Carcinoma
Abdominal neoplasm
Abdominal cancer
Abdominal carcinoma
Digestive organ tumor
Cancer of secretory structure
Digestive system cancer
Adenocarcinoma
Abdominal adenocarcinoma
Head and neck tumor
Gastrointestinal tract cancer
Head and neck cancer
Head and neck carcinoma
Malignant neoplasm of large intestine
Gastrointestinal carcinoma
Endocrine gland tumor
Thyroid gland tumor
−2
0
2
Activation.z.score

Orange Module
Module Size = 76 biotypes
r-value = 0.4
p= 0.03
Positive Association with Age

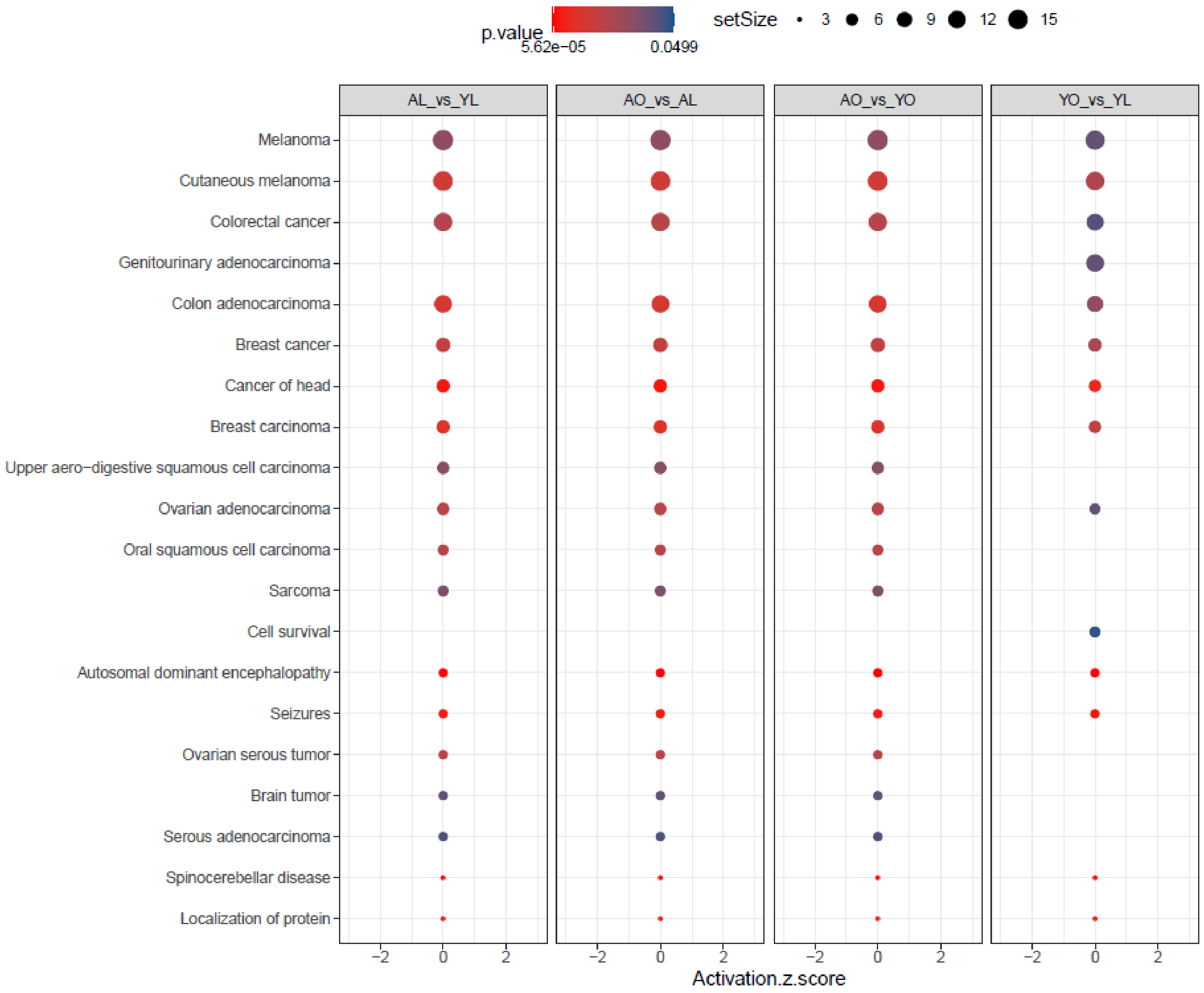

Pale Turquoise
Module Size = 41 biotypes
r-value = 0.4
p= 0.03
Positive Association with Age

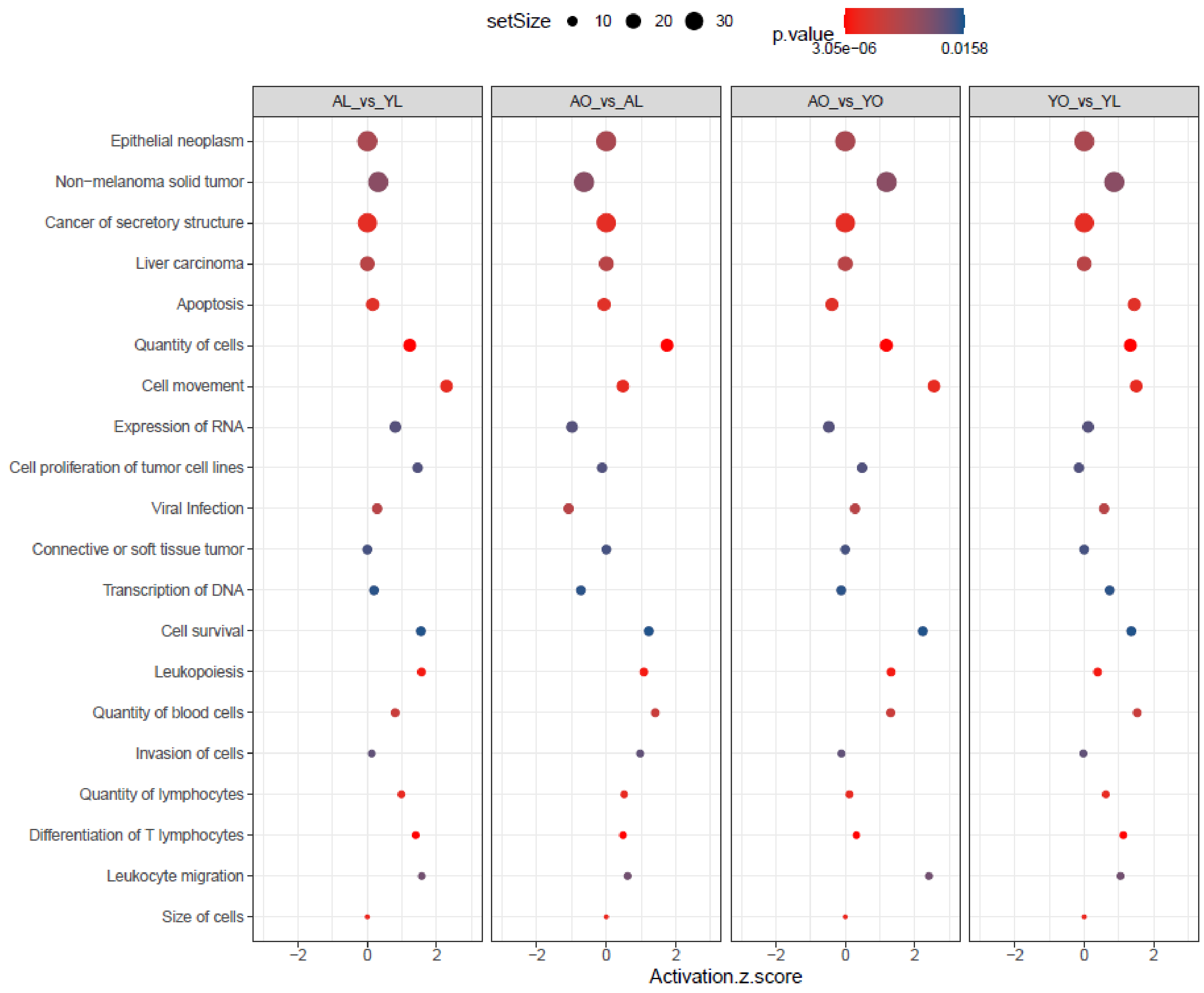

Cyan
Module Size = 331 biotypes
r-value = 0.39
p= 0.03
Positive Association with Age

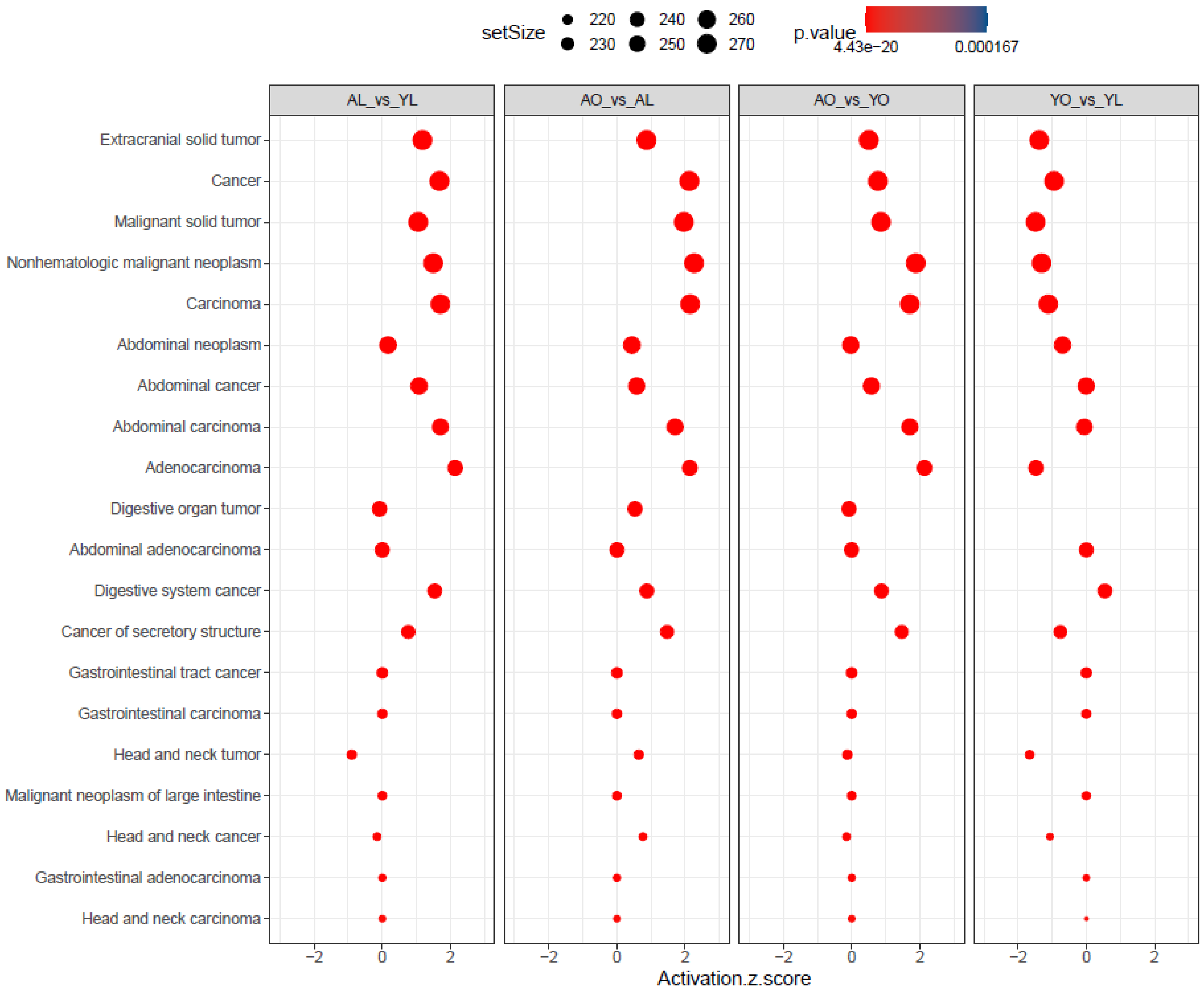

Royal Blue
Module Size = 136 biotypes
r-value = 0.36
p= 0.05
Positive Association with Age

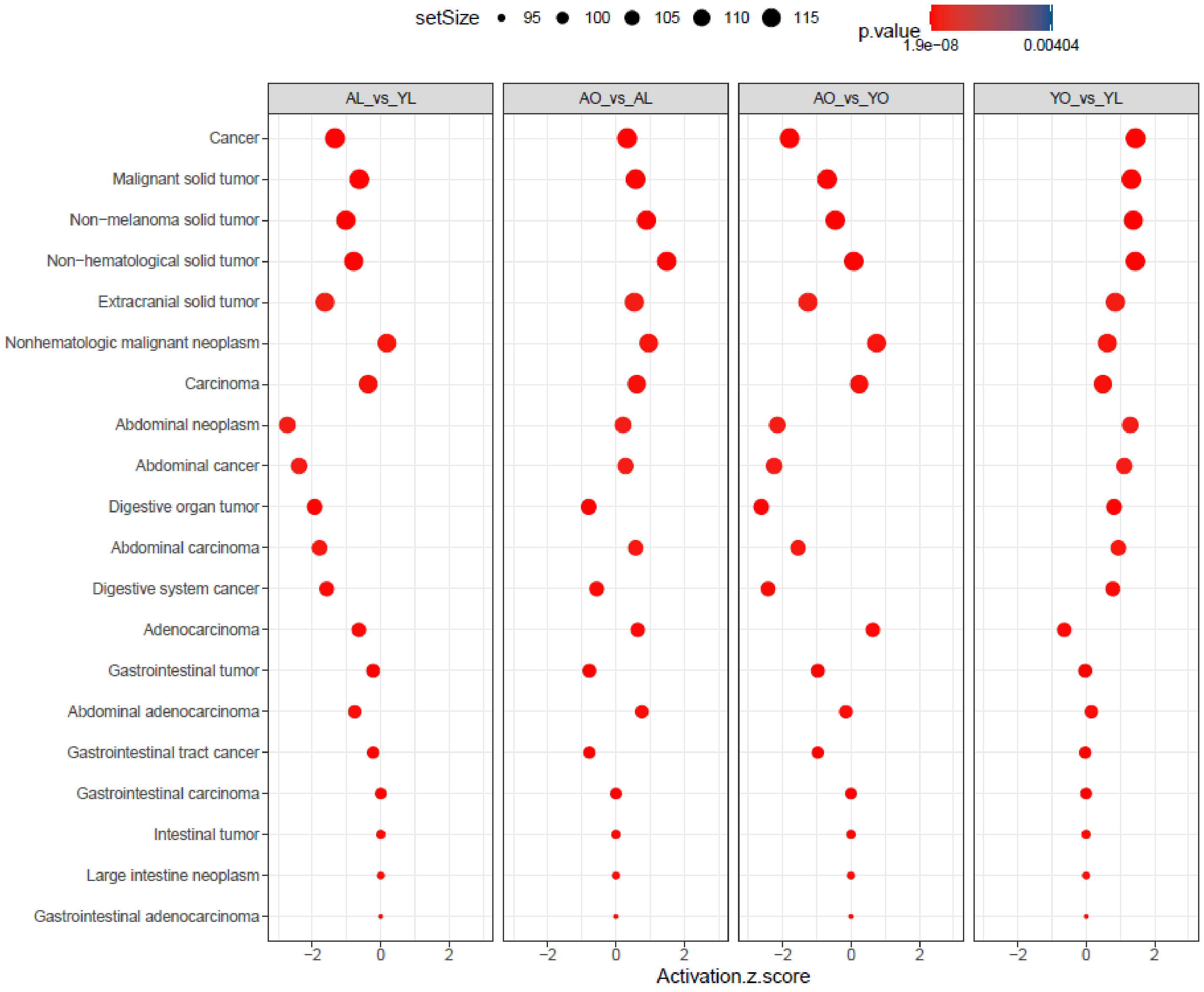

B. Negative Association with Age

Sky Blue
Module Size = 61 biotypes
r-value = -0.53
p= 0.003
Negative Association with Age

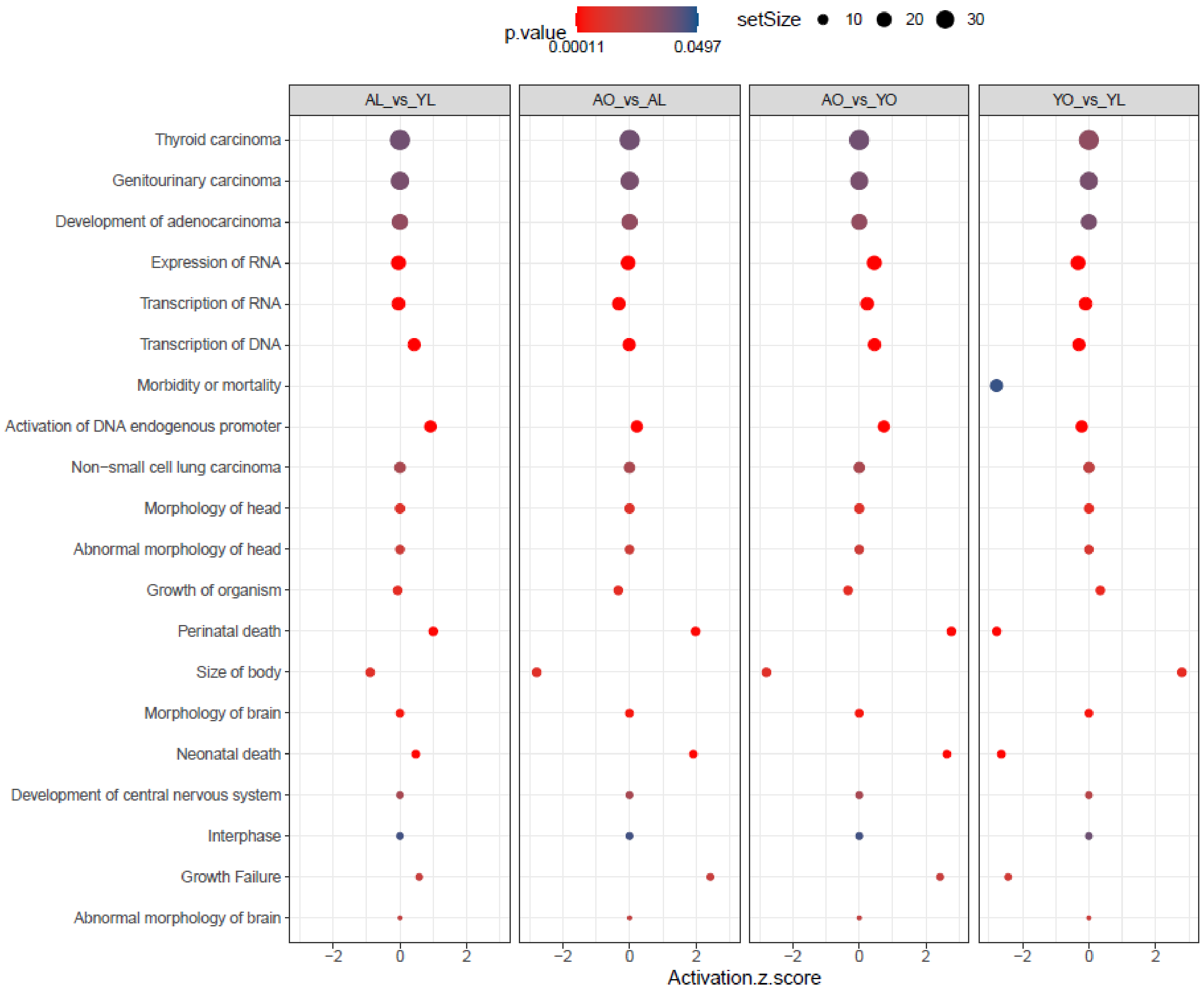

Dark Red
Module Size = 122 biotypes
r-value = -0.5
p= 0.005
Negative Association with Age

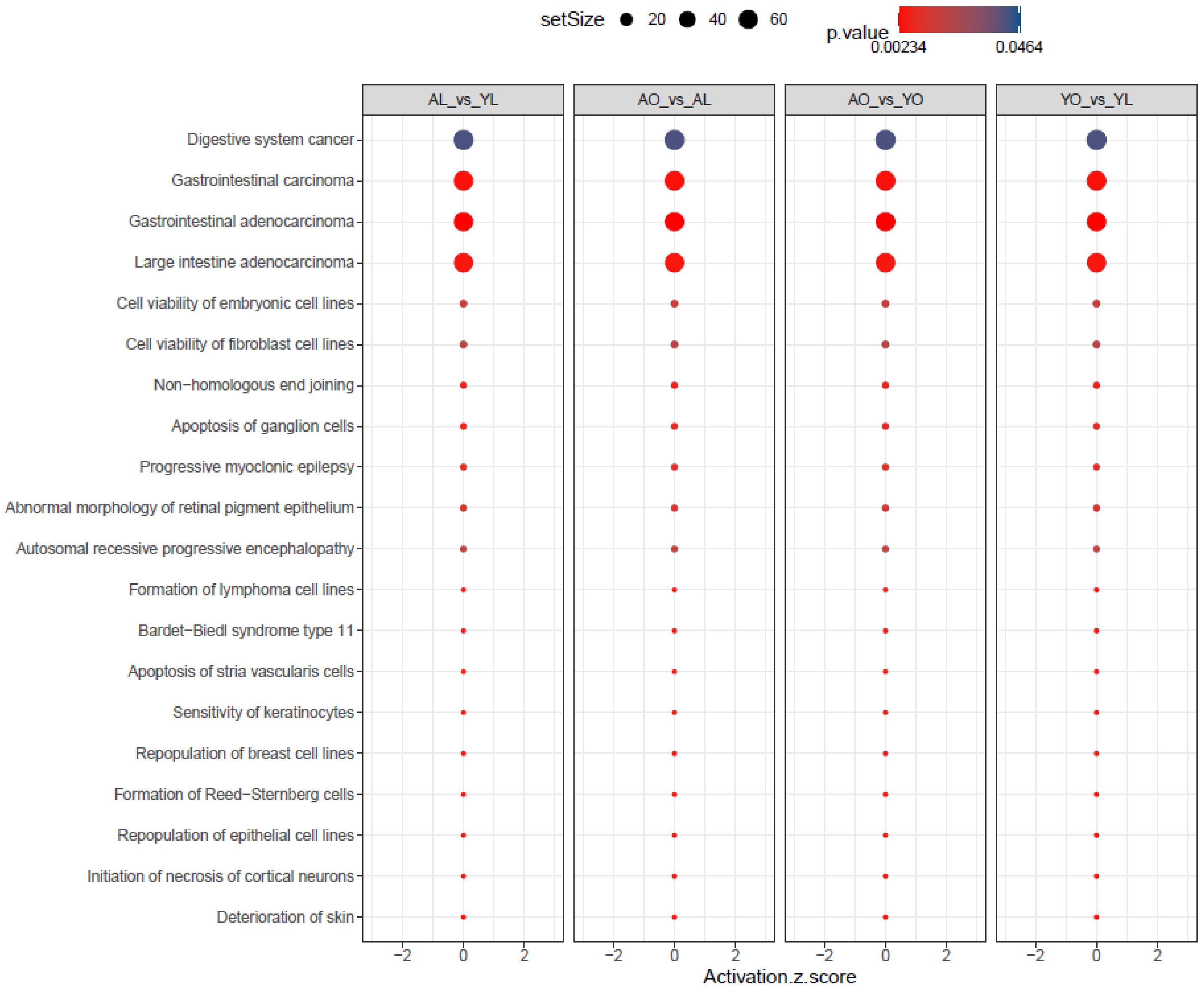

Dark Green
Module Size = 120 biotypes
r-value = -0.47
p= 0.008
Negative Association with Age

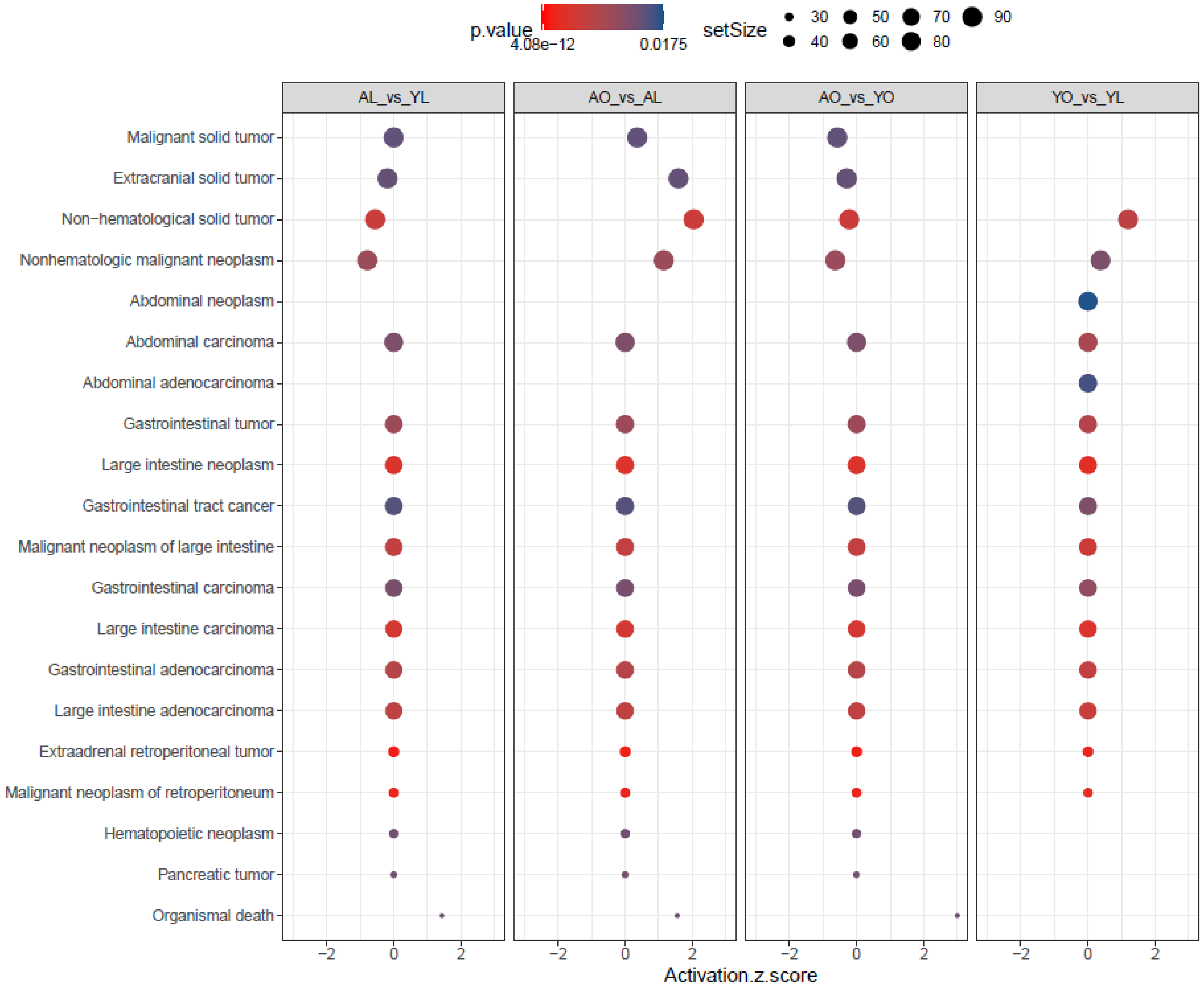

Magenta
Module Size = 557 biotypes
r-value = -0.44
p= 0.02
Negative Association with Age

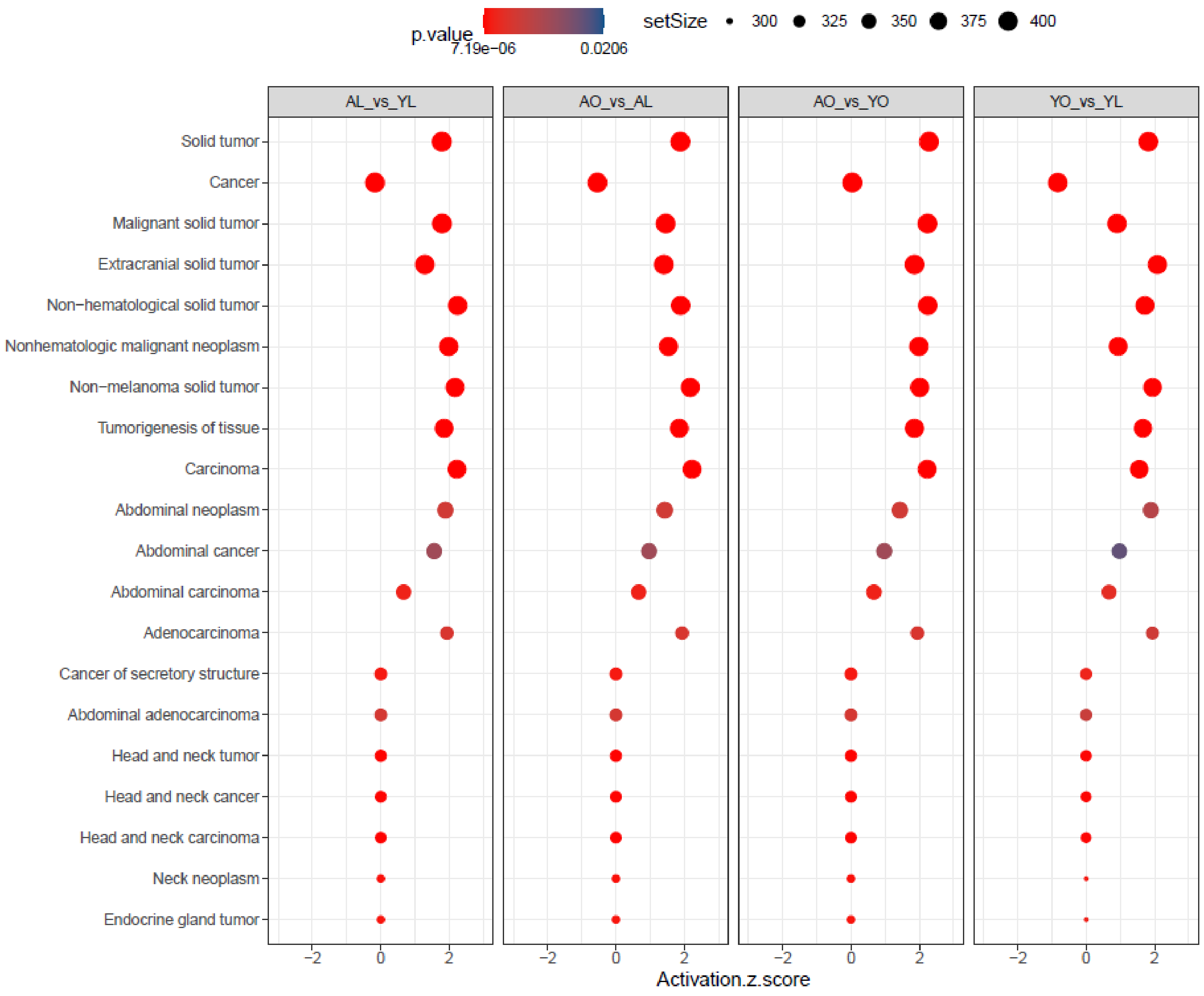

Green
Module Size = 785 biotypes
r-value = -0.43
p= 0.02
Negative Association with Age

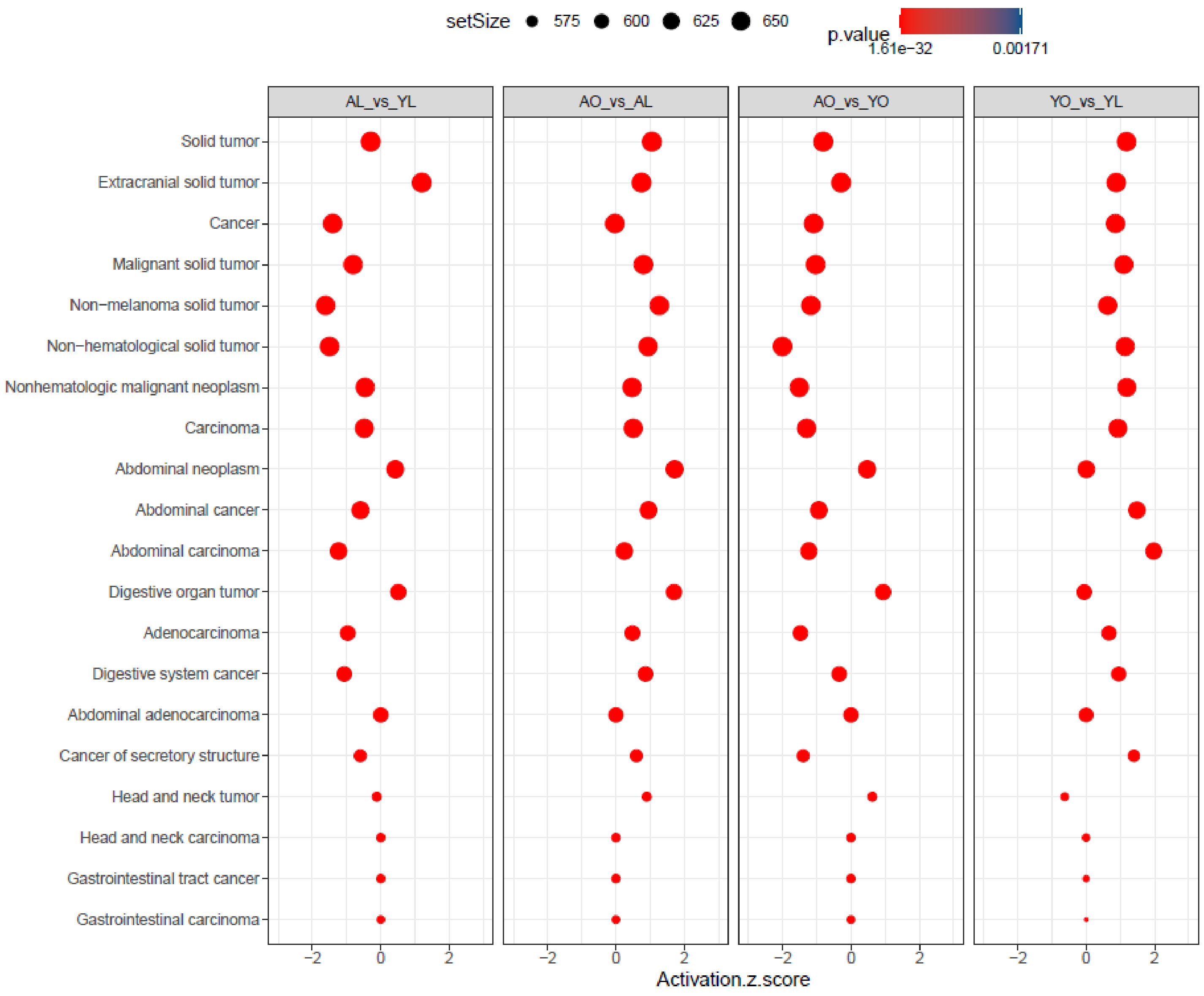

C. Negative Association with Diet

Grey
Module Size = 513 biotypes
r-value = -0.43
p= 0.02
Negative Association with Diet

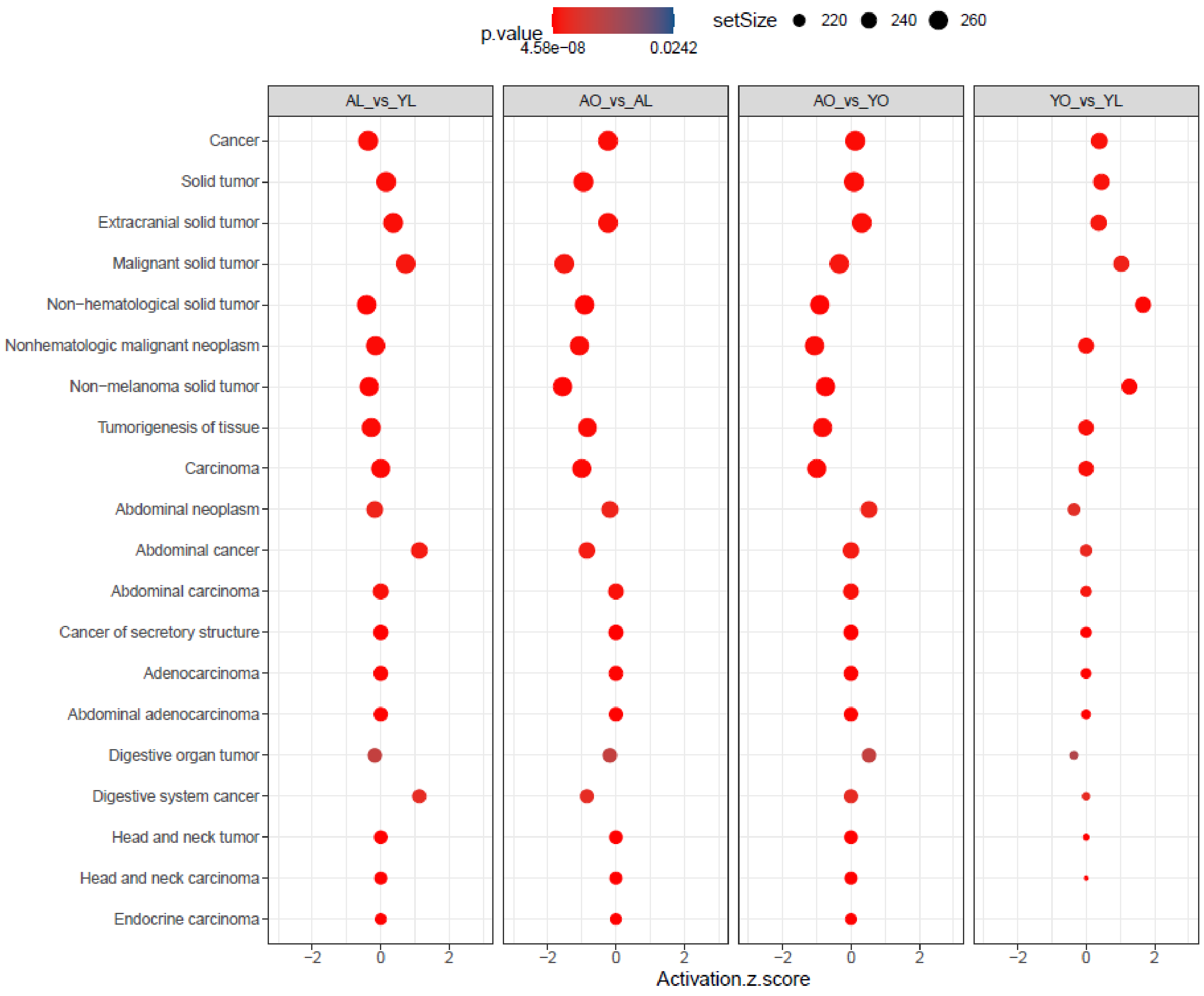

Sky Blue
Module Size = 61 biotypes
r-value = -0.53
p= 0.003
Negative Association with Diet

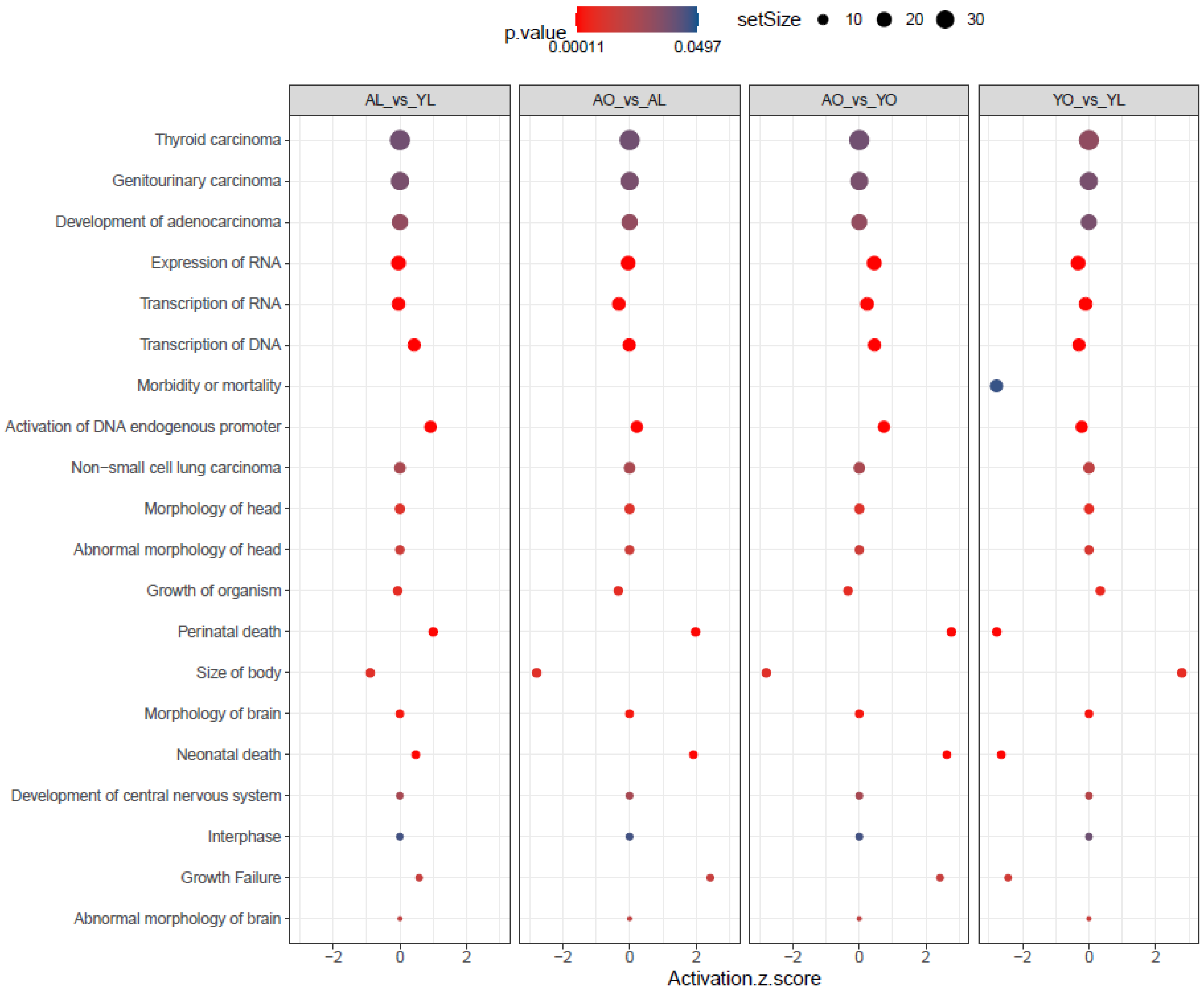

UpSet Plots for WCGNA modules associated with bodyweight at euthanasia in aged mice ordered by r-value with 'Top 20' Disease and Functions from Ingenuity Pathways Analysis identified and ranked by significance and by predicted activation z-score. Set size represents number of in-module biotypes associated with that disease or function.

D. Negative Association with Bodyweight

Midnight Blue
Module Size = 185 biotypes
r-value = -0.65
p= 0.006
Negative Association with Bodyweight

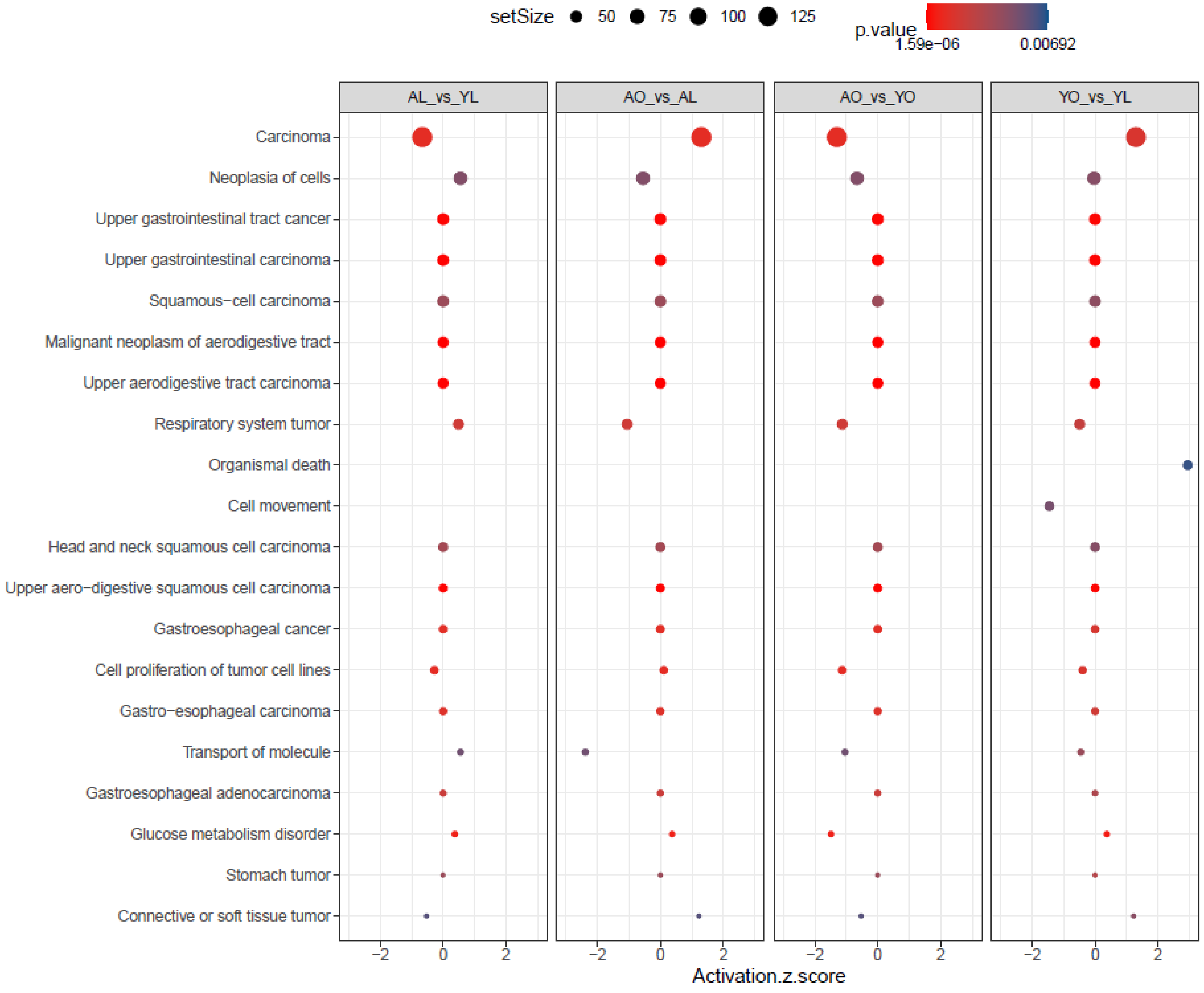

Sky Blue
Module Size = 79 biotypes
r-value = -0.63
p= 0.02
Negative Association with Bodyweight

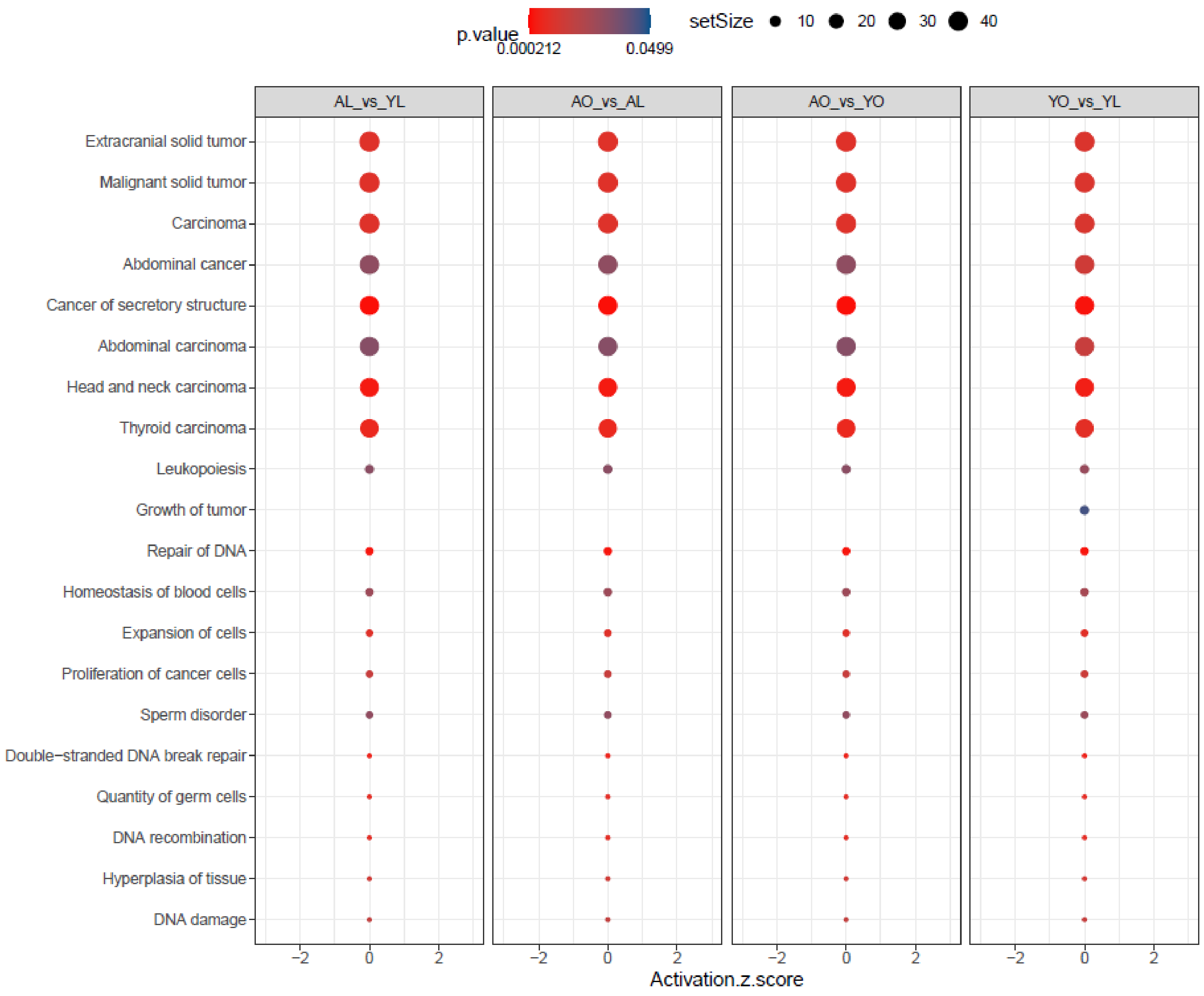

Pale Turquoise
Module Size = 68 biotypes
r-value = -0.63
p= 0.02
Negative Association with Bodyweight

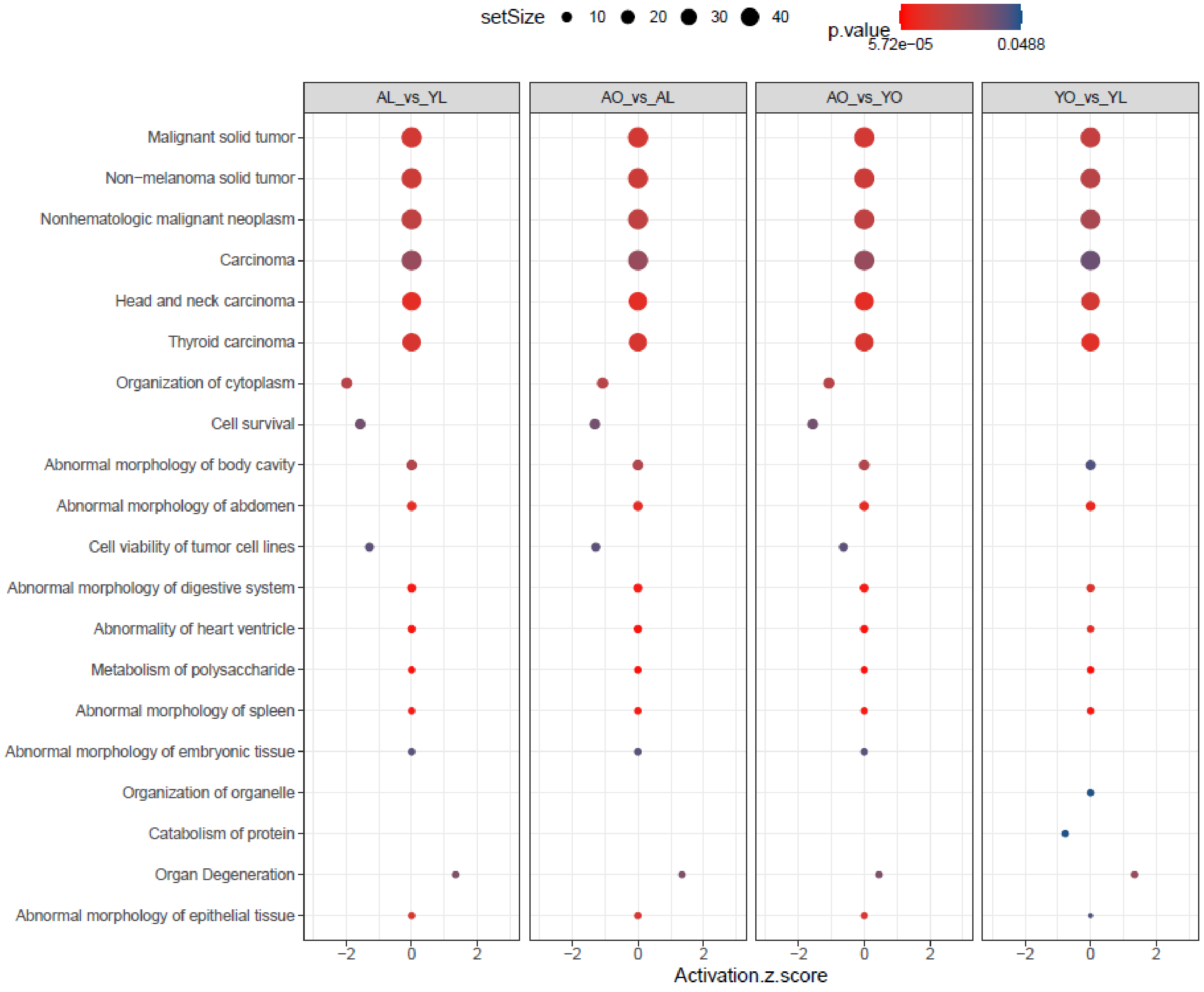

Black
Module Size = 390 biotypes
r-value = -0.58
p= 0.03
Negative Association with Bodyweight

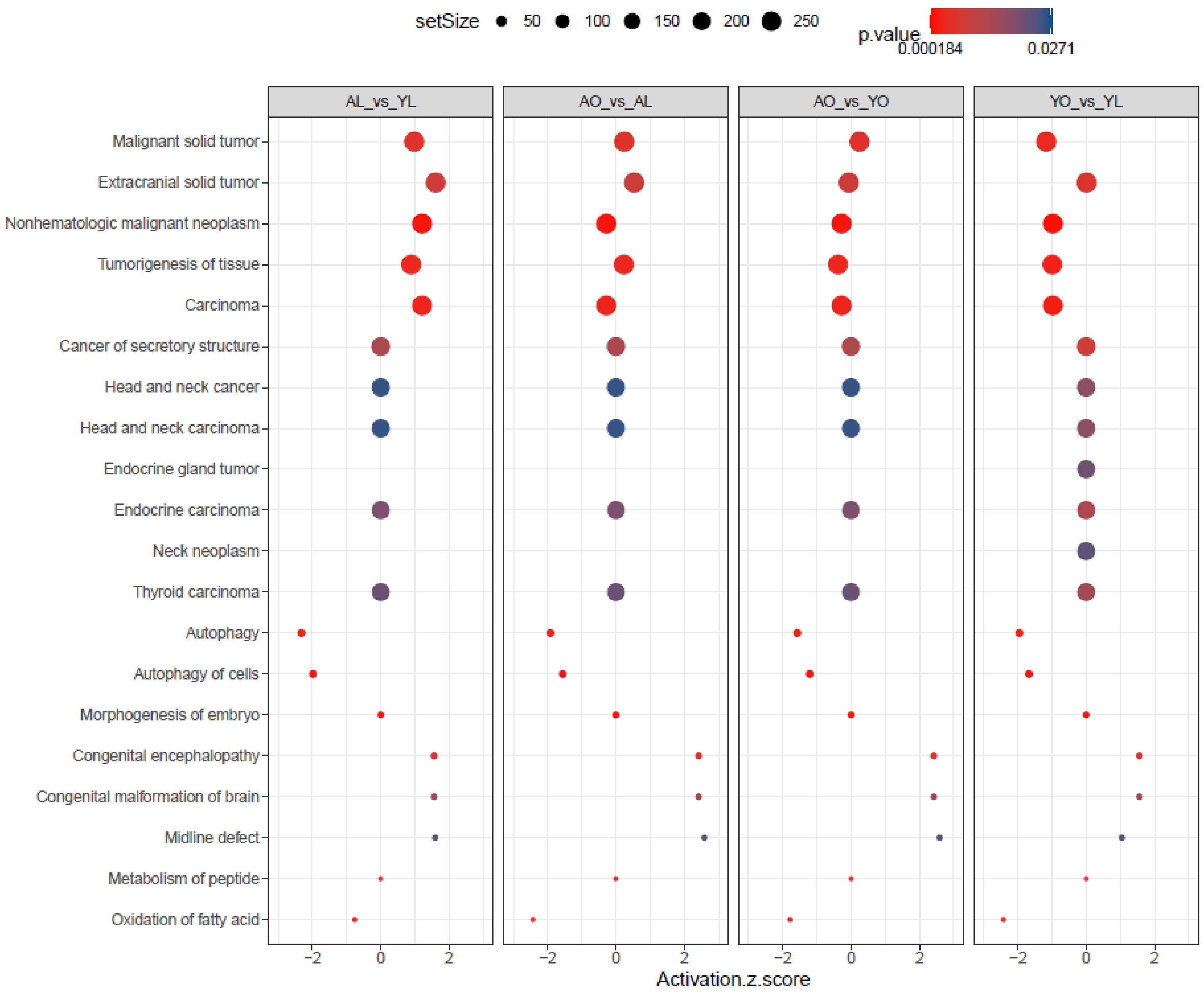

Pink
Module Size = 320 biotypes
r-value = -0.55
p= 0.04
Negative Association with Bodyweight

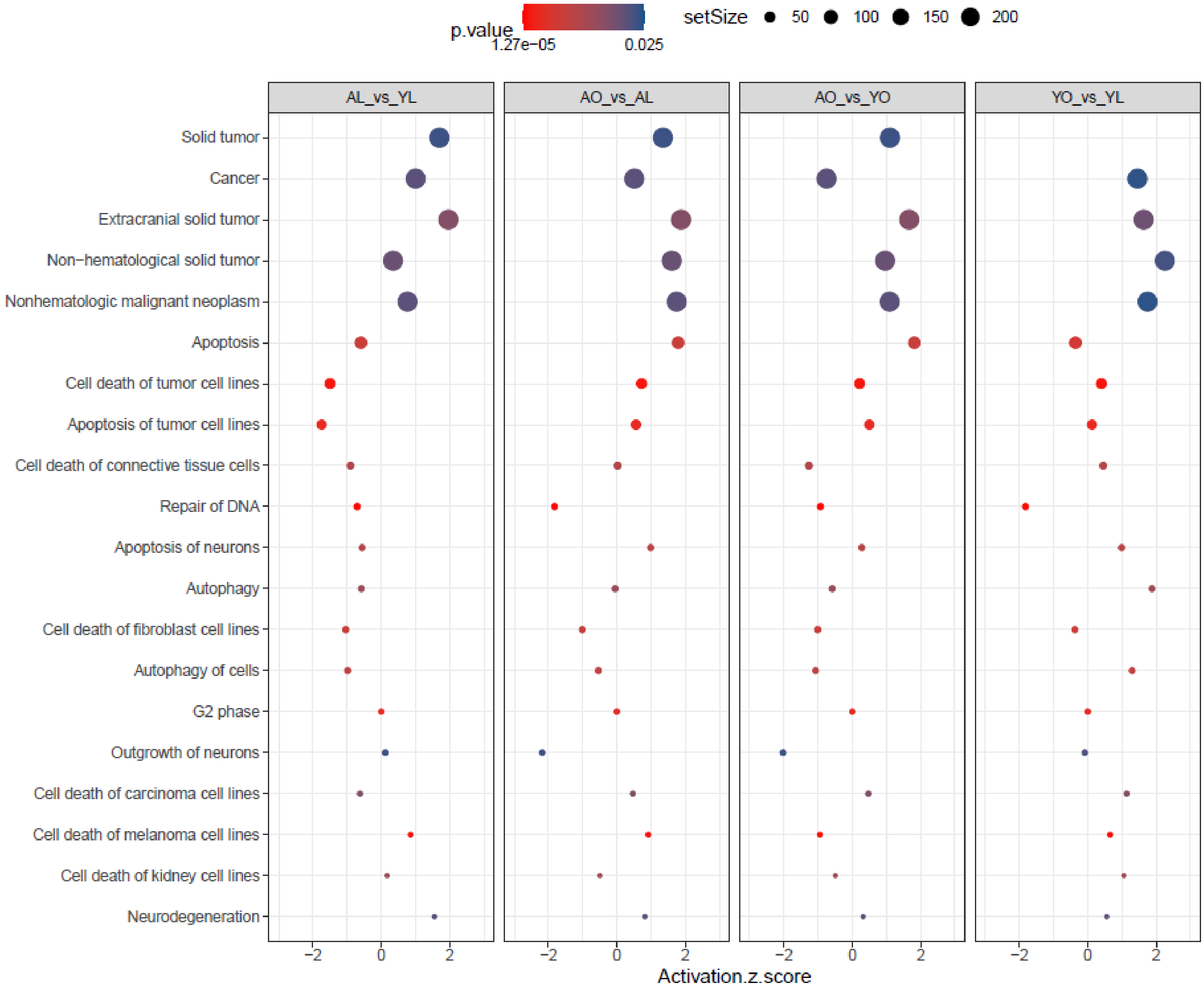

E. Positive association with Bodyweight at euthanasia

Cyan
Module Size = 206 biotypes
r-value = 0.70
p= 0.005
Positive Association with Bodyweight

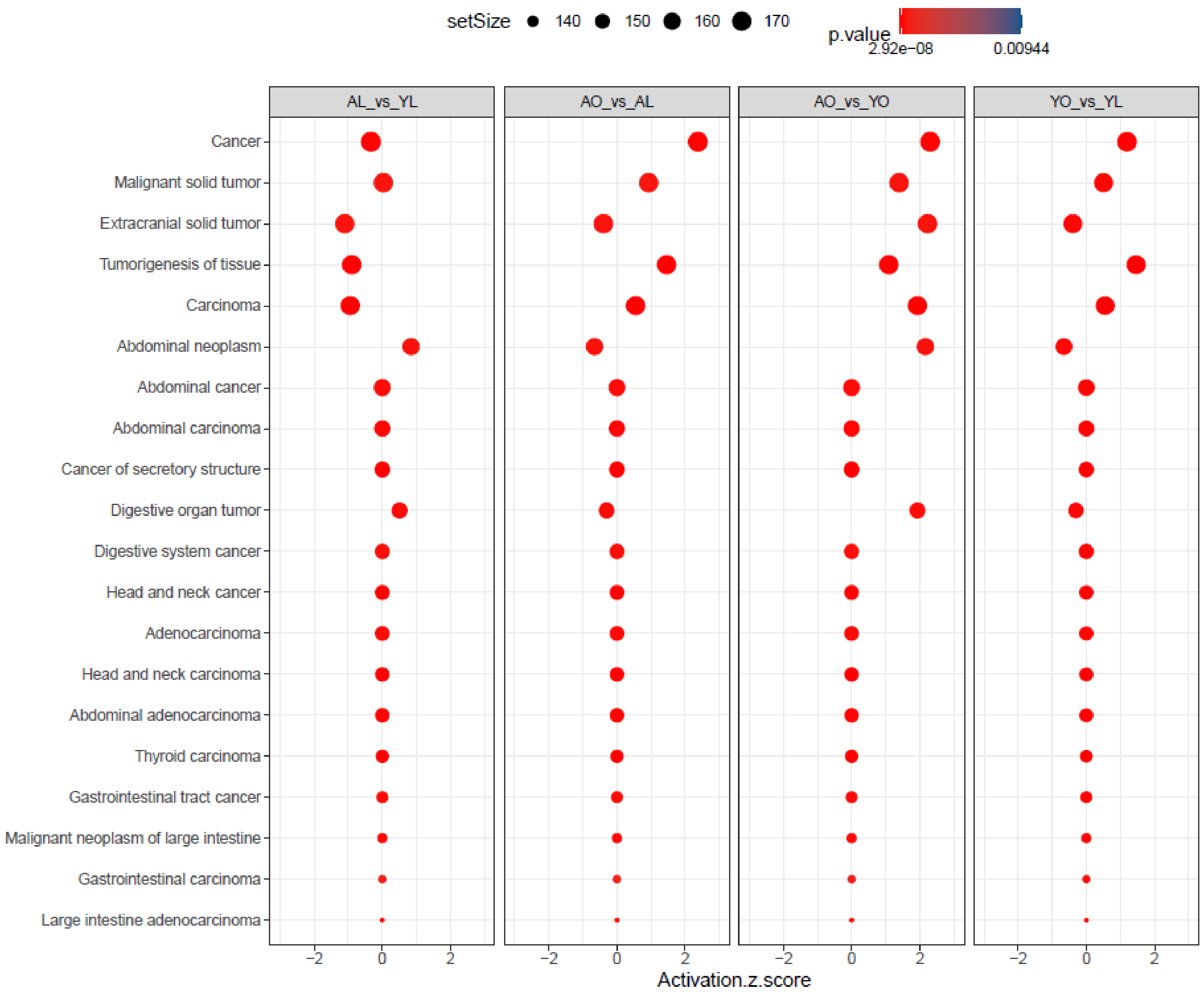

Salmon
Module Size = 219 biotypes
r-value = 0.62
p= 0.02
Positive Association with Bodyweight

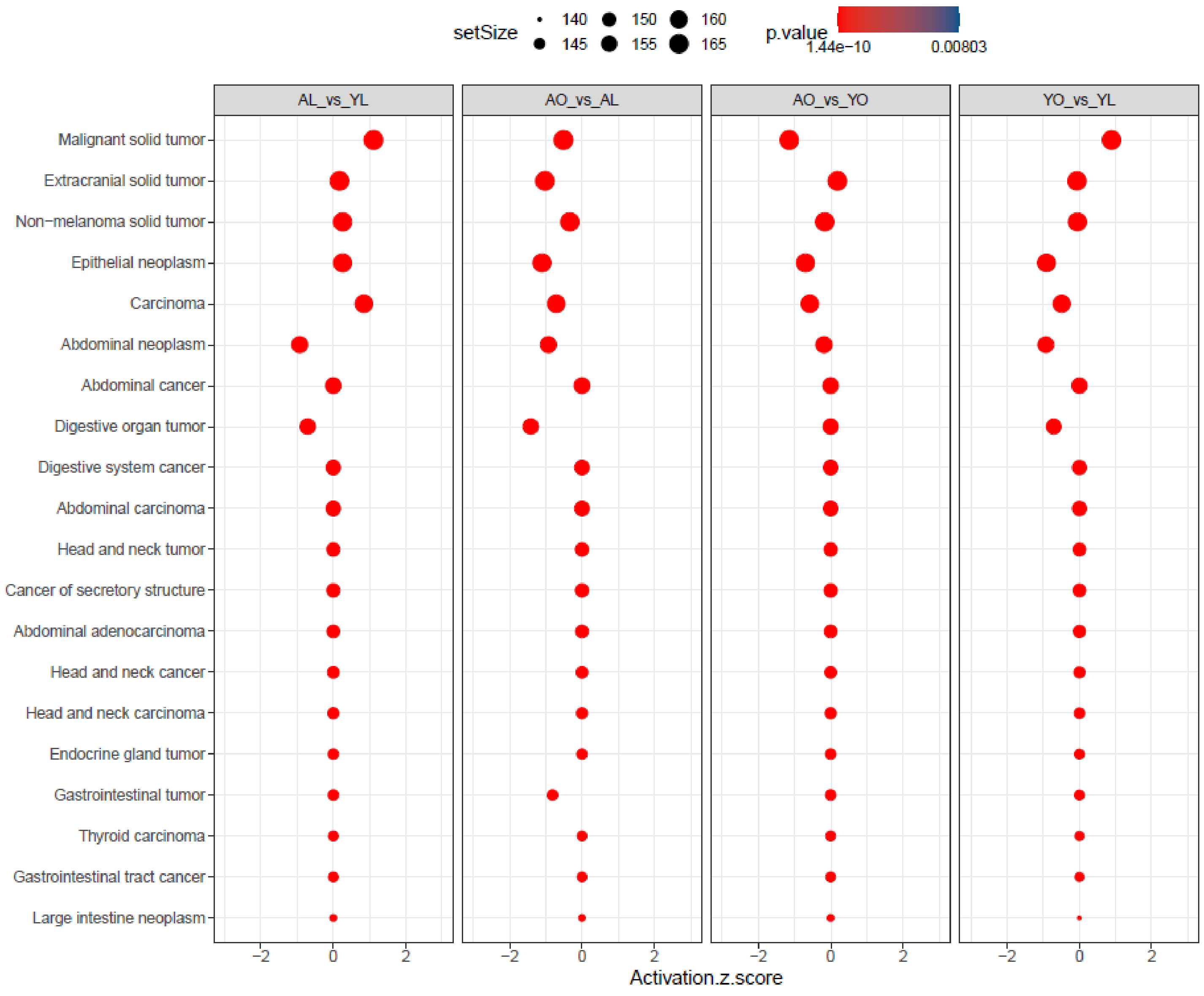

Green
Module Size = 840 biotypes
r-value = 0.56
p= 0.04
Positive Association with Bodyweight

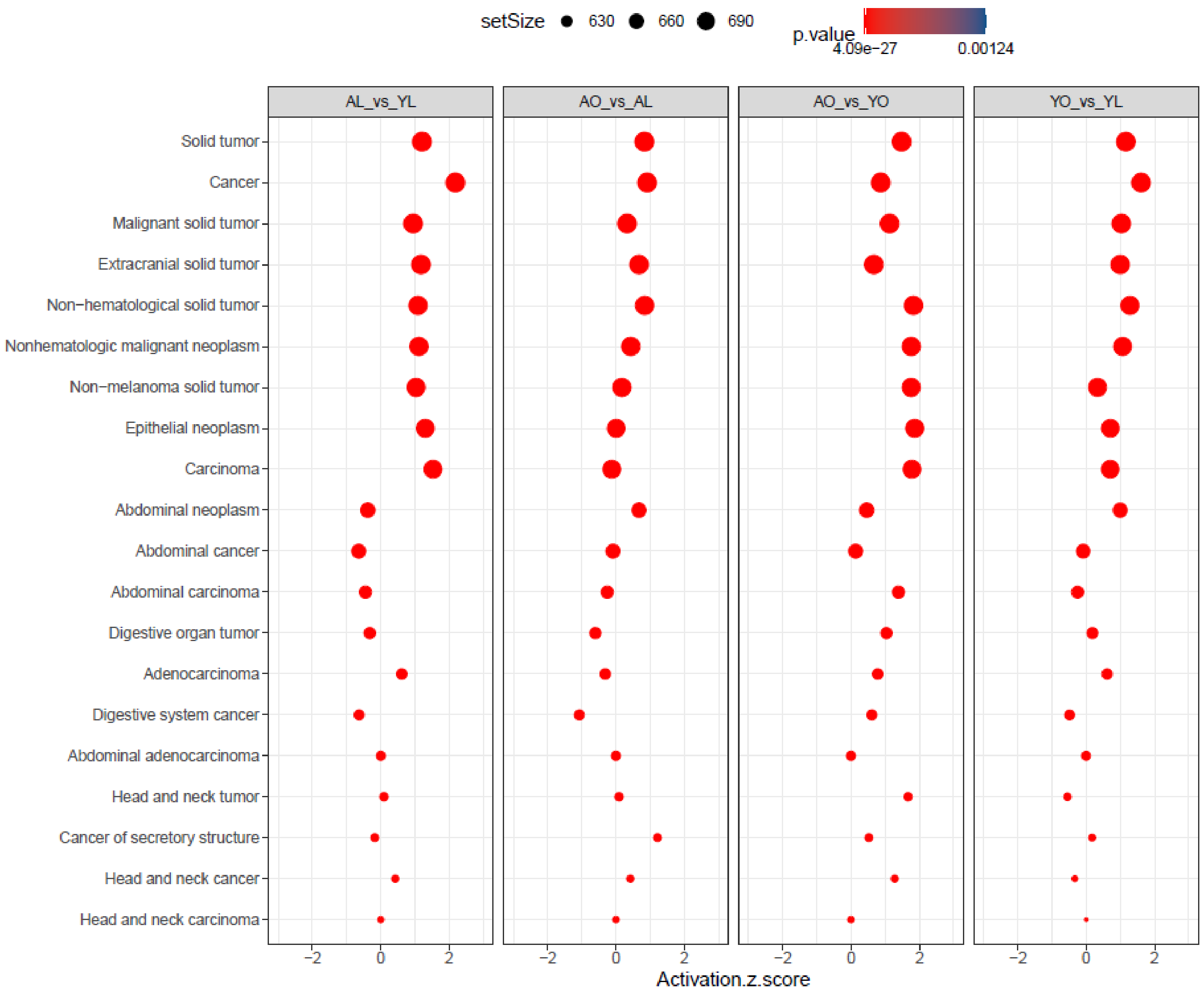

Light Yellow
Module Size = 143 biotypes
r-value = 0.54
p= 0.04
Positive Association with Bodyweight

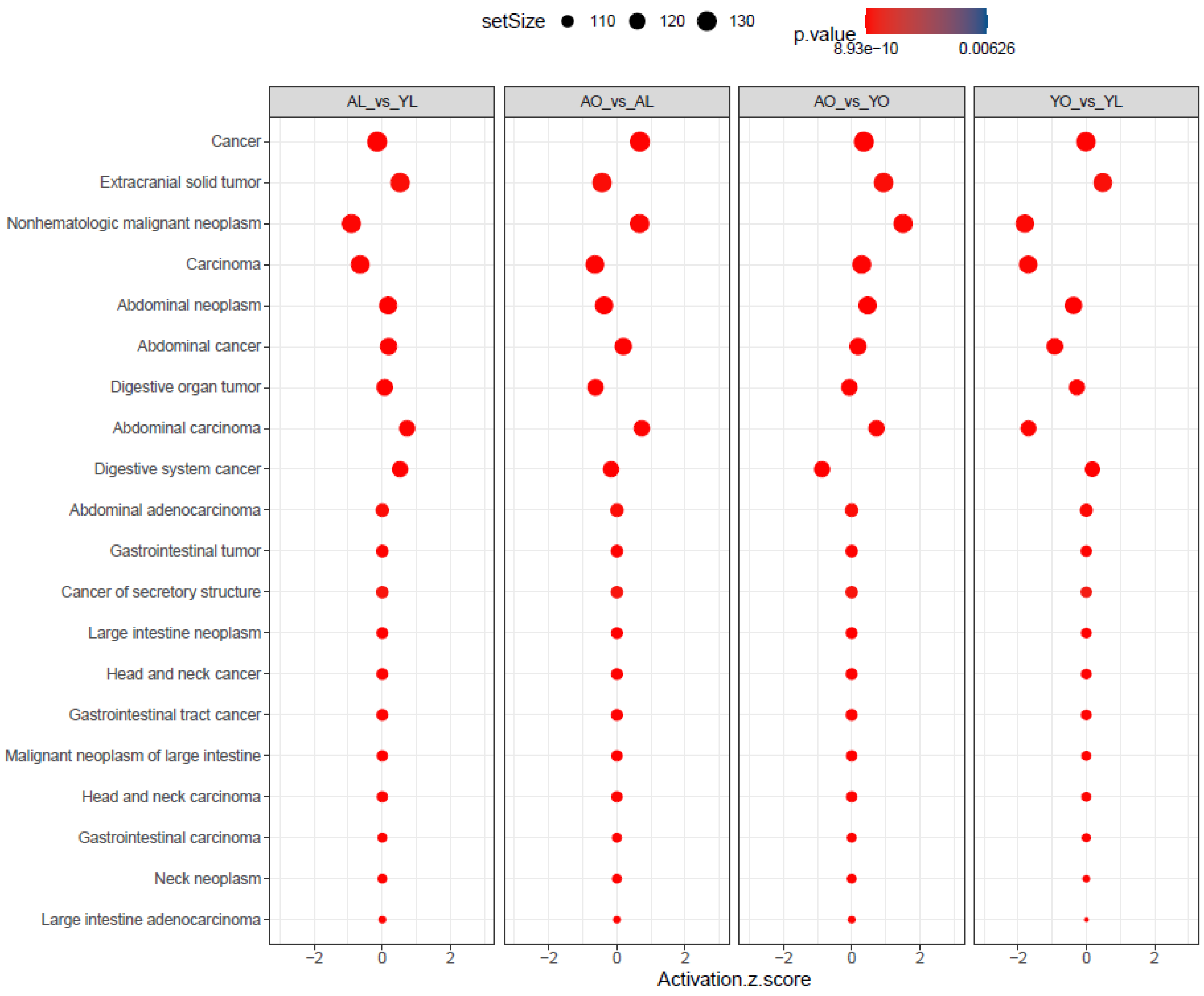
