## Supplemental Figure 4 for "Aging and Obesity Prime the Methylome and Transcriptome of Adipose Stem Cells for Disease and Dysfunction"

Tcfl5
ENSMUSG00000038932
VIP: 2.78, $p_{adj}$: 2.66E-11

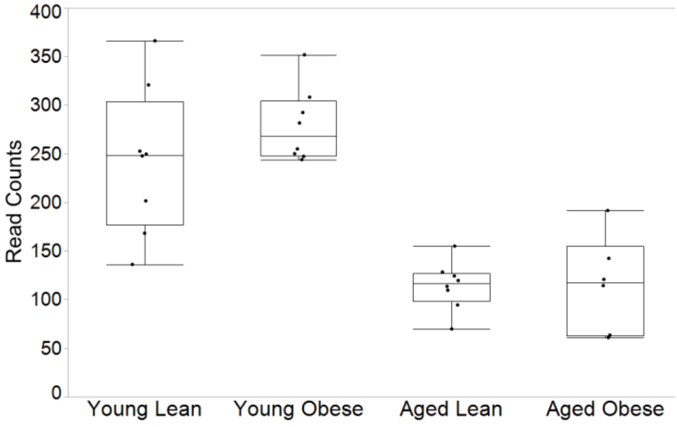


Vhl
ENSMUSG00000033933
VIP: 2.50, $p_{adj}$: 6.89E-06

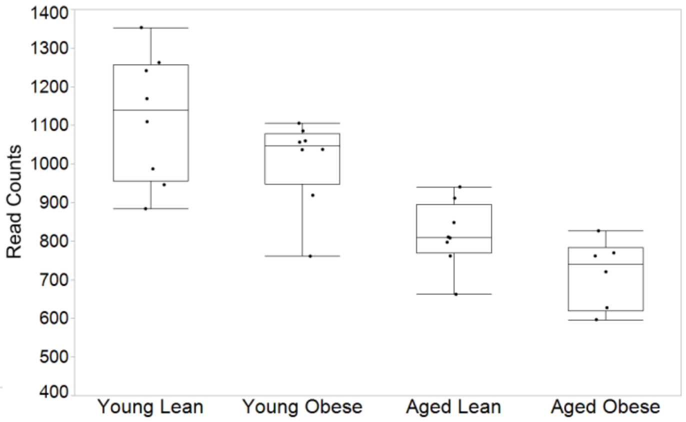


Rflna
ENSMUSG00000037962
VIP: 2.13, $p_{adj}$: 8.09E-06

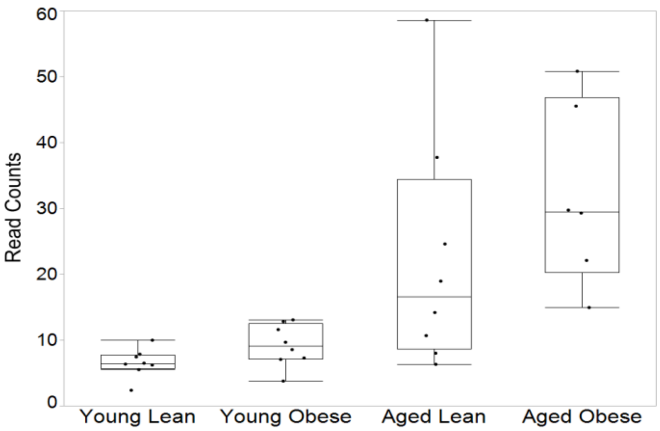


Shox2
ENSMUSG0000027833VIP:
2.40, $p_{adj}$: 2.92E-05

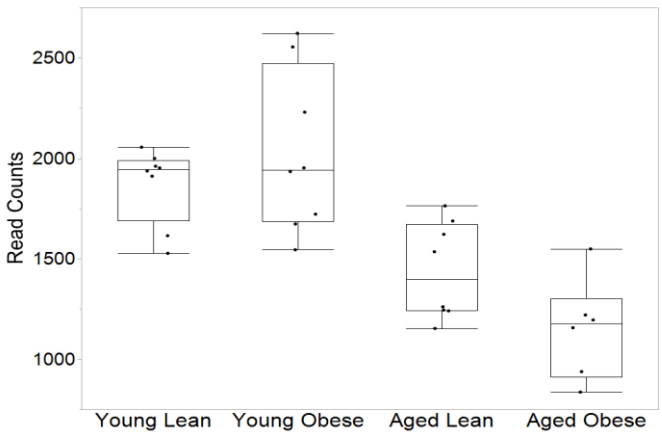


Stc2
ENSMUSG00000020303
VIP: 2.30, $p_{adj}$: 2.92E-05

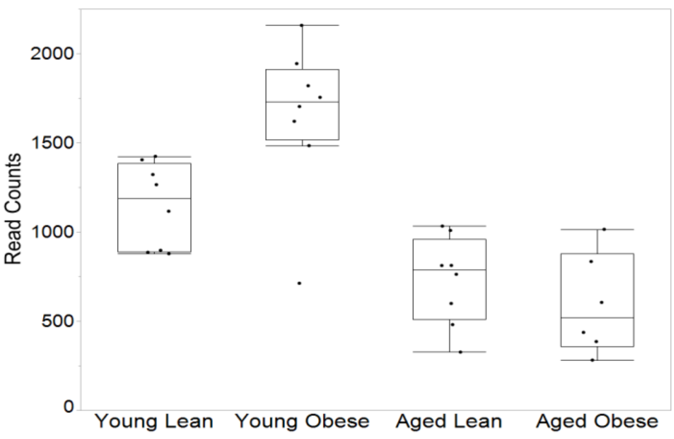


Klra4
ENSMUSG00000079852
VIP: 2.73, $p_{adj}$: 8.54E-05

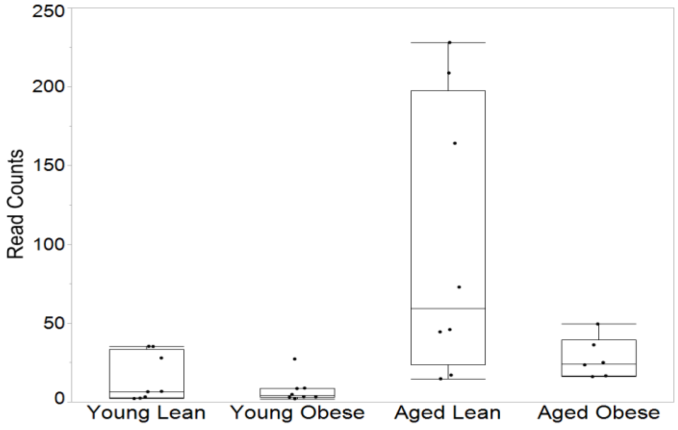

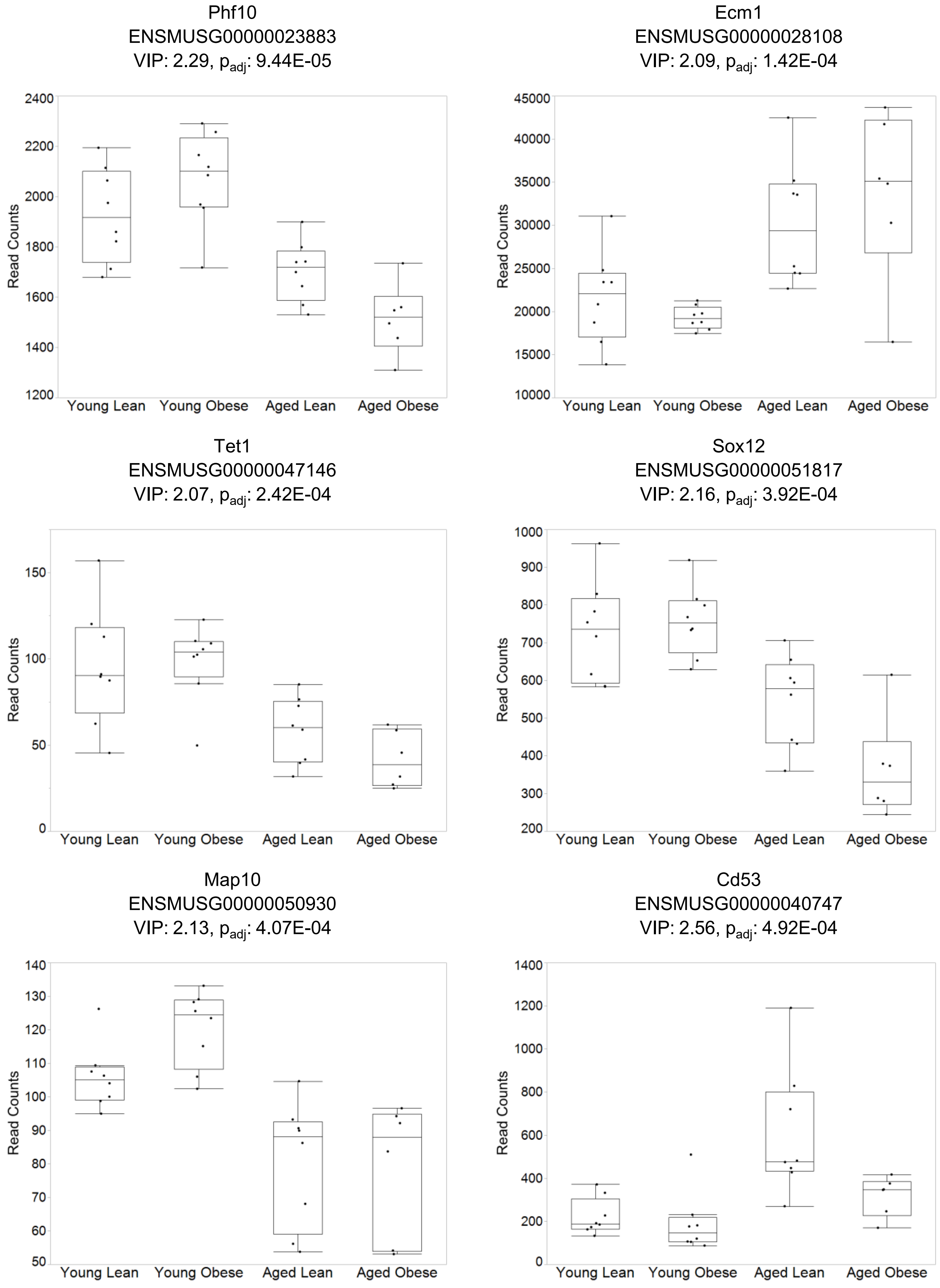

Phf10
ENSMUSG00000023883
VIP: 2.29, $p_{adj}$: 9.44E-05
Read Counts
1200 1400 1600 1800 2000 2200 2400
Young Lean Young Obese Aged Lean Aged Obese
Ecm1
ENSMUSG00000028108
VIP: 2.09, $p_{adj}$: 1.42E-04
Read Counts
10000 15000 20000 25000 30000 35000 40000 45000
Young Lean Young Obese Aged Lean Aged Obese
Tet1
ENSMUSG00000047146
VIP: 2.07, $p_{adj}$: 2.42E-04
Read Counts
0 50 100 150
Young Lean Young Obese Aged Lean Aged Obese
Sox12
ENSMUSG00000051817
VIP: 2.16, $p_{adj}$: 3.92E-04
Read Counts
200 300 400 500 600 700 800 900 1000
Young Lean Young Obese Aged Lean Aged Obese
Map10
ENSMUSG00000050930
VIP: 2.13, $p_{adj}$: 4.07E-04
Read Counts
50 60 70 80 90 100 110 120 130 140
Young Lean Young Obese Aged Lean Aged Obese
Cd53
ENSMUSG00000040747
VIP: 2.56, $p_{adj}$: 4.92E-04
Read Counts
0 200 400 600 800 1000 1200 1400
Young Lean Young Obese Aged Lean Aged Obese

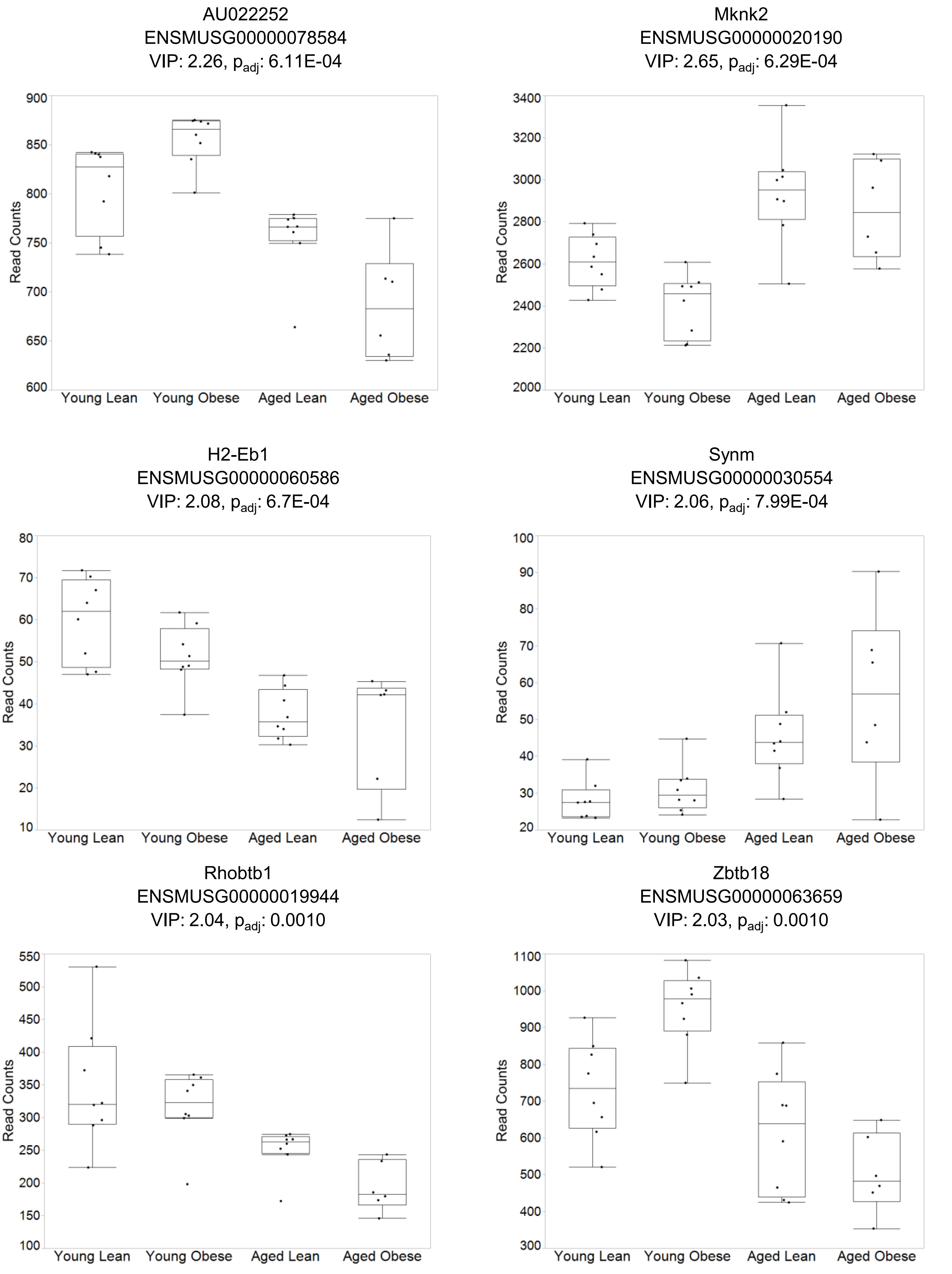
AU022252
ENSMUSG00000078584
VIP: 2.26, p_adj: 6.11E-04
Read Counts
900
850
800
750
700
650
600
Young Lean
Young Obese
Aged Lean
Aged Obese
Mknk2
ENSMUSG00000020190
VIP: 2.65, p_adj: 6.29E-04
Read Counts
3400
3200
3000
2800
2600
2400
2200
2000
Young Lean
Young Obese
Aged Lean
Aged Obese
H2-Eb1
ENSMUSG00000060586
VIP: 2.08, p_adj: 6.7E-04
Read Counts
80
70
60
50
40
30
20
10
Young Lean
Young Obese
Aged Lean
Aged Obese
Synm
ENSMUSG00000030554
VIP: 2.06, p_adj: 7.99E-04
Read Counts
100
90
80
70
60
50
40
30
20
Young Lean
Young Obese
Aged Lean
Aged Obese
Rhobtb1
ENSMUSG00000019944
VIP: 2.04, p_adj: 0.0010
Read Counts
550
500
450
400
350
300
250
200
150
100
Young Lean
Young Obese
Aged Lean
Aged Obese
Zbtb18
ENSMUSG00000063659
VIP: 2.03, p_adj: 0.0010
Read Counts
1100
1000
900
800
700
600
500
400
300
Young Lean
Young Obese
Aged Lean
Aged Obese

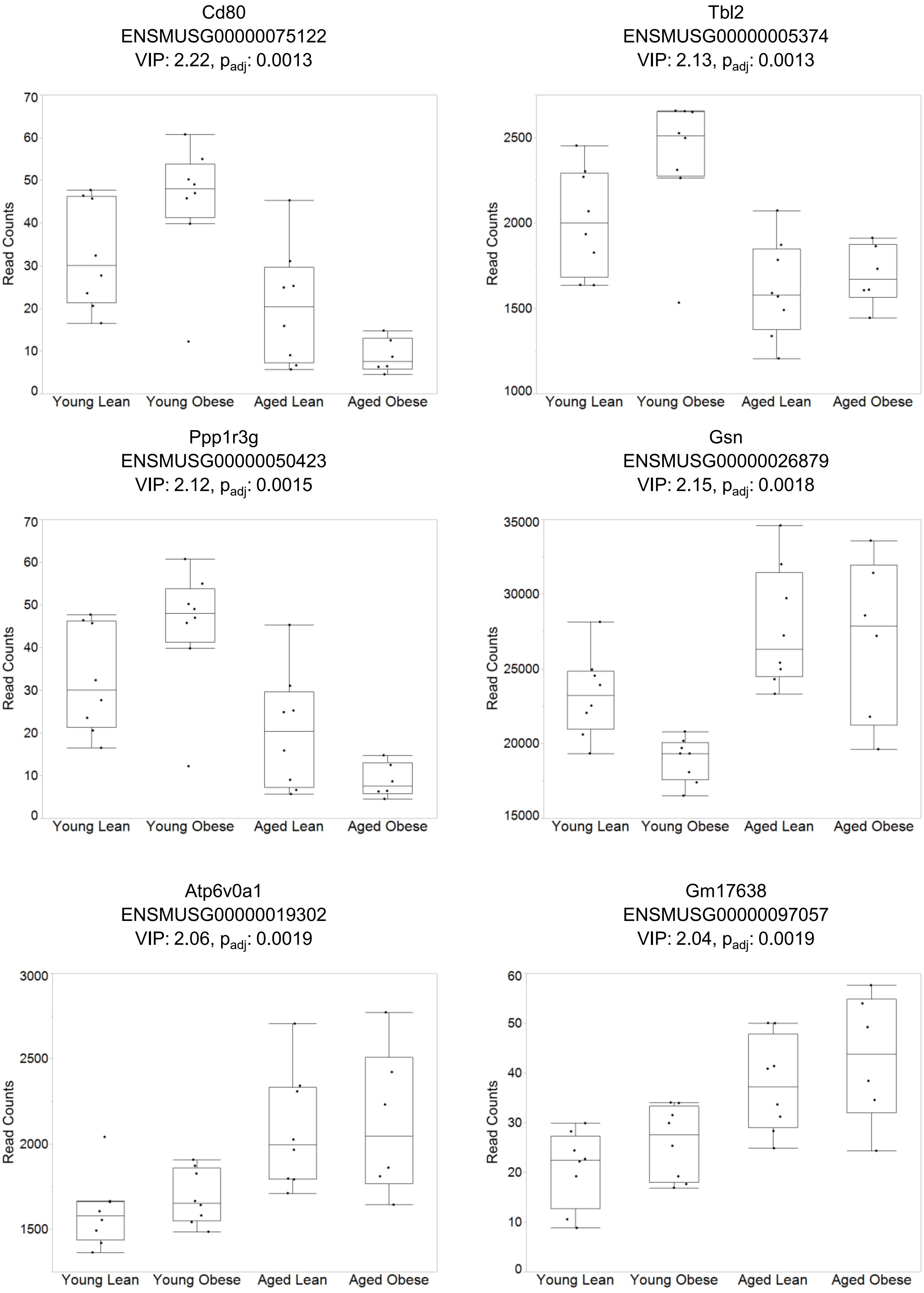
Cd80
ENSMUSG00000075122
VIP: 2.22, p_adj: 0.0013
Read Counts
0 10 20 30 40 50 60 70
Young Lean Young Obese Aged Lean Aged Obese
Tbl2
ENSMUSG00000005374
VIP: 2.13, p_adj: 0.0013
Read Counts
1000 1500 2000 2500
Young Lean Young Obese Aged Lean Aged Obese
Ppp1r3g
ENSMUSG00000050423
VIP: 2.12, p_adj: 0.0015
Read Counts
0 10 20 30 40 50 60 70
Young Lean Young Obese Aged Lean Aged Obese
Gsn
ENSMUSG00000026879
VIP: 2.15, p_adj: 0.0018
Read Counts
15000 20000 25000 30000 35000
Young Lean Young Obese Aged Lean Aged Obese
Atp6v0a1
ENSMUSG00000019302
VIP: 2.06, p_adj: 0.0019
Read Counts
1500 2000 2500 3000
Young Lean Young Obese Aged Lean Aged Obese
Gm17638
ENSMUSG00000097057
VIP: 2.04, p_adj: 0.0019
Read Counts
0 10 20 30 40 50 60
Young Lean Young Obese Aged Lean Aged Obese

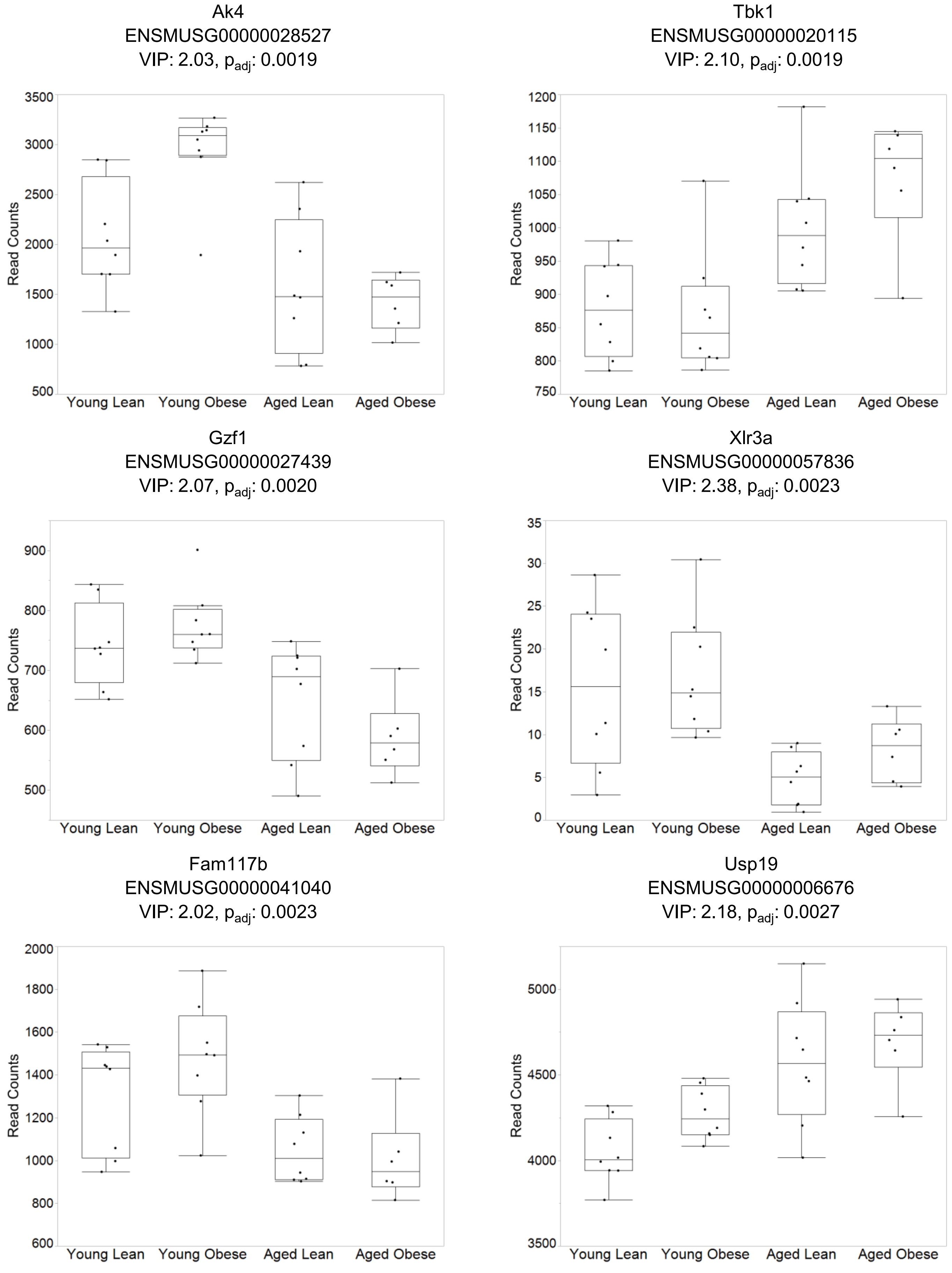
Ak4
ENSMUSG00000028527
VIP: 2.03, $p_{adj}$: 0.0019
Read Counts
500 1000 1500 2000 2500 3000 3500
Young Lean Young Obese Aged Lean Aged Obese
Tbk1
ENSMUSG00000020115
VIP: 2.10, $p_{adj}$: 0.0019
Read Counts
750 800 850 900 950 1000 1050 1100 1150 1200
Young Lean Young Obese Aged Lean Aged Obese
Gzf1
ENSMUSG00000027439
VIP: 2.07, $p_{adj}$: 0.0020
Read Counts
500 600 700 800 900
Young Lean Young Obese Aged Lean Aged Obese
Xlr3a
ENSMUSG00000057836
VIP: 2.38, $p_{adj}$: 0.0023
Read Counts
0 5 10 15 20 25 30 35
Young Lean Young Obese Aged Lean Aged Obese
Fam117b
ENSMUSG00000041040
VIP: 2.02, $p_{adj}$: 0.0023
Read Counts
600 800 1000 1200 1400 1600 1800 2000
Young Lean Young Obese Aged Lean Aged Obese
Usp19
ENSMUSG00000006676
VIP: 2.18, $p_{adj}$: 0.0027
Read Counts
3500 4000 4500 5000
Young Lean Young Obese Aged Lean Aged Obese

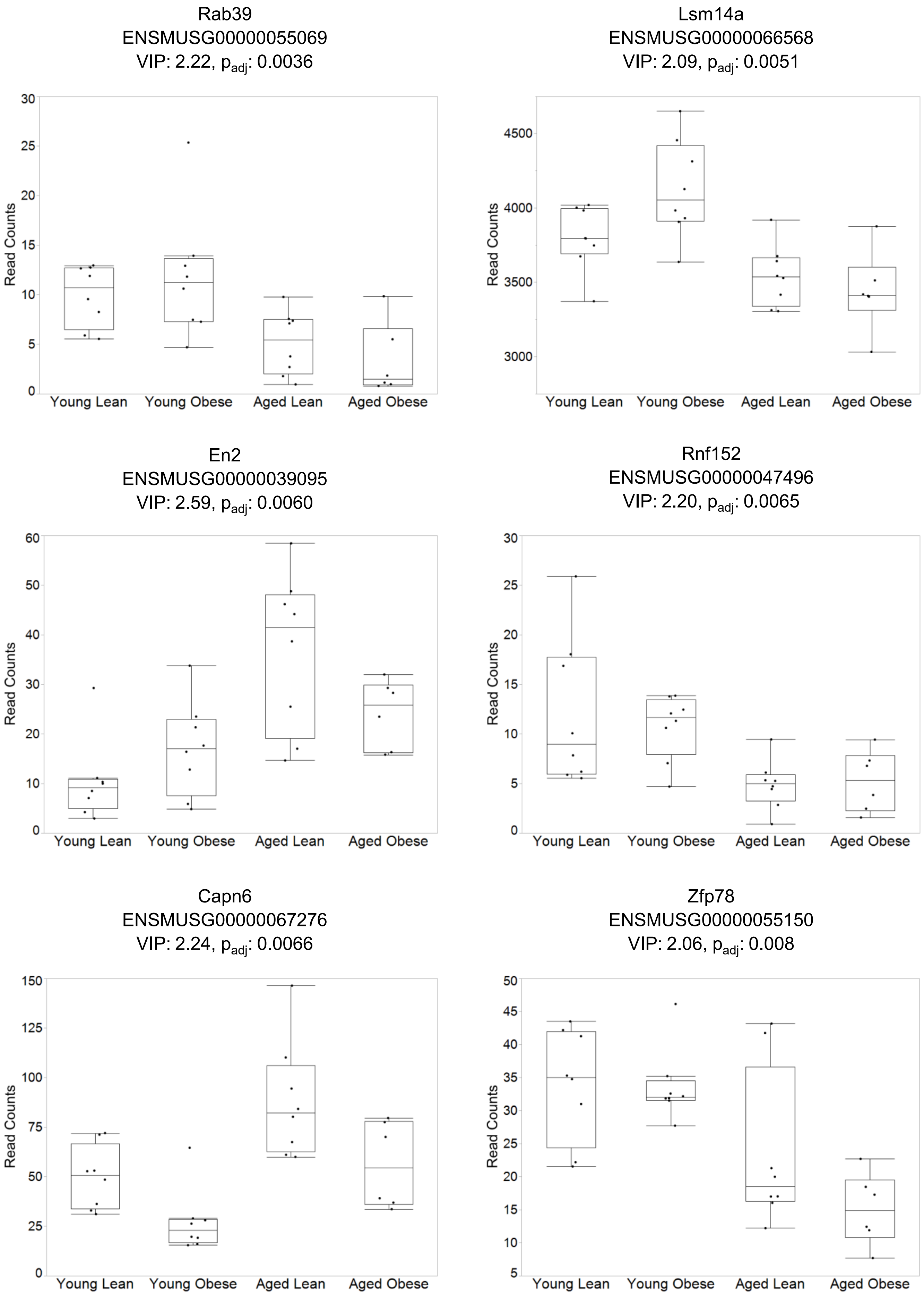
Rab39
ENSMUSG00000055069
VIP: 2.22, p_adj: 0.0036
Lsm14a
ENSMUSG00000066568
VIP: 2.09, p_adj: 0.0051
En2
ENSMUSG00000039095
VIP: 2.59, p_adj: 0.0060
Rnf152
ENSMUSG00000047496
VIP: 2.20, p_adj: 0.0065
Capn6
ENSMUSG00000067276
VIP: 2.24, p_adj: 0.0066
Zfp78
ENSMUSG00000055150
VIP: 2.06, p_adj: 0.008
Read Counts
Young Lean
Young Obese
Aged Lean
Aged Obese

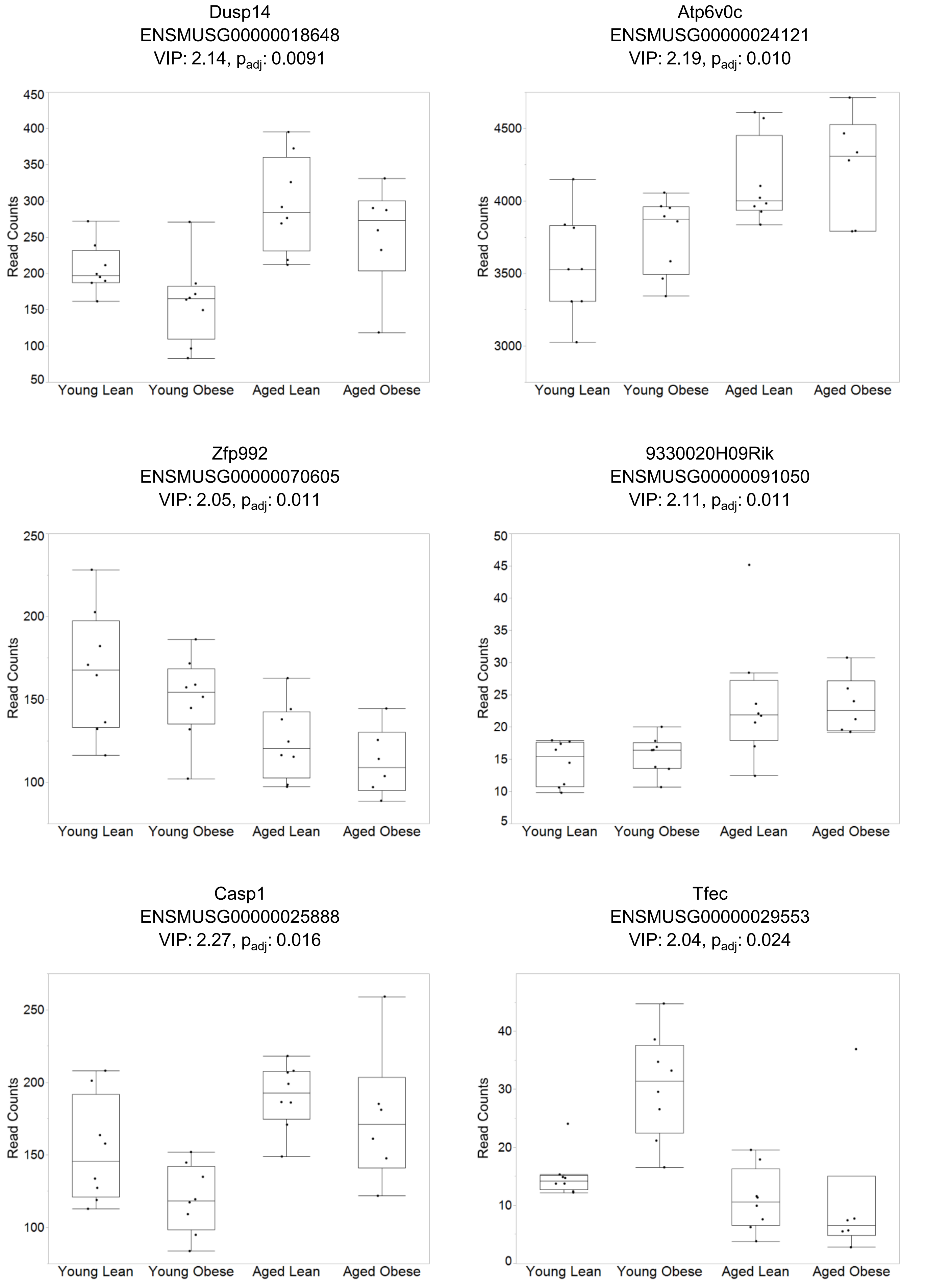
Dusp14
ENSMUSG00000018648
VIP: 2.14, $p_{adj}$: 0.0091
Read Counts
450
400
350
300
250
200
150
100
50
Young Lean
Young Obese
Aged Lean
Aged Obese
Atp6v0c
ENSMUSG00000024121
VIP: 2.19, $p_{adj}$: 0.010
Read Counts
4500
4000
3500
3000
Young Lean
Young Obese
Aged Lean
Aged Obese
Zfp992
ENSMUSG00000070605
VIP: 2.05, $p_{adj}$: 0.011
Read Counts
250
200
150
100
Young Lean
Young Obese
Aged Lean
Aged Obese
9330020H09Rik
ENSMUSG00000091050
VIP: 2.11, $p_{adj}$: 0.011
Read Counts
50
45
40
35
30
25
20
15
10
5
Young Lean
Young Obese
Aged Lean
Aged Obese
Casp1
ENSMUSG00000025888
VIP: 2.27, $p_{adj}$: 0.016
Read Counts
250
200
150
100
Young Lean
Young Obese
Aged Lean
Aged Obese
Tfec
ENSMUSG00000029553
VIP: 2.04, $p_{adj}$: 0.024
Read Counts
40
30
20
10
0
Young Lean
Young Obese
Aged Lean
Aged Obese

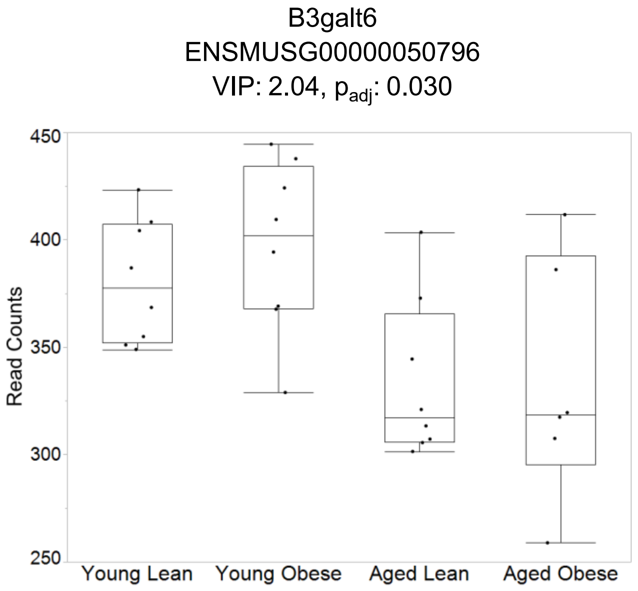
B3galt6
ENSMUSG00000050796
VIP: 2.04, $p_{adj}$: 0.030
Read Counts
450
400
350
300
250
Young Lean
Young Obese
Aged Lean
Aged Obese

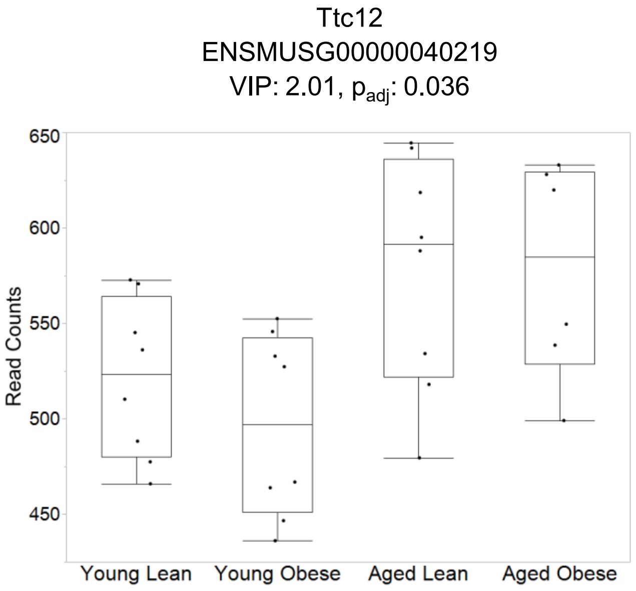
Ttc12
ENSMUSG00000040219
VIP: 2.01, $p_{adj}$: 0.036
Read Counts
650
600
550
500
450
Young Lean
Young Obese
Aged Lean
Aged Obese

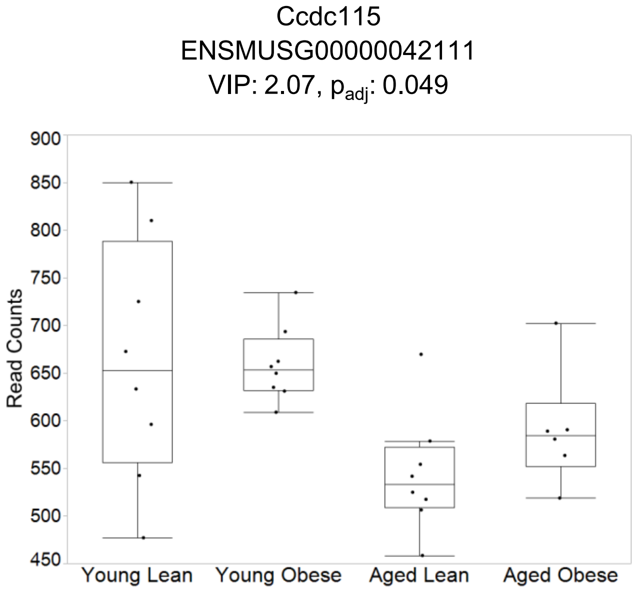
Ccdc115
ENSMUSG00000042111
VIP: 2.07, $p_{adj}$: 0.049
Read Counts
900
850
800
750
700
650
600
550
500
450
Young Lean
Young Obese
Aged Lean
Aged Obese
