## Supplemental Figure 5 for "Aging and Obesity Prime the Methylome and Transcriptome of Adipose Stem Cells for Disease and Dysfunction"

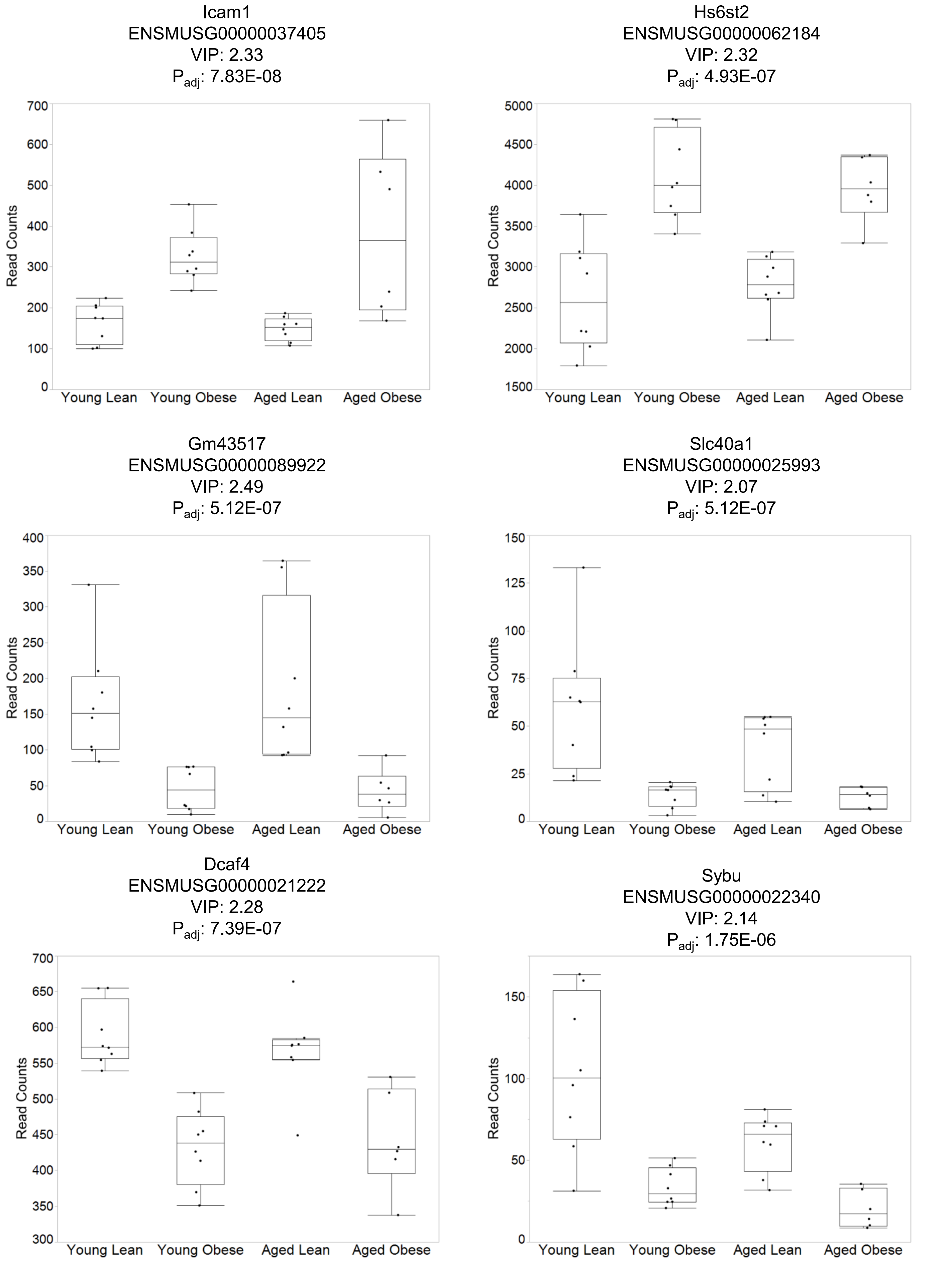
Icam1
ENSMUSG00000037405
VIP: 2.33
$P_{adj}$: 7.83E-08
Read Counts
0
100
200
300
400
500
600
700
Young Lean
Young Obese
Aged Lean
Aged Obese
Hs6st2
ENSMUSG00000062184
VIP: 2.32
$P_{adj}$: 4.93E-07
Read Counts
1500
2000
2500
3000
3500
4000
4500
5000
Young Lean
Young Obese
Aged Lean
Aged Obese
Gm43517
ENSMUSG00000089922
VIP: 2.49
$P_{adj}$: 5.12E-07
Read Counts
0
50
100
150
200
250
300
350
400
Young Lean
Young Obese
Aged Lean
Aged Obese
Slc40a1
ENSMUSG00000025993
VIP: 2.07
$P_{adj}$: 5.12E-07
Read Counts
0
25
50
75
100
125
150
Young Lean
Young Obese
Aged Lean
Aged Obese
Dcaf4
ENSMUSG00000021222
VIP: 2.28
$P_{adj}$: 7.39E-07
Read Counts
300
350
400
450
500
550
600
650
700
Young Lean
Young Obese
Aged Lean
Aged Obese
Sybu
ENSMUSG00000022340
VIP: 2.14
$P_{adj}$: 1.75E-06
Read Counts
0
50
100
150
Young Lean
Young Obese
Aged Lean
Aged Obese

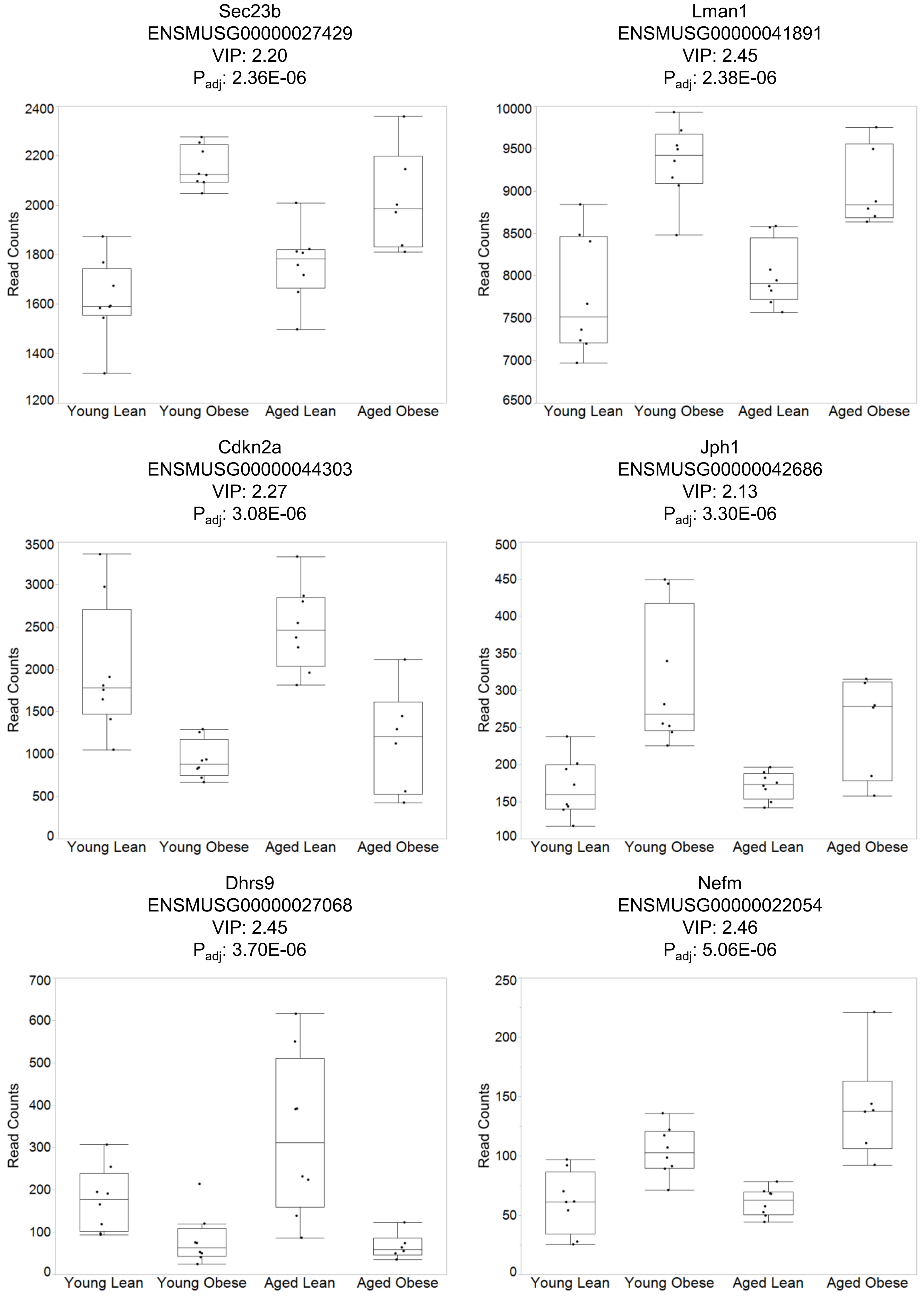

Sec23b
ENSMUSG00000027429
VIP: 2.20
P_adj: 2.36E-06
Read Counts
1200
1400
1600
1800
2000
2200
2400
Young Lean
Young Obese
Aged Lean
Aged Obese
Lman1
ENSMUSG00000041891
VIP: 2.45
P_adj: 2.38E-06
Read Counts
6500
7000
7500
8000
8500
9000
9500
10000
Young Lean
Young Obese
Aged Lean
Aged Obese
Cdkn2a
ENSMUSG00000044303
VIP: 2.27
P_adj: 3.08E-06
Read Counts
0
500
1000
1500
2000
2500
3000
3500
Young Lean
Young Obese
Aged Lean
Aged Obese
Jph1
ENSMUSG00000042686
VIP: 2.13
P_adj: 3.30E-06
Read Counts
100
150
200
250
300
350
400
450
500
Young Lean
Young Obese
Aged Lean
Aged Obese
Dhrs9
ENSMUSG00000027068
VIP: 2.45
P_adj: 3.70E-06
Read Counts
0
100
200
300
400
500
600
700
Young Lean
Young Obese
Aged Lean
Aged Obese
Nefm
ENSMUSG00000022054
VIP: 2.46
P_adj: 5.06E-06
Read Counts
0
50
100
150
200
250
Young Lean
Young Obese
Aged Lean
Aged Obese

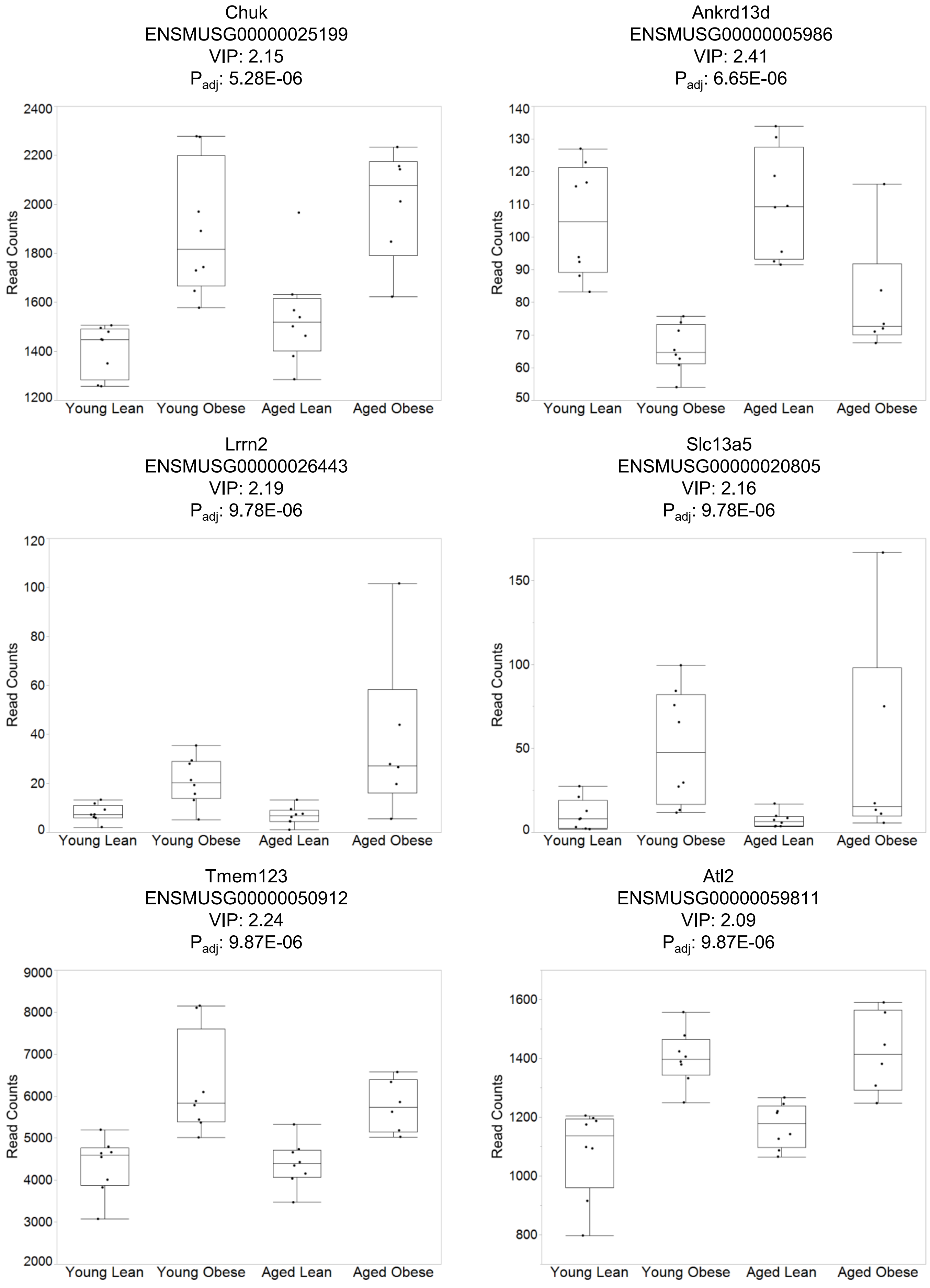
Chuk
ENSMUSG00000025199
VIP: 2.15
Padj: 5.28E-06
Read Counts
2400
2200
2000
1800
1600
1400
1200
Young Lean
Young Obese
Aged Lean
Aged Obese
Ankrd13d
ENSMUSG00000005986
VIP: 2.41
Padj: 6.65E-06
Read Counts
140
130
120
110
100
90
80
70
60
50
Young Lean
Young Obese
Aged Lean
Aged Obese
Lrrn2
ENSMUSG00000026443
VIP: 2.19
Padj: 9.78E-06
Read Counts
120
100
80
60
40
20
0
Young Lean
Young Obese
Aged Lean
Aged Obese
Slc13a5
ENSMUSG00000020805
VIP: 2.16
Padj: 9.78E-06
Read Counts
150
100
50
0
Young Lean
Young Obese
Aged Lean
Aged Obese
Tmem123
ENSMUSG00000050912
VIP: 2.24
Padj: 9.87E-06
Read Counts
9000
8000
7000
6000
5000
4000
3000
2000
Young Lean
Young Obese
Aged Lean
Aged Obese
Atl2
ENSMUSG00000059811
VIP: 2.09
Padj: 9.87E-06
Read Counts
1600
1400
1200
1000
800
Young Lean
Young Obese
Aged Lean
Aged Obese

Echdc3
ENSMUSG00000039063
VIP: 2.05
$P_{adj}$: 1.36E-05

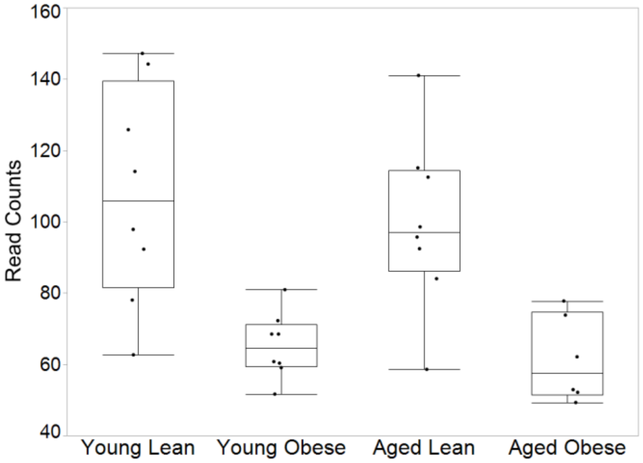


Khk
ENSMUSG00000029162
VIP: 2.12
$P_{adj}$: 1.59E-05

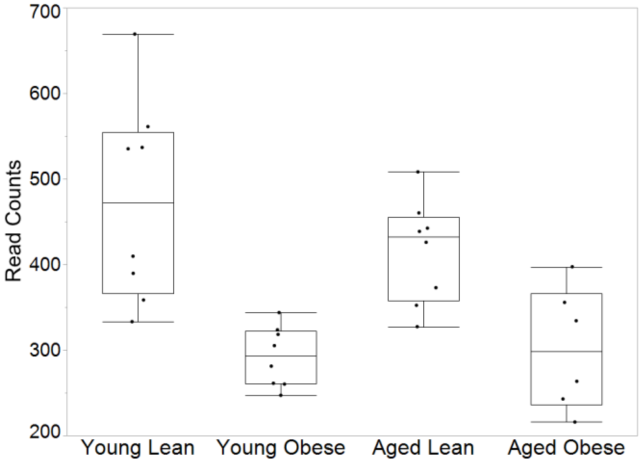


Rell1
ENSMUSG00000047881
VIP: 2.09
$P_{adj}$: 1.68E-05

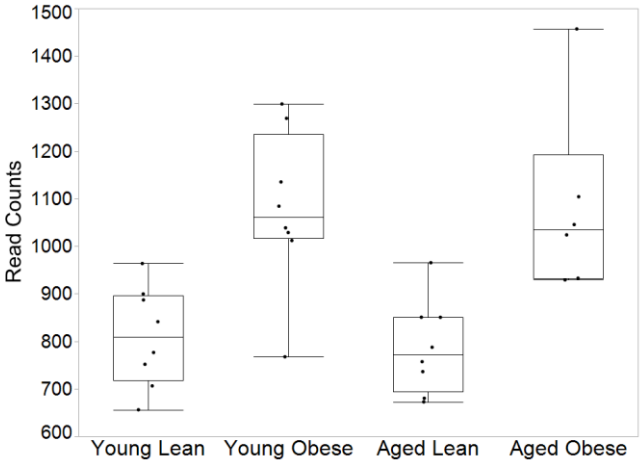


Ephx1
ENSMUSG00000038776
VIP: 2.20
$P_{adj}$: 1.71E-05

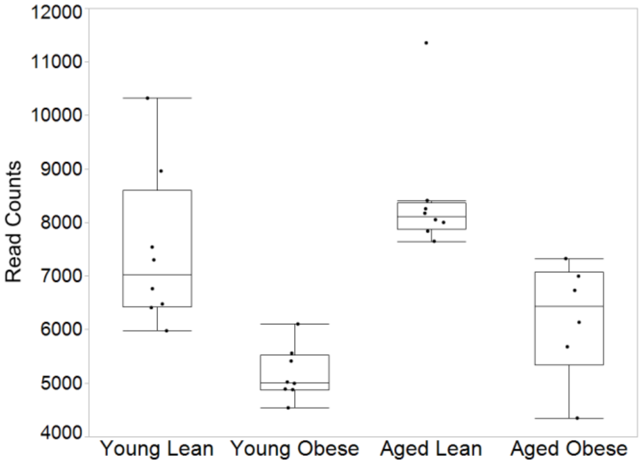


Foxg1
ENSMUSG00000020950
VIP: 2.26
$P_{adj}$: 2.13E-05

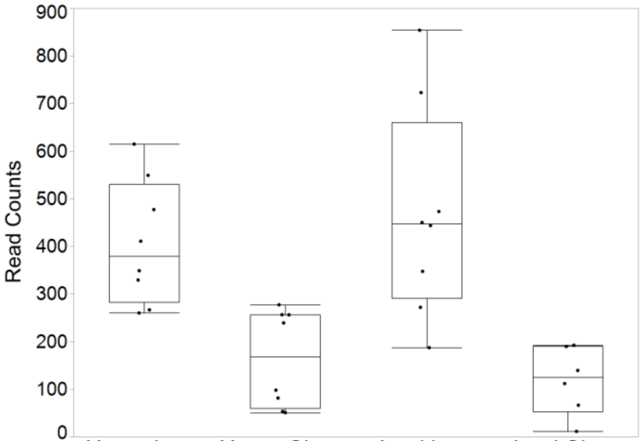


Oas1c
ENSMUSG00000001166
VIP: 2.06
$P_{adj}$: 2.78E-05

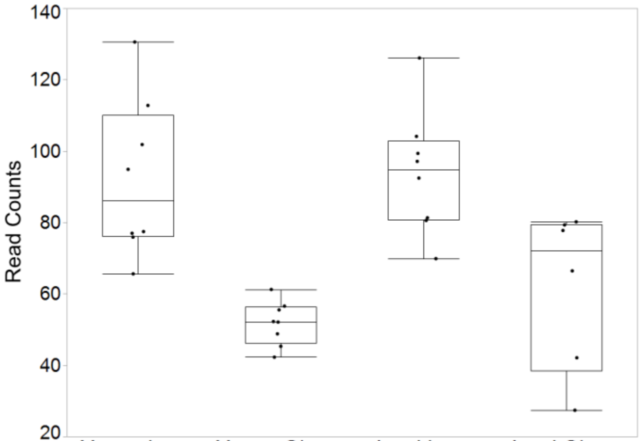

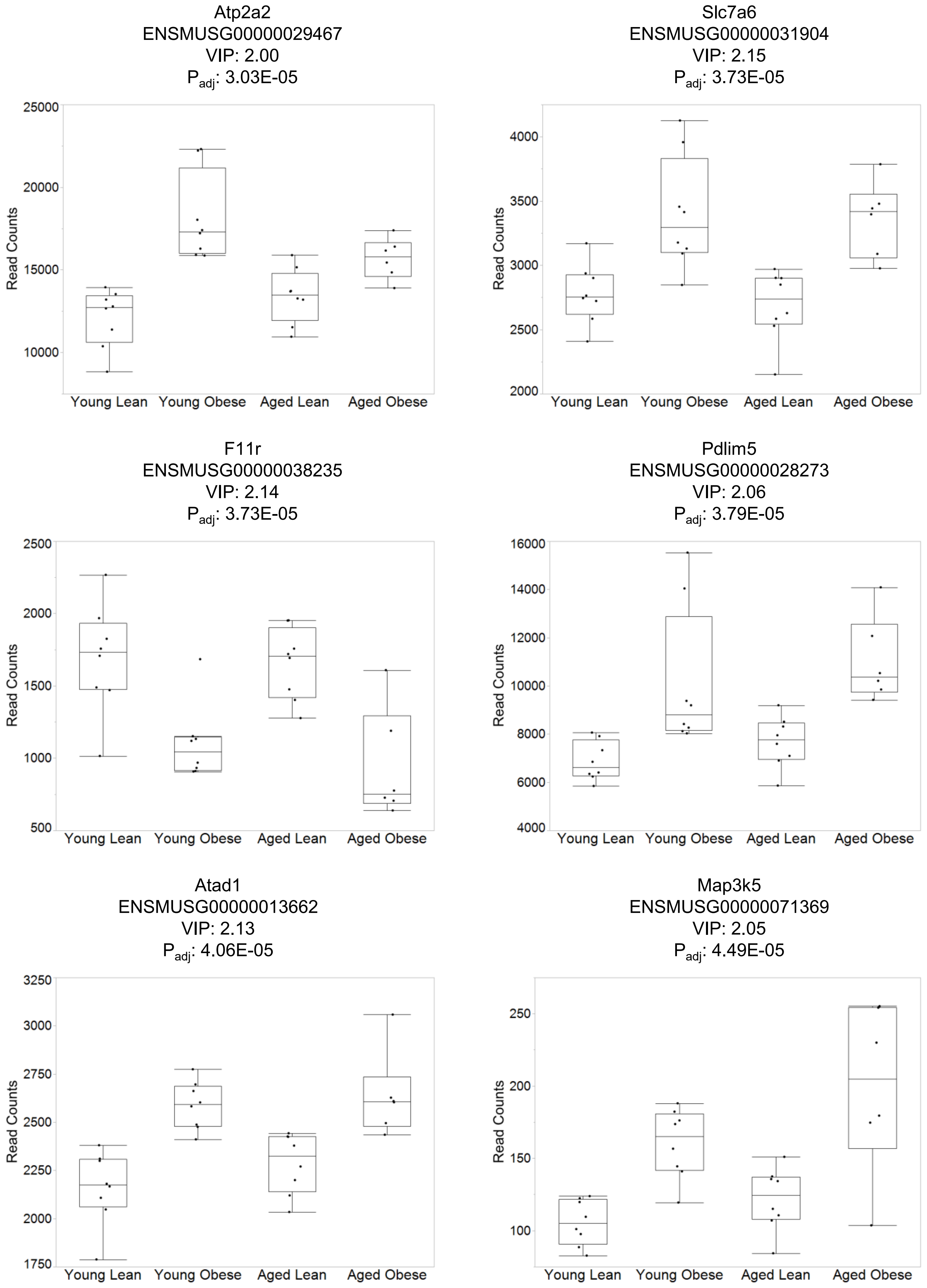

Atp2a2
ENSMUSG00000029467
VIP: 2.00
P adj: 3.03E-05
Read Counts
25000
20000
15000
10000
Young Lean
Young Obese
Aged Lean
Aged Obese
Slc7a6
ENSMUSG00000031904
VIP: 2.15
P adj: 3.73E-05
Read Counts
4000
3500
3000
2500
2000
Young Lean
Young Obese
Aged Lean
Aged Obese
F11r
ENSMUSG00000038235
VIP: 2.14
P adj: 3.73E-05
Read Counts
2500
2000
1500
1000
500
Young Lean
Young Obese
Aged Lean
Aged Obese
Pdlim5
ENSMUSG00000028273
VIP: 2.06
P adj: 3.79E-05
Read Counts
16000
14000
12000
10000
8000
6000
4000
Young Lean
Young Obese
Aged Lean
Aged Obese
Atad1
ENSMUSG00000013662
VIP: 2.13
P adj: 4.06E-05
Read Counts
3250
3000
2750
2500
2250
2000
1750
Young Lean
Young Obese
Aged Lean
Aged Obese
Map3k5
ENSMUSG00000071369
VIP: 2.05
P adj: 4.49E-05
Read Counts
250
200
150
100
Young Lean
Young Obese
Aged Lean
Aged Obese

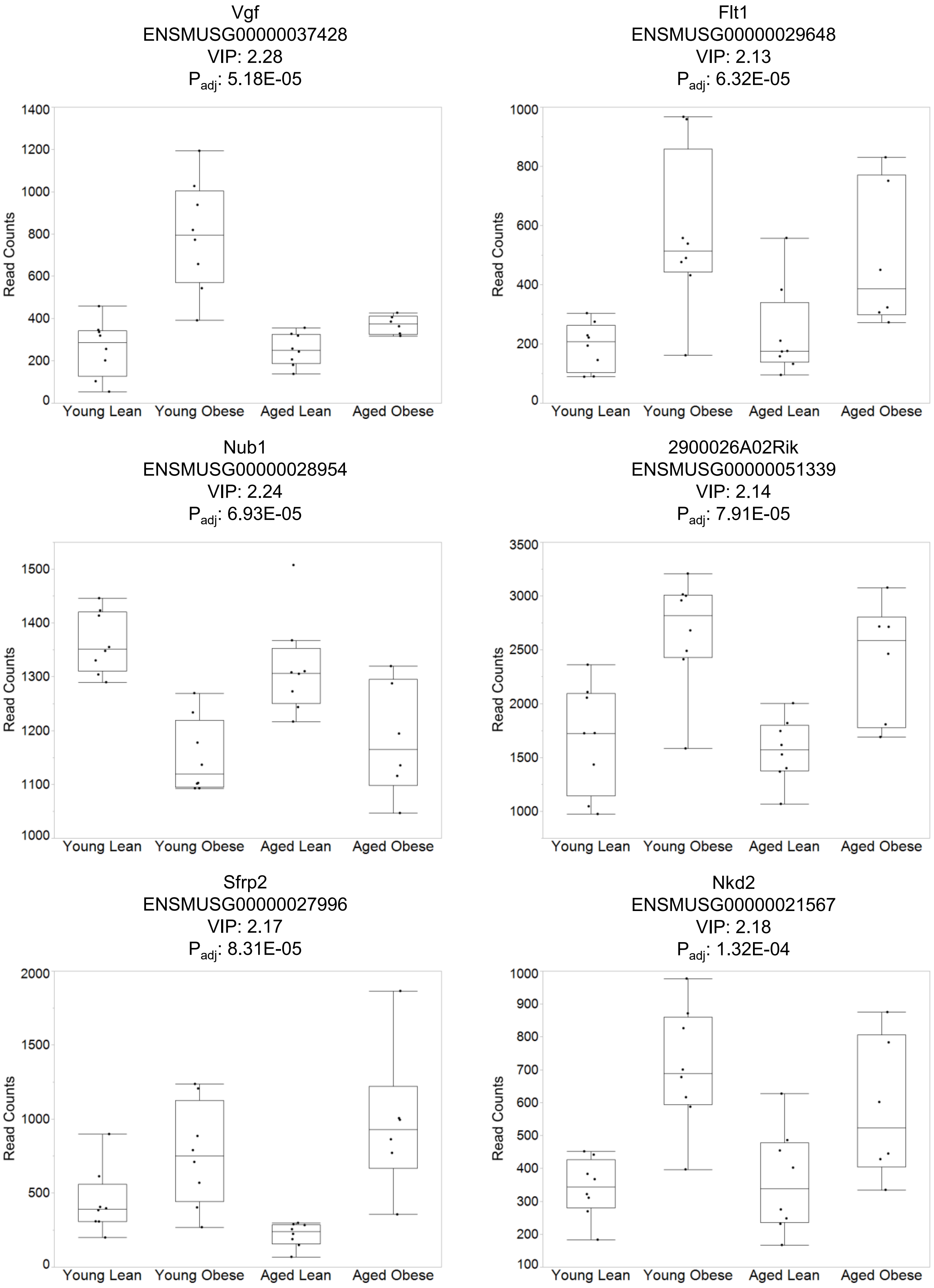
Vgf
ENSMUSG00000037428
VIP: 2.28
Padj: 5.18E-05
Read Counts
Young Lean
Young Obese
Aged Lean
Aged Obese
Flt1
ENSMUSG00000029648
VIP: 2.13
Padj: 6.32E-05
Nub1
ENSMUSG00000028954
VIP: 2.24
Padj: 6.93E-05
2900026A02Rik
ENSMUSG00000051339
VIP: 2.14
Padj: 7.91E-05
Sfrp2
ENSMUSG00000027996
VIP: 2.17
Padj: 8.31E-05
Nkd2
ENSMUSG00000021567
VIP: 2.18
Padj: 1.32E-04

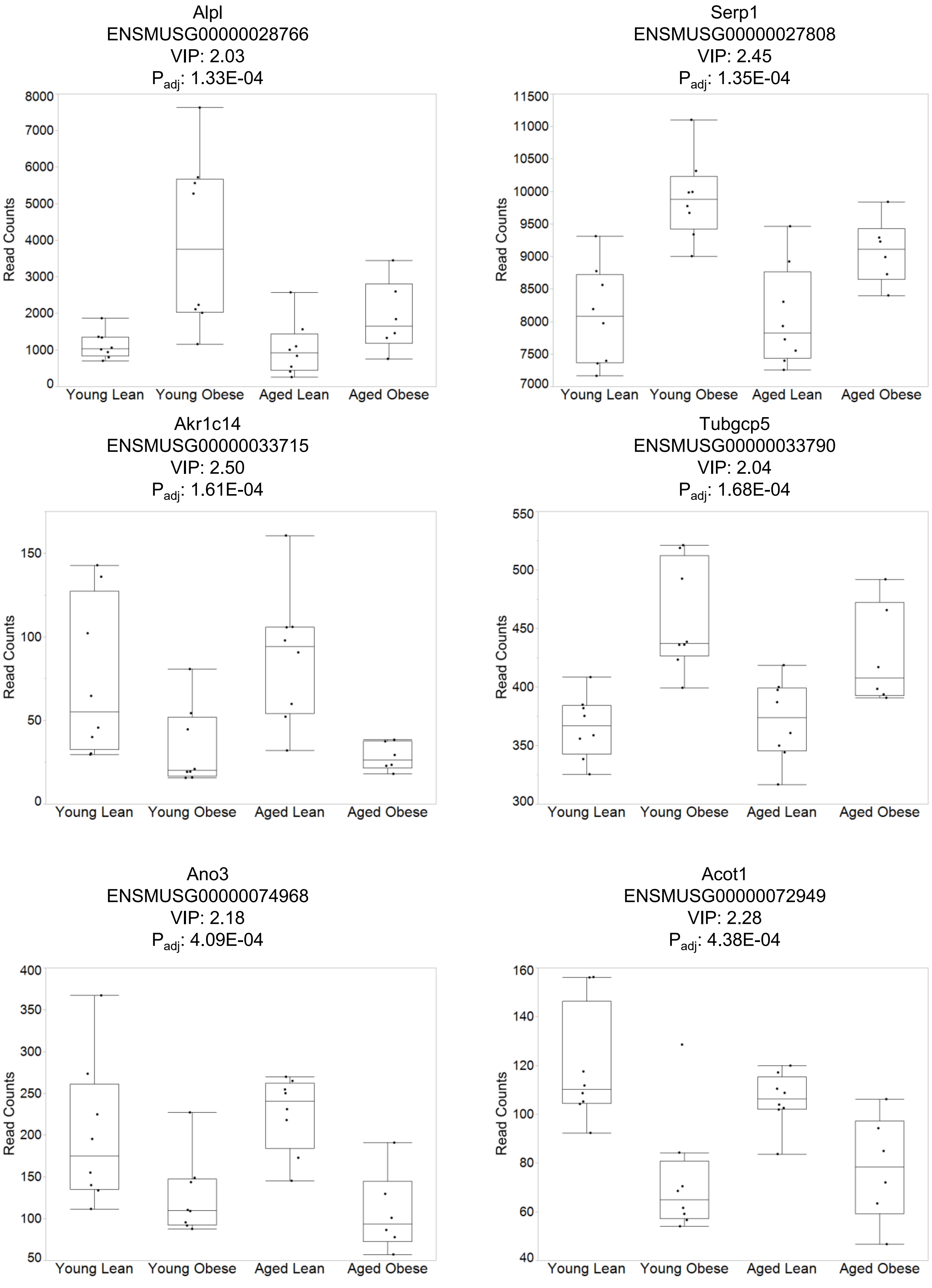
Alpl
ENSMUSG00000028766
VIP: 2.03
P_adj: 1.33E-04
Read Counts
0
1000
2000
3000
4000
5000
6000
7000
8000
Young Lean
Young Obese
Aged Lean
Aged Obese
Serp1
ENSMUSG00000027808
VIP: 2.45
P_adj: 1.35E-04
Read Counts
7000
7500
8000
8500
9000
9500
10000
10500
11000
11500
Young Lean
Young Obese
Aged Lean
Aged Obese
Akr1c14
ENSMUSG00000033715
VIP: 2.50
P_adj: 1.61E-04
Read Counts
0
50
100
150
Young Lean
Young Obese
Aged Lean
Aged Obese
Tubgcp5
ENSMUSG00000033790
VIP: 2.04
P_adj: 1.68E-04
Read Counts
300
350
400
450
500
550
Young Lean
Young Obese
Aged Lean
Aged Obese
Ano3
ENSMUSG00000074968
VIP: 2.18
P_adj: 4.09E-04
Read Counts
50
100
150
200
250
300
350
400
Young Lean
Young Obese
Aged Lean
Aged Obese
Acot1
ENSMUSG00000072949
VIP: 2.28
P_adj: 4.38E-04
Read Counts
40
60
80
100
120
140
160
Young Lean
Young Obese
Aged Lean
Aged Obese

Trpc6
ENSMUSG00000031997
VIP: 2.18
$P_{adj}$: 4.91E-04

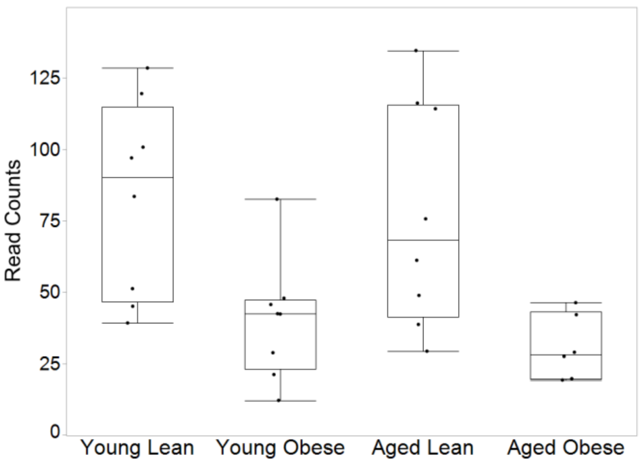


Acaa2
ENSMUSG00000036880
VIP: 2.09
$P_{adj}$: 5.42E-04

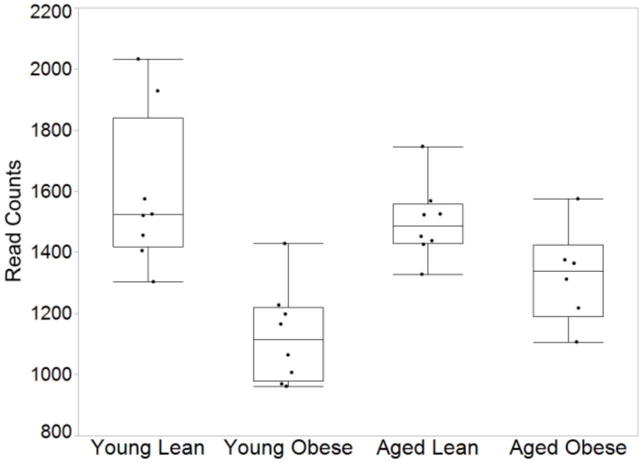


Tmem47
ENSMUSG00000025666
VIP: 2.22
$P_{adj}$: 6.43E-04

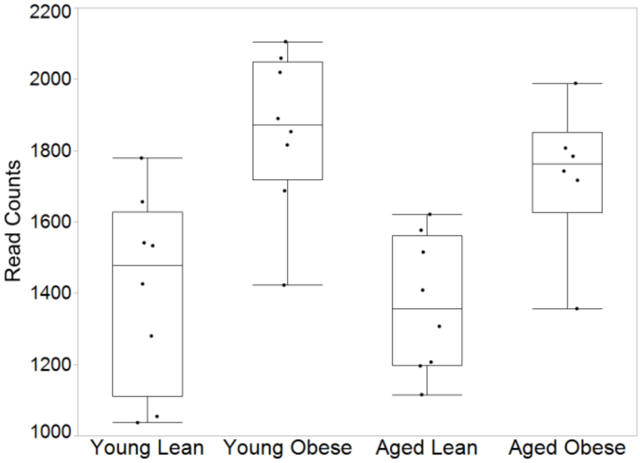


A630033H20Rik
ENSMUSG00000054293
VIP: 2.11
$P_{adj}$: 7.85E-04

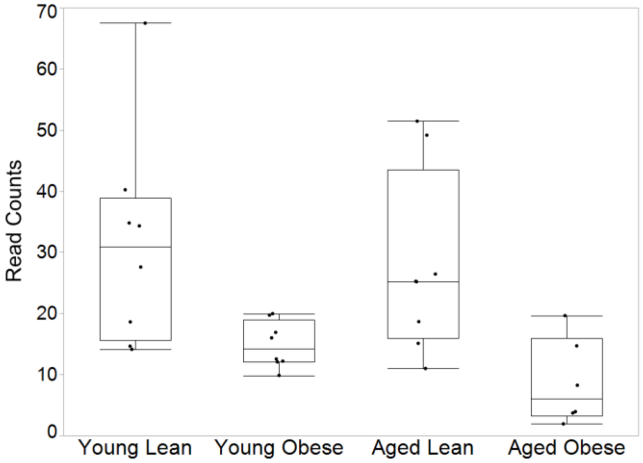


Marc1
ENSMUSG00000026621
VIP: 2.30
$P_{adj}$: 7.99E-04

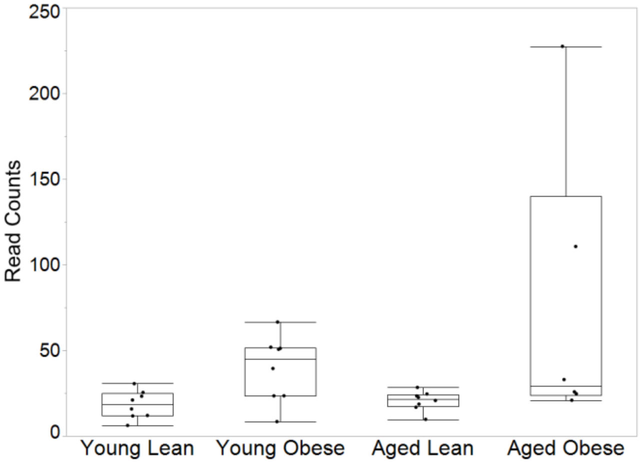


Psrc1
ENSMUSG00000068744
VIP: 2.09
$P_{adj}$: 8.28E-04

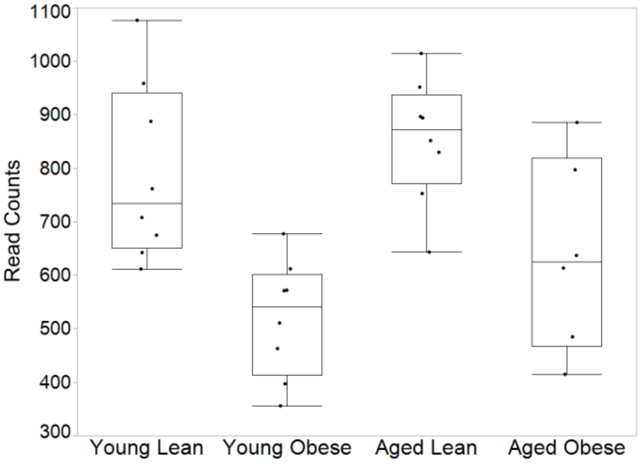

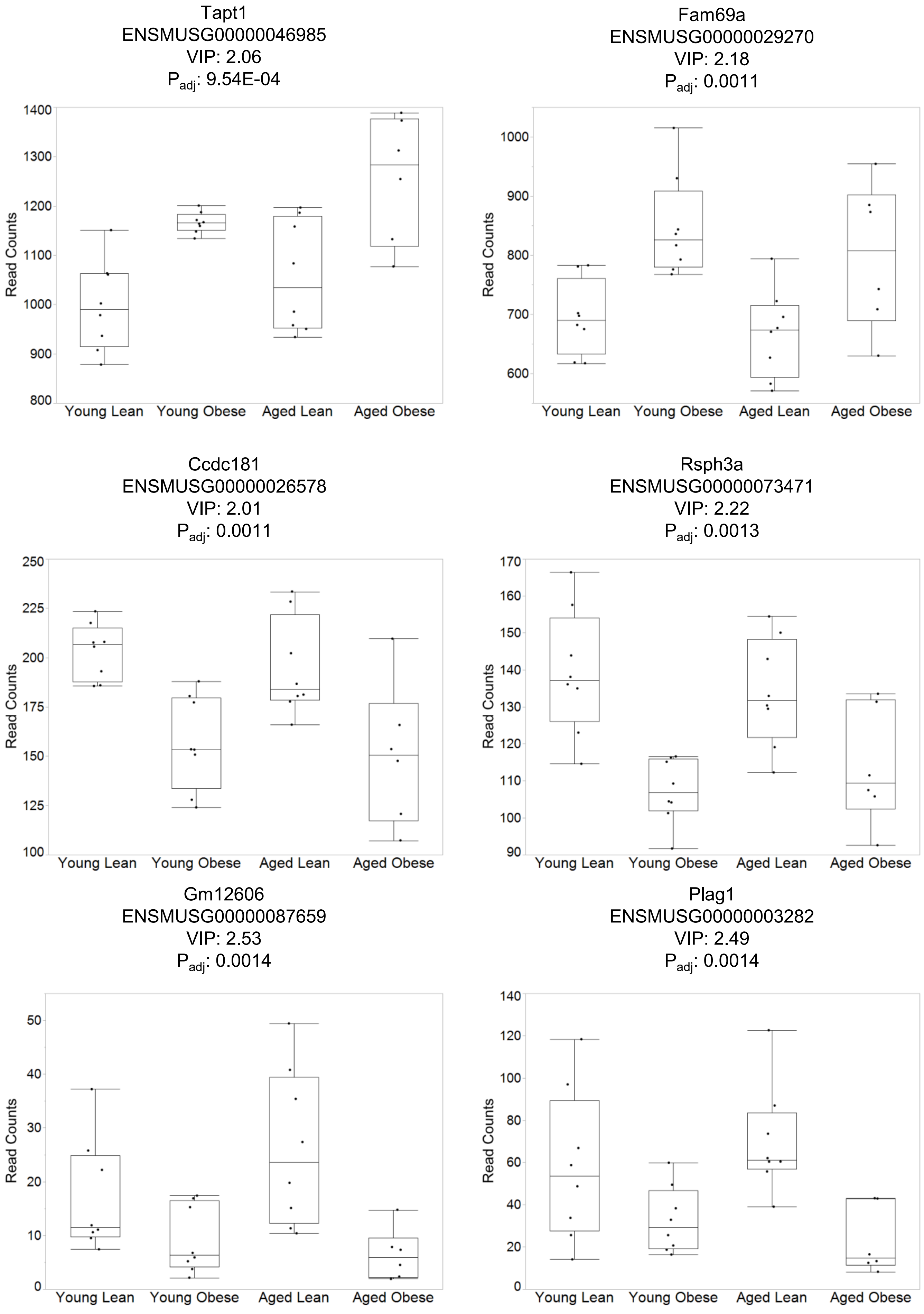
Tapt1
ENSMUSG00000046985
VIP: 2.06
P_adj: 9.54E-04
Read Counts
1400
1300
1200
1100
1000
900
800
Young Lean
Young Obese
Aged Lean
Aged Obese
Fam69a
ENSMUSG00000029270
VIP: 2.18
P_adj: 0.0011
Read Counts
1000
900
800
700
600
Young Lean
Young Obese
Aged Lean
Aged Obese
Ccdc181
ENSMUSG00000026578
VIP: 2.01
P_adj: 0.0011
Read Counts
250
225
200
175
150
125
100
Young Lean
Young Obese
Aged Lean
Aged Obese
Rsph3a
ENSMUSG00000073471
VIP: 2.22
P_adj: 0.0013
Read Counts
170
160
150
140
130
120
110
100
90
Young Lean
Young Obese
Aged Lean
Aged Obese
Gm12606
ENSMUSG00000087659
VIP: 2.53
P_adj: 0.0014
Read Counts
50
40
30
20
10
0
Young Lean
Young Obese
Aged Lean
Aged Obese
Plag1
ENSMUSG00000003282
VIP: 2.49
P_adj: 0.0014
Read Counts
140
120
100
80
60
40
20
0
Young Lean
Young Obese
Aged Lean
Aged Obese

Alcam
ENSMUSG00000022636
VIP: 2.16
$P_{adj}$: 0.0014

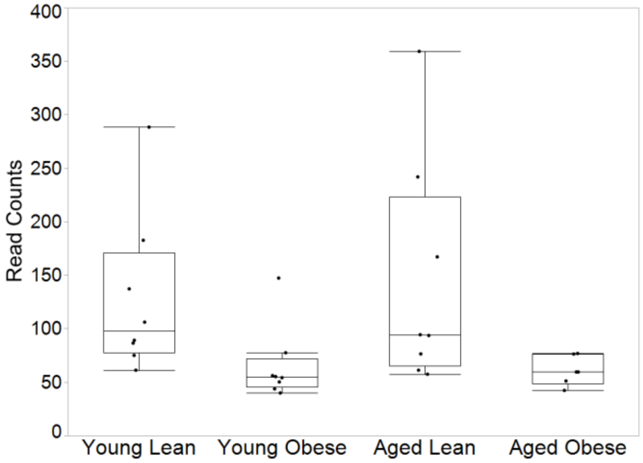


M6pr
ENSMUSG00000007458
VIP: 2.16
$P_{adj}$: 0.0015

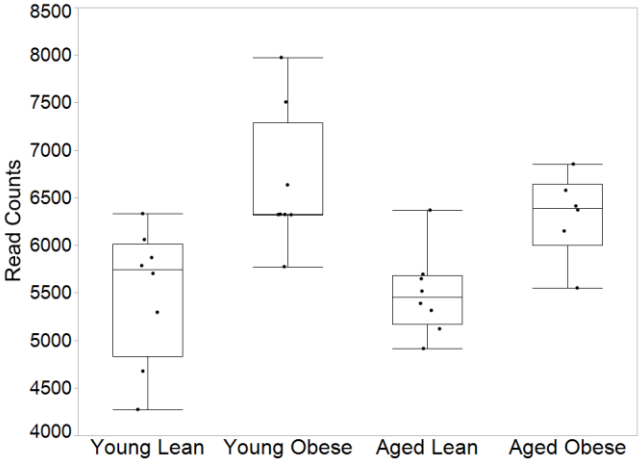


Hyou1
ENSMUSG00000032115
VIP: 2.32
$P_{adj}$: 0.0015

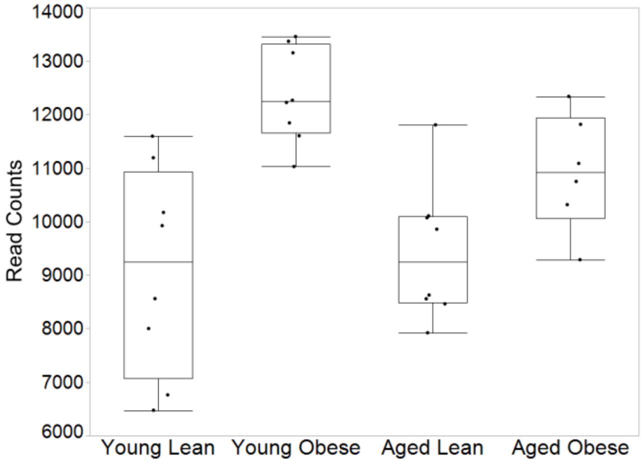


Ube2f
ENSMUSG00000034343
VIP: 2.12
$P_{adj}$: 0.0015

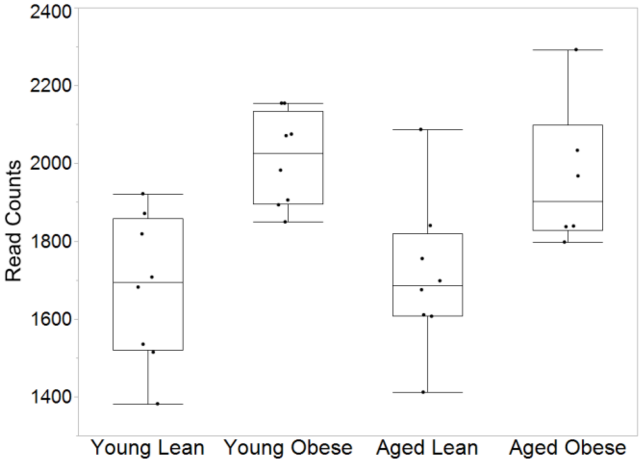


Gzmd
ENSMUSG00000059256
VIP: 2.07
$P_{adj}$: 0.0016

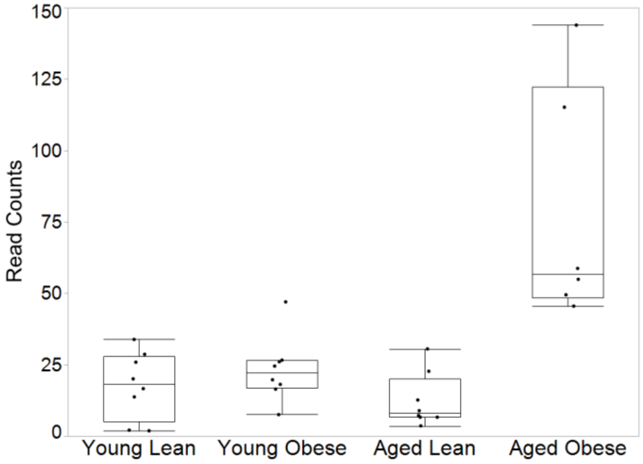


Pax3
ENSMUSG00000004872
VIP: 2.12
$P_{adj}$: 0.0018

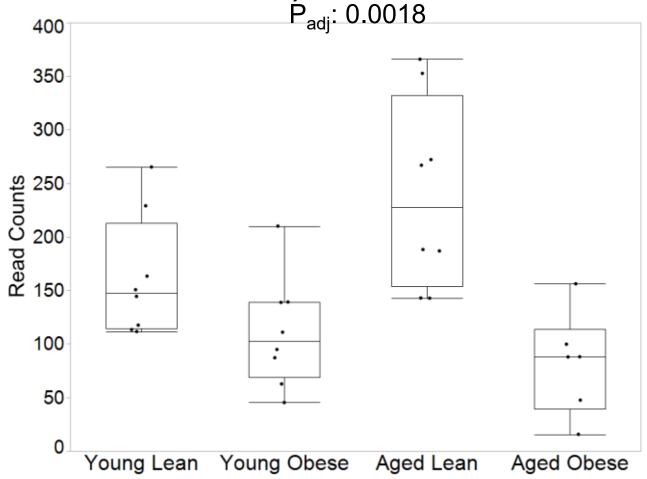

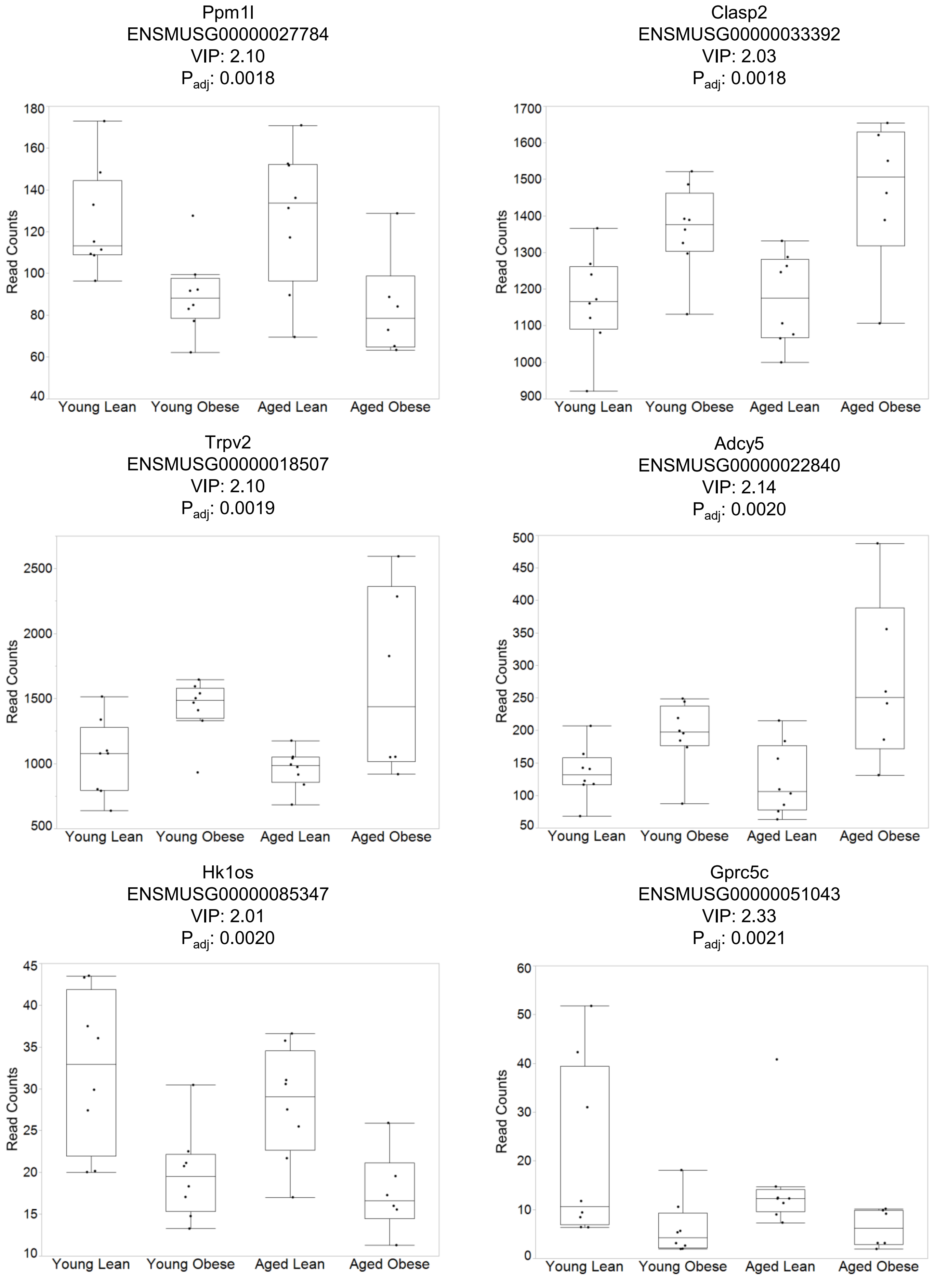
Ppm1l
ENSMUSG00000027784
VIP: 2.10
Padj: 0.0018
Read Counts
180
160
140
120
100
80
60
40
Young Lean
Young Obese
Aged Lean
Aged Obese
Clasp2
ENSMUSG00000033392
VIP: 2.03
Padj: 0.0018
Read Counts
1700
1600
1500
1400
1300
1200
1100
1000
900
Young Lean
Young Obese
Aged Lean
Aged Obese
Trpv2
ENSMUSG00000018507
VIP: 2.10
Padj: 0.0019
Read Counts
2500
2000
1500
1000
500
Young Lean
Young Obese
Aged Lean
Aged Obese
Adcy5
ENSMUSG00000022840
VIP: 2.14
Padj: 0.0020
Read Counts
500
450
400
350
300
250
200
150
100
50
Young Lean
Young Obese
Aged Lean
Aged Obese
Hk1os
ENSMUSG00000085347
VIP: 2.01
Padj: 0.0020
Read Counts
45
40
35
30
25
20
15
10
Young Lean
Young Obese
Aged Lean
Aged Obese
Gprc5c
ENSMUSG00000051043
VIP: 2.33
Padj: 0.0021
Read Counts
60
50
40
30
20
10
0
Young Lean
Young Obese
Aged Lean
Aged Obese

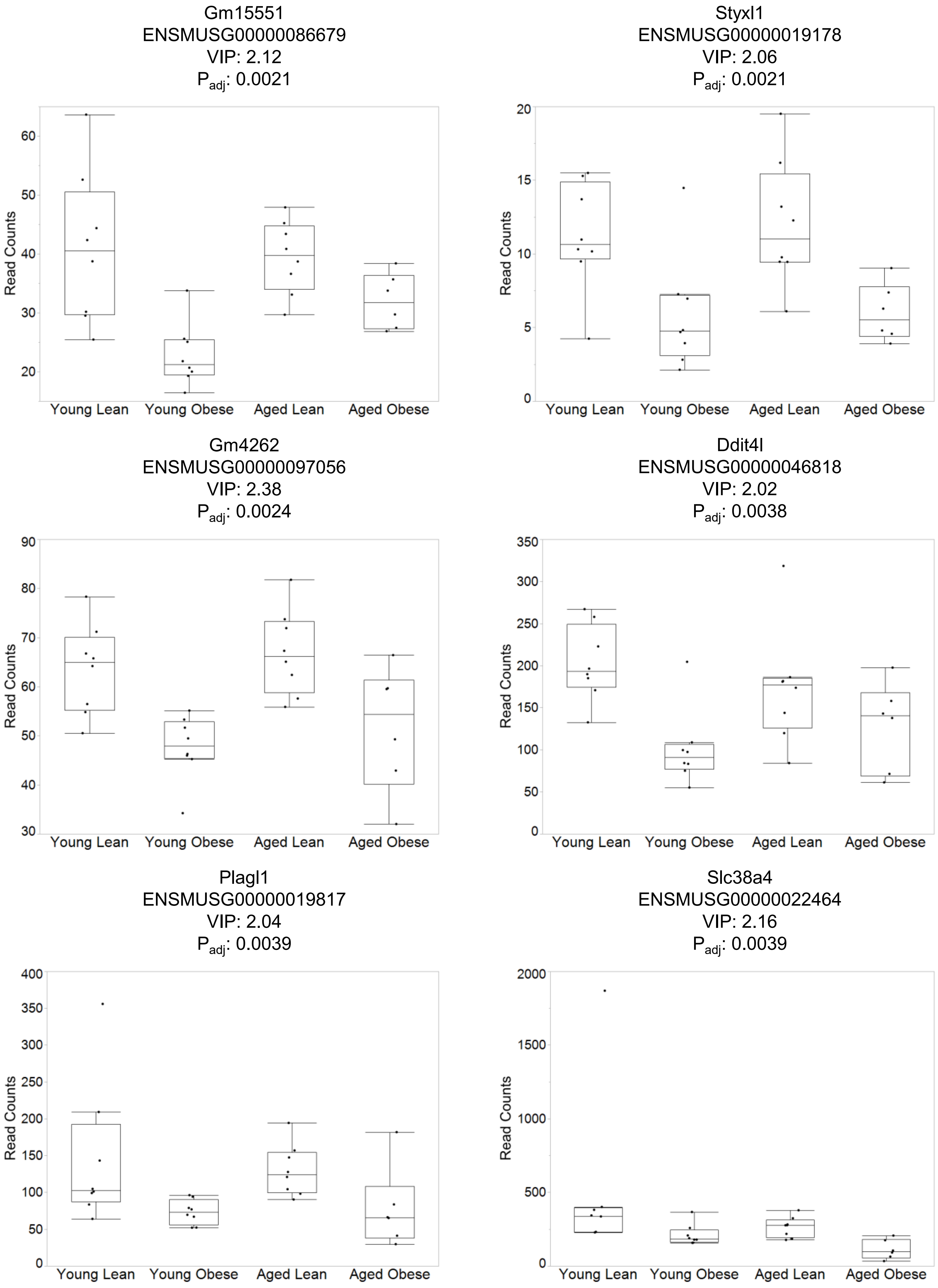
Gm15551
ENSMUSG00000086679
VIP: 2.12
$P_{adj}$: 0.0021
Read Counts
20
30
40
50
60
Young Lean
Young Obese
Aged Lean
Aged Obese
Styxl1
ENSMUSG00000019178
VIP: 2.06
$P_{adj}$: 0.0021
Read Counts
0
5
10
15
20
Young Lean
Young Obese
Aged Lean
Aged Obese
Gm4262
ENSMUSG00000097056
VIP: 2.38
$P_{adj}$: 0.0024
Read Counts
30
40
50
60
70
80
90
Young Lean
Young Obese
Aged Lean
Aged Obese
Ddit4l
ENSMUSG00000046818
VIP: 2.02
$P_{adj}$: 0.0038
Read Counts
0
50
100
150
200
250
300
350
Young Lean
Young Obese
Aged Lean
Aged Obese
Plagl1
ENSMUSG00000019817
VIP: 2.04
$P_{adj}$: 0.0039
Read Counts
0
50
100
150
200
250
300
350
400
Young Lean
Young Obese
Aged Lean
Aged Obese
Slc38a4
ENSMUSG00000022464
VIP: 2.16
$P_{adj}$: 0.0039
Read Counts
0
500
1000
1500
2000
Young Lean
Young Obese
Aged Lean
Aged Obese

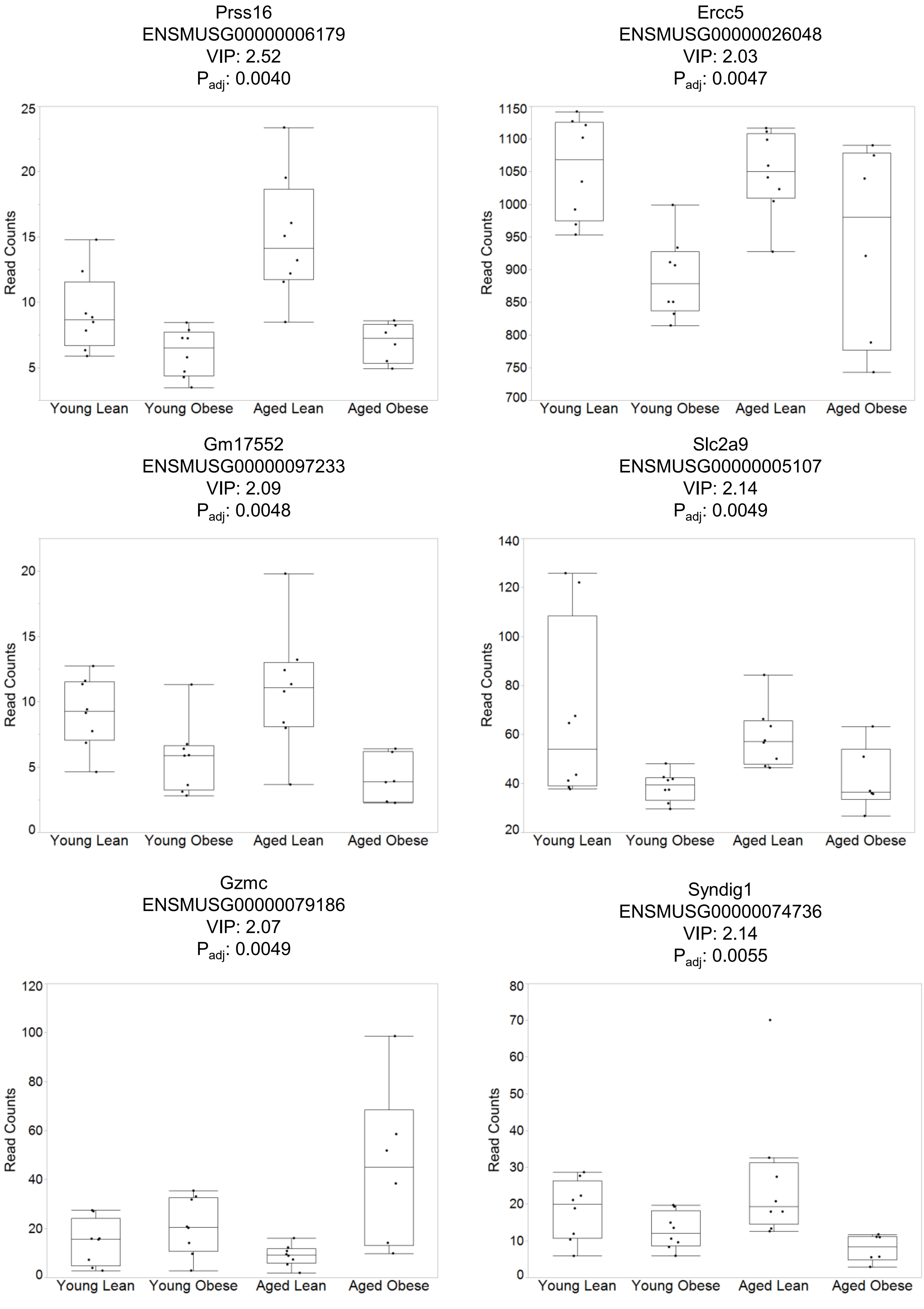
Prss16
ENSMUSG00000006179
VIP: 2.52
Padj: 0.0040
Read Counts
Young Lean
Young Obese
Aged Lean
Aged Obese
Ercc5
ENSMUSG00000026048
VIP: 2.03
Padj: 0.0047
Gm17552
ENSMUSG00000097233
VIP: 2.09
Padj: 0.0048
Slc2a9
ENSMUSG00000005107
VIP: 2.14
Padj: 0.0049
Gzmc
ENSMUSG00000079186
VIP: 2.07
Padj: 0.0049
Syndig1
ENSMUSG00000074736
VIP: 2.14
Padj: 0.0055

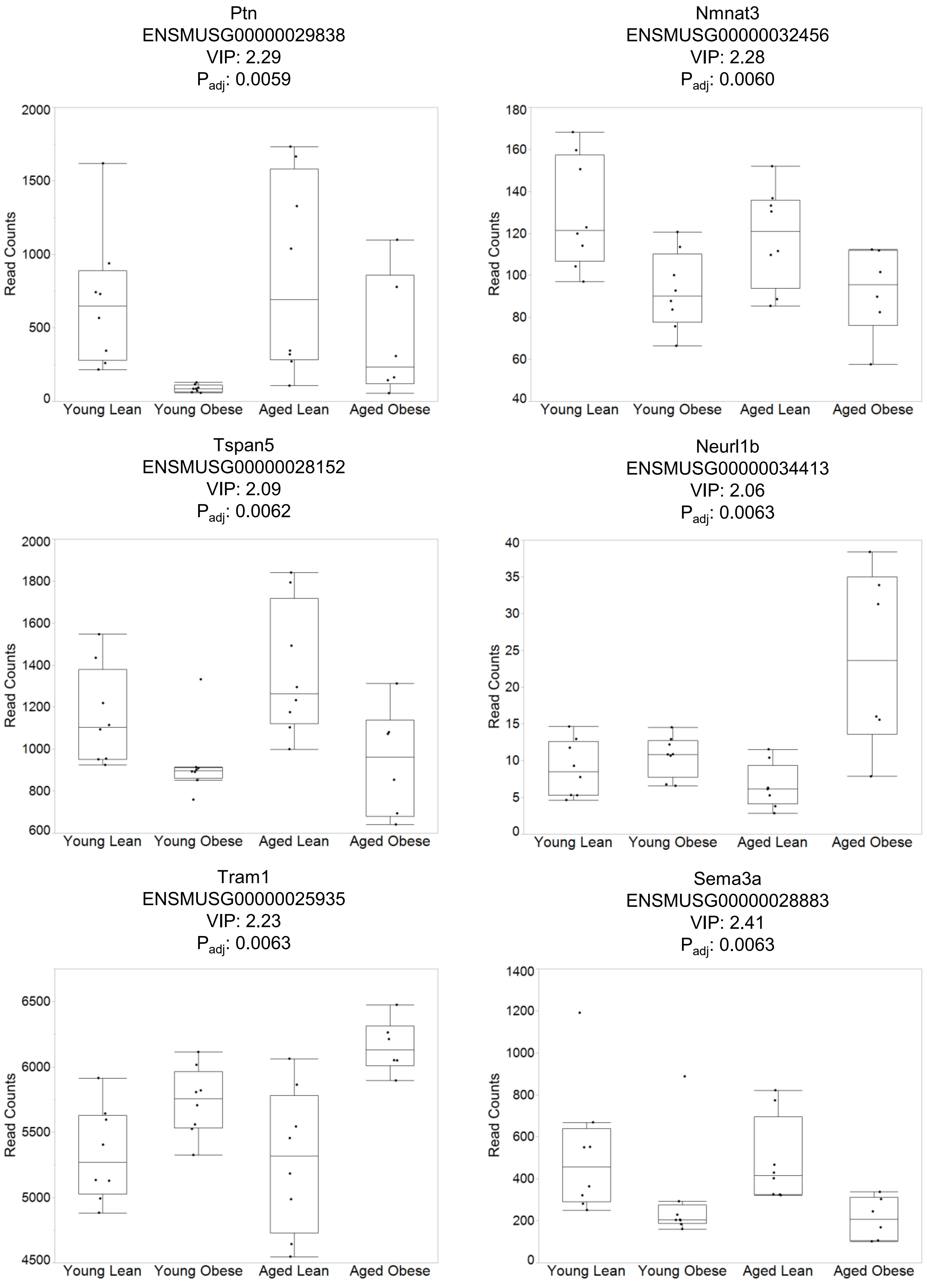
Ptn
ENSMUSG00000029838
VIP: 2.29
$P_{adj}$: 0.0059
Nmnat3
ENSMUSG00000032456
VIP: 2.28
$P_{adj}$: 0.0060
Tspan5
ENSMUSG00000028152
VIP: 2.09
$P_{adj}$: 0.0062
Neurl1b
ENSMUSG00000034413
VIP: 2.06
$P_{adj}$: 0.0063
Tram1
ENSMUSG00000025935
VIP: 2.23
$P_{adj}$: 0.0063
Sema3a
ENSMUSG00000028883
VIP: 2.41
$P_{adj}$: 0.0063
Read Counts
Young Lean
Young Obese
Aged Lean
Aged Obese

Dlk1
ENSMUSG00000040856
VIP: 2.05
$P_{adj}$: 0.0066

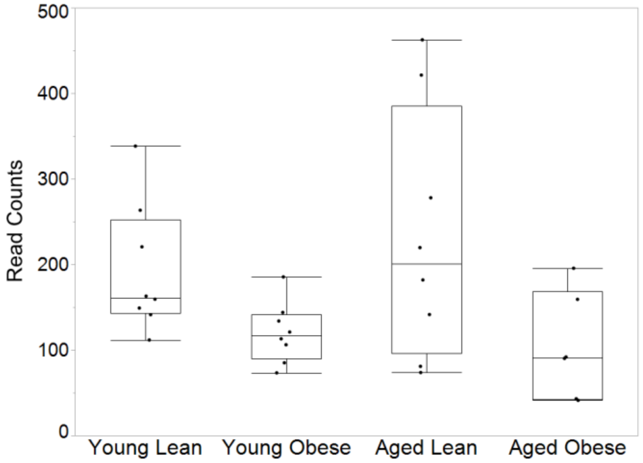


Gmppb
ENSMUSG00000070284
VIP: 2.10
$P_{adj}$: 0.0076

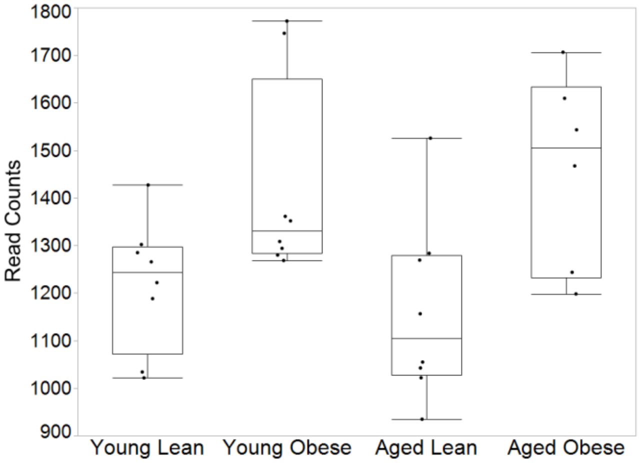


Slc43a1
ENSMUSG00000027075
VIP: 2.03
$P_{adj}$: 0.0081

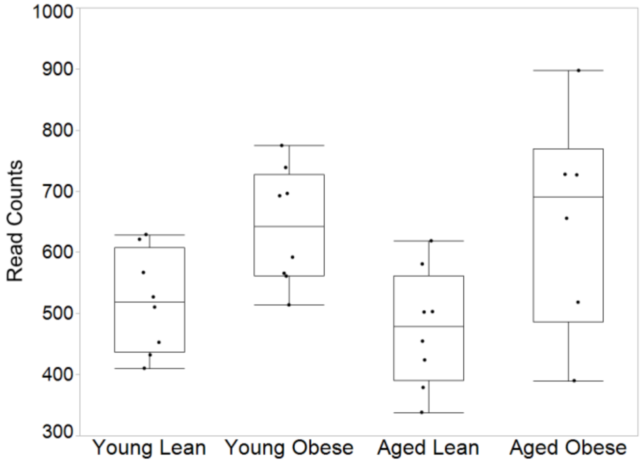


Gm9949
ENSMUSG00000054589
VIP: 2.04
$P_{adj}$: 0.0084

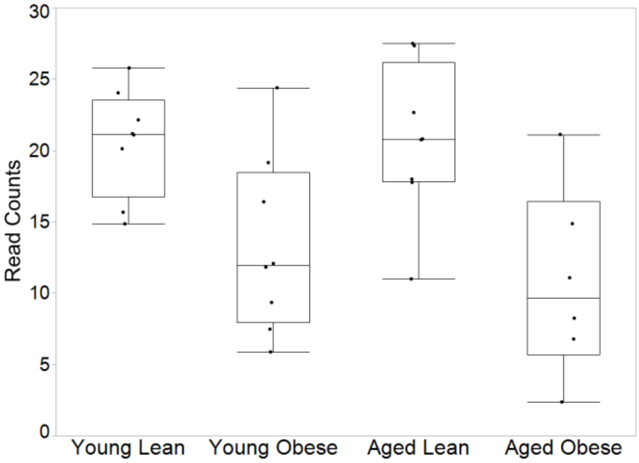


Gsto1
ENSMUSG00000025068
VIP: 2.13
$P_{adj}$: 0.0085

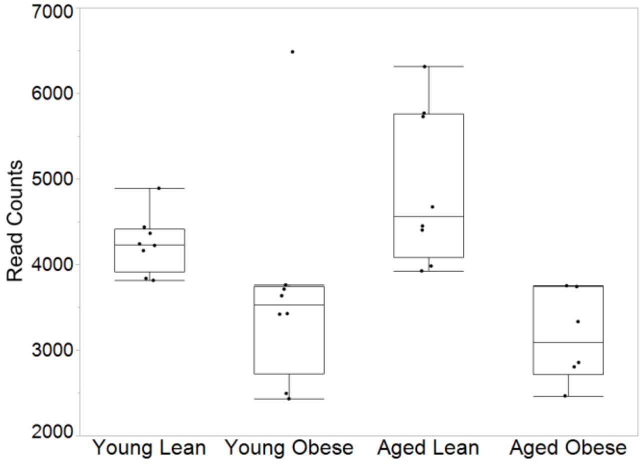


Wnk4
ENSMUSG00000035112
VIP: 2.34
$P_{adj}$: 0.010

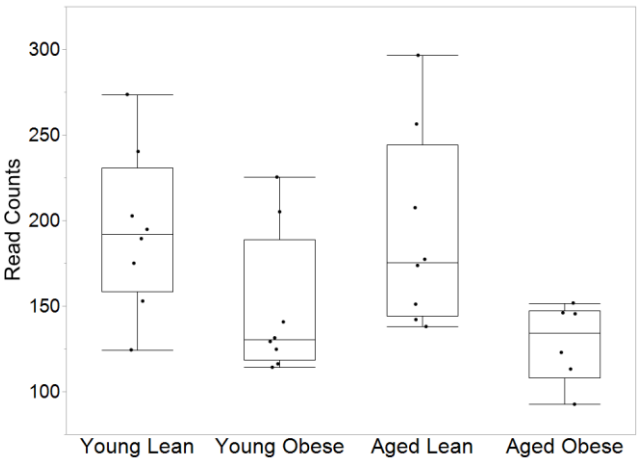

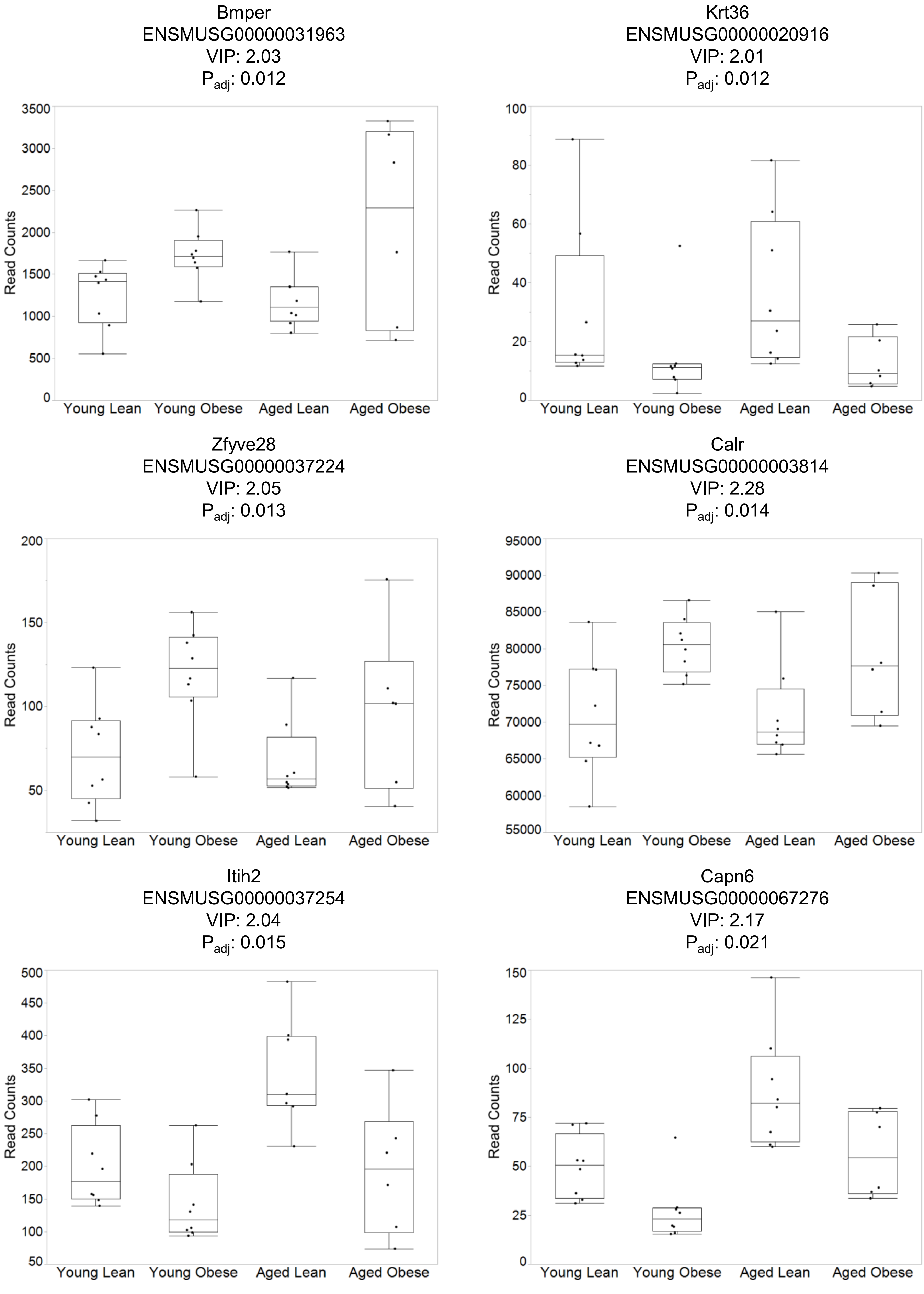
Bmper
ENSMUSG00000031963
VIP: 2.03
$P_{adj}$: 0.012
Read Counts
0
500
1000
1500
2000
2500
3000
3500
Young Lean
Young Obese
Aged Lean
Aged Obese
Krt36
ENSMUSG00000020916
VIP: 2.01
$P_{adj}$: 0.012
Read Counts
0
20
40
60
80
100
Young Lean
Young Obese
Aged Lean
Aged Obese
Zfyve28
ENSMUSG00000037224
VIP: 2.05
$P_{adj}$: 0.013
Read Counts
50
100
150
200
Young Lean
Young Obese
Aged Lean
Aged Obese
Calr
ENSMUSG00000003814
VIP: 2.28
$P_{adj}$: 0.014
Read Counts
55000
60000
65000
70000
75000
80000
85000
90000
95000
Young Lean
Young Obese
Aged Lean
Aged Obese
Itih2
ENSMUSG00000037254
VIP: 2.04
$P_{adj}$: 0.015
Read Counts
50
100
150
200
250
300
350
400
450
500
Young Lean
Young Obese
Aged Lean
Aged Obese
Capn6
ENSMUSG00000067276
VIP: 2.17
$P_{adj}$: 0.021
Read Counts
0
25
50
75
100
125
150
Young Lean
Young Obese
Aged Lean
Aged Obese

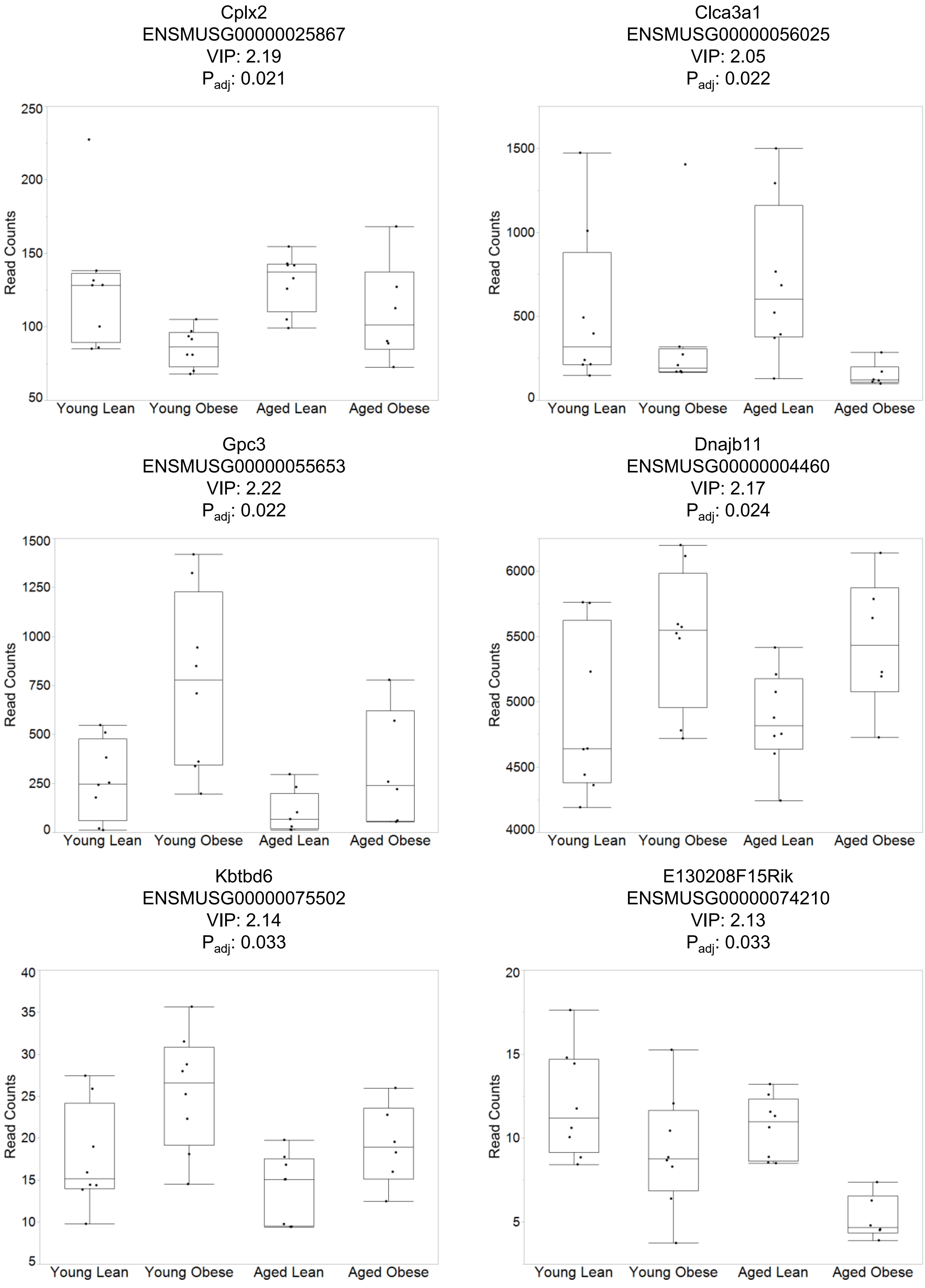
Cplx2
ENSMUSG00000025867
VIP: 2.19
P adj: 0.021
Clca3a1
ENSMUSG00000056025
VIP: 2.05
P adj: 0.022
Gpc3
ENSMUSG00000055653
VIP: 2.22
P adj: 0.022
Dnajb11
ENSMUSG00000004460
VIP: 2.17
P adj: 0.024
Kbtbd6
ENSMUSG00000075502
VIP: 2.14
P adj: 0.033
E130208F15Rik
ENSMUSG00000074210
VIP: 2.13
P adj: 0.033
Read Counts
Young Lean
Young Obese
Aged Lean
Aged Obese

Pdia4
ENSMUSG00000025823
VIP: 2.23
$P_{adj}$: 0.042

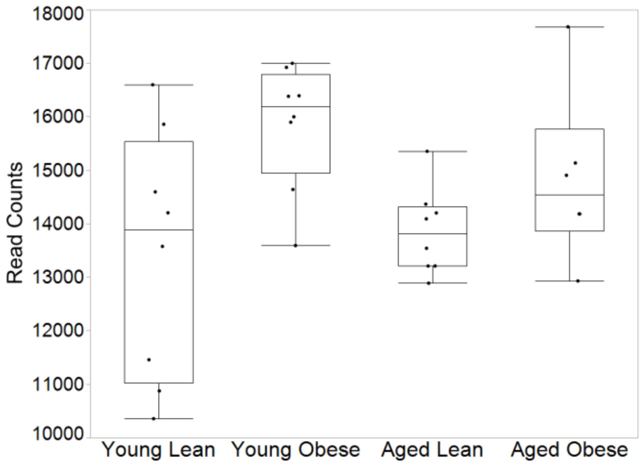


Gm10658
ENSMUSG00000074284
VIP: 2.27
$P_{adj}$: 0.042

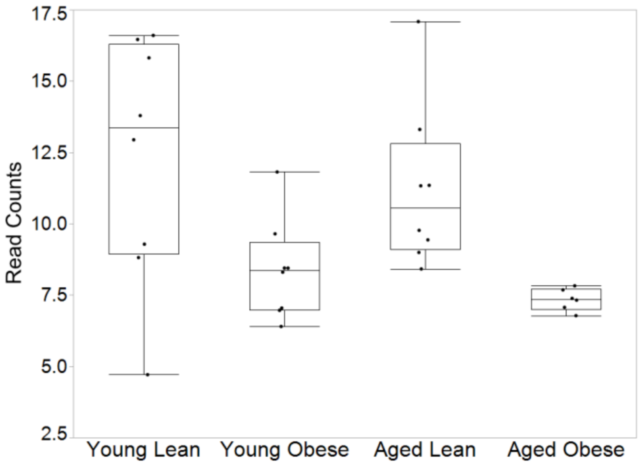


Oxa1l
ENSMUSG00000000959
VIP: 2.02
$P_{adj}$: 0.043

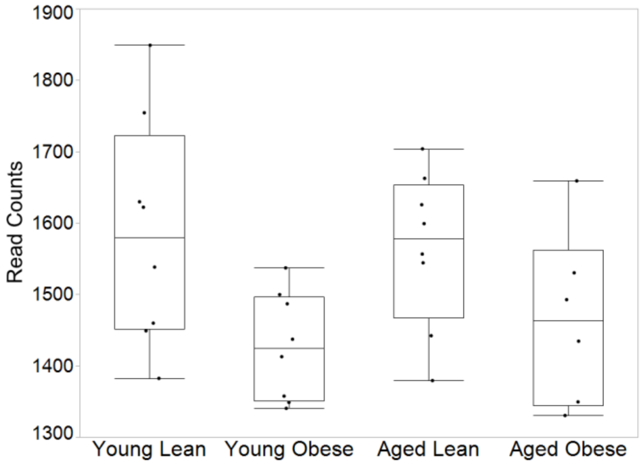
