## Supplemental Figure 7 for "Aging and Obesity Prime the Methylome and Transcriptome of Adipose Stem Cells for Disease and Dysfunction"

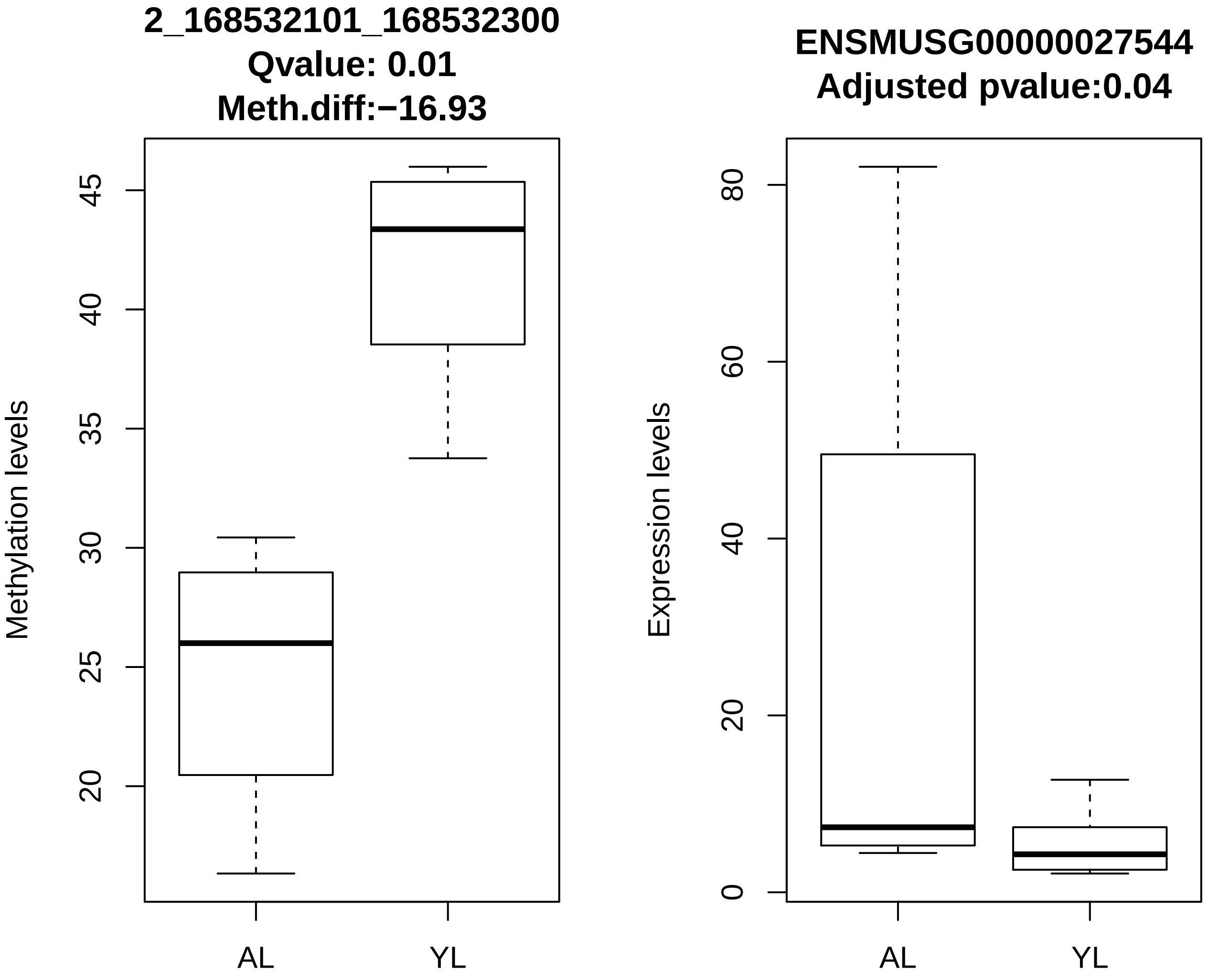
2_168532101_168532300
Qvalue: 0.01
Meth.diff:−16.93
Methylation levels
20
25
30
35
40
45
AL
YL
ENSMUSG00000027544
Adjusted pvalue:0.04
Expression levels
0
20
40
60
80
AL
YL

Nfatc2

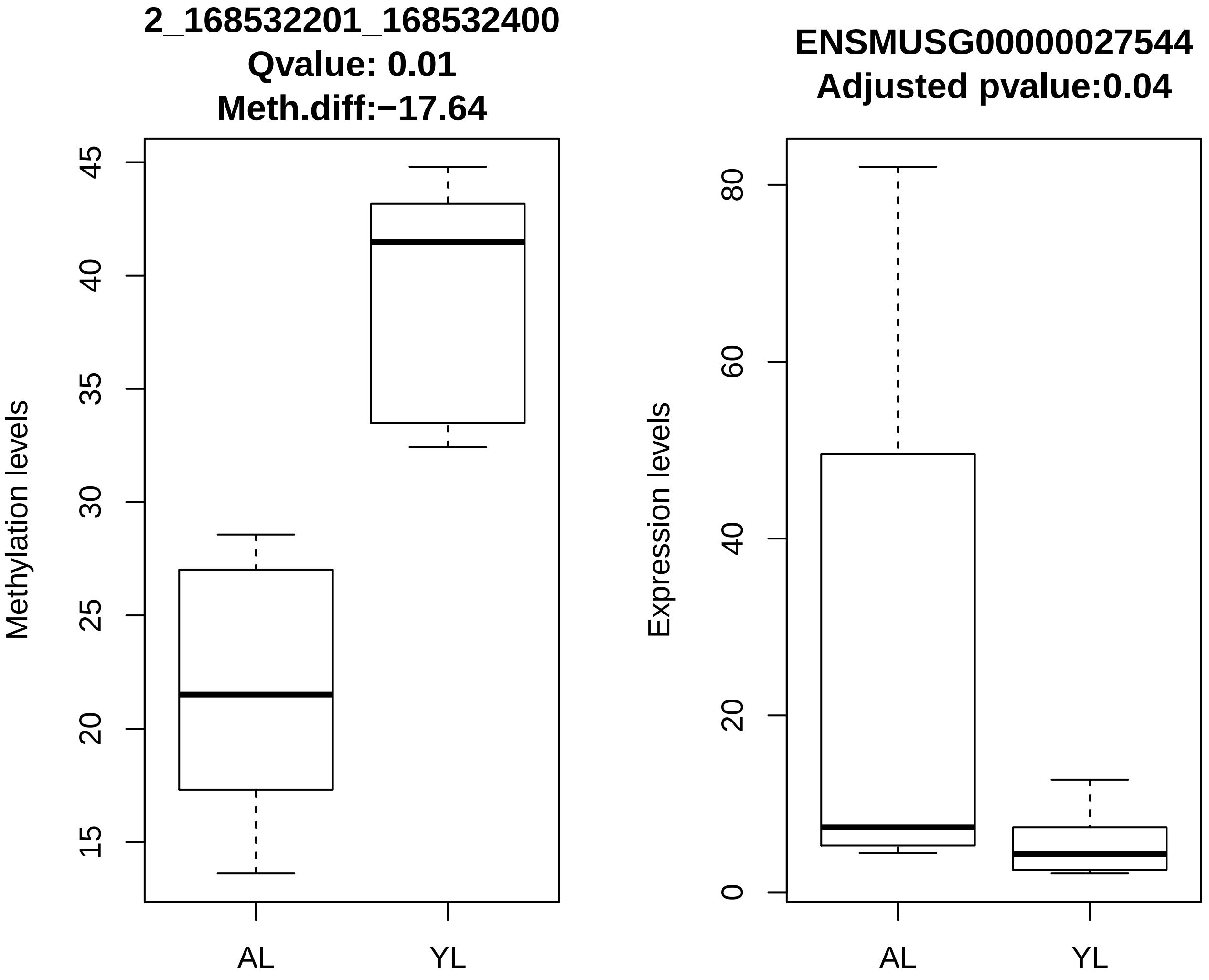
2_168532201_168532400
Qvalue: 0.01
Meth.diff:−17.64
Methylation levels
45
40
35
30
25
20
15
AL
YL
ENSMUSG00000027544
Adjusted pvalue:0.04
Expression levels
80
60
40
20
0
AL
YL

Nfatc2

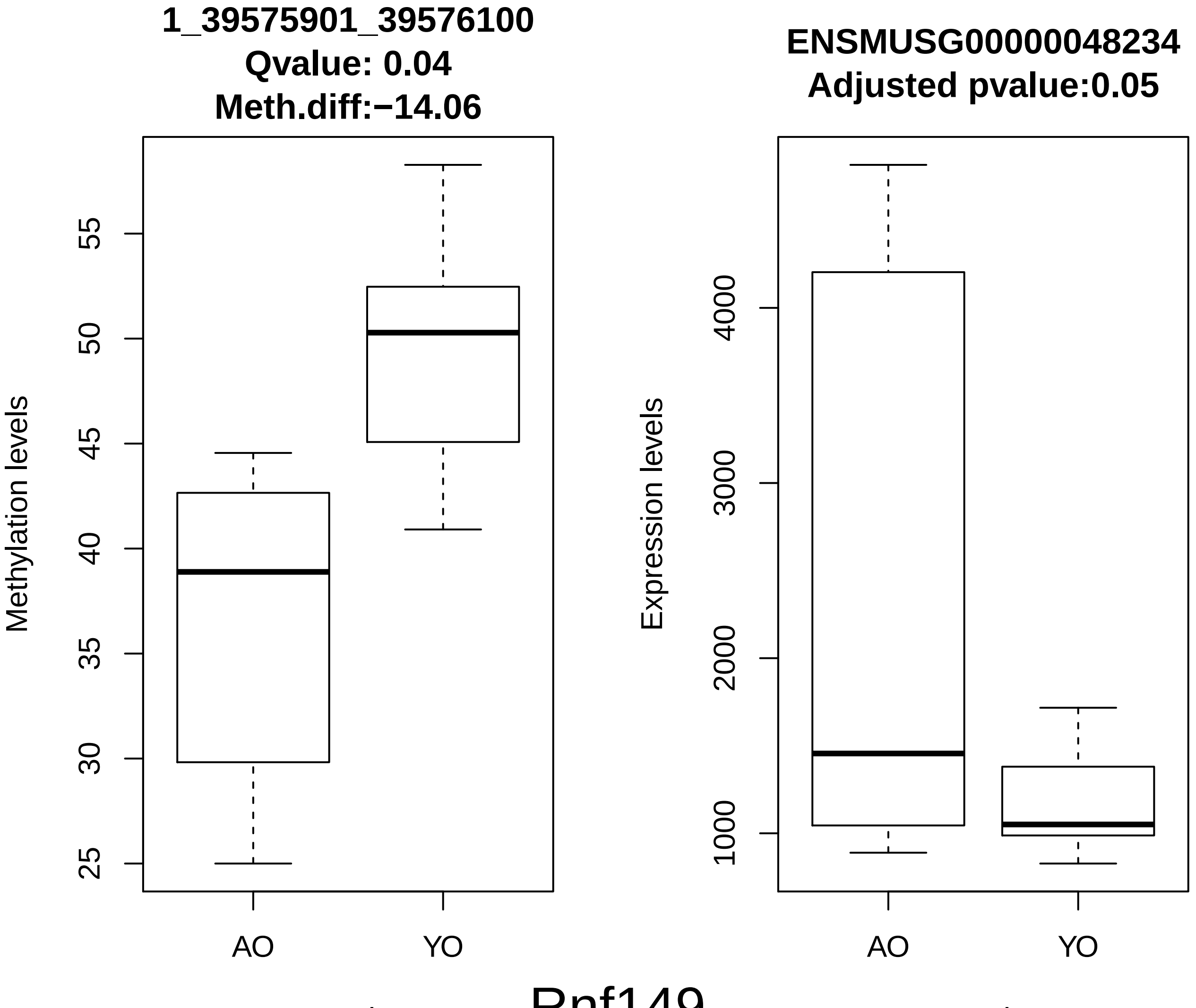
1_39575901_39576100
Qvalue: 0.04
Meth.diff:−14.06
Methylation levels
25
30
35
40
45
50
55
AO
YO
Samples
ENSMUSG00000048234
Adjusted pvalue:0.05
Expression levels
1000
2000
3000
4000
AO
YO
Samples
Rnf149

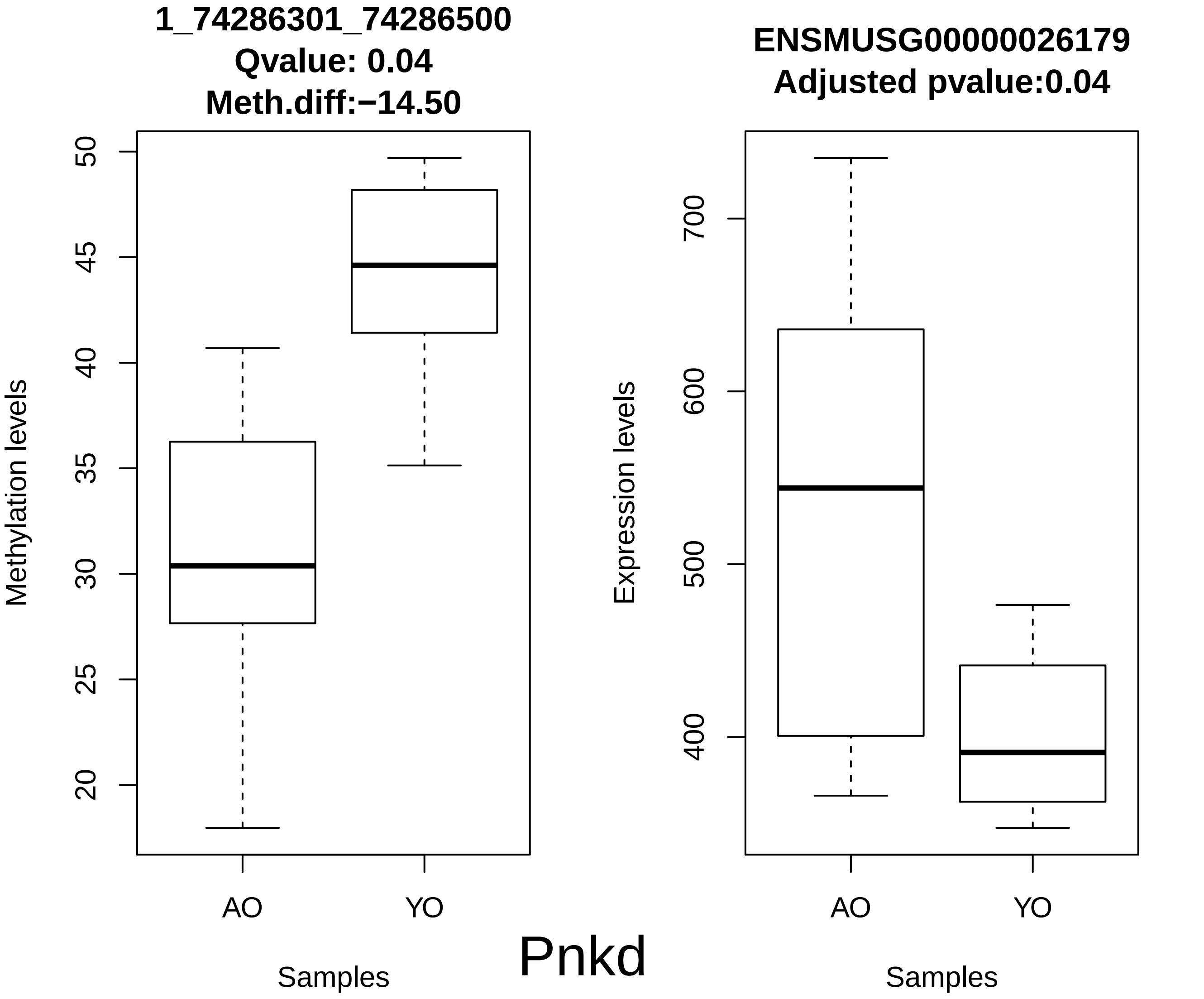
1_74286301_74286500
Qvalue: 0.04
Meth.diff:−14.50
Methylation levels
50
45
40
35
30
25
20
AO
YO
Samples
ENSMUSG00000026179
Adjusted pvalue:0.04
Expression levels
700
600
500
400
AO
YO
Samples
Pnkd

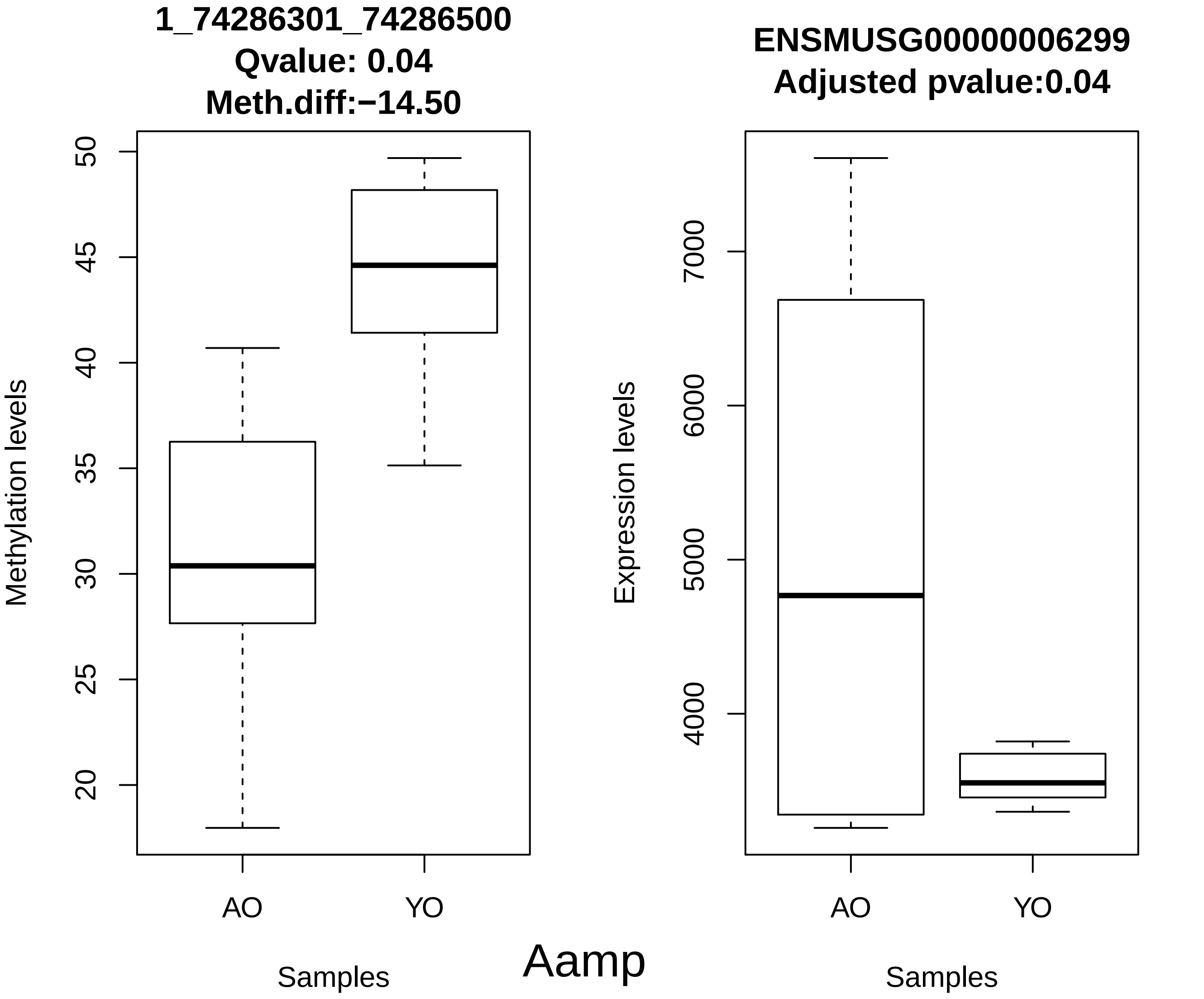
1_74286301_74286500
Qvalue: 0.04
Meth.diff:−14.50
Methylation levels
50
45
40
35
30
25
20
AO
YO
Samples
ENSMUSG00000006299
Adjusted pvalue:0.04
Expression levels
7000
6000
5000
4000
AO
YO
Samples
Aamp

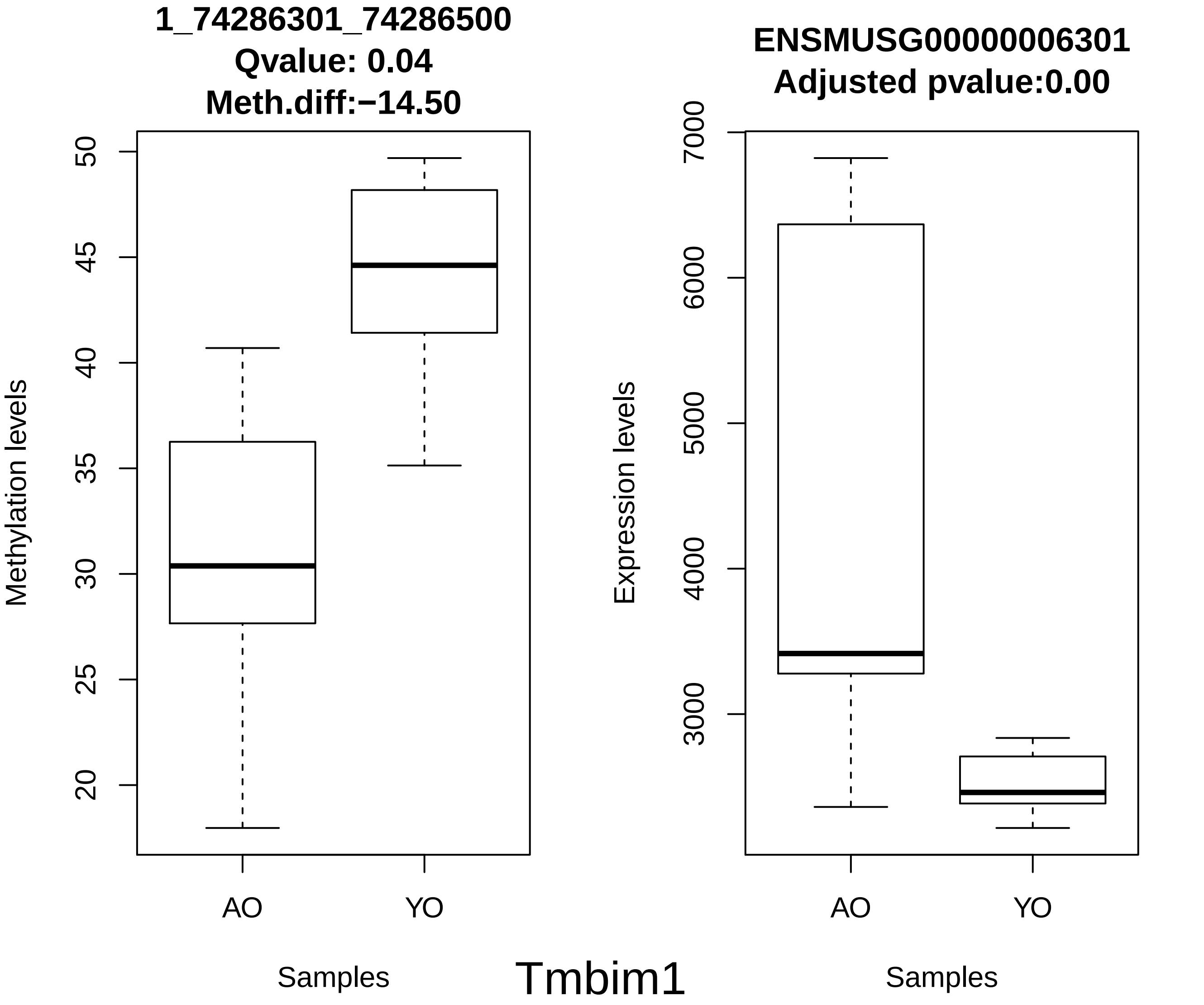
1_74286301_74286500
Qvalue: 0.04
Meth.diff:−14.50
Methylation levels
50
45
40
35
30
25
20
AO
YO
Samples
ENSMUSG00000006301
Adjusted pvalue:0.00
Expression levels
7000
6000
5000
4000
3000
AO
YO
Samples
Tmbim1

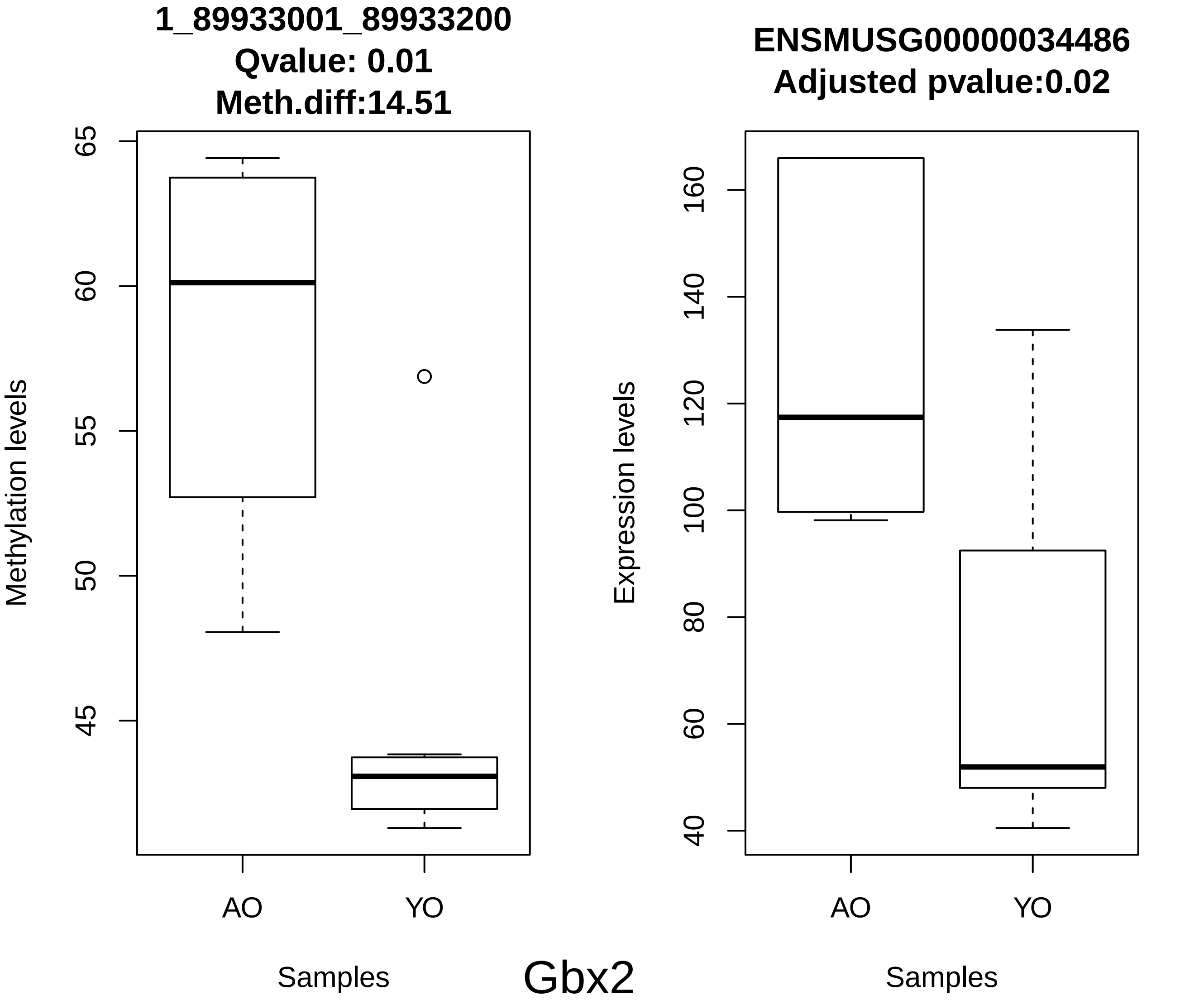
1_89933001_89933200
Qvalue: 0.01
Meth.diff:14.51
Methylation levels
65
60
55
50
45
AO
YO
Samples
ENSMUSG00000034486
Adjusted pvalue:0.02
Expression levels
160
140
120
100
80
60
40
AO
YO
Samples
Gbx2

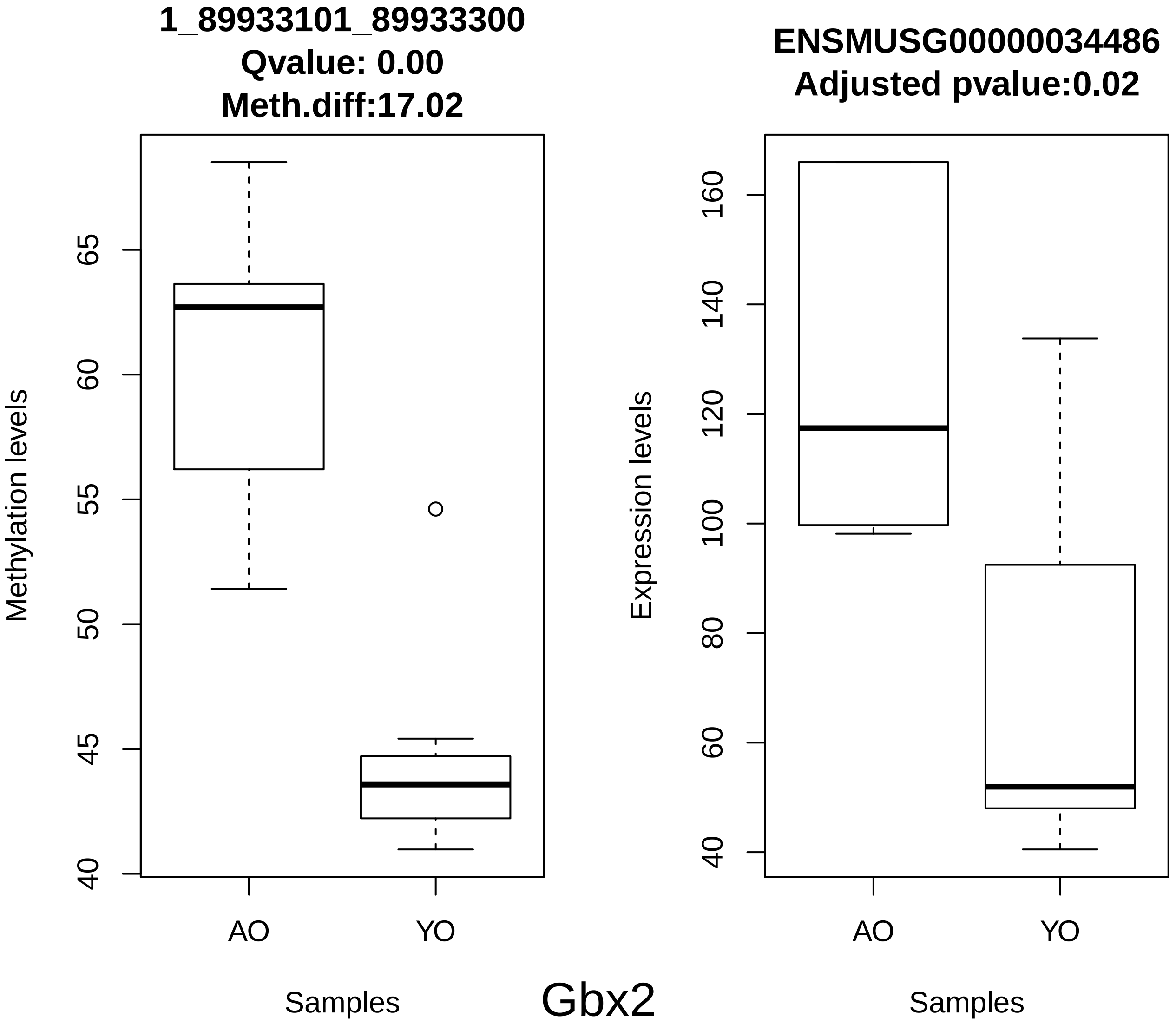
1_89933101_89933300
Qvalue: 0.00
Meth.diff:17.02
Methylation levels
65
60
55
50
45
40
AO
YO
Samples
ENSMUSG00000034486
Adjusted pvalue:0.02
Expression levels
160
140
120
100
80
60
40
AO
YO
Samples

Gbx2

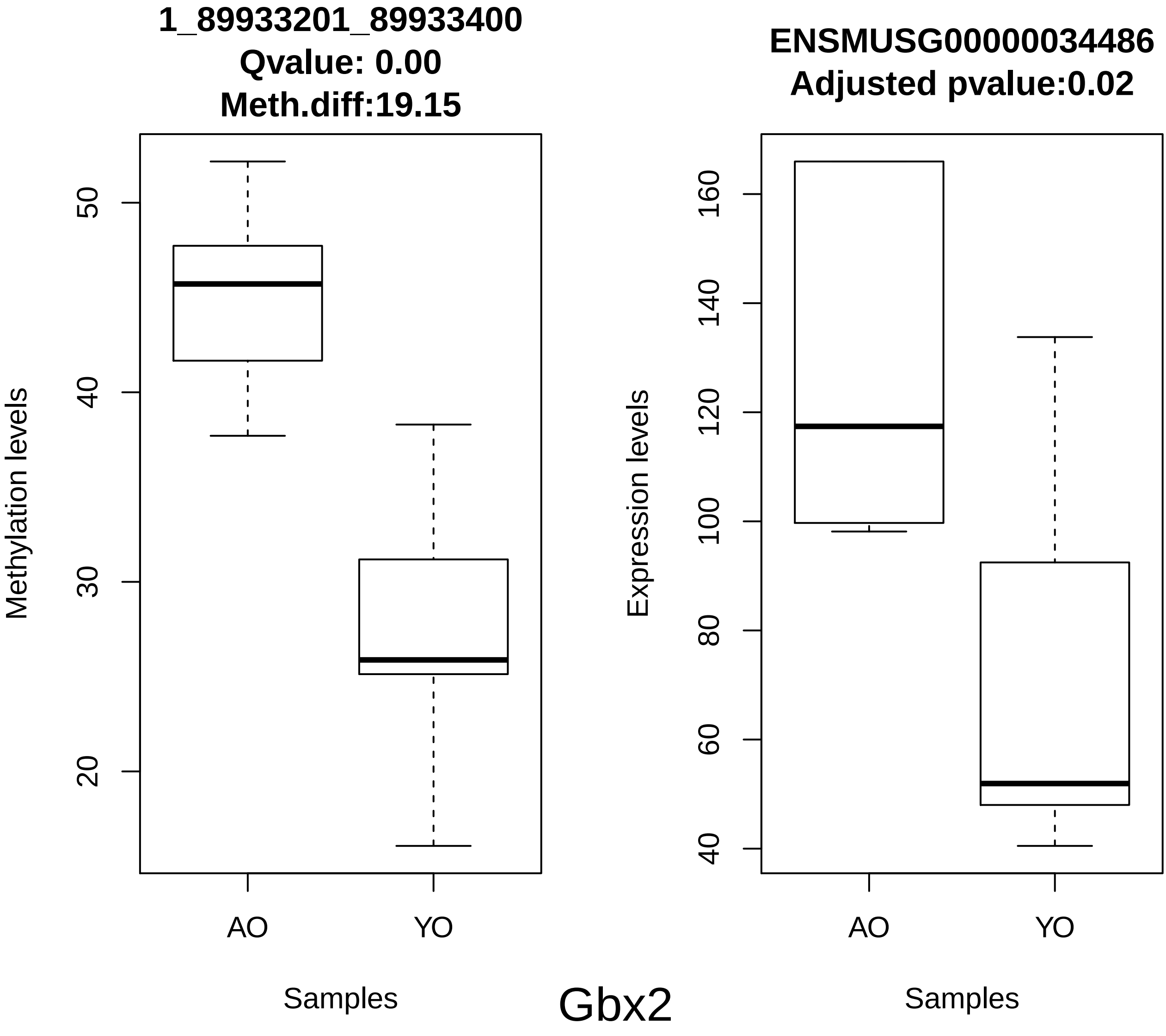
1_89933201_89933400
Qvalue: 0.00
Meth.diff:19.15
Methylation levels
50
40
30
20
AO
YO
Samples
ENSMUSG00000034486
Adjusted pvalue:0.02
Expression levels
160
140
120
100
80
60
40
AO
YO
Samples
Gbx2

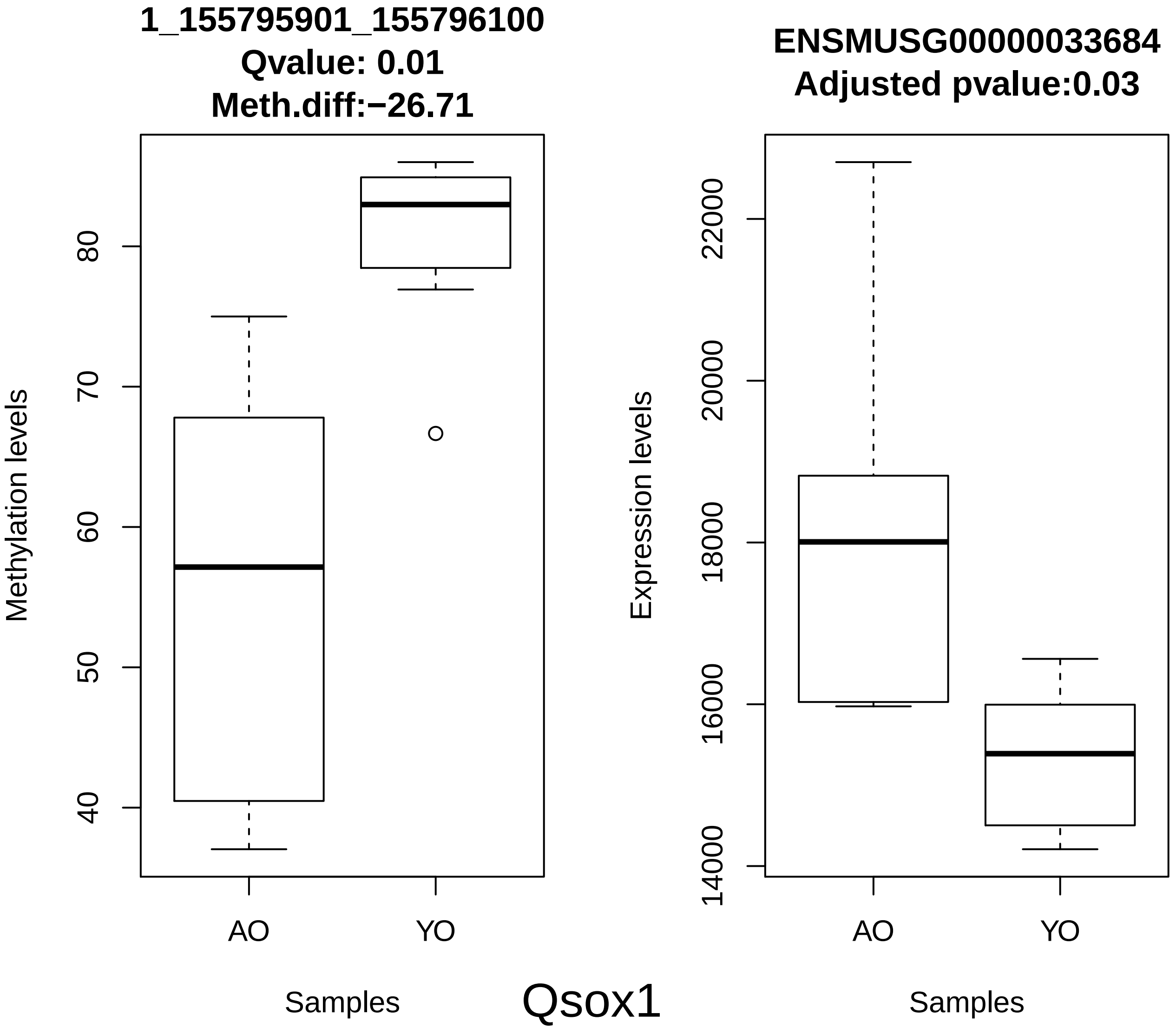
1_155795901_155796100
Qvalue: 0.01
Meth.diff:−26.71
Methylation levels
40
50
60
70
80
AO
YO
Samples
ENSMUSG00000033684
Adjusted pvalue:0.03
Expression levels
14000
16000
18000
20000
22000
AO
YO
Samples

Qsox1

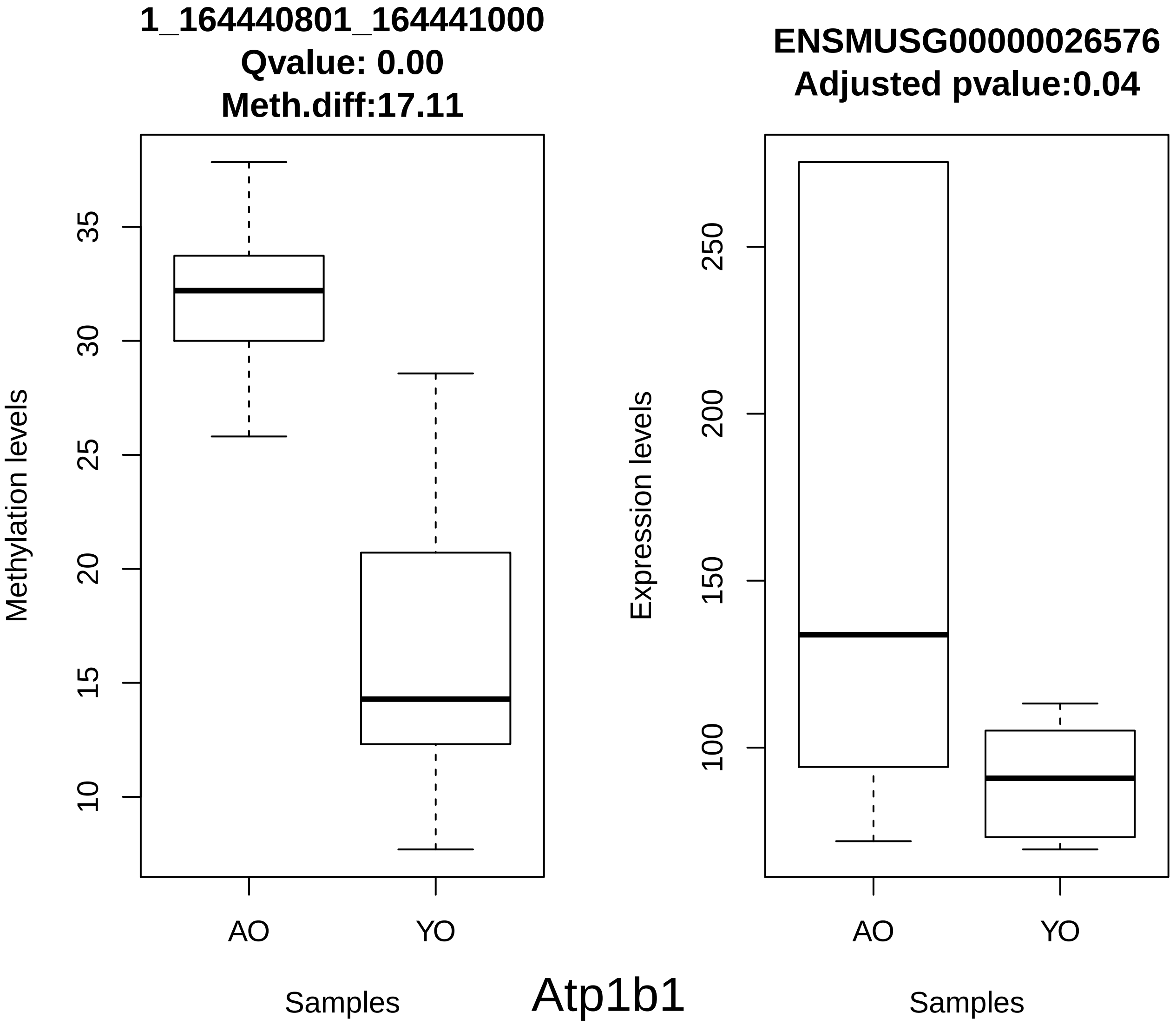
1_164440801_164441000
Qvalue: 0.00
Meth.diff:17.11
Methylation levels
35
30
25
20
15
10
AO
YO
Samples
ENSMUSG00000026576
Adjusted pvalue:0.04
Expression levels
250
200
150
100
AO
YO
Samples
Atp1b1

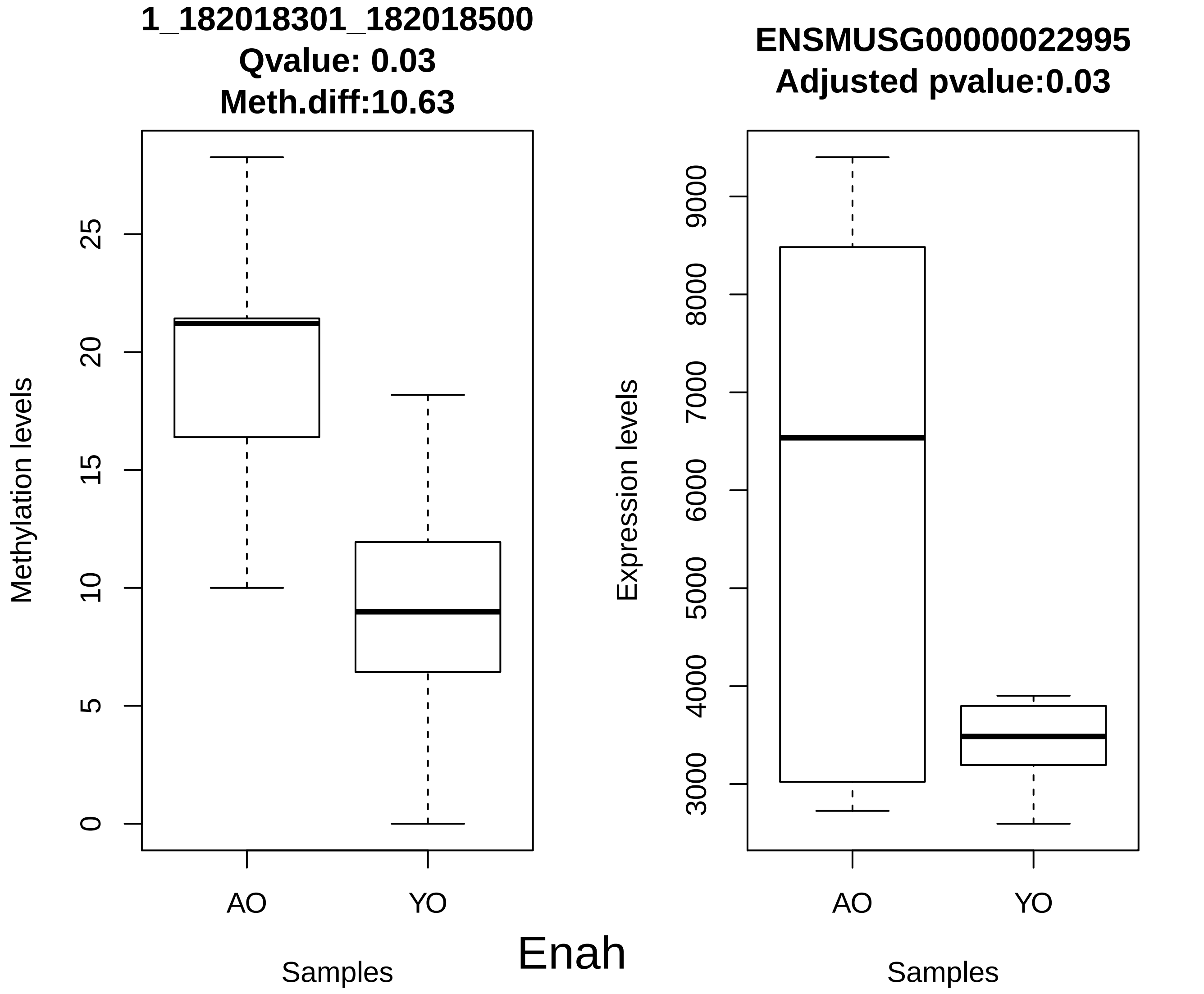
1_182018301_182018500
Qvalue: 0.03
Meth.diff:10.63
Methylation levels
0
5
10
15
20
25
AO
YO
Samples
ENSMUSG00000022995
Adjusted pvalue:0.03
Expression levels
3000
4000
5000
6000
7000
8000
9000
AO
YO
Samples
Enah

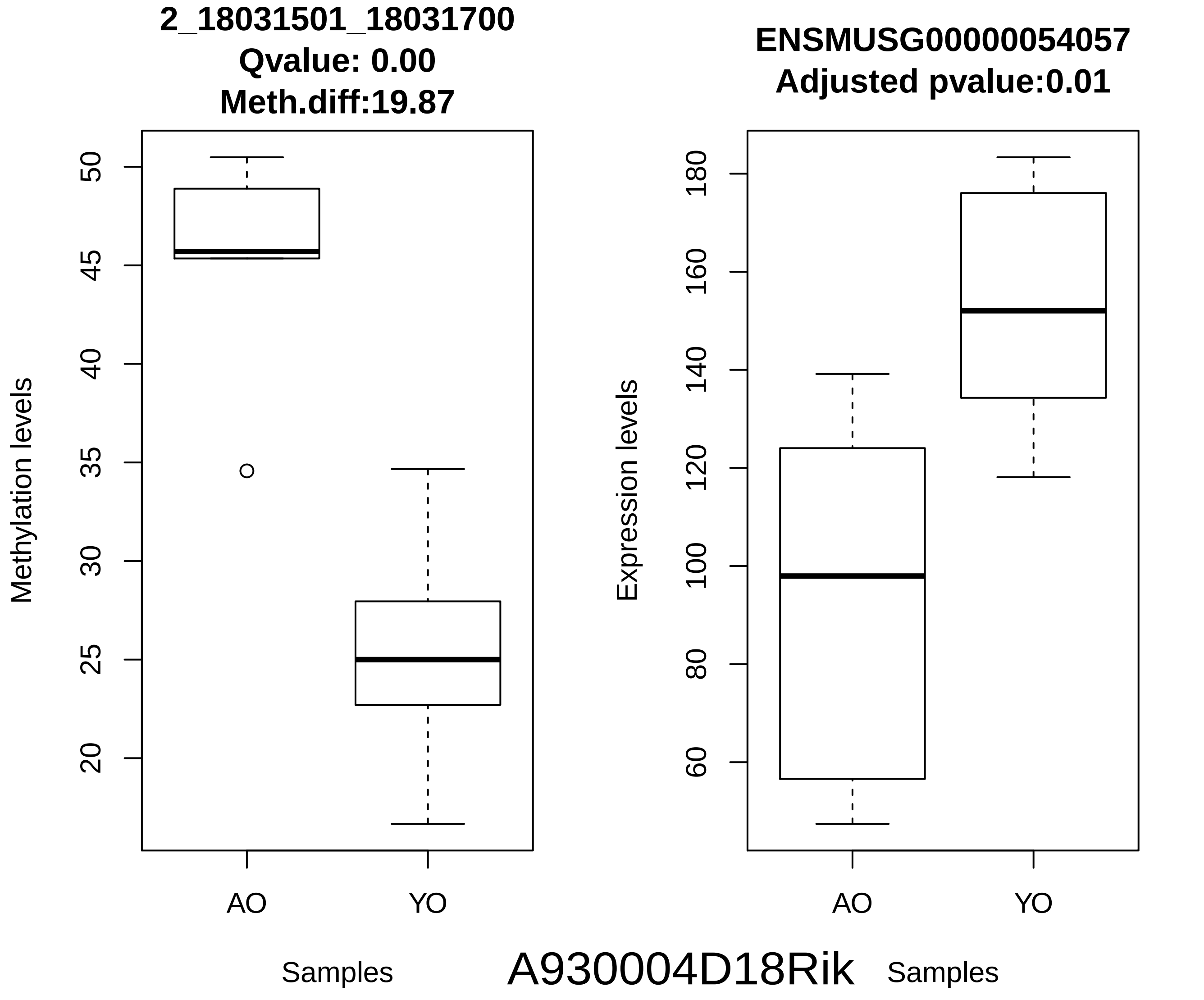

2_18031501_18031700
Qvalue: 0.00
Meth.diff:19.87
Methylation levels
50
45
40
35
30
25
20
AO
YO
Samples
ENSMUSG00000054057
Adjusted pvalue:0.01
Expression levels
180
160
140
120
100
80
60
AO
YO
Samples
A930004D18Rik

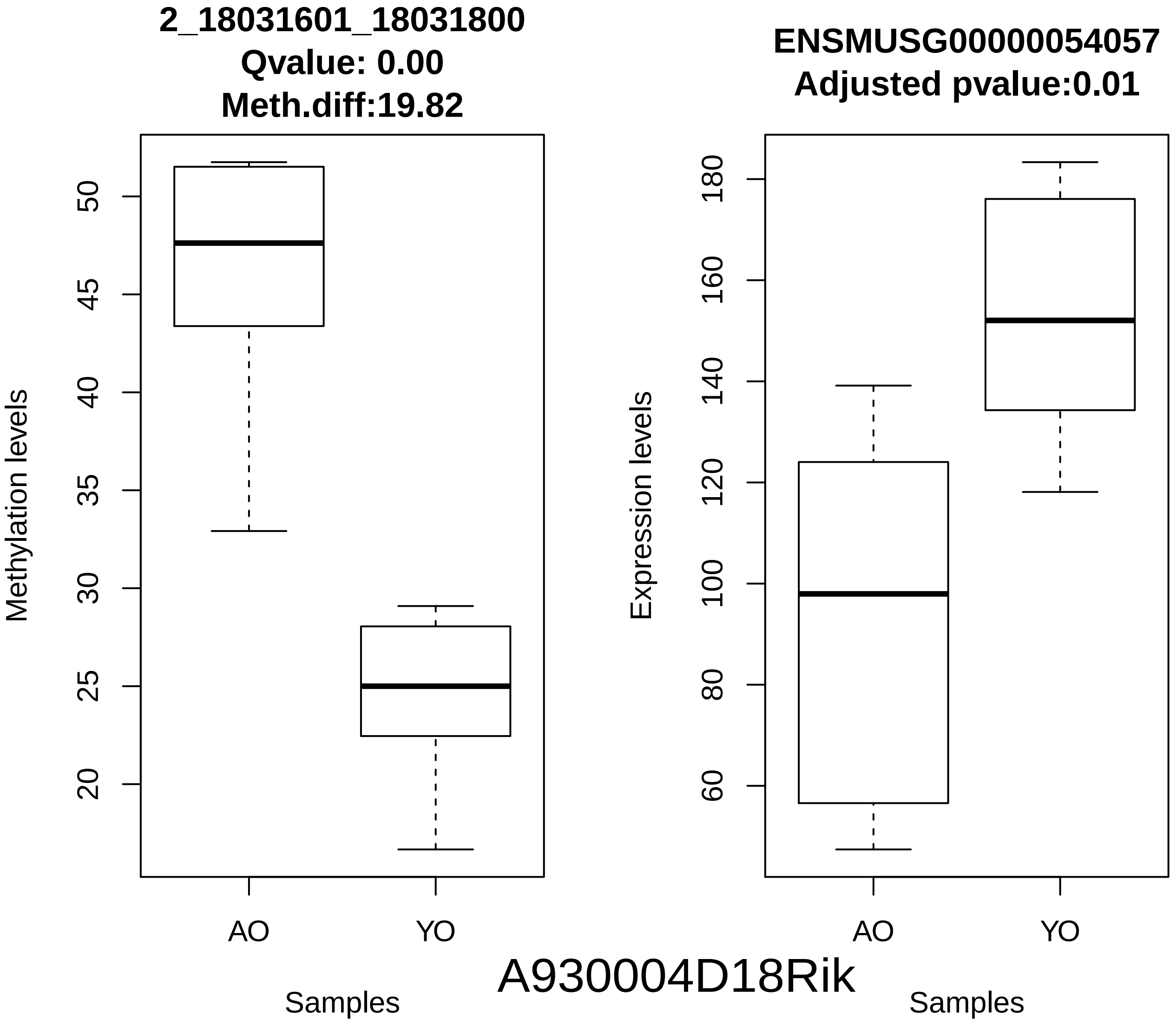
2_18031601_18031800
Qvalue: 0.00
Meth.diff:19.82
Methylation levels
50
45
40
35
30
25
20
AO
YO
Samples
ENSMUSG00000054057
Adjusted pvalue:0.01
Expression levels
180
160
140
120
100
80
60
AO
YO
Samples
A930004D18Rik

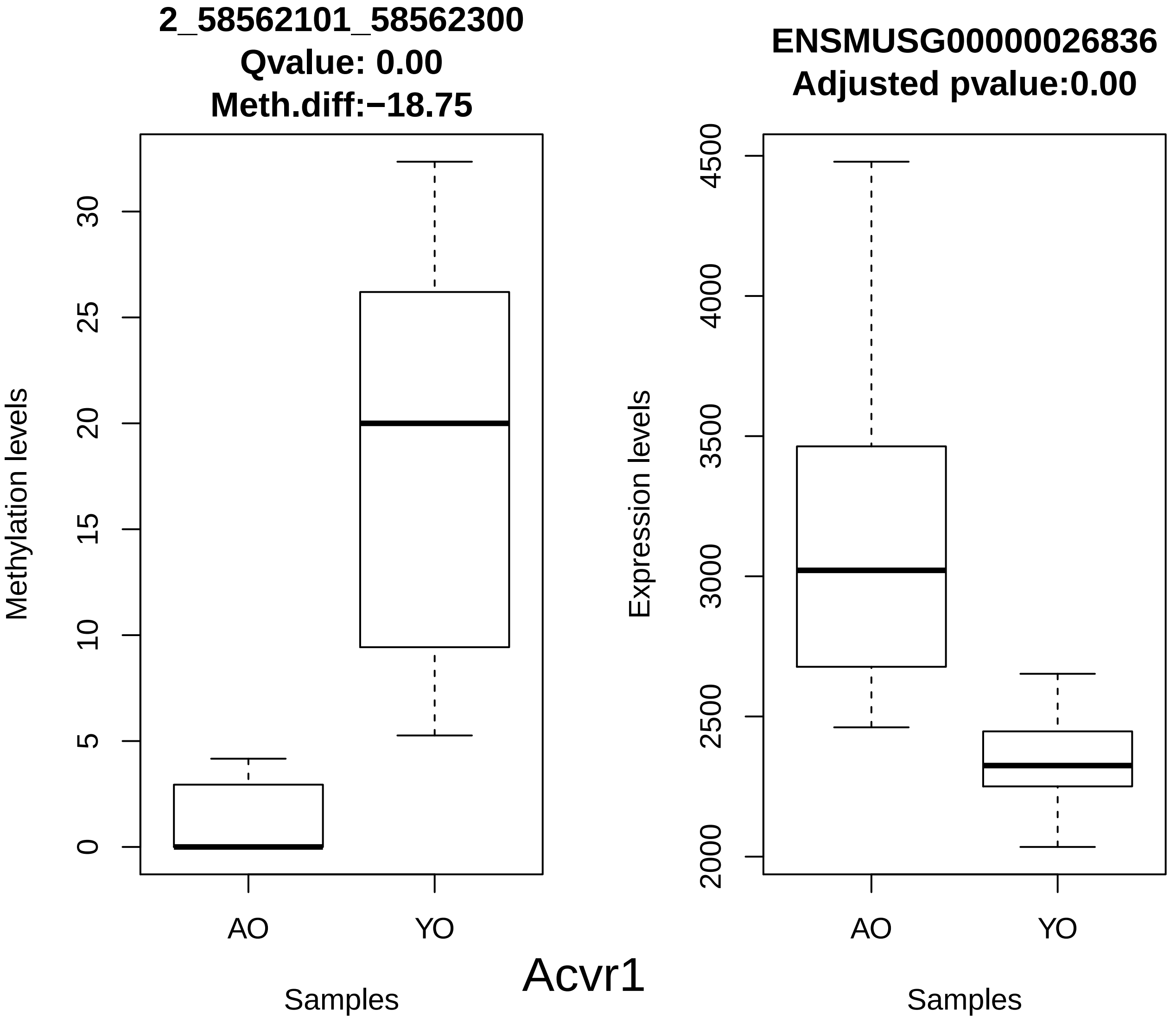
2_58562101_58562300
Qvalue: 0.00
Meth.diff:−18.75
Methylation levels
0
5
10
15
20
25
30
AO
YO
Samples
ENSMUSG00000026836
Adjusted pvalue:0.00
Expression levels
2000
2500
3000
3500
4000
4500
AO
YO
Samples
Acvr1

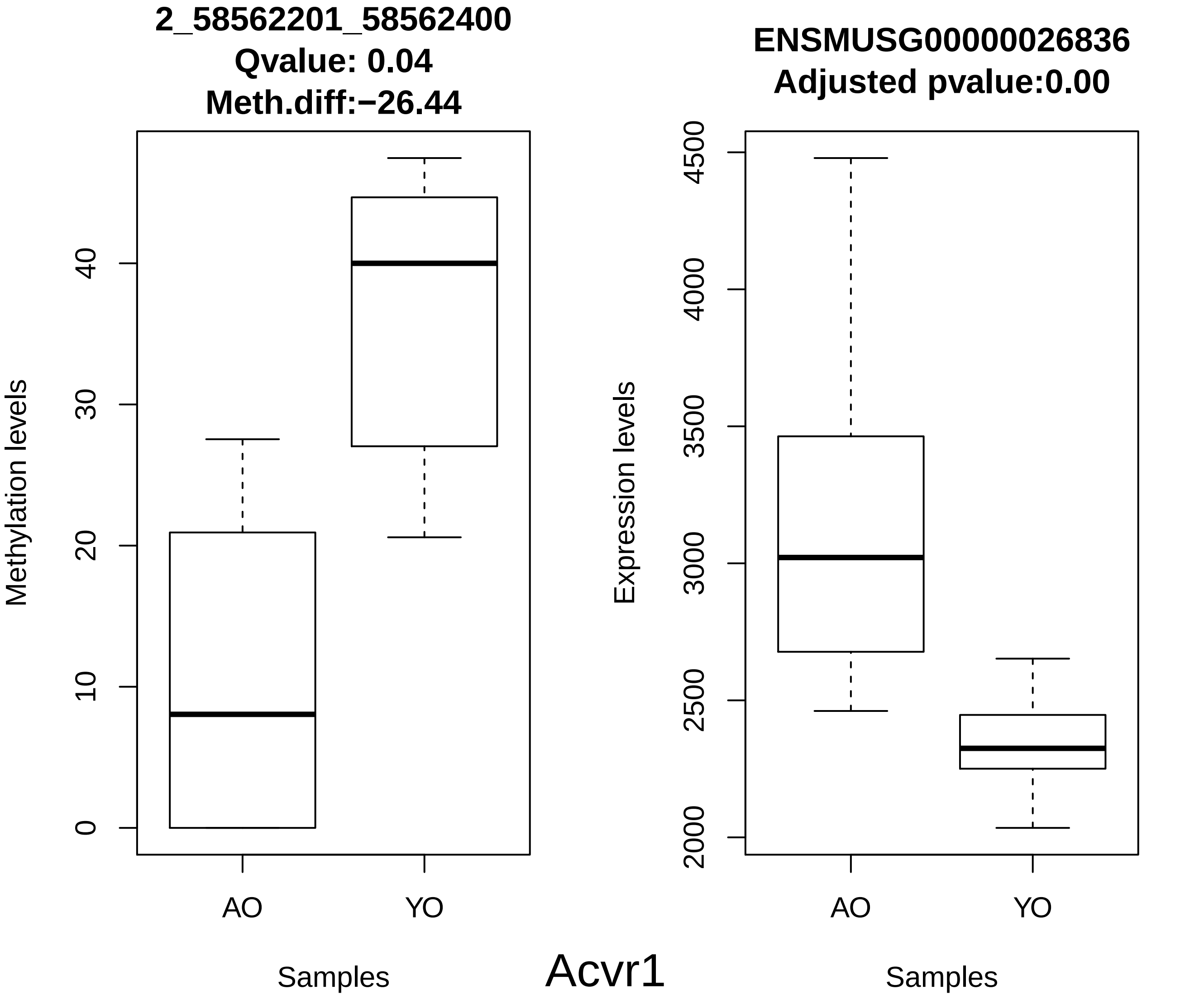
2_58562201_58562400
Qvalue: 0.04
Meth.diff:−26.44
Methylation levels
0
10
20
30
40
AO
YO
Samples
ENSMUSG00000026836
Adjusted pvalue:0.00
Expression levels
2000
2500
3000
3500
4000
4500
AO
YO
Samples

Acvr1

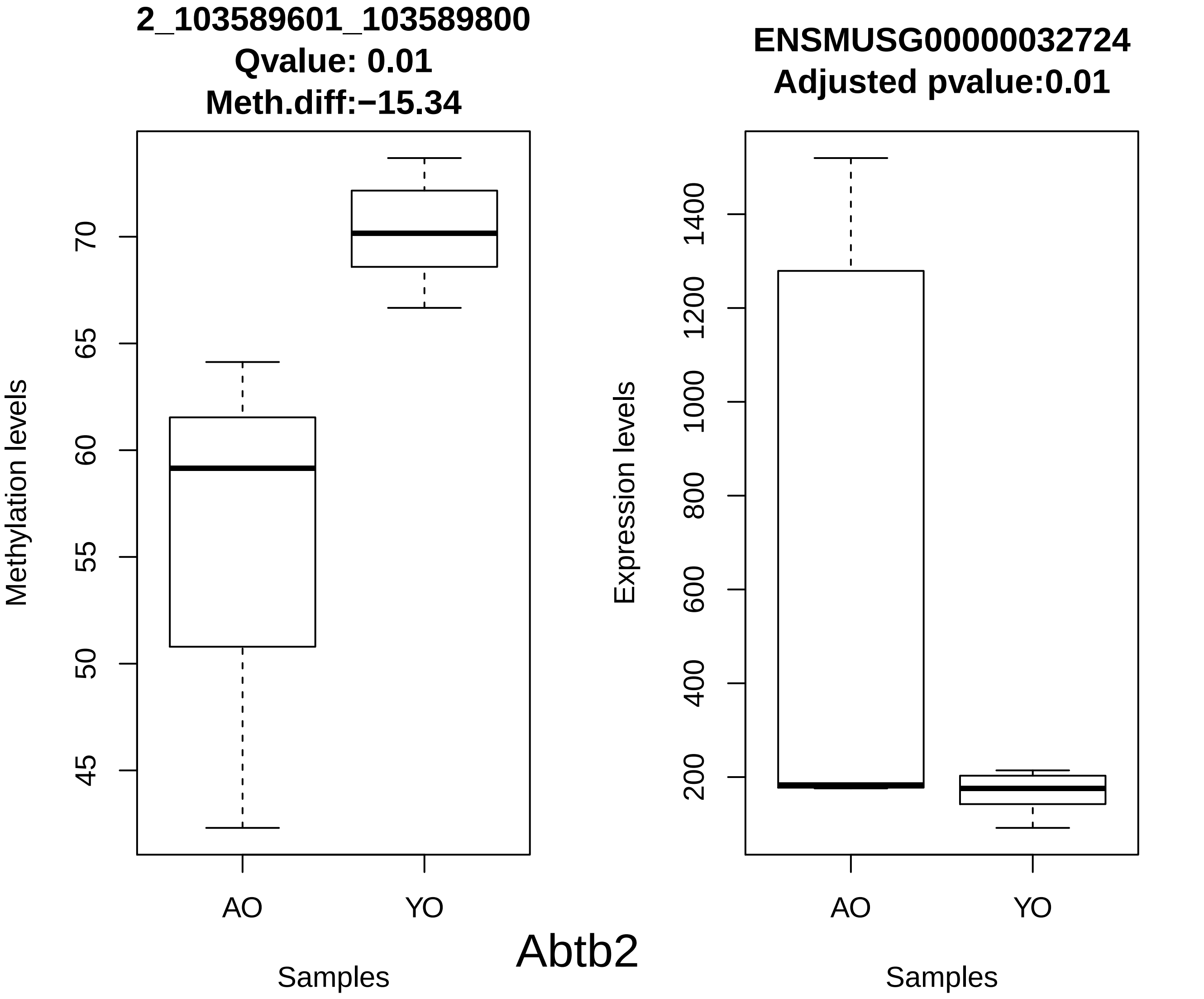
2_103589601_103589800
Qvalue: 0.01
Meth.diff:−15.34
Methylation levels
45
50
55
60
65
70
AO
YO
Samples
ENSMUSG00000032724
Adjusted pvalue:0.01
Expression levels
200
400
600
800
1000
1200
1400
AO
YO
Samples
Abtb2

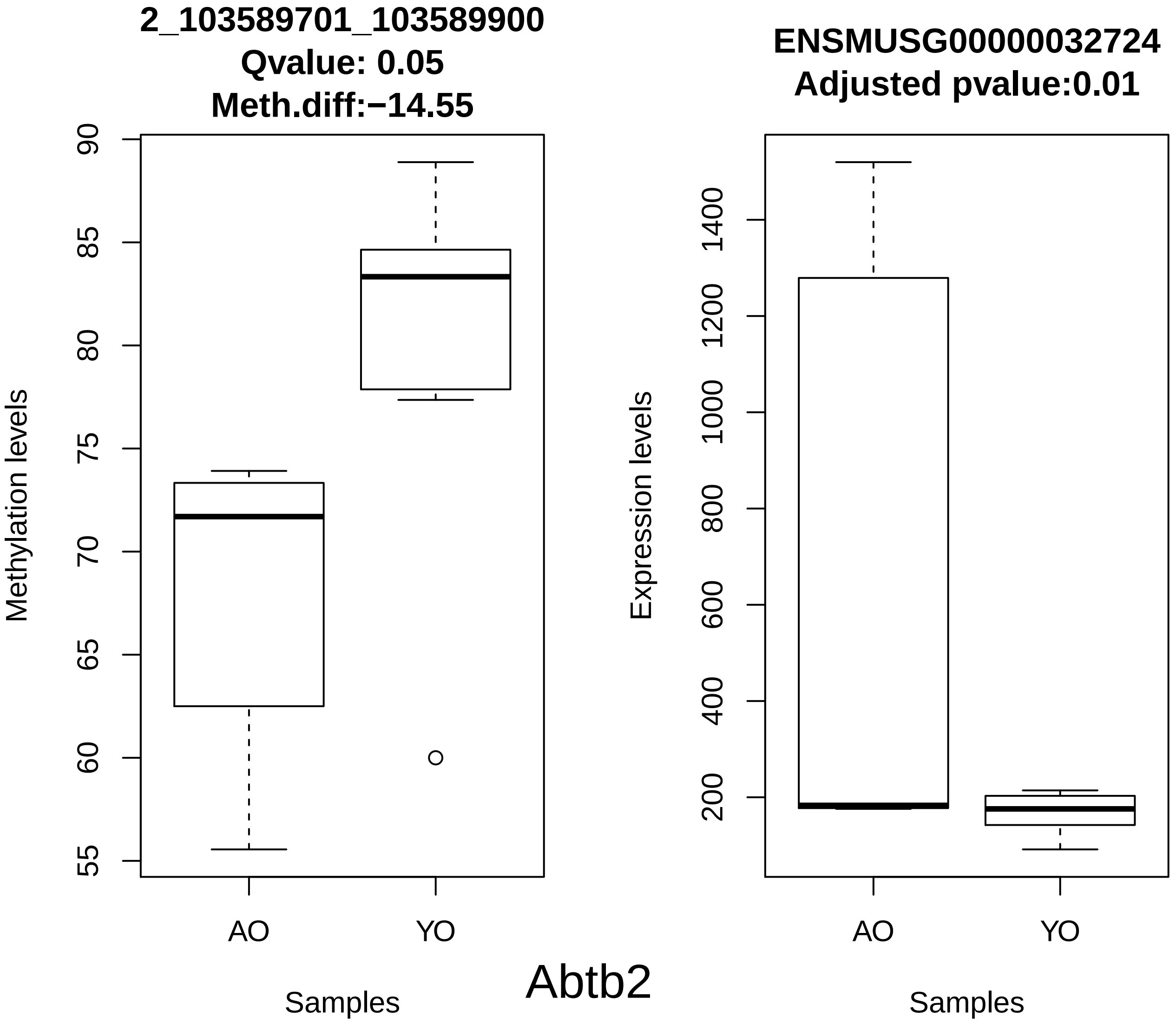
2_103589701_103589900
Qvalue: 0.05
Meth.diff:−14.55
Methylation levels
90
85
80
75
70
65
60
55
AO
YO
Samples
ENSMUSG00000032724
Adjusted pvalue:0.01
Expression levels
1400
1200
1000
800
600
400
200
AO
YO
Samples
Abtb2

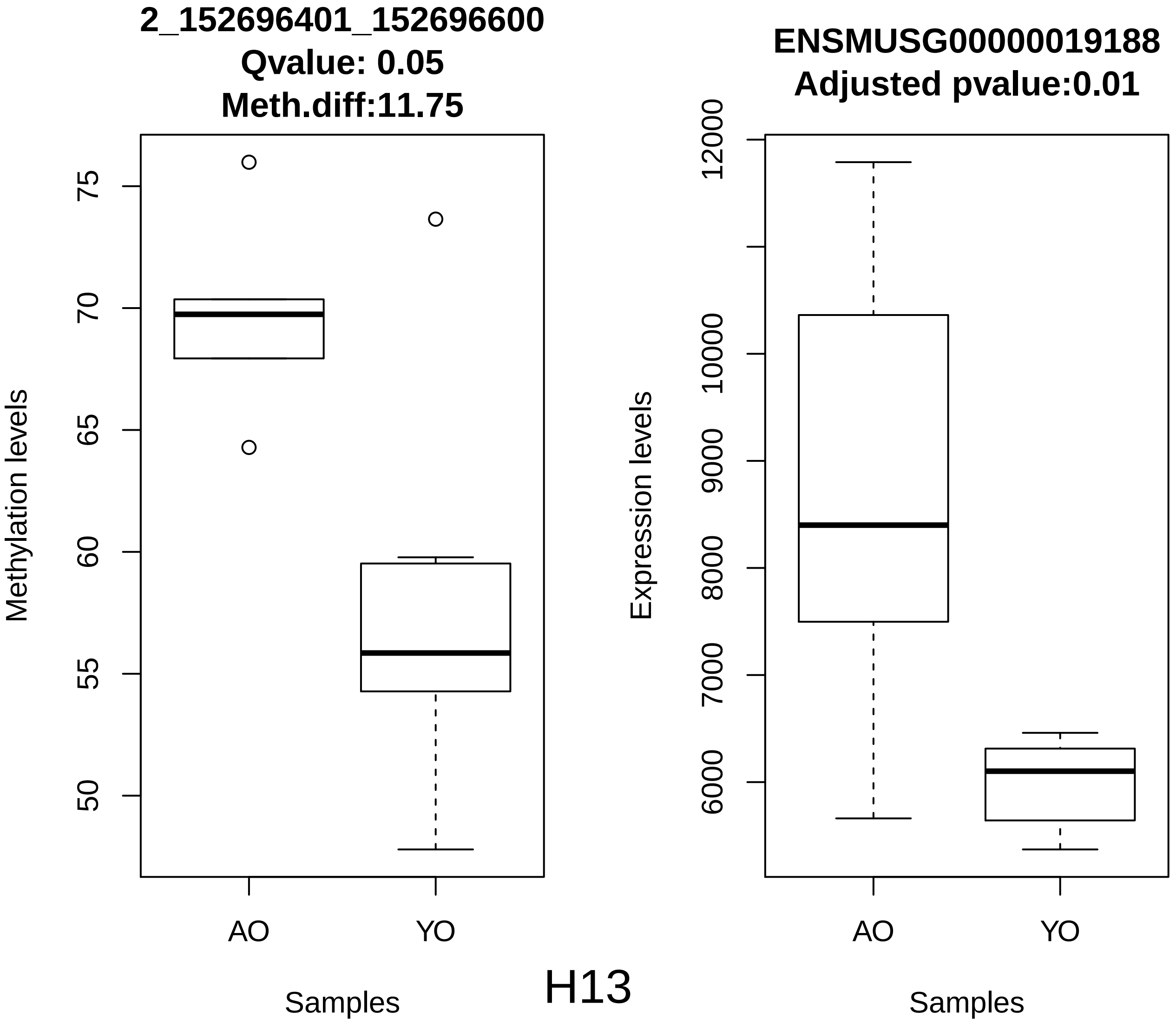
2_152696401_152696600
Qvalue: 0.05
Meth.diff:11.75
Methylation levels
75
70
65
60
55
50
AO
YO
Samples
ENSMUSG00000019188
Adjusted pvalue:0.01
Expression levels
12000
10000
9000
8000
7000
6000
AO
YO
Samples

H13

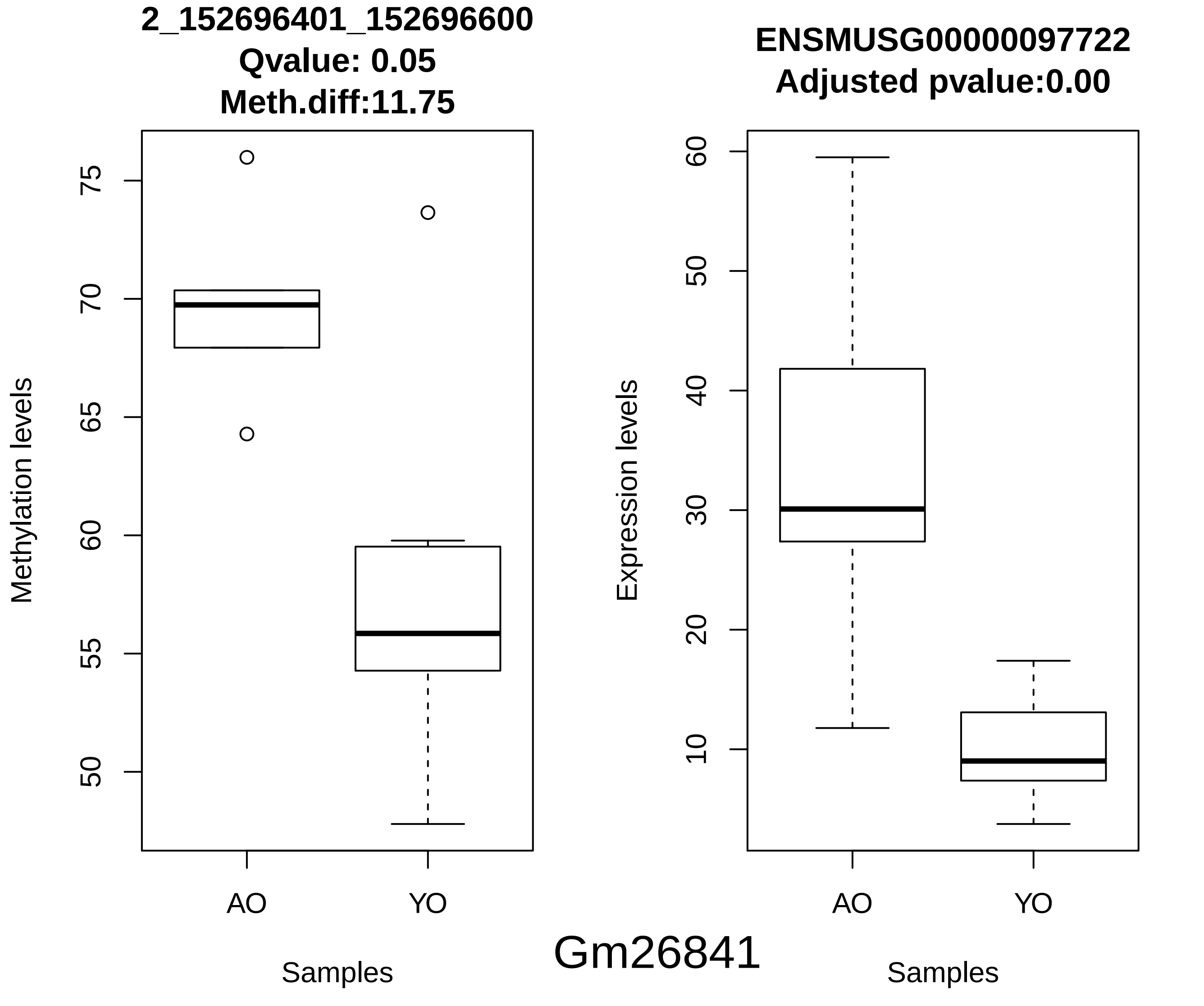
2_152696401_152696600
Qvalue: 0.05
Meth.diff:11.75
Methylation levels
75
70
65
60
55
50
AO
YO
Samples
ENSMUSG00000097722
Adjusted pvalue:0.00
Expression levels
60
50
40
30
20
10
AO
YO
Samples
Gm26841

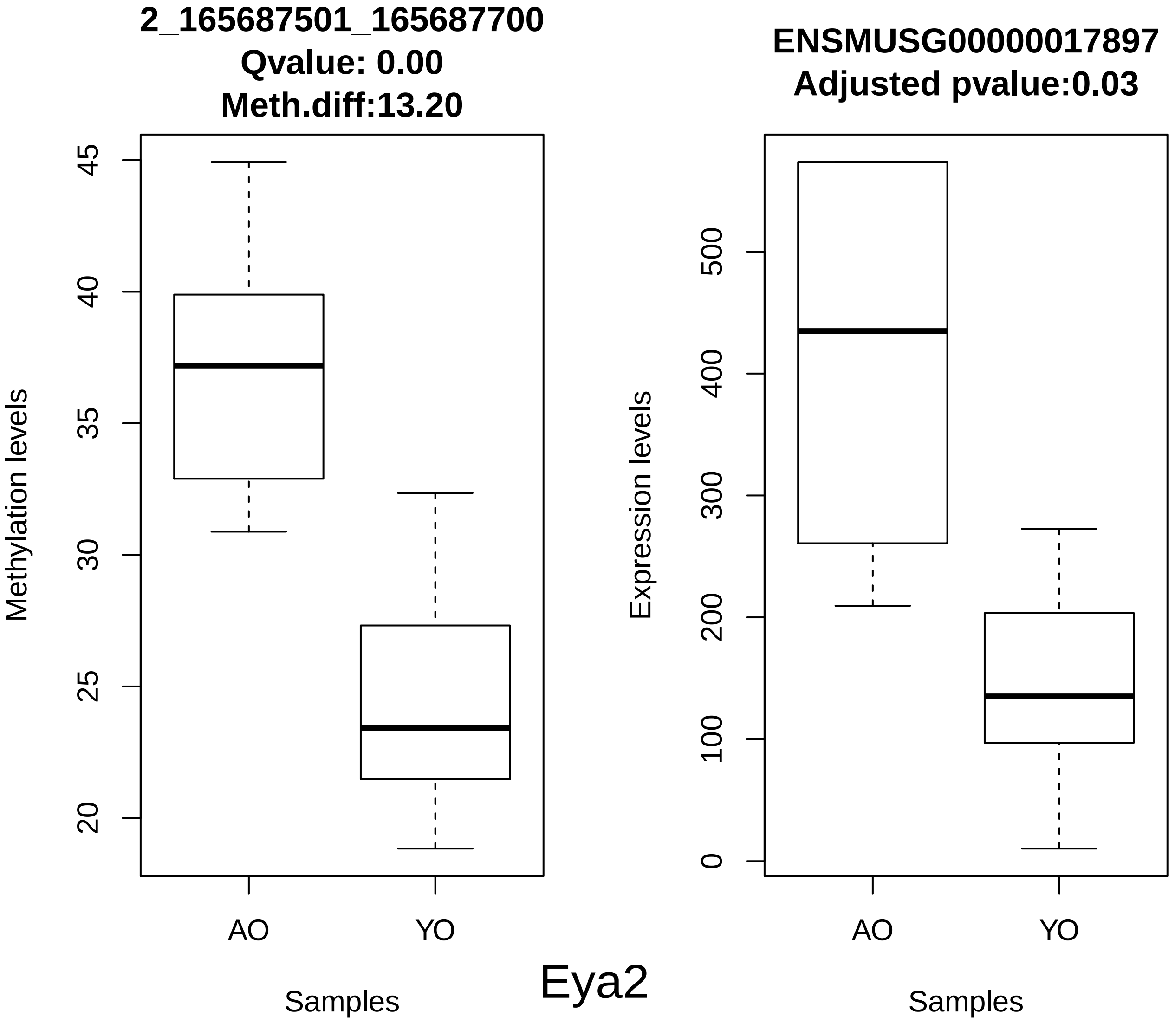

2_165687501_165687700
Qvalue: 0.00
Meth.diff:13.20
Methylation levels
45
40
35
30
25
20
AO
YO
Samples
ENSMUSG00000017897
Adjusted pvalue:0.03
Expression levels
500
400
300
200
100
0
AO
YO
Samples
Eya2

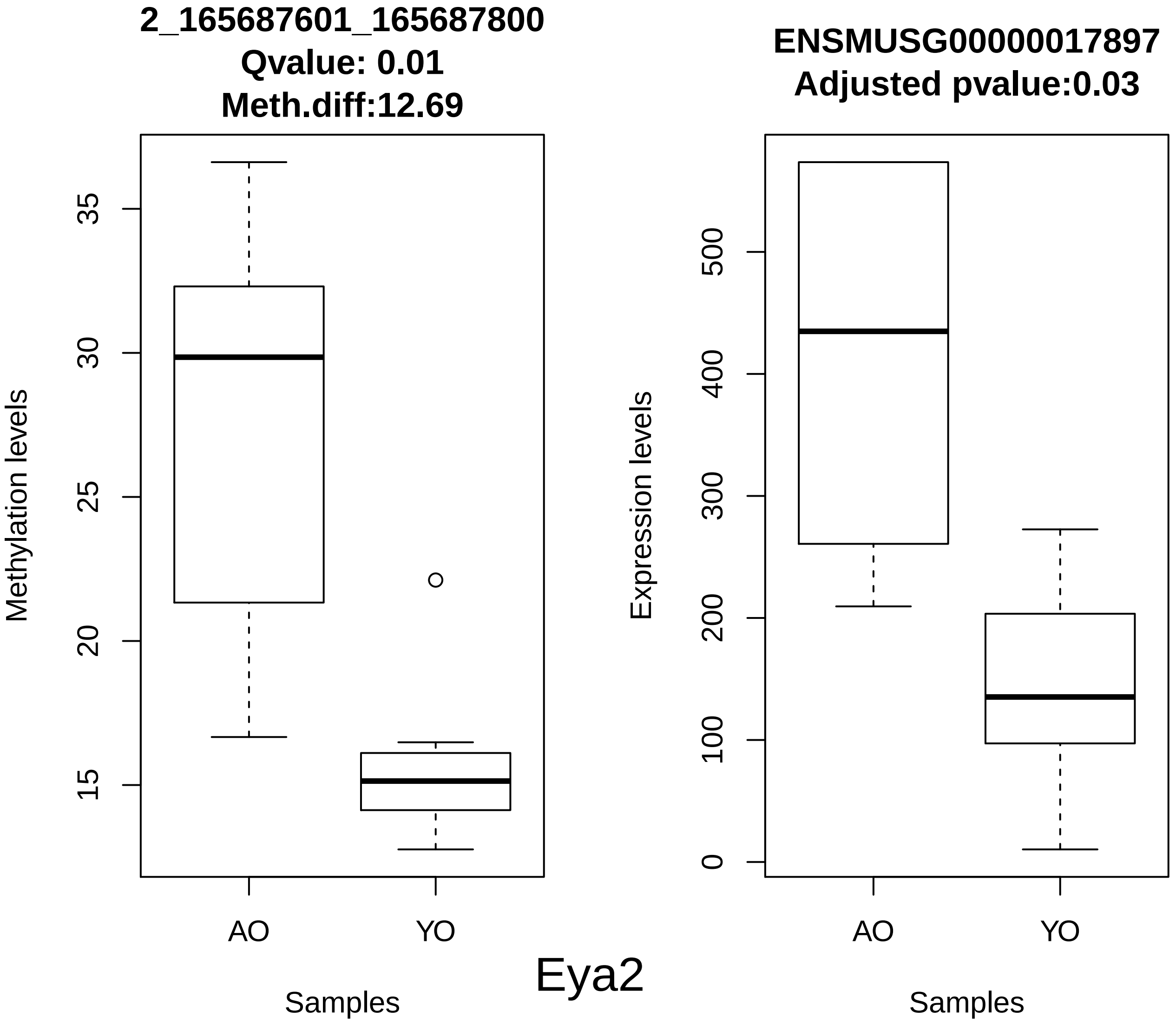
2_165687601_165687800
Qvalue: 0.01
Meth.diff:12.69
Methylation levels
35
30
25
20
15
AO
YO
Samples
Eya2
ENSMUSG00000017897
Adjusted pvalue:0.03
Expression levels
500
400
300
200
100
0
AO
YO
Samples

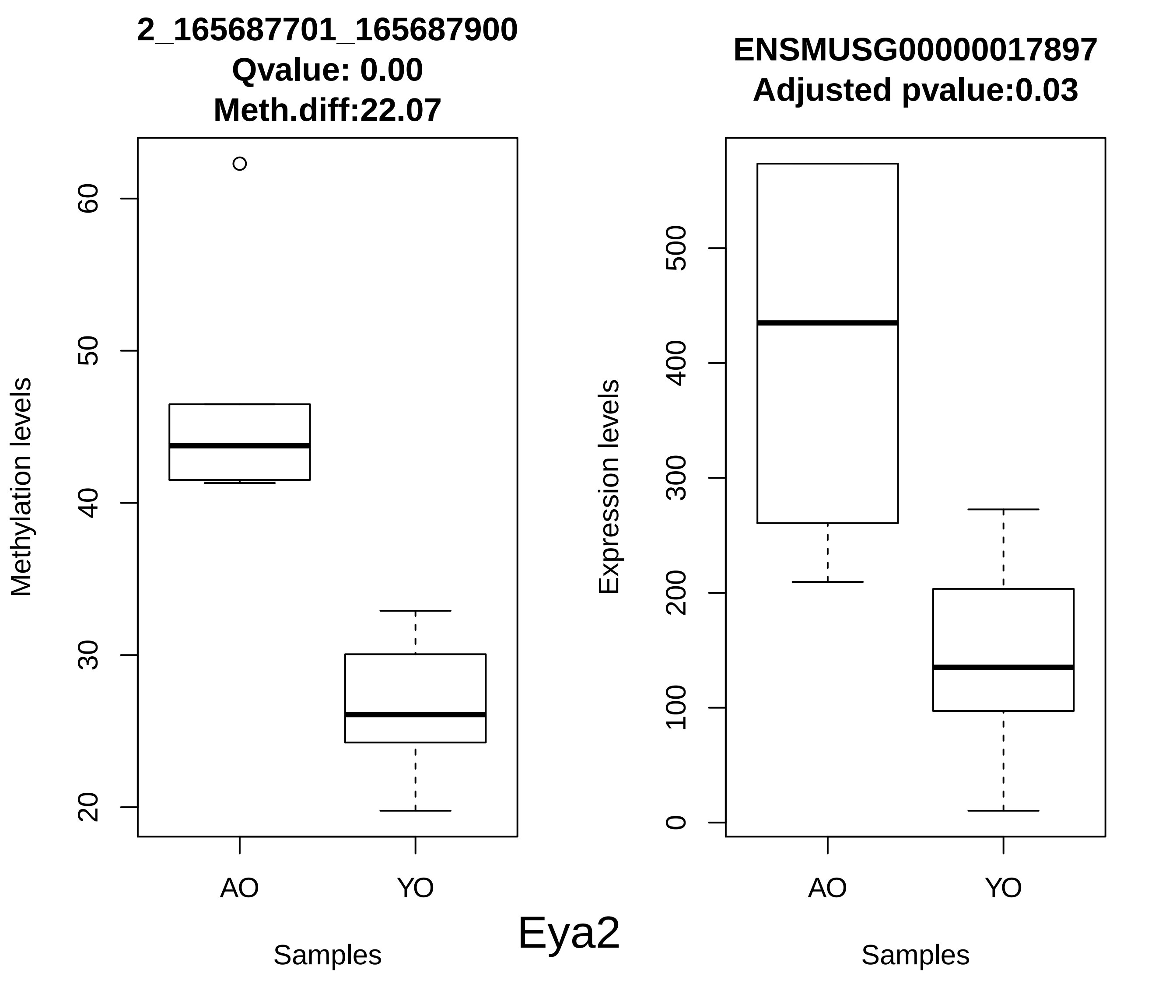

2_165687701_165687900
Qvalue: 0.00
Meth.diff:22.07
Methylation levels
60
50
40
30
20
AO
YO
Samples
ENSMUSG00000017897
Adjusted pvalue:0.03
Expression levels
500
400
300
200
100
0
AO
YO
Samples
Eya2

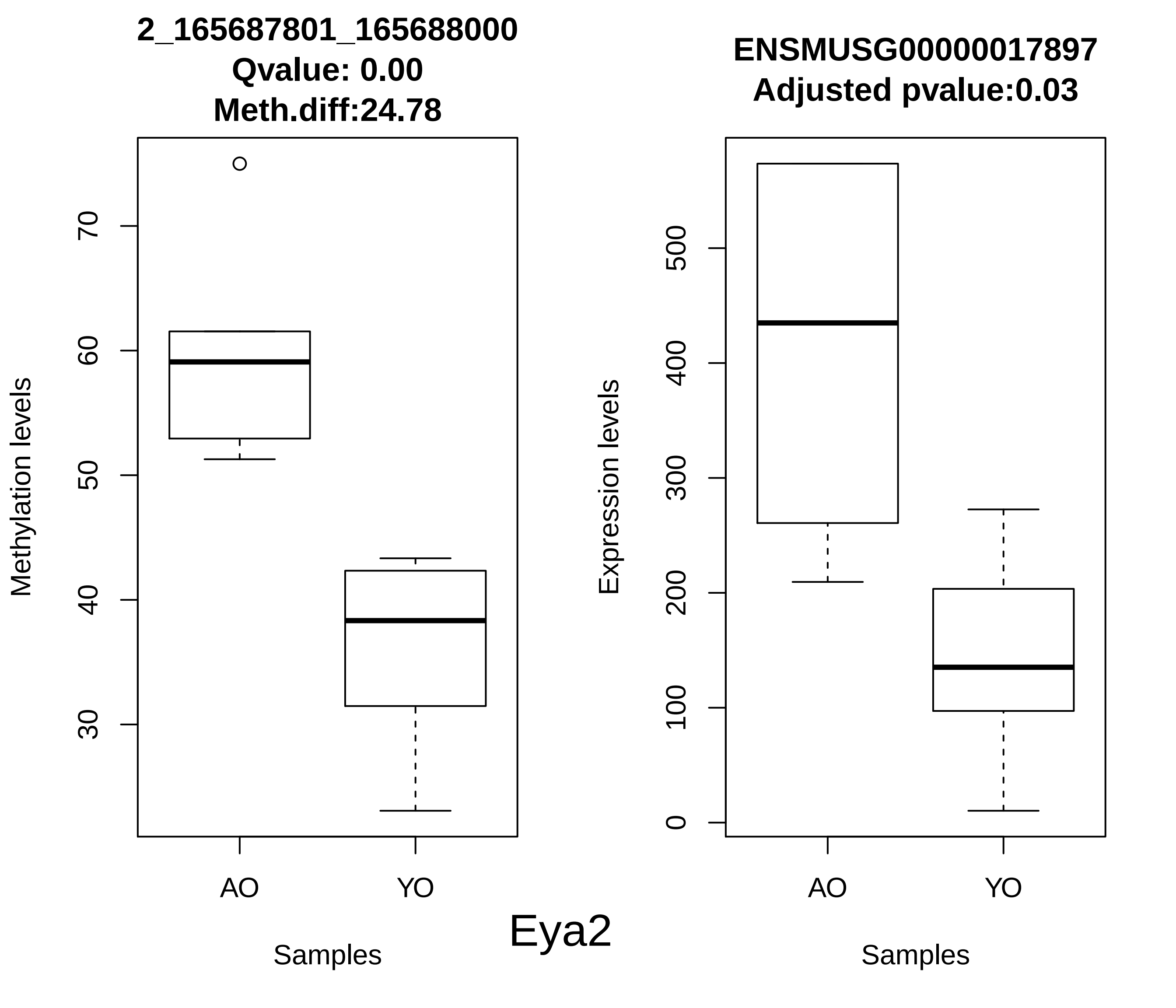

2_165687801_165688000
Qvalue: 0.00
Meth.diff:24.78
Methylation levels
70
60
50
40
30
AO
YO
Samples
ENSMUSG00000017897
Adjusted pvalue:0.03
Expression levels
500
400
300
200
100
0
AO
YO
Samples
Eya2

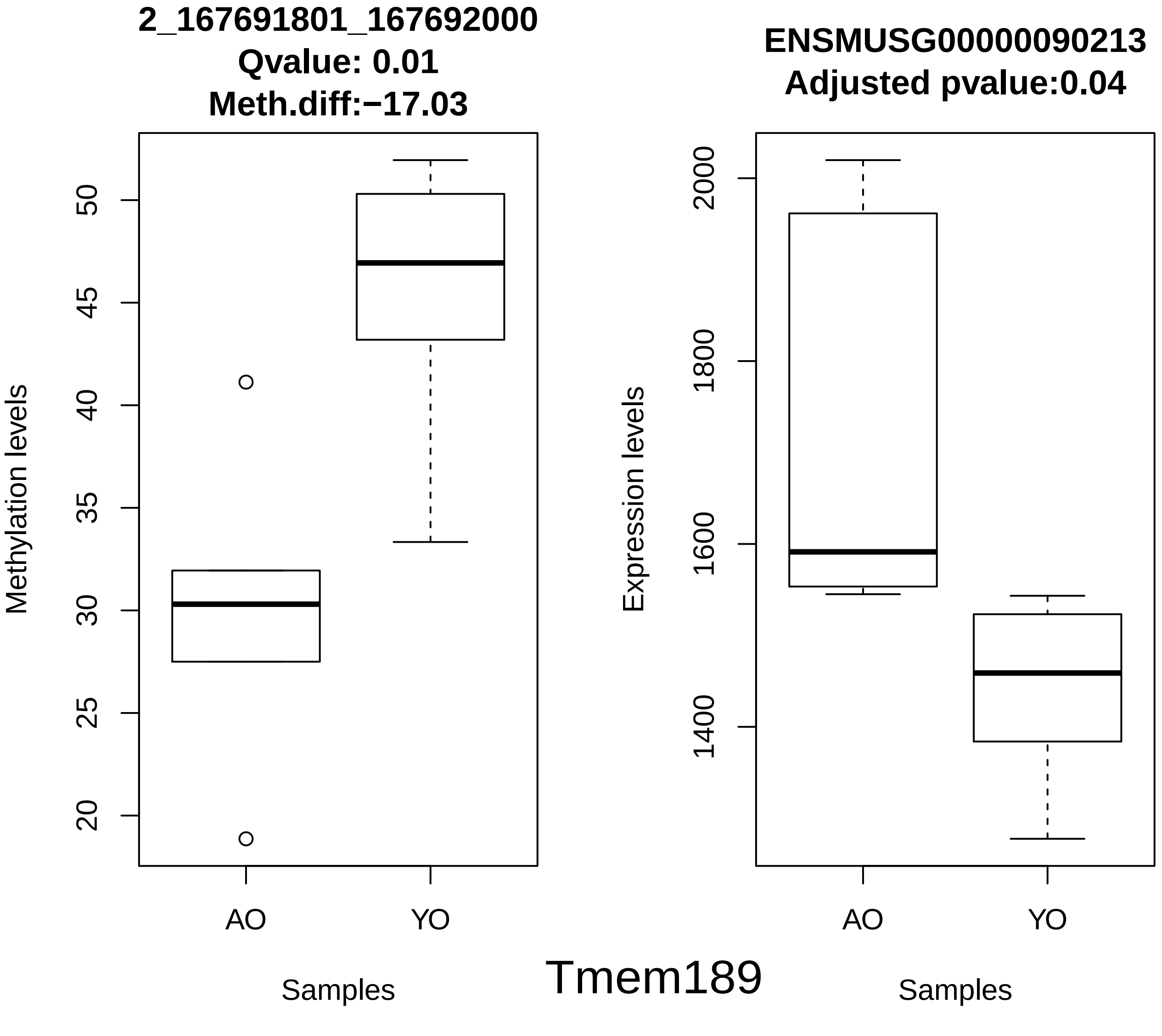
2_167691801_167692000
Qvalue: 0.01
Meth.diff:−17.03
Methylation levels
50
45
40
35
30
25
20
AO
YO
Samples
ENSMUSG00000090213
Adjusted pvalue:0.04
Expression levels
2000
1800
1600
1400
AO
YO
Samples
Tmem189

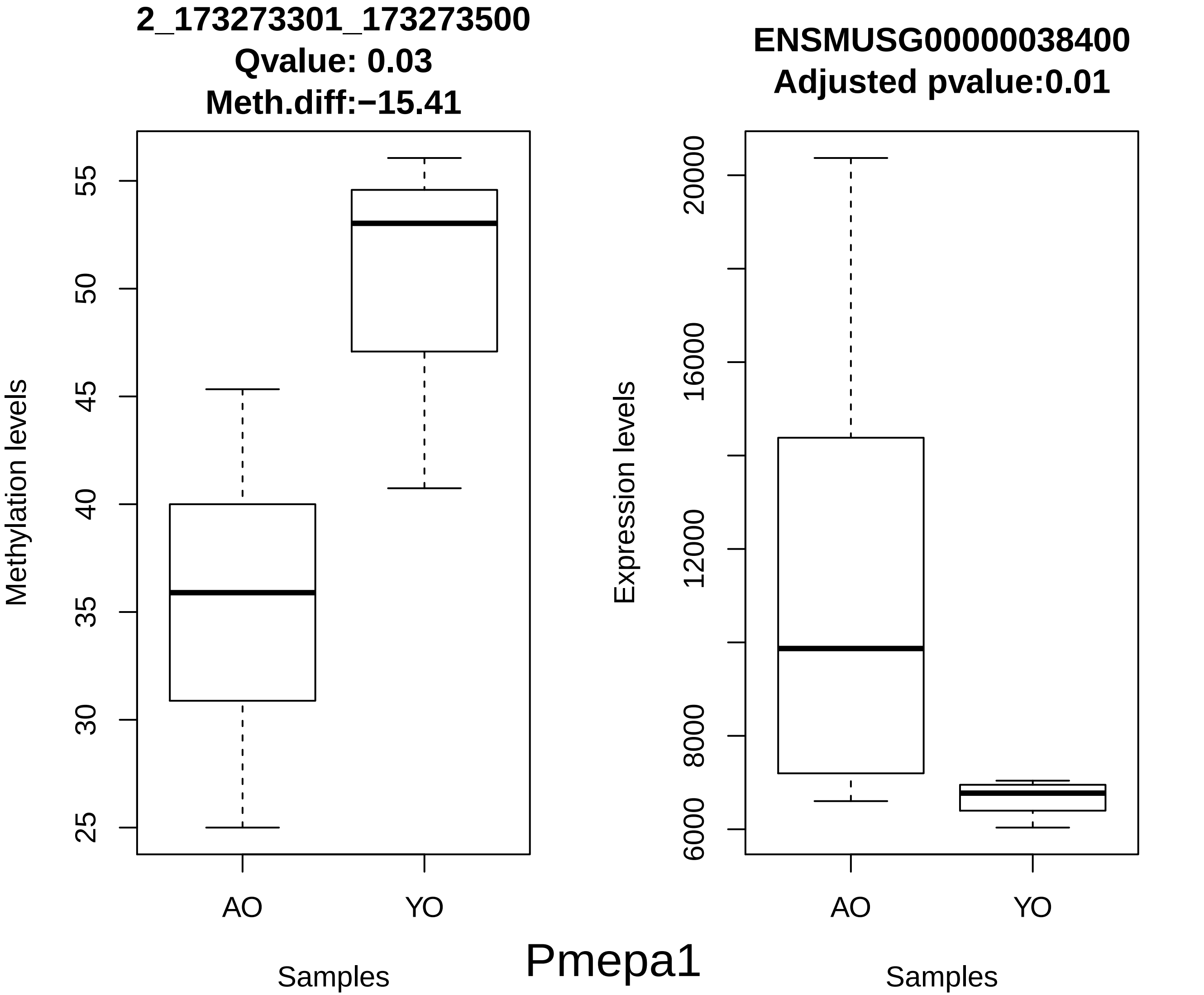

2_173273301_173273500
Qvalue: 0.03
Meth.diff:−15.41
Methylation levels
55
50
45
40
35
30
25
AO
YO
Samples
ENSMUSG00000038400
Adjusted pvalue:0.01
Expression levels
20000
16000
12000
8000
6000
AO
YO
Samples
Pmepa1

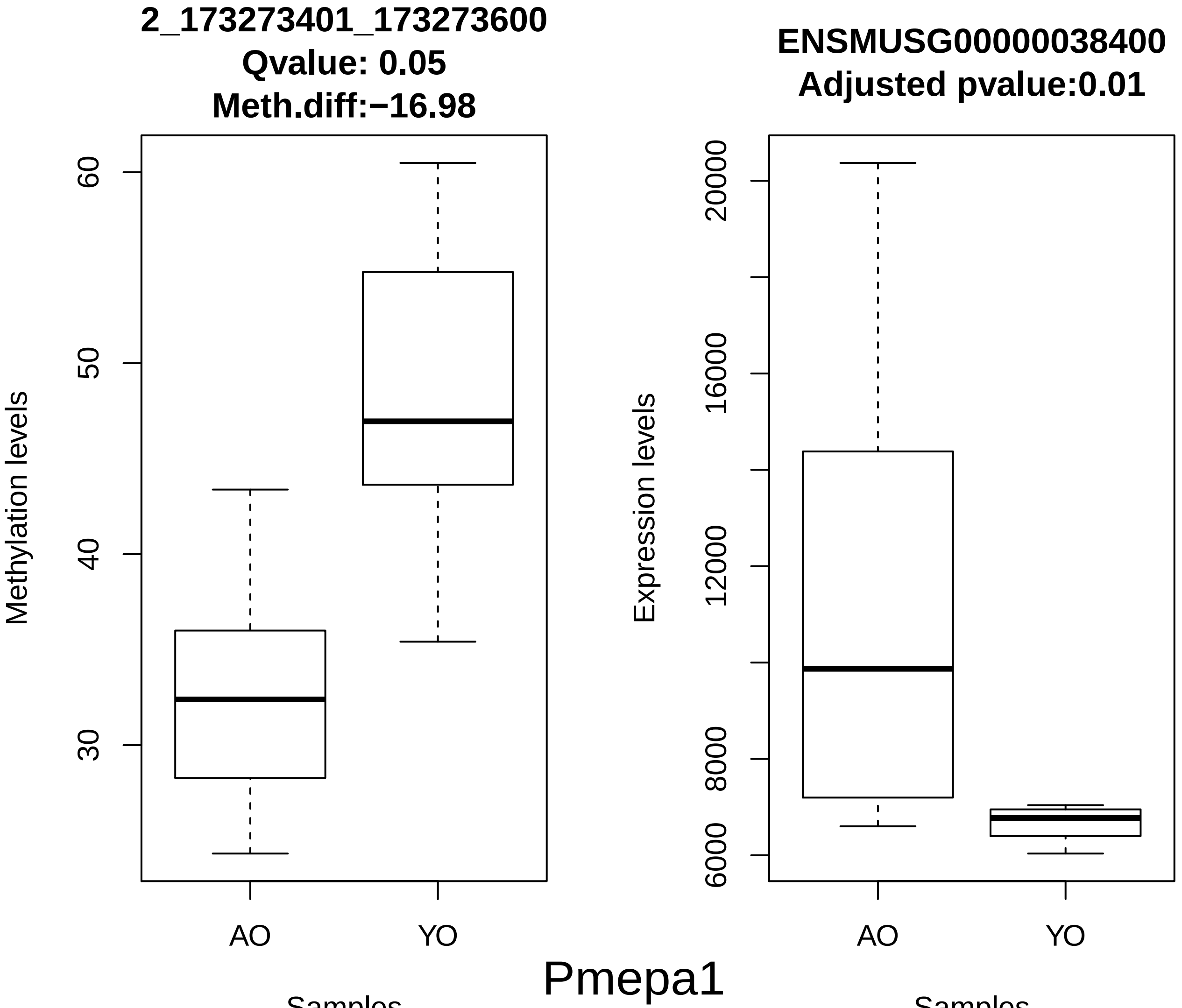
2_173273401_173273600
Qvalue: 0.05
Meth.diff:−16.98
Methylation levels
60
50
40
30
AO
YO
Samples
ENSMUSG00000038400
Adjusted pvalue:0.01
Expression levels
20000
16000
12000
8000
6000
AO
YO
Samples
Pmepa1

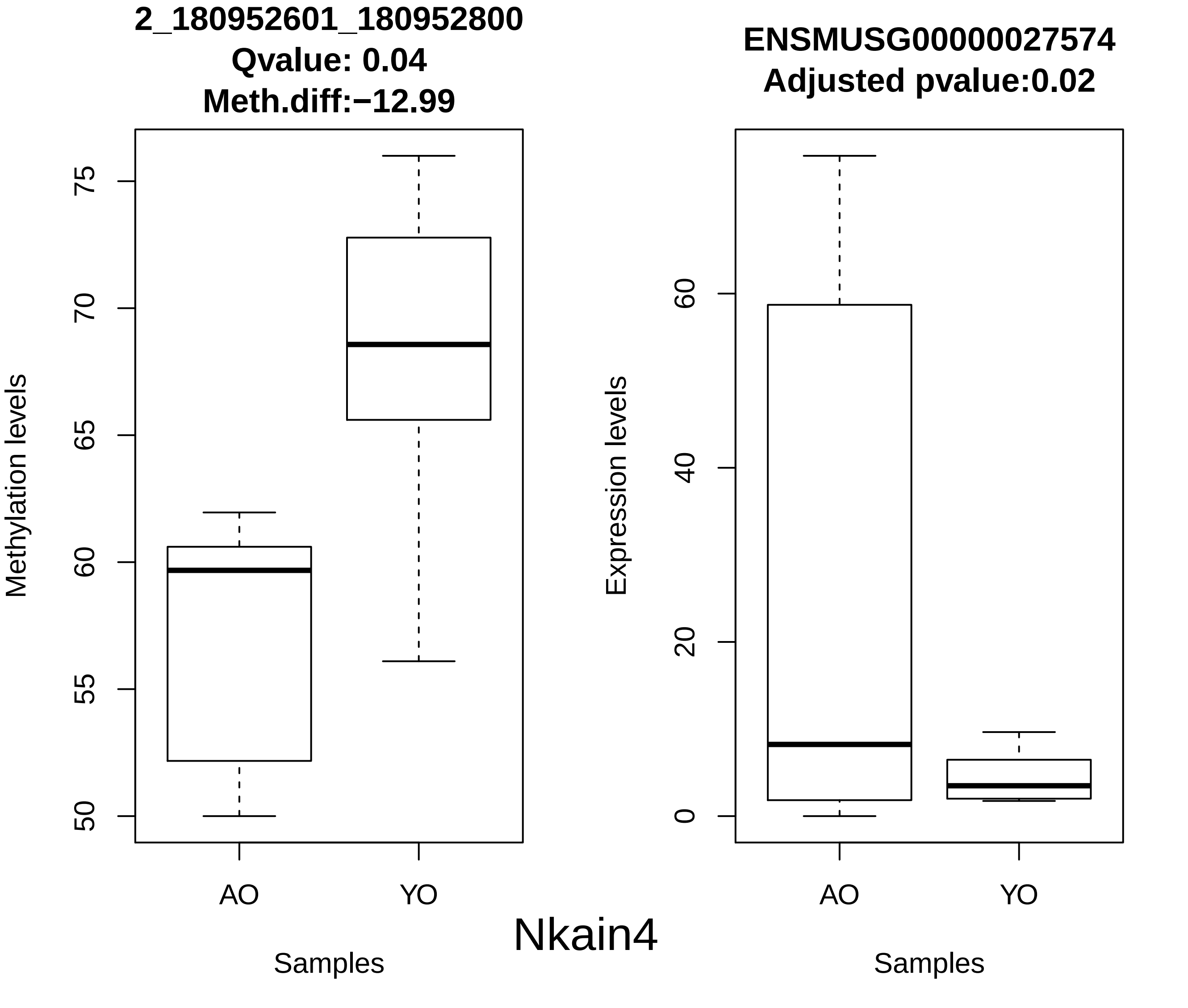

2_180952601_180952800
Qvalue: 0.04
Meth.diff:−12.99
Methylation levels
50
55
60
65
70
75
AO
YO
Samples
ENSMUSG00000027574
Adjusted pvalue:0.02
Expression levels
0
20
40
60
AO
YO
Samples
Nkain4

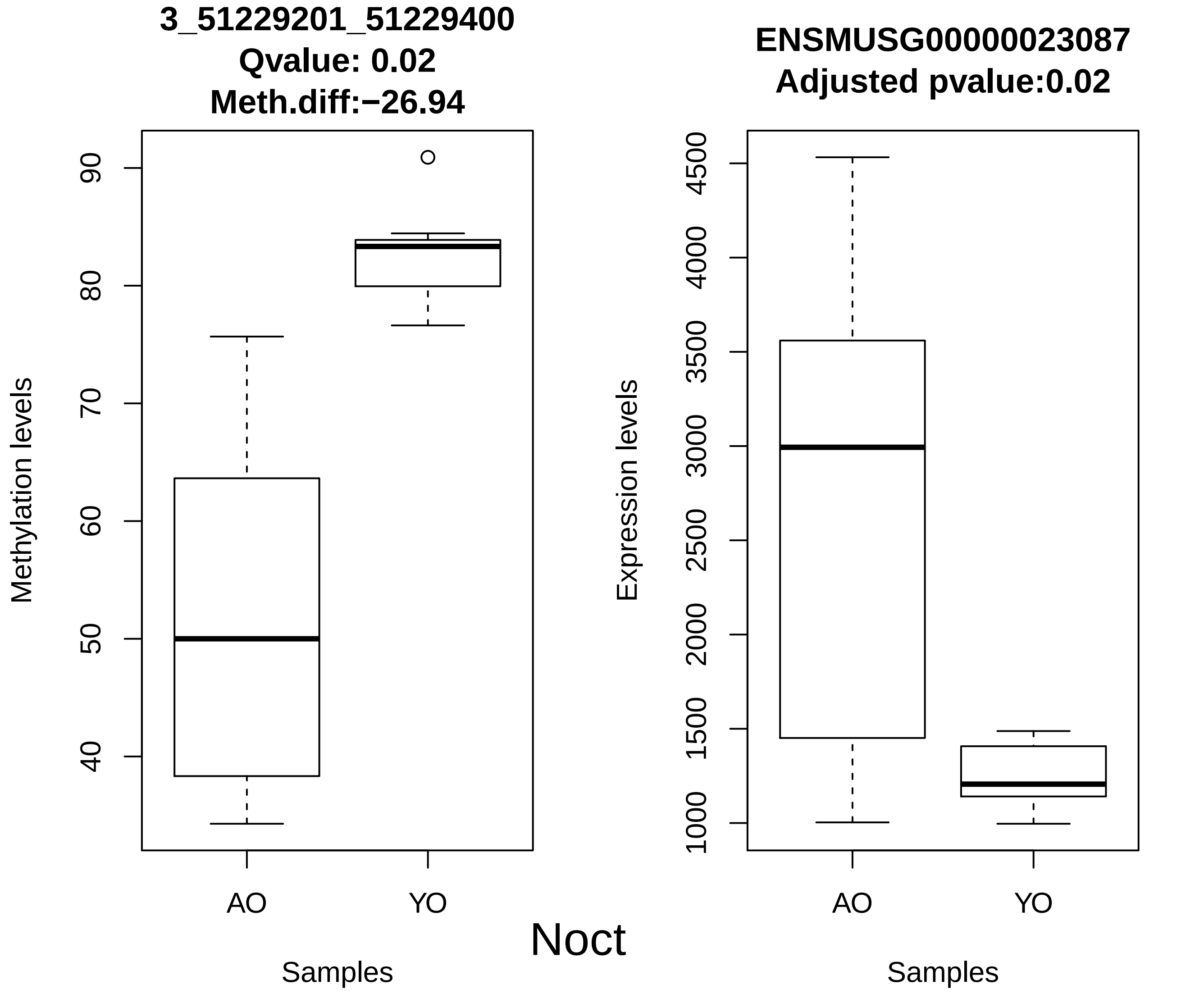

3_51229201_51229400
Qvalue: 0.02
Meth.diff:−26.94
Methylation levels
90
80
70
60
50
40
AO
YO
Samples
ENSMUSG00000023087
Adjusted pvalue:0.02
Expression levels
4500
4000
3500
3000
2500
2000
1500
1000
AO
YO
Samples
Noct

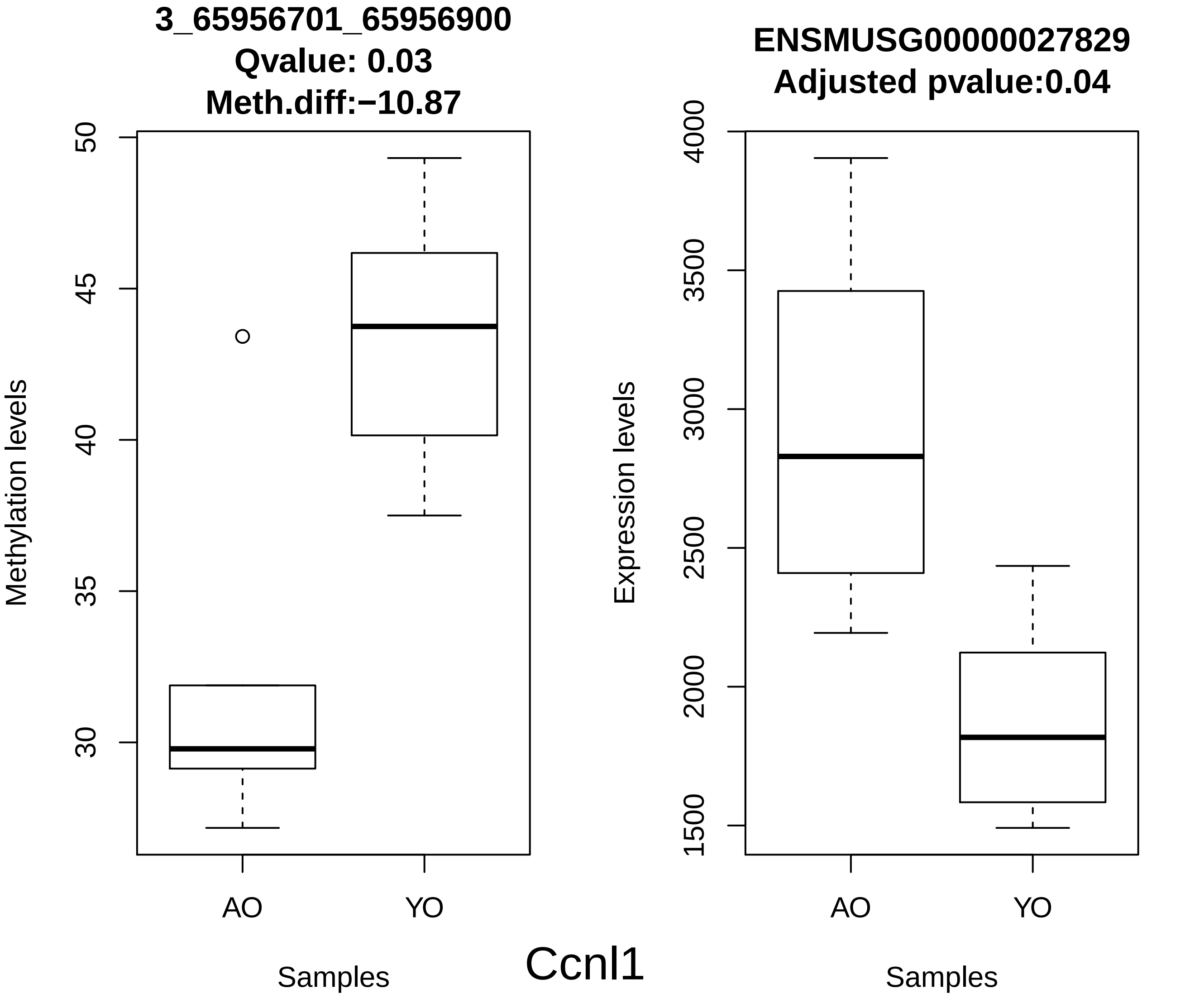

3_65956701_65956900
Qvalue: 0.03
Meth.diff:−10.87
Methylation levels
50
45
40
35
30
AO
YO
Samples
ENSMUSG00000027829
Adjusted pvalue:0.04
Expression levels
4000
3500
3000
2500
2000
1500
AO
YO
Samples
Ccnl1

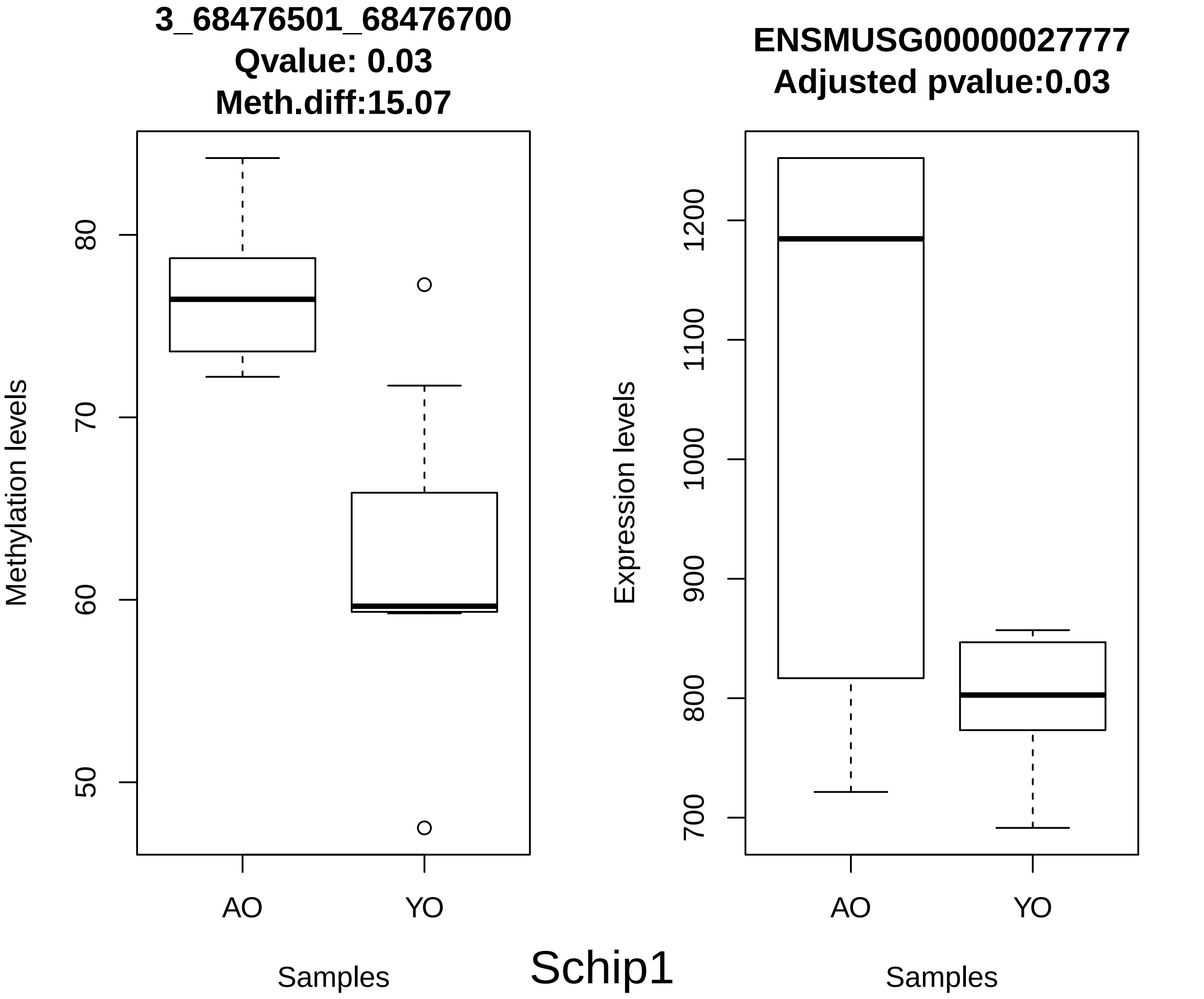

3_68476501_68476700
Qvalue: 0.03
Meth.diff:15.07
Methylation levels
80
70
60
50
AO
YO
Samples
ENSMUSG00000027777
Adjusted pvalue:0.03
Expression levels
1200
1100
1000
900
800
700
AO
YO
Samples
Schip1

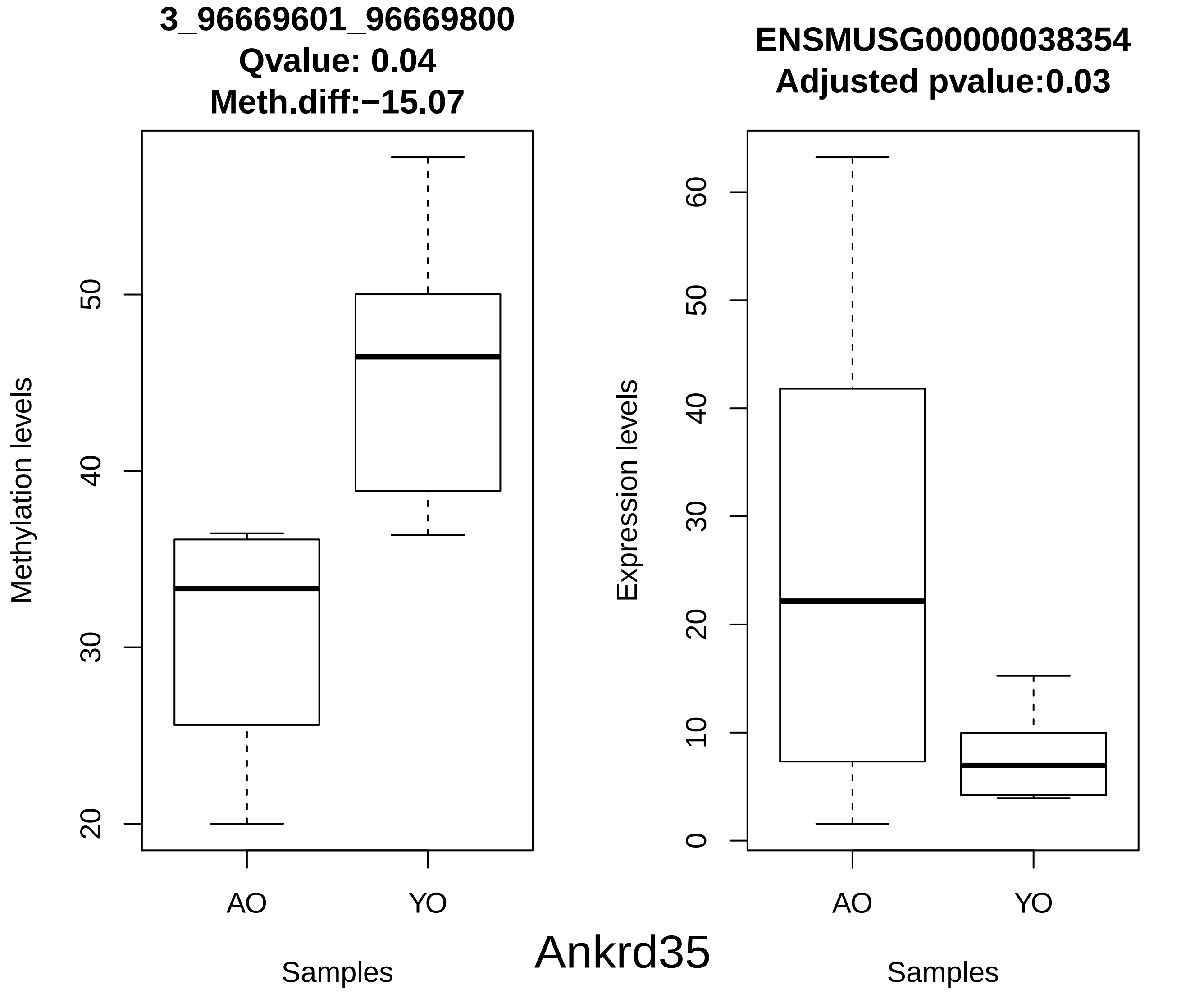
3_96669601_96669800
Qvalue: 0.04
Meth.diff:−15.07
Methylation levels
20
30
40
50
AO
YO
Samples
ENSMUSG00000038354
Adjusted pvalue:0.03
Expression levels
0
10
20
30
40
50
60
AO
YO
Samples
Ankrd35

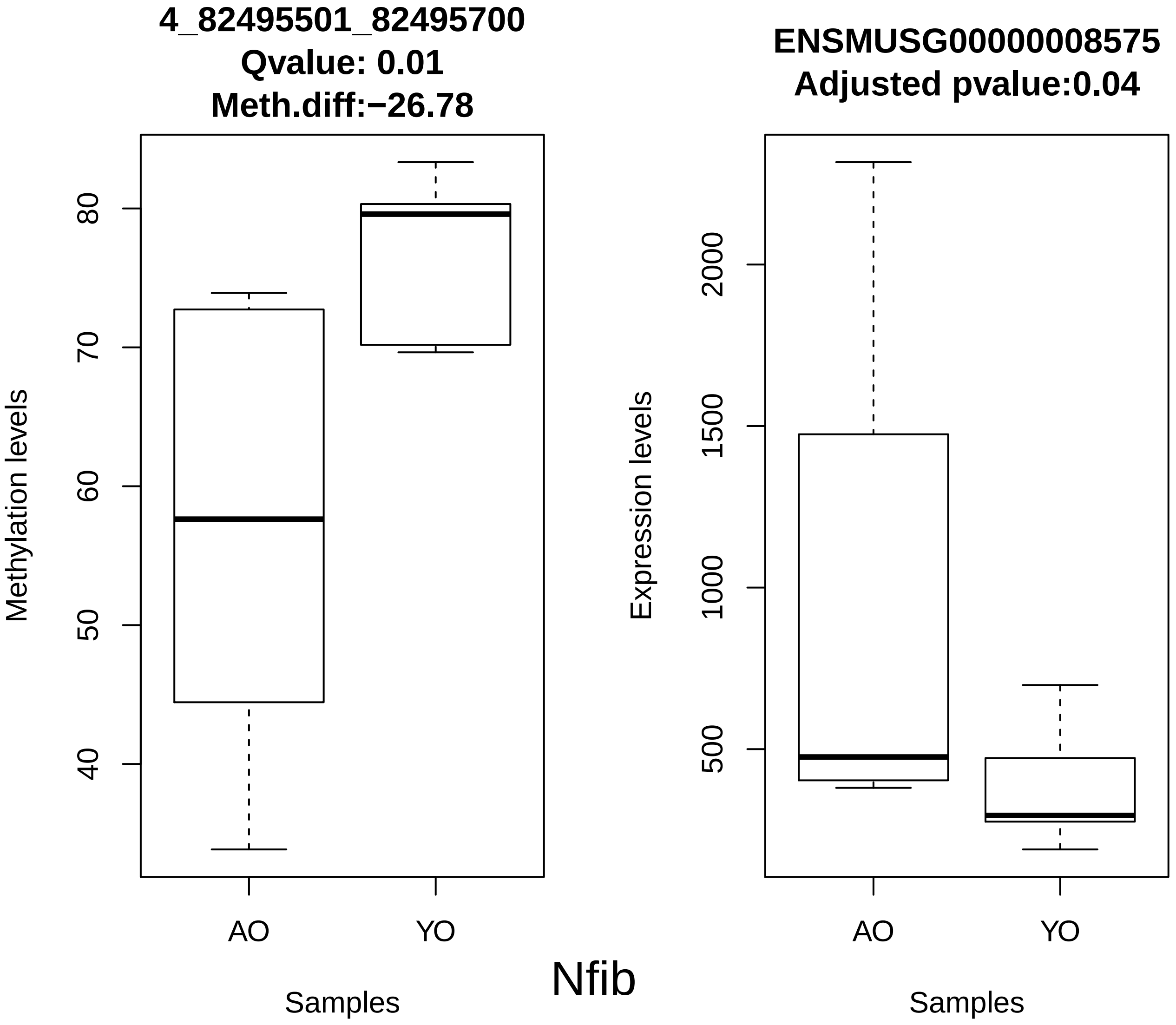
4_82495501_82495700
Qvalue: 0.01
Meth.diff:−26.78
Methylation levels
80
70
60
50
40
AO
YO
Samples
ENSMUSG00000008575
Adjusted pvalue:0.04
Expression levels
2000
1500
1000
500
AO
YO
Samples
Nfib

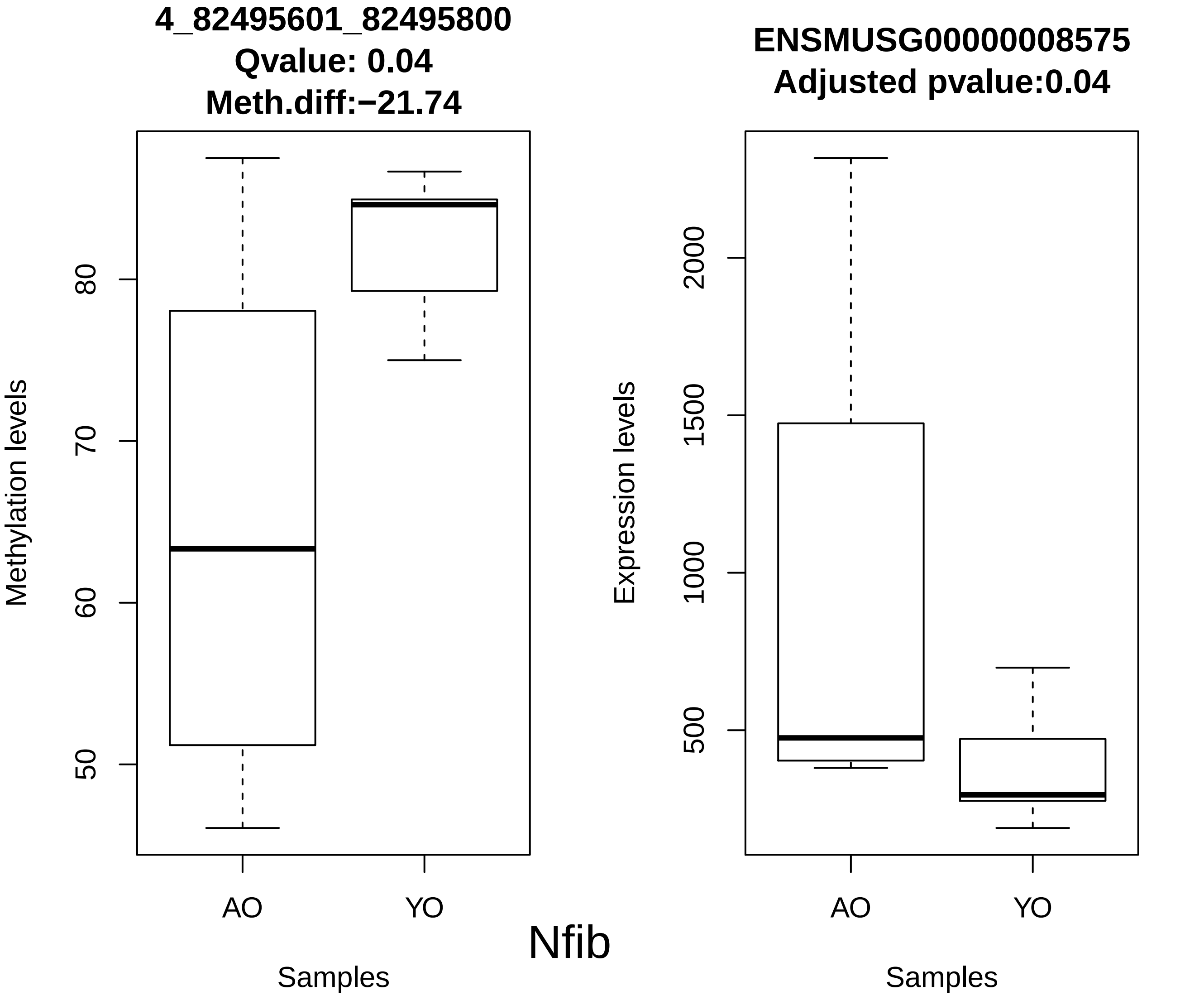
4_82495601_82495800
Qvalue: 0.04
Meth.diff:−21.74
Methylation levels
50
60
70
80
AO
YO
Samples
ENSMUSG00000008575
Adjusted pvalue:0.04
Expression levels
500
1000
1500
2000
AO
YO
Samples
Nfib

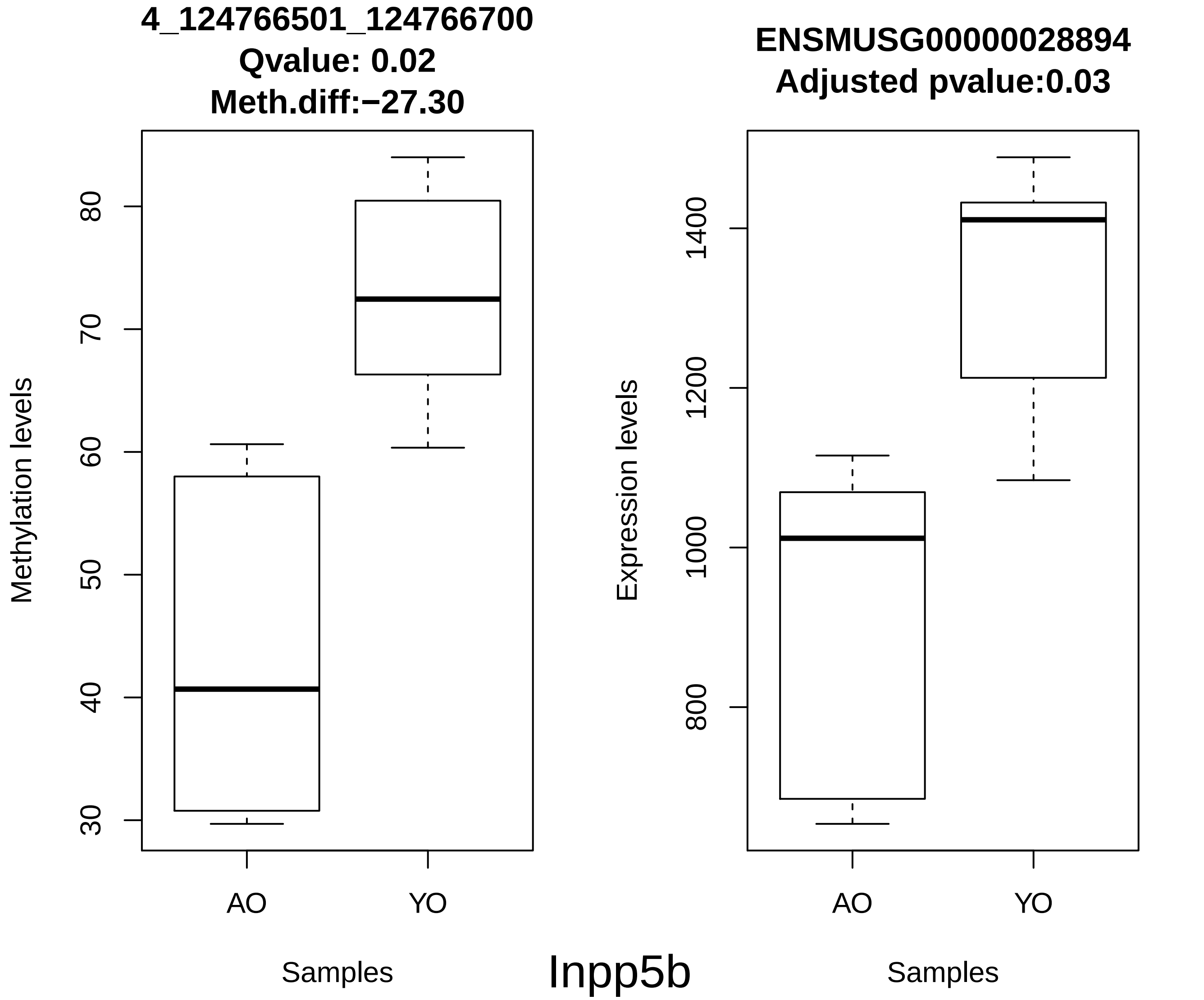
4_124766501_124766700
Qvalue: 0.02
Meth.diff:−27.30
Methylation levels
30
40
50
60
70
80
AO
YO
Samples
ENSMUSG00000028894
Adjusted pvalue:0.03
Expression levels
800
1000
1200
1400
AO
YO
Samples
Inpp5b

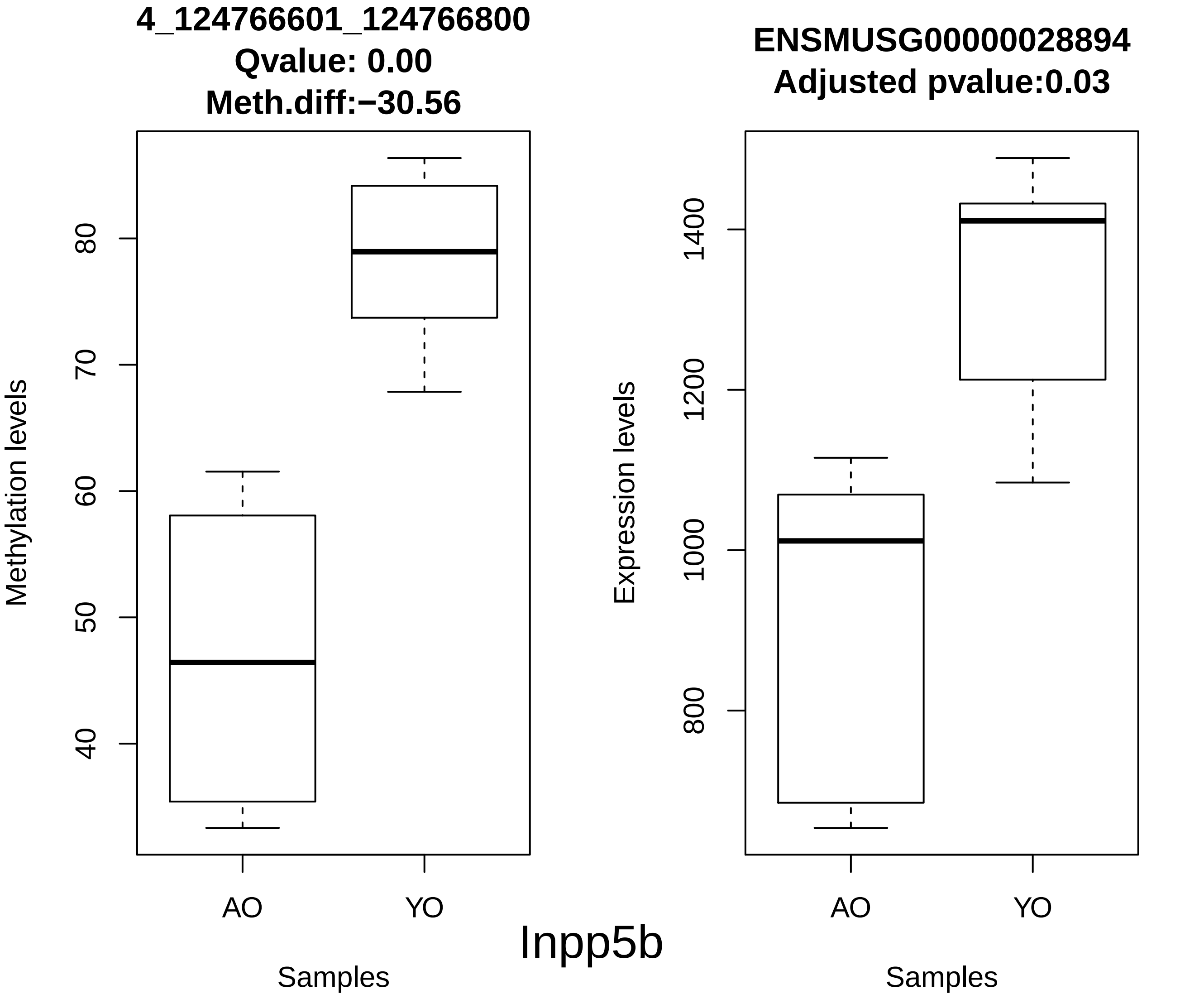
4_124766601_124766800
Qvalue: 0.00
Meth.diff:−30.56
Methylation levels
40
50
60
70
80
AO
YO
Samples
ENSMUSG00000028894
Adjusted pvalue:0.03
Expression levels
800
1000
1200
1400
AO
YO
Samples
Inpp5b

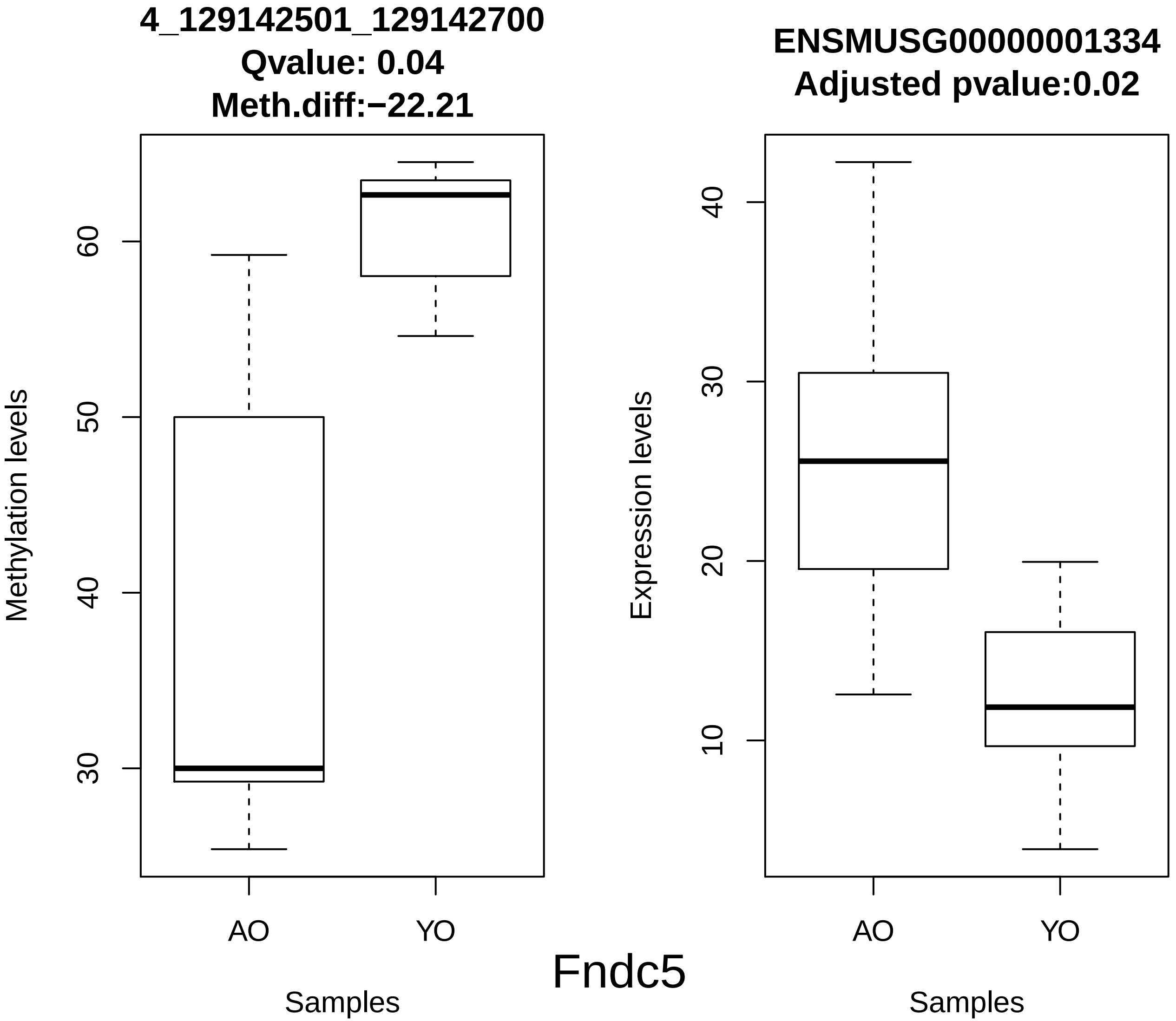
4_129142501_129142700
Qvalue: 0.04
Meth.diff:−22.21
Methylation levels
30
40
50
60
AO
YO
Samples
ENSMUSG00000001334
Adjusted pvalue:0.02
Expression levels
10
20
30
40
AO
YO
Samples
Fndc5

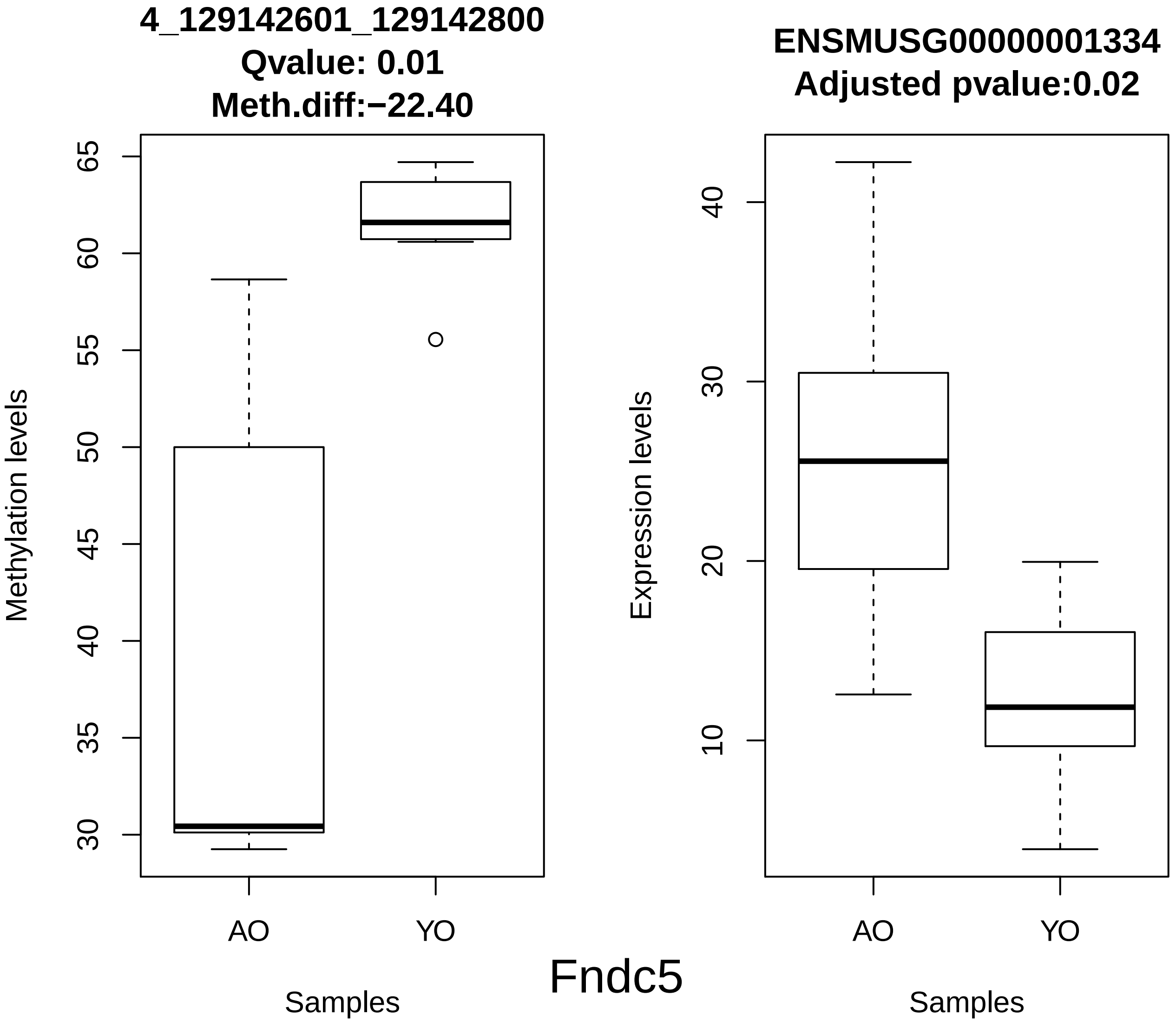
4_129142601_129142800
Qvalue: 0.01
Meth.diff:−22.40
Methylation levels
65
60
55
50
45
40
35
30
AO
YO
Samples
ENSMUSG00000001334
Adjusted pvalue:0.02
Expression levels
40
30
20
10
AO
YO
Samples
Fndc5

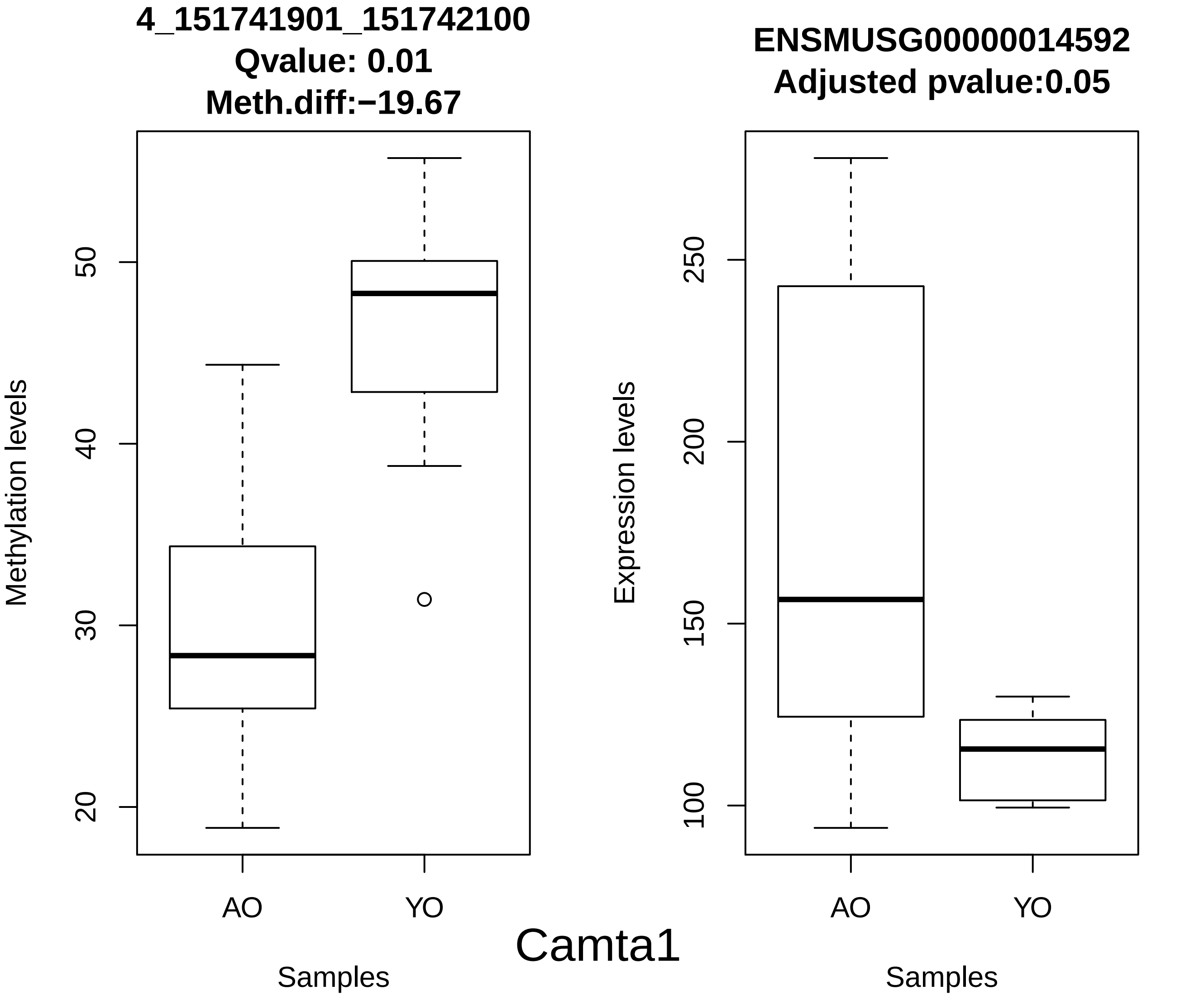
4_151741901_151742100
Qvalue: 0.01
Meth.diff:−19.67
Methylation levels
20
30
40
50
AO
YO
Samples
ENSMUSG00000014592
Adjusted pvalue:0.05
Expression levels
100
150
200
250
AO
YO
Samples
Camta1

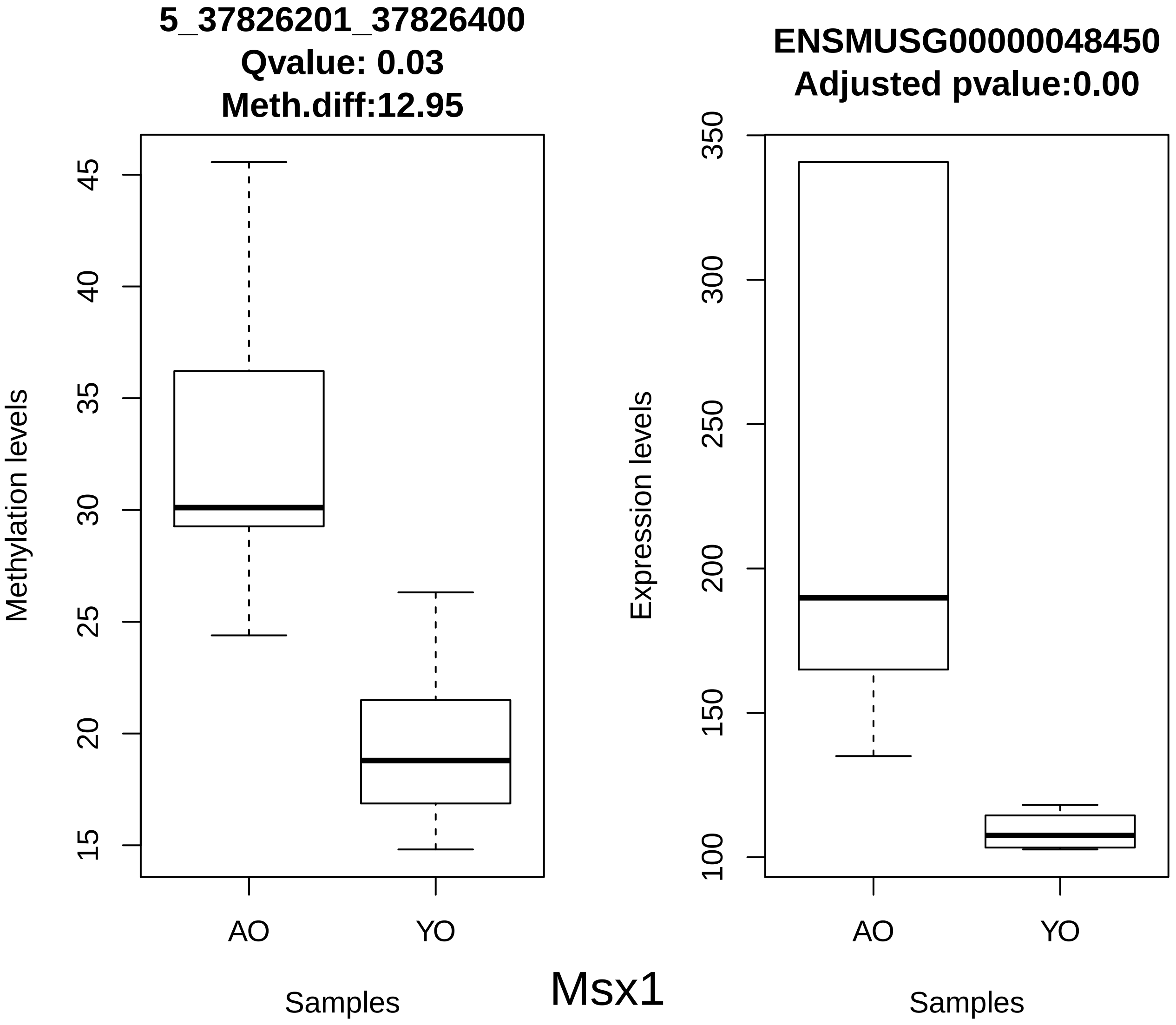
5_37826201_37826400
Qvalue: 0.03
Meth.diff:12.95
Methylation levels
15
20
25
30
35
40
45
AO
YO
Samples
ENSMUSG00000048450
Adjusted pvalue:0.00
Expression levels
100
150
200
250
300
350
AO
YO
Samples

Msx1

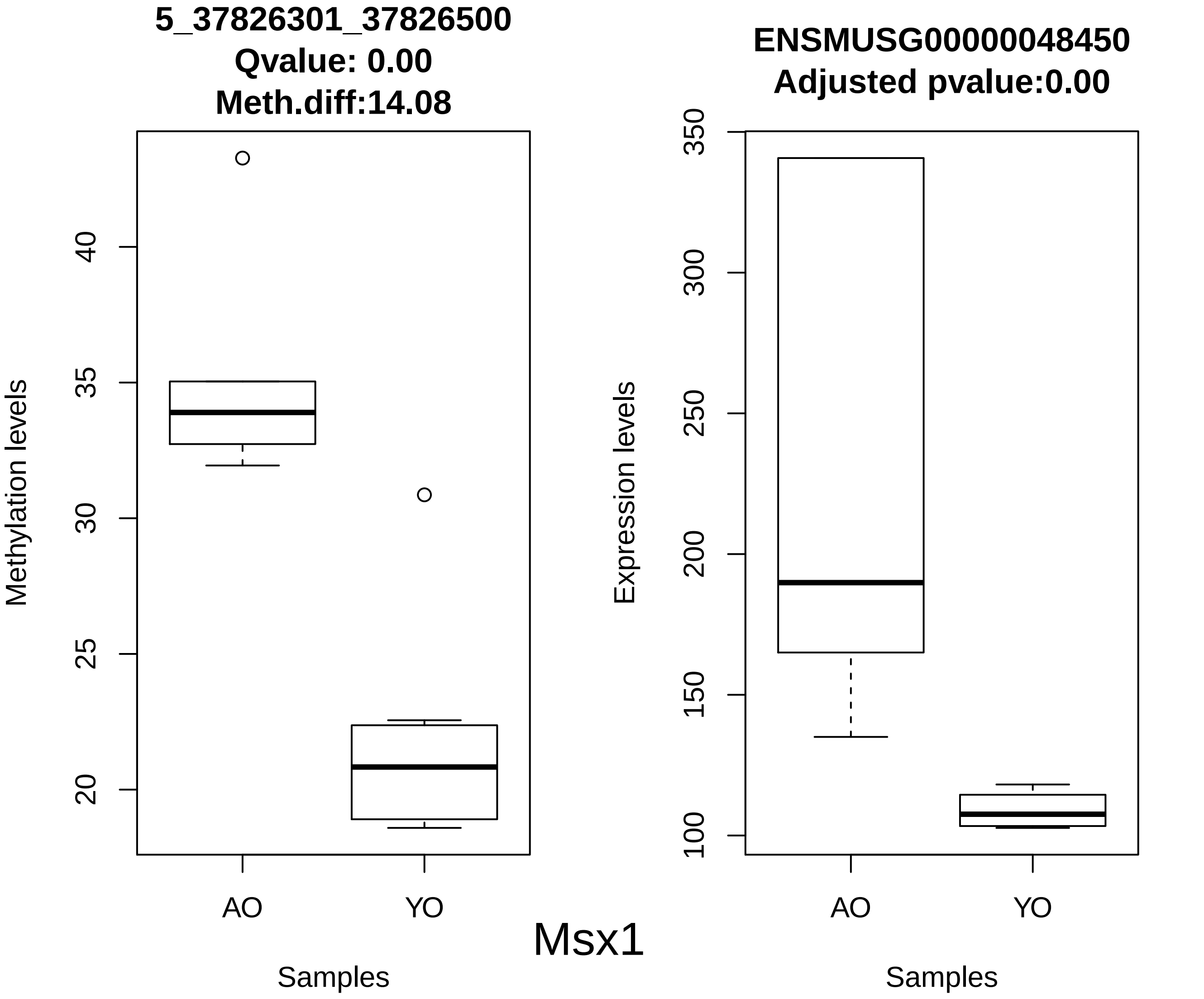
5_37826301_37826500
Qvalue: 0.00
Meth.diff:14.08
Methylation levels
40
35
30
25
20
AO
YO
Samples
ENSMUSG00000048450
Adjusted pvalue:0.00
Expression levels
350
300
250
200
150
100
AO
YO
Samples
Msx1

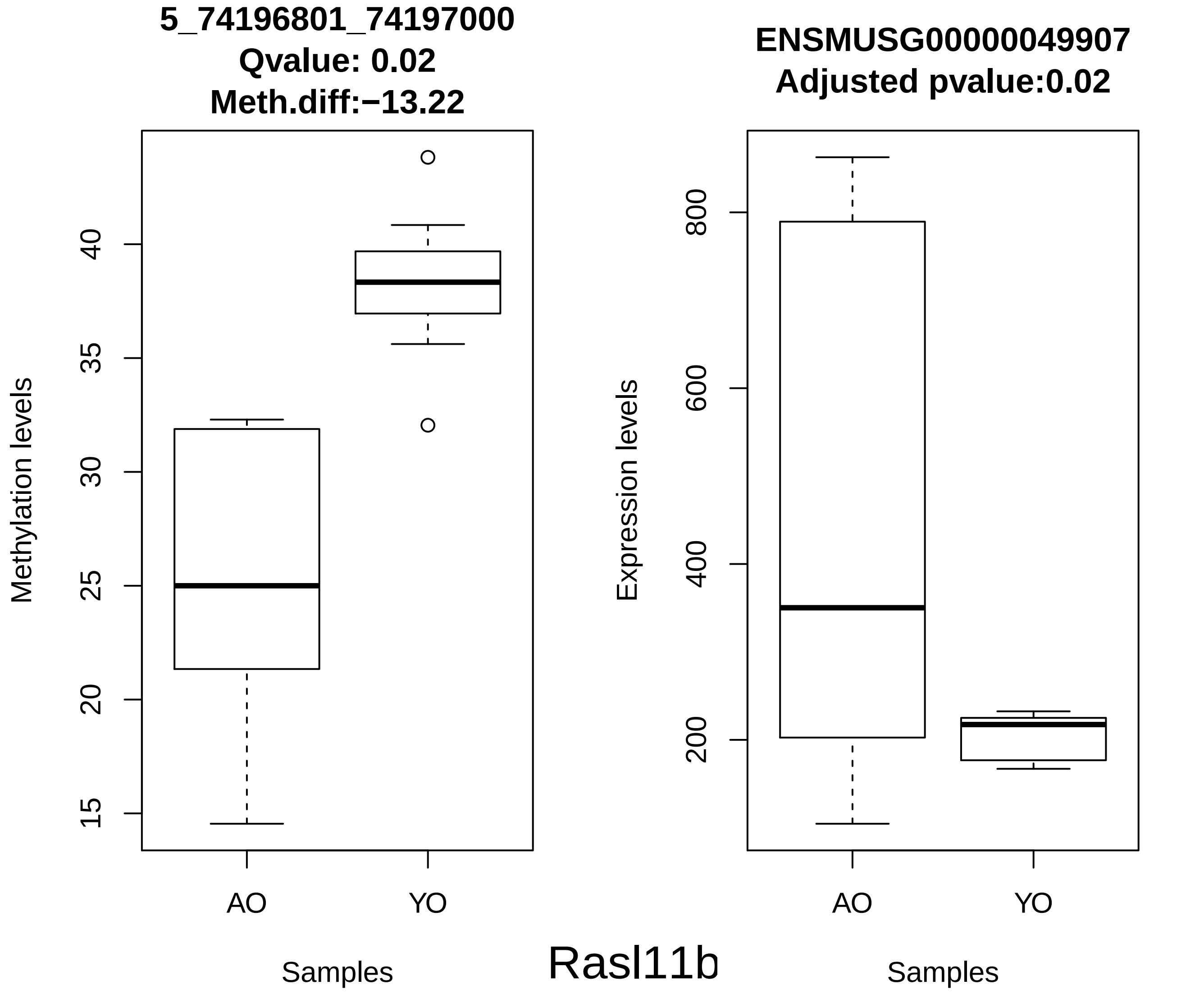

5_74196801_74197000
Qvalue: 0.02
Meth.diff:−13.22
Methylation levels
15
20
25
30
35
40
AO
YO
Samples
ENSMUSG00000049907
Adjusted pvalue:0.02
Expression levels
200
400
600
800
AO
YO
Samples
Rasl11b

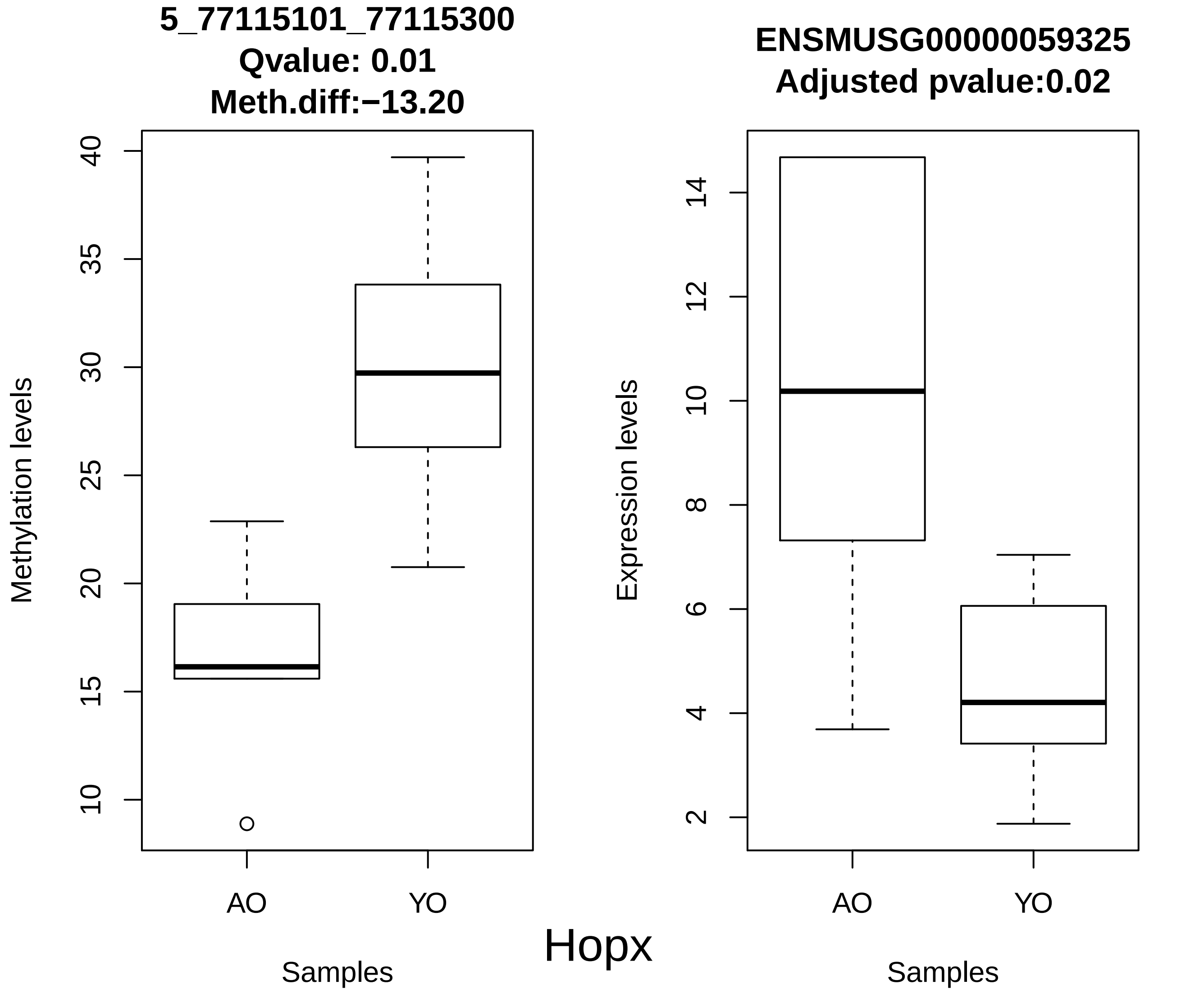
5_77115101_77115300
Qvalue: 0.01
Meth.diff:−13.20
Methylation levels
40
35
30
25
20
15
10
AO
YO
Samples
ENSMUSG00000059325
Adjusted pvalue:0.02
Expression levels
14
12
10
8
6
4
2
AO
YO
Samples
Hopx

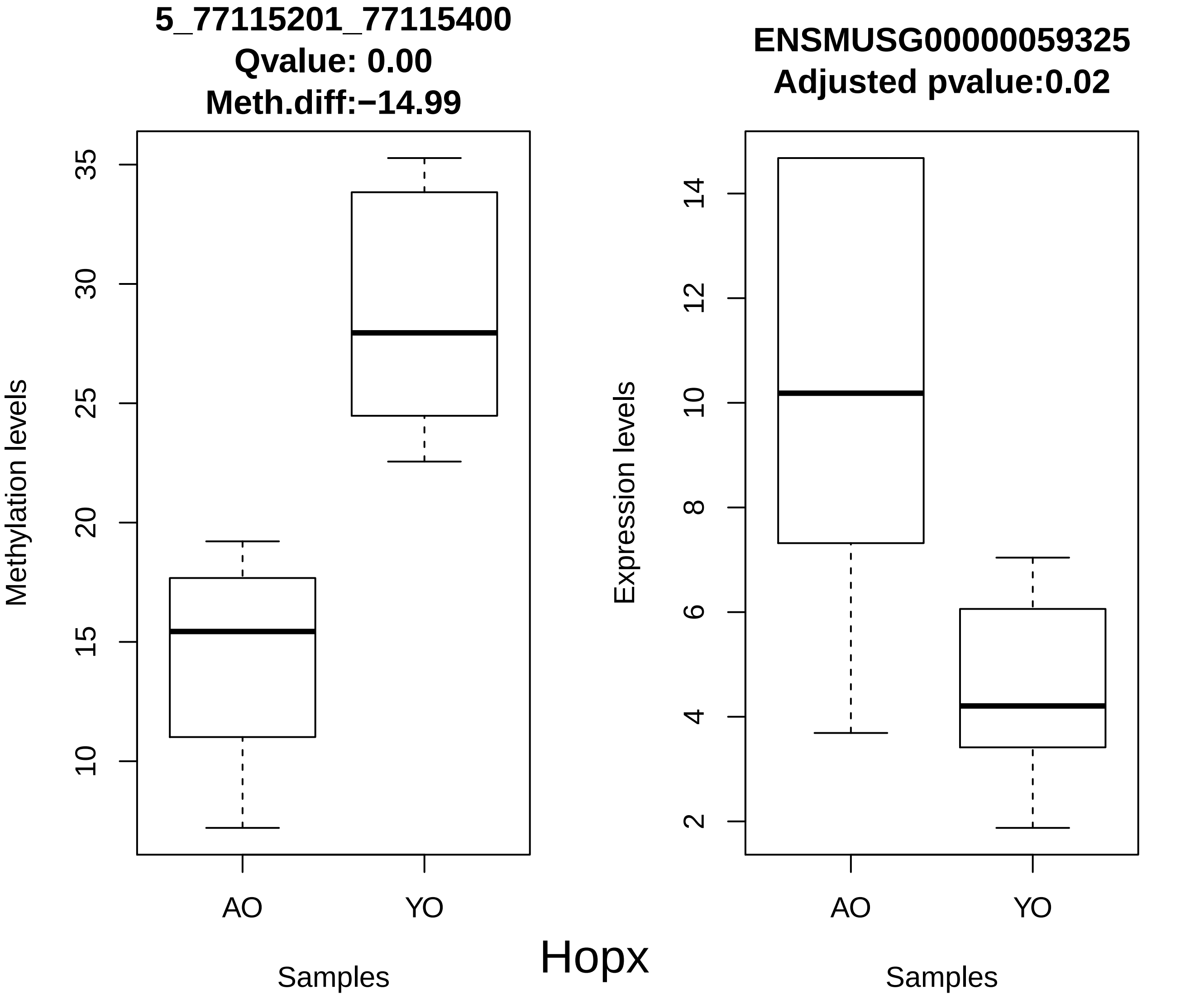
5_77115201_77115400
Qvalue: 0.00
Meth.diff:−14.99
Methylation levels
35
30
25
20
15
10
AO
YO
Samples
ENSMUSG00000059325
Adjusted pvalue:0.02
Expression levels
14
12
10
8
6
4
2
AO
YO
Samples
Hopx

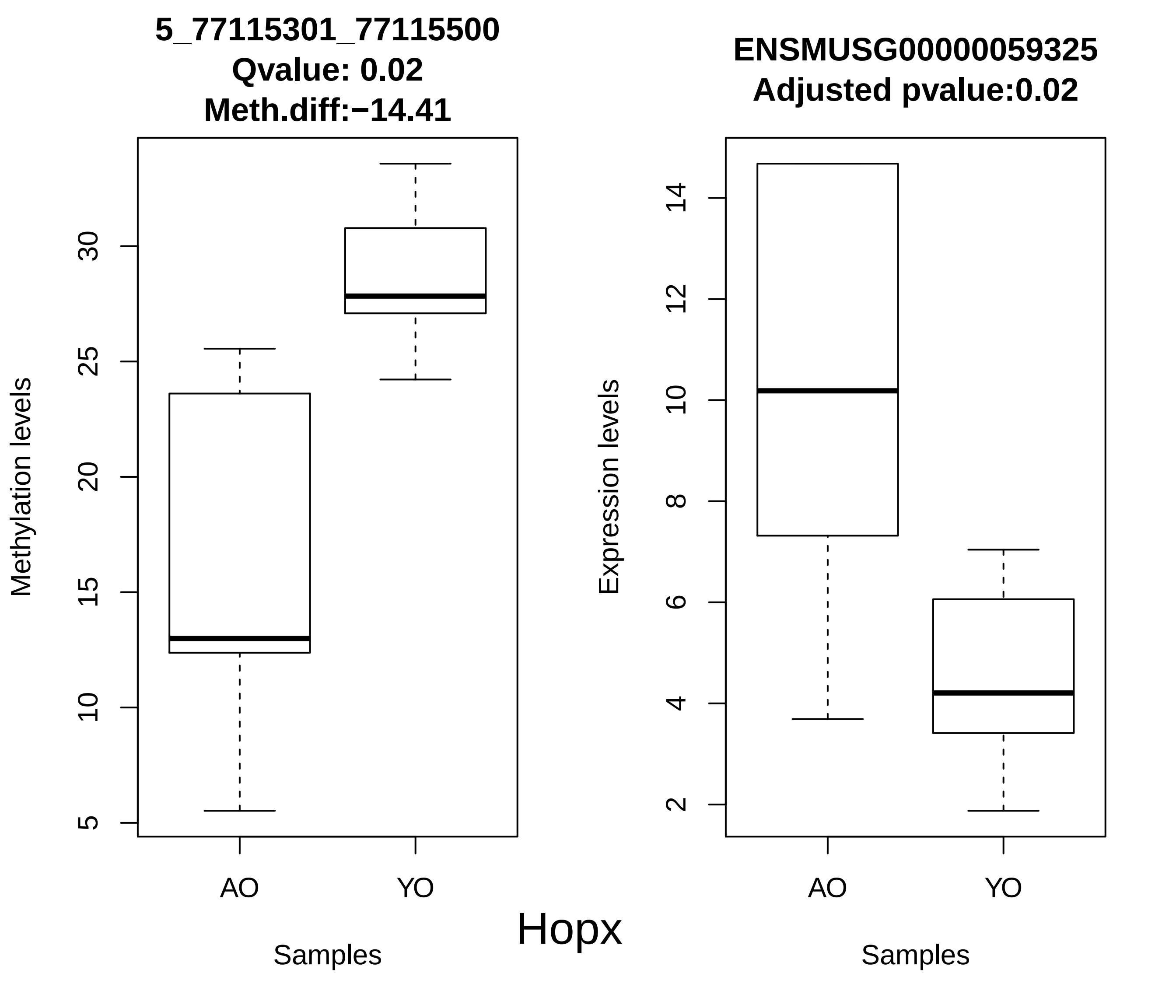
5_77115301_77115500
Qvalue: 0.02
Meth.diff:−14.41
Methylation levels
5
10
15
20
25
30
AO
YO
Samples
ENSMUSG00000059325
Adjusted pvalue:0.02
Expression levels
2
4
6
8
10
12
14
AO
YO
Samples
Hopx

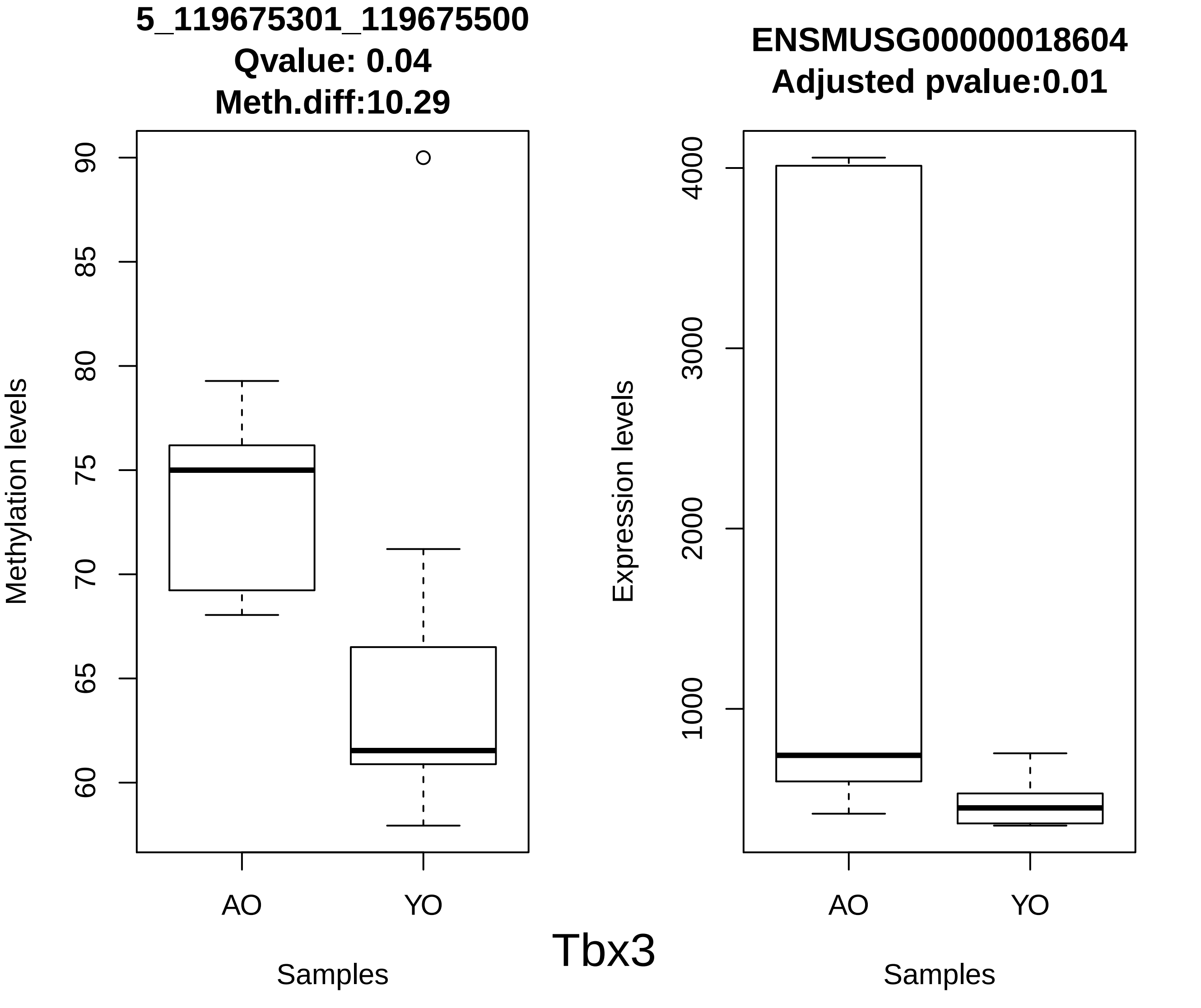
5_119675301_119675500
Qvalue: 0.04
Meth.diff:10.29
Methylation levels
90
85
80
75
70
65
60
AO
YO
Samples
ENSMUSG00000018604
Adjusted pvalue:0.01
Expression levels
4000
3000
2000
1000
AO
YO
Samples
Tbx3

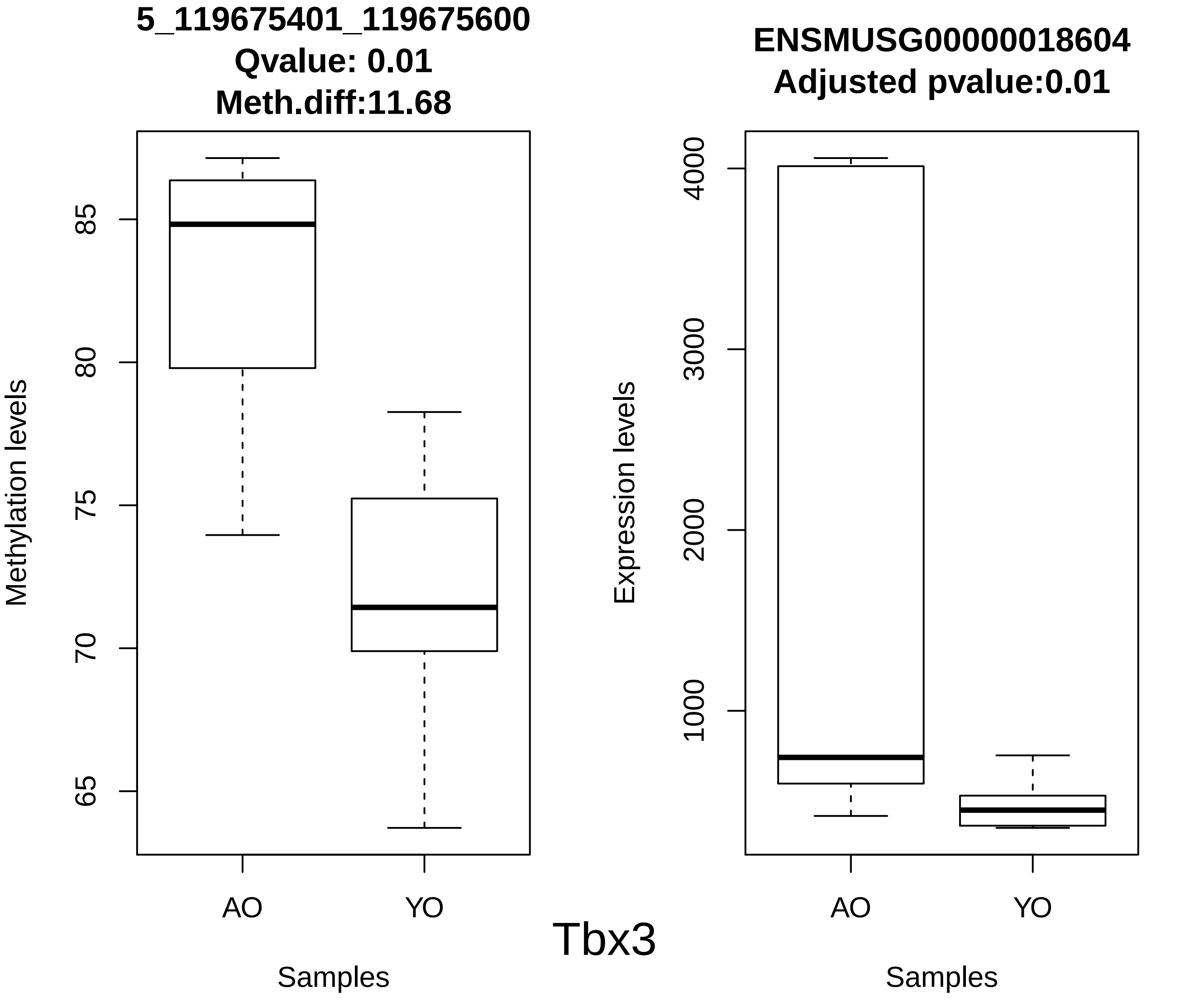
5_119675401_119675600
Qvalue: 0.01
Meth.diff:11.68
Methylation levels
85
80
75
70
65
AO
YO
Samples
ENSMUSG00000018604
Adjusted pvalue:0.01
Expression levels
4000
3000
2000
1000
AO
YO
Samples

Tbx3

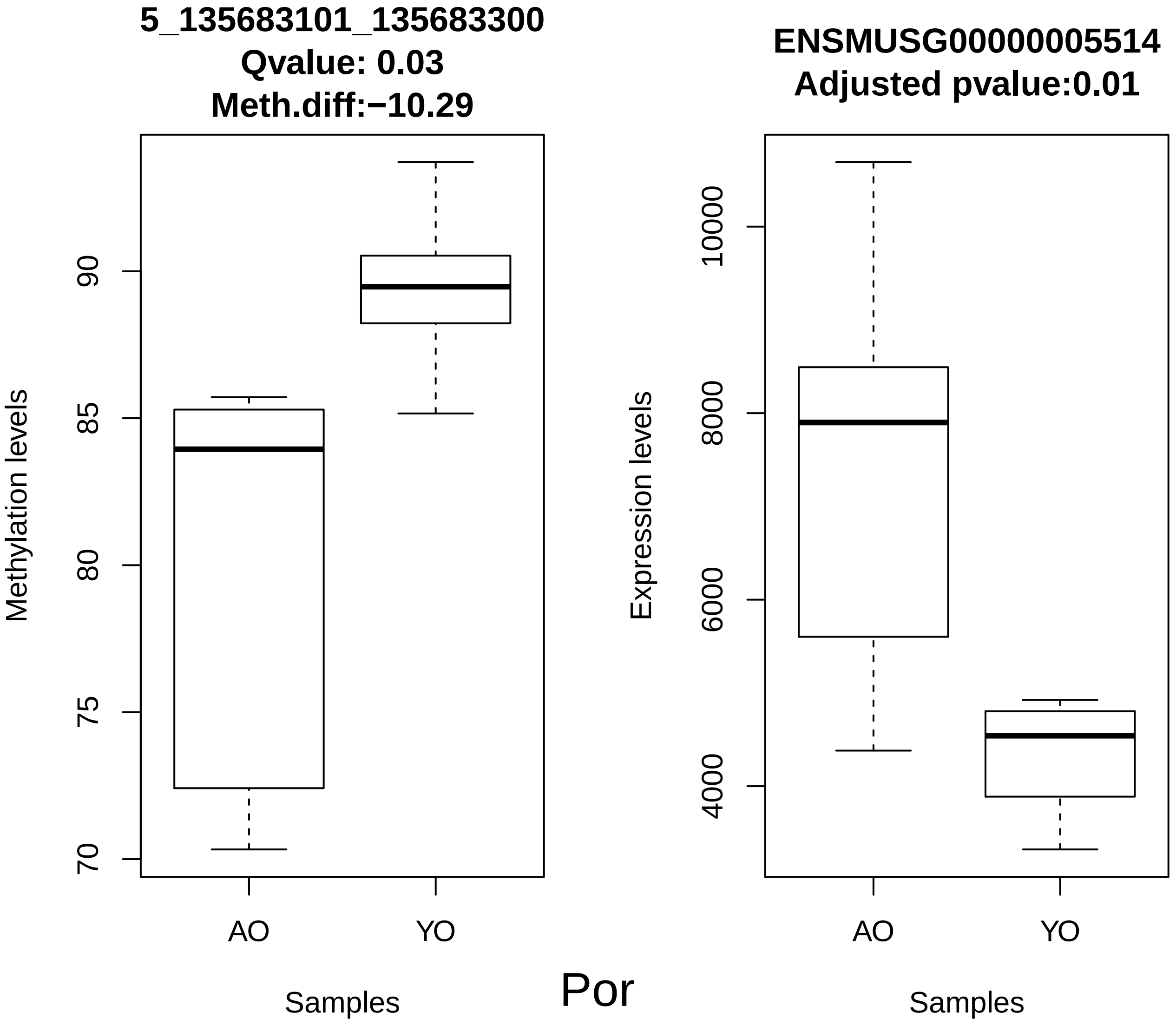
5_135683101_135683300
Qvalue: 0.03
Meth.diff:−10.29
Methylation levels
70
75
80
85
90
AO
YO
Samples
ENSMUSG00000005514
Adjusted pvalue:0.01
Expression levels
4000
6000
8000
10000
AO
YO
Samples
Por

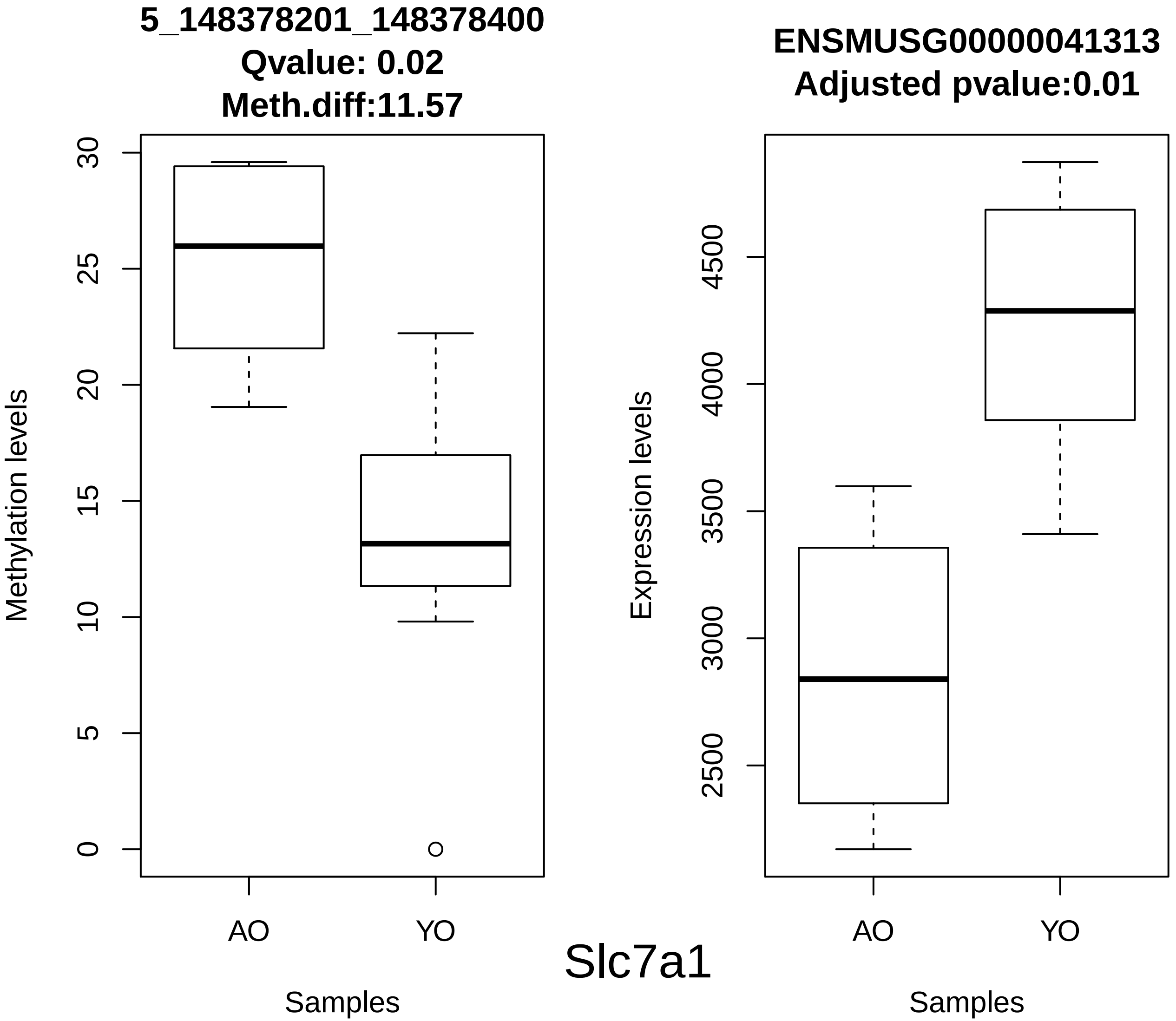
5_148378201_148378400
Qvalue: 0.02
Meth.diff:11.57
Methylation levels
0
5
10
15
20
25
30
AO
YO
Samples
ENSMUSG00000041313
Adjusted pvalue:0.01
Expression levels
2500
3000
3500
4000
4500
AO
YO
Samples

Slc7a1

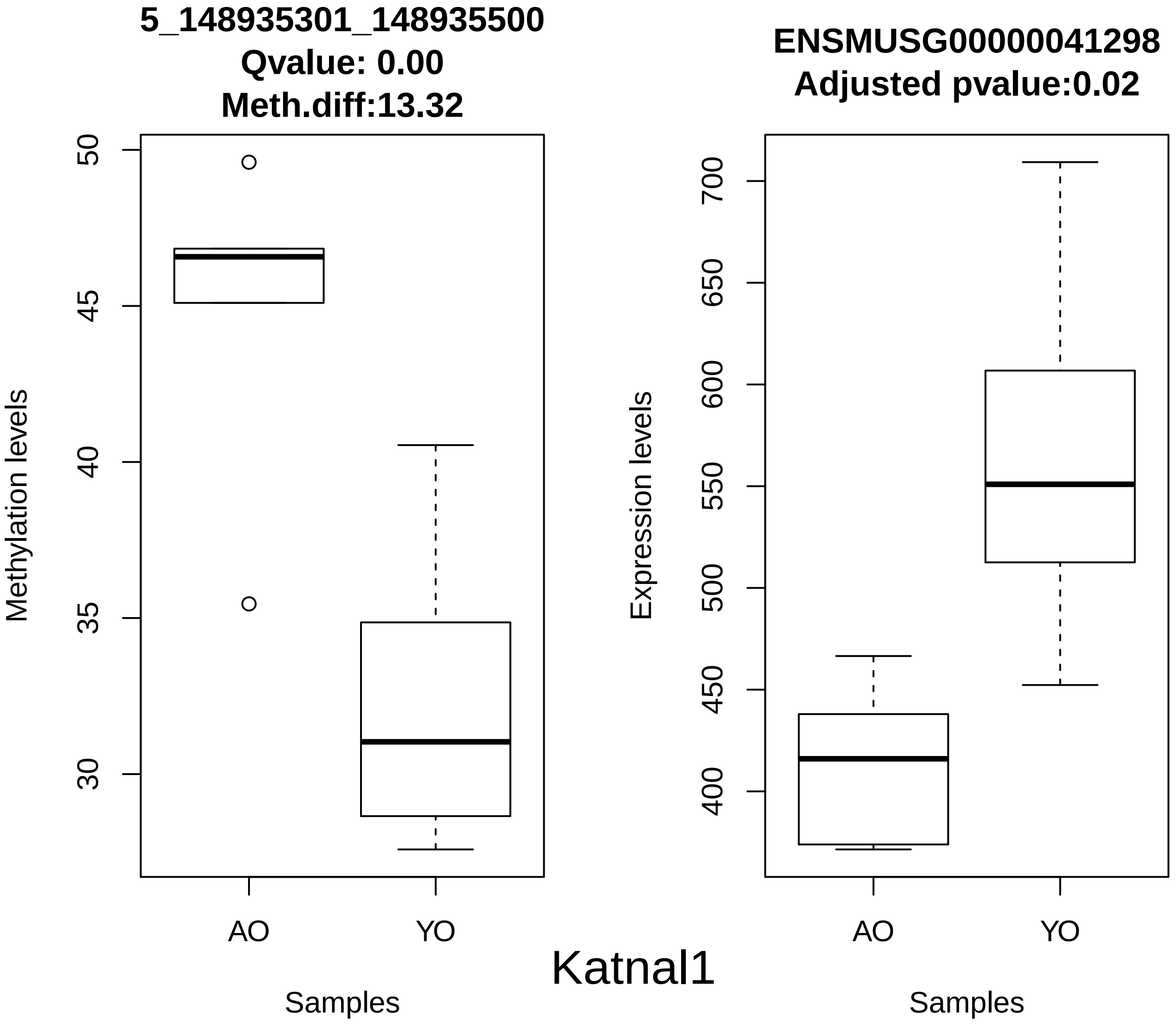

5_148935301_148935500
Qvalue: 0.00
Meth.diff:13.32
Methylation levels
50
45
40
35
30
AO
YO
Samples
ENSMUSG00000041298
Adjusted pvalue:0.02
Expression levels
700
650
600
550
500
450
400
AO
YO
Samples


Katnal1

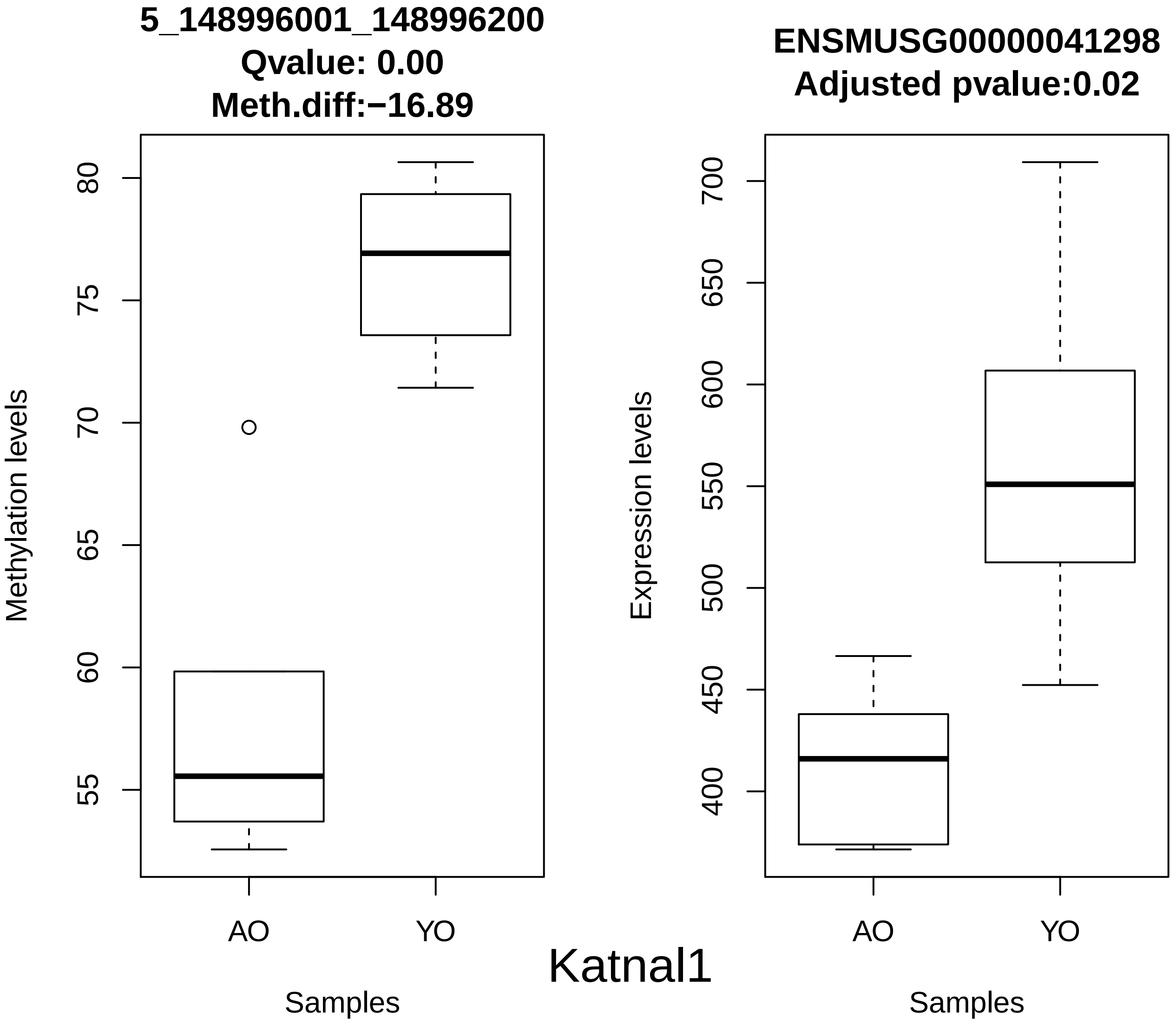
5_148996001_148996200
Qvalue: 0.00
Meth.diff:−16.89
Methylation levels
80
75
70
65
60
55
AO
YO
Samples
ENSMUSG00000041298
Adjusted pvalue:0.02
Expression levels
700
650
600
550
500
450
400
AO
YO
Samples
Katnal1

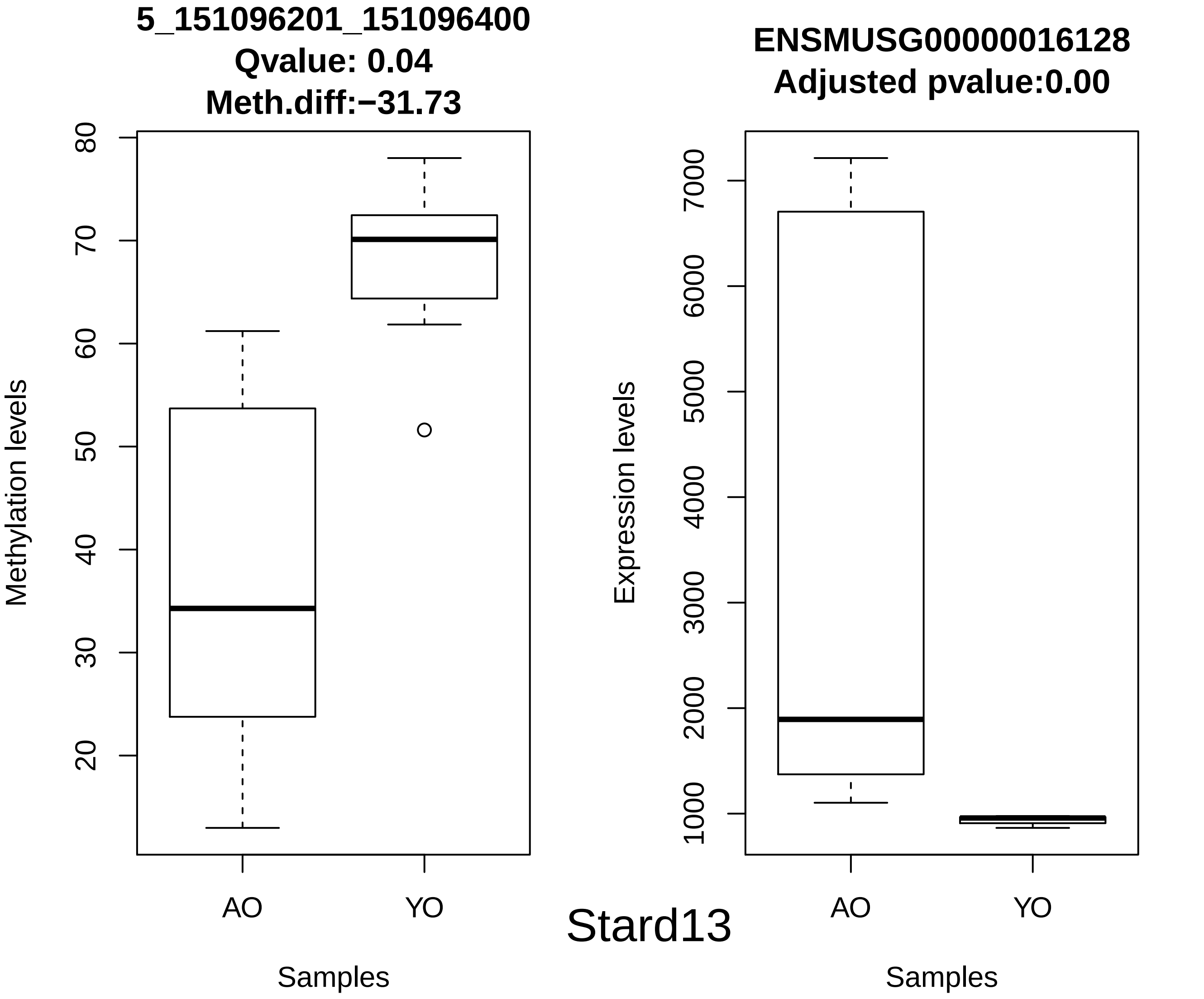
5_151096201_151096400
Qvalue: 0.04
Meth.diff:−31.73
Methylation levels
80
70
60
50
40
30
20
AO
YO
Samples
ENSMUSG00000016128
Adjusted pvalue:0.00
Expression levels
7000
6000
5000
4000
3000
2000
1000
AO
YO
Samples
Stard13

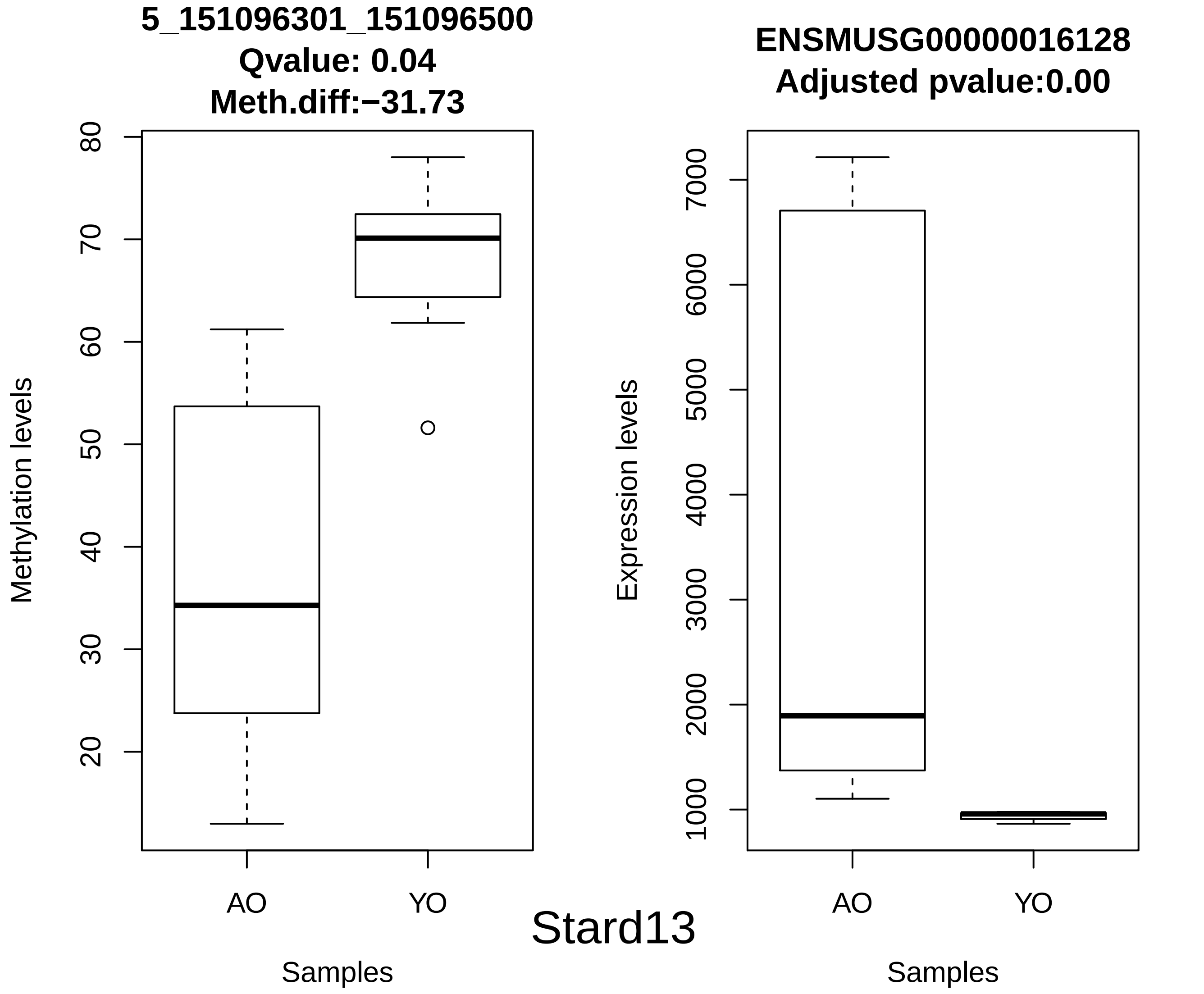
5_151096301_151096500
Qvalue: 0.04
Meth.diff:−31.73
Methylation levels
80
70
60
50
40
30
20
AO
YO
Samples
Stard13
ENSMUSG00000016128
Adjusted pvalue:0.00
Expression levels
7000
6000
5000
4000
3000
2000
1000
AO
YO
Samples

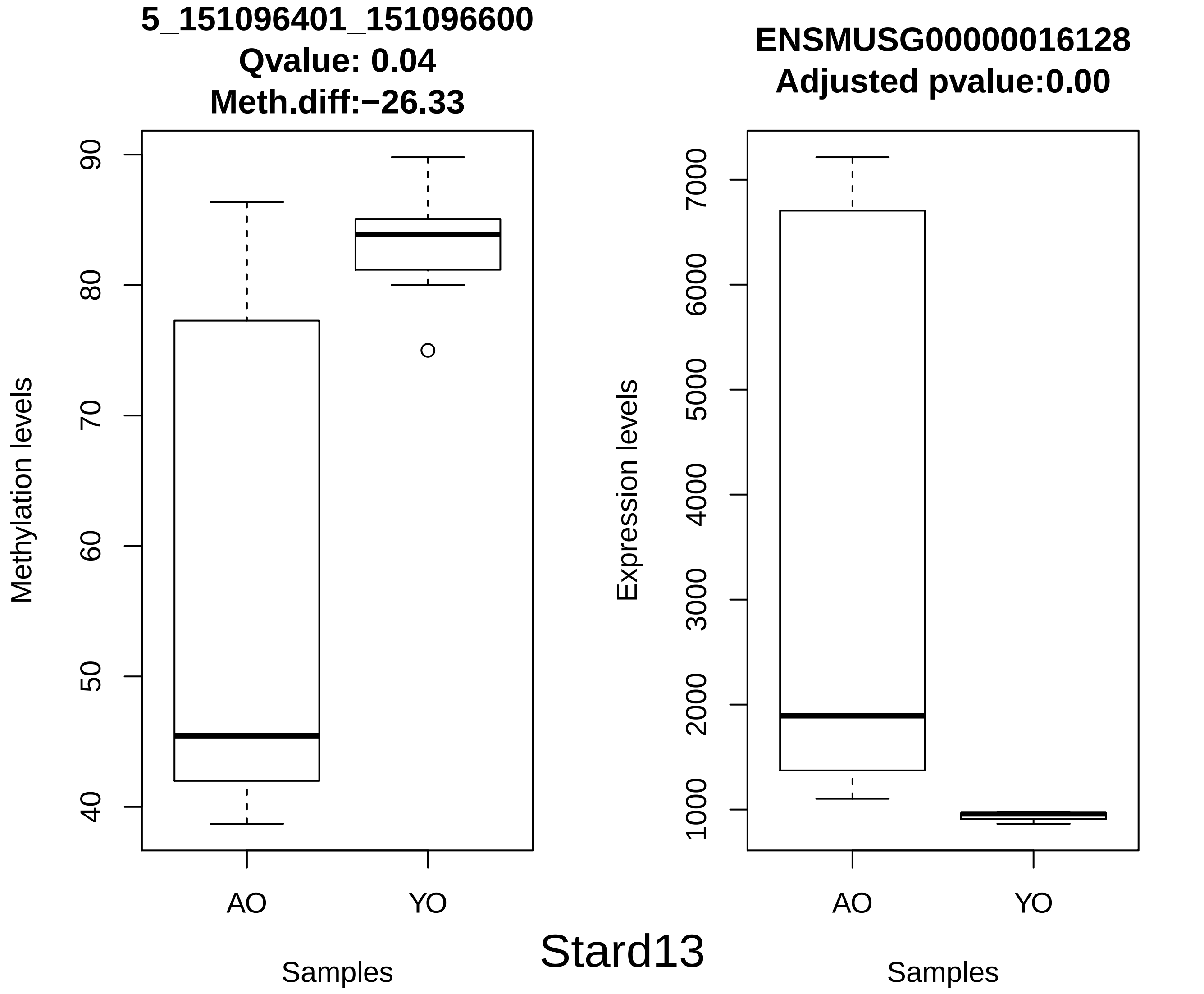
5_151096401_151096600
Qvalue: 0.04
Meth.diff:−26.33
Methylation levels
90
80
70
60
50
40
AO
YO
Samples
ENSMUSG00000016128
Adjusted pvalue:0.00
Expression levels
7000
6000
5000
4000
3000
2000
1000
AO
YO
Samples
Stard13

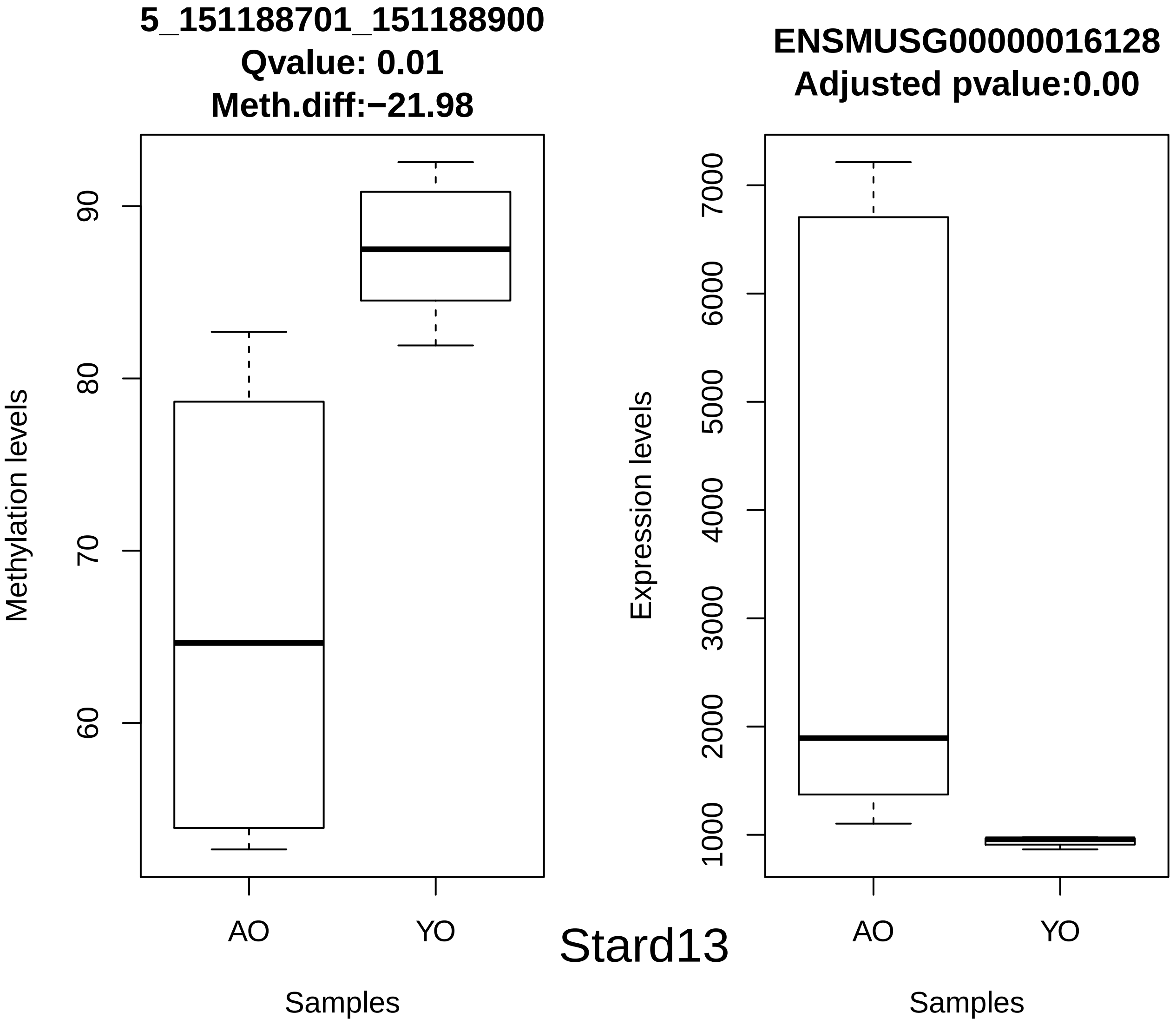
5_151188701_151188900
Qvalue: 0.01
Meth.diff:−21.98
Methylation levels
60
70
80
90
AO
YO
Samples
Stard13
ENSMUSG00000016128
Adjusted pvalue:0.00
Expression levels
1000
2000
3000
4000
5000
6000
7000
AO
YO
Samples

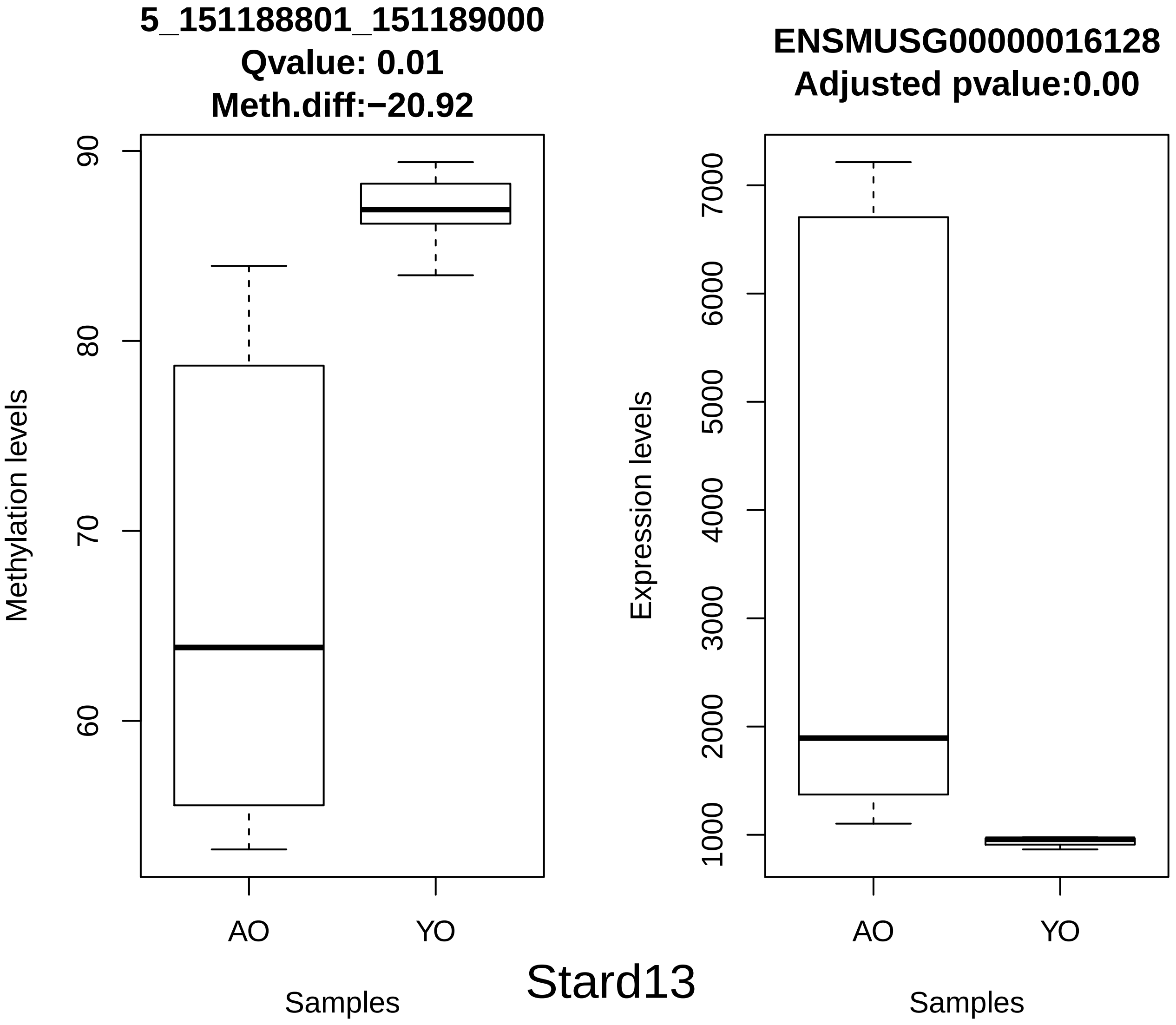
5_151188801_151189000
Qvalue: 0.01
Meth.diff:−20.92
Methylation levels
90
80
70
60
AO
YO
Samples
ENSMUSG00000016128
Adjusted pvalue:0.00
Expression levels
7000
6000
5000
4000
3000
2000
1000
AO
YO
Samples

Stard13

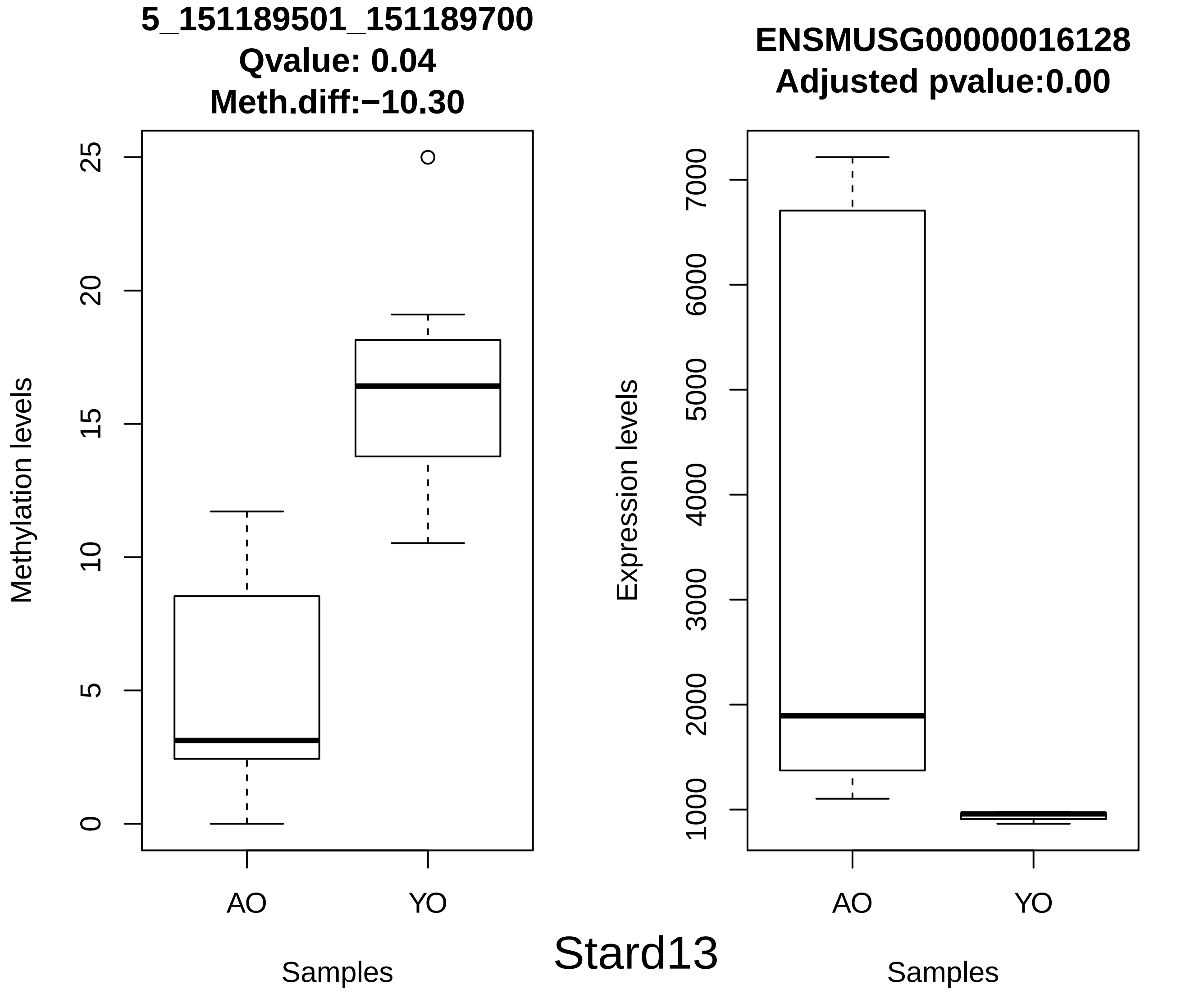
5_151189501_151189700
Qvalue: 0.04
Meth.diff:−10.30
Methylation levels
0
5
10
15
20
25
AO
YO
Samples
ENSMUSG00000016128
Adjusted pvalue:0.00
Expression levels
1000
2000
3000
4000
5000
6000
7000
AO
YO
Samples
Stard13

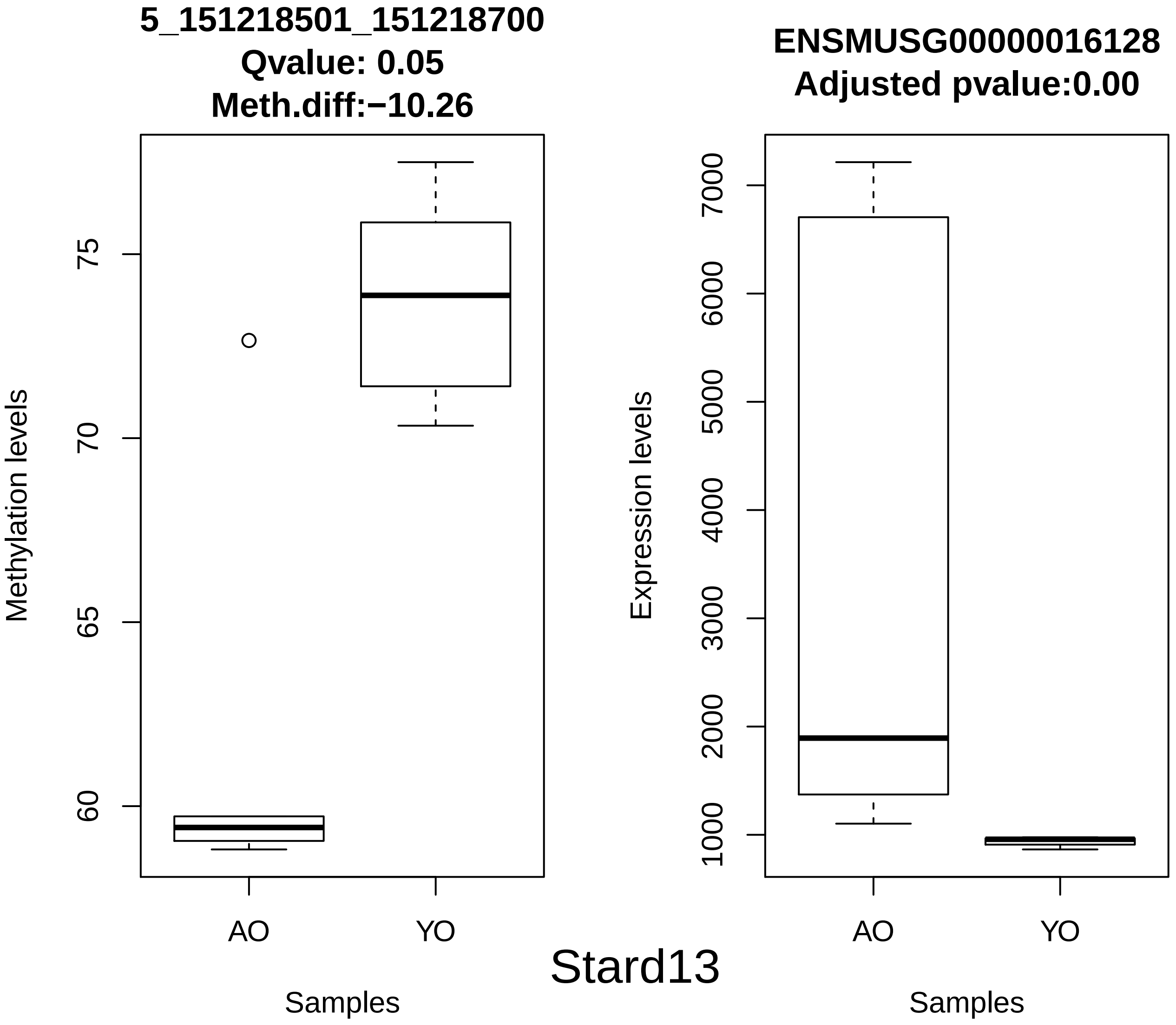
5_151218501_151218700
Qvalue: 0.05
Meth.diff:−10.26
Methylation levels
75
70
65
60
AO
YO
Samples
ENSMUSG00000016128
Adjusted pvalue:0.00
Expression levels
7000
6000
5000
4000
3000
2000
1000
AO
YO
Samples
Stard13

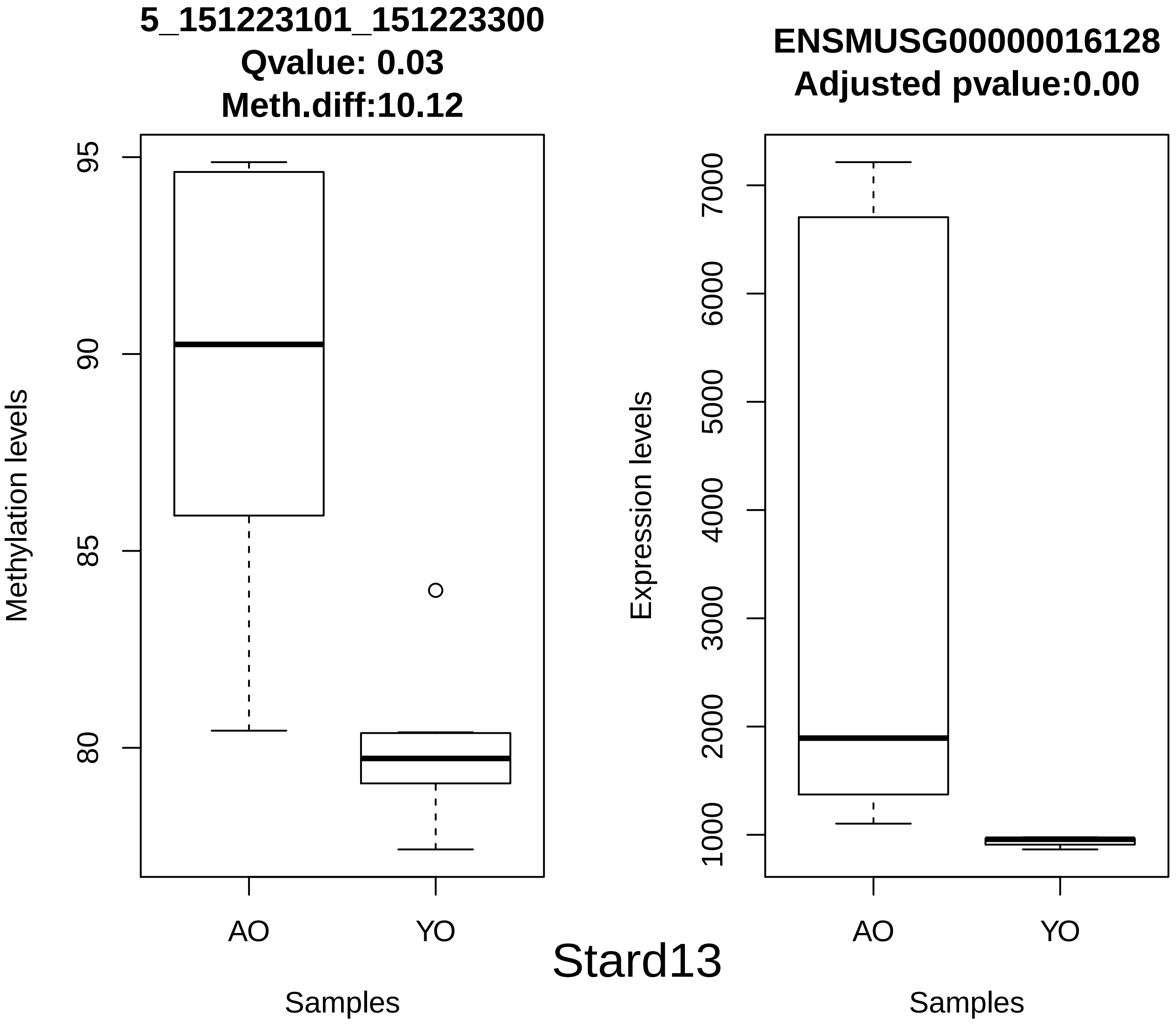
5_151223101_151223300
Qvalue: 0.03
Meth.diff:10.12
Methylation levels
95
90
85
80
AO
YO
Samples
ENSMUSG00000016128
Adjusted pvalue:0.00
Expression levels
7000
6000
5000
4000
3000
2000
1000
AO
YO
Samples

Stard13

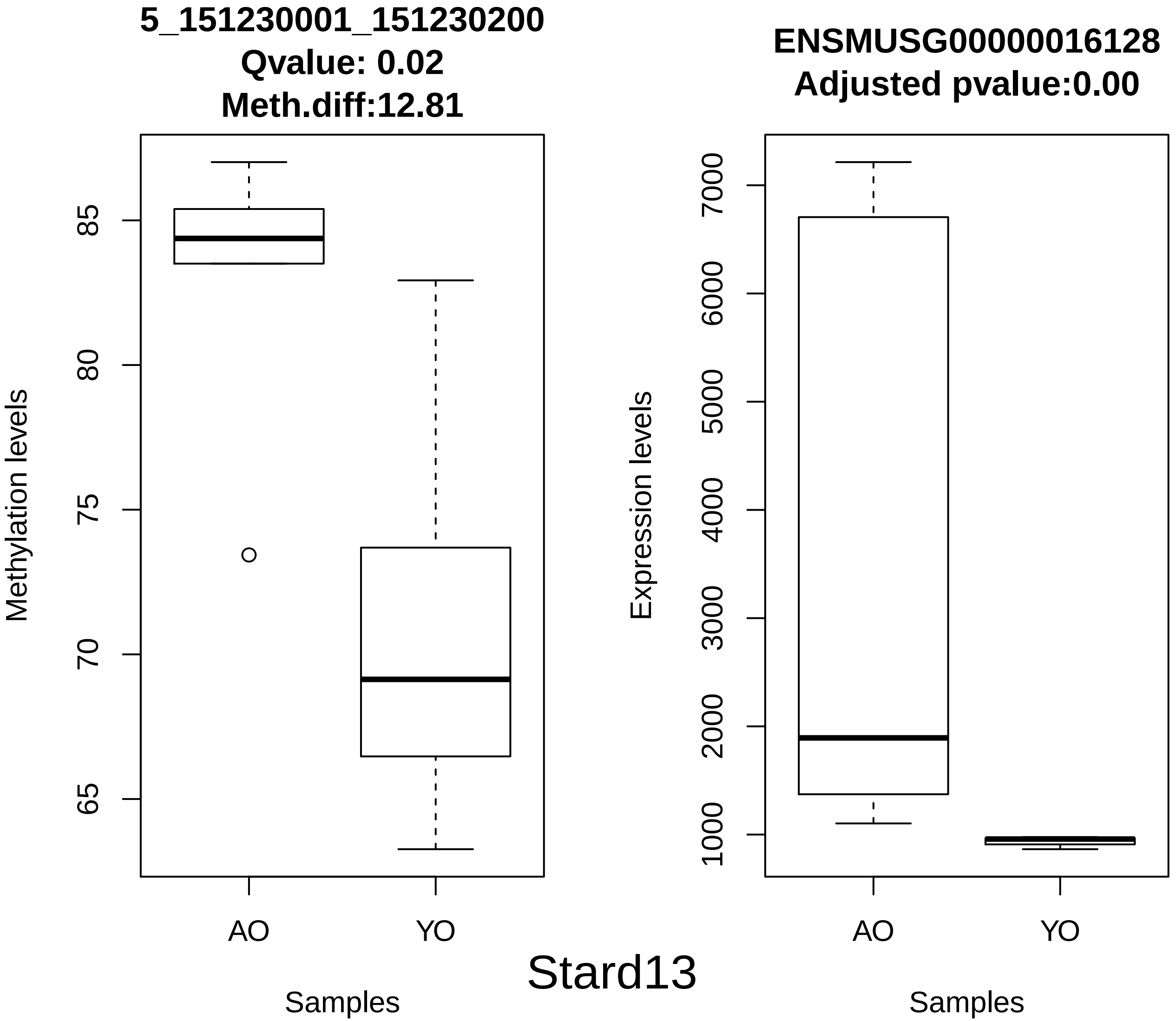
5_151230001_151230200
Qvalue: 0.02
Meth.diff:12.81
Methylation levels
85
80
75
70
65
AO
YO
Samples
ENSMUSG00000016128
Adjusted pvalue:0.00
Expression levels
7000
6000
5000
4000
3000
2000
1000
AO
YO
Samples
Stard13

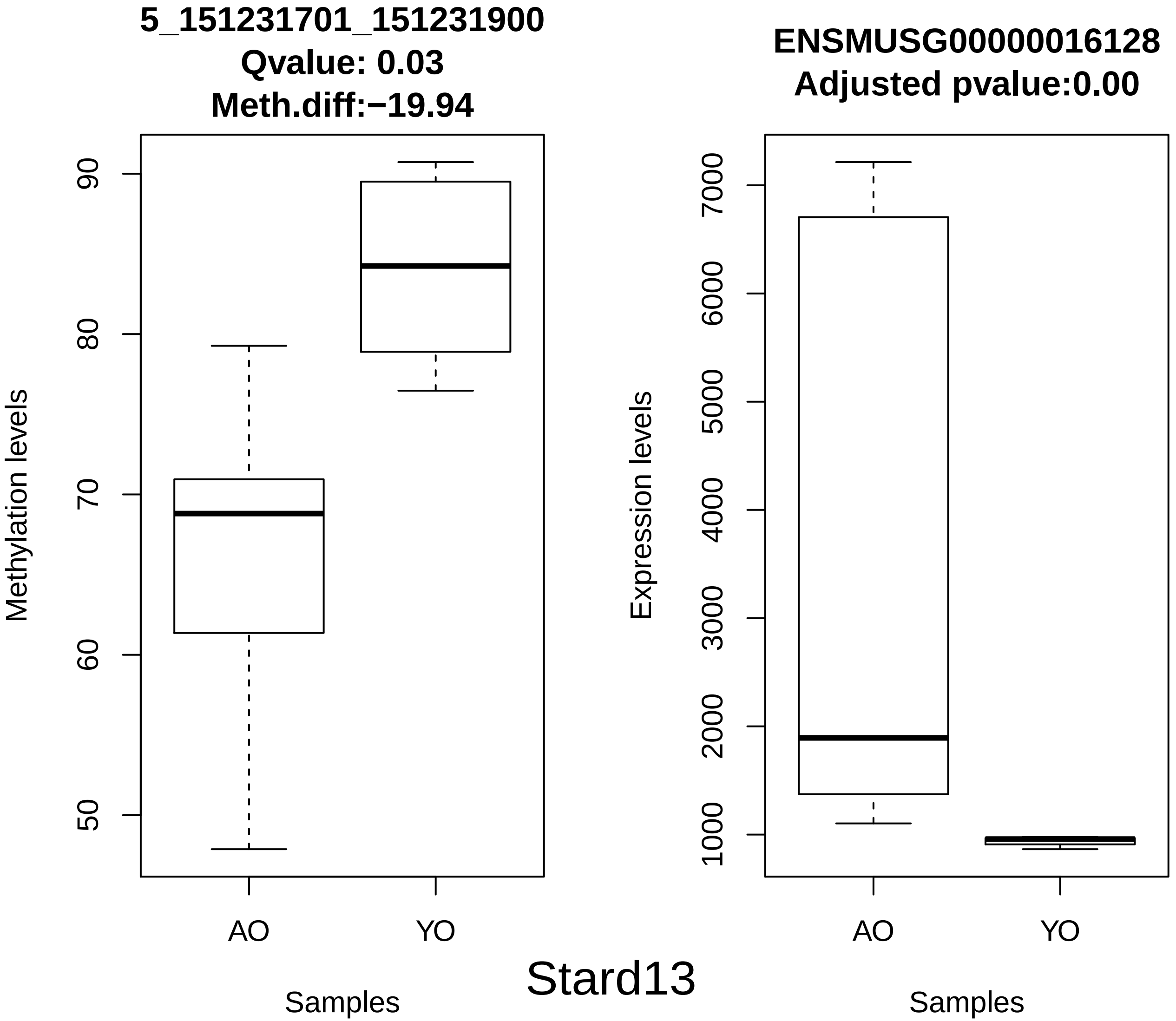
5_151231701_151231900
Qvalue: 0.03
Meth.diff:−19.94
Methylation levels
90
80
70
60
50
AO
YO
Samples
ENSMUSG00000016128
Adjusted pvalue:0.00
Expression levels
7000
6000
5000
4000
3000
2000
1000
AO
YO
Samples
Stard13

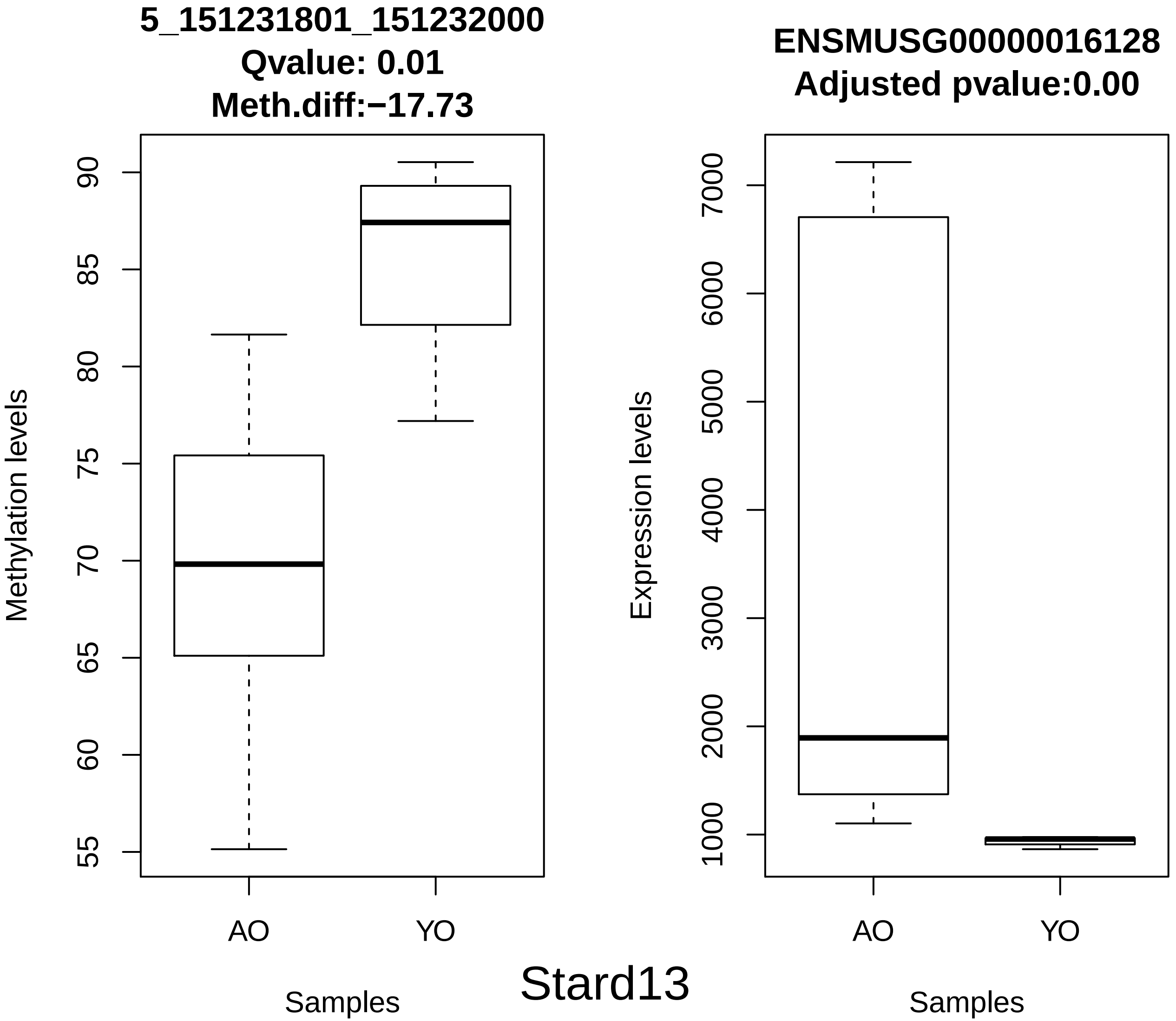
5_151231801_151232000
Qvalue: 0.01
Meth.diff:−17.73
Methylation levels
90
85
80
75
70
65
60
55
AO
YO
Samples
ENSMUSG00000016128
Adjusted pvalue:0.00
Expression levels
7000
6000
5000
4000
3000
2000
1000
AO
YO
Samples
Stard13

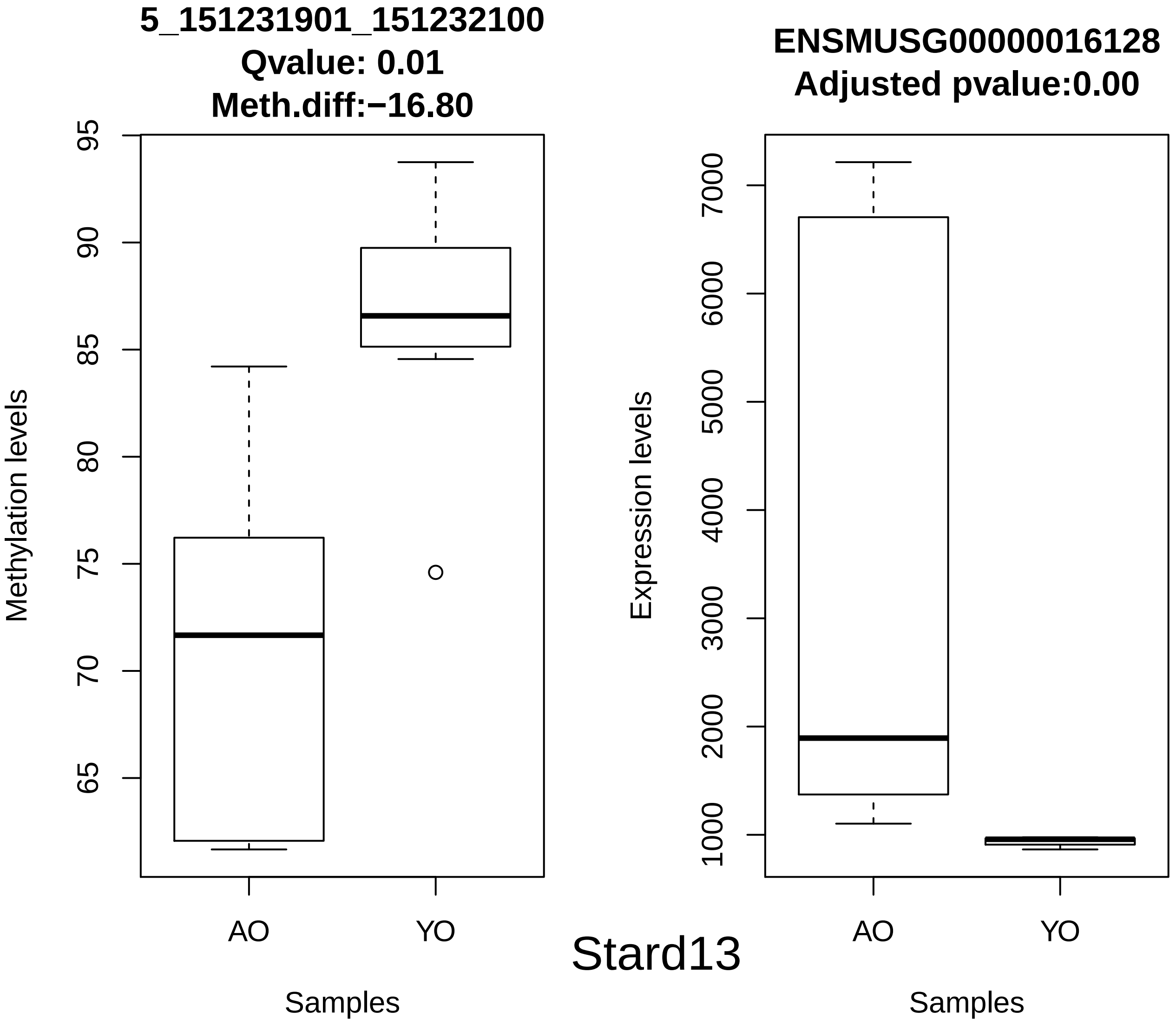
5_151231901_151232100
Qvalue: 0.01
Meth.diff:−16.80
Methylation levels
95
90
85
80
75
70
65
AO
YO
Samples
ENSMUSG00000016128
Adjusted pvalue:0.00
Expression levels
7000
6000
5000
4000
3000
2000
1000
AO
YO
Samples
Stard13

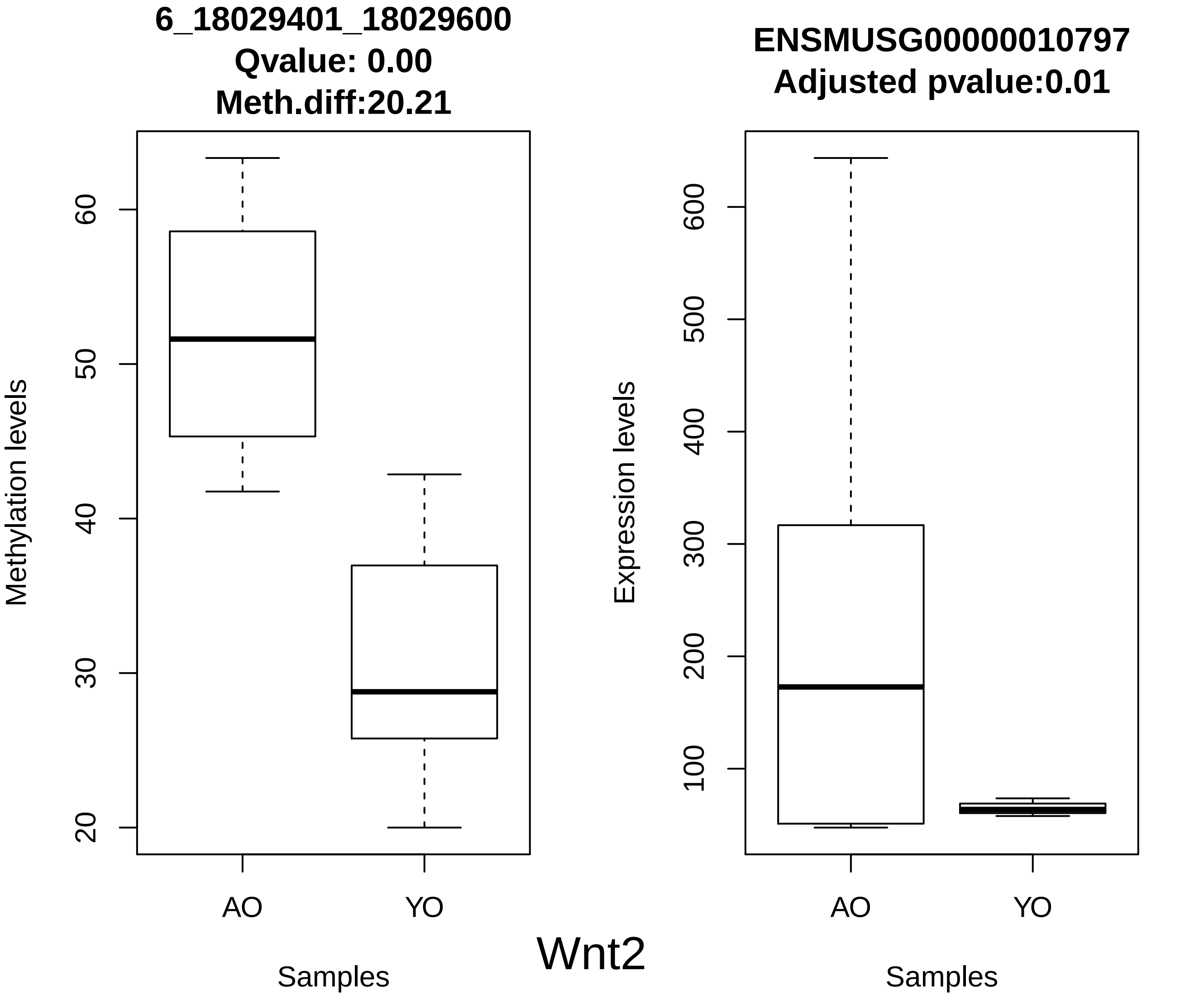
6_18029401_18029600
Qvalue: 0.00
Meth.diff:20.21
Methylation levels
20
30
40
50
60
AO
YO
Samples
ENSMUSG00000010797
Adjusted pvalue:0.01
Expression levels
100
200
300
400
500
600
AO
YO
Samples
Wnt2

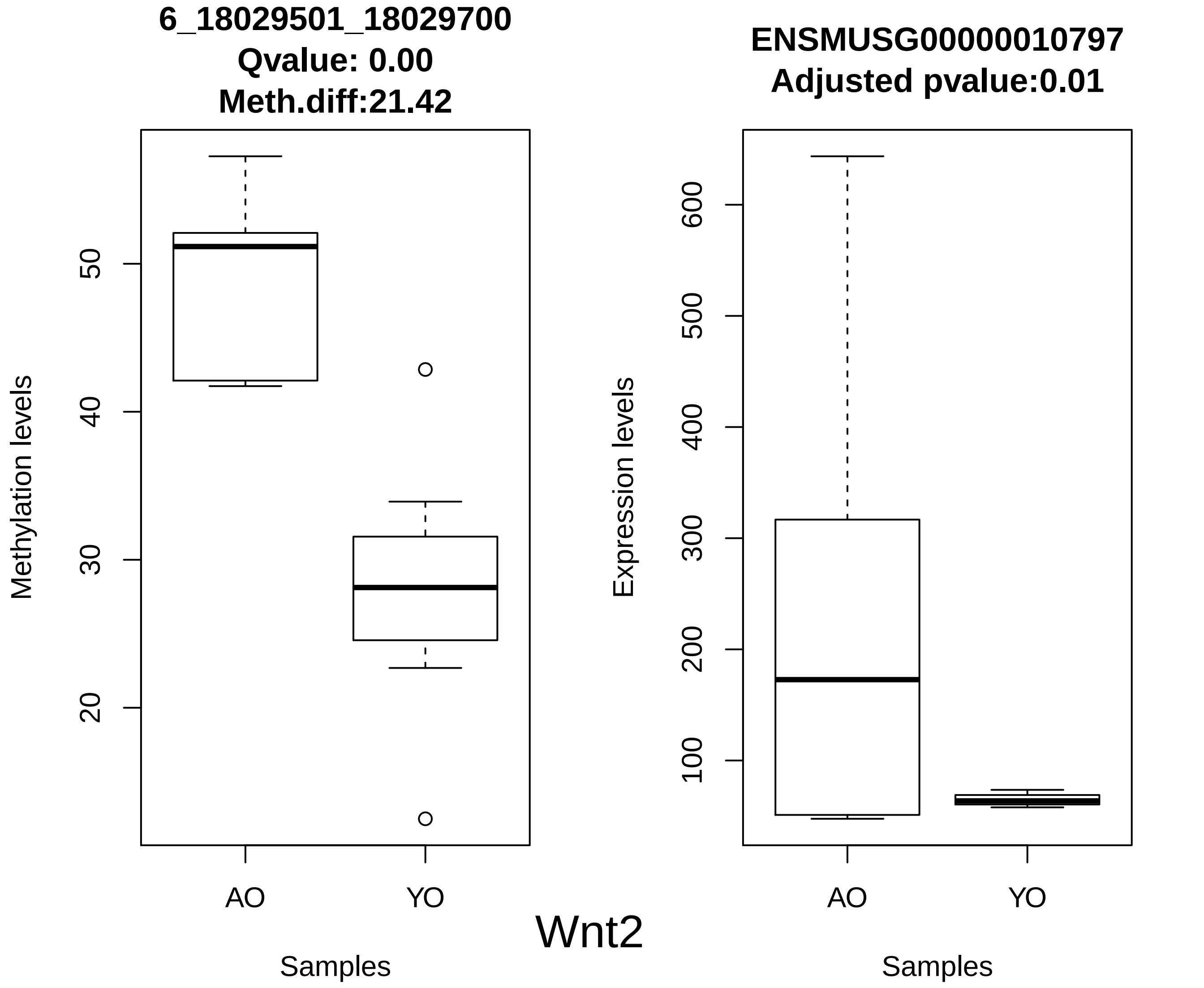
6_18029501_18029700
Qvalue: 0.00
Meth.diff:21.42
Methylation levels
20
30
40
50
AO
YO
Samples
ENSMUSG00000010797
Adjusted pvalue:0.01
Expression levels
100
200
300
400
500
600
AO
YO
Samples
Wnt2

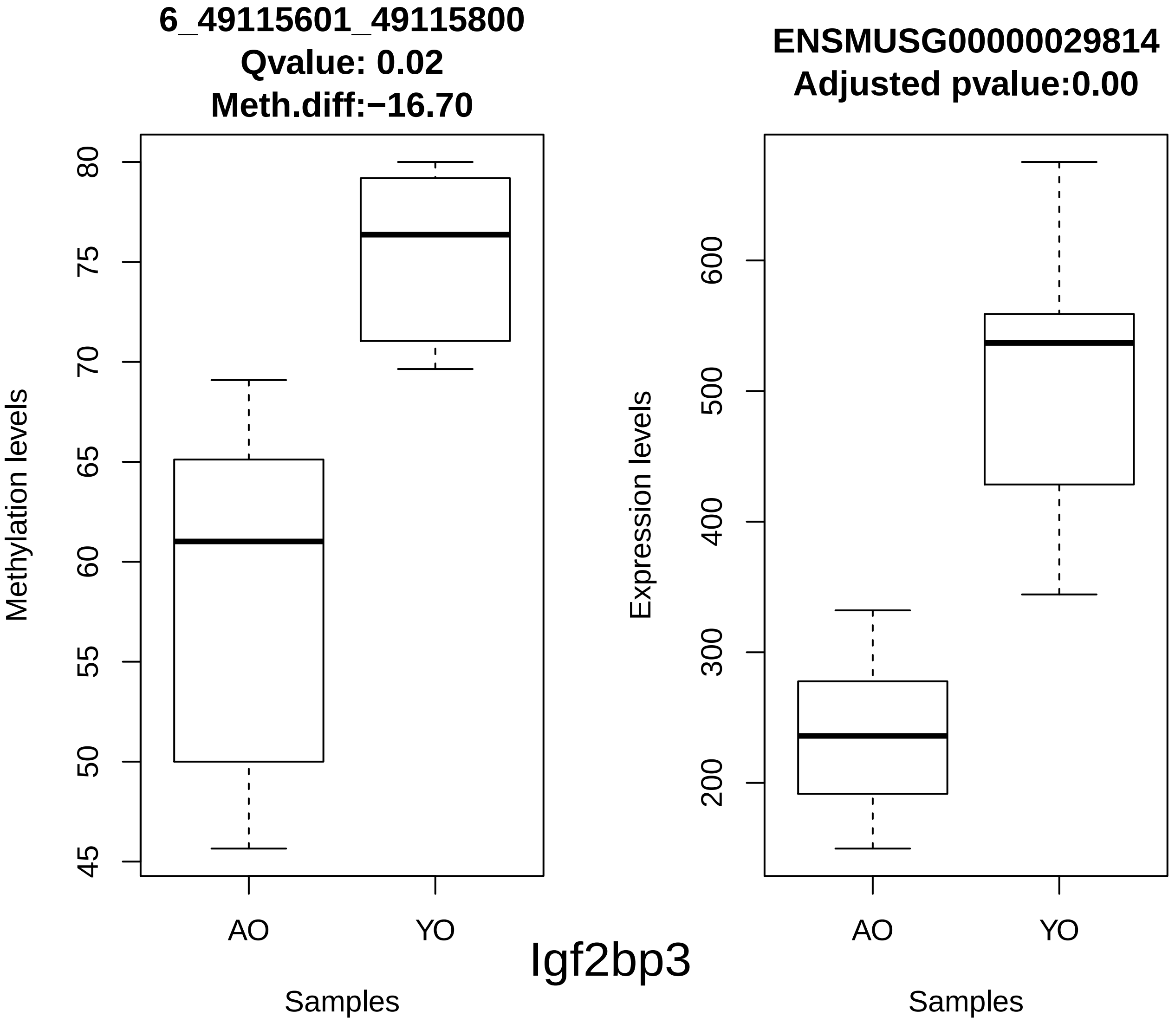
6_49115601_49115800
Qvalue: 0.02
Meth.diff:−16.70
Methylation levels
80
75
70
65
60
55
50
45
AO
YO
Samples
ENSMUSG00000029814
Adjusted pvalue:0.00
Expression levels
600
500
400
300
200
AO
YO
Samples
Igf2bp3

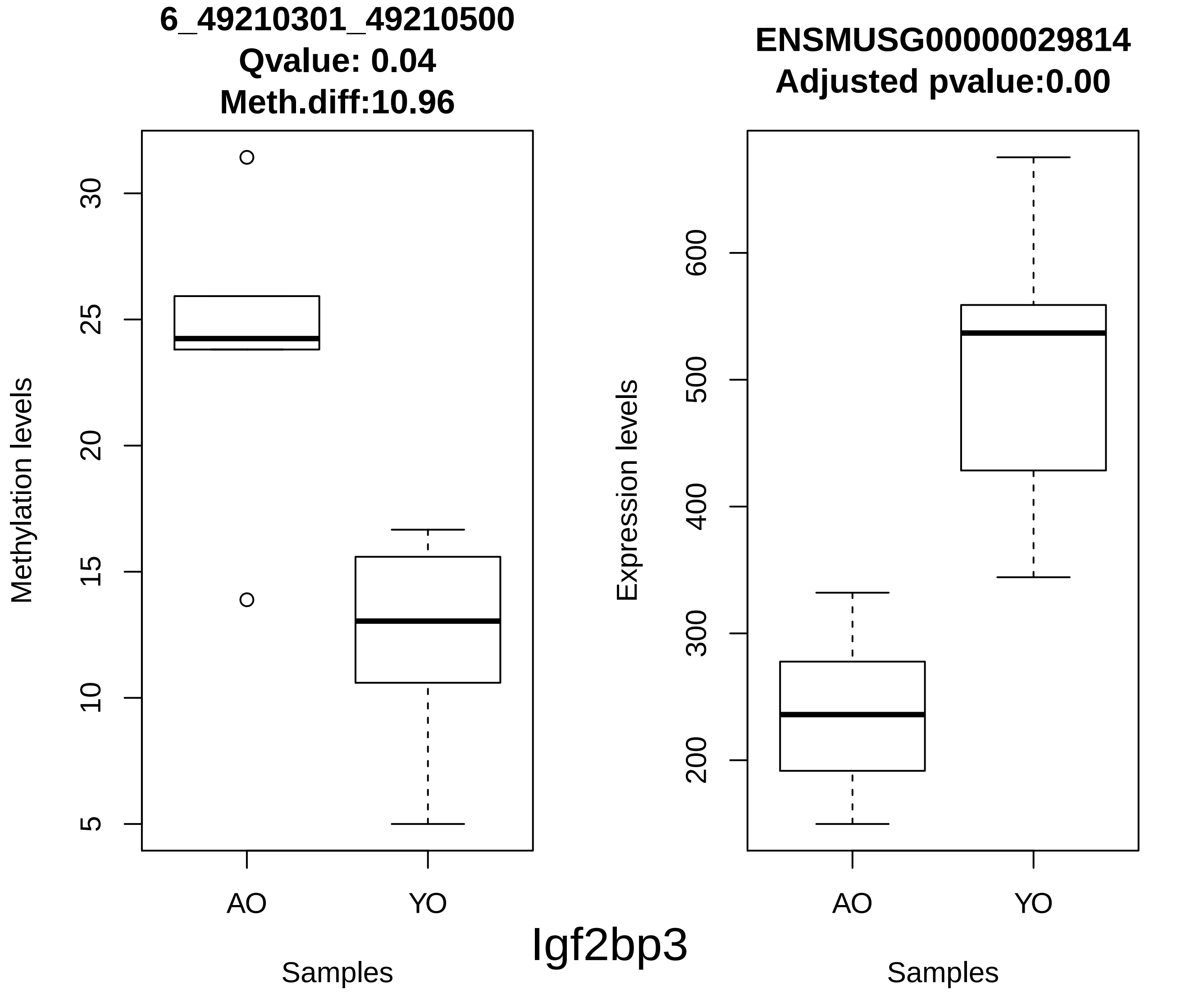
6_49210301_49210500
Qvalue: 0.04
Meth.diff:10.96
Methylation levels
30
25
20
15
10
5
AO
YO
Samples
ENSMUSG00000029814
Adjusted pvalue:0.00
Expression levels
600
500
400
300
200
AO
YO
Samples
Igf2bp3

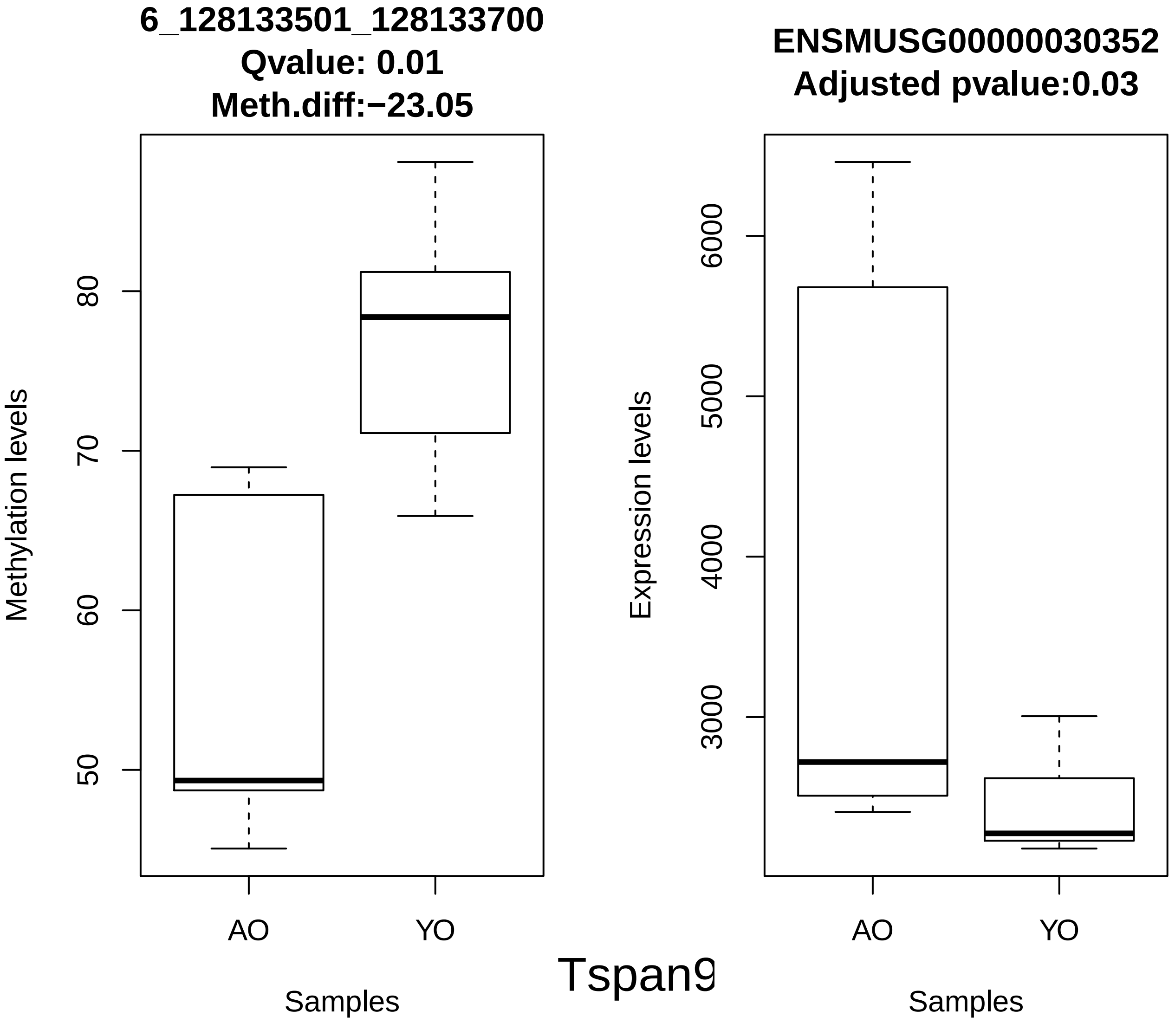
6_128133501_128133700
Qvalue: 0.01
Meth.diff:−23.05
Methylation levels
50
60
70
80
AO
YO
Samples
ENSMUSG00000030352
Adjusted pvalue:0.03
Expression levels
3000
4000
5000
6000
AO
YO
Samples
Tspan9

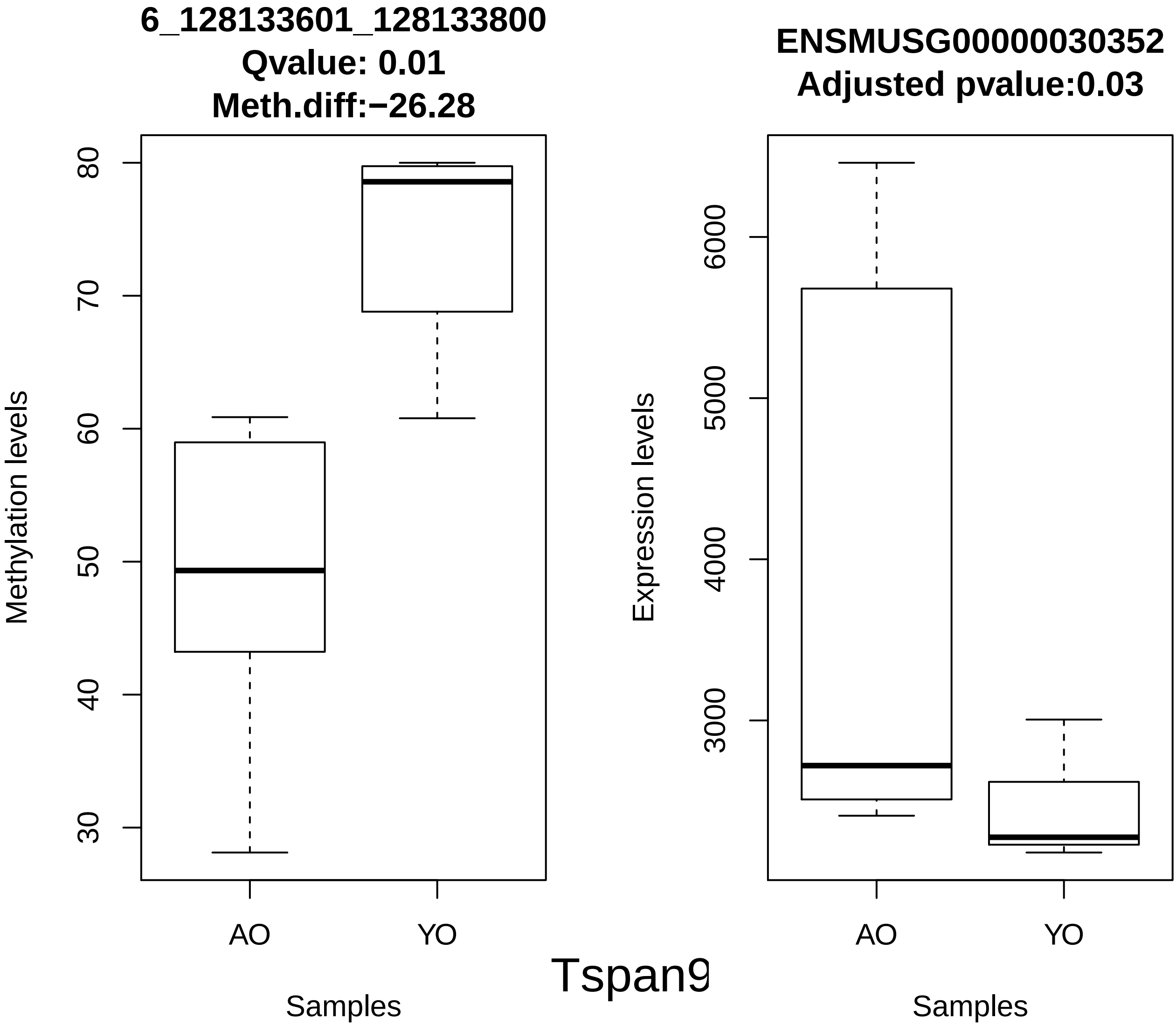
6_128133601_128133800
Qvalue: 0.01
Meth.diff:−26.28
Methylation levels
80
70
60
50
40
30
AO
YO
Samples
ENSMUSG00000030352
Adjusted pvalue:0.03
Expression levels
6000
5000
4000
3000
AO
YO
Samples
Tspan9

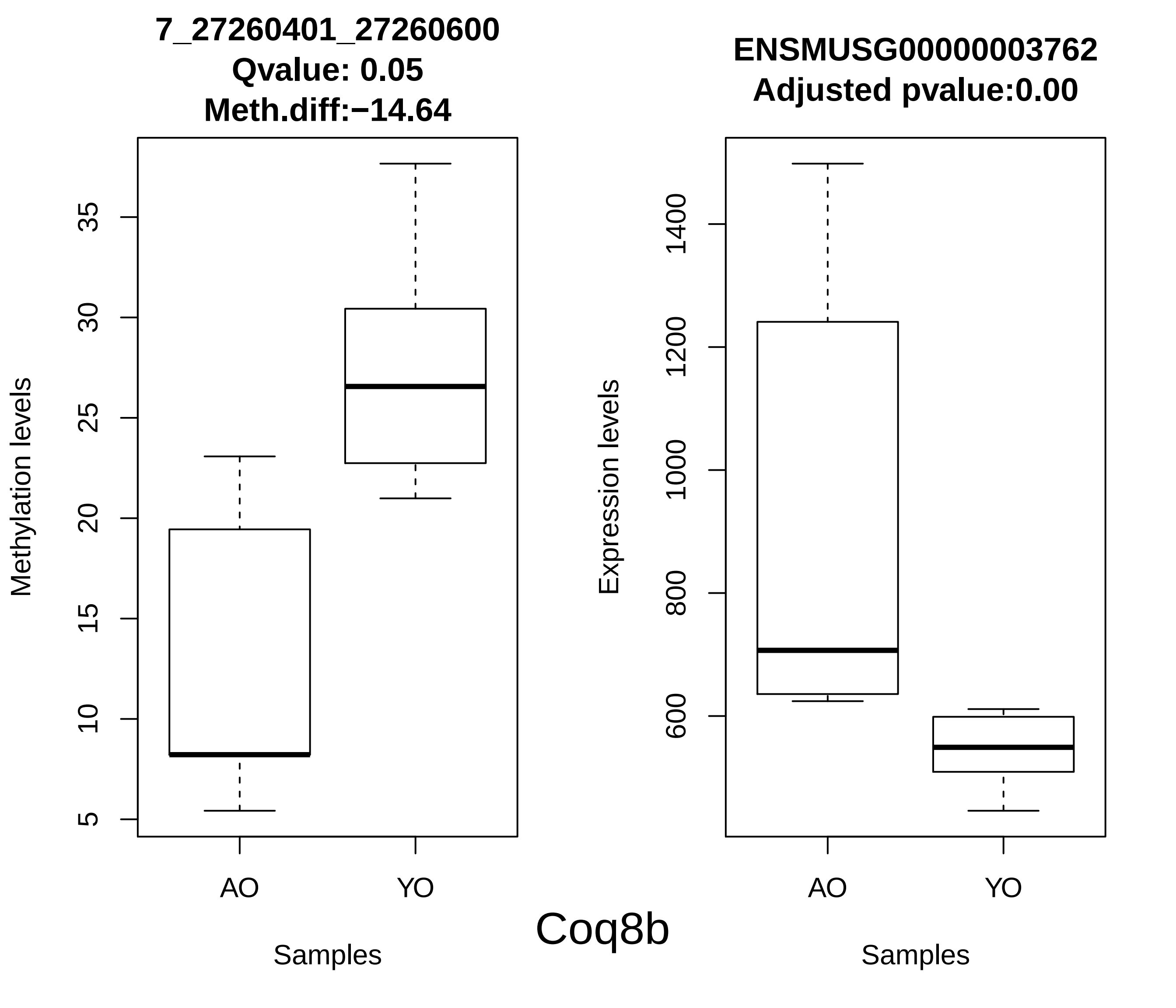
7_27260401_27260600
Qvalue: 0.05
Meth.diff:−14.64
Methylation levels
5
10
15
20
25
30
35
AO
YO
Samples
ENSMUSG00000003762
Adjusted pvalue:0.00
Expression levels
600
800
1000
1200
1400
AO
YO
Samples
Coq8b

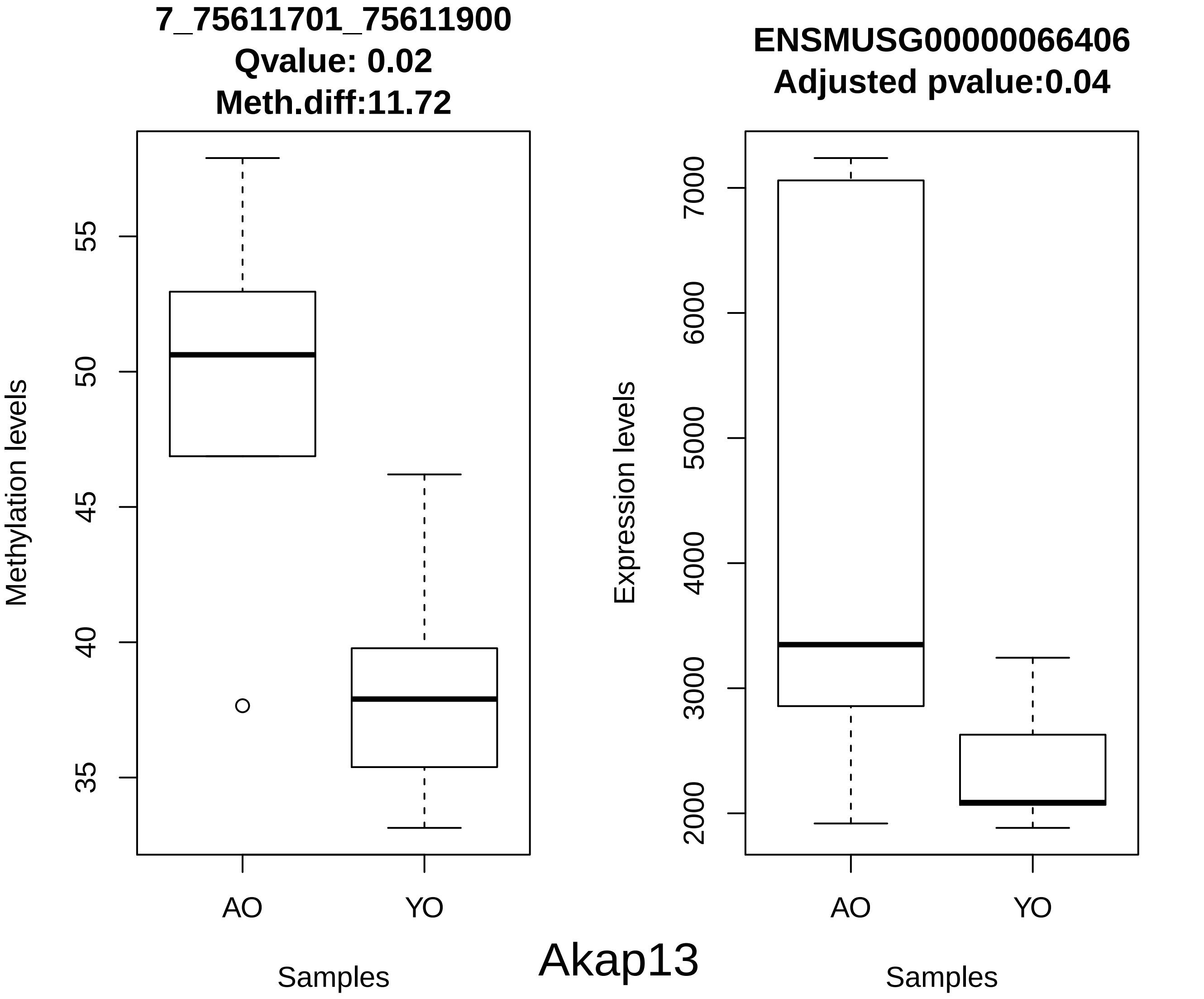
7_75611701_75611900
Qvalue: 0.02
Meth.diff:11.72
Methylation levels
55
50
45
40
35
AO
YO
Samples
ENSMUSG00000066406
Adjusted pvalue:0.04
Expression levels
7000
6000
5000
4000
3000
2000
AO
YO
Samples
Akap13

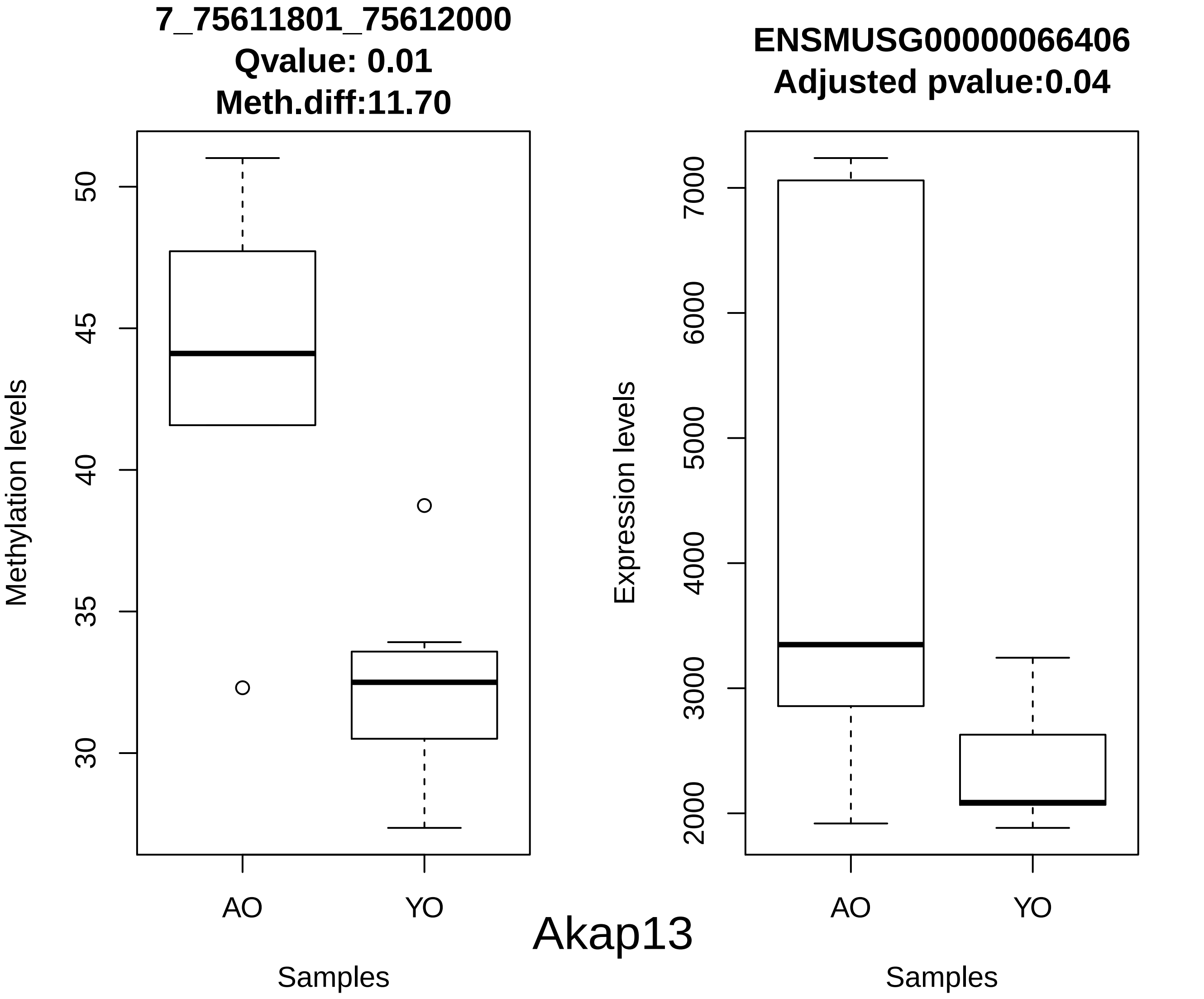
7_75611801_75612000
Qvalue: 0.01
Meth.diff:11.70
Methylation levels
50
45
40
35
30
AO
YO
Samples
ENSMUSG00000066406
Adjusted pvalue:0.04
Expression levels
7000
6000
5000
4000
3000
2000
AO
YO
Samples
Akap13

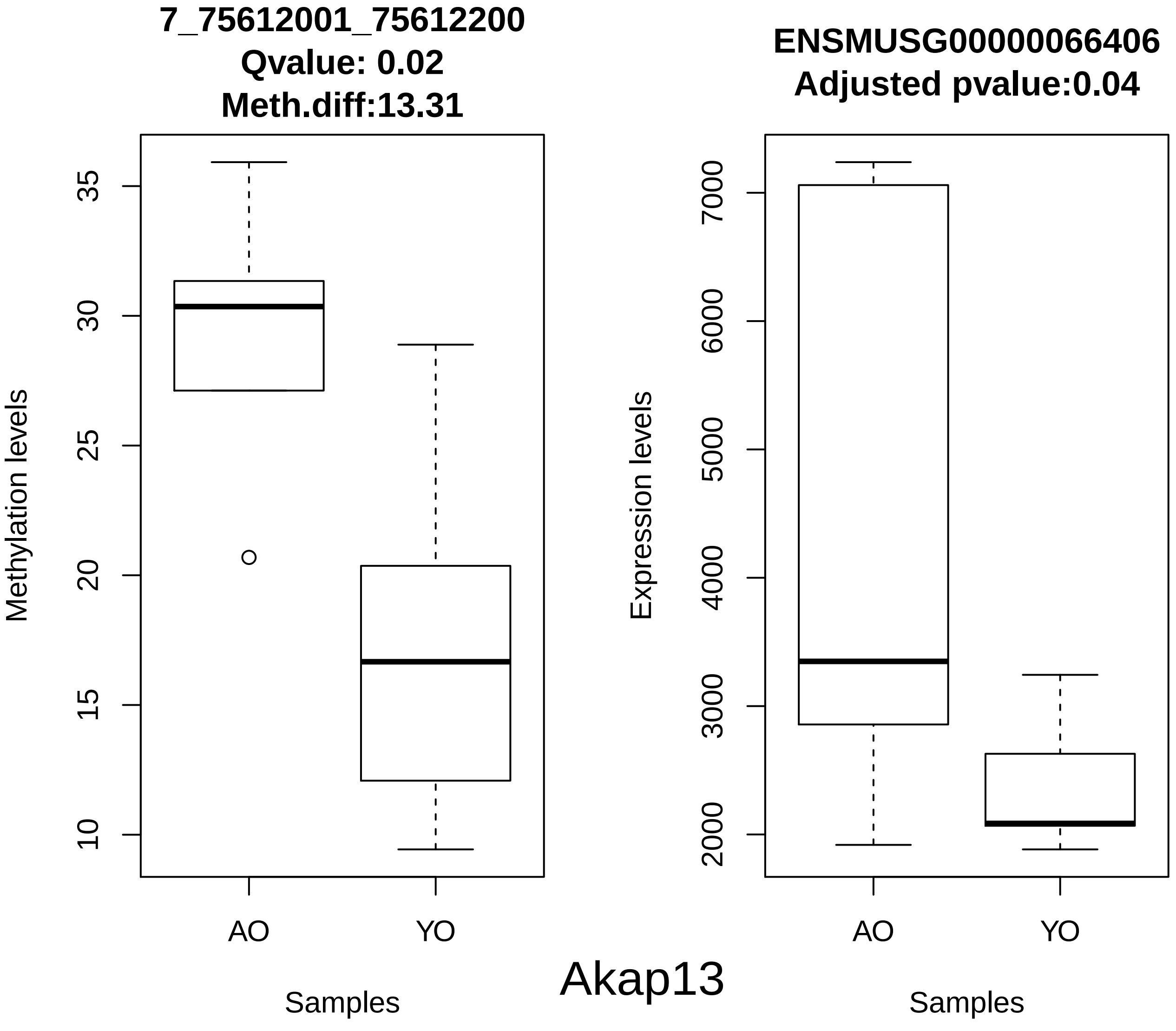
7_75612001_75612200
Qvalue: 0.02
Meth.diff:13.31
Methylation levels
10
15
20
25
30
35
AO
YO
Samples
ENSMUSG00000066406
Adjusted pvalue:0.04
Expression levels
2000
3000
4000
5000
6000
7000
AO
YO
Samples
Akap13

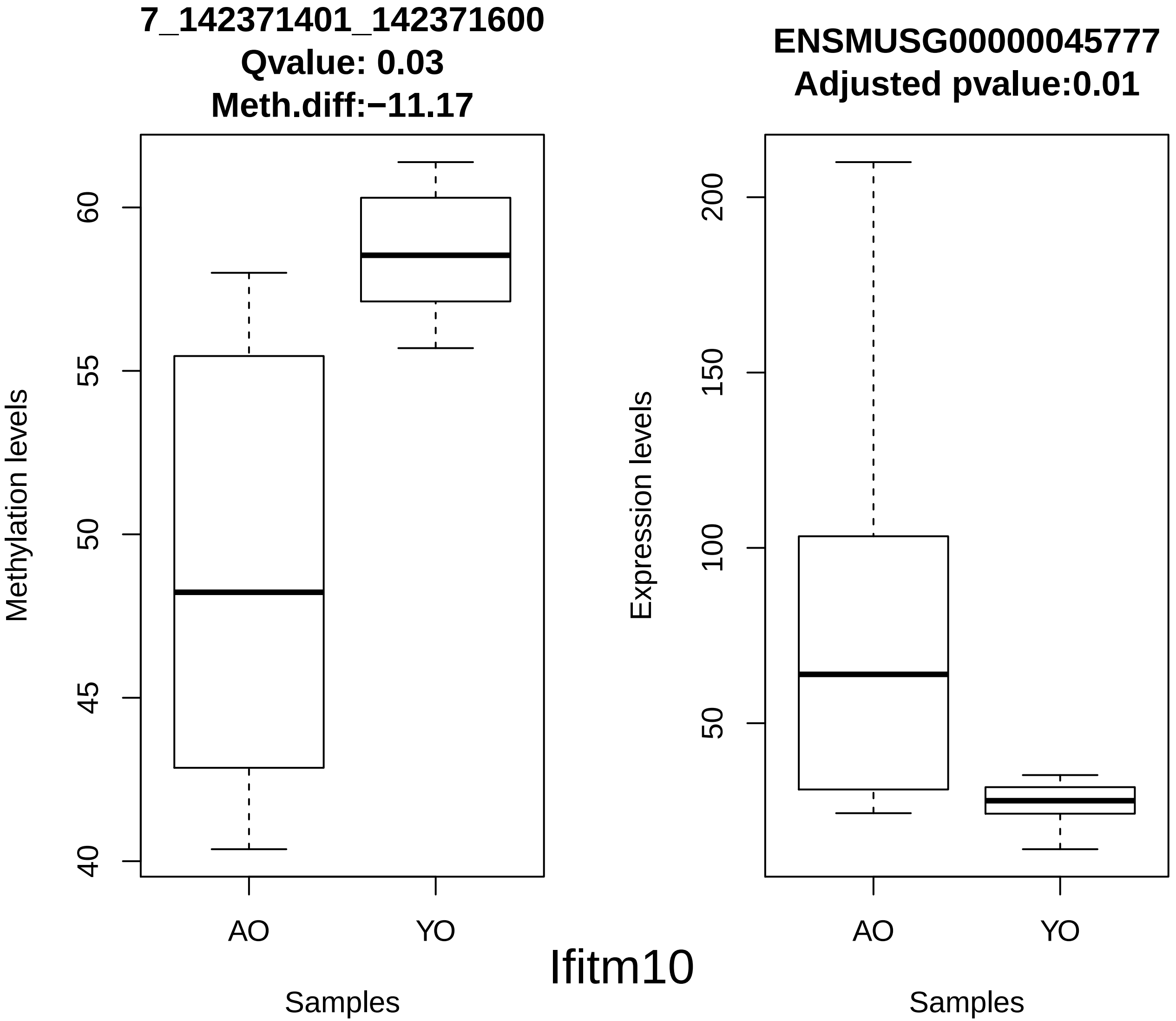
7_142371401_142371600
Qvalue: 0.03
Meth.diff:−11.17
Methylation levels
60
55
50
45
40
AO
YO
Samples
ENSMUSG00000045777
Adjusted pvalue:0.01
Expression levels
200
150
100
50
AO
YO
Samples

Ifitm10

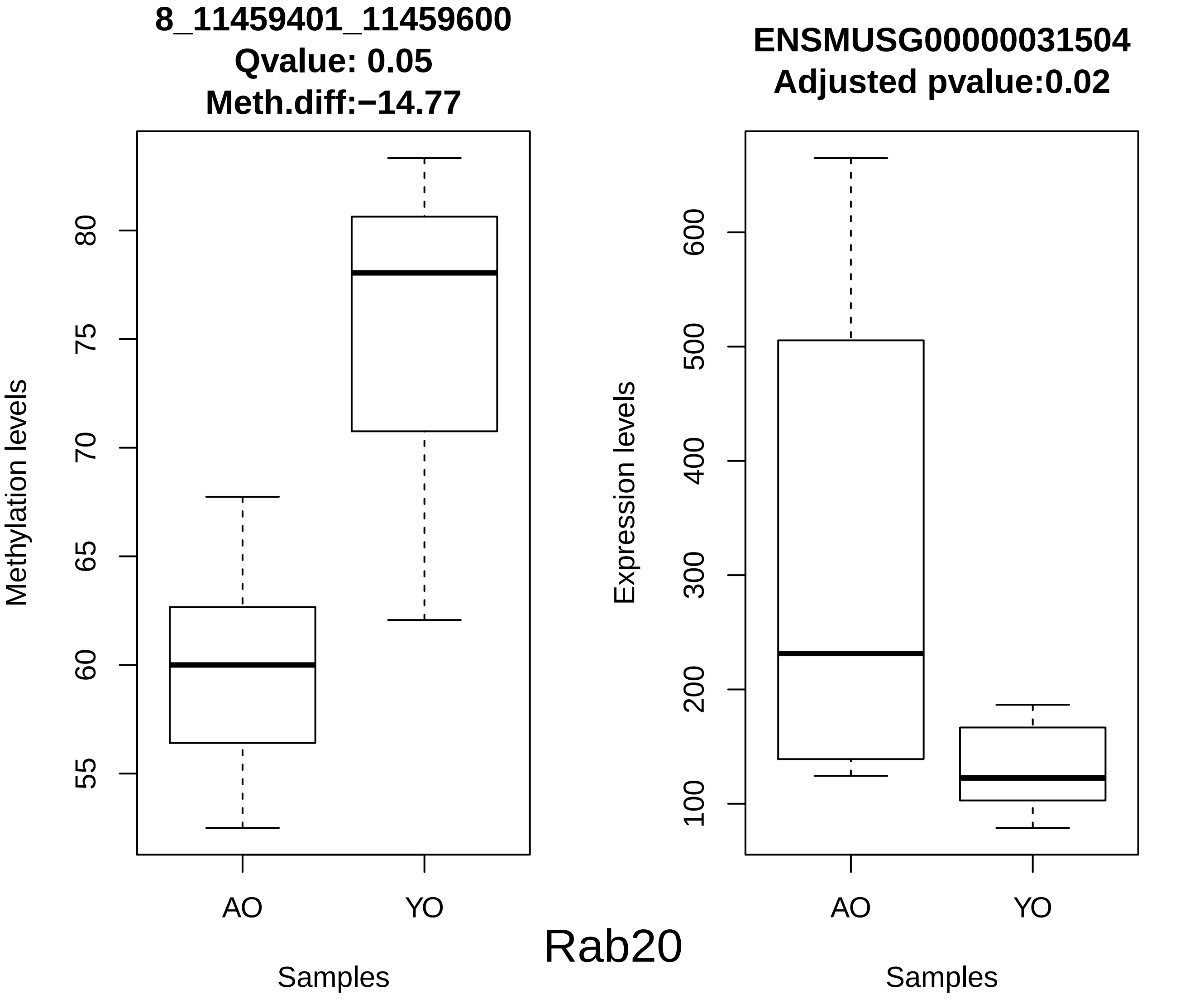
8_11459401_11459600
Qvalue: 0.05
Meth.diff:−14.77
Methylation levels
80
75
70
65
60
55
AO
YO
Samples
ENSMUSG00000031504
Adjusted pvalue:0.02
Expression levels
600
500
400
300
200
100
AO
YO
Samples
Rab20

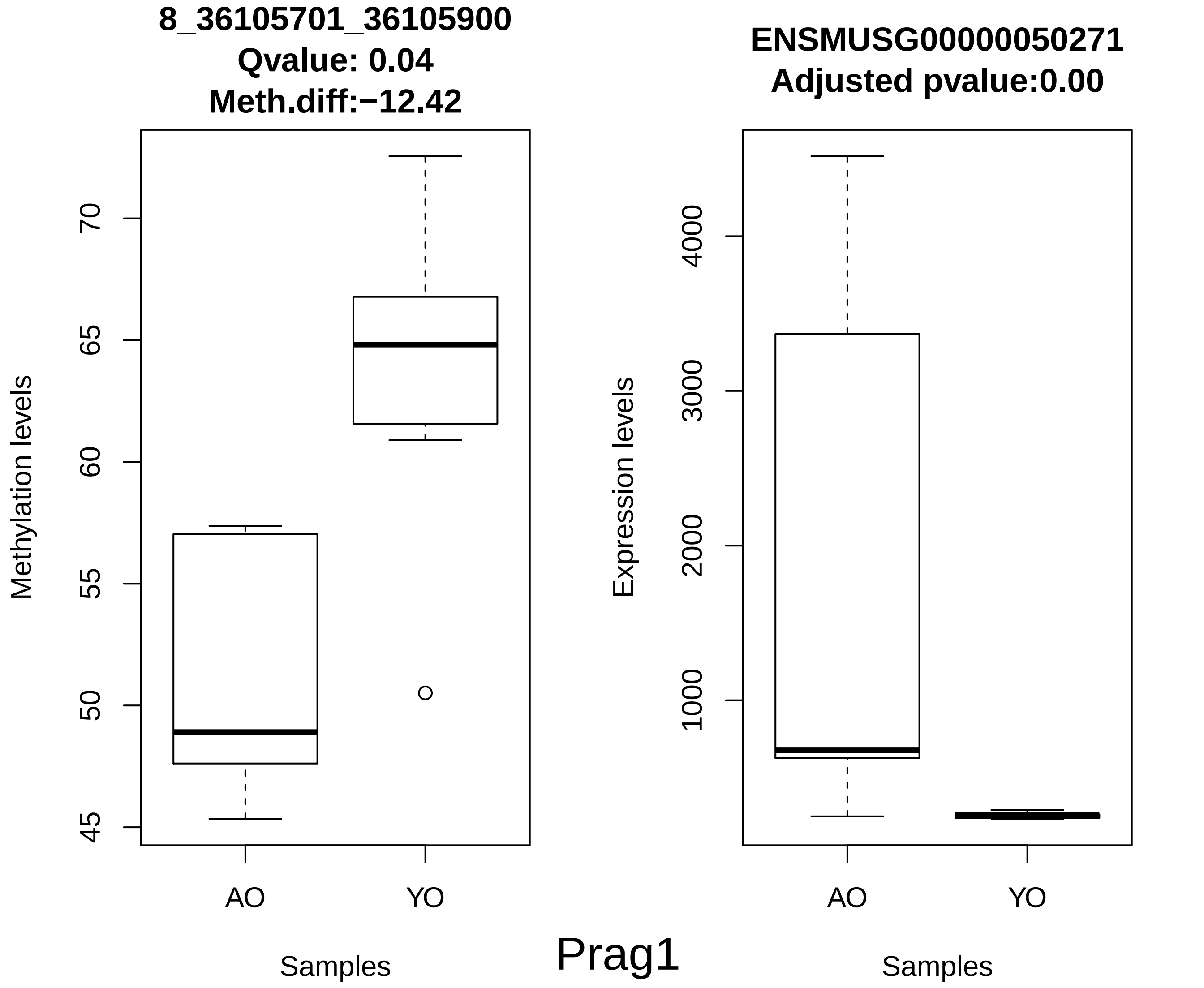
8_36105701_36105900
Qvalue: 0.04
Meth.diff:−12.42
Methylation levels
45
50
55
60
65
70
AO
YO
Samples
ENSMUSG00000050271
Adjusted pvalue:0.00
Expression levels
1000
2000
3000
4000
AO
YO
Samples
Prag1

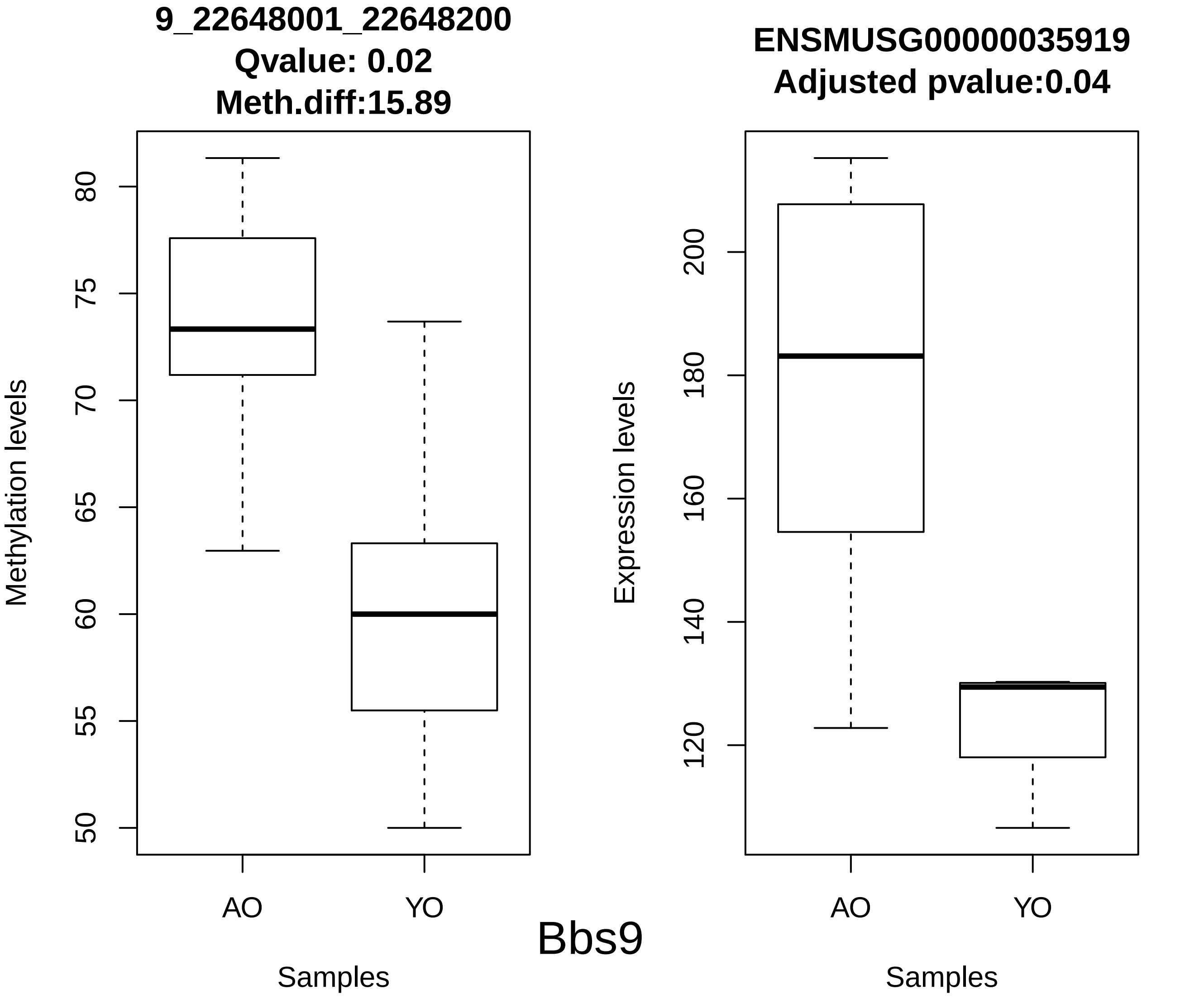
9_22648001_22648200
Qvalue: 0.02
Meth.diff:15.89
Methylation levels
50
55
60
65
70
75
80
AO
YO
Samples
ENSMUSG00000035919
Adjusted pvalue:0.04
Expression levels
120
140
160
180
200
AO
YO
Samples
Bbs9

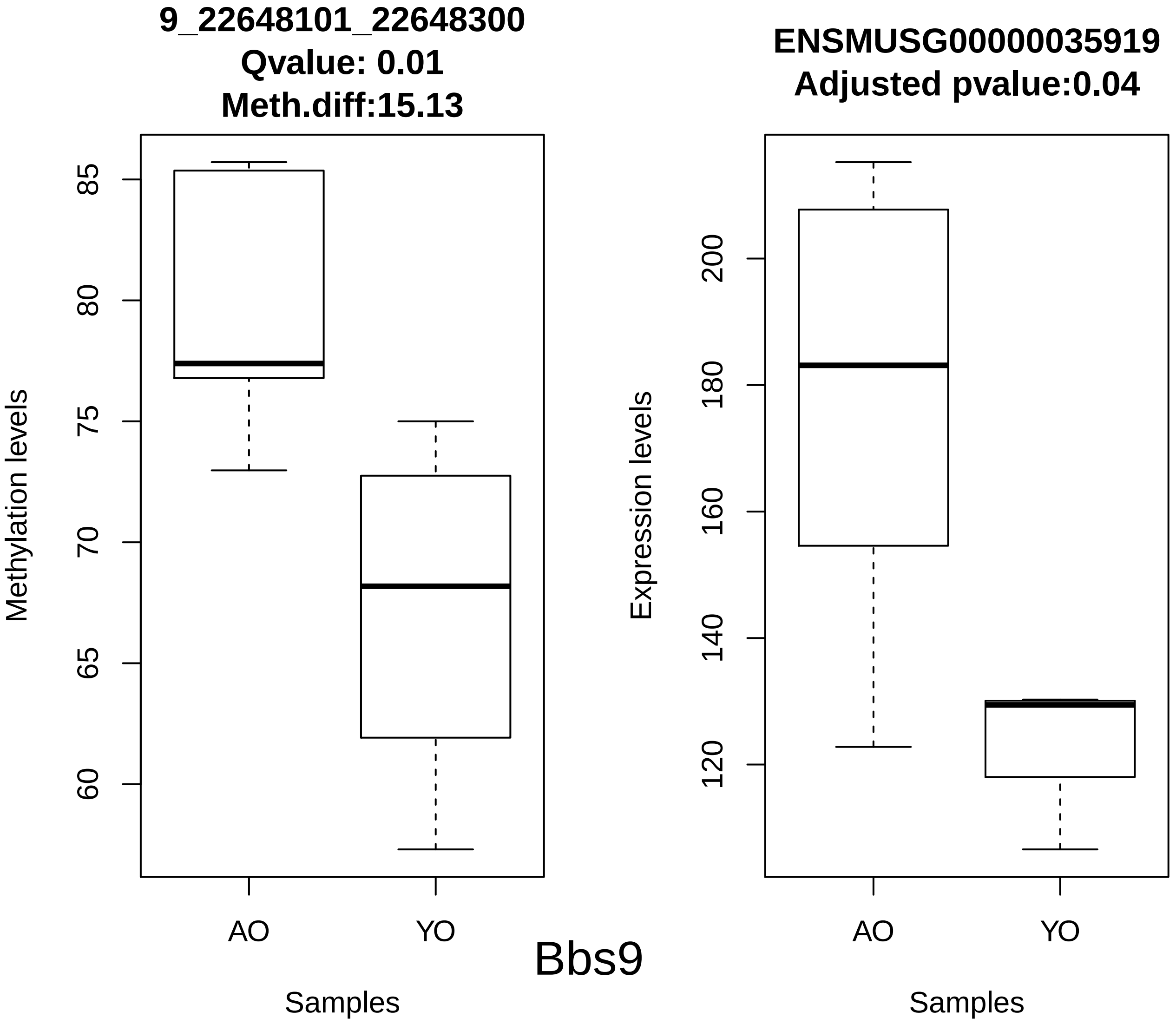

9_22648101_22648300
Qvalue: 0.01
Meth.diff:15.13
Methylation levels
85
80
75
70
65
60
AO
YO
Samples
ENSMUSG00000035919
Adjusted pvalue:0.04
Expression levels
200
180
160
140
120
AO
YO
Samples
Bbs9

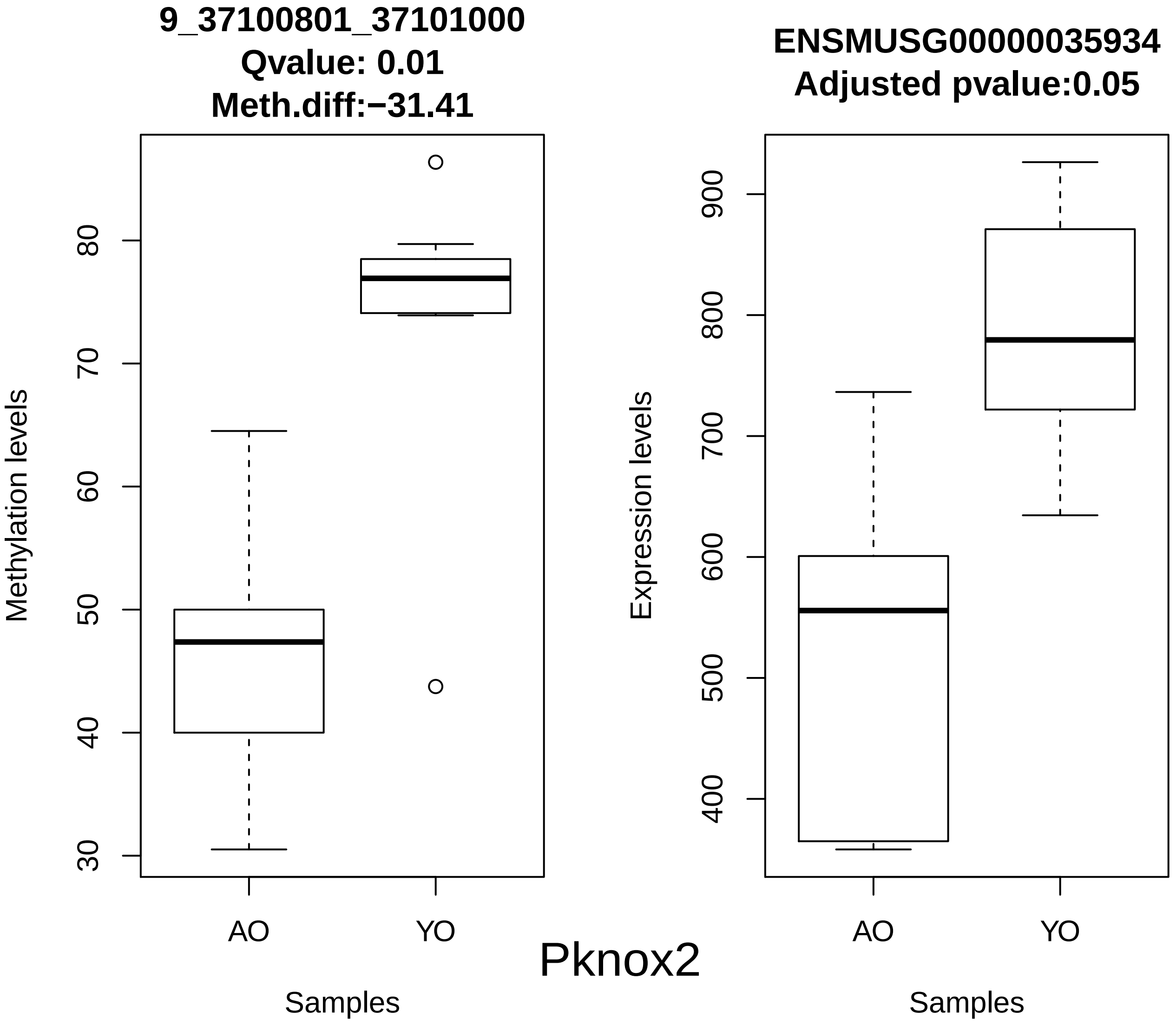
9_37100801_37101000
Qvalue: 0.01
Meth.diff:−31.41
Methylation levels
30
40
50
60
70
80
AO
YO
Samples
ENSMUSG00000035934
Adjusted pvalue:0.05
Expression levels
400
500
600
700
800
900
AO
YO
Samples
Pknox2

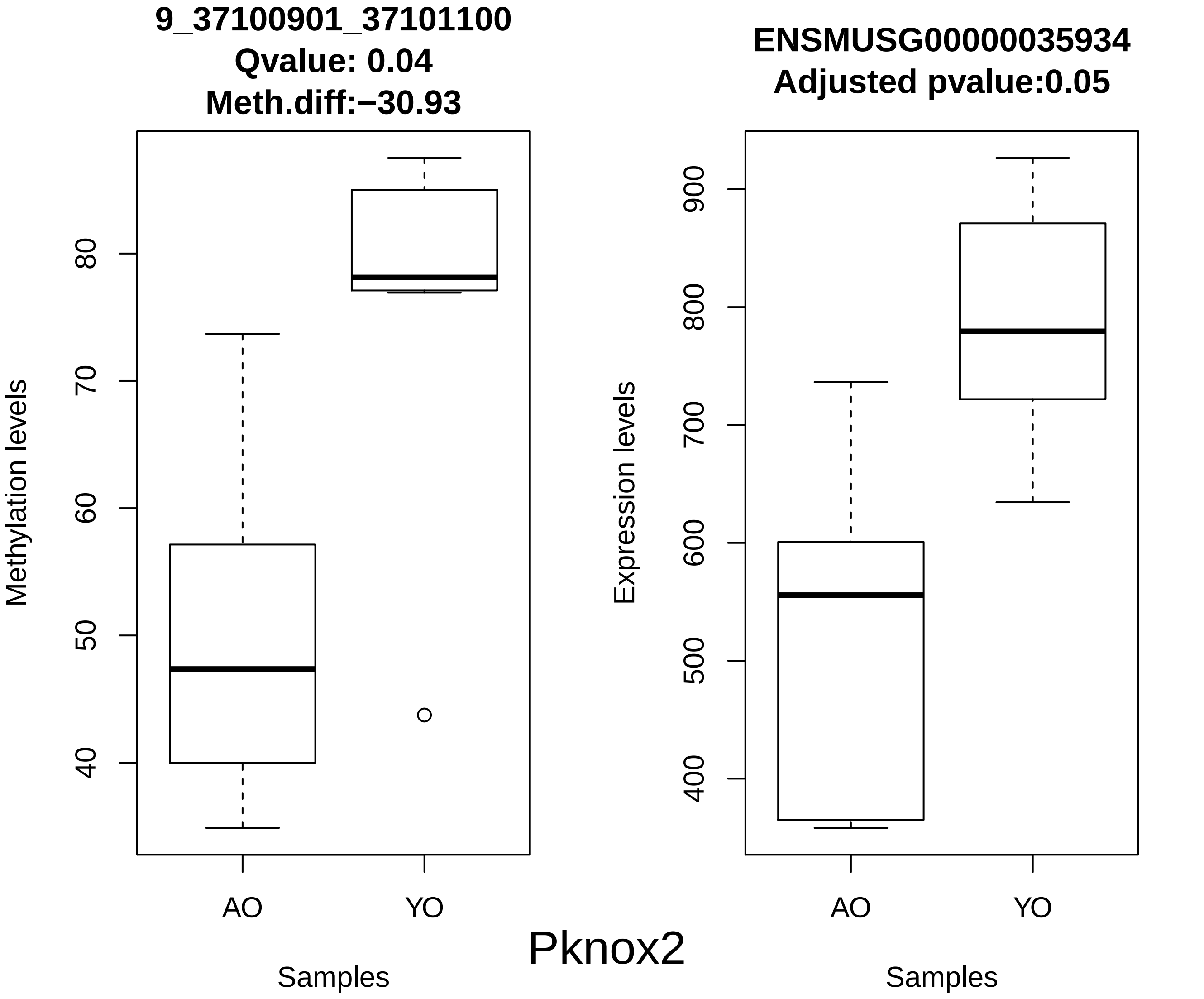
9_37100901_37101100
Qvalue: 0.04
Meth.diff:−30.93
Methylation levels
80
70
60
50
40
AO
YO
Samples
ENSMUSG00000035934
Adjusted pvalue:0.05
Expression levels
900
800
700
600
500
400
AO
YO
Samples

Pknox2

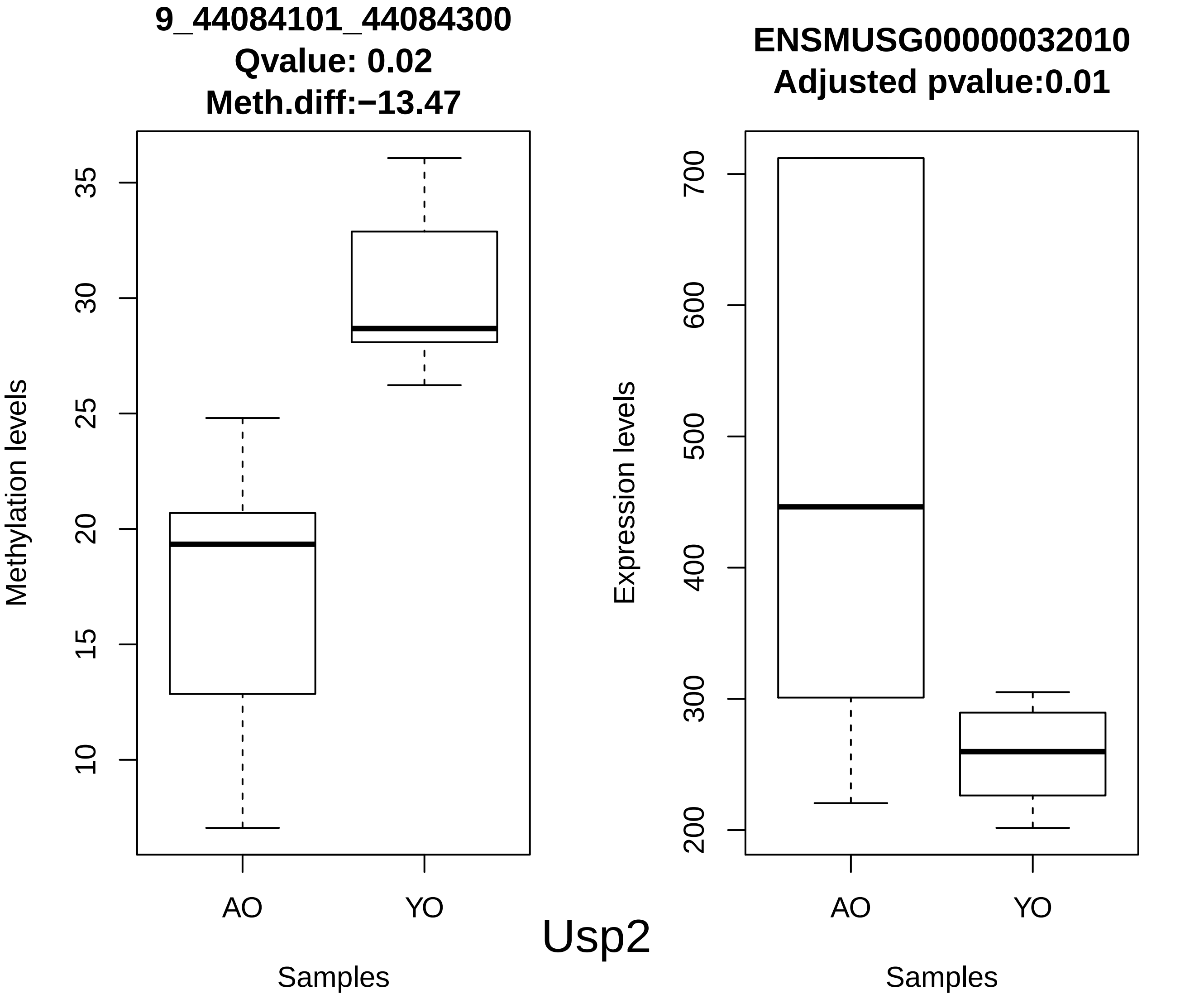

9_44084101_44084300
Qvalue: 0.02
Meth.diff:−13.47
Methylation levels
35
30
25
20
15
10
AO
YO
Samples
ENSMUSG00000032010
Adjusted pvalue:0.01
Expression levels
700
600
500
400
300
200
AO
YO
Samples


Usp2

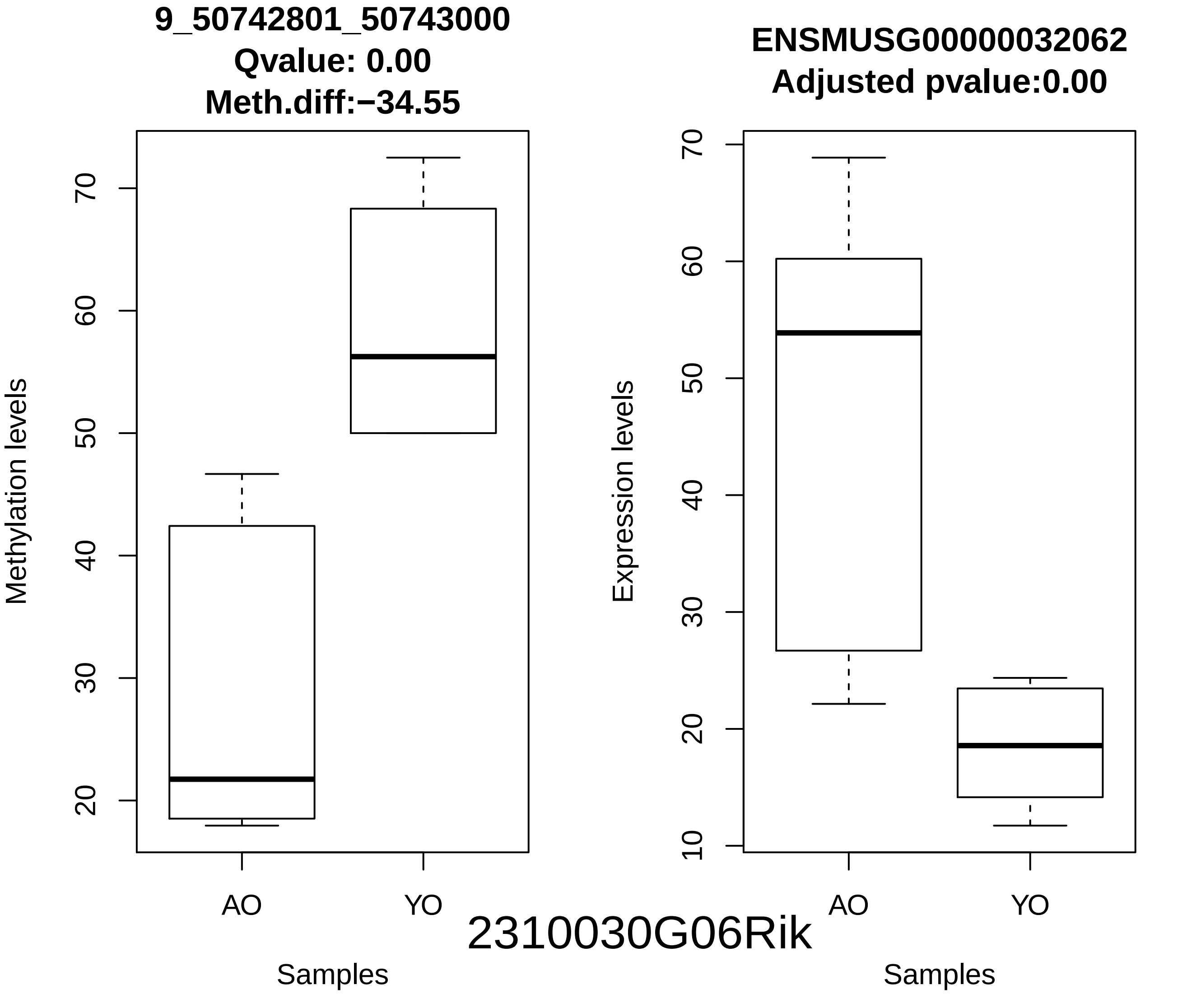
9_50742801_50743000
Qvalue: 0.00
Meth.diff:−34.55
Methylation levels
70
60
50
40
30
20
AO
YO
Samples
ENSMUSG00000032062
Adjusted pvalue:0.00
Expression levels
70
60
50
40
30
20
10
AO
YO
Samples
2310030G06Rik

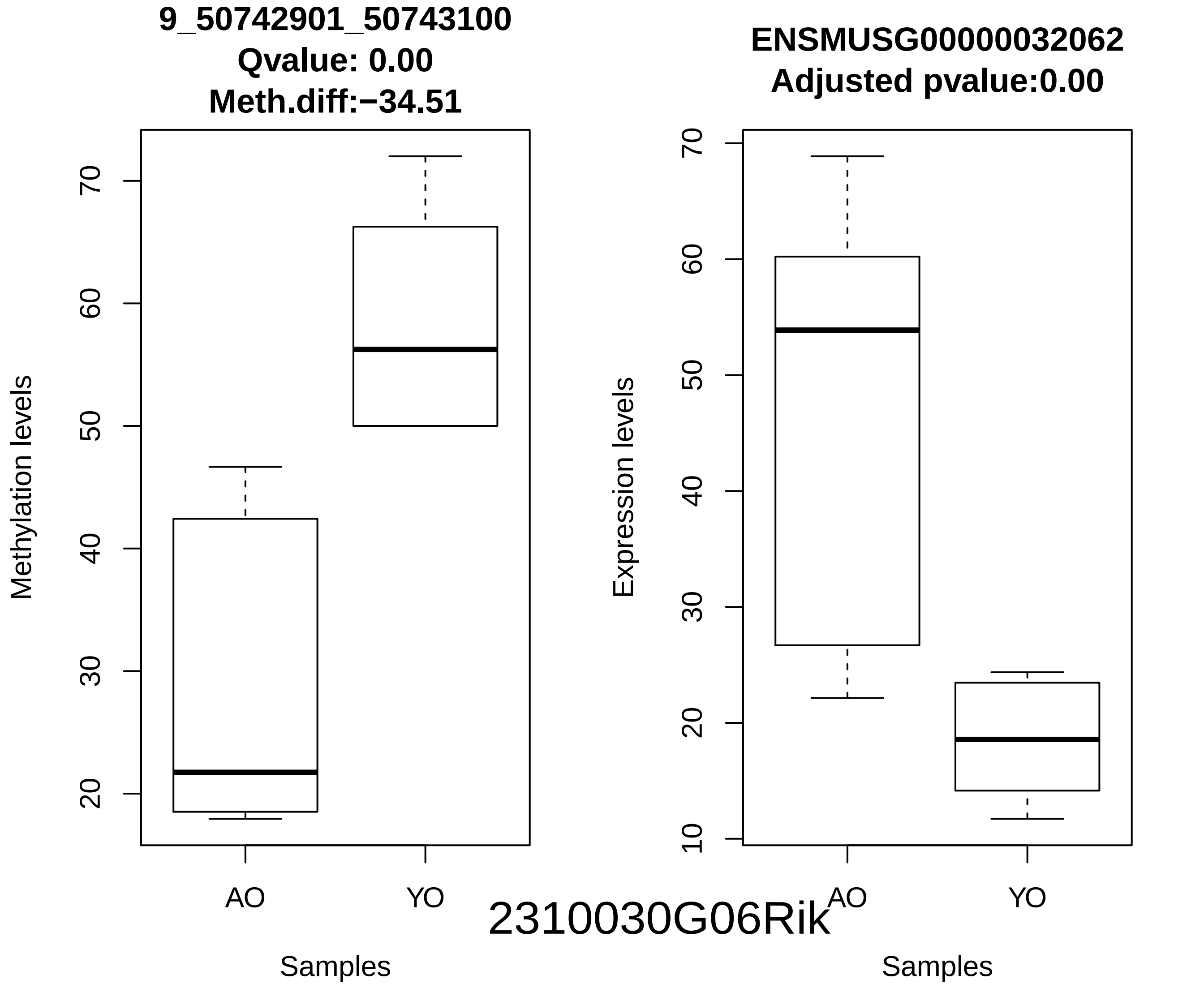

9_50742901_50743100
Qvalue: 0.00
Meth.diff:−34.51
Methylation levels
20
30
40
50
60
70
AO
YO
Samples
ENSMUSG00000032062
Adjusted pvalue:0.00
Expression levels
10
20
30
40
50
60
70
AO
YO
Samples
2310030G06Rik

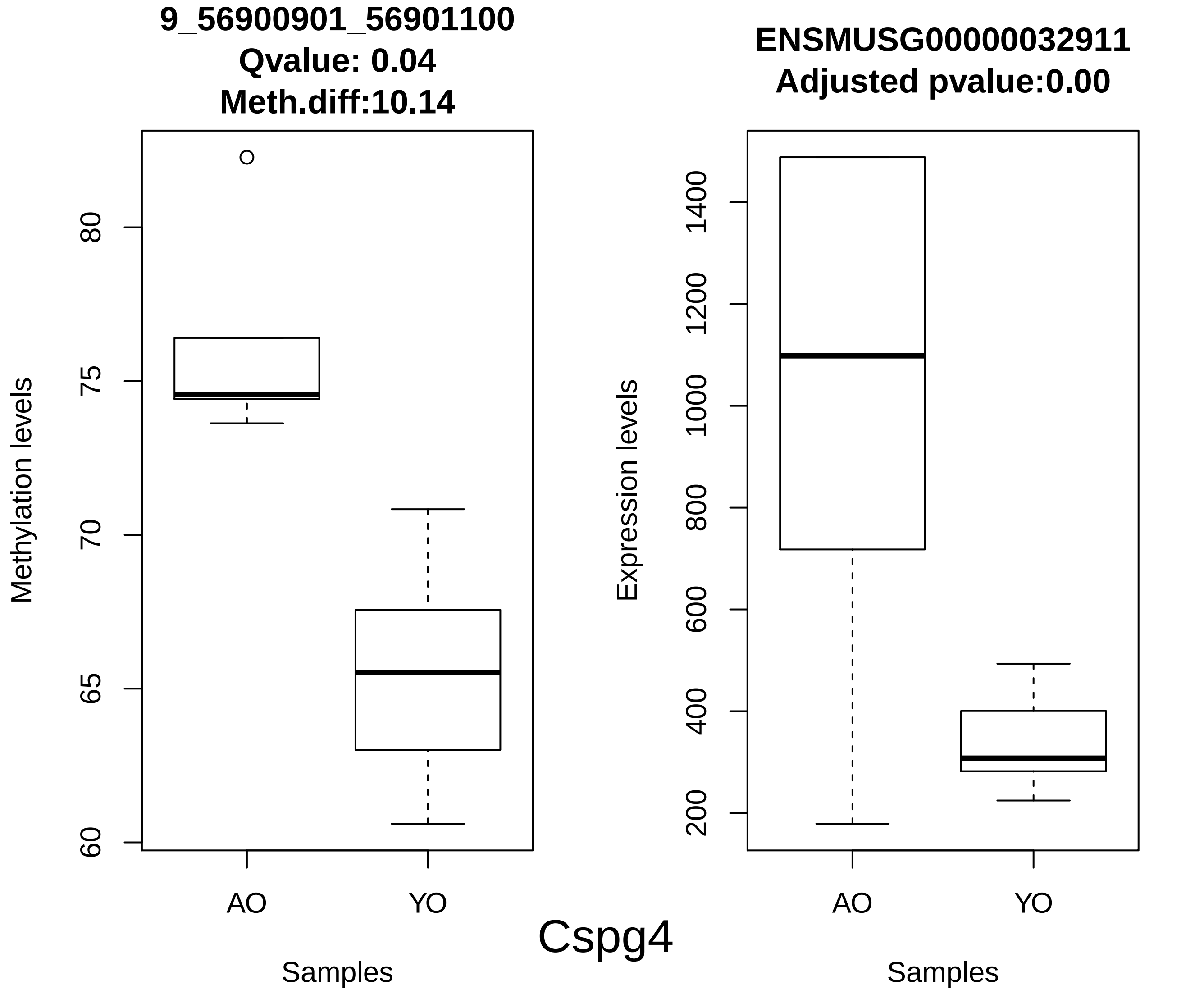

9_56900901_56901100
Qvalue: 0.04
Meth.diff:10.14
Methylation levels
60
65
70
75
80
AO
YO
Samples
ENSMUSG00000032911
Adjusted pvalue:0.00
Expression levels
200
400
600
800
1000
1200
1400
AO
YO
Samples
Cspg4

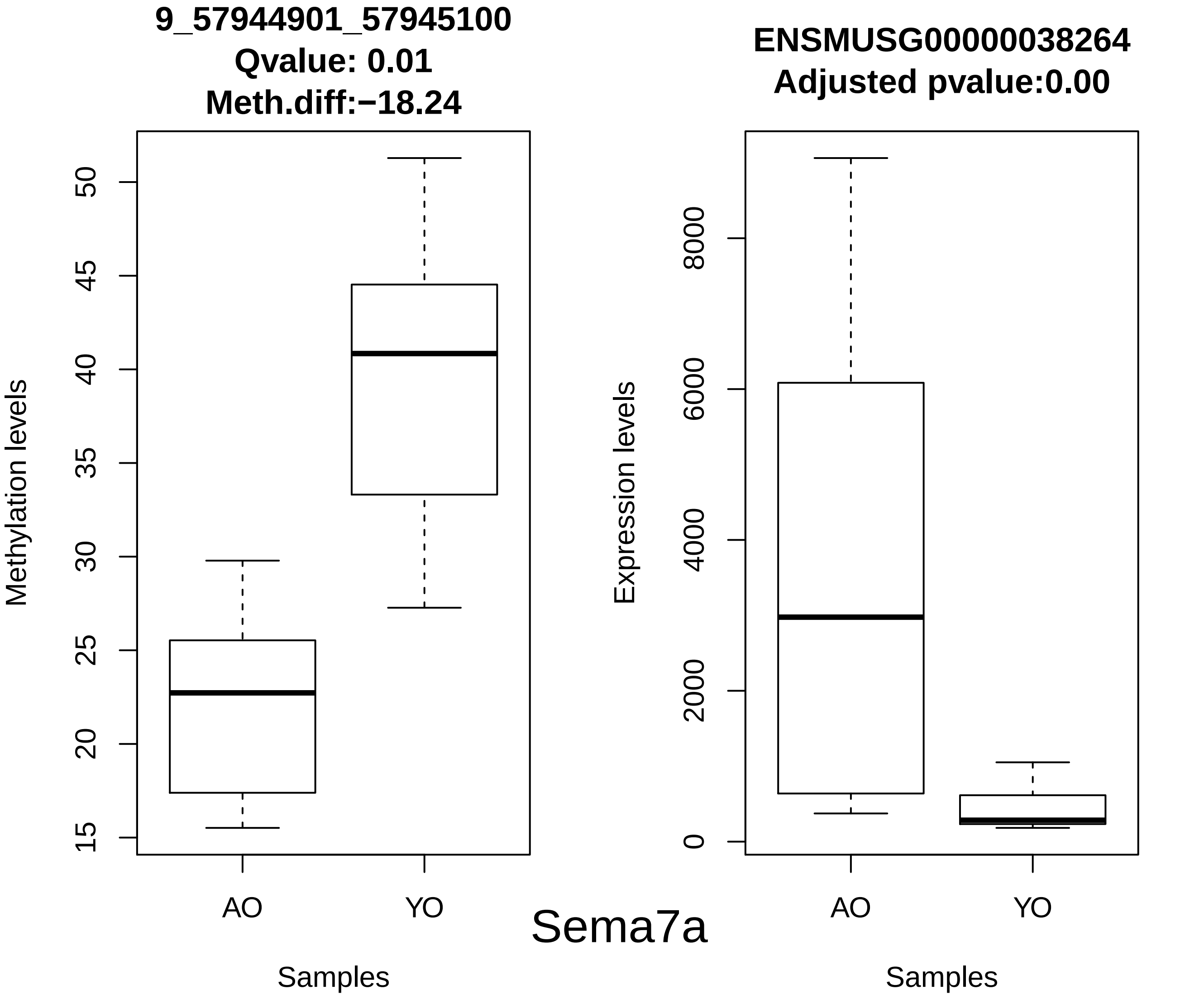
9_57944901_57945100
Qvalue: 0.01
Meth.diff:−18.24
Methylation levels
15
20
25
30
35
40
45
50
AO
YO
Samples
Sema7a
ENSMUSG00000038264
Adjusted pvalue:0.00
Expression levels
0
2000
4000
6000
8000
AO
YO
Samples

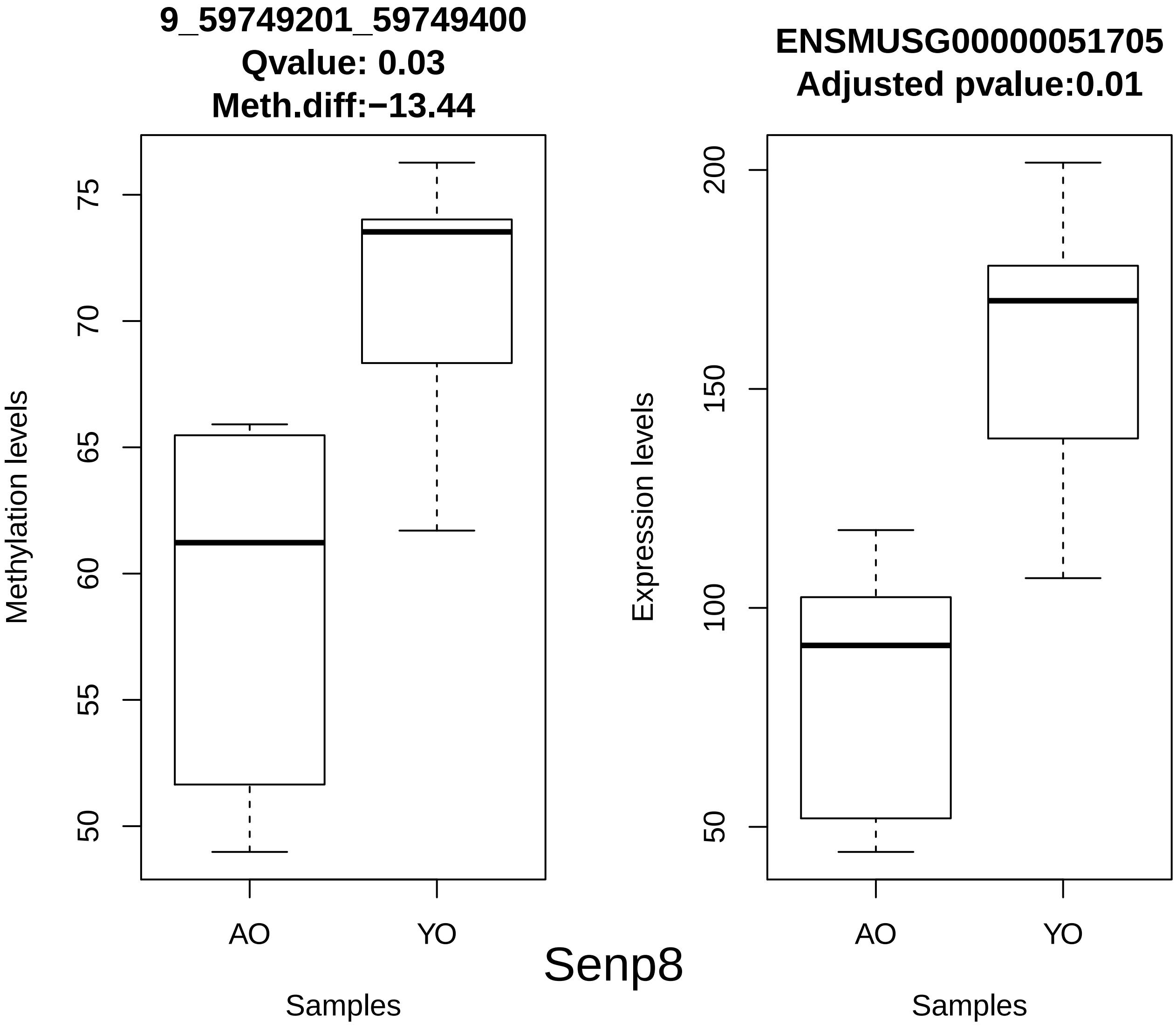
9_59749201_59749400
Qvalue: 0.03
Meth.diff:−13.44
Methylation levels
75
70
65
60
55
50
AO
YO
Samples
ENSMUSG00000051705
Adjusted pvalue:0.01
Expression levels
200
150
100
50
AO
YO
Samples
Senp8

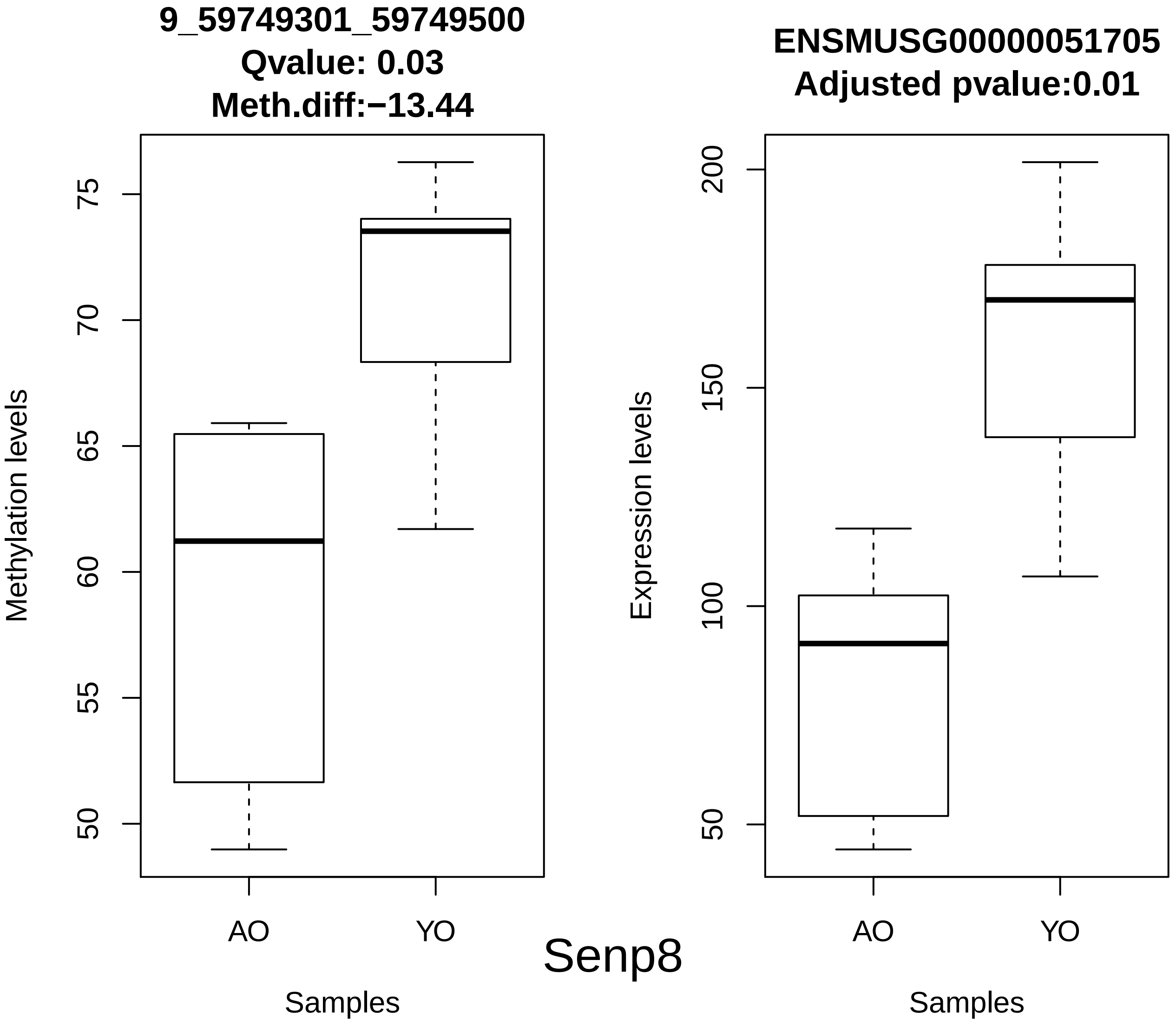

9_59749301_59749500
Qvalue: 0.03
Meth.diff:−13.44
Methylation levels
75
70
65
60
55
50
AO
YO
Samples
ENSMUSG00000051705
Adjusted pvalue:0.01
Expression levels
200
150
100
50
AO
YO
Samples


Senp8

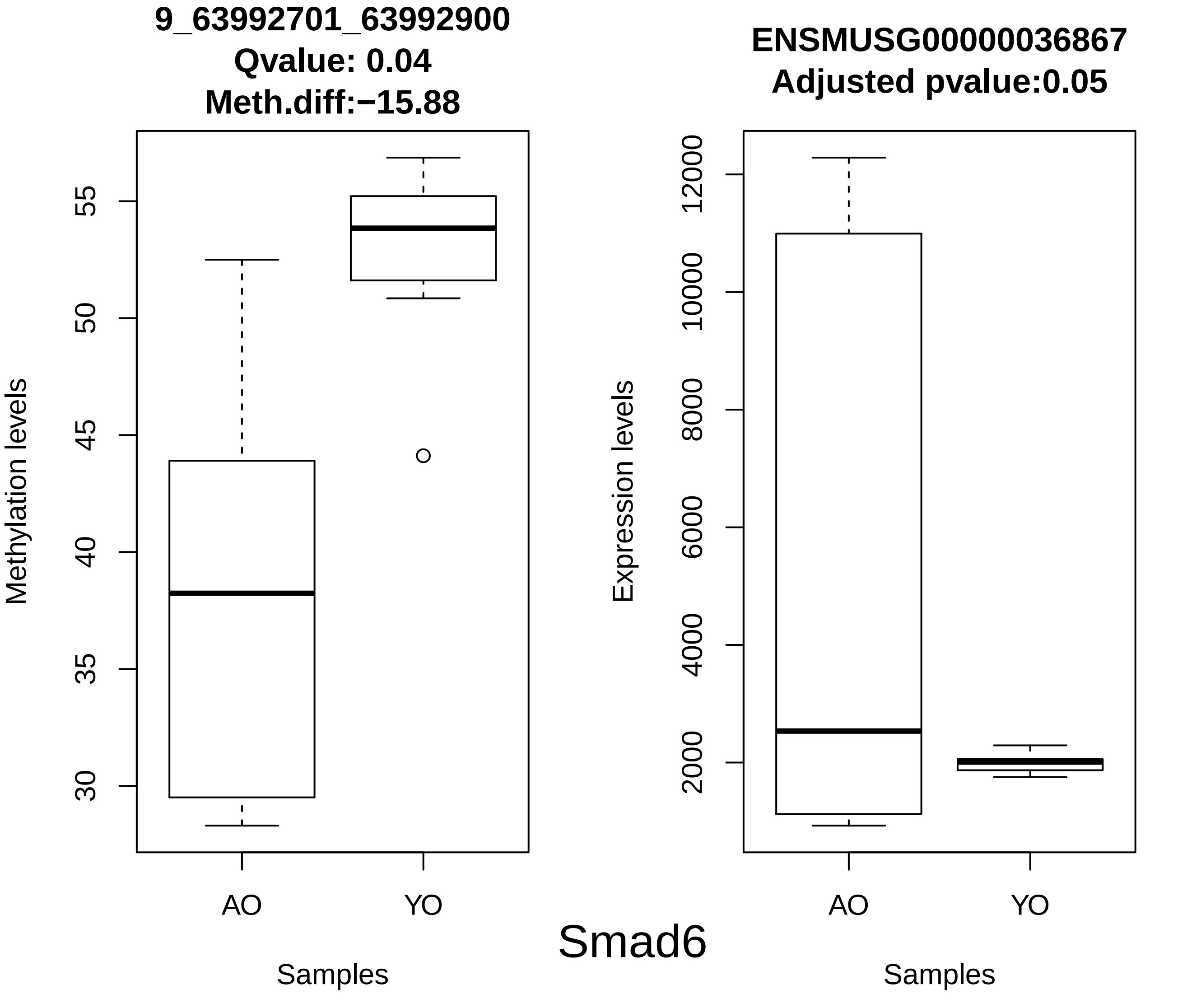
9_63992701_63992900
Qvalue: 0.04
Meth.diff:−15.88
Methylation levels
55
50
45
40
35
30
AO
YO
Samples
ENSMUSG00000036867
Adjusted pvalue:0.05
Expression levels
12000
10000
8000
6000
4000
2000
AO
YO
Samples
Smad6

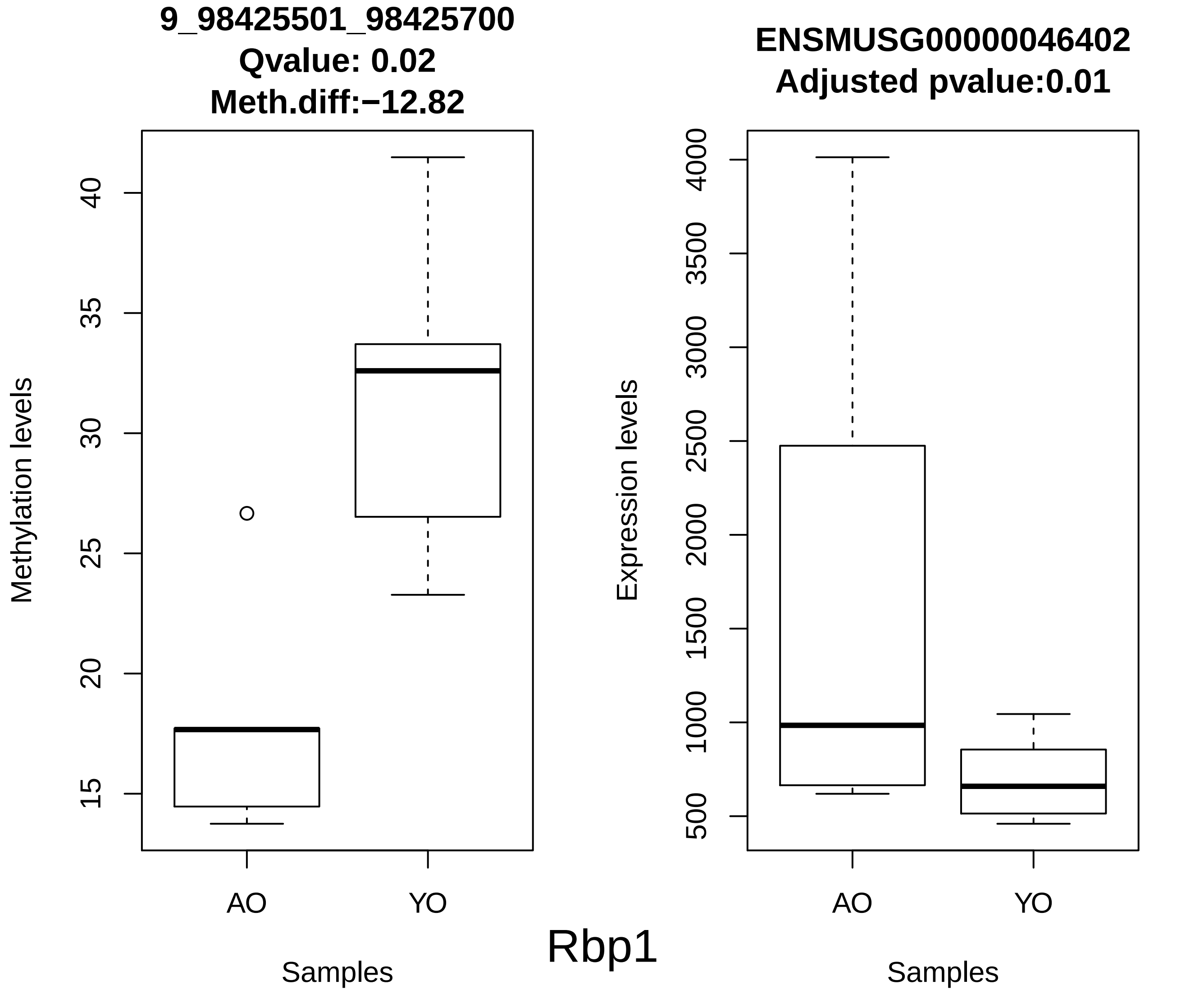
9_98425501_98425700
Qvalue: 0.02
Meth.diff:−12.82
Methylation levels
40
35
30
25
20
15
AO
YO
Samples
ENSMUSG00000046402
Adjusted pvalue:0.01
Expression levels
4000
3500
3000
2500
2000
1500
1000
500
AO
YO
Samples
Rbp1

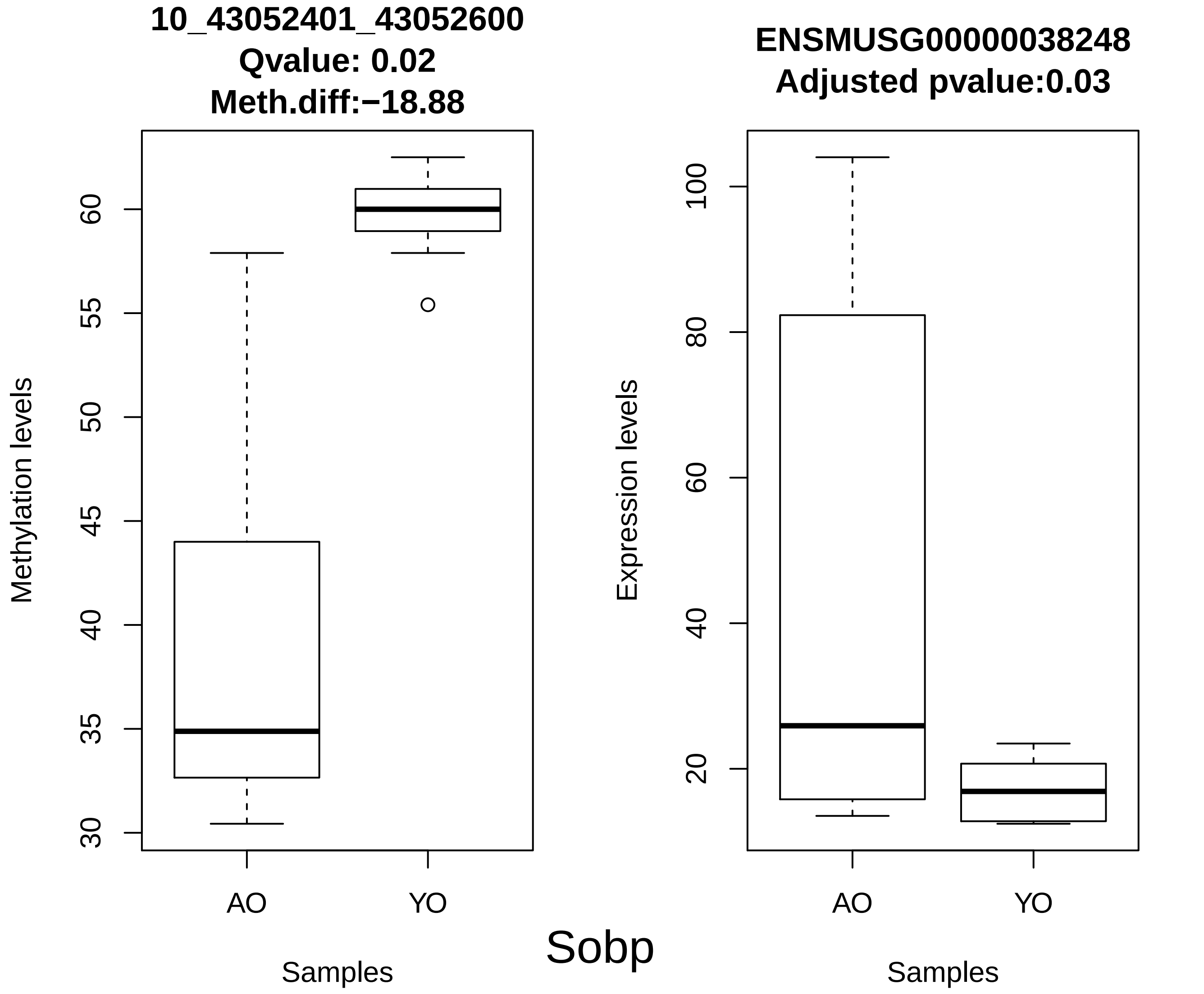

10_43052401_43052600
Qvalue: 0.02
Meth.diff:−18.88
Methylation levels
30
35
40
45
50
55
60
AO
YO
Samples
ENSMUSG00000038248
Adjusted pvalue:0.03
Expression levels
20
40
60
80
100
AO
YO
Samples
Sobp

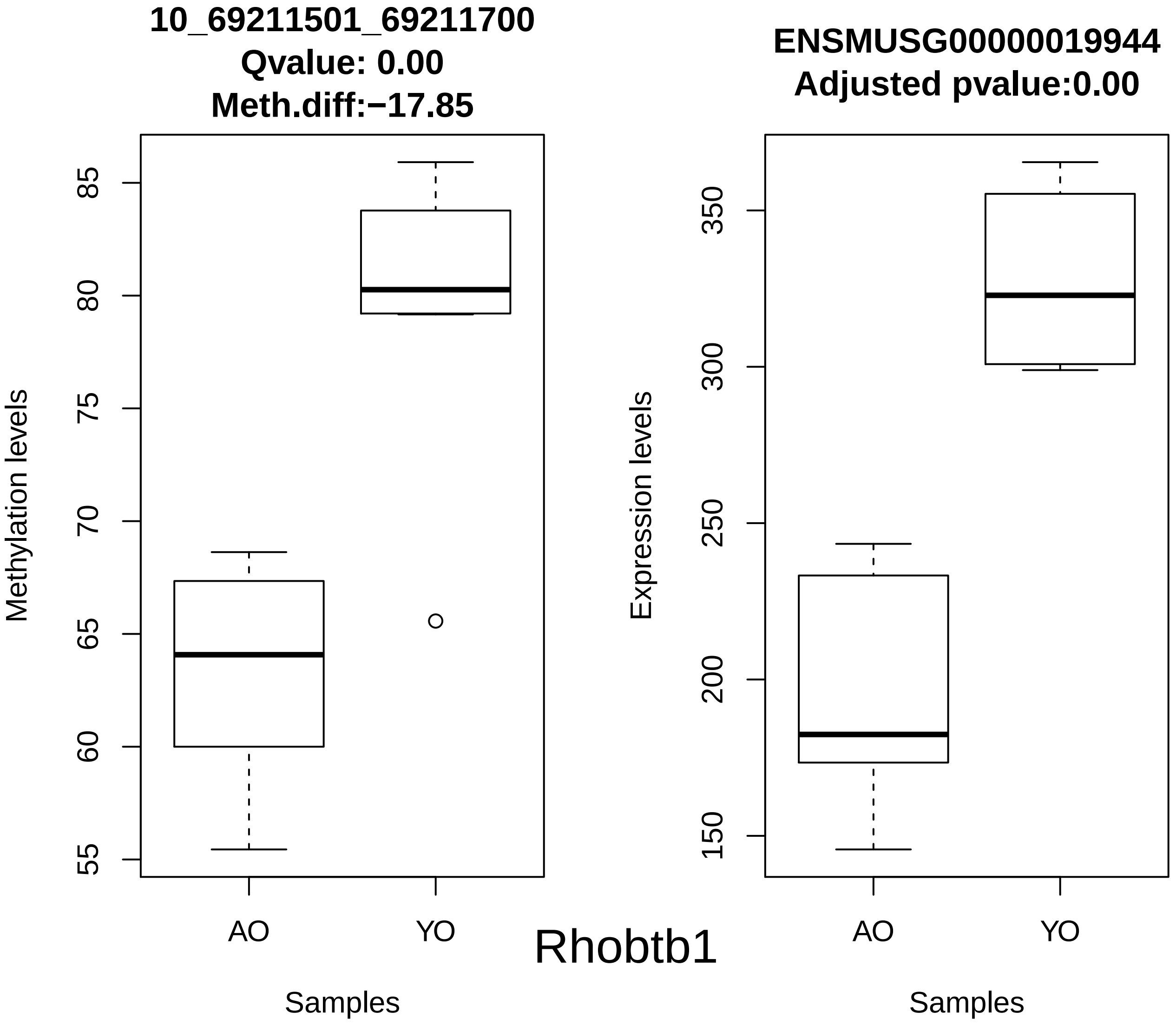

10_69211501_69211700
Qvalue: 0.00
Meth.diff:−17.85
Methylation levels
55
60
65
70
75
80
85
AO
YO
Samples
Rhobtb1
ENSMUSG00000019944
Adjusted pvalue:0.00
Expression levels
150
200
250
300
350
AO
YO
Samples

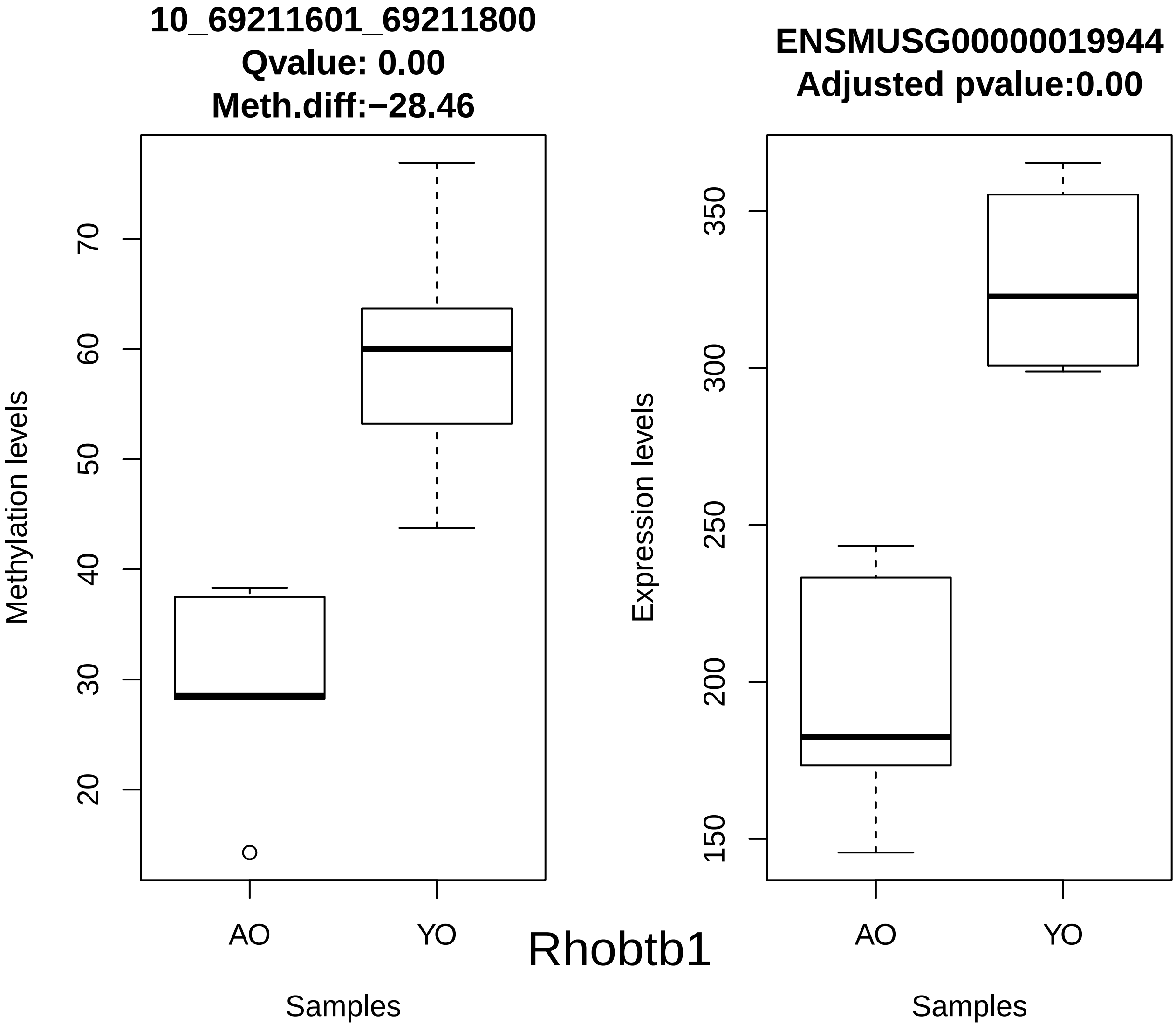
10_69211601_69211800
Qvalue: 0.00
Meth.diff:−28.46
Methylation levels
20
30
40
50
60
70
AO
YO
Samples
Rhobtb1
ENSMUSG00000019944
Adjusted pvalue:0.00
Expression levels
150
200
250
300
350
AO
YO
Samples

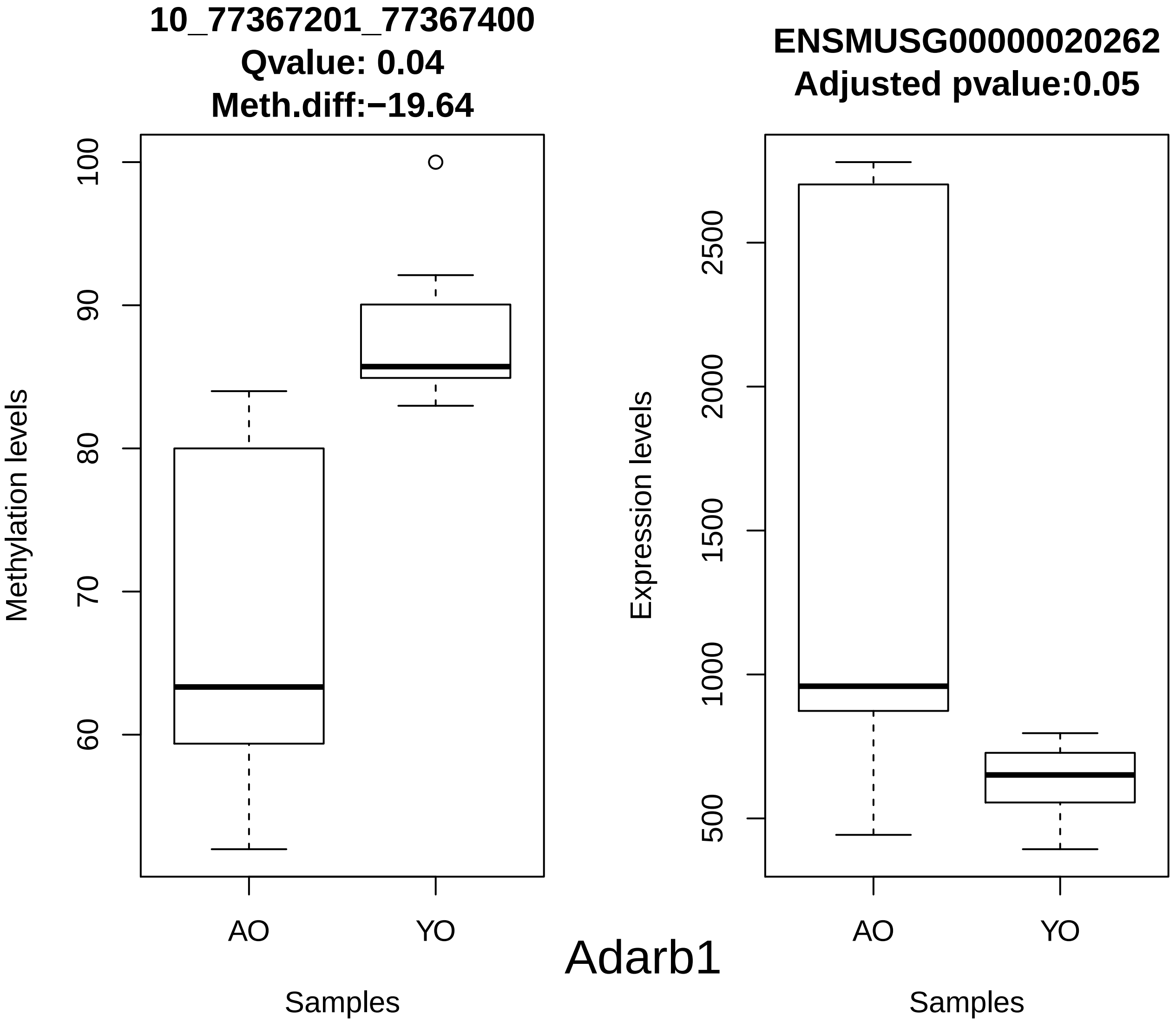

10_77367201_77367400
Qvalue: 0.04
Meth.diff:−19.64
Methylation levels
100
90
80
70
60
AO
YO
Samples
ENSMUSG00000020262
Adjusted pvalue:0.05
Expression levels
2500
2000
1500
1000
500
AO
YO
Samples
Adarb1

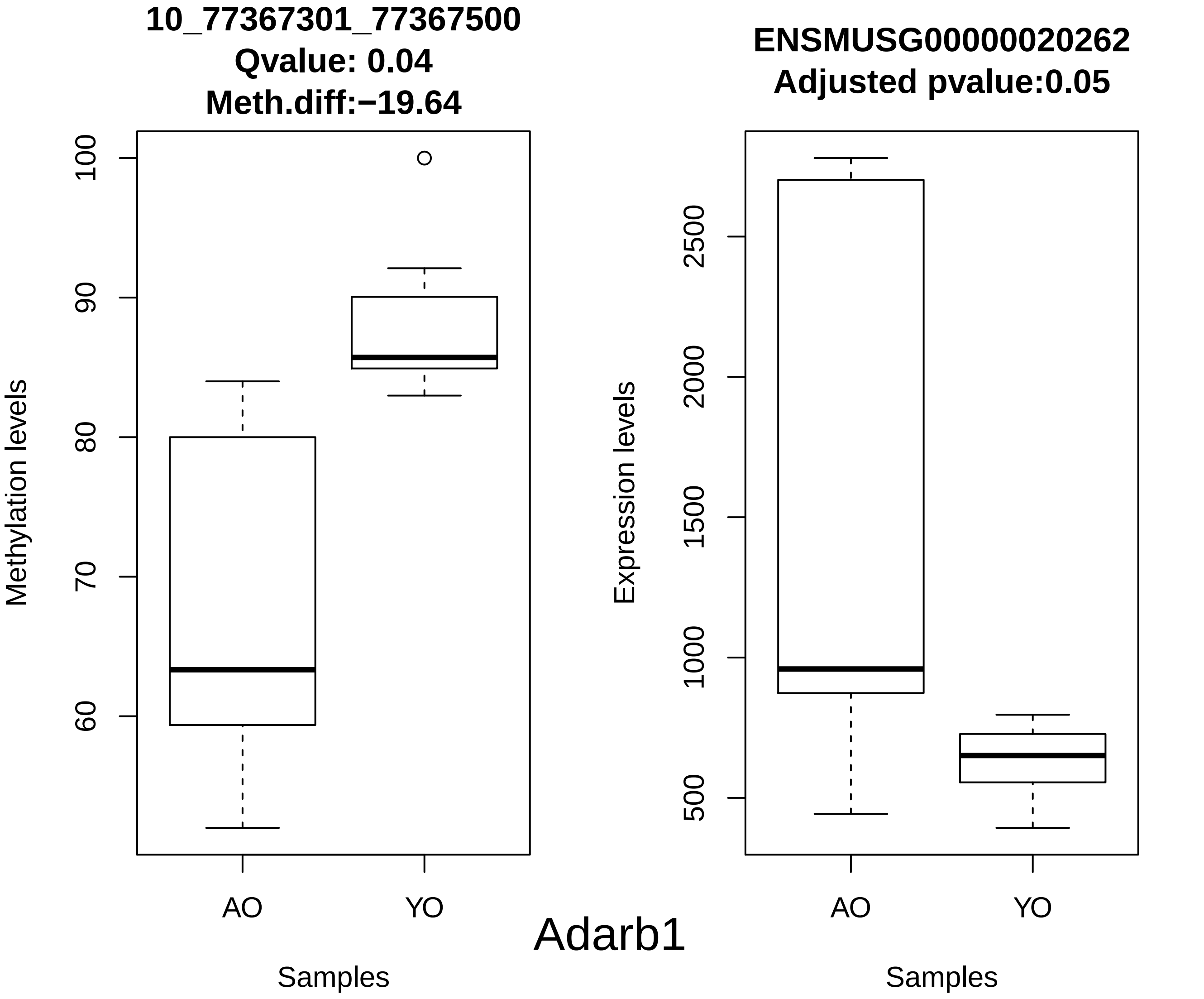
10_77367301_77367500
Qvalue: 0.04
Meth.diff:−19.64
Methylation levels
100
90
80
70
60
AO
YO
Samples
ENSMUSG00000020262
Adjusted pvalue:0.05
Expression levels
2500
2000
1500
1000
500
AO
YO
Samples
Adarb1

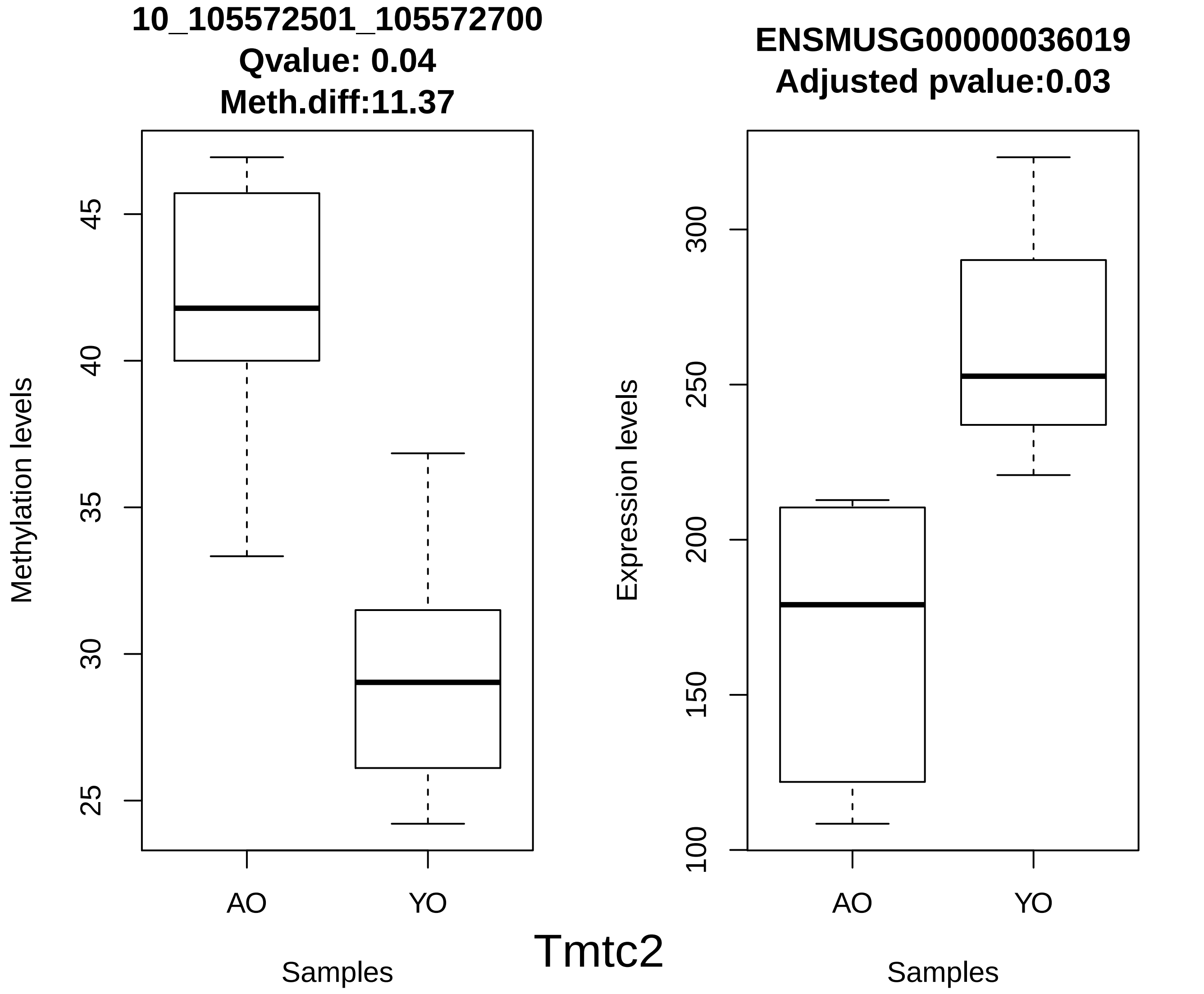

10_105572501_105572700
Qvalue: 0.04
Meth.diff:11.37
Methylation levels
45
40
35
30
25
AO
YO
Samples
ENSMUSG00000036019
Adjusted pvalue:0.03
Expression levels
300
250
200
150
100
AO
YO
Samples
Tmtc2

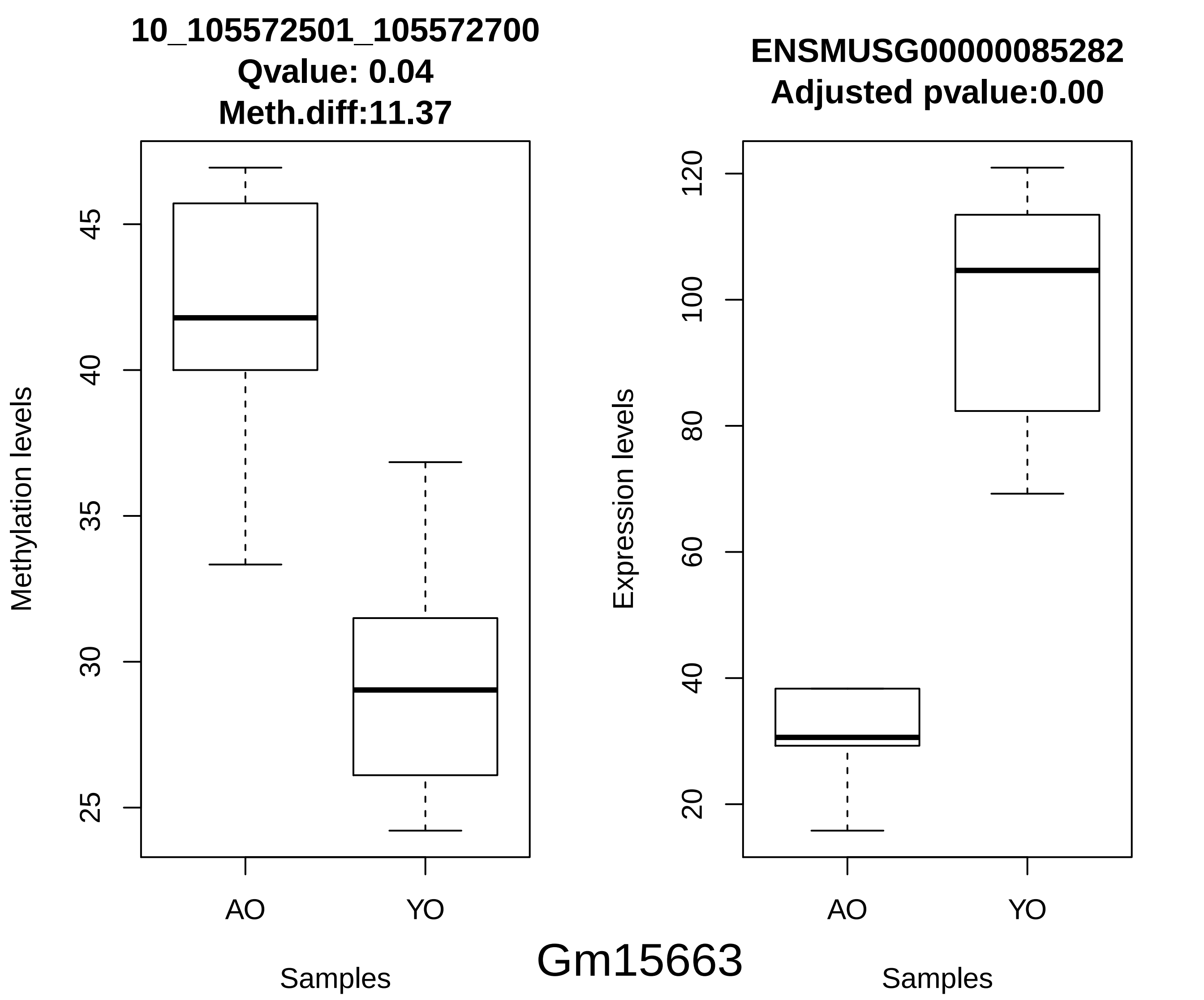
10_105572501_105572700
Qvalue: 0.04
Meth.diff:11.37
Methylation levels
45
40
35
30
25
AO
YO
Samples
ENSMUSG00000085282
Adjusted pvalue:0.00
Expression levels
120
100
80
60
40
20
AO
YO
Samples
Gm15663

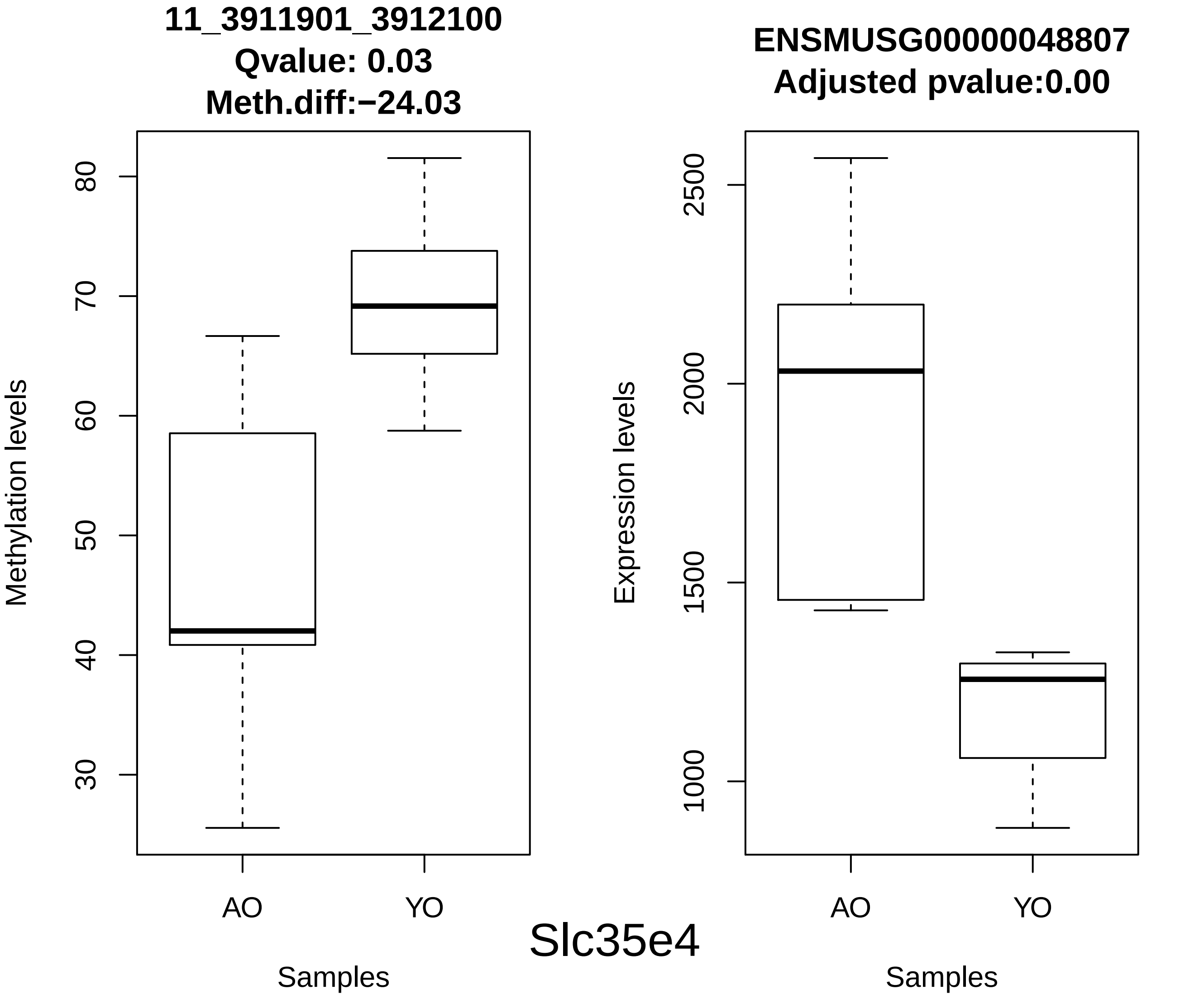
11_3911901_3912100
Qvalue: 0.03
Meth.diff:−24.03
Methylation levels
80
70
60
50
40
30
AO
YO
Samples
ENSMUSG00000048807
Adjusted pvalue:0.00
Expression levels
2500
2000
1500
1000
AO
YO
Samples
Slc35e4

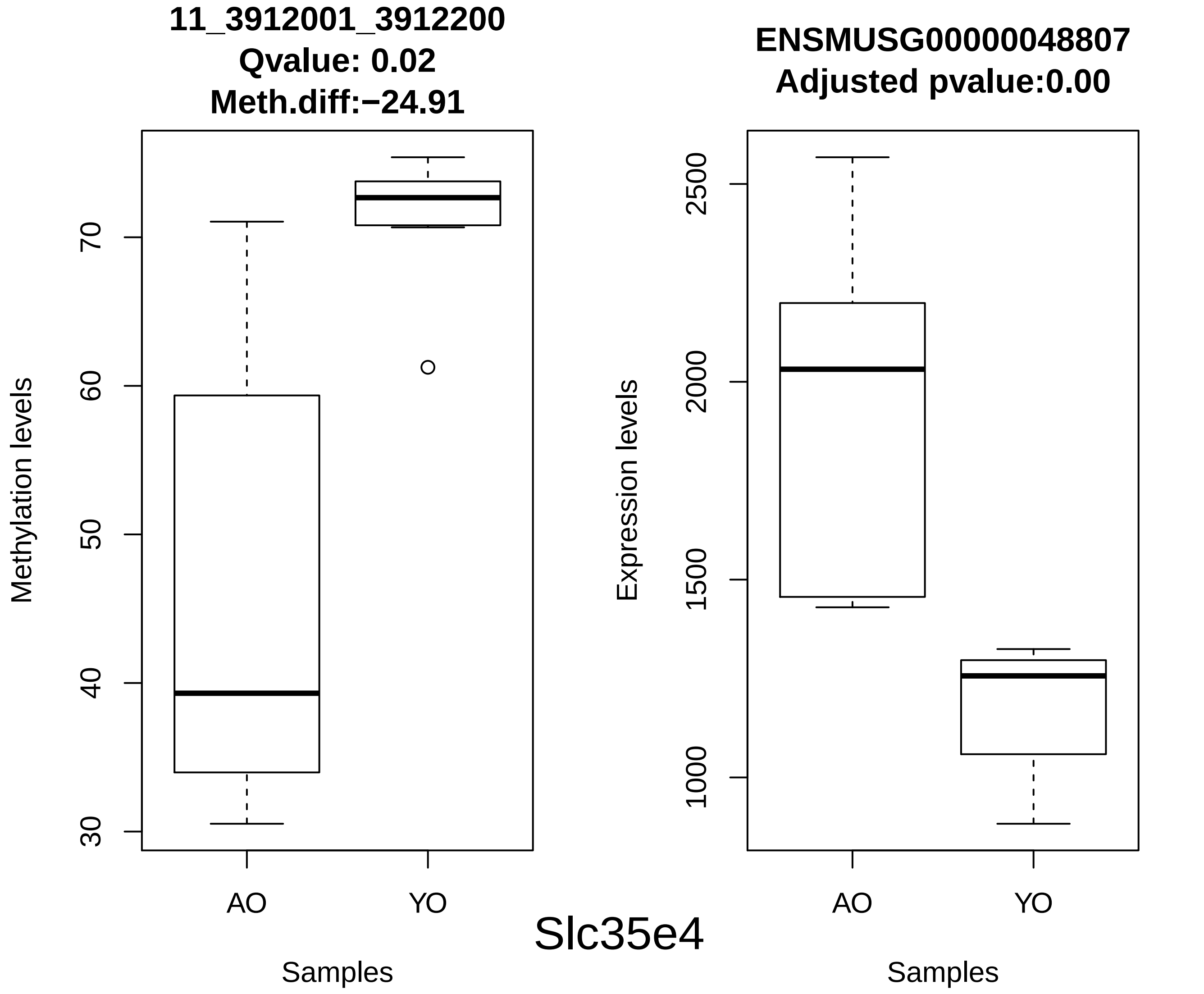
11_3912001_3912200
Qvalue: 0.02
Meth.diff:−24.91
Methylation levels
30
40
50
60
70
AO
YO
Samples
ENSMUSG00000048807
Adjusted pvalue:0.00
Expression levels
1000
1500
2000
2500
AO
YO
Samples
Slc35e4

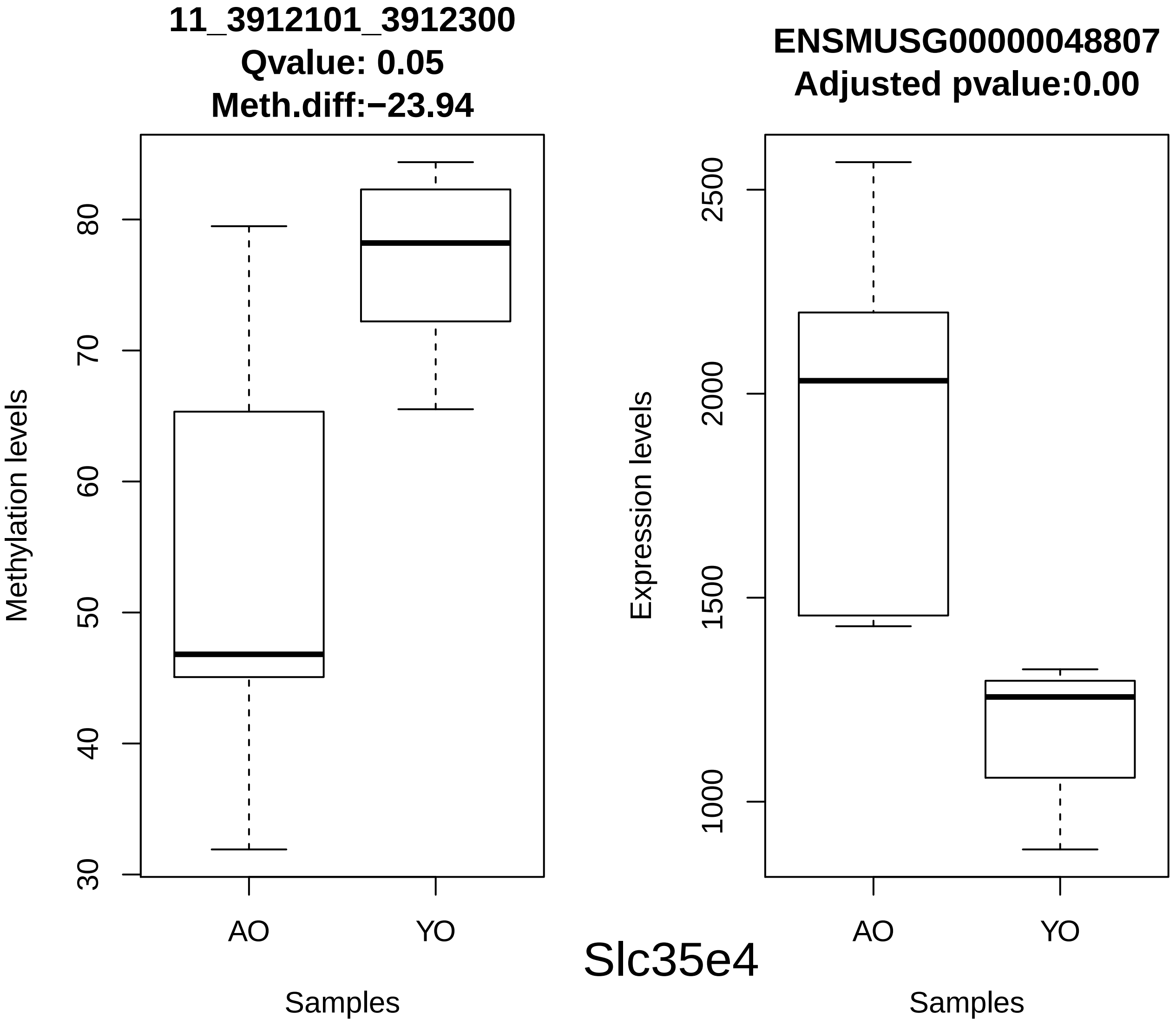
11_3912101_3912300
Qvalue: 0.05
Meth.diff:−23.94
Methylation levels
30
40
50
60
70
80
AO
YO
Samples
ENSMUSG00000048807
Adjusted pvalue:0.00
Expression levels
1000
1500
2000
2500
AO
YO
Samples
Slc35e4

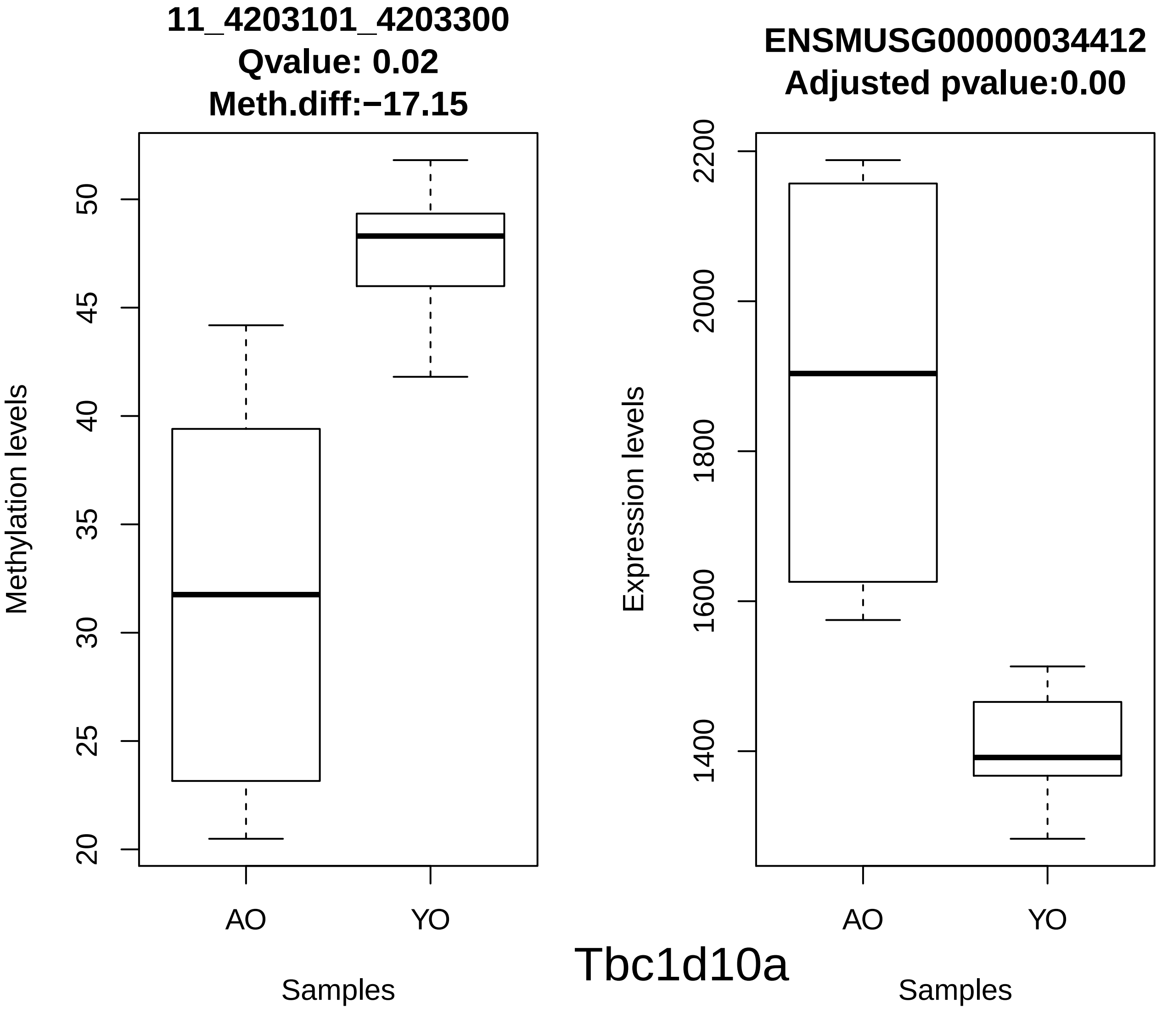
11_4203101_4203300
Qvalue: 0.02
Meth.diff:−17.15
Methylation levels
50
45
40
35
30
25
20
AO
YO
Samples
ENSMUSG00000034412
Adjusted pvalue:0.00
Expression levels
2200
2000
1800
1600
1400
AO
YO
Samples
Tbc1d10a

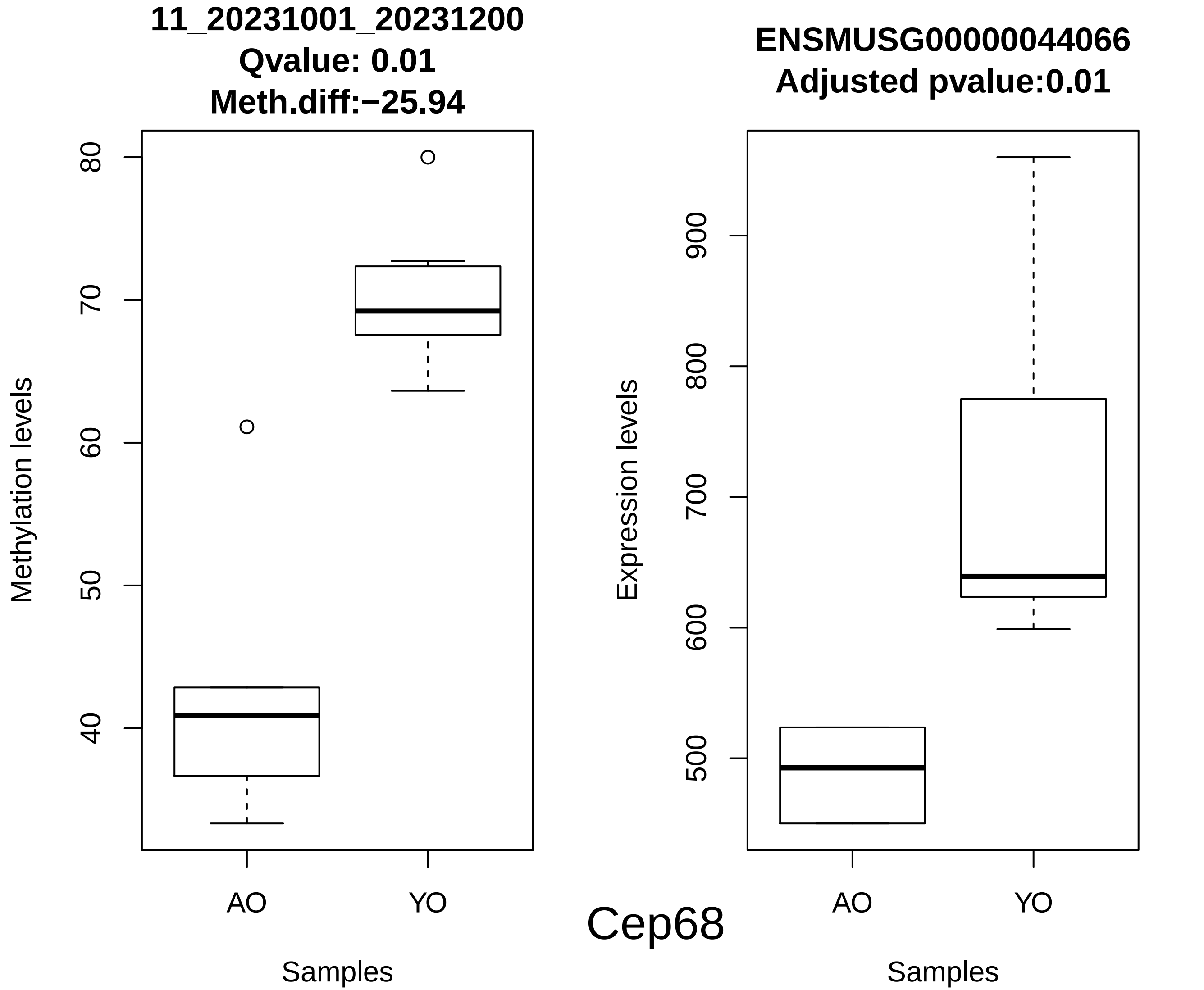
11_20231001_20231200
Qvalue: 0.01
Meth.diff:−25.94
Methylation levels
80
70
60
50
40
AO
YO
Samples
ENSMUSG00000044066
Adjusted pvalue:0.01
Expression levels
900
800
700
600
500
AO
YO
Samples
Cep68

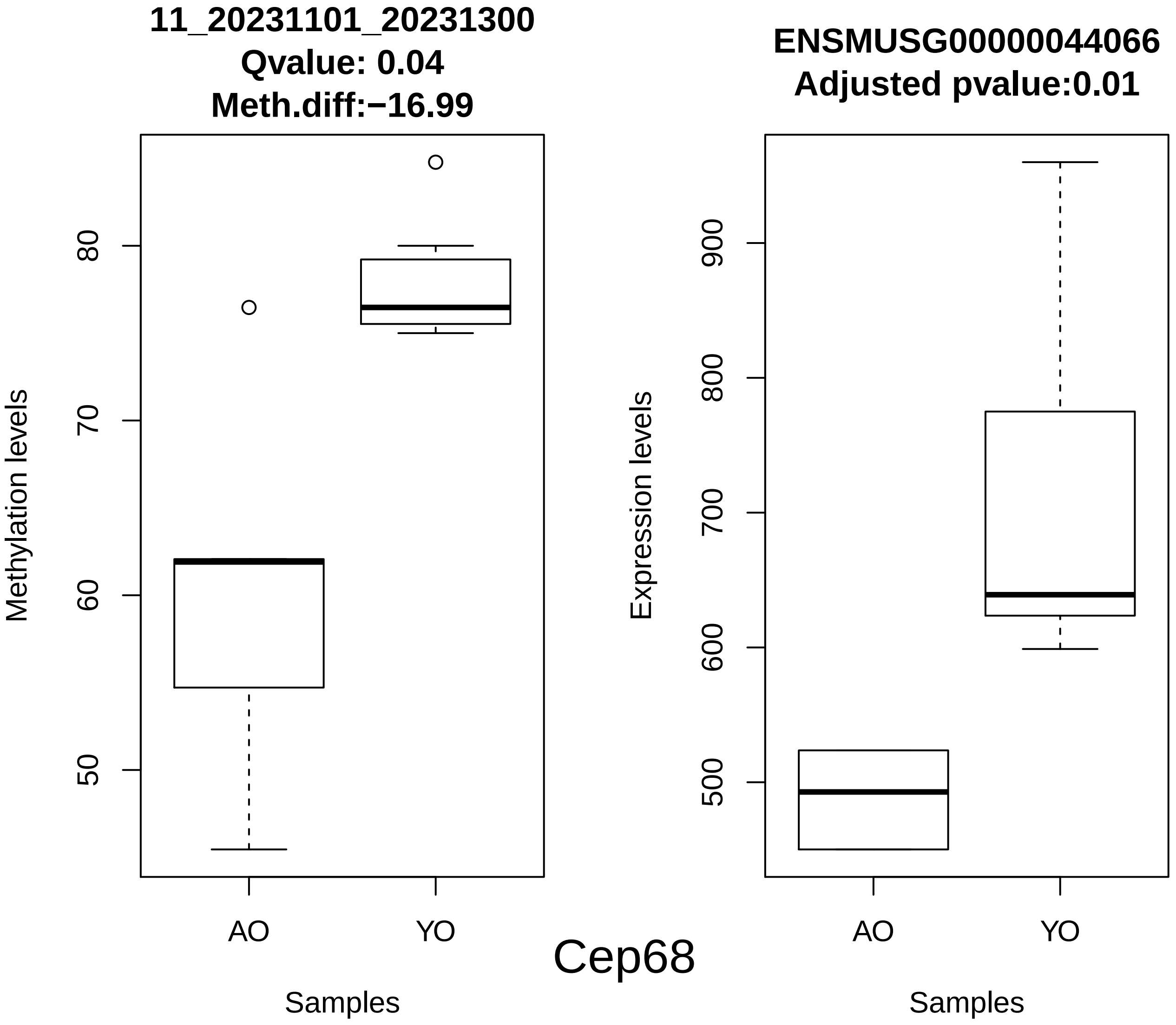
11_20231101_20231300
Qvalue: 0.04
Meth.diff:−16.99
Methylation levels
80
70
60
50
AO
YO
Samples
ENSMUSG00000044066
Adjusted pvalue:0.01
Expression levels
900
800
700
600
500
AO
YO
Samples
Cep68

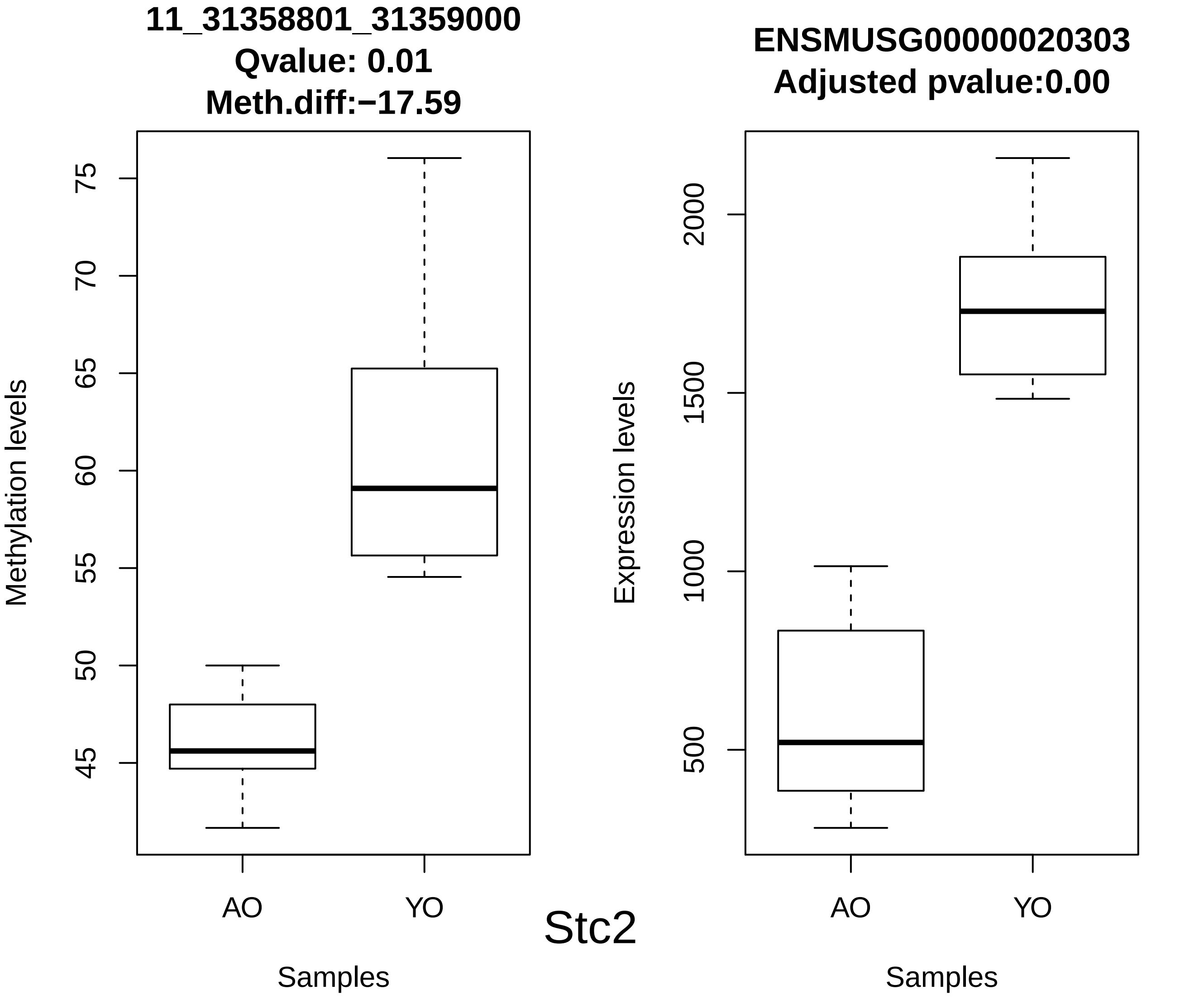
11_31358801_31359000
Qvalue: 0.01
Meth.diff:−17.59
Methylation levels
45
50
55
60
65
70
75
AO
YO
Samples
ENSMUSG00000020303
Adjusted pvalue:0.00
Expression levels
500
1000
1500
2000
AO
YO
Samples
Stc2

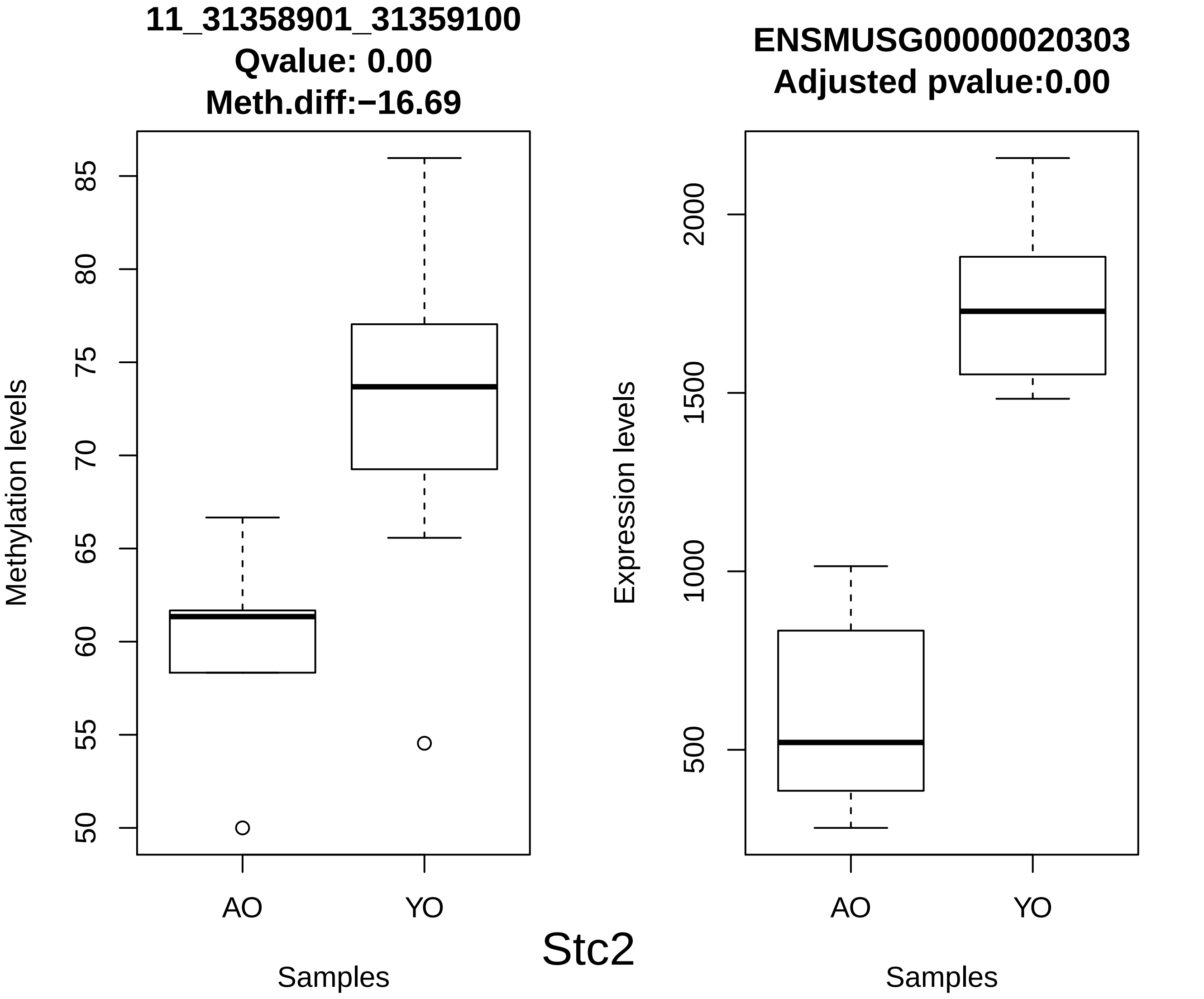
11_31358901_31359100
Qvalue: 0.00
Meth.diff:−16.69
Methylation levels
50
55
60
65
70
75
80
85
AO
YO
Samples
ENSMUSG00000020303
Adjusted pvalue:0.00
Expression levels
500
1000
1500
2000
AO
YO
Samples
Stc2

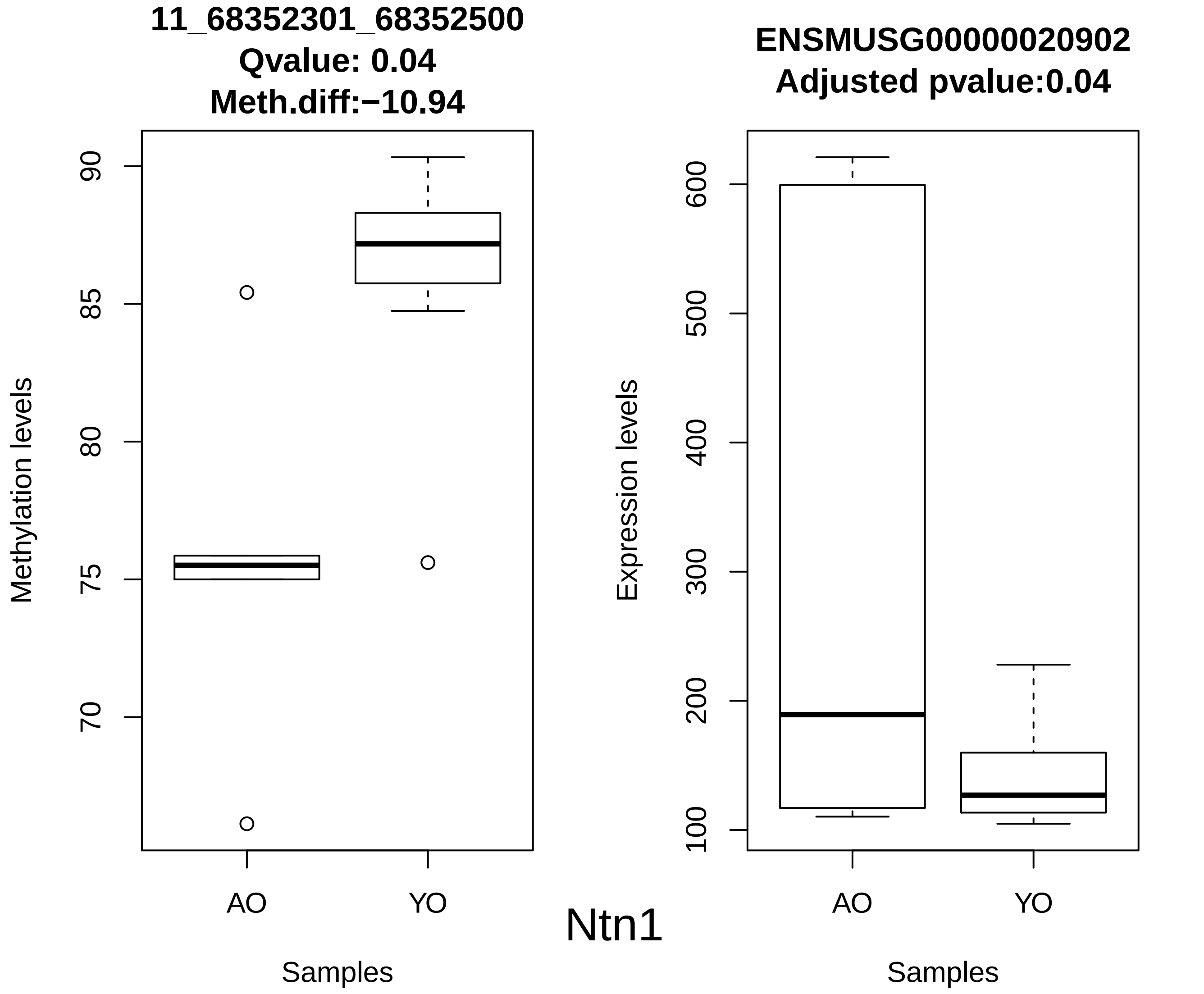
11_68352301_68352500
Qvalue: 0.04
Meth.diff:−10.94
Methylation levels
90
85
80
75
70
AO
YO
Samples
ENSMUSG00000020902
Adjusted pvalue:0.04
Expression levels
600
500
400
300
200
100
AO
YO
Samples
Ntn1

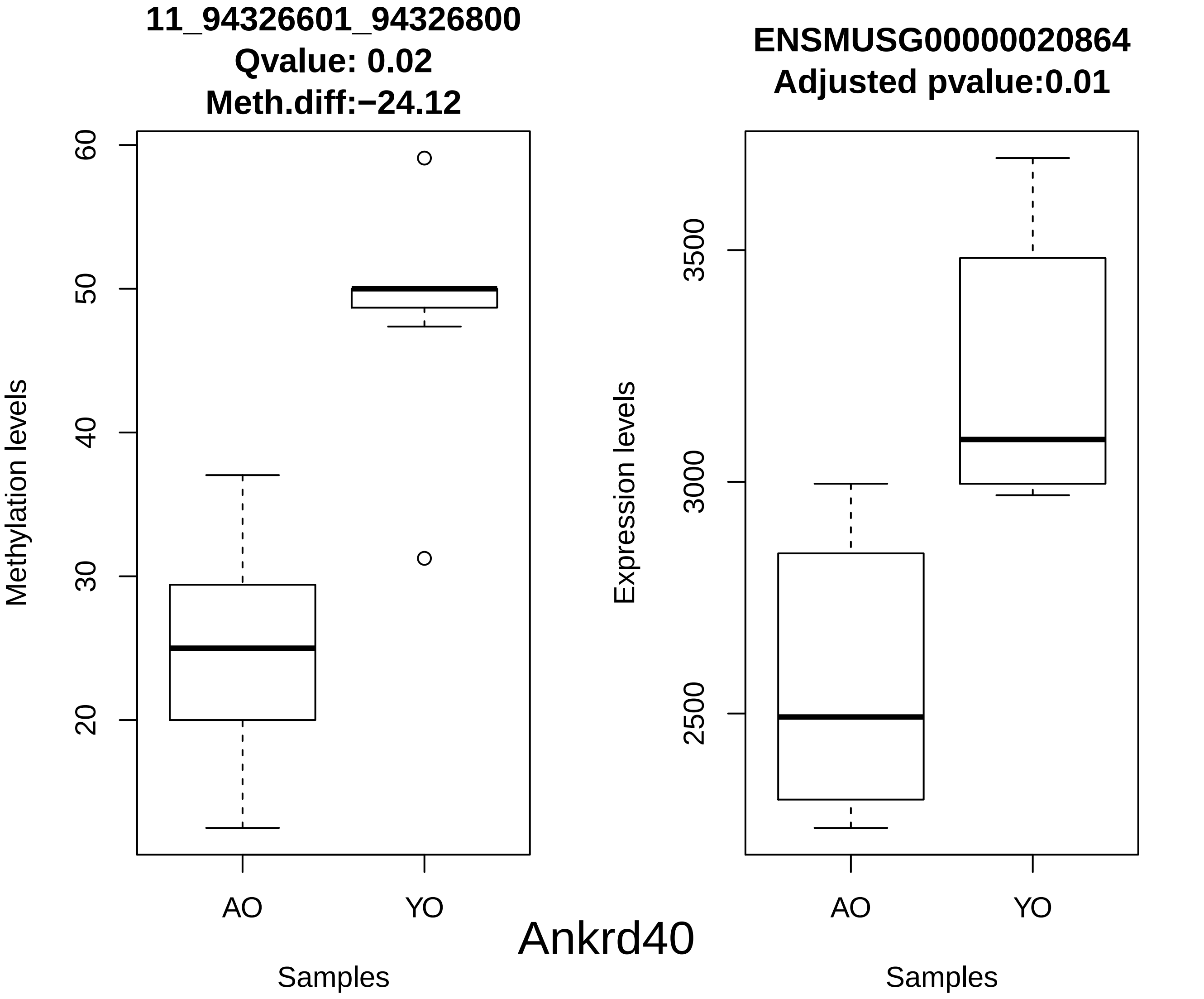
11_94326601_94326800
Qvalue: 0.02
Meth.diff:−24.12
Methylation levels
60
50
40
30
20
AO
YO
Samples
ENSMUSG00000020864
Adjusted pvalue:0.01
Expression levels
3500
3000
2500
AO
YO
Samples
Ankrd40

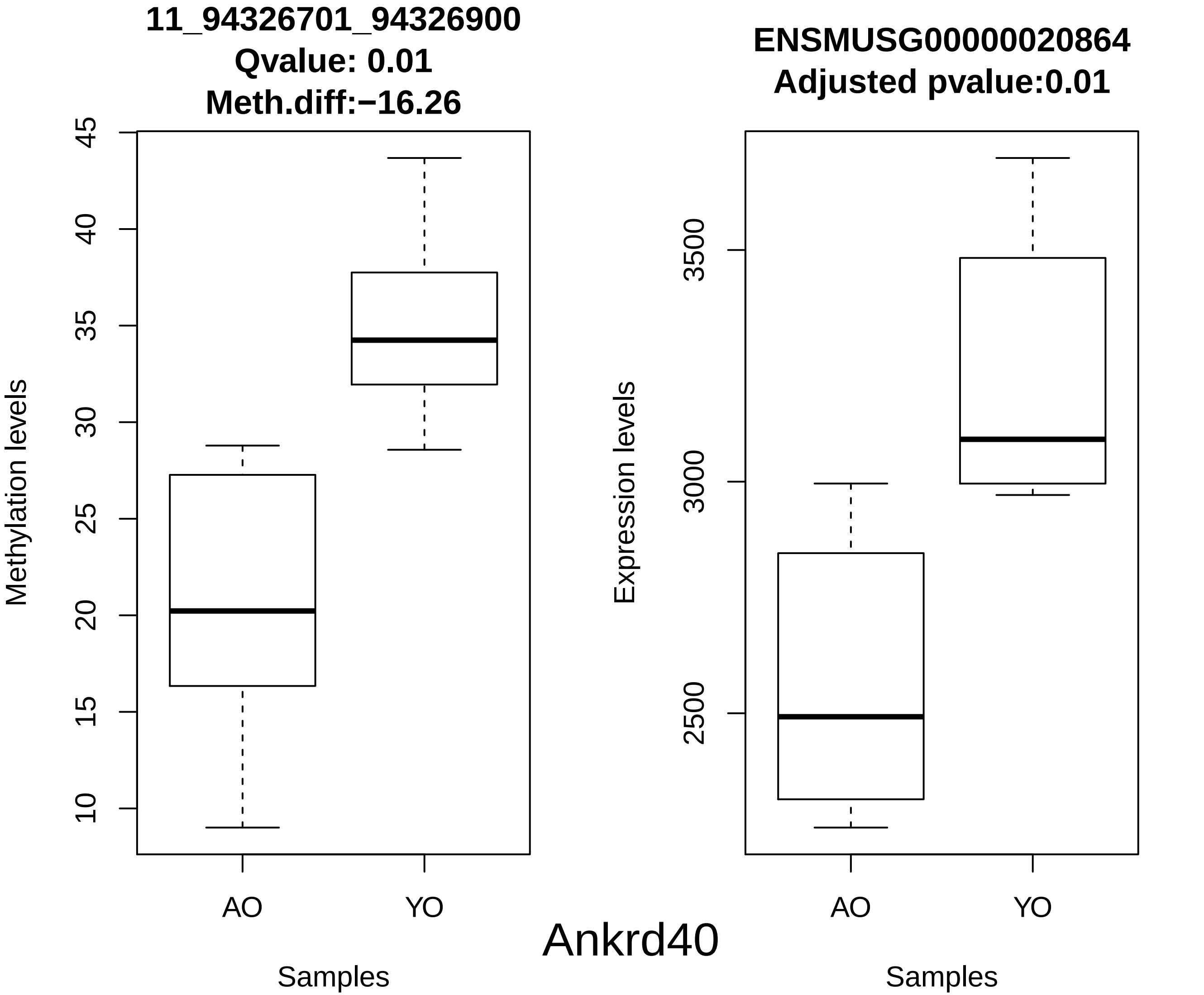
11_94326701_94326900
Qvalue: 0.01
Meth.diff:−16.26
Methylation levels
45
40
35
30
25
20
15
10
AO
YO
Samples
ENSMUSG00000020864
Adjusted pvalue:0.01
Expression levels
3500
3000
2500
AO
YO
Samples
Ankrd40

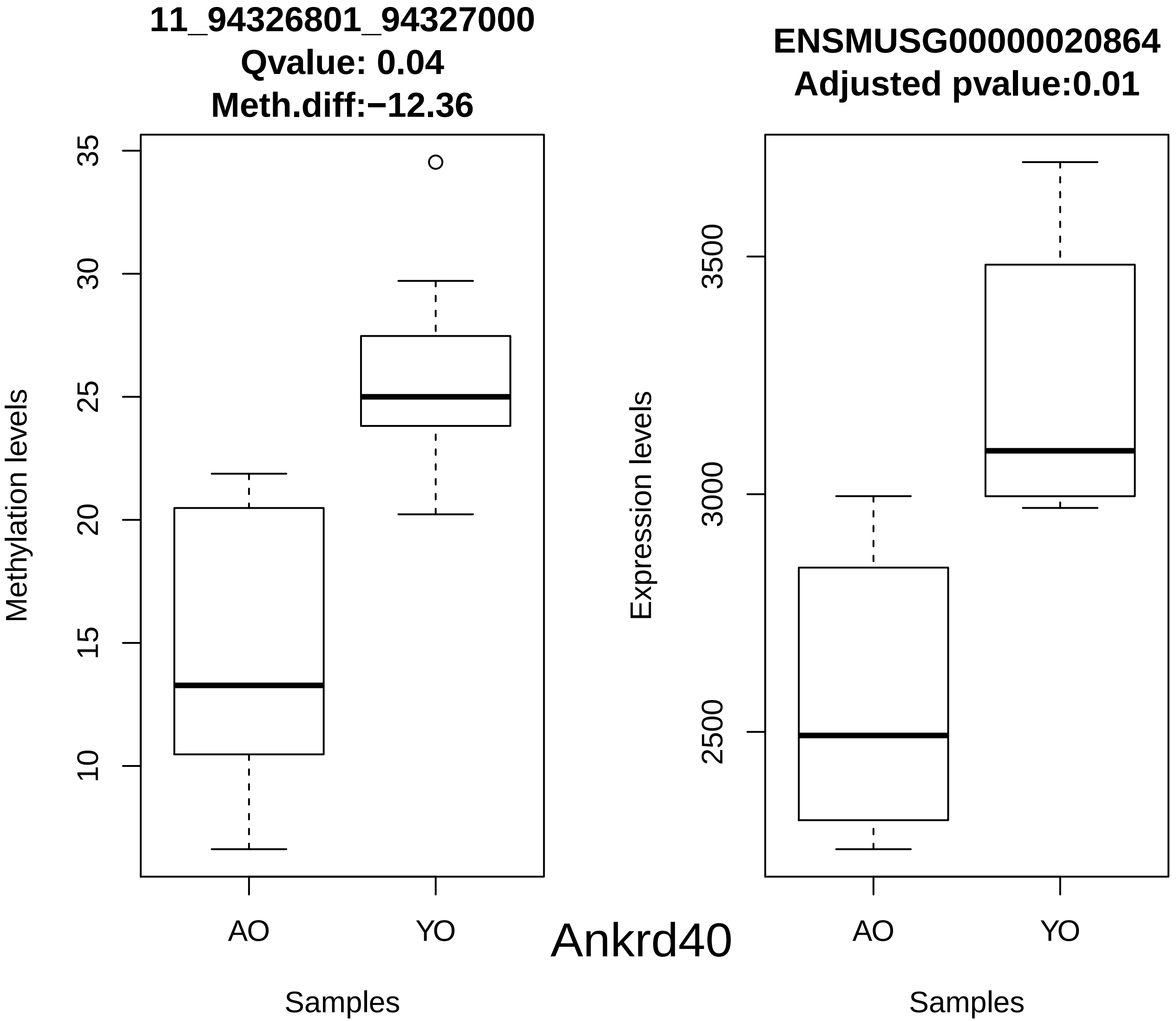

11_94326801_94327000
Qvalue: 0.04
Meth.diff:−12.36
Methylation levels
35
30
25
20
15
10
AO
YO
Samples
ENSMUSG00000020864
Adjusted pvalue:0.01
Expression levels
3500
3000
2500
AO
YO
Samples


Ankrd40

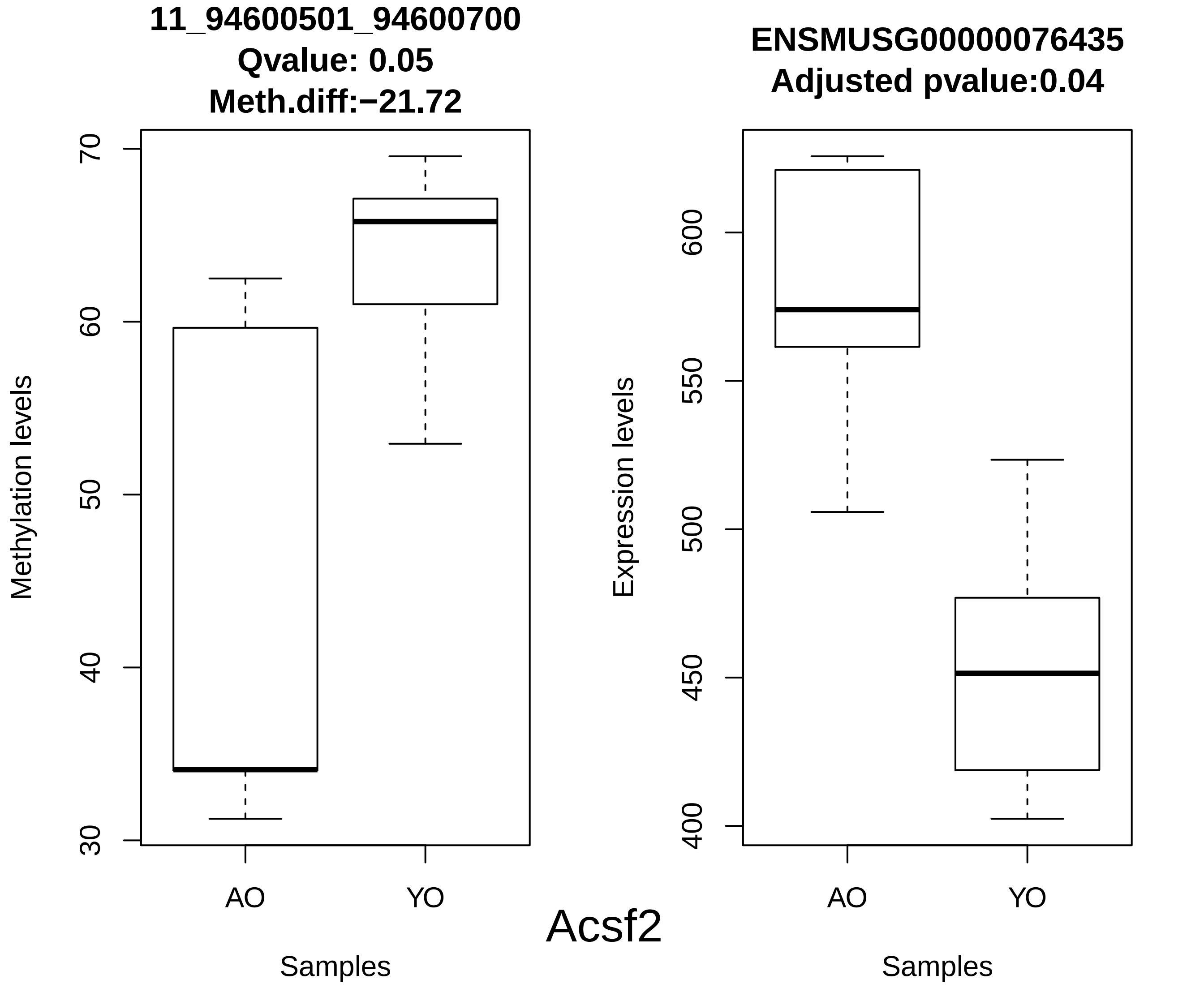
11_94600501_94600700
Qvalue: 0.05
Meth.diff:−21.72
Methylation levels
70
60
50
40
30
AO
YO
Samples
ENSMUSG00000076435
Adjusted pvalue:0.04
Expression levels
600
550
500
450
400
AO
YO
Samples
Acsf2

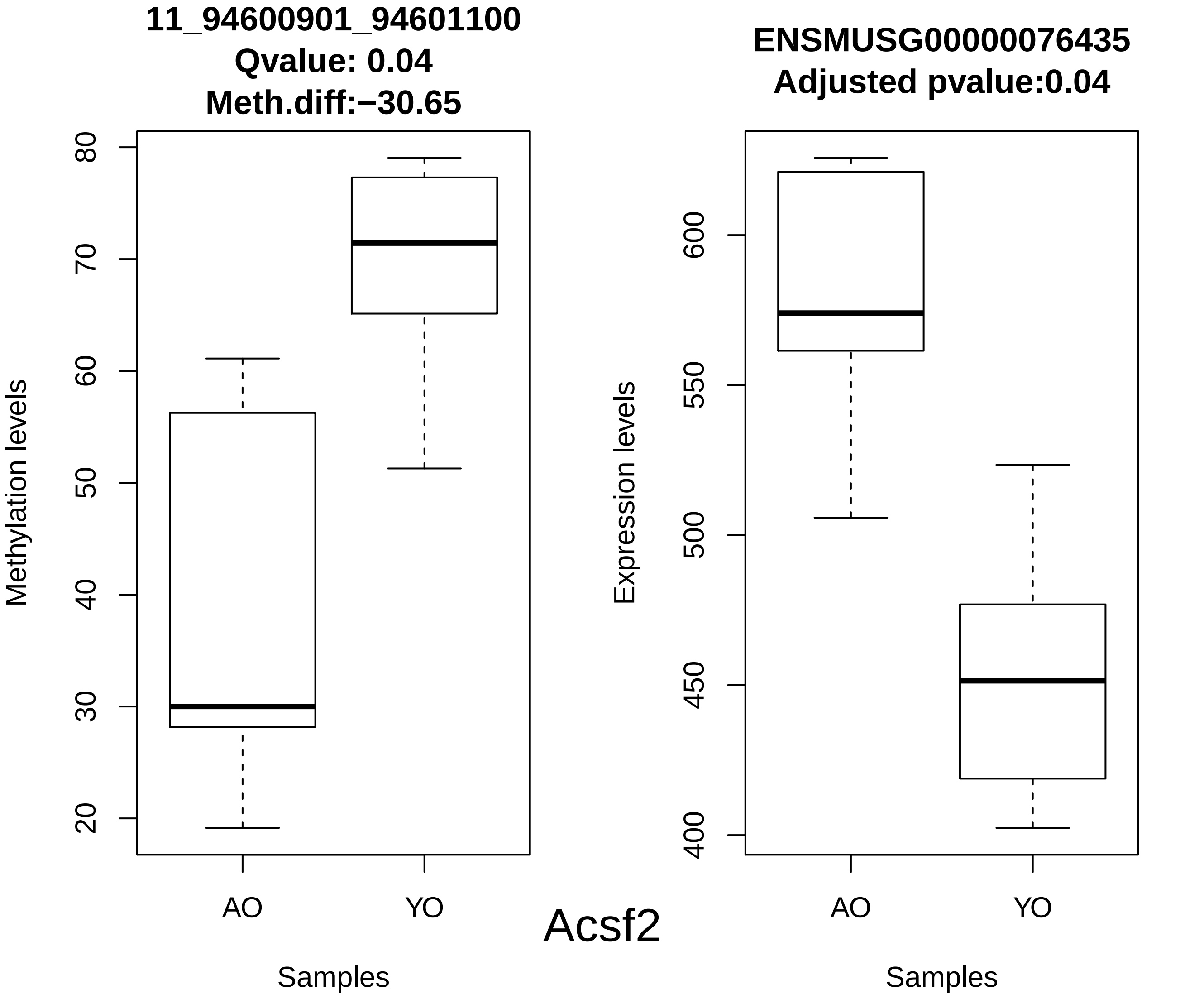
11_94600901_94601100
Qvalue: 0.04
Meth.diff:−30.65
Methylation levels
80
70
60
50
40
30
20
AO
YO
Samples
Acsf2
ENSMUSG00000076435
Adjusted pvalue:0.04
Expression levels
600
550
500
450
400
AO
YO
Samples

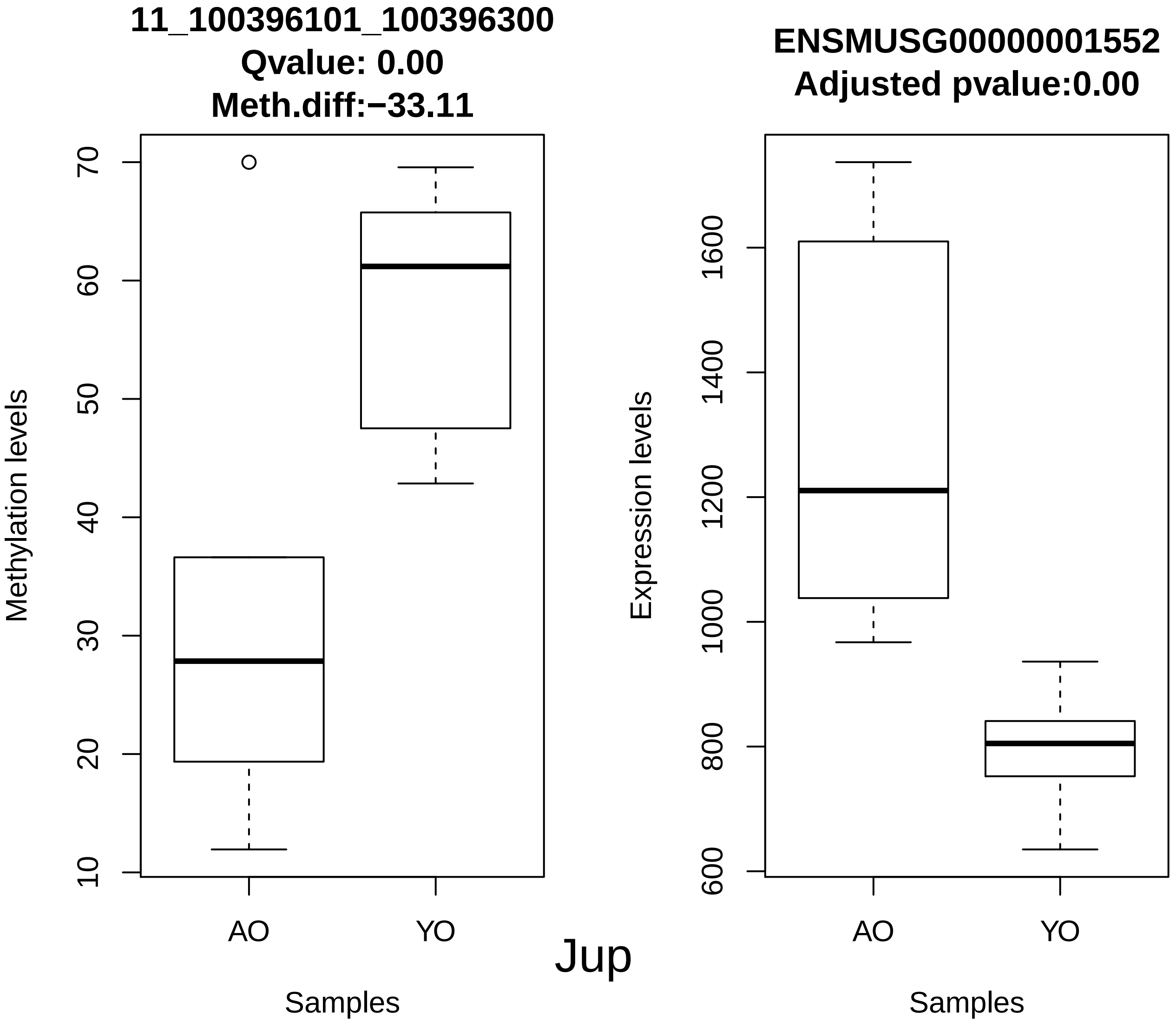
11_100396101_100396300
Qvalue: 0.00
Meth.diff:−33.11
Methylation levels
10
20
30
40
50
60
70
AO
YO
Samples
ENSMUSG00000001552
Adjusted pvalue:0.00
Expression levels
600
800
1000
1200
1400
1600
AO
YO
Samples
Jup

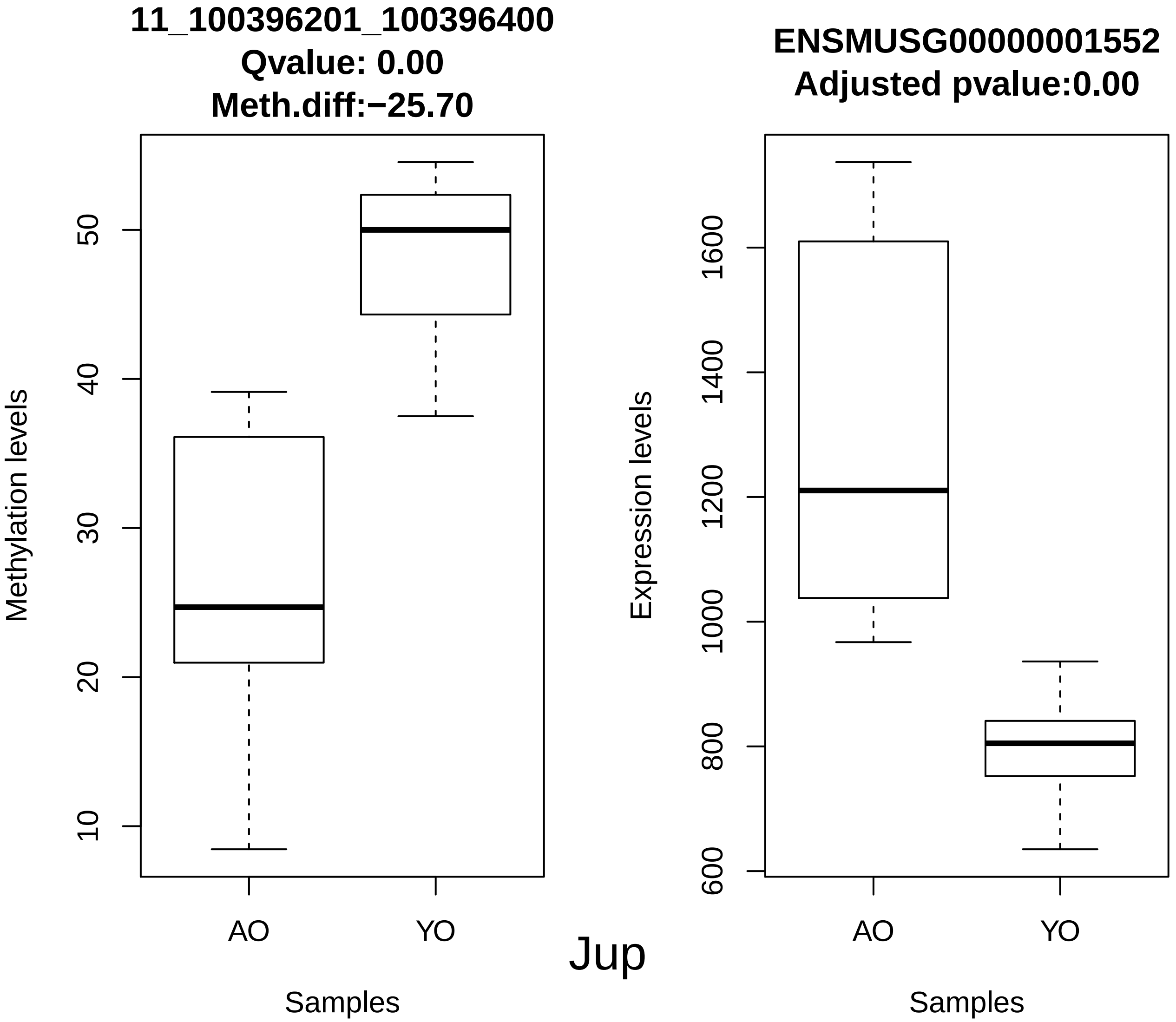

11_100396201_100396400
Qvalue: 0.00
Meth.diff:−25.70
Methylation levels
10
20
30
40
50
AO
YO
Samples
ENSMUSG00000001552
Adjusted pvalue:0.00
Expression levels
600
800
1000
1200
1400
1600
AO
YO
Samples


Jup

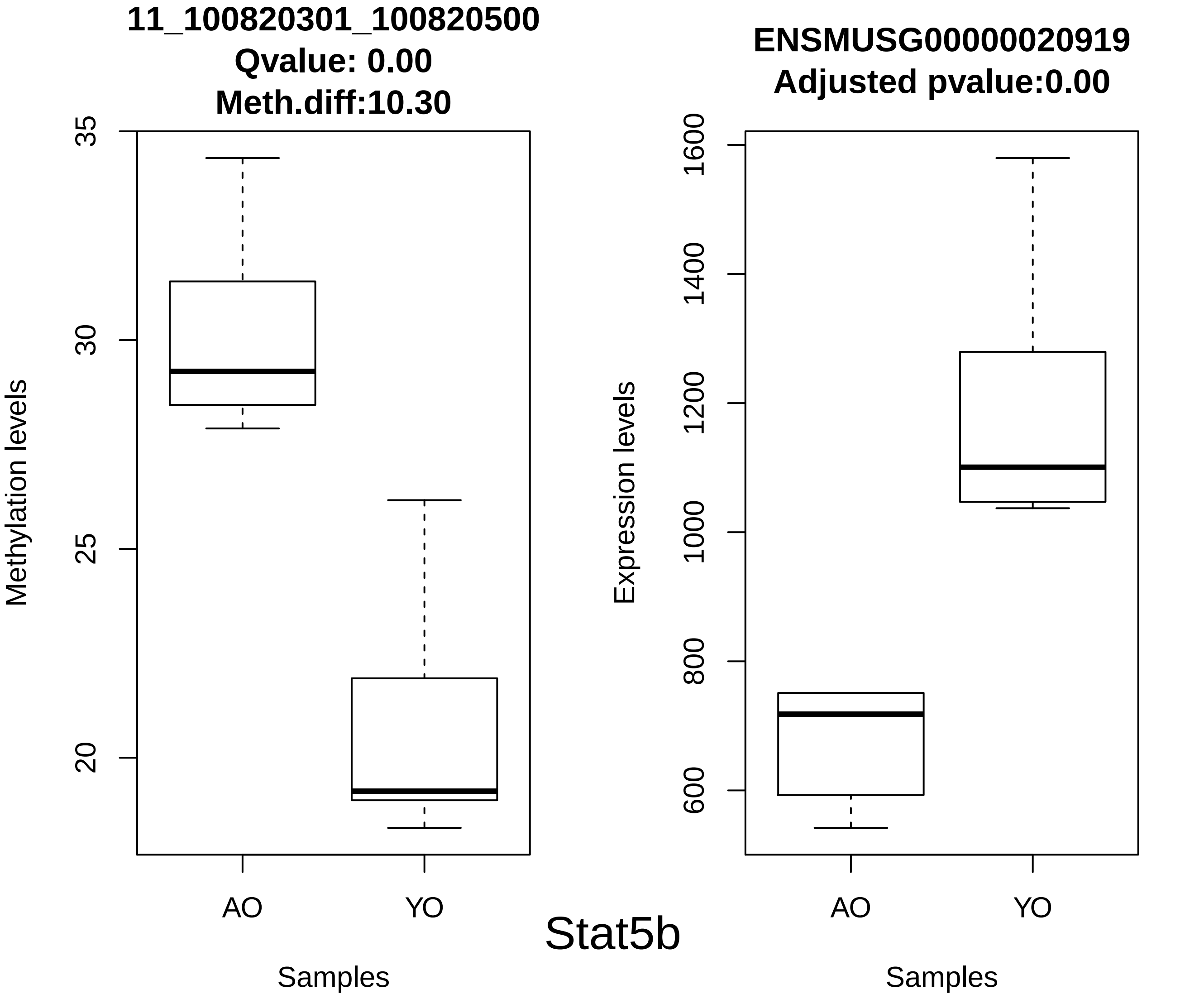
11_100820301_100820500
Qvalue: 0.00
Meth.diff:10.30
Methylation levels
35
30
25
20
AO
YO
Samples
ENSMUSG00000020919
Adjusted pvalue:0.00
Expression levels
1600
1400
1200
1000
800
600
AO
YO
Samples
Stat5b

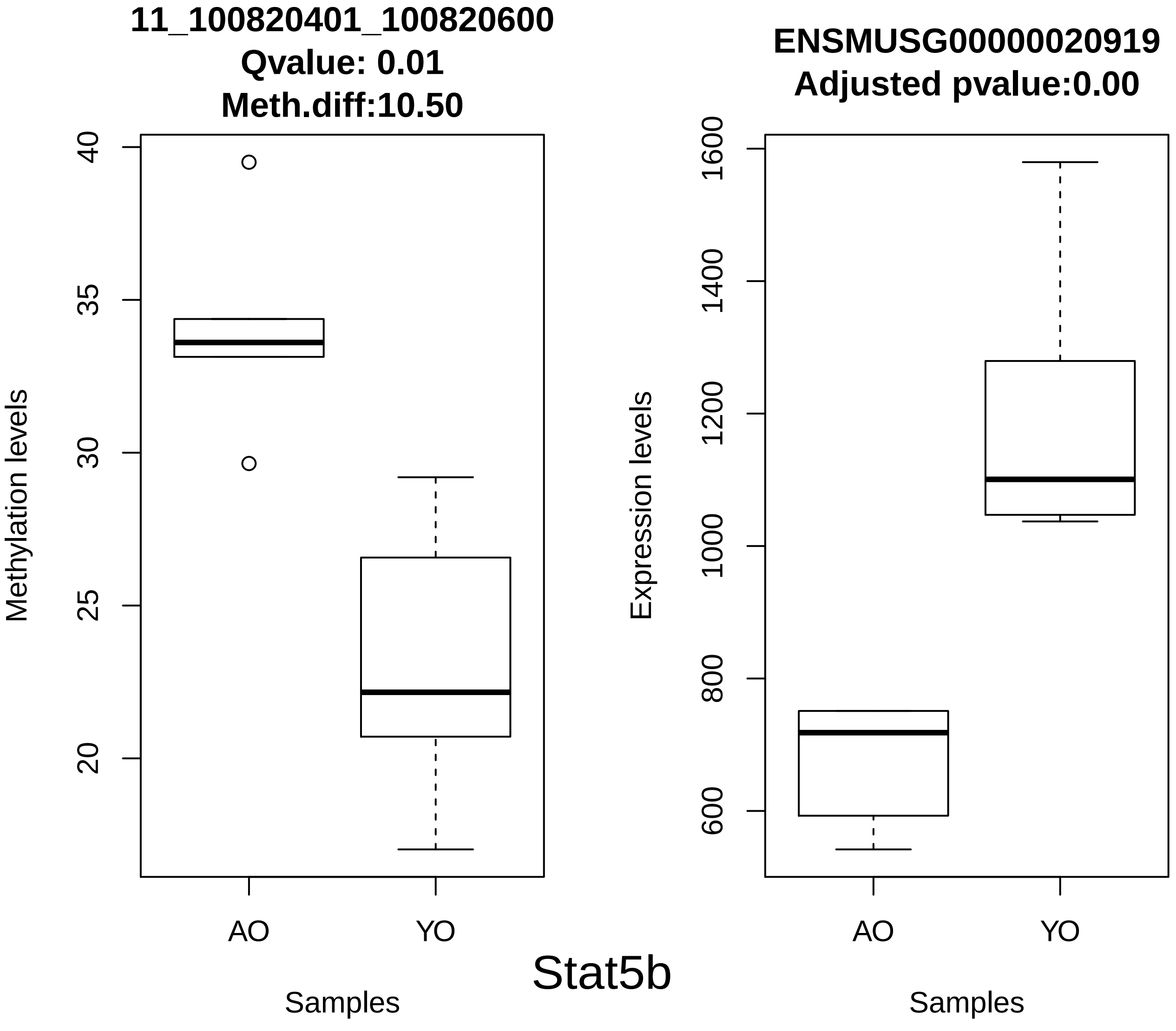
11_100820401_100820600
Qvalue: 0.01
Meth.diff:10.50
Methylation levels
40
35
30
25
20
AO
YO
Samples
ENSMUSG00000020919
Adjusted pvalue:0.00
Expression levels
1600
1400
1200
1000
800
600
AO
YO
Samples

Stat5b

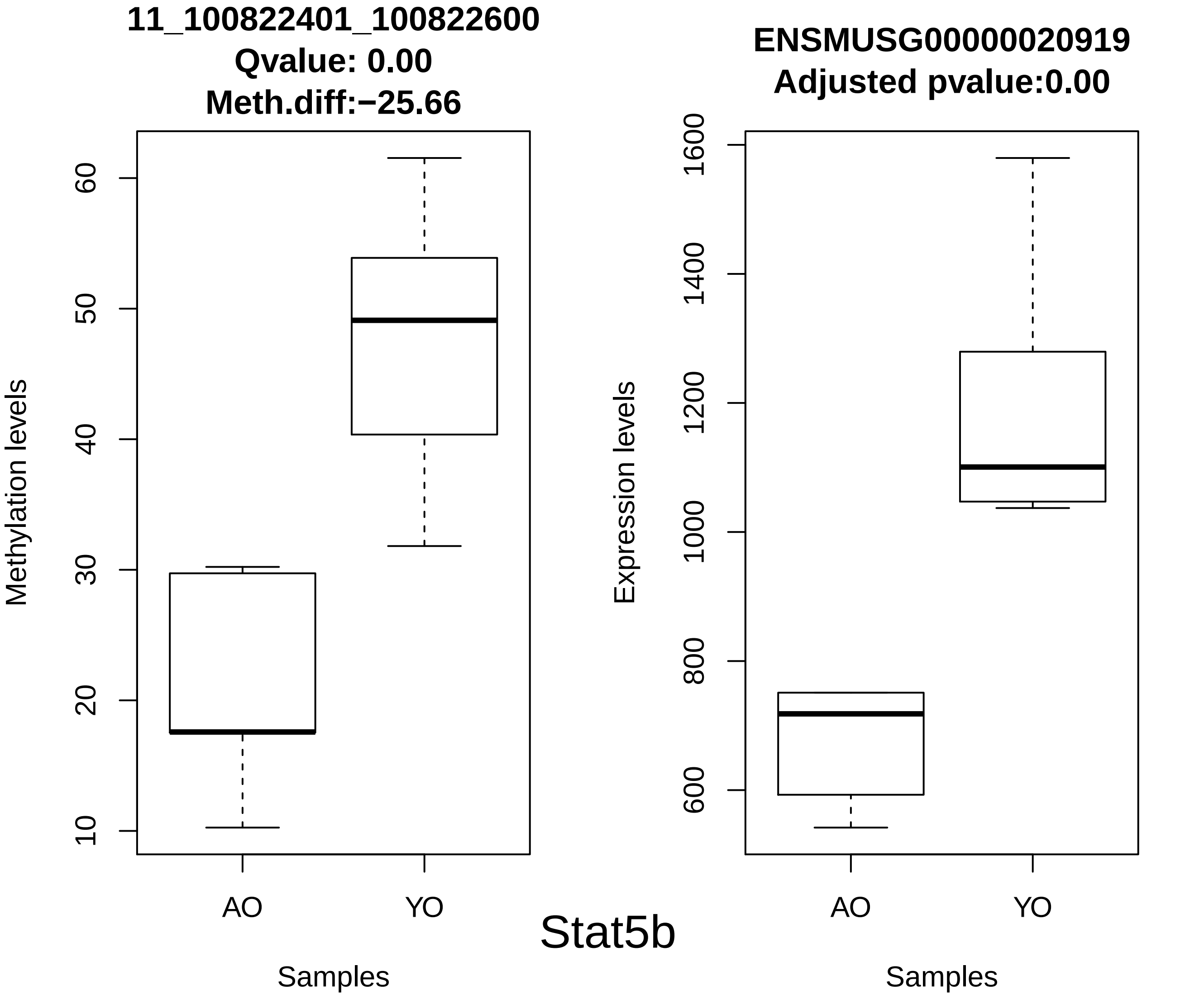
11_100822401_100822600
Qvalue: 0.00
Meth.diff:−25.66
Methylation levels
10
20
30
40
50
60
AO
YO
Samples
ENSMUSG00000020919
Adjusted pvalue:0.00
Expression levels
600
800
1000
1200
1400
1600
AO
YO
Samples

Stat5b

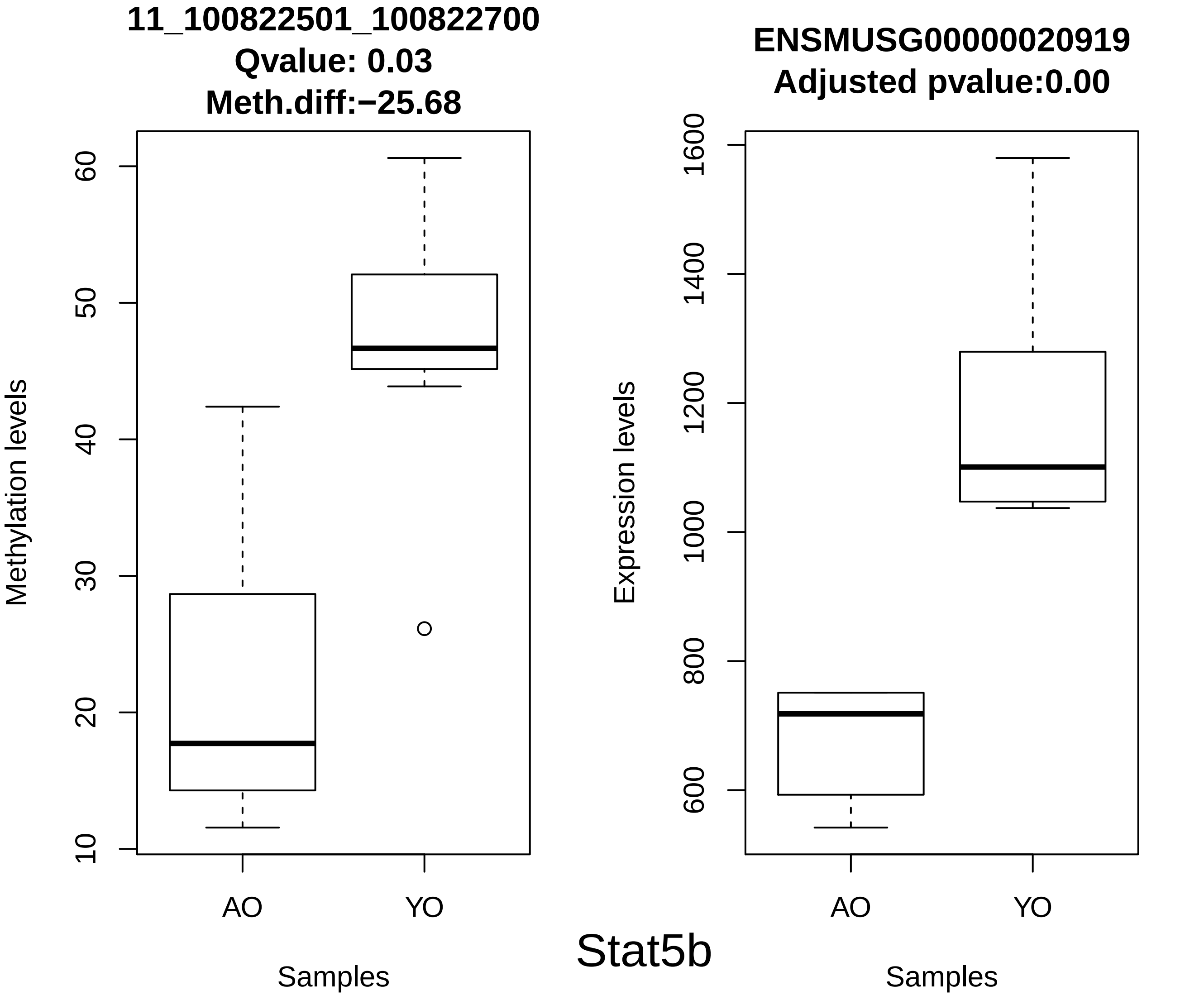
11_100822501_100822700
Qvalue: 0.03
Meth.diff:−25.68
Methylation levels
10
20
30
40
50
60
AO
YO
Samples
ENSMUSG00000020919
Adjusted pvalue:0.00
Expression levels
600
800
1000
1200
1400
1600
AO
YO
Samples
Stat5b

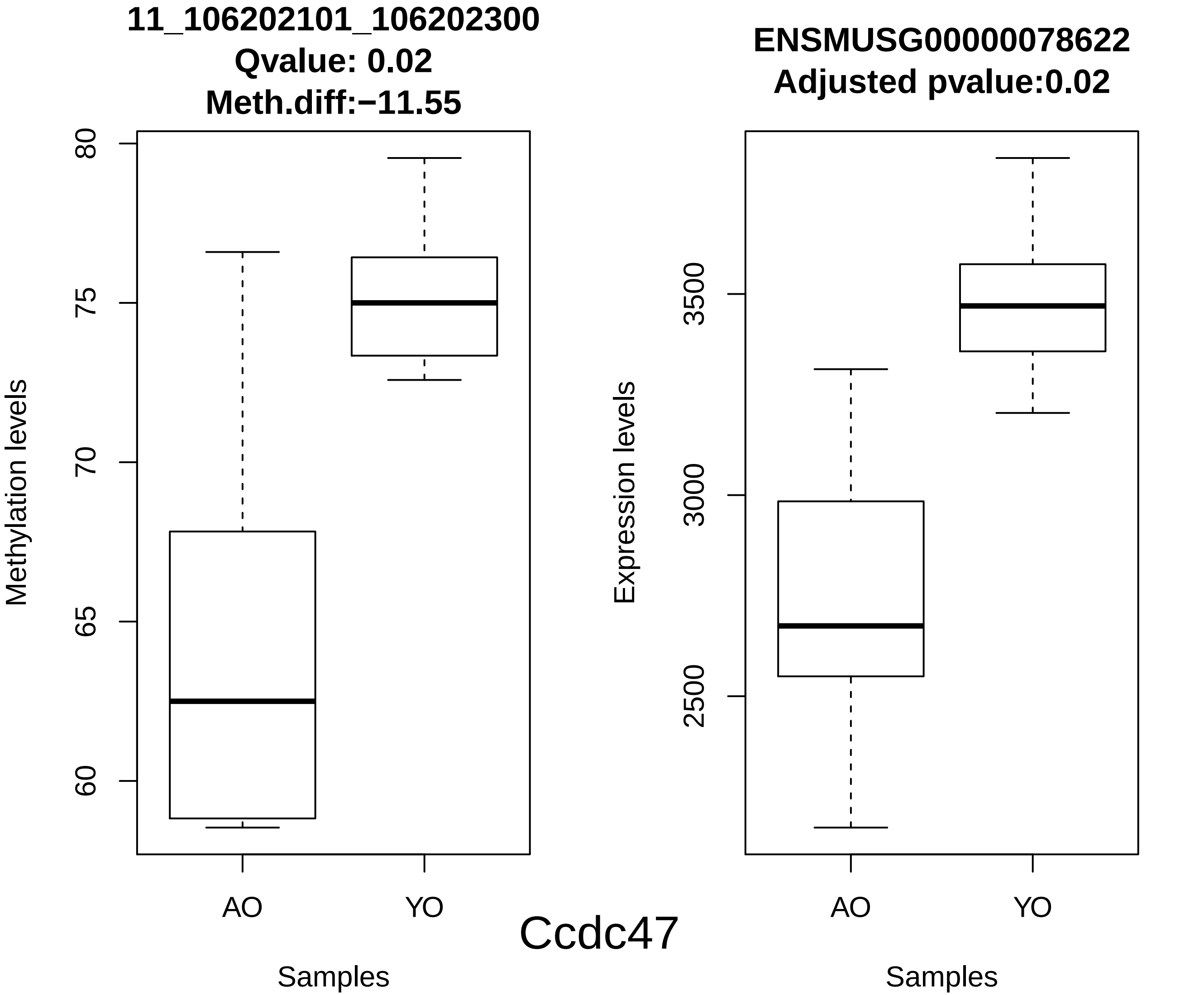
11_106202101_106202300
Qvalue: 0.02
Meth.diff:−11.55
Methylation levels
80
75
70
65
60
AO
YO
Samples
ENSMUSG00000078622
Adjusted pvalue:0.02
Expression levels
3500
3000
2500
AO
YO
Samples
Ccdc47

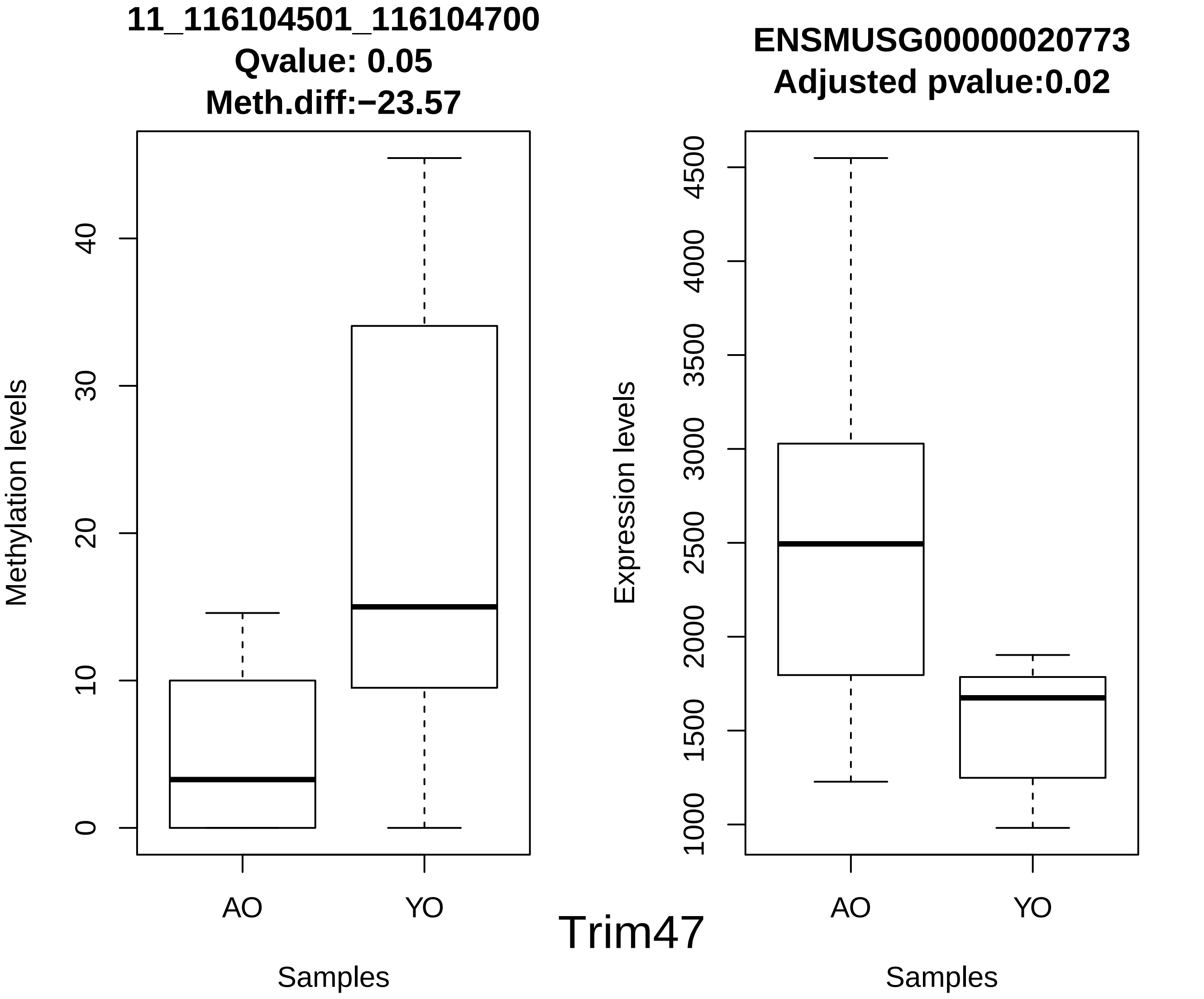
11_116104501_116104700
Qvalue: 0.05
Meth.diff:−23.57
Methylation levels
0
10
20
30
40
AO
YO
Samples
ENSMUSG00000020773
Adjusted pvalue:0.02
Expression levels
1000
1500
2000
2500
3000
3500
4000
4500
AO
YO
Samples
Trim47

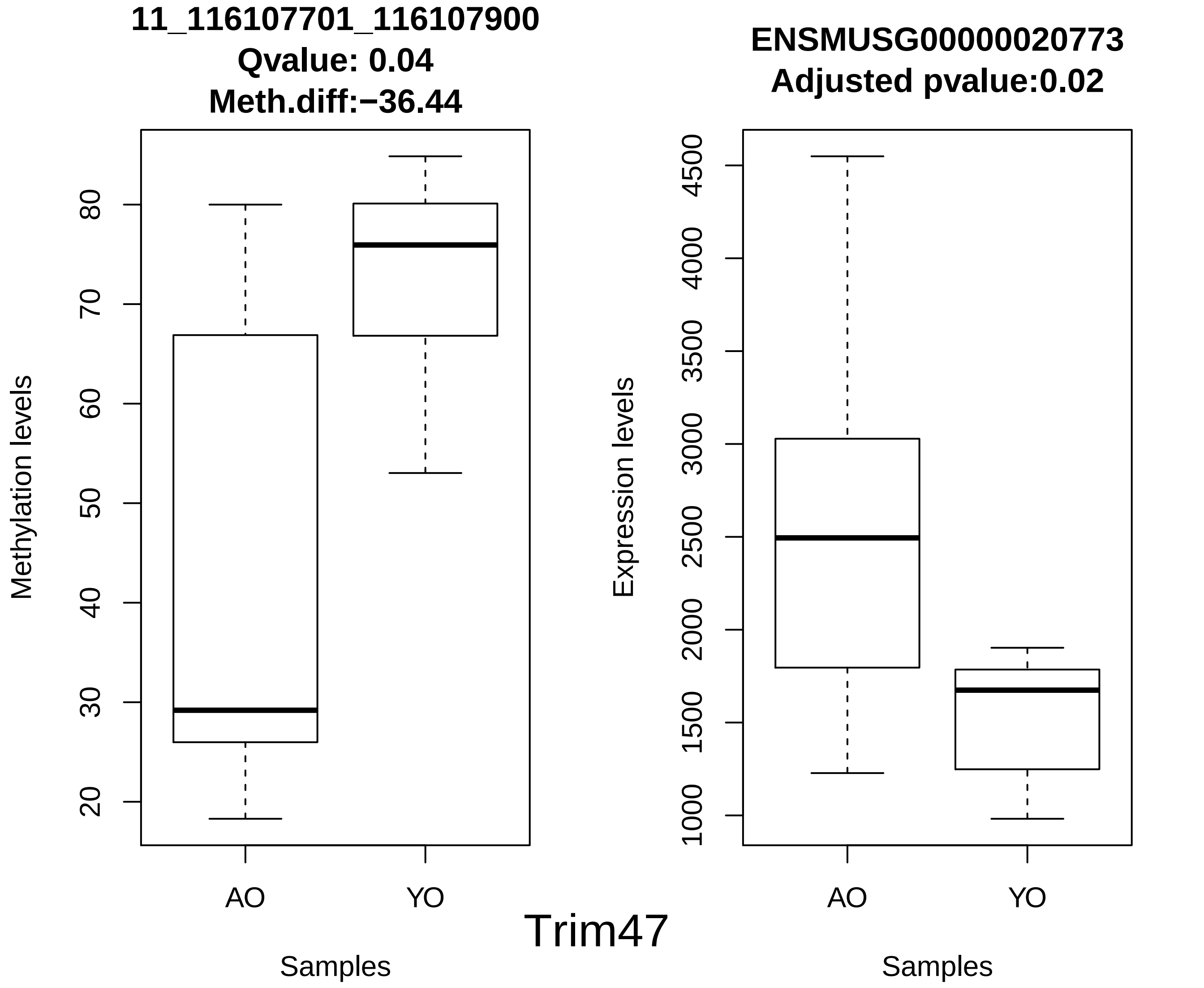
11_116107701_116107900
Qvalue: 0.04
Meth.diff:−36.44
Methylation levels
20
30
40
50
60
70
80
AO
YO
Samples
ENSMUSG00000020773
Adjusted pvalue:0.02
Expression levels
1000
1500
2000
2500
3000
3500
4000
4500
AO
YO
Samples
Trim47

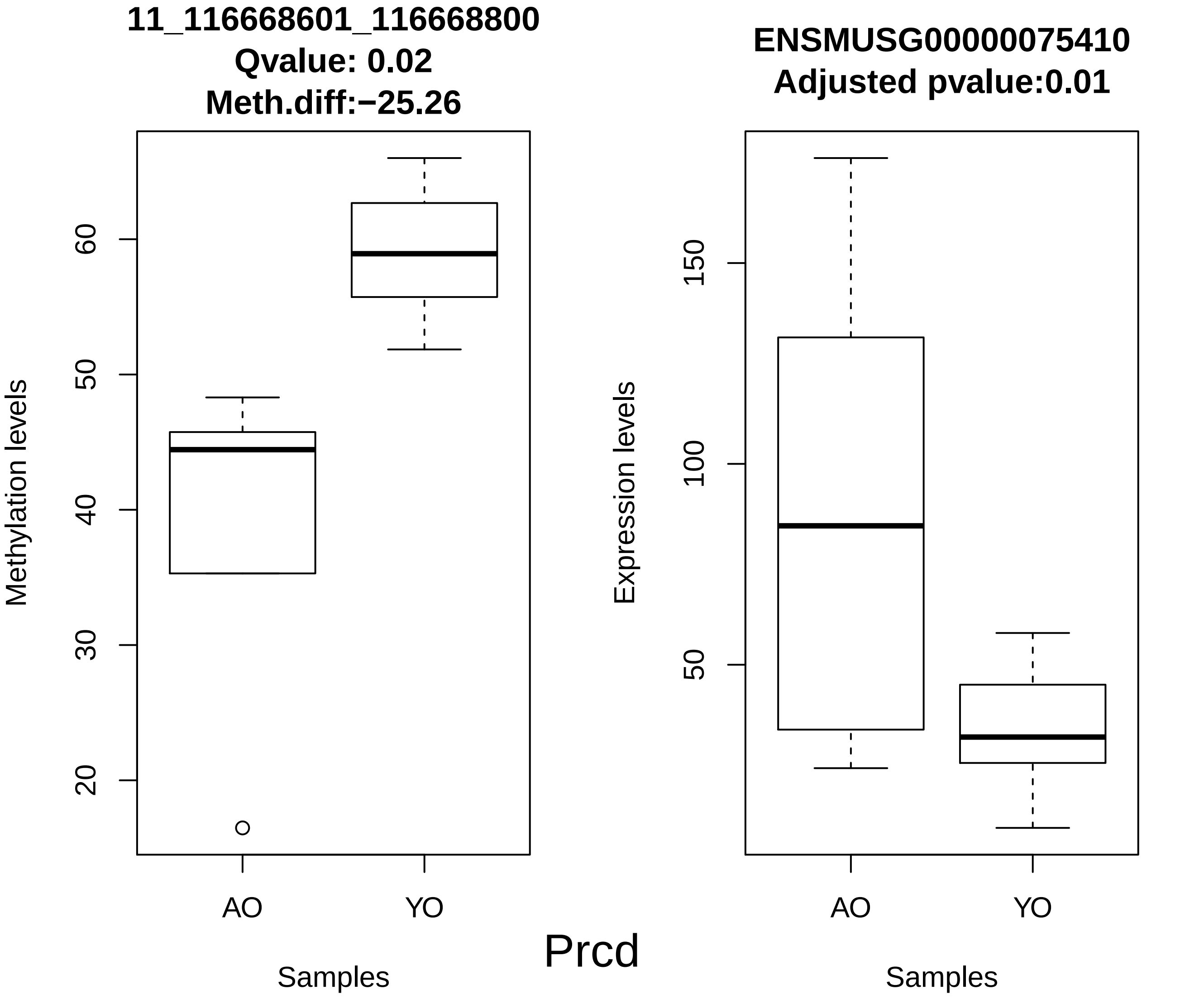
11_116668601_116668800
Qvalue: 0.02
Meth.diff:−25.26
Methylation levels
20
30
40
50
60
AO
YO
Samples
ENSMUSG00000075410
Adjusted pvalue:0.01
Expression levels
50
100
150
AO
YO
Samples
Prcd

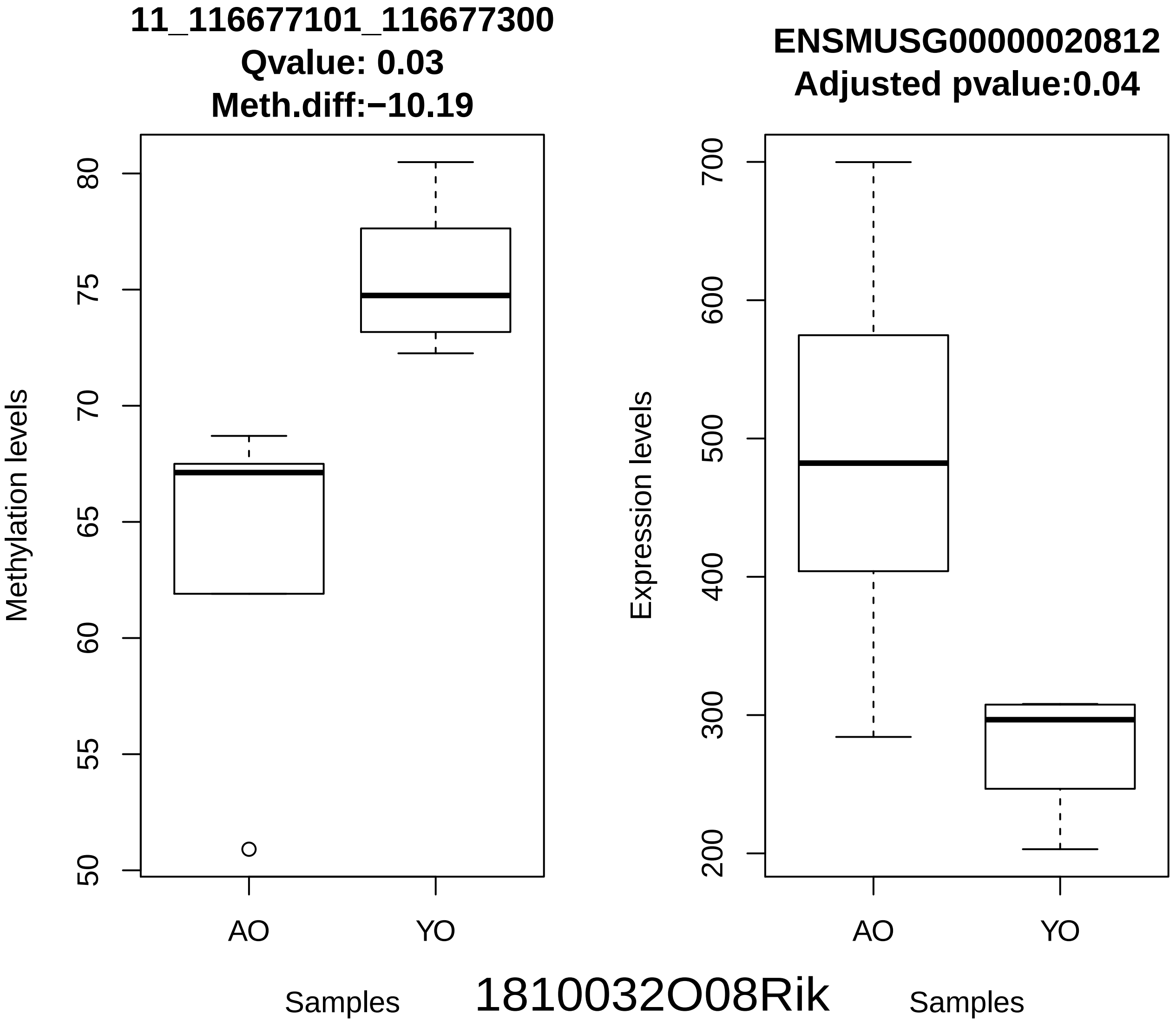
11_116677101_116677300
Qvalue: 0.03
Meth.diff:−10.19
Methylation levels
80
75
70
65
60
55
50
AO
YO
Samples
ENSMUSG00000020812
Adjusted pvalue:0.04
Expression levels
700
600
500
400
300
200
AO
YO
Samples
1810032O08Rik

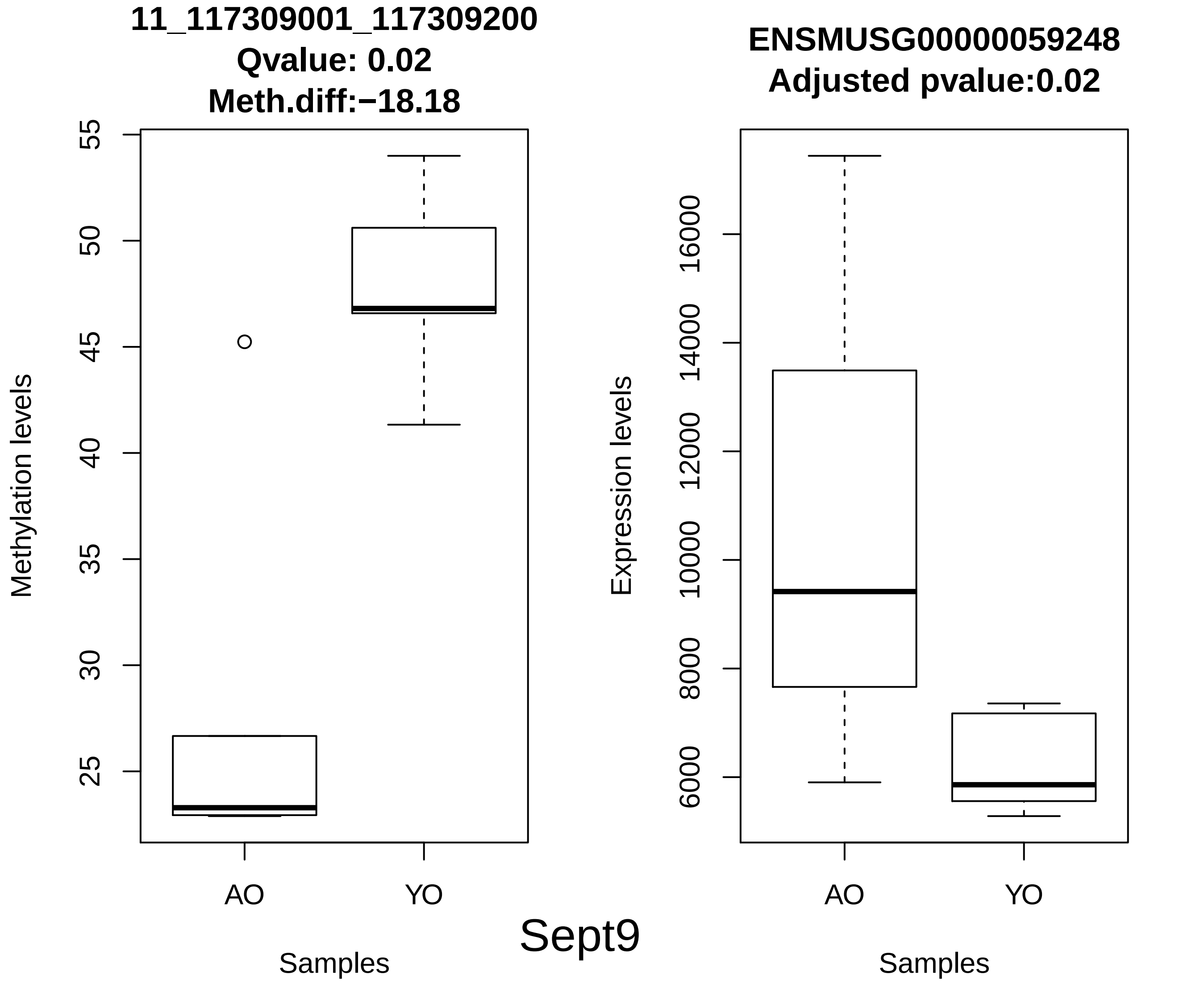
11_117309001_117309200
Qvalue: 0.02
Meth.diff:−18.18
Methylation levels
55
50
45
40
35
30
25
AO
YO
Samples
ENSMUSG00000059248
Adjusted pvalue:0.02
Expression levels
16000
14000
12000
10000
8000
6000
AO
YO
Samples
Sept9

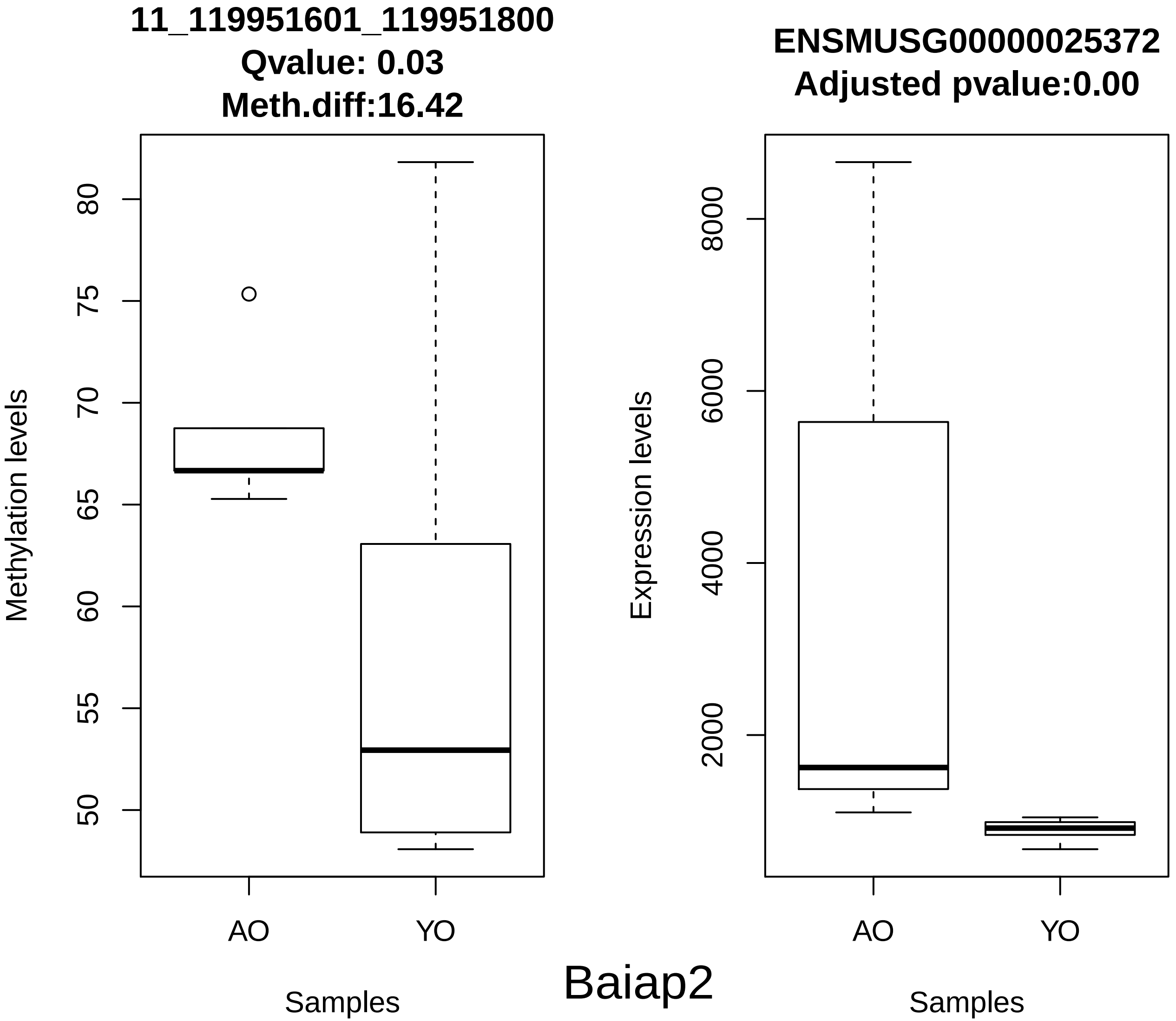
11_119951601_119951800
Qvalue: 0.03
Meth.diff:16.42
Methylation levels
80
75
70
65
60
55
50
AO
YO
Samples
ENSMUSG00000025372
Adjusted pvalue:0.00
Expression levels
8000
6000
4000
2000
AO
YO
Samples
Baiap2

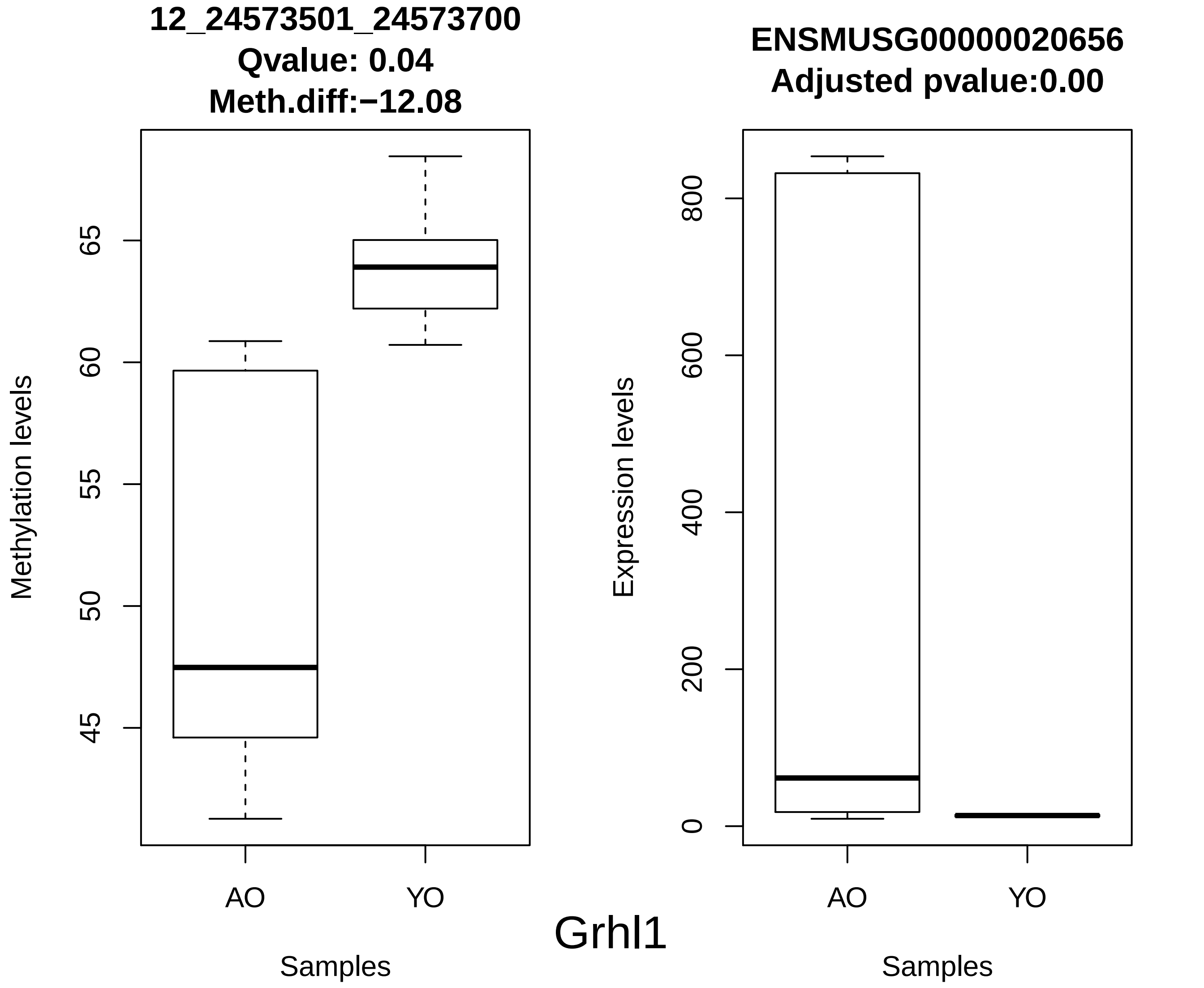
12_24573501_24573700
Qvalue: 0.04
Meth.diff:−12.08
Methylation levels
65
60
55
50
45
AO
YO
Samples
ENSMUSG00000020656
Adjusted pvalue:0.00
Expression levels
800
600
400
200
0
AO
YO
Samples
Grhl1

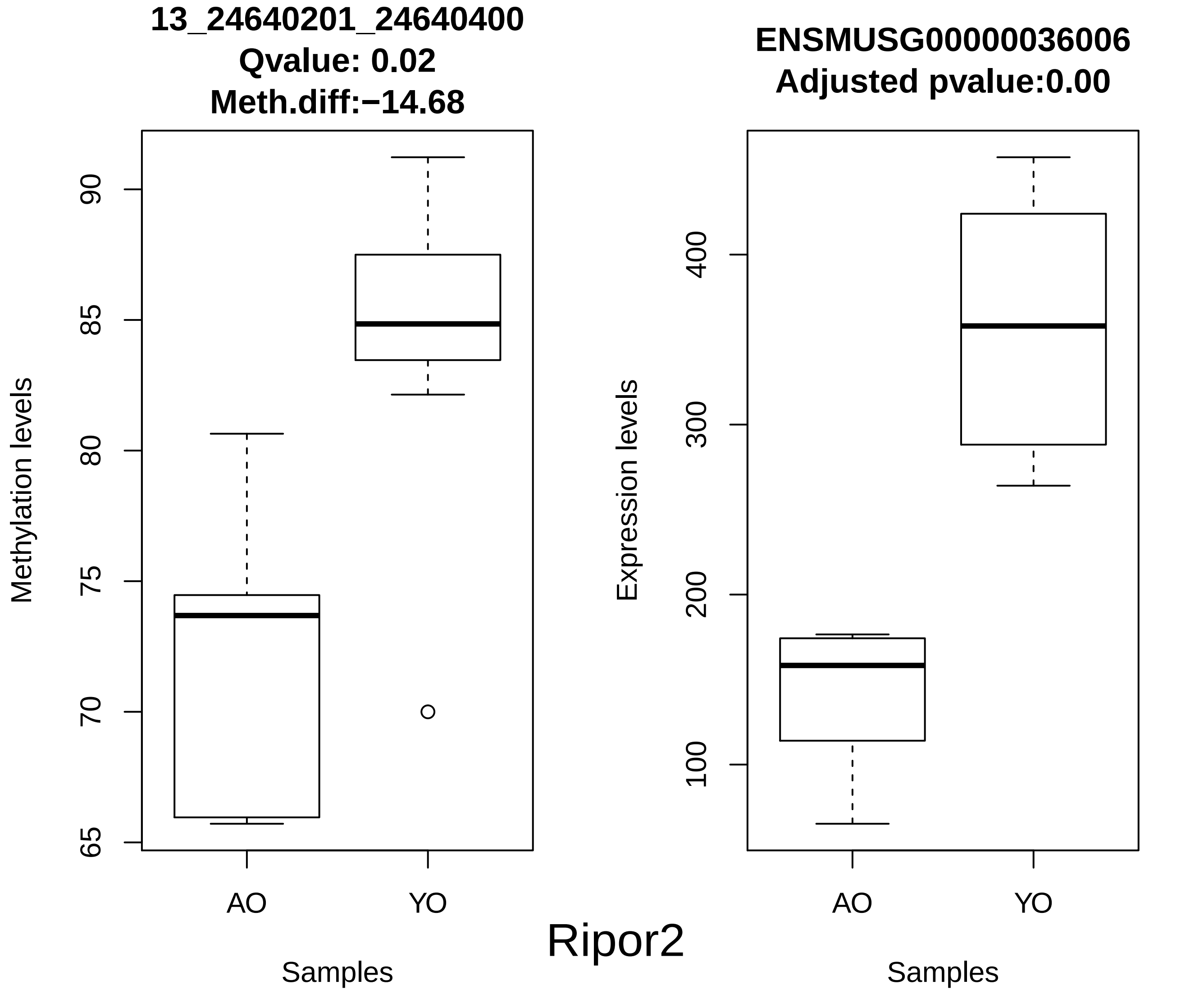
13_24640201_24640400
Qvalue: 0.02
Meth.diff:−14.68
Methylation levels
65
70
75
80
85
90
AO
YO
Samples
ENSMUSG00000036006
Adjusted pvalue:0.00
Expression levels
100
200
300
400
AO
YO
Samples
Ripor2

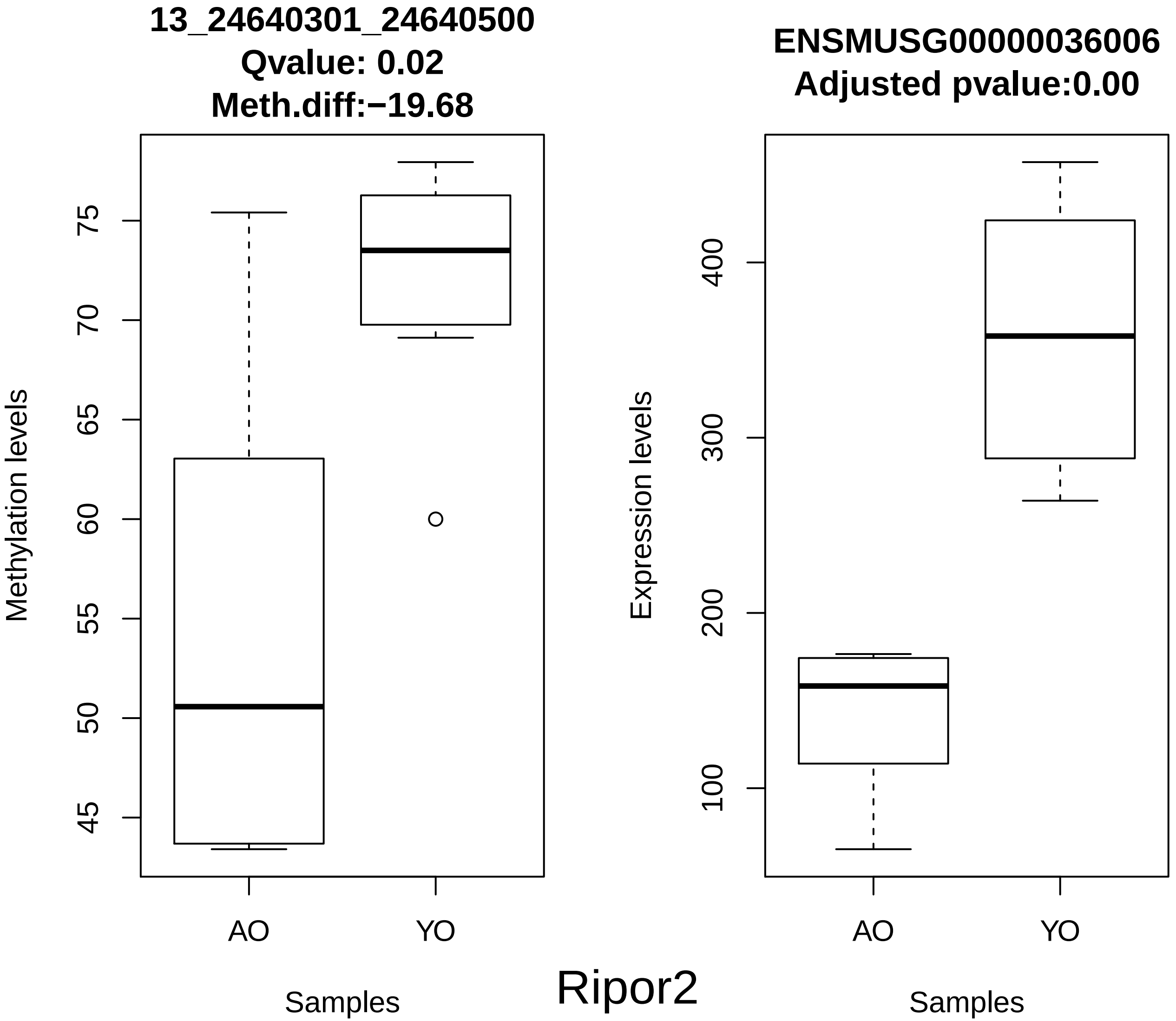
13_24640301_24640500
Qvalue: 0.02
Meth.diff:−19.68
Methylation levels
75
70
65
60
55
50
45
AO
YO
Samples
ENSMUSG00000036006
Adjusted pvalue:0.00
Expression levels
400
300
200
100
AO
YO
Samples
Ripor2

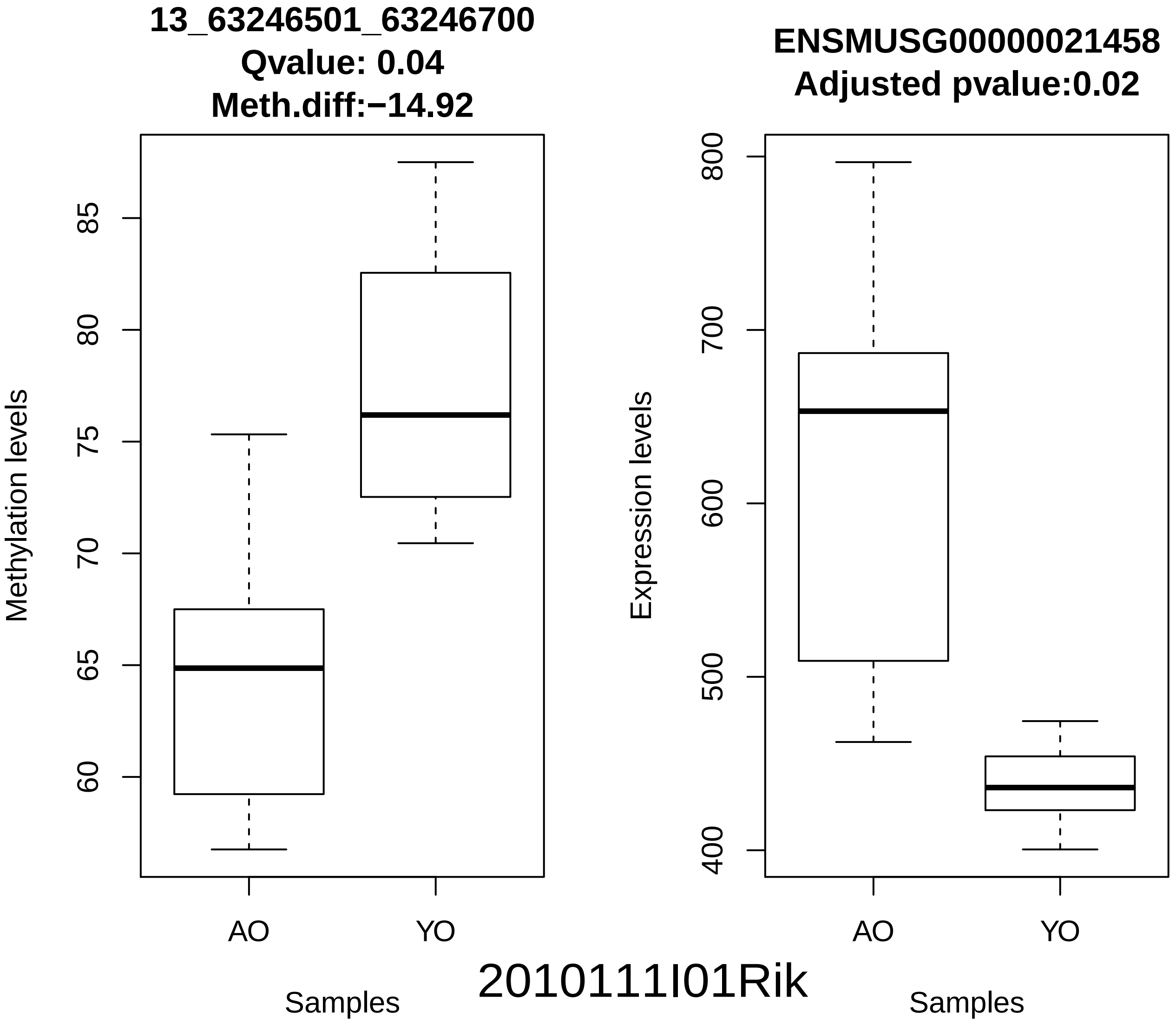
13_63246501_63246700
Qvalue: 0.04
Meth.diff:−14.92
Methylation levels
60
65
70
75
80
85
AO
YO
Samples
ENSMUSG00000021458
Adjusted pvalue:0.02
Expression levels
400
500
600
700
800
AO
YO
Samples
2010111I01Rik

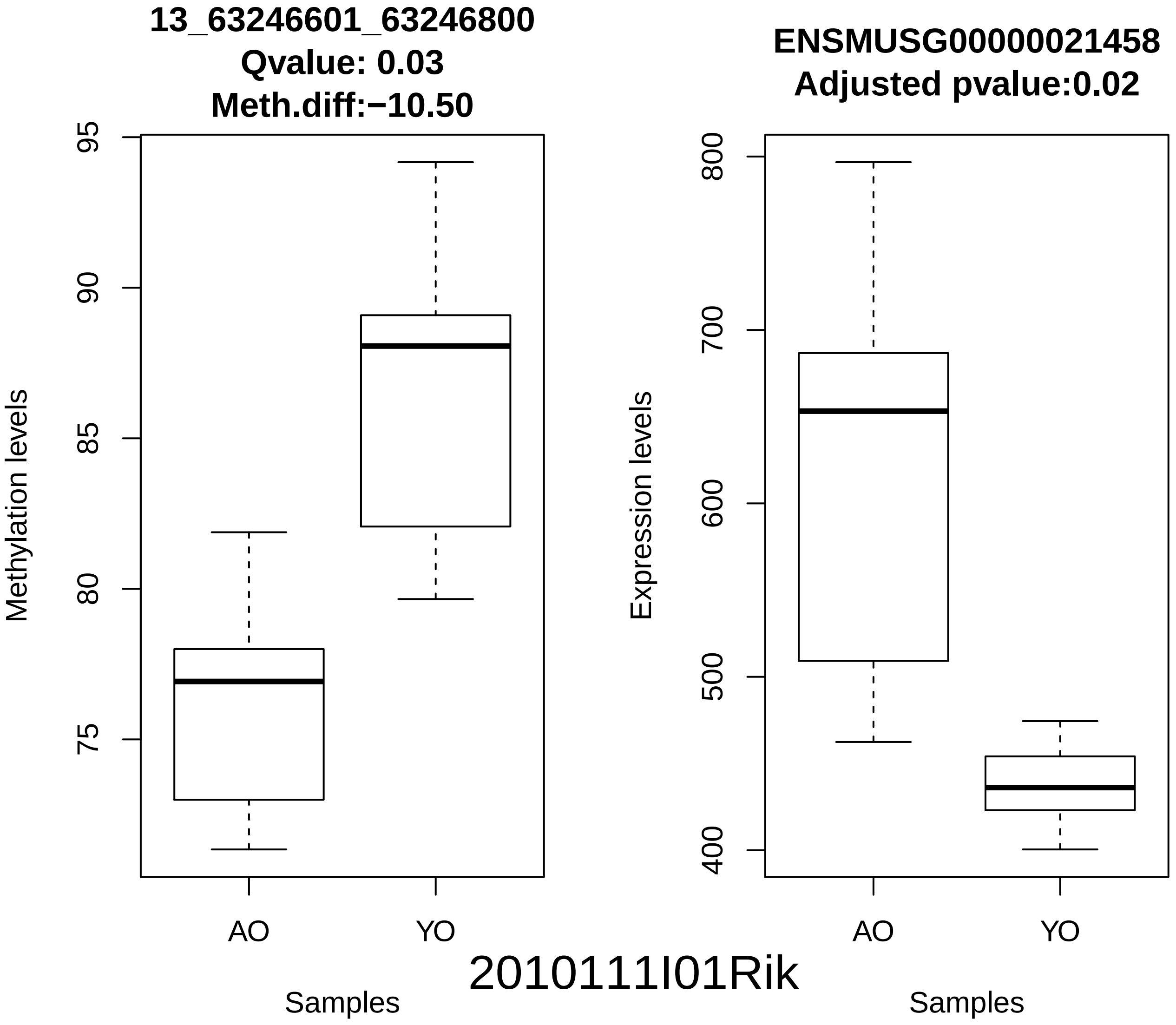
13_63246601_63246800
Qvalue: 0.03
Meth.diff:−10.50
Methylation levels
95
90
85
80
75
AO
YO
Samples
ENSMUSG00000021458
Adjusted pvalue:0.02
Expression levels
800
700
600
500
400
AO
YO
Samples
2010111I01Rik

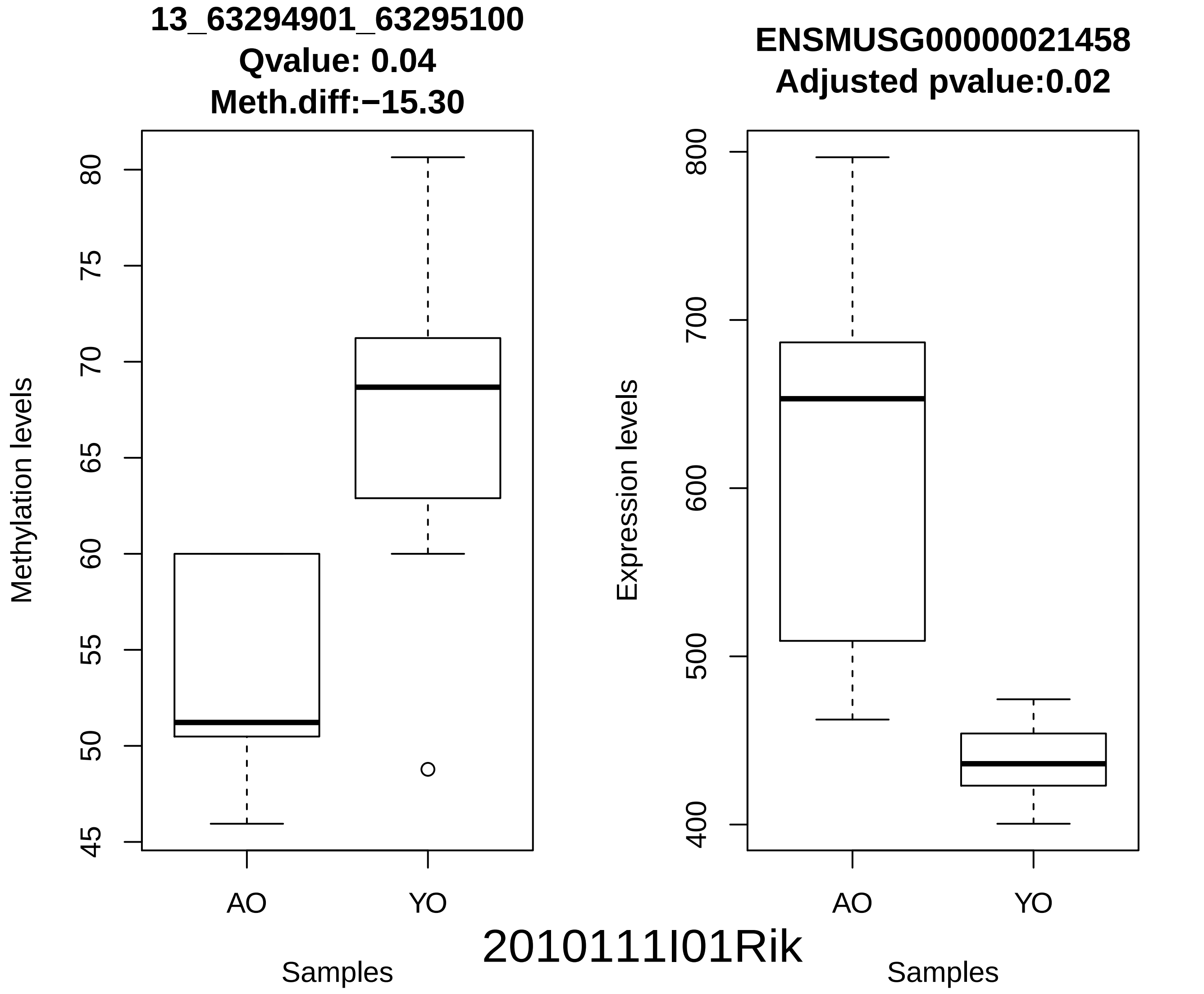
13_63294901_63295100
Qvalue: 0.04
Meth.diff:−15.30
Methylation levels
80
75
70
65
60
55
50
45
AO
YO
Samples
ENSMUSG00000021458
Adjusted pvalue:0.02
Expression levels
800
700
600
500
400
AO
YO
Samples
2010111I01Rik

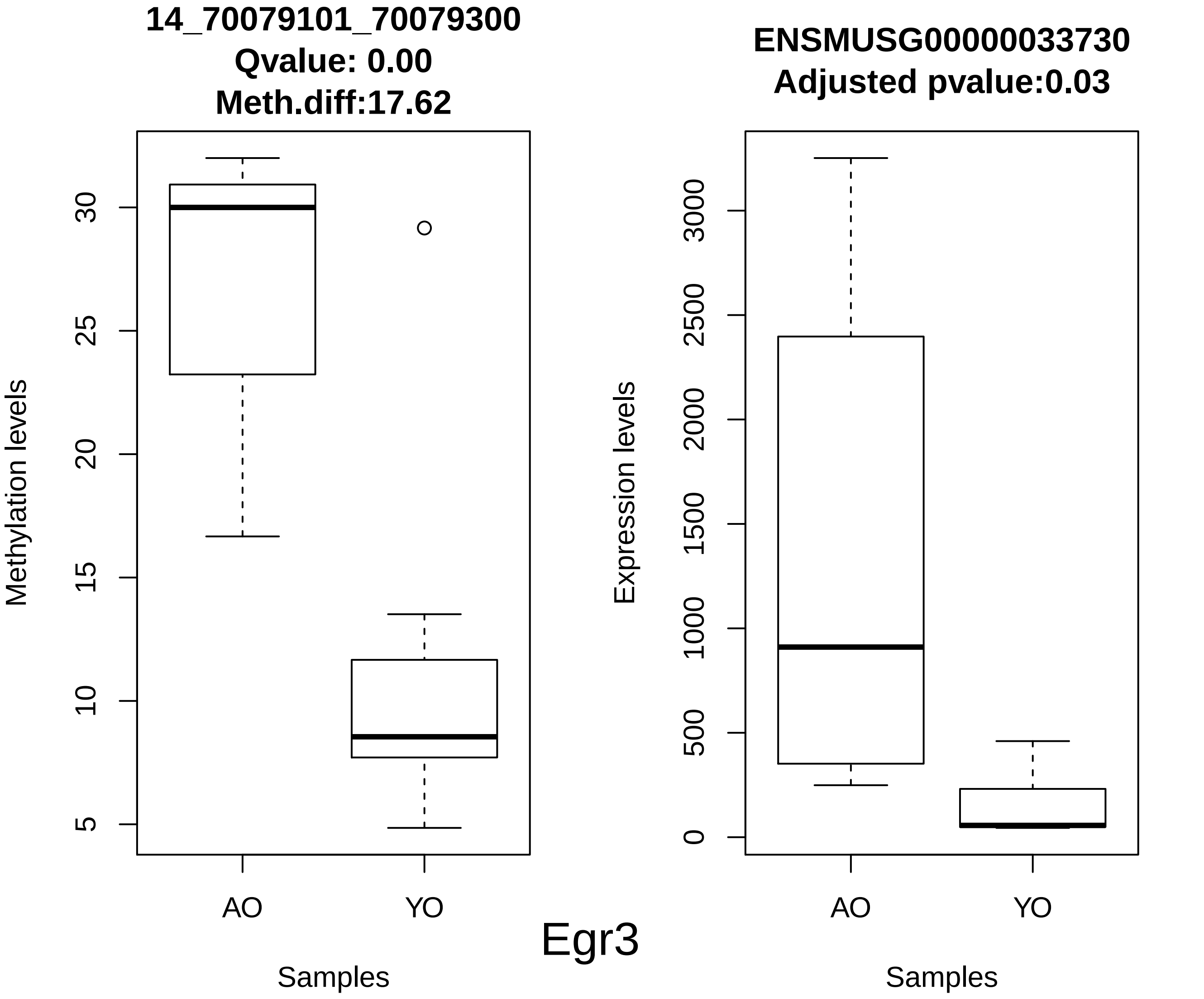
14_70079101_70079300
Qvalue: 0.00
Meth.diff:17.62
Methylation levels
5
10
15
20
25
30
AO
YO
Samples
ENSMUSG00000033730
Adjusted pvalue:0.03
Expression levels
0
500
1000
1500
2000
2500
3000
AO
YO
Samples
Egr3

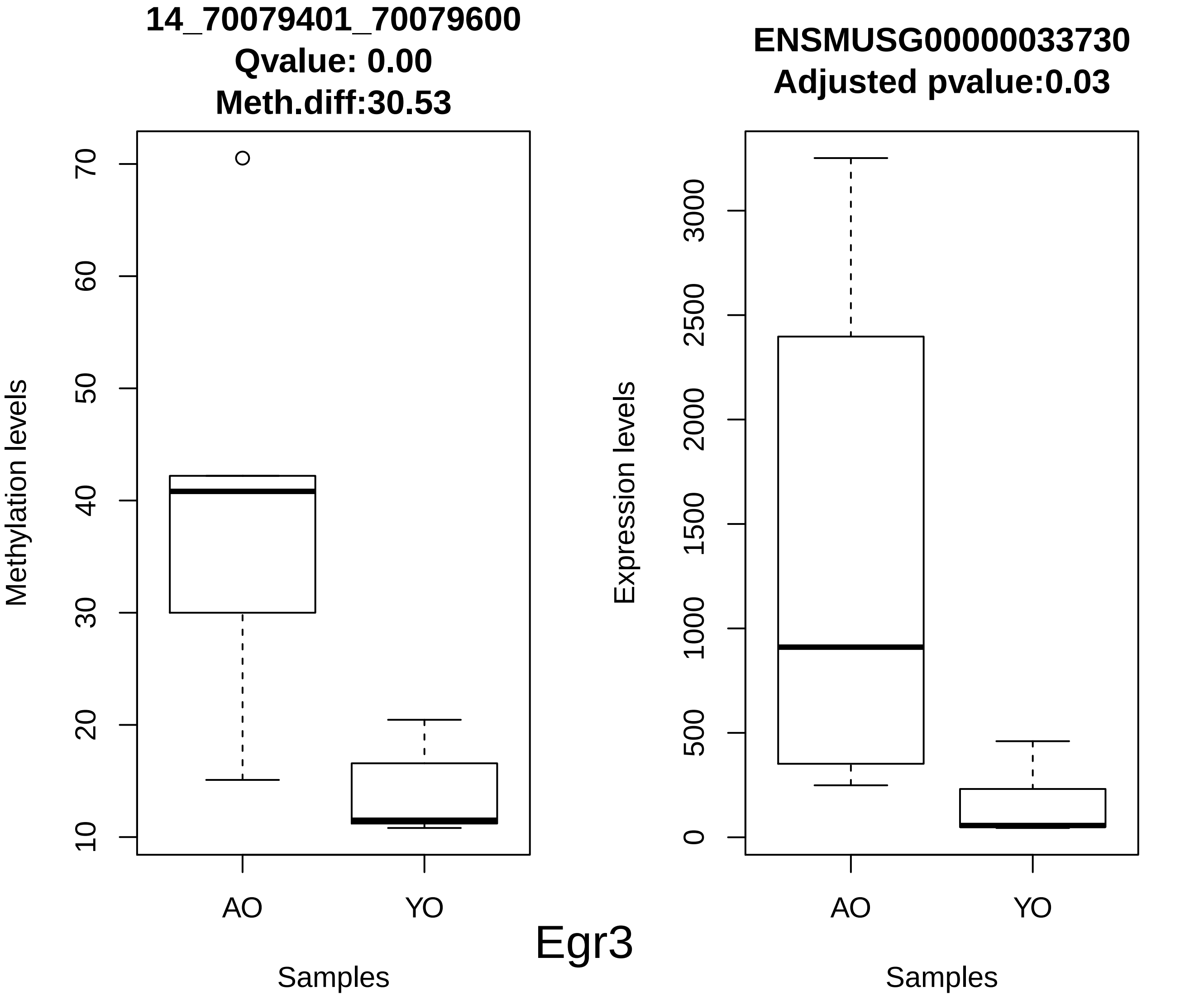
14_70079401_70079600
Qvalue: 0.00
Meth.diff:30.53
Methylation levels
10
20
30
40
50
60
70
AO
YO
Samples
ENSMUSG00000033730
Adjusted pvalue:0.03
Expression levels
0
500
1000
1500
2000
2500
3000
AO
YO
Samples

Egr3

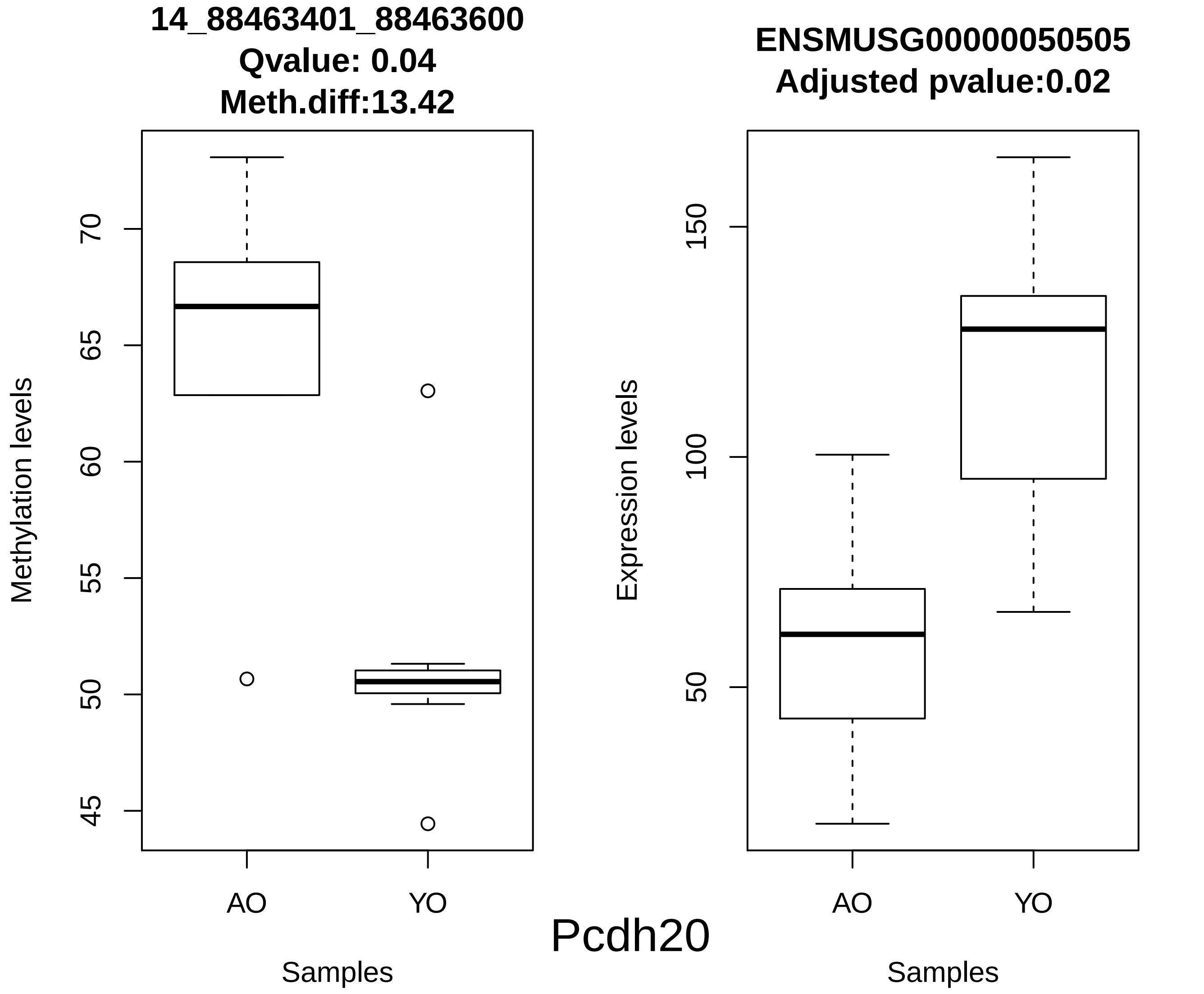
14_88463401_88463600
Qvalue: 0.04
Meth.diff:13.42
Methylation levels
70
65
60
55
50
45
AO
YO
Samples
ENSMUSG00000050505
Adjusted pvalue:0.02
Expression levels
150
100
50
AO
YO
Samples
Pcdh20

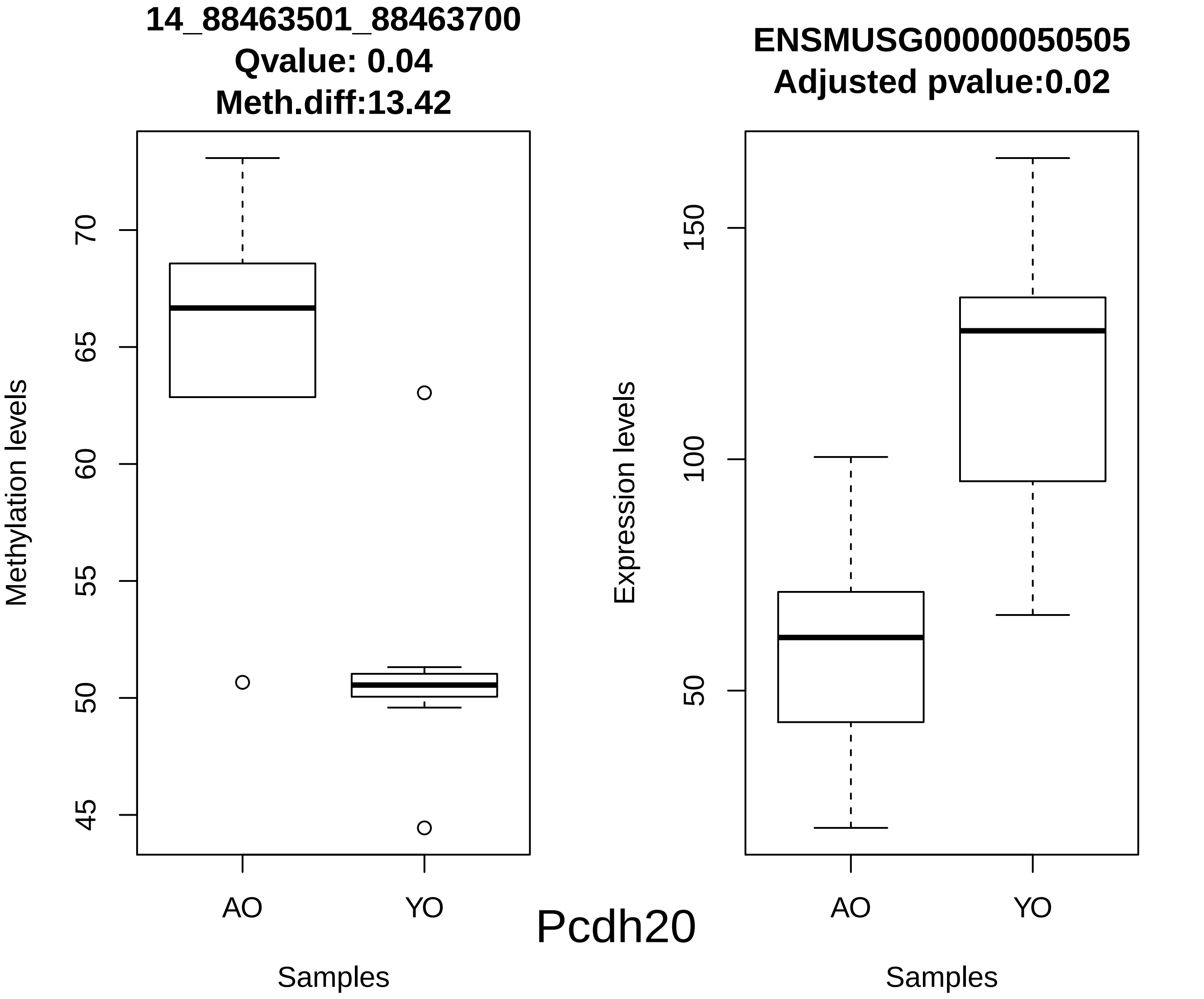
14_88463501_88463700
Qvalue: 0.04
Meth.diff:13.42
Methylation levels
70
65
60
55
50
45
AO
YO
Samples
Pcdh20
ENSMUSG00000050505
Adjusted pvalue:0.02
Expression levels
150
100
50
AO
YO
Samples

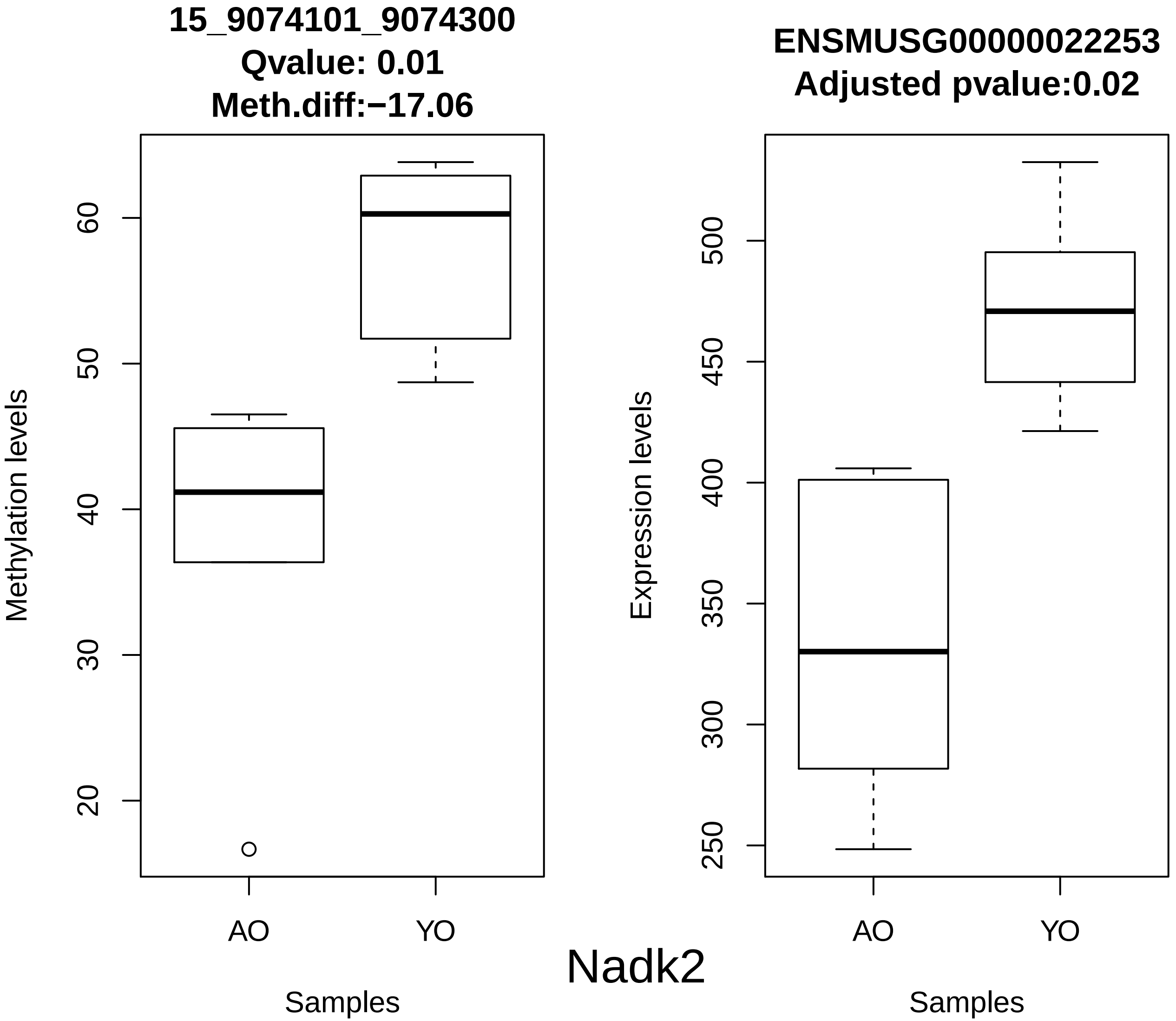
15_9074101_9074300
Qvalue: 0.01
Meth.diff:−17.06
Methylation levels
20
30
40
50
60
AO
YO
Samples
ENSMUSG00000022253
Adjusted pvalue:0.02
Expression levels
250
300
350
400
450
500
AO
YO
Samples
Nadk2

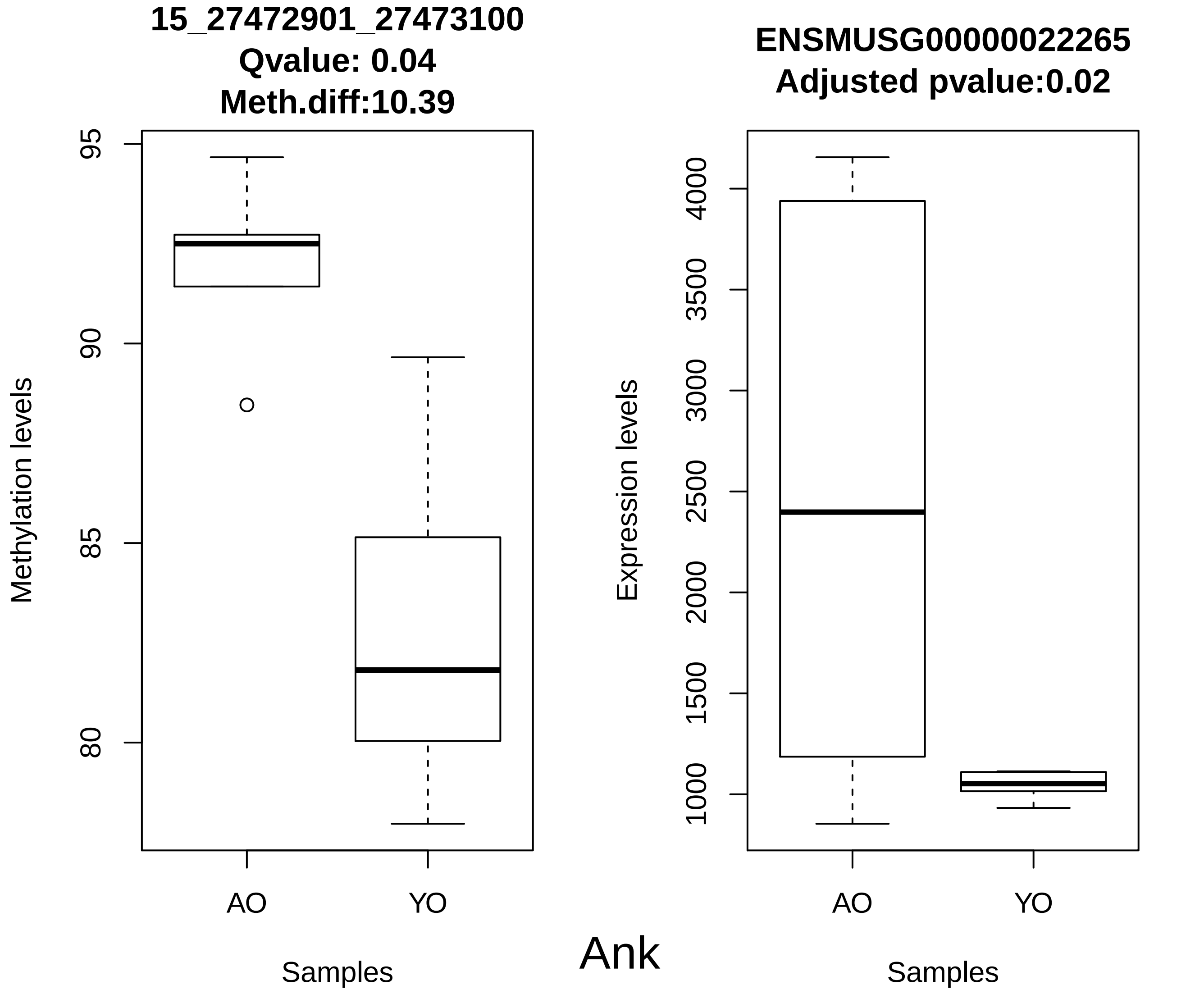
15_27472901_27473100
Qvalue: 0.04
Meth.diff:10.39
Methylation levels
95
90
85
80
AO
YO
Samples
ENSMUSG00000022265
Adjusted pvalue:0.02
Expression levels
4000
3500
3000
2500
2000
1500
1000
AO
YO
Samples
Ank

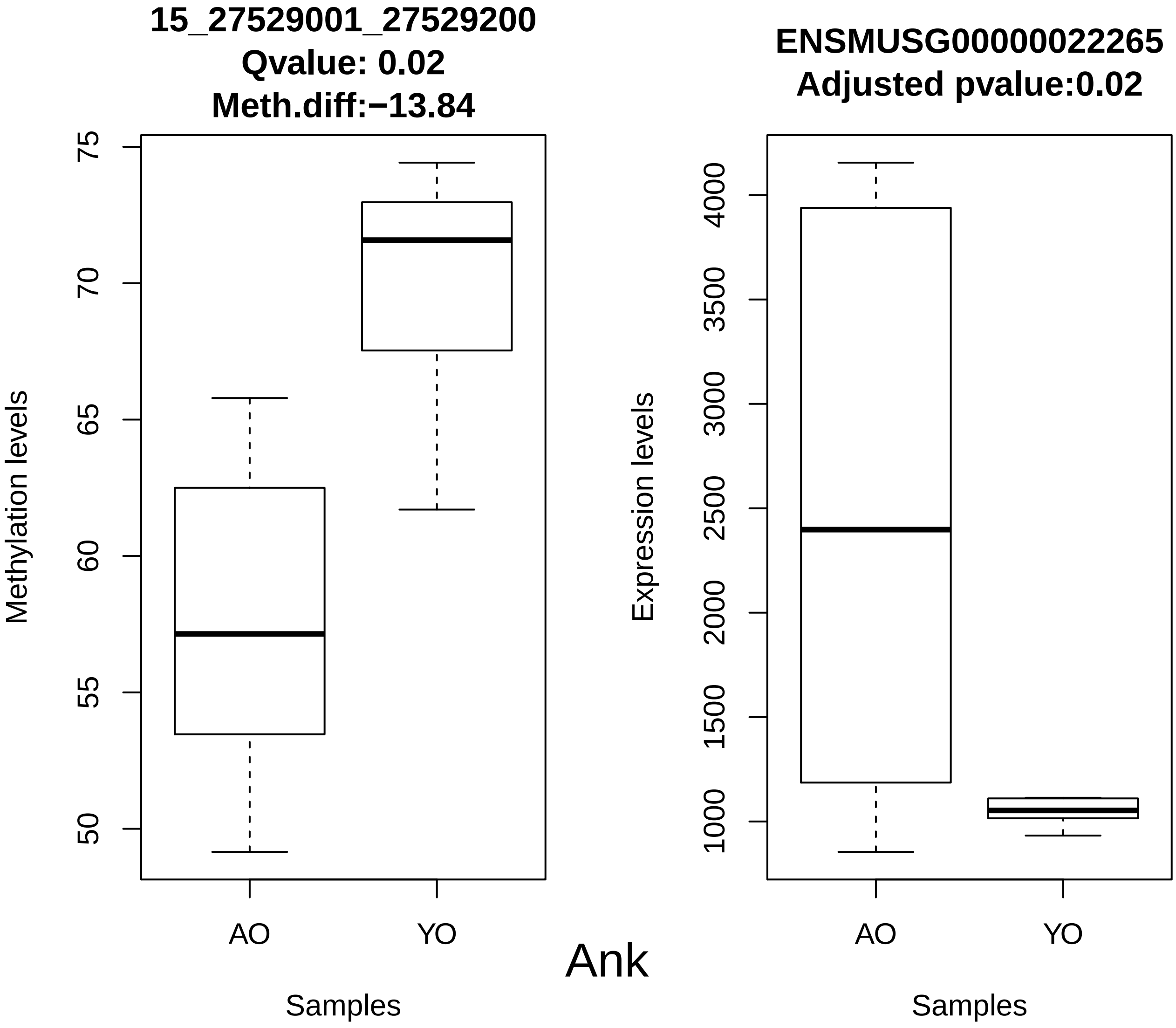
15_27529001_27529200
Qvalue: 0.02
Meth.diff:−13.84
Methylation levels
75
70
65
60
55
50
AO
YO
Samples
ENSMUSG00000022265
Adjusted pvalue:0.02
Expression levels
4000
3500
3000
2500
2000
1500
1000
AO
YO
Samples

Ank

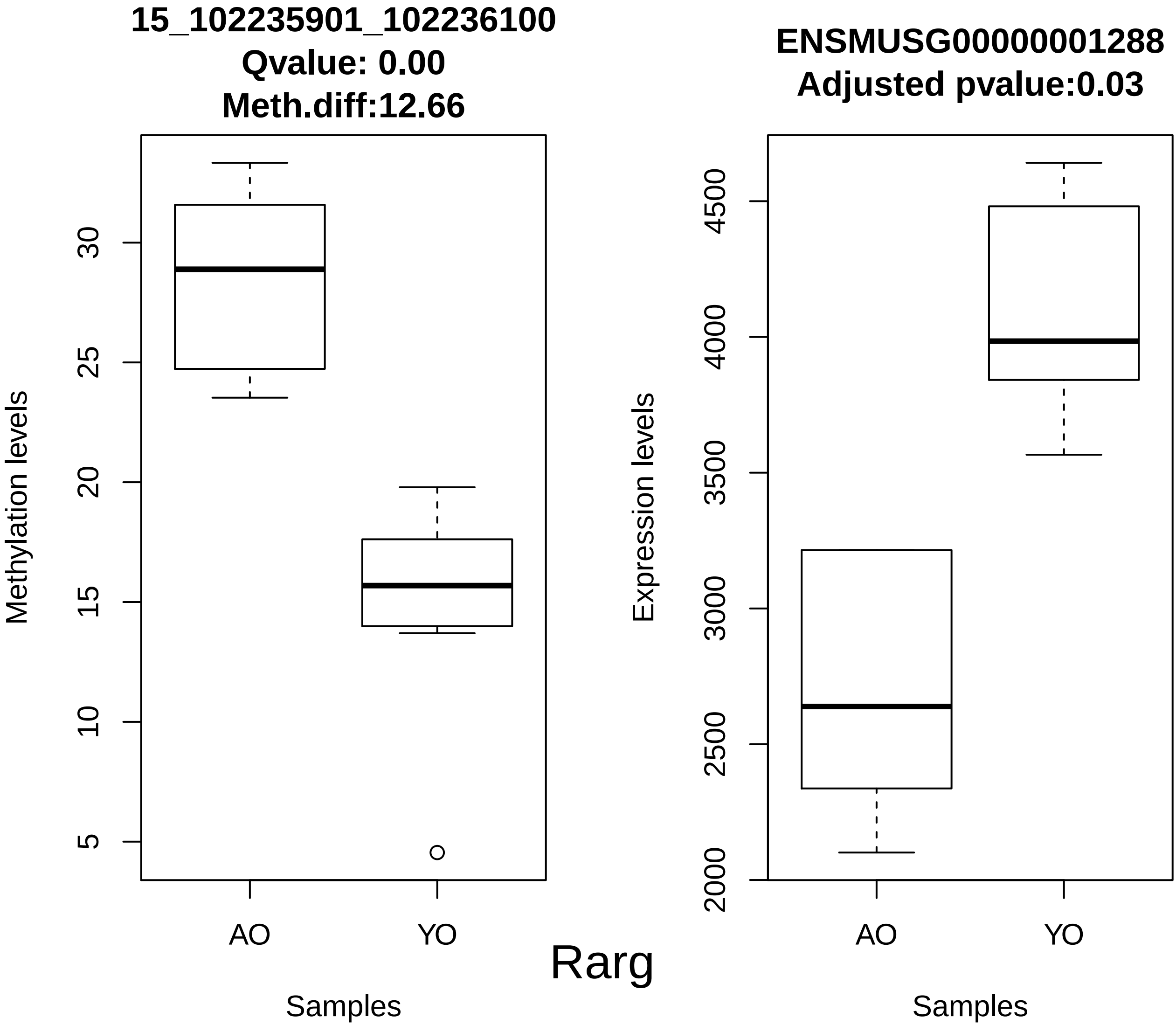

15_102235901_102236100
Qvalue: 0.00
Meth.diff:12.66
Methylation levels
5
10
15
20
25
30
AO
YO
Samples
ENSMUSG00000001288
Adjusted pvalue:0.03
Expression levels
2000
2500
3000
3500
4000
4500
AO
YO
Samples
Rarg

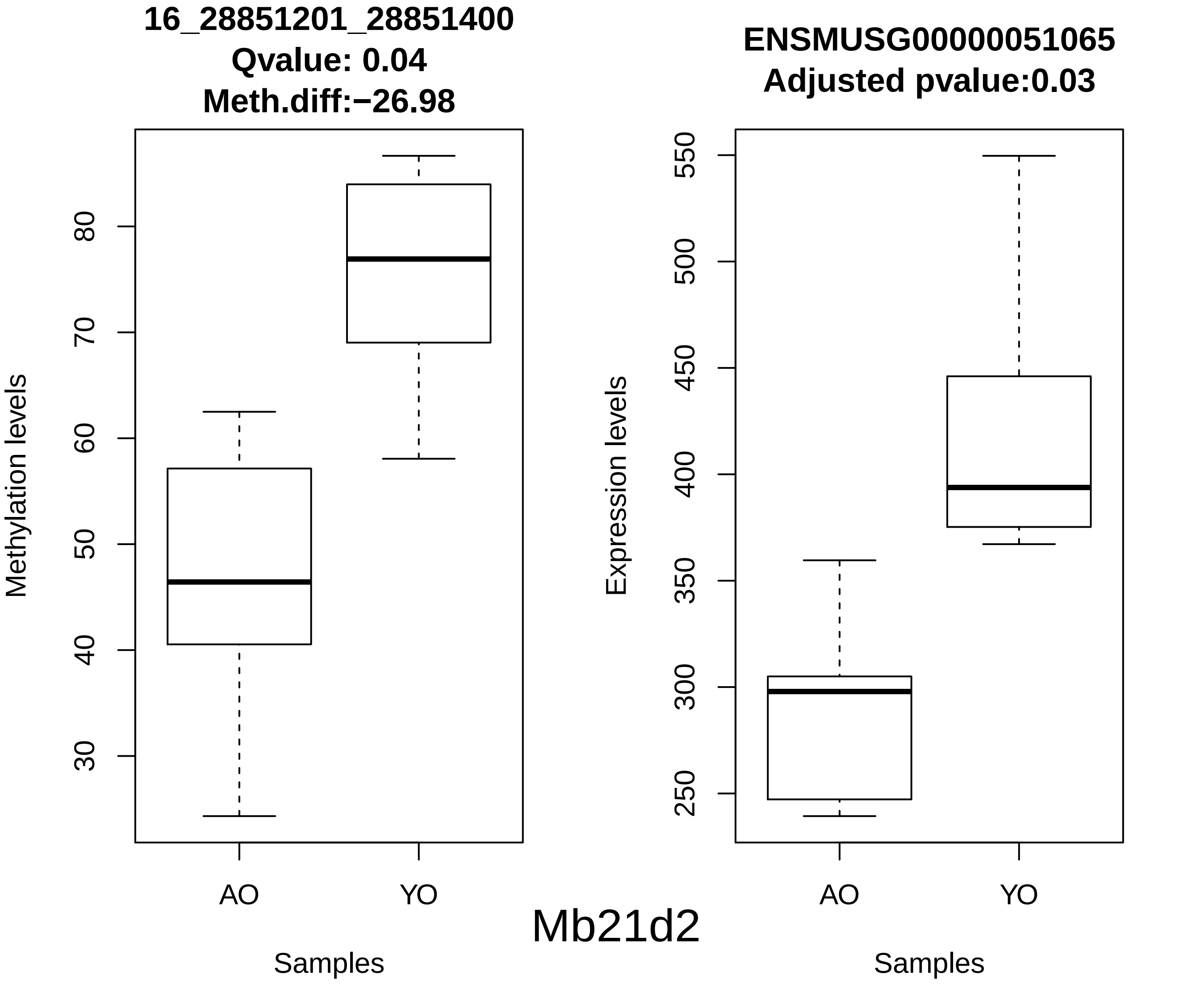
16_28851201_28851400
Qvalue: 0.04
Meth.diff:−26.98
Methylation levels
30
40
50
60
70
80
AO
YO
Samples
ENSMUSG00000051065
Adjusted pvalue:0.03
Expression levels
250
300
350
400
450
500
550
AO
YO
Samples
Mb21d2

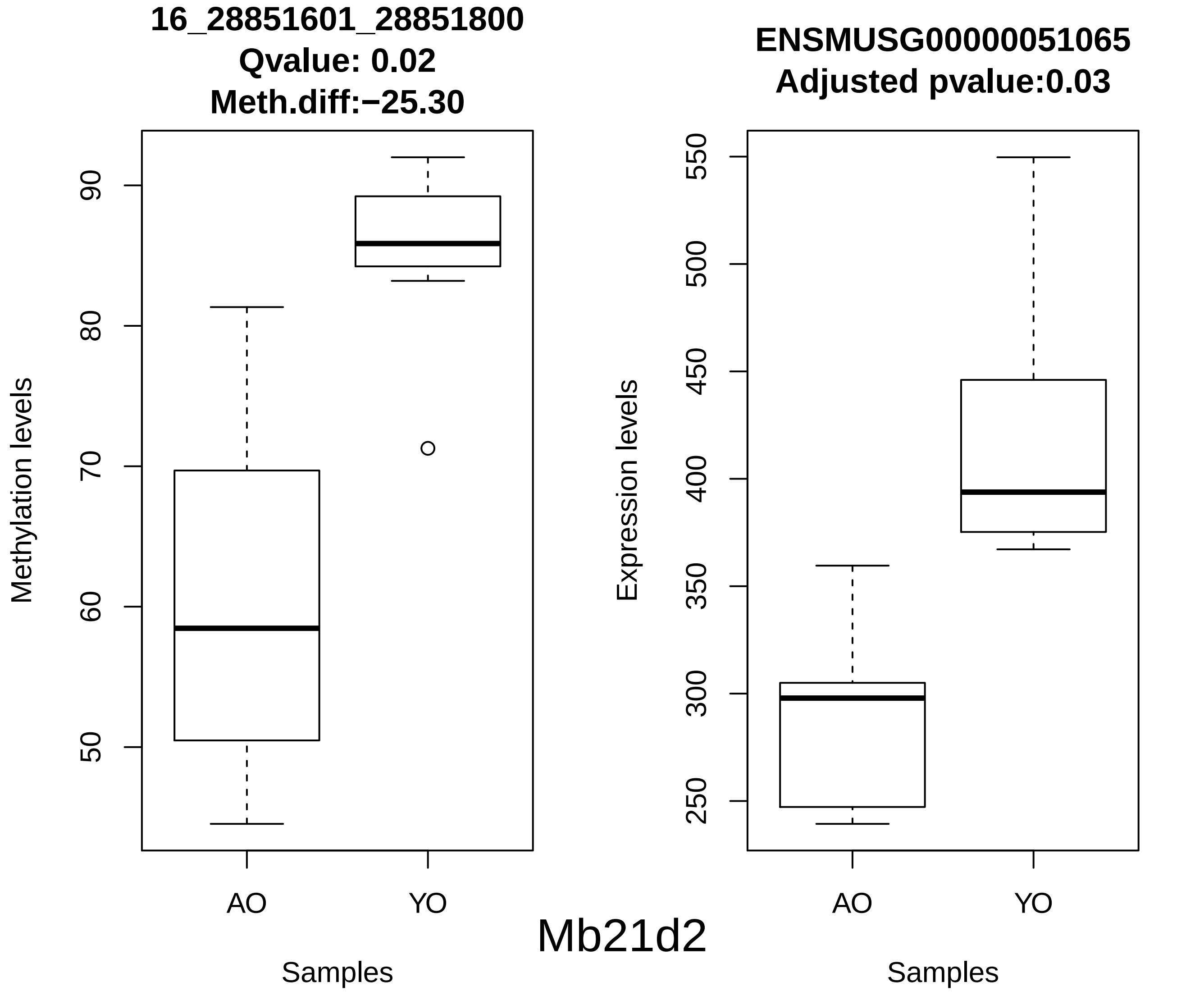
16_28851601_28851800
Qvalue: 0.02
Meth.diff:−25.30
Methylation levels
50
60
70
80
90
AO
YO
Samples
ENSMUSG00000051065
Adjusted pvalue:0.03
Expression levels
250
300
350
400
450
500
550
AO
YO
Samples
Mb21d2

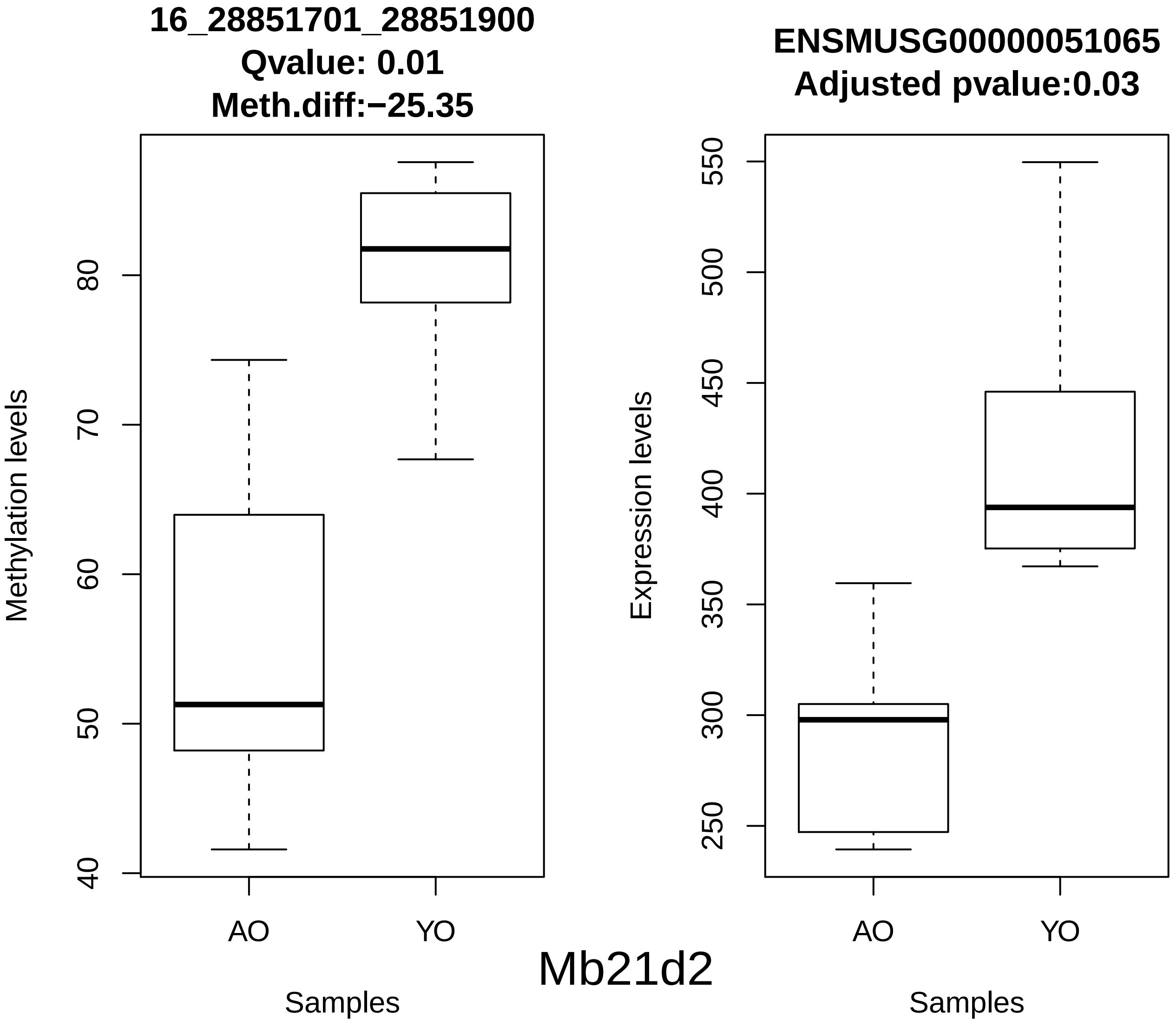
16_28851701_28851900
Qvalue: 0.01
Meth.diff:−25.35
Methylation levels
40
50
60
70
80
AO
YO
Samples
ENSMUSG00000051065
Adjusted pvalue:0.03
Expression levels
250
300
350
400
450
500
550
AO
YO
Samples
Mb21d2

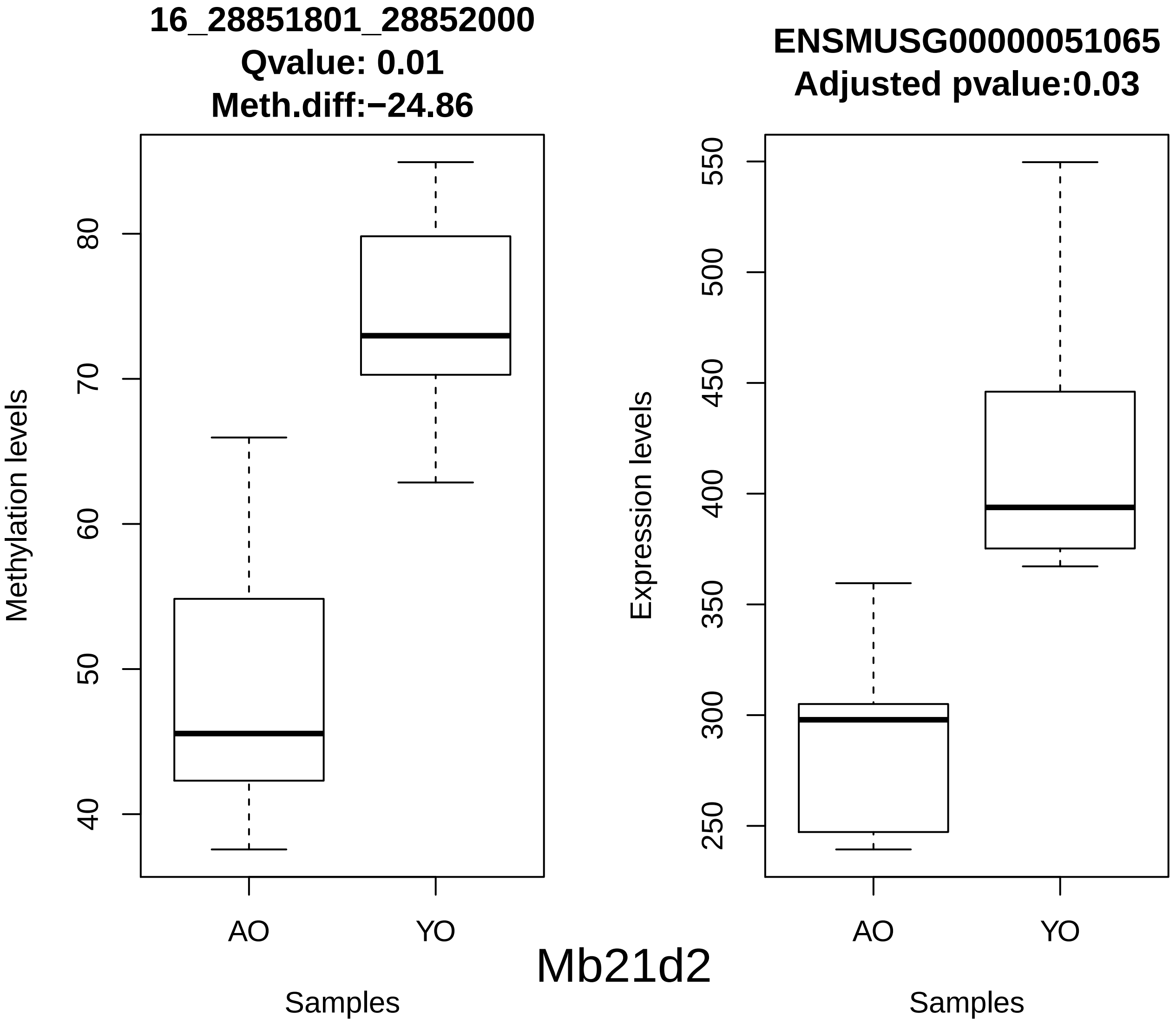
16_28851801_28852000
Qvalue: 0.01
Meth.diff:−24.86
Methylation levels
40
50
60
70
80
AO
YO
Samples
ENSMUSG00000051065
Adjusted pvalue:0.03
Expression levels
250
300
350
400
450
500
550
AO
YO
Samples
Mb21d2

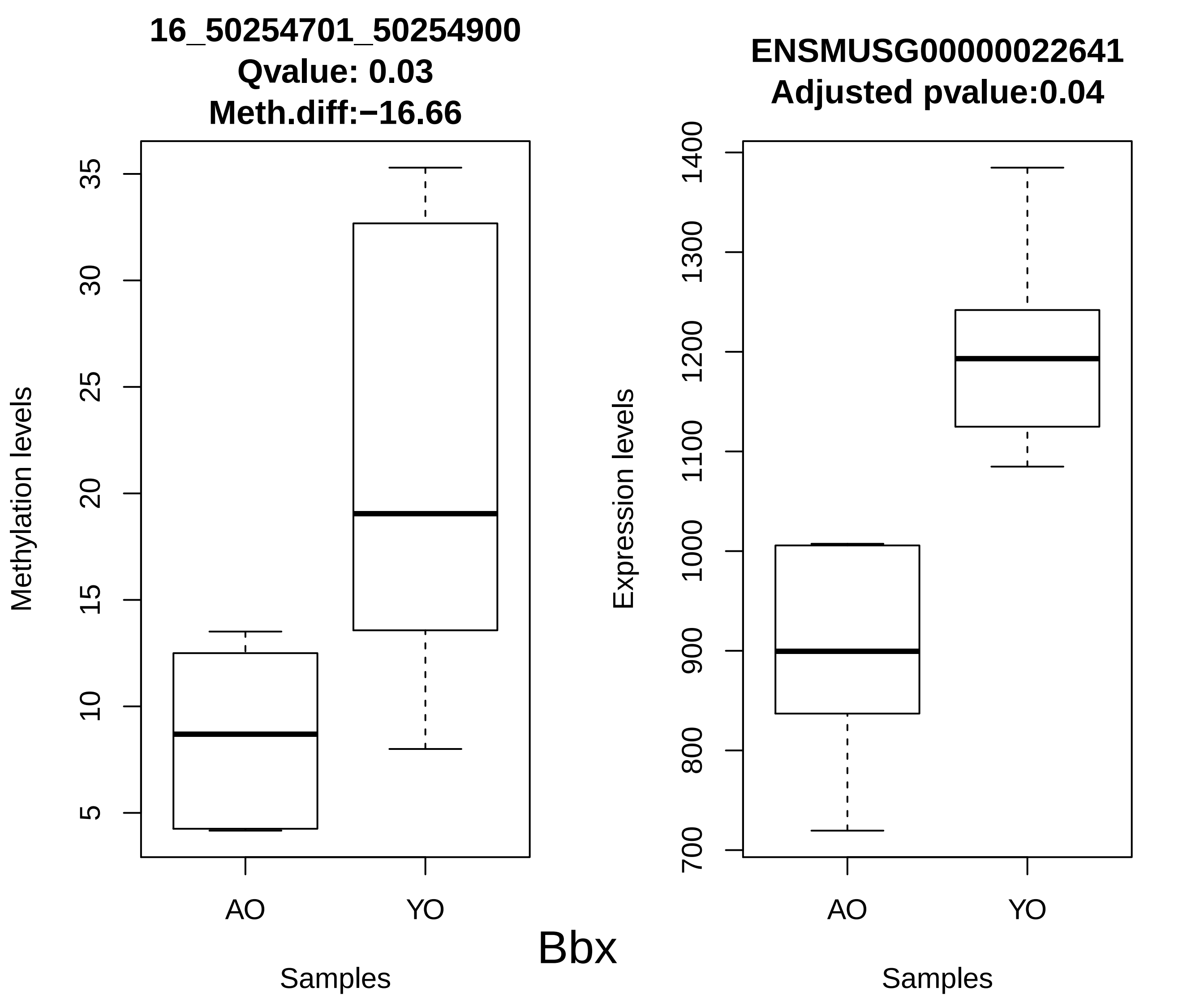
16_50254701_50254900
Qvalue: 0.03
Meth.diff:−16.66
Methylation levels
5
10
15
20
25
30
35
AO
YO
Samples
ENSMUSG00000022641
Adjusted pvalue:0.04
Expression levels
700
800
900
1000
1100
1200
1300
1400
AO
YO
Samples
Bbx

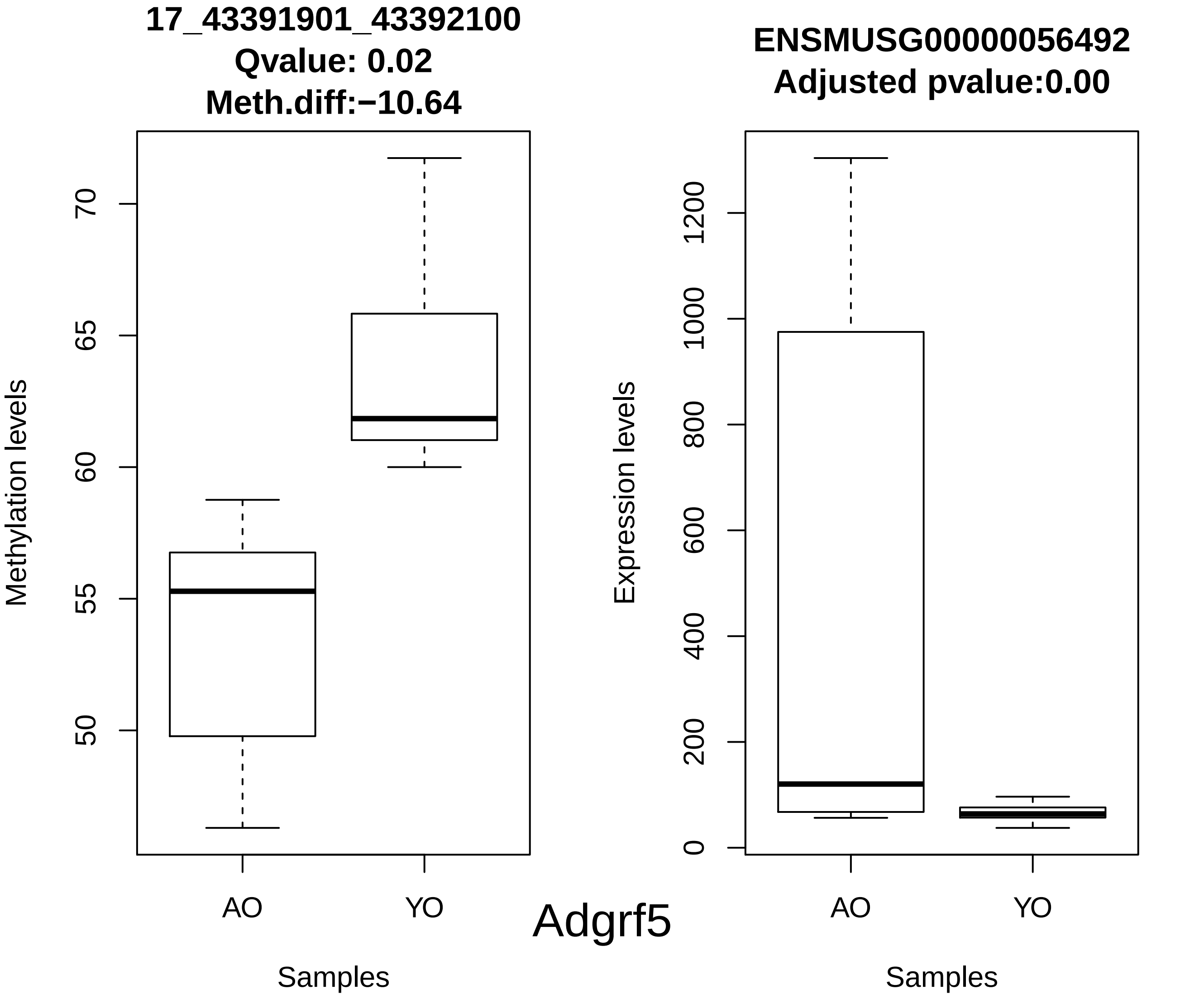
17_43391901_43392100
Qvalue: 0.02
Meth.diff:−10.64
Methylation levels
50
55
60
65
70
AO
YO
Samples
ENSMUSG00000056492
Adjusted pvalue:0.00
Expression levels
0
200
400
600
800
1000
1200
AO
YO
Samples
Adgrf5

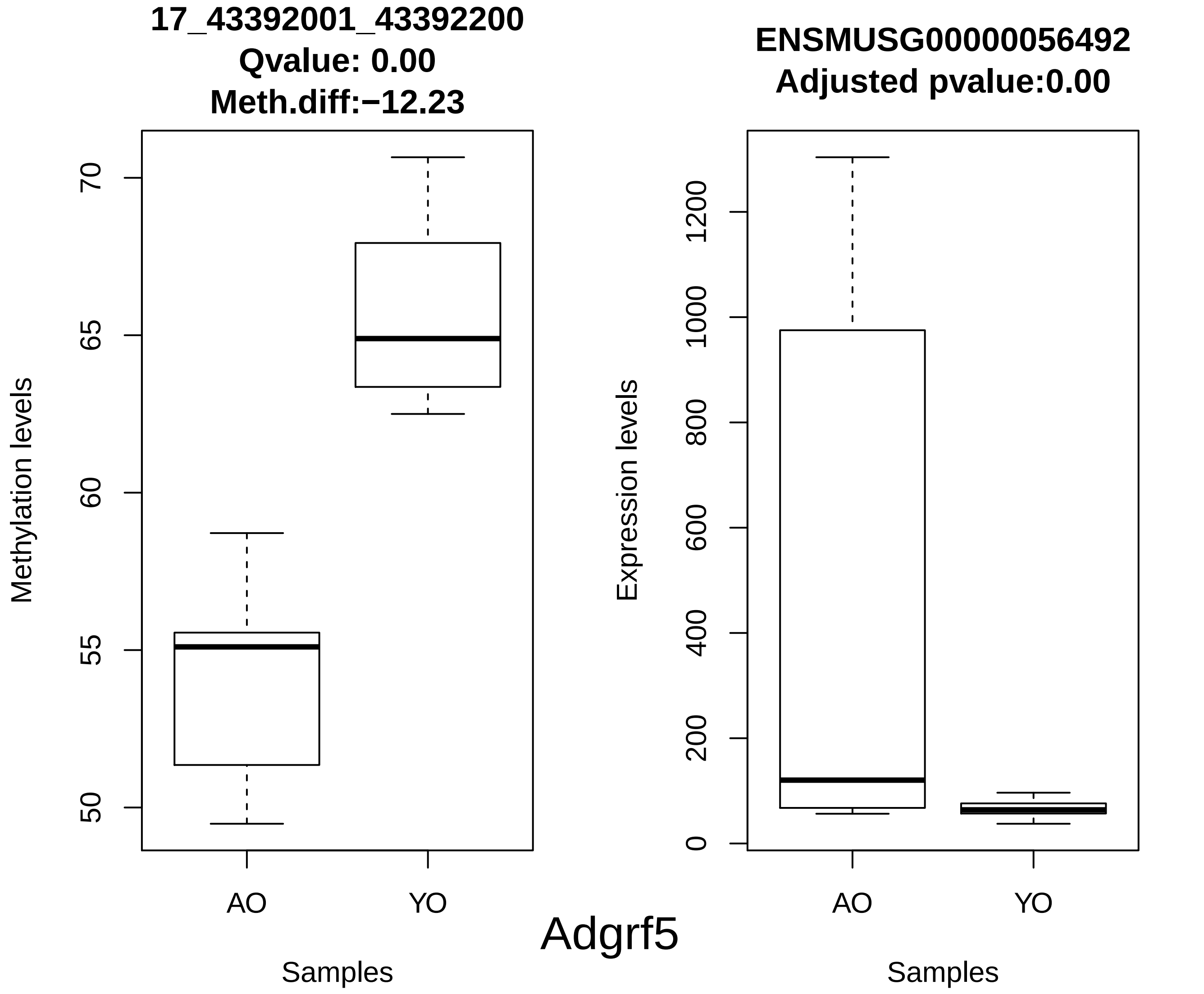

17_43392001_43392200
Qvalue: 0.00
Meth.diff:−12.23
Methylation levels
70
65
60
55
50
AO
YO
Samples
ENSMUSG00000056492
Adjusted pvalue:0.00
Expression levels
1200
1000
800
600
400
200
0
AO
YO
Samples
Adgrf5

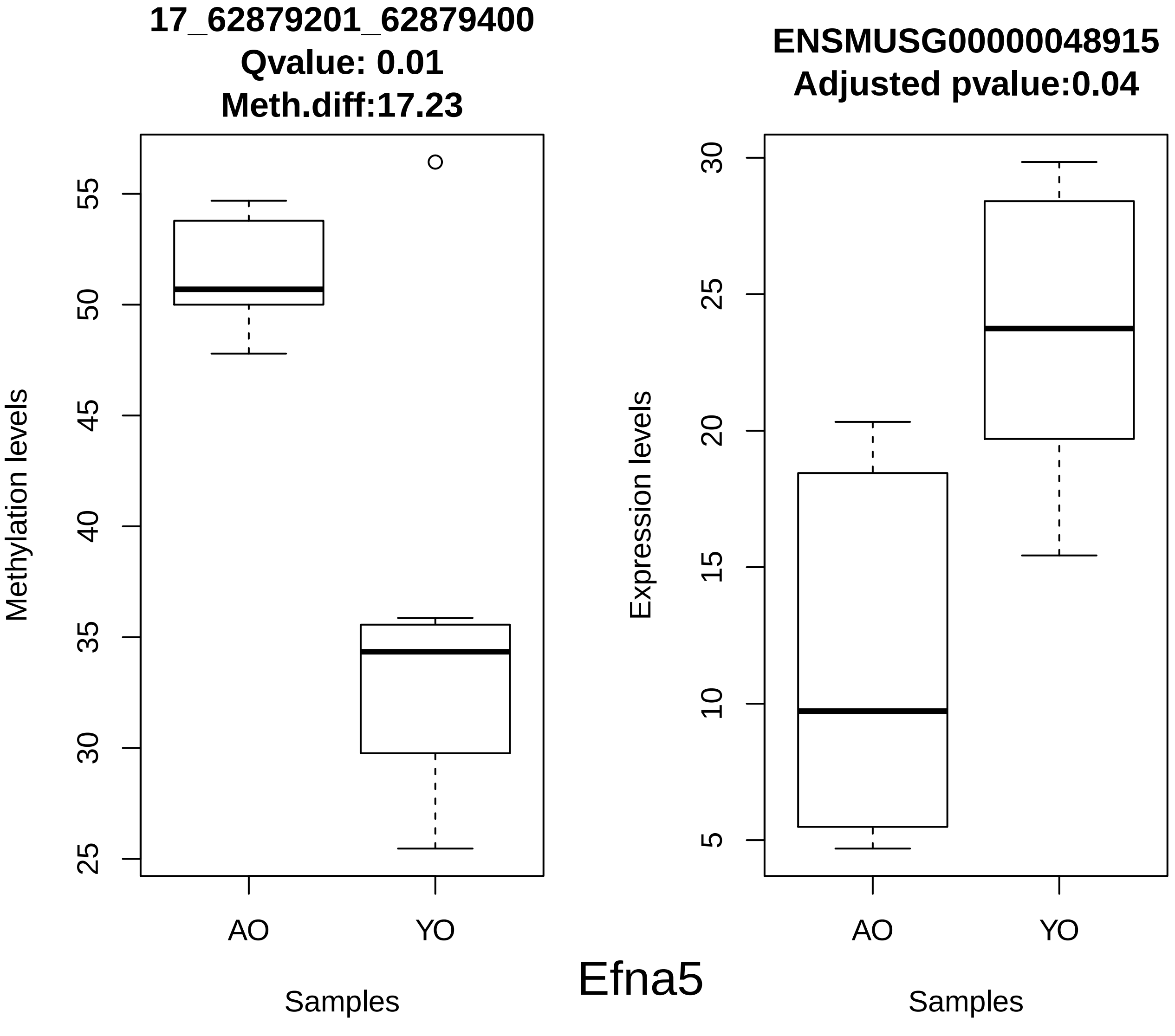
17_62879201_62879400
Qvalue: 0.01
Meth.diff:17.23
Methylation levels
55
50
45
40
35
30
25
AO
YO
Samples
ENSMUSG00000048915
Adjusted pvalue:0.04
Expression levels
30
25
20
15
10
5
AO
YO
Samples
Efna5

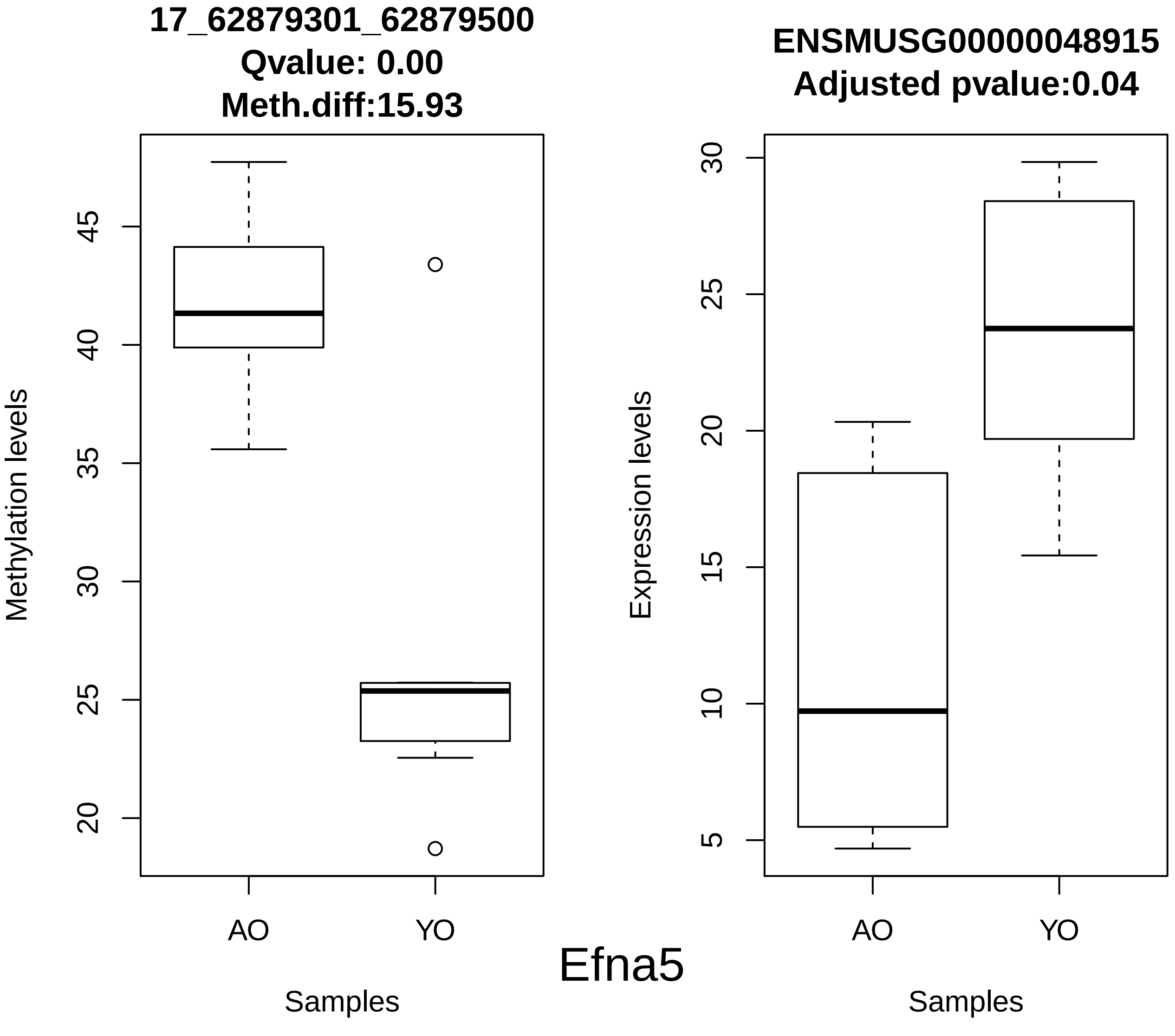
17_62879301_62879500
Qvalue: 0.00
Meth.diff:15.93
Methylation levels
45
40
35
30
25
20
AO
YO
Samples
ENSMUSG00000048915
Adjusted pvalue:0.04
Expression levels
30
25
20
15
10
5
AO
YO
Samples
Efna5

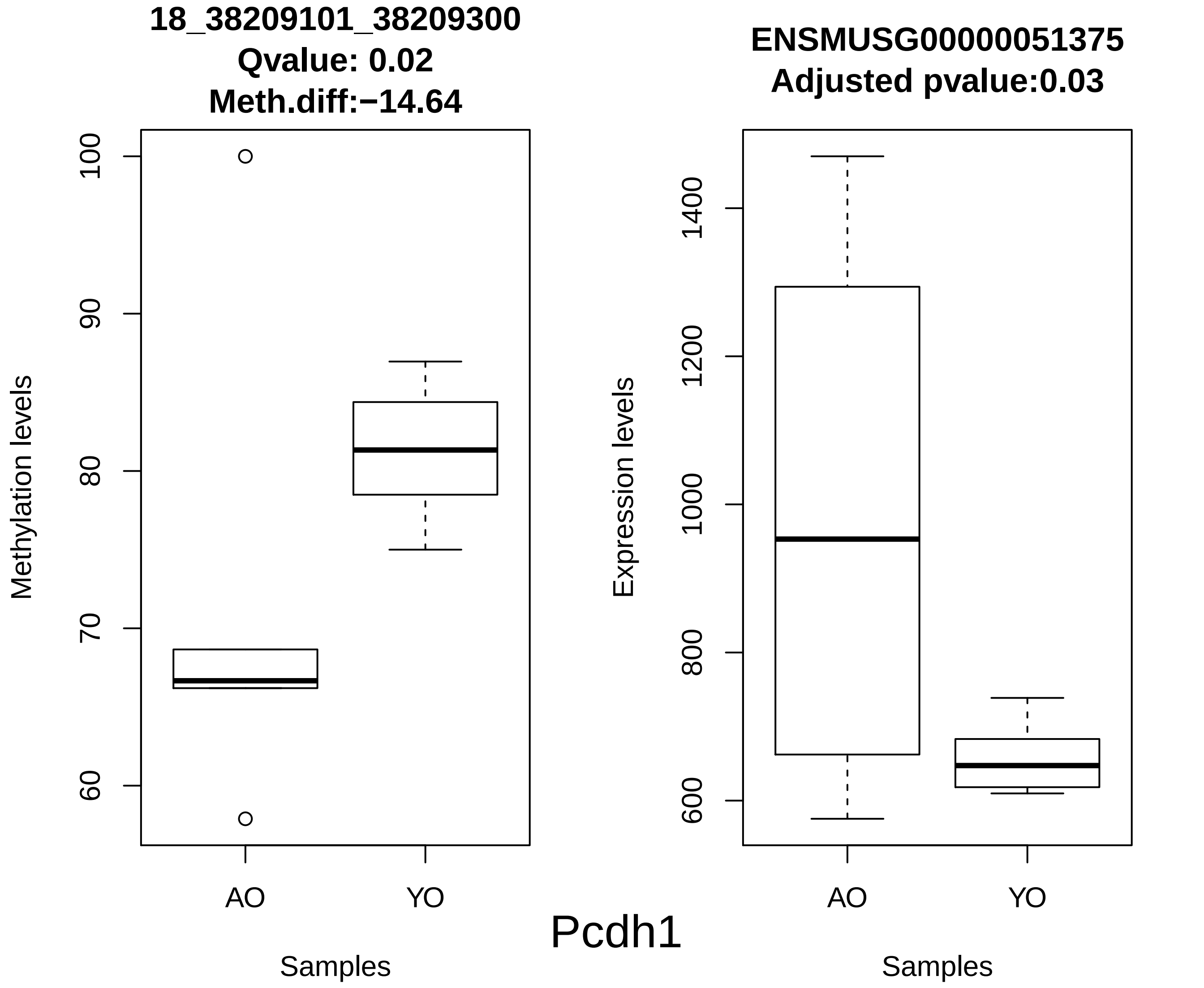

18_38209101_38209300
Qvalue: 0.02
Meth.diff:−14.64
Methylation levels
100
90
80
70
60
AO
YO
Samples
ENSMUSG00000051375
Adjusted pvalue:0.03
Expression levels
1400
1200
1000
800
600
AO
YO
Samples
Pcdh1

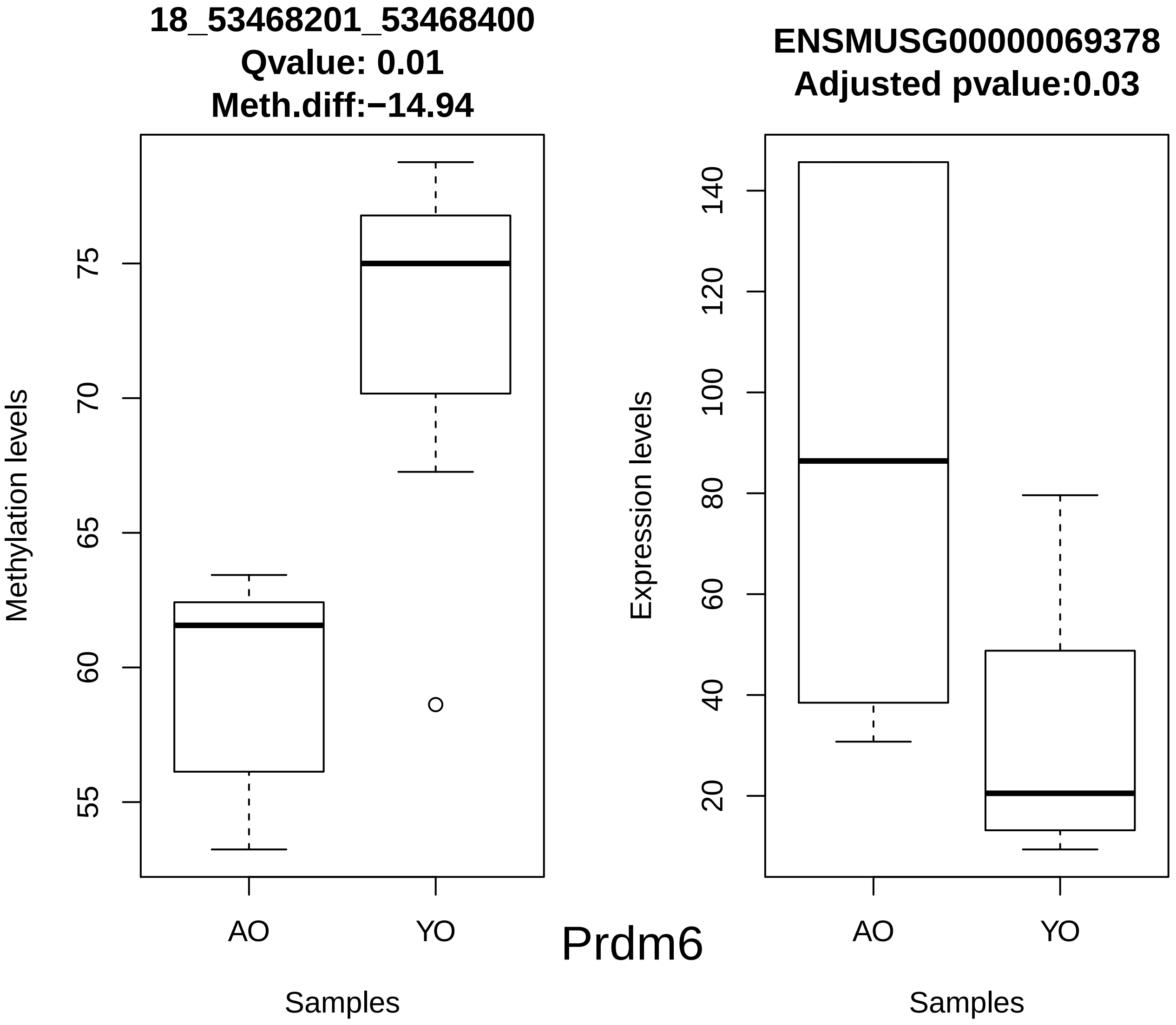
18_53468201_53468400
Qvalue: 0.01
Meth.diff:−14.94
Methylation levels
55
60
65
70
75
AO
YO
Samples
Prdm6
ENSMUSG00000069378
Adjusted pvalue:0.03
Expression levels
20
40
60
80
100
120
140
AO
YO
Samples

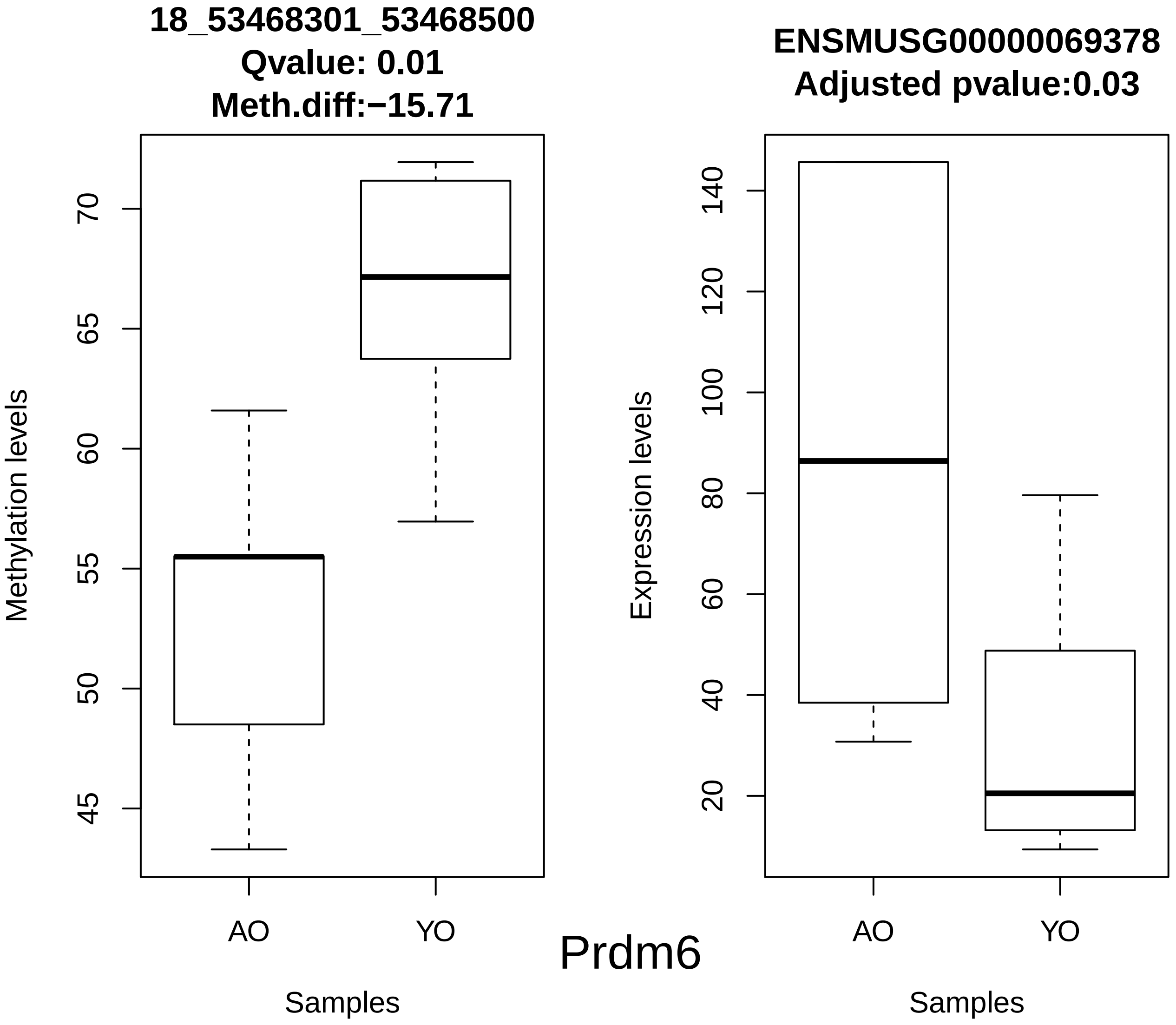
18_53468301_53468500
Qvalue: 0.01
Meth.diff:−15.71
Methylation levels
45
50
55
60
65
70
AO
YO
Samples
ENSMUSG00000069378
Adjusted pvalue:0.03
Expression levels
20
40
60
80
100
120
140
AO
YO
Samples
Prdm6

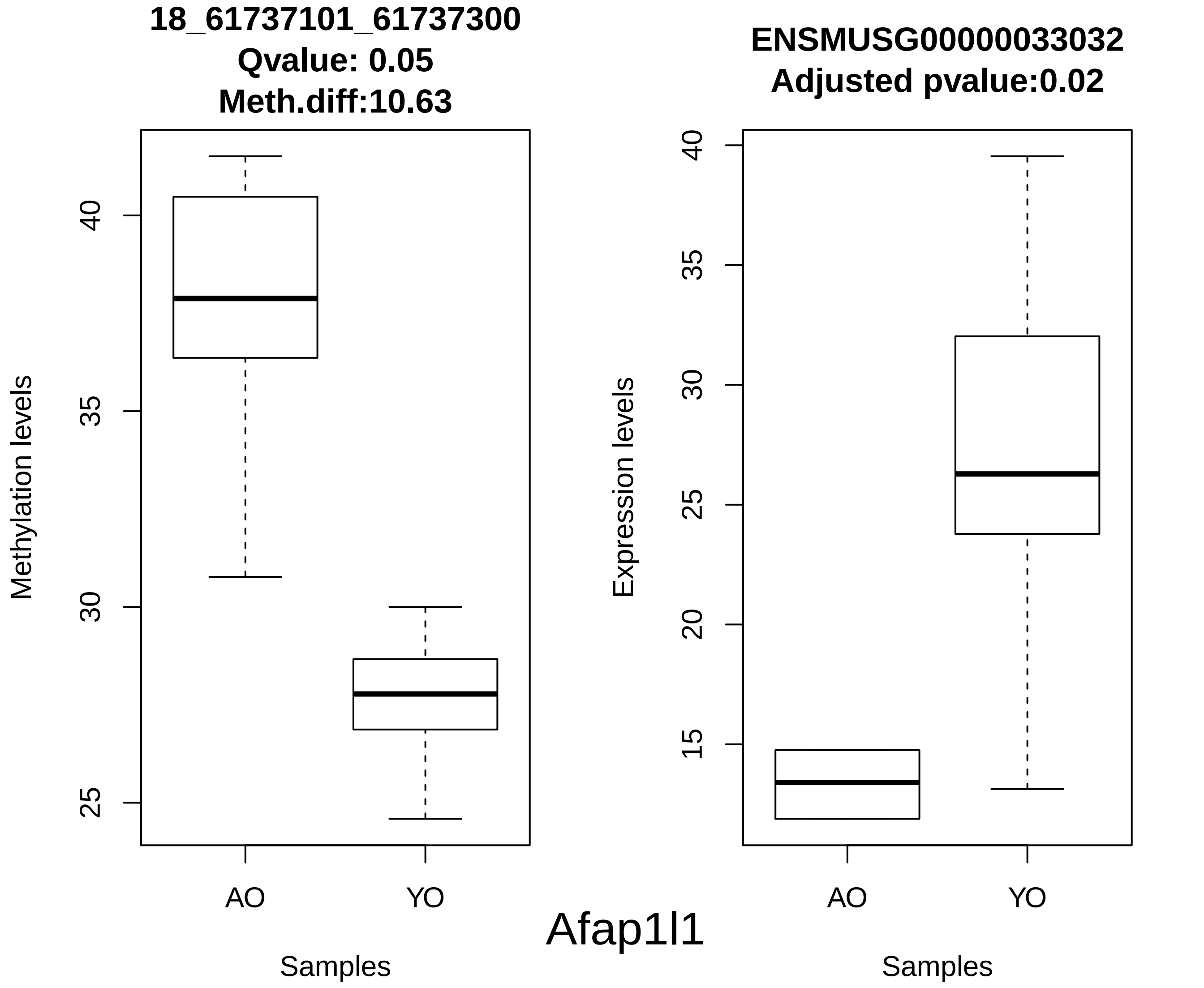
18_61737101_61737300
Qvalue: 0.05
Meth.diff:10.63
Methylation levels
25
30
35
40
AO
YO
Samples
ENSMUSG00000033032
Adjusted pvalue:0.02
Expression levels
15
20
25
30
35
40
AO
YO
Samples
Afap1l1

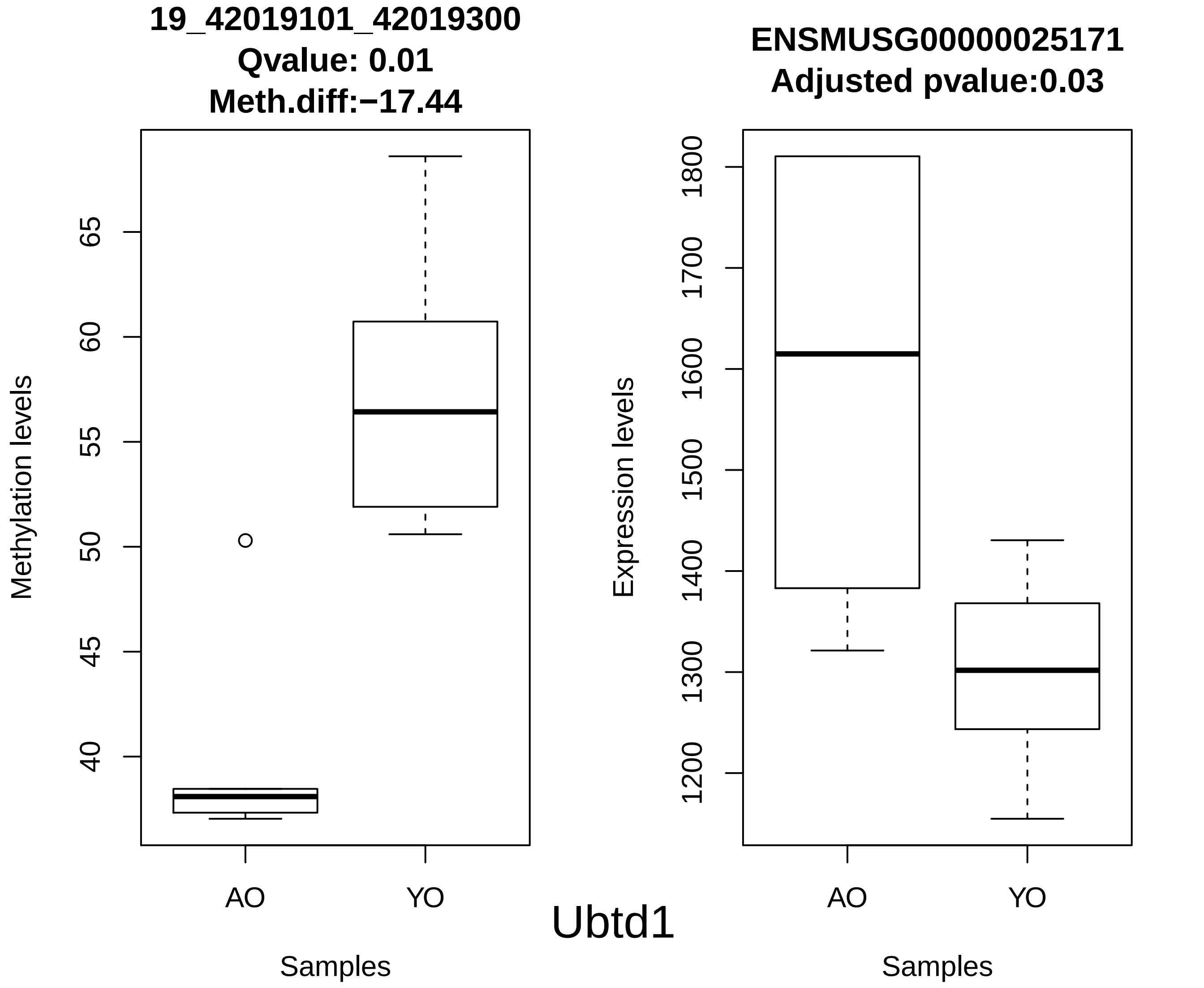
19_42019101_42019300
Qvalue: 0.01
Meth.diff:−17.44
Methylation levels
40
45
50
55
60
65
AO
YO
Samples
ENSMUSG00000025171
Adjusted pvalue:0.03
Expression levels
1200
1300
1400
1500
1600
1700
1800
AO
YO
Samples
Ubtd1

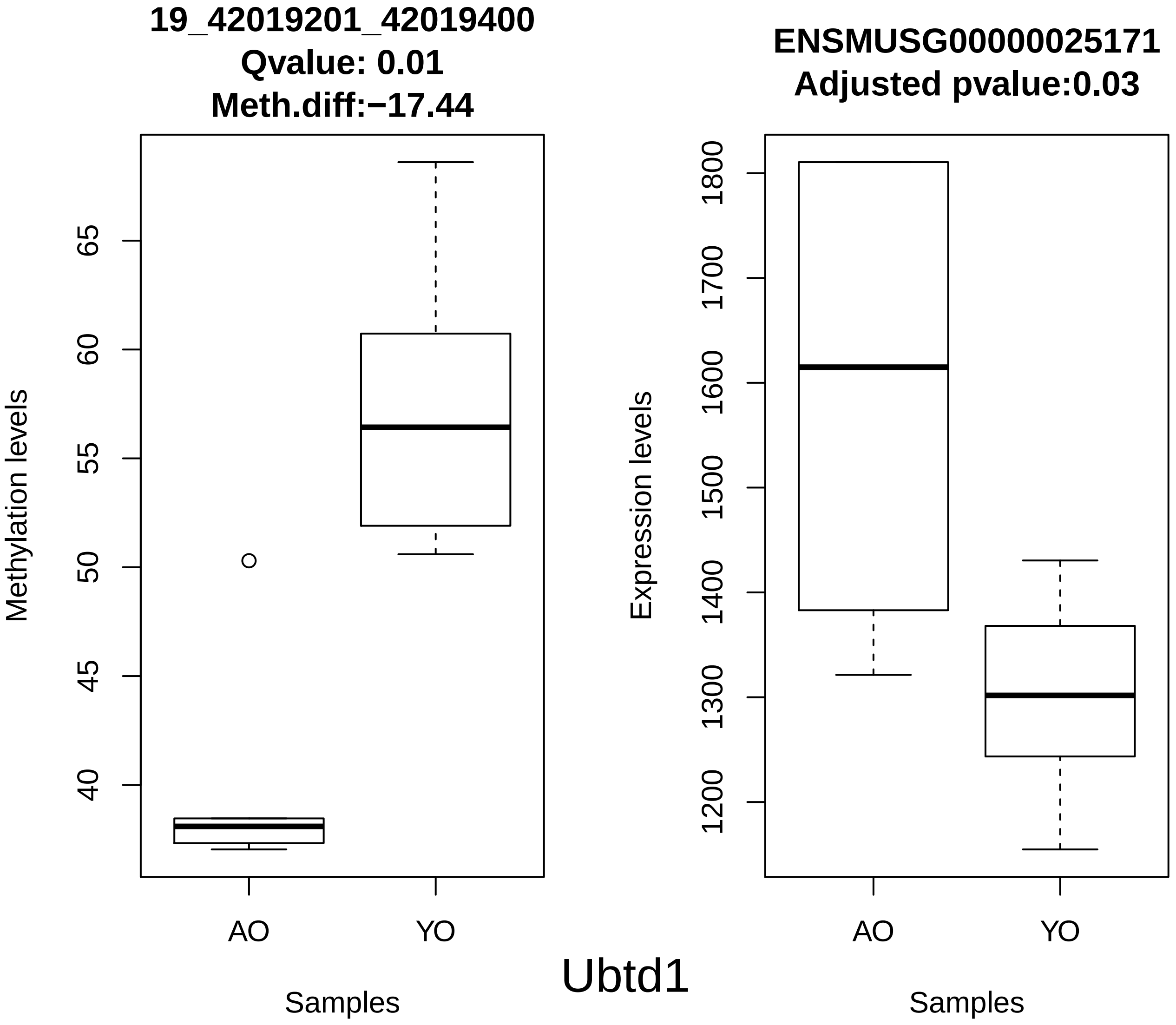
19_42019201_42019400
Qvalue: 0.01
Meth.diff:−17.44
Methylation levels
40
45
50
55
60
65
AO
YO
Samples
ENSMUSG00000025171
Adjusted pvalue:0.03
Expression levels
1200
1300
1400
1500
1600
1700
1800
AO
YO
Samples
Ubtd1

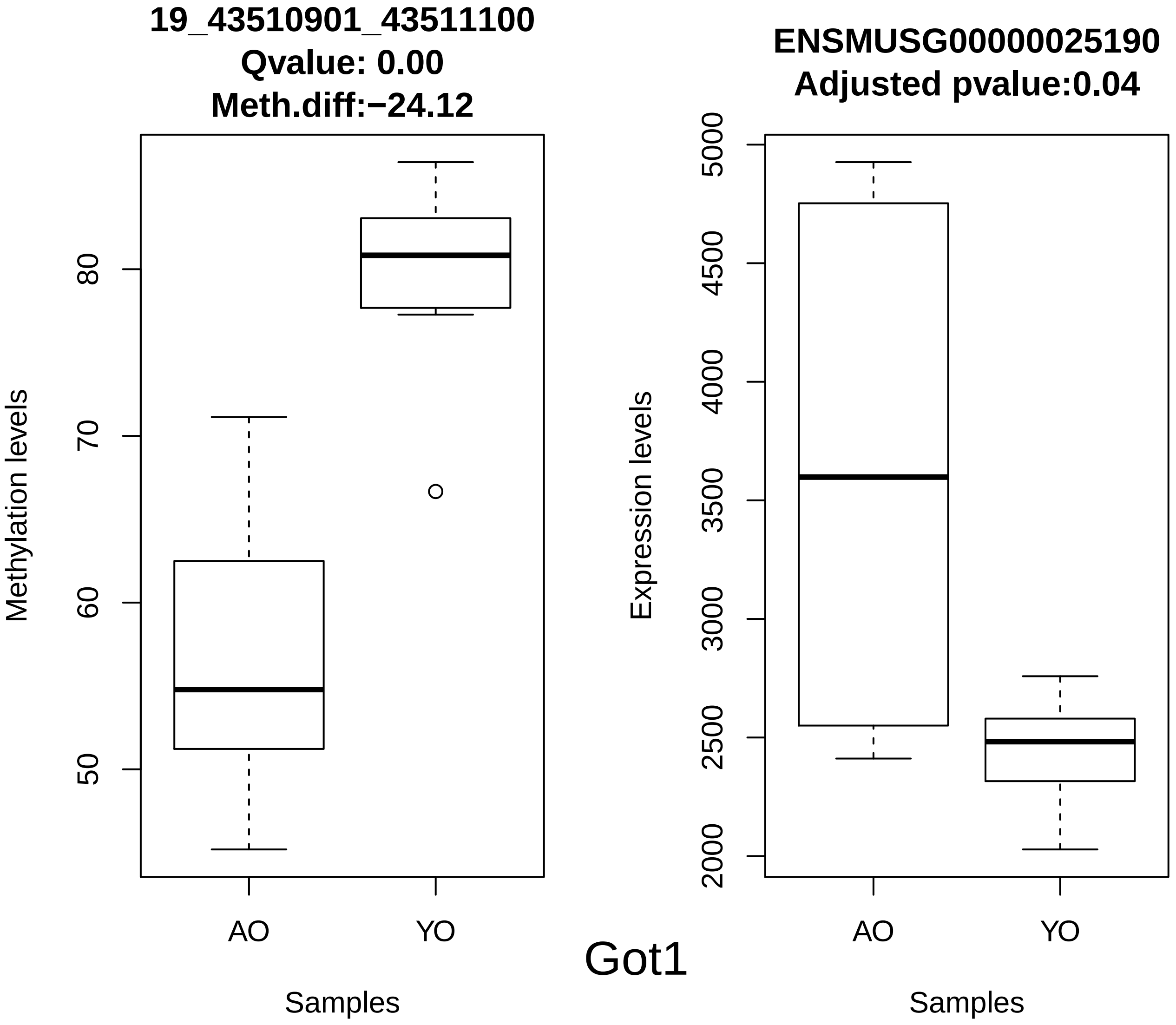
19_43510901_43511100
Qvalue: 0.00
Meth.diff:−24.12
Methylation levels
80
70
60
50
AO
YO
Samples
ENSMUSG00000025190
Adjusted pvalue:0.04
Expression levels
5000
4500
4000
3500
3000
2500
2000
AO
YO
Samples
Got1

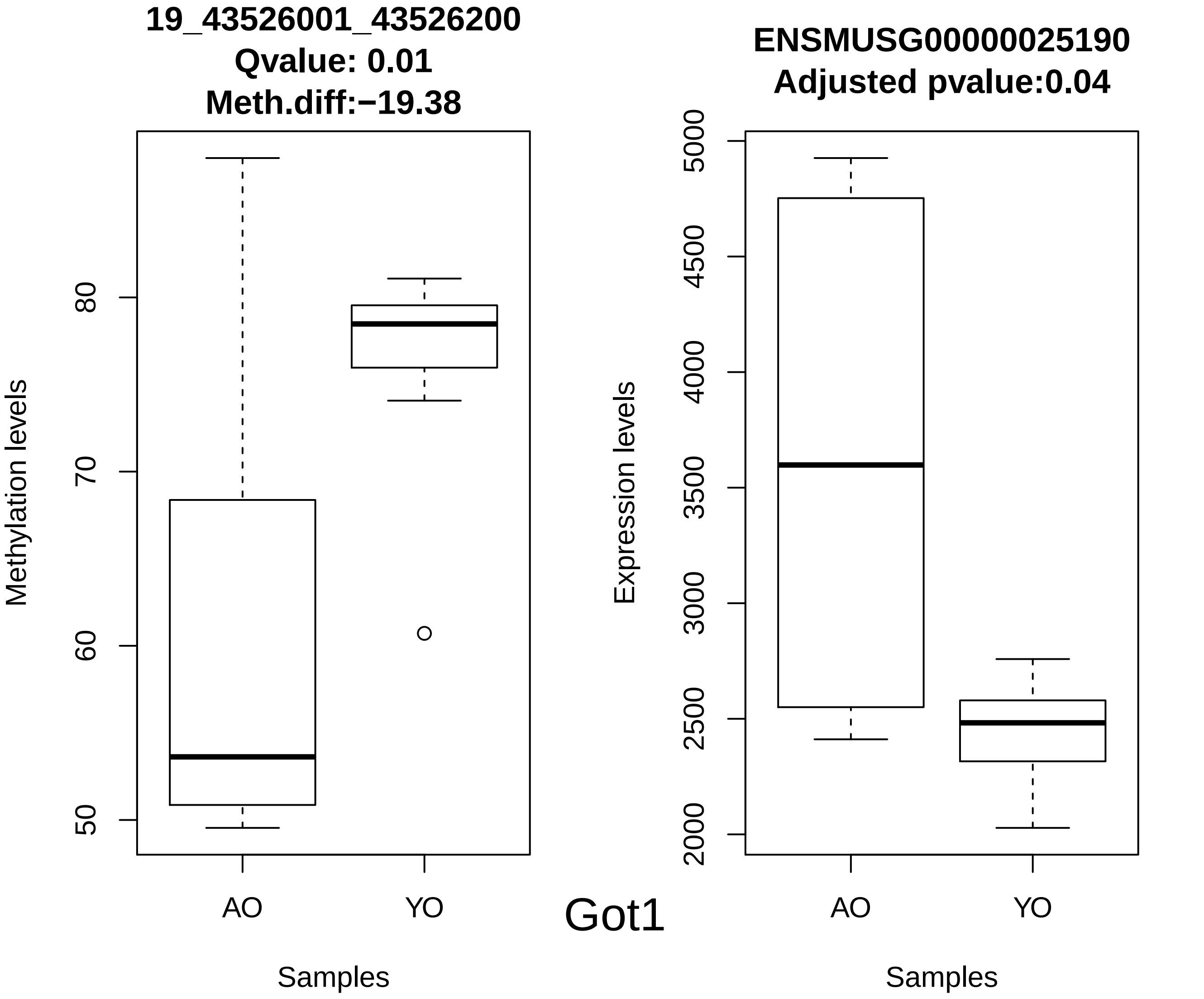

19_43526001_43526200
Qvalue: 0.01
Meth.diff:−19.38
Methylation levels
50
60
70
80
AO
YO
Samples
ENSMUSG00000025190
Adjusted pvalue:0.04
Expression levels
2000
2500
3000
3500
4000
4500
5000
AO
YO
Samples
Got1

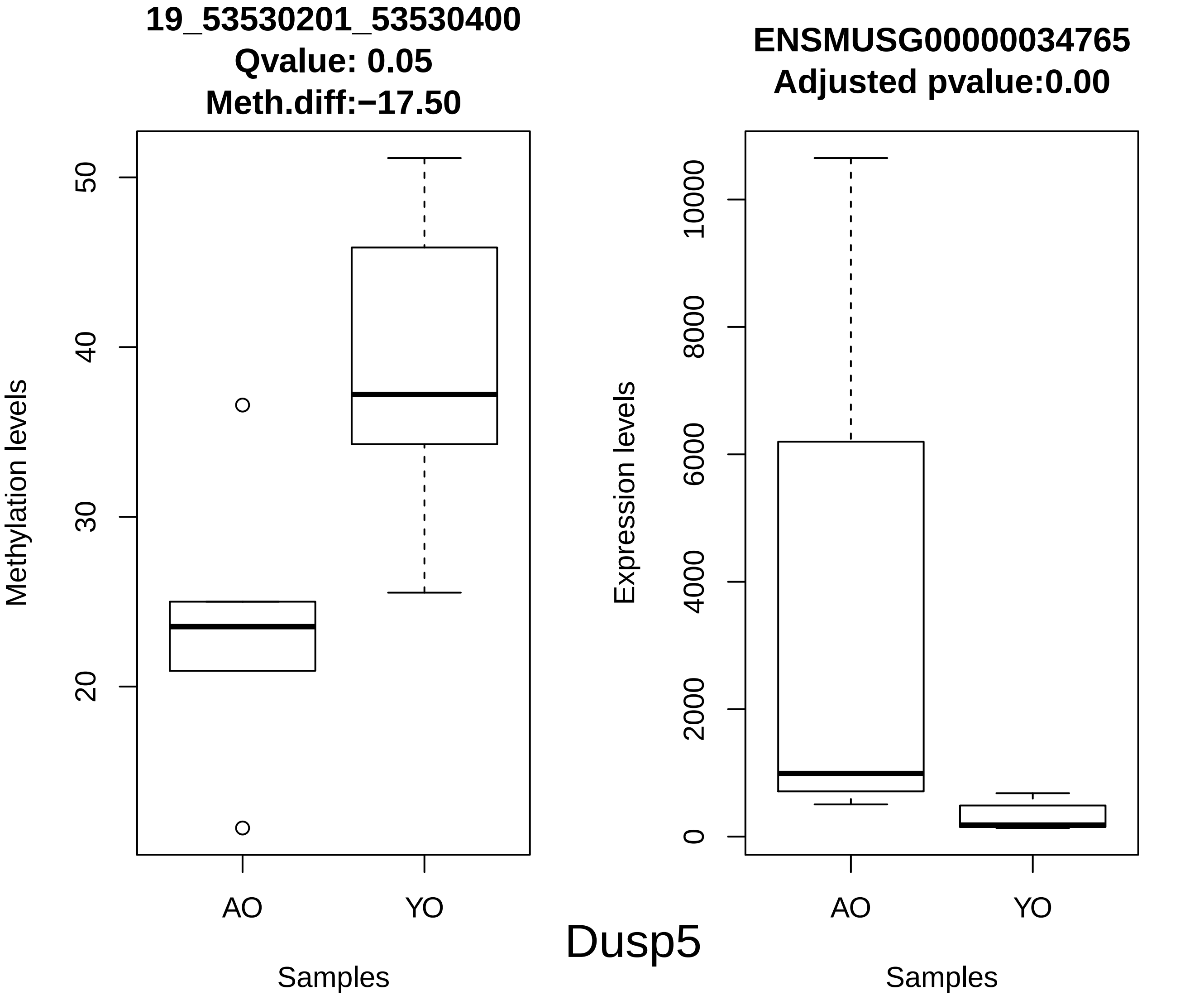
19_53530201_53530400
Qvalue: 0.05
Meth.diff:−17.50
Methylation levels
50
40
30
20
AO
YO
Samples
ENSMUSG00000034765
Adjusted pvalue:0.00
Expression levels
10000
8000
6000
4000
2000
0
AO
YO
Samples
Dusp5

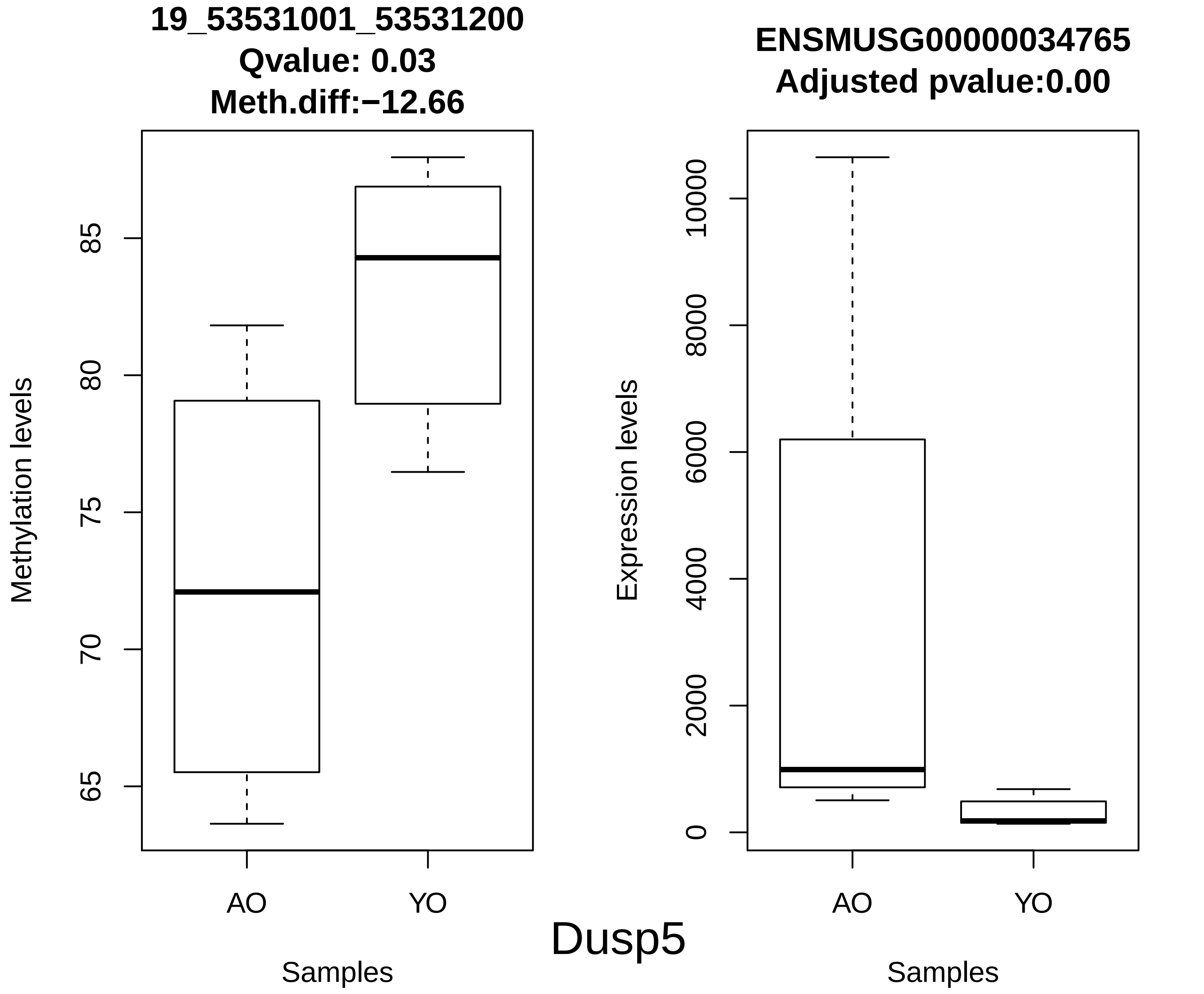
19_53531001_53531200
Qvalue: 0.03
Meth.diff:−12.66
Methylation levels
65
70
75
80
85
AO
YO
Samples
ENSMUSG00000034765
Adjusted pvalue:0.00
Expression levels
0
2000
4000
6000
8000
10000
AO
YO
Samples
Dusp5

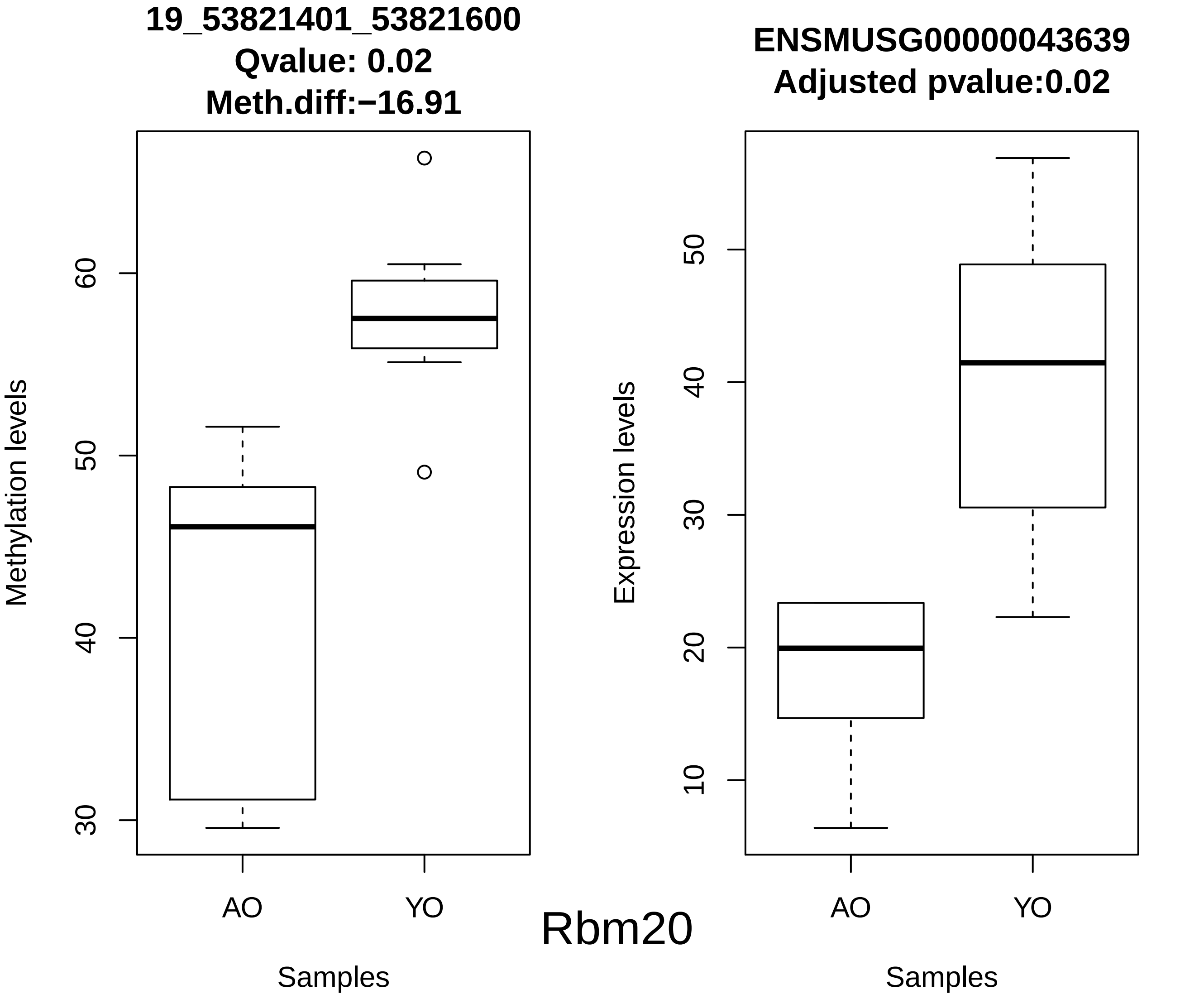
19_53821401_53821600
Qvalue: 0.02
Meth.diff:−16.91
Methylation levels
30
40
50
60
AO
YO
Samples
Rbm20
ENSMUSG00000043639
Adjusted pvalue:0.02
Expression levels
10
20
30
40
50
AO
YO
Samples

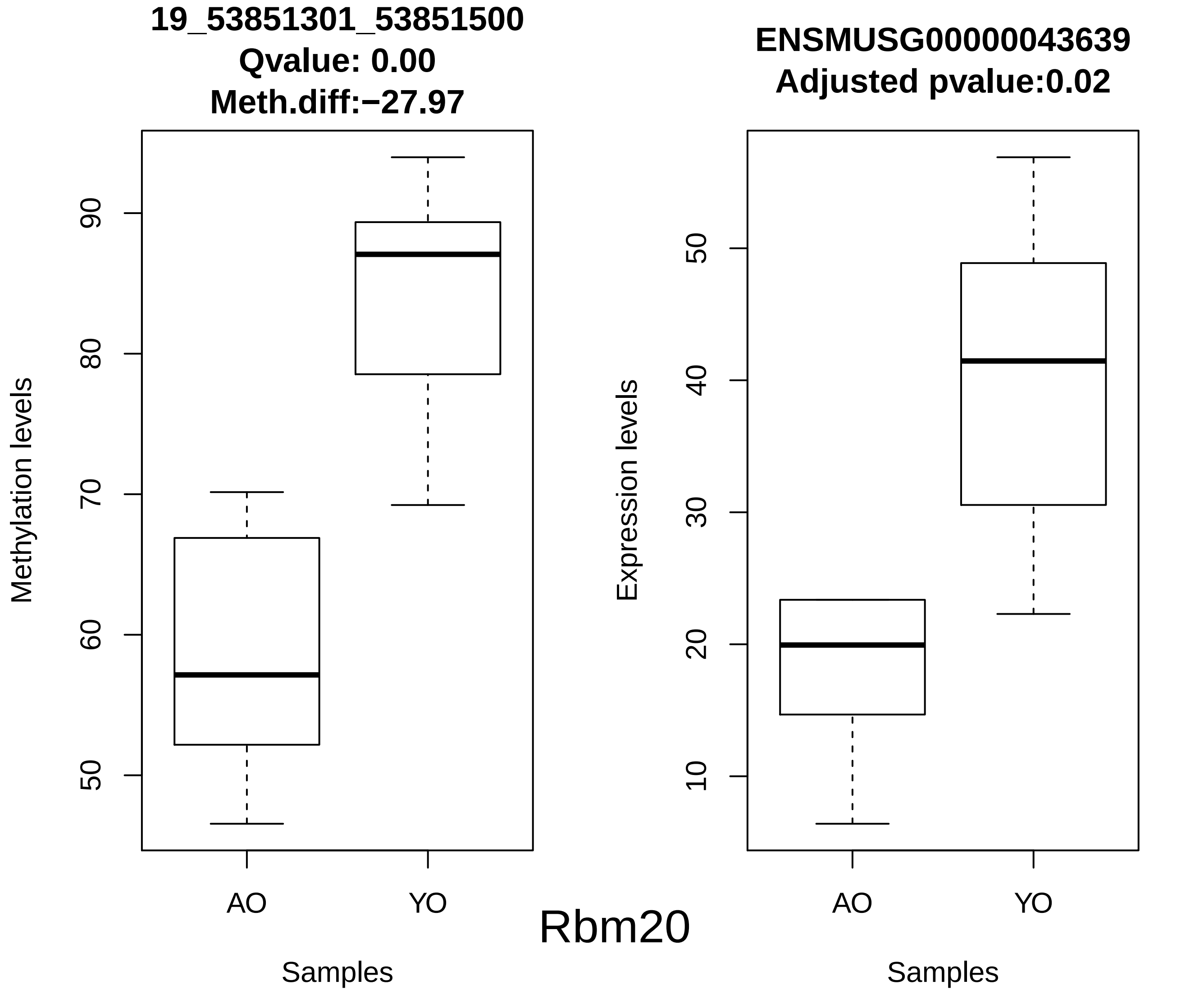
19_53851301_53851500
Qvalue: 0.00
Meth.diff:−27.97
Methylation levels
90
80
70
60
50
AO
YO
Samples
ENSMUSG00000043639
Adjusted pvalue:0.02
Expression levels
50
40
30
20
10
AO
YO
Samples
Rbm20

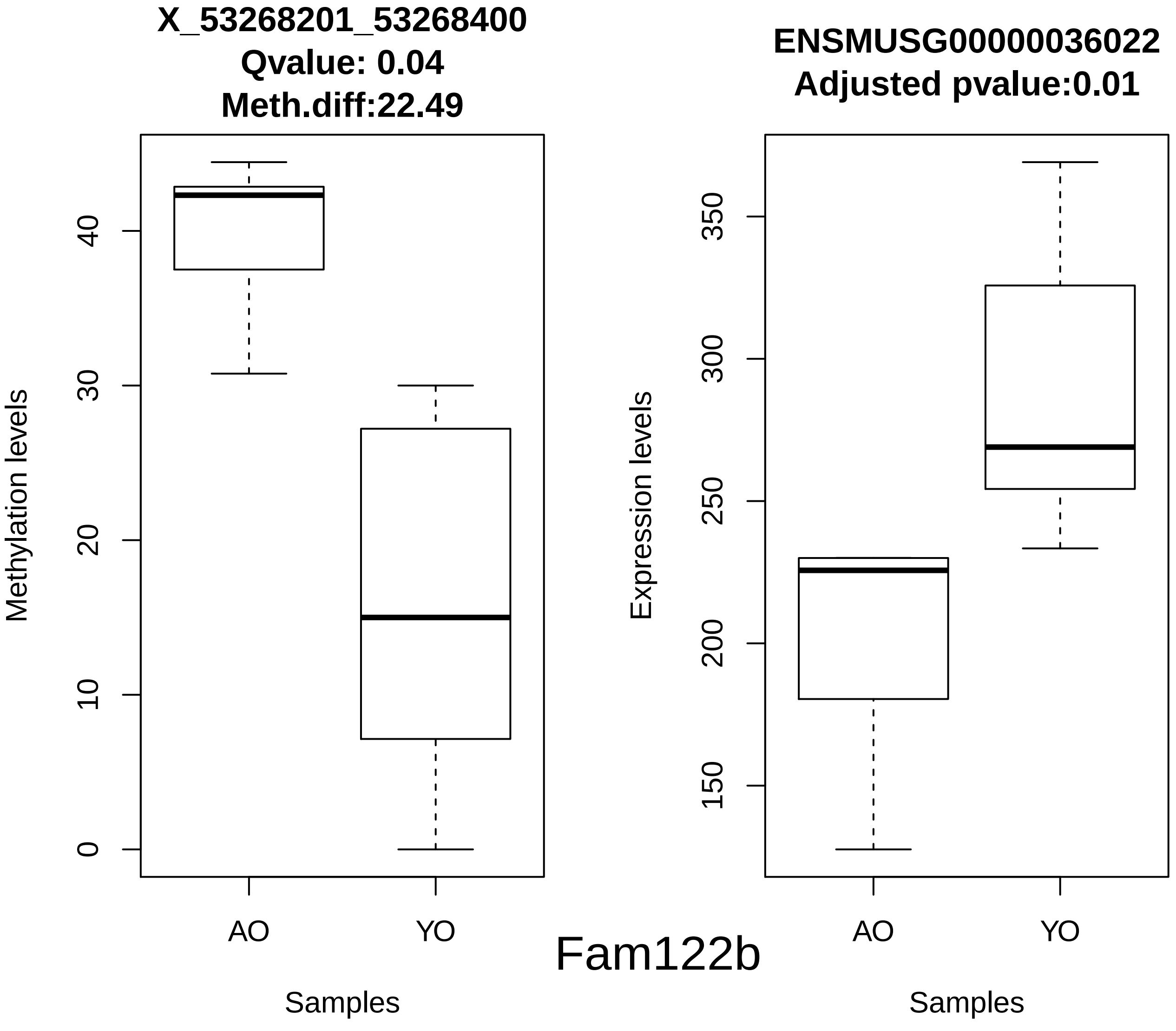
X_53268201_53268400
Qvalue: 0.04
Meth.diff:22.49
Methylation levels
0
10
20
30
40
AO
YO
Samples
ENSMUSG00000036022
Adjusted pvalue:0.01
Expression levels
150
200
250
300
350
AO
YO
Samples

Fam122b

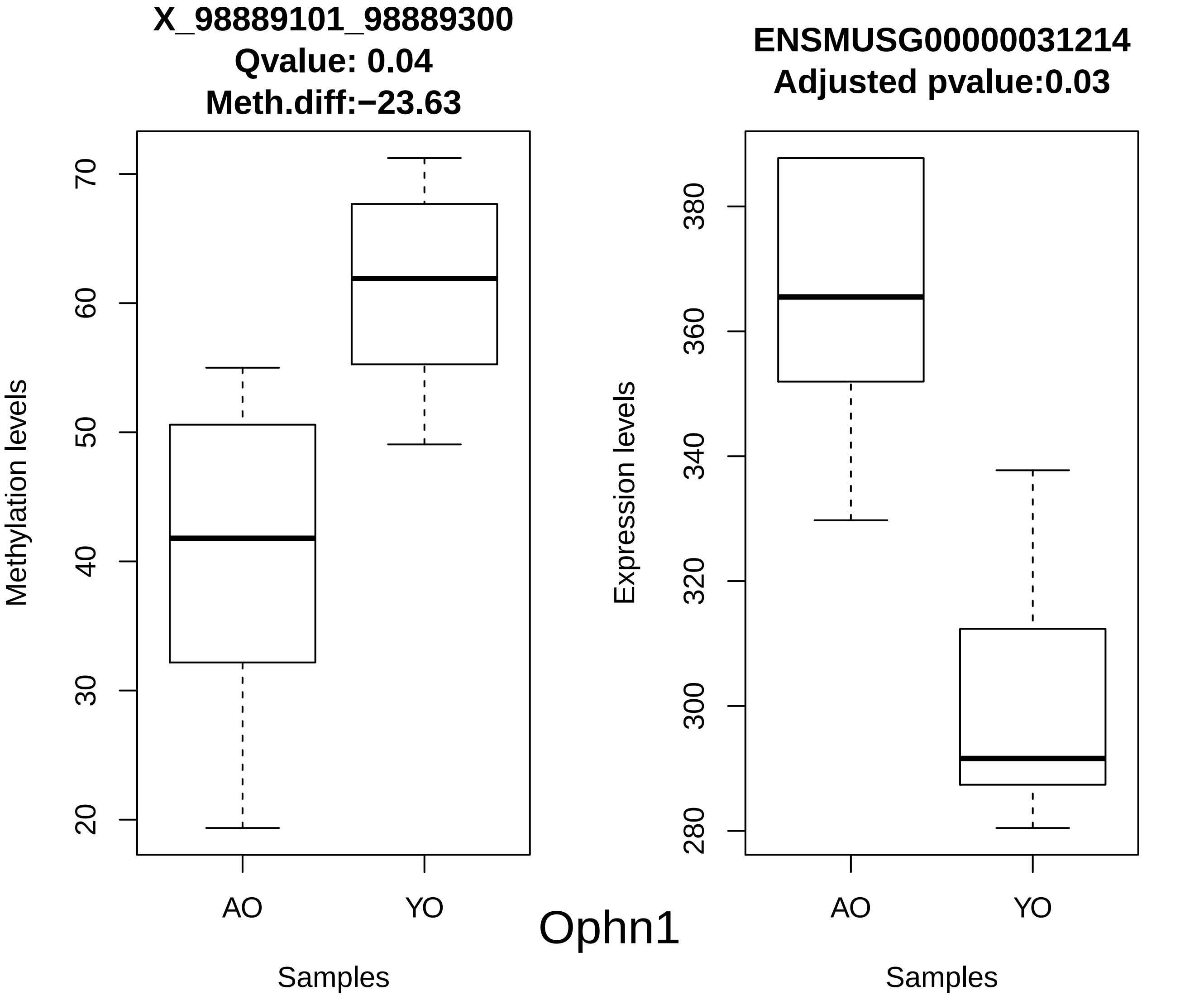
X_98889101_98889300
Qvalue: 0.04
Meth.diff:−23.63
Methylation levels
70
60
50
40
30
20
AO
YO
Samples
ENSMUSG00000031214
Adjusted pvalue:0.03
Expression levels
380
360
340
320
300
280
AO
YO
Samples
Ophn1

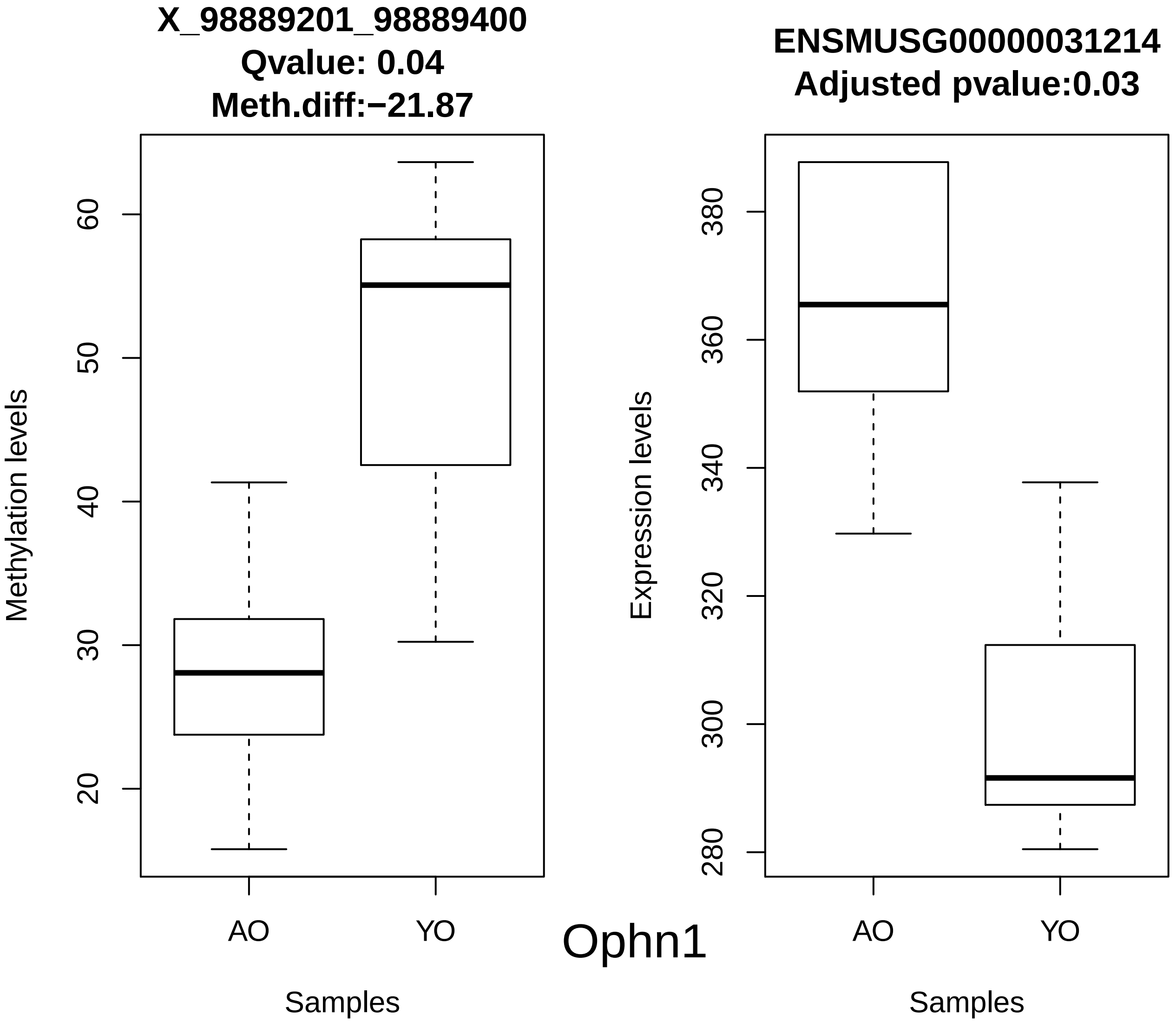
X_98889201_98889400
Qvalue: 0.04
Meth.diff:−21.87
Methylation levels
60
50
40
30
20
AO
YO
Samples
ENSMUSG00000031214
Adjusted pvalue:0.03
Expression levels
380
360
340
320
300
280
AO
YO
Samples
Ophn1

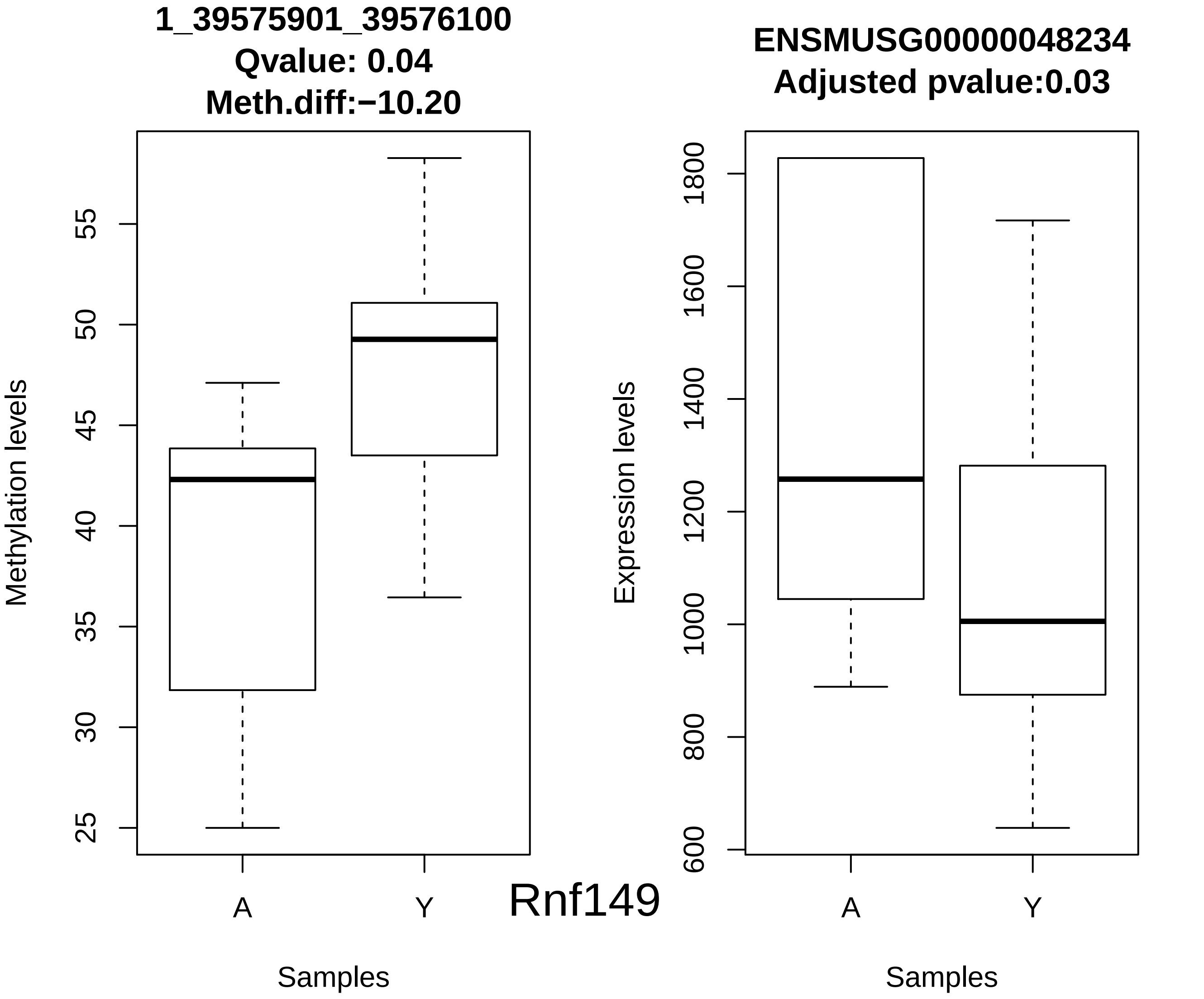
1_39575901_39576100
Qvalue: 0.04
Meth.diff:−10.20
Methylation levels
25
30
35
40
45
50
55
A
Y
Samples
Rnf149
ENSMUSG00000048234
Adjusted pvalue:0.03
Expression levels
600
800
1000
1200
1400
1600
1800
A
Y
Samples

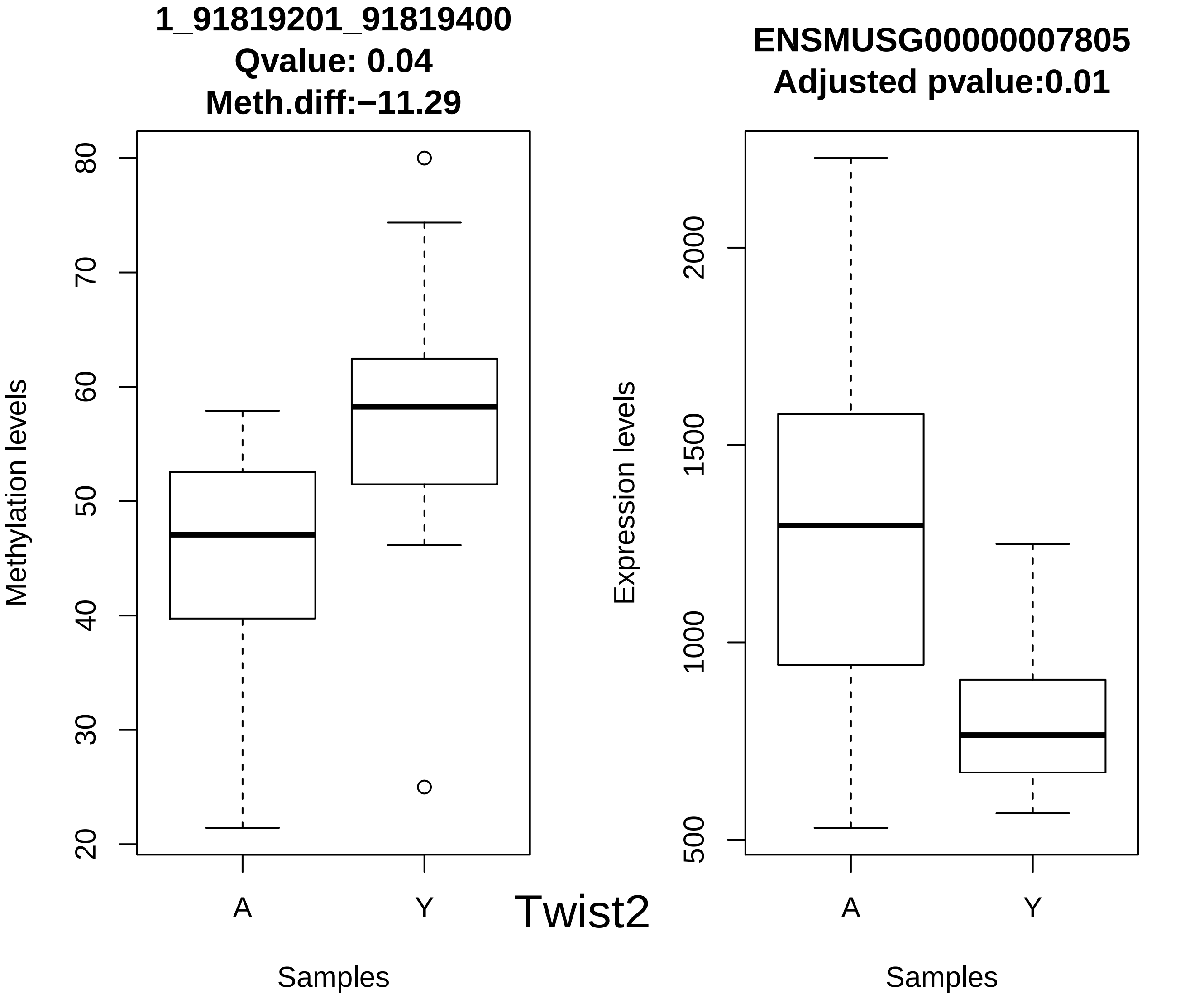
1_91819201_91819400
Qvalue: 0.04
Meth.diff:−11.29
Methylation levels
20
30
40
50
60
70
80
A
Y
Samples
Twist2
ENSMUSG00000007805
Adjusted pvalue:0.01
Expression levels
500
1000
1500
2000
A
Y
Samples

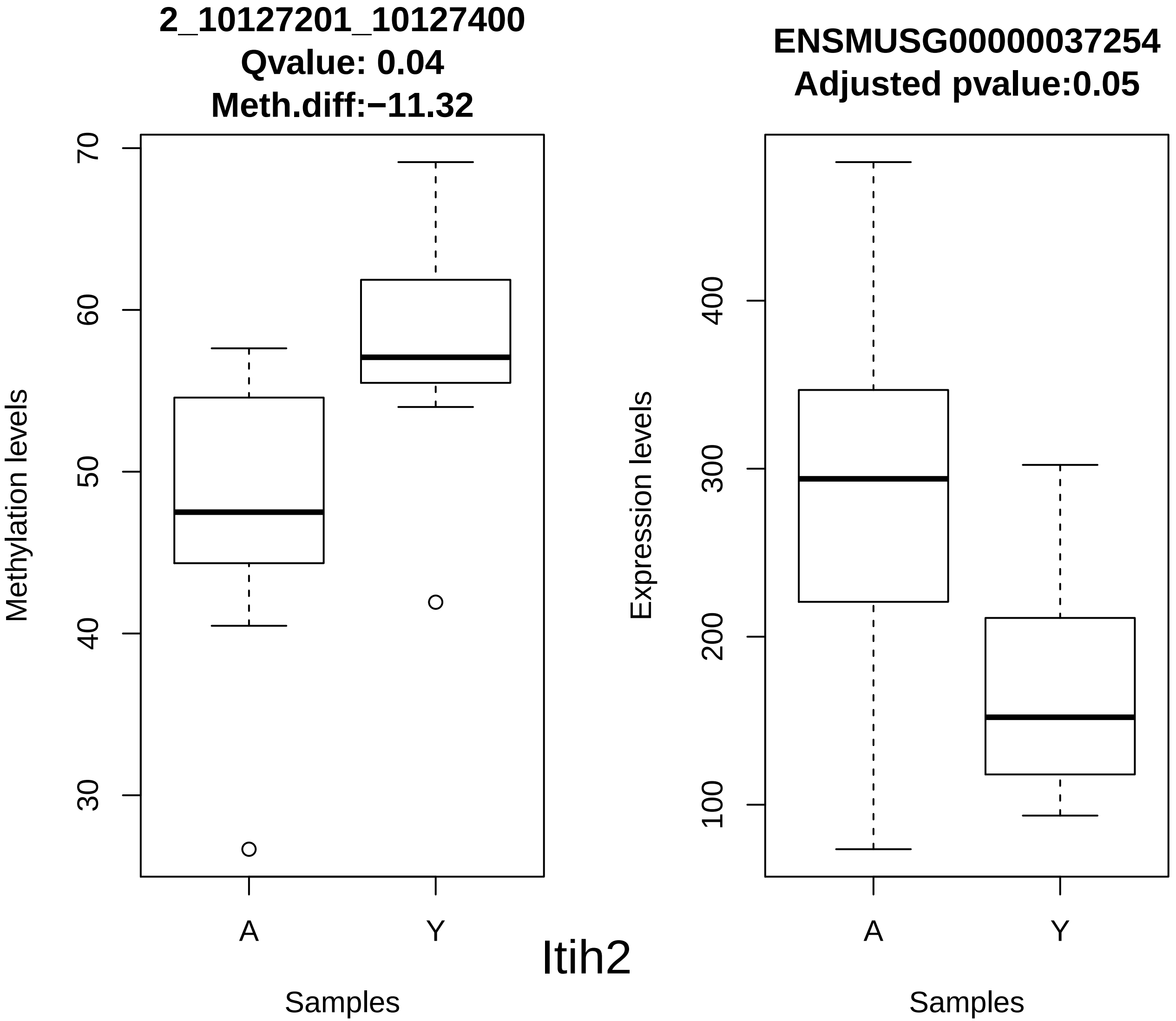
2_10127201_10127400
Qvalue: 0.04
Meth.diff:−11.32
Methylation levels
70
60
50
40
30
A
Y
Samples
ENSMUSG00000037254
Adjusted pvalue:0.05
Expression levels
400
300
200
100
A
Y
Samples
Itih2

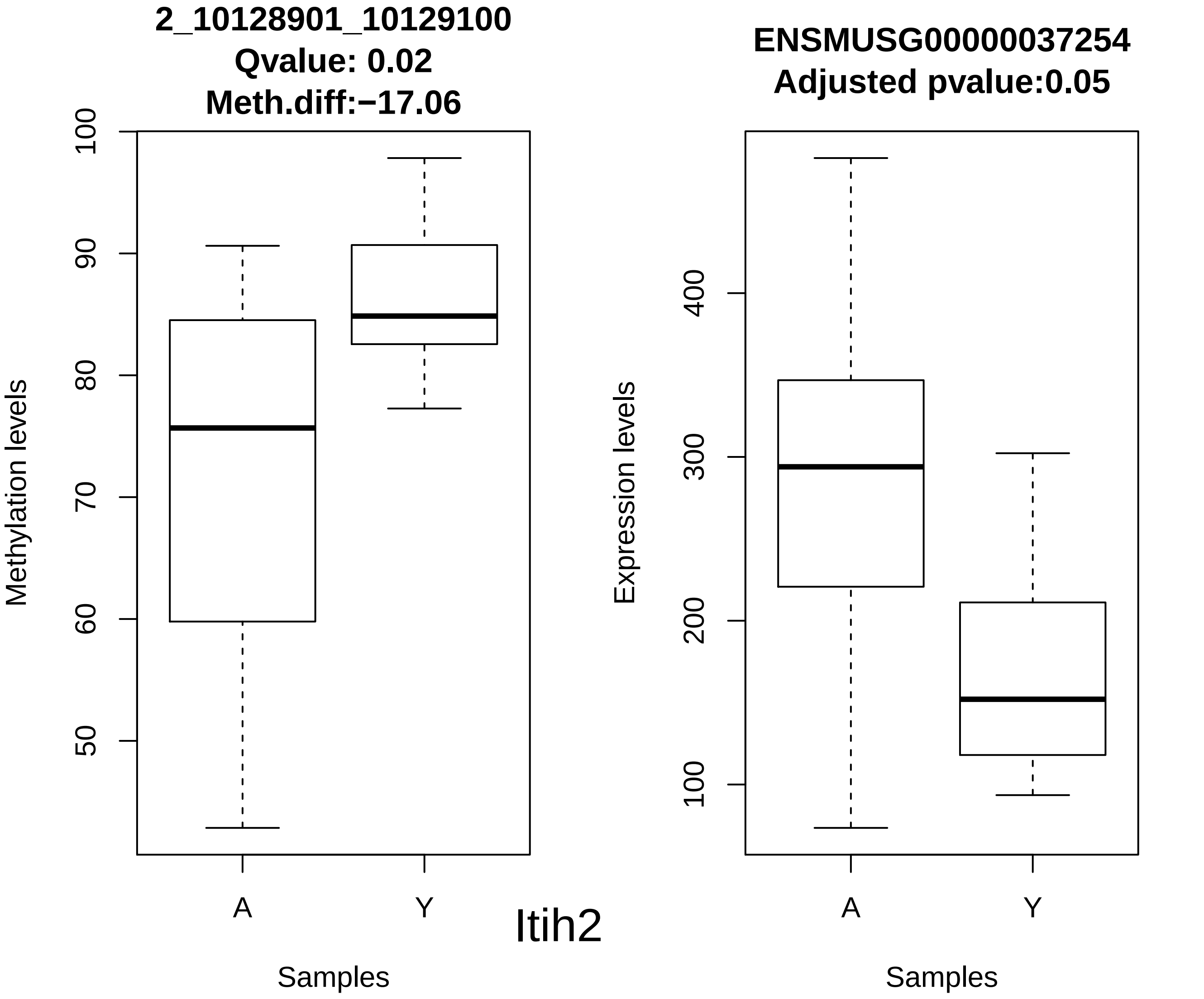
2_10128901_10129100
Qvalue: 0.02
Meth.diff:−17.06
Methylation levels
100
90
80
70
60
50
A
Y
Samples
ENSMUSG00000037254
Adjusted pvalue:0.05
Expression levels
400
300
200
100
A
Y
Samples

Itih2

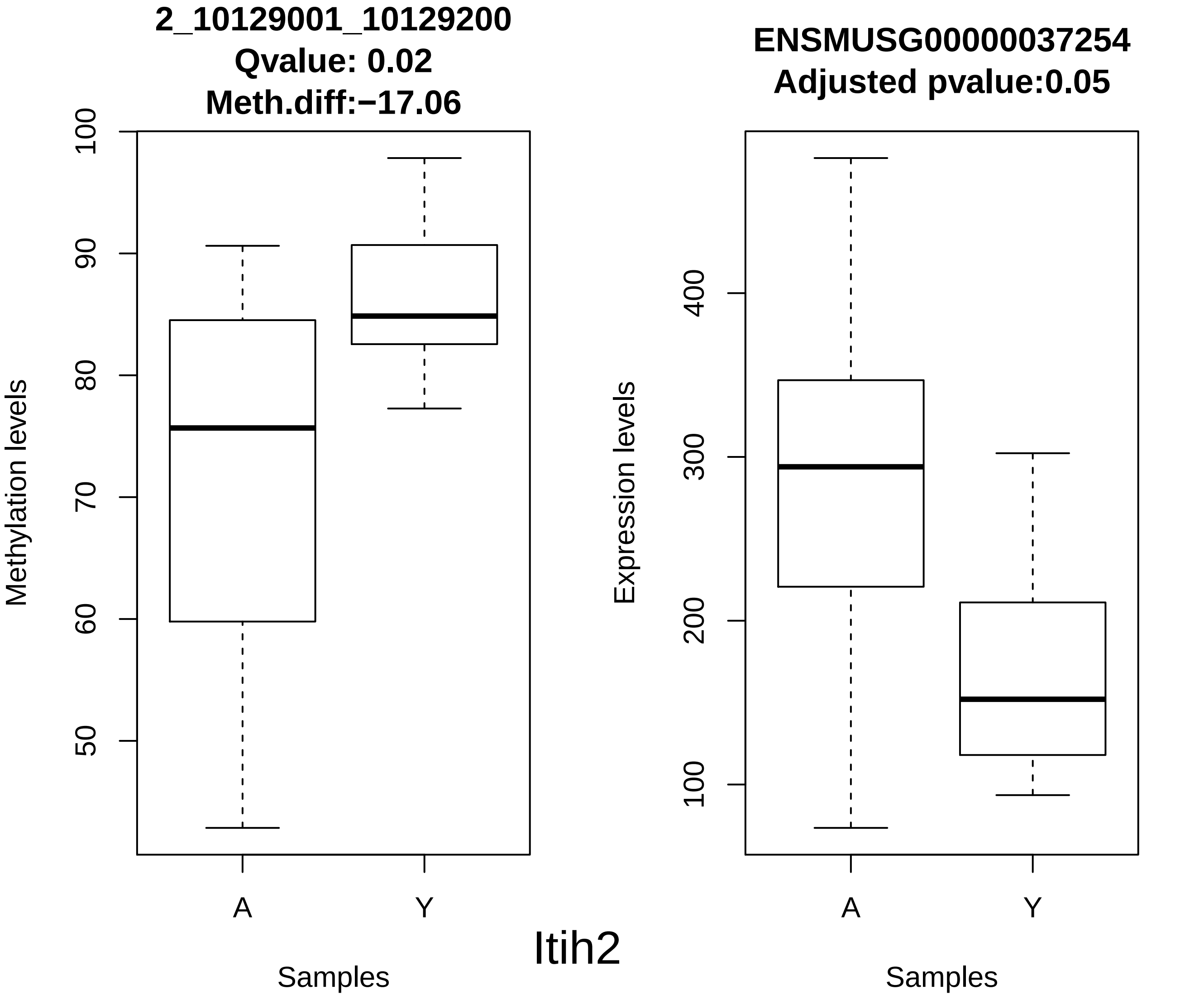
2_10129001_10129200
Qvalue: 0.02
Meth.diff:−17.06
Methylation levels
100
90
80
70
60
50
A
Y
Samples
ENSMUSG00000037254
Adjusted pvalue:0.05
Expression levels
400
300
200
100
A
Y
Samples
Itih2

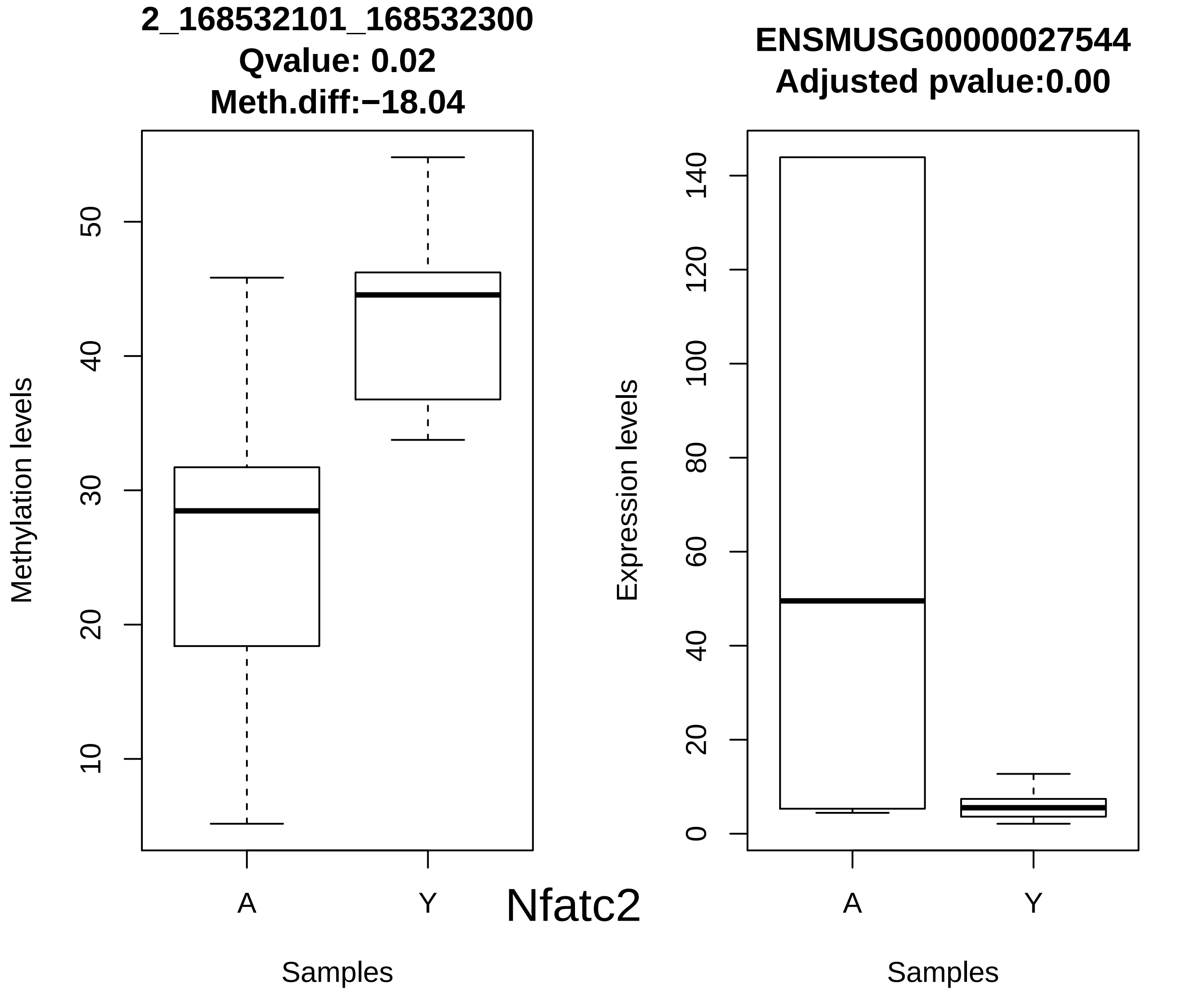
2_168532101_168532300
Qvalue: 0.02
Meth.diff:−18.04
Methylation levels
10
20
30
40
50
A
Y
Samples
ENSMUSG00000027544
Adjusted pvalue:0.00
Expression levels
0
20
40
60
80
100
120
140
A
Y
Samples
Nfatc2

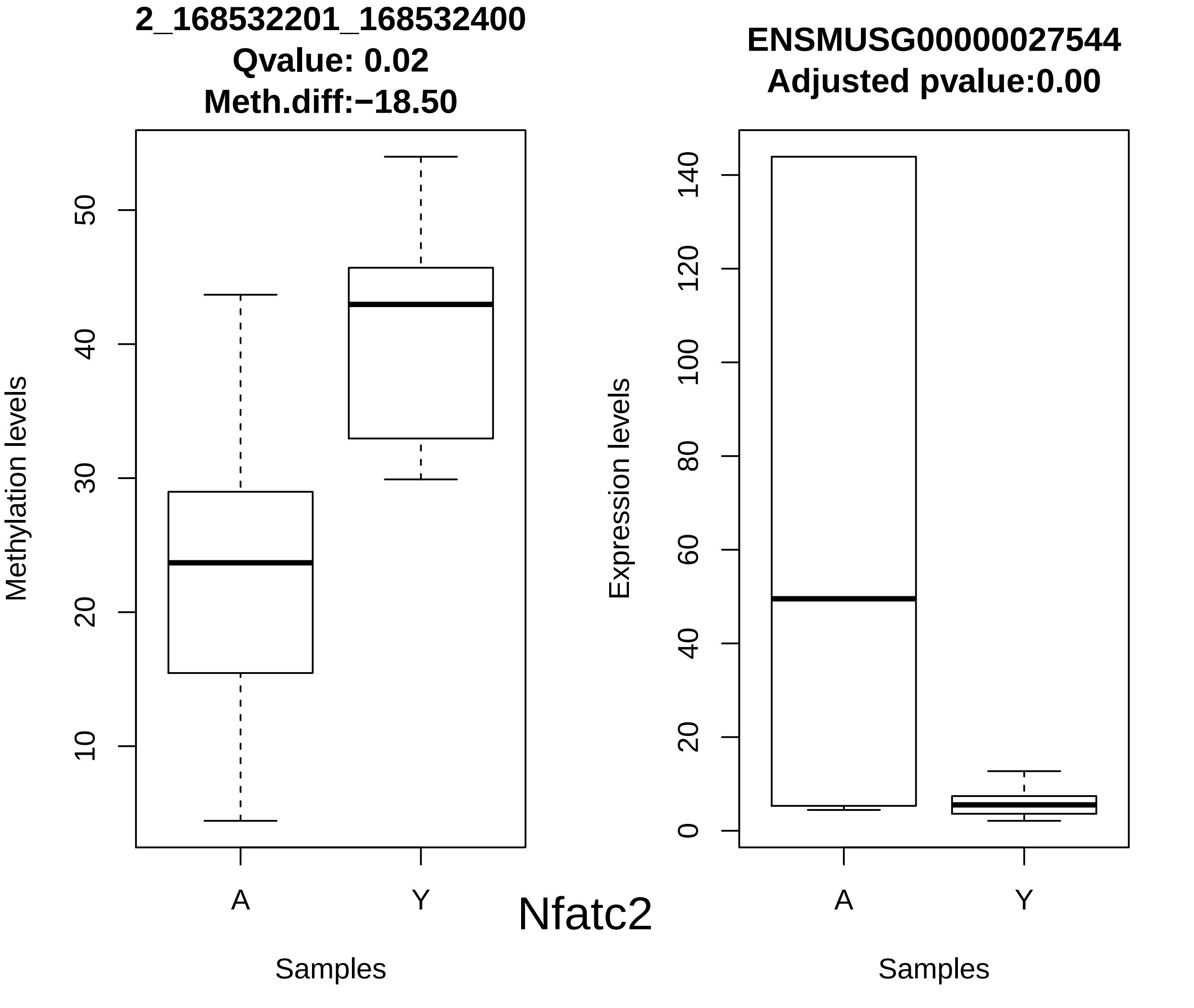
2_168532201_168532400
Qvalue: 0.02
Meth.diff:−18.50
Methylation levels
10
20
30
40
50
A
Y
Samples
ENSMUSG00000027544
Adjusted pvalue:0.00
Expression levels
0
20
40
60
80
100
120
140
A
Y
Samples

Nfatc2

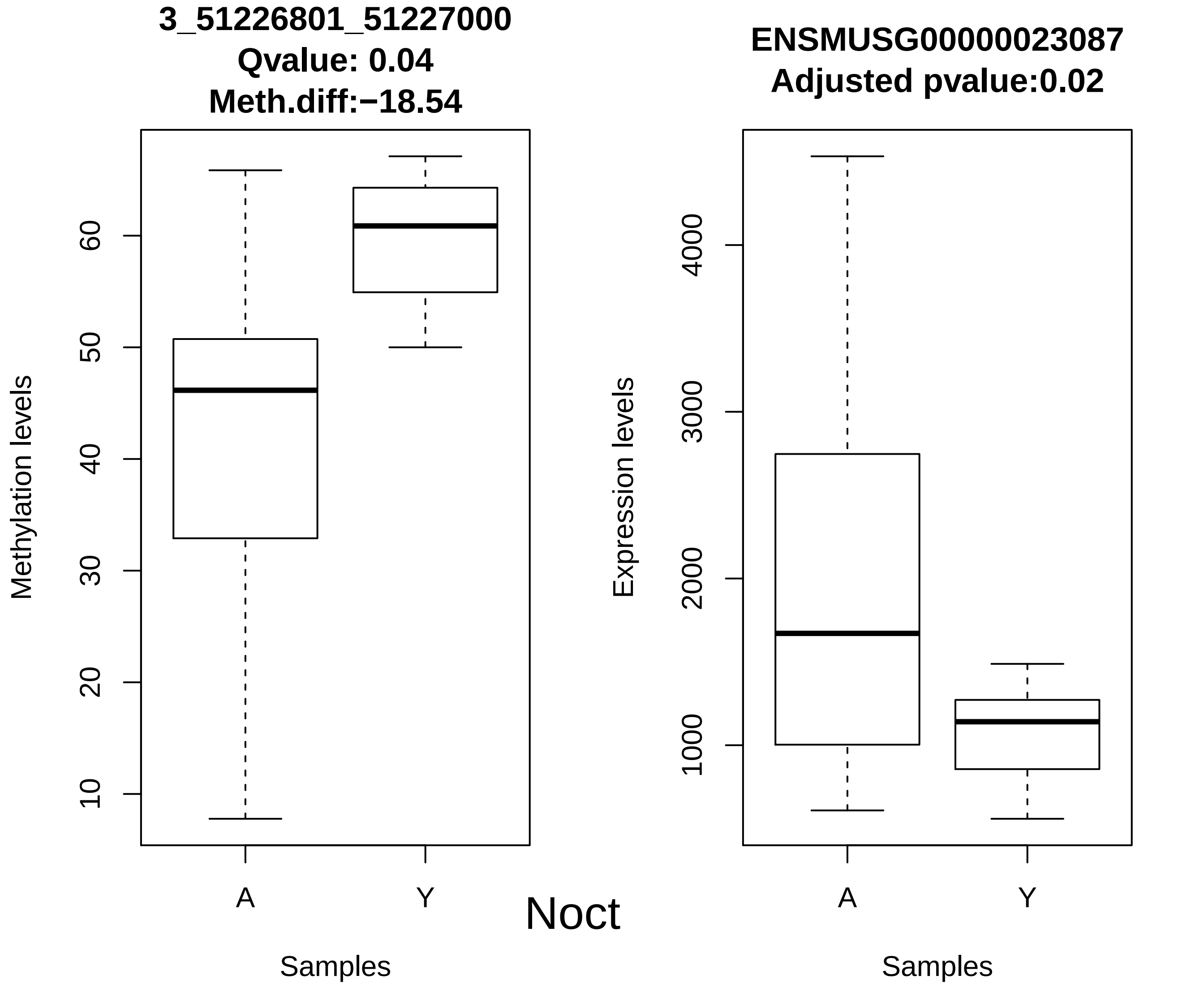
3_51226801_51227000
Qvalue: 0.04
Meth.diff:−18.54
Methylation levels
10
20
30
40
50
60
A
Y
Samples
Noct
ENSMUSG00000023087
Adjusted pvalue:0.02
Expression levels
1000
2000
3000
4000
A
Y
Samples

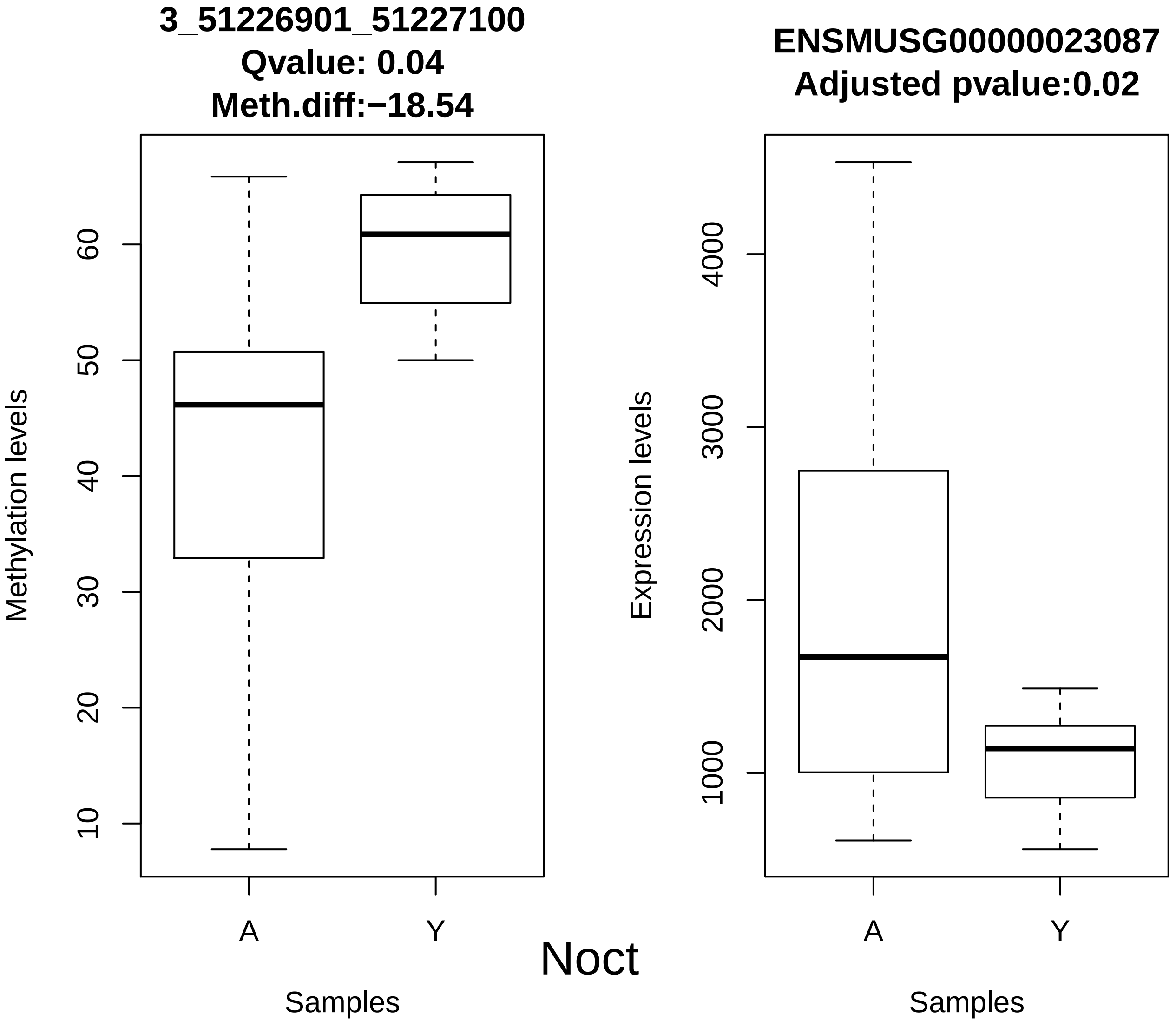
3_51226901_51227100
Qvalue: 0.04
Meth.diff:−18.54
Methylation levels
10
20
30
40
50
60
A
Y
Samples
ENSMUSG00000023087
Adjusted pvalue:0.02
Expression levels
1000
2000
3000
4000
A
Y
Samples
Noct

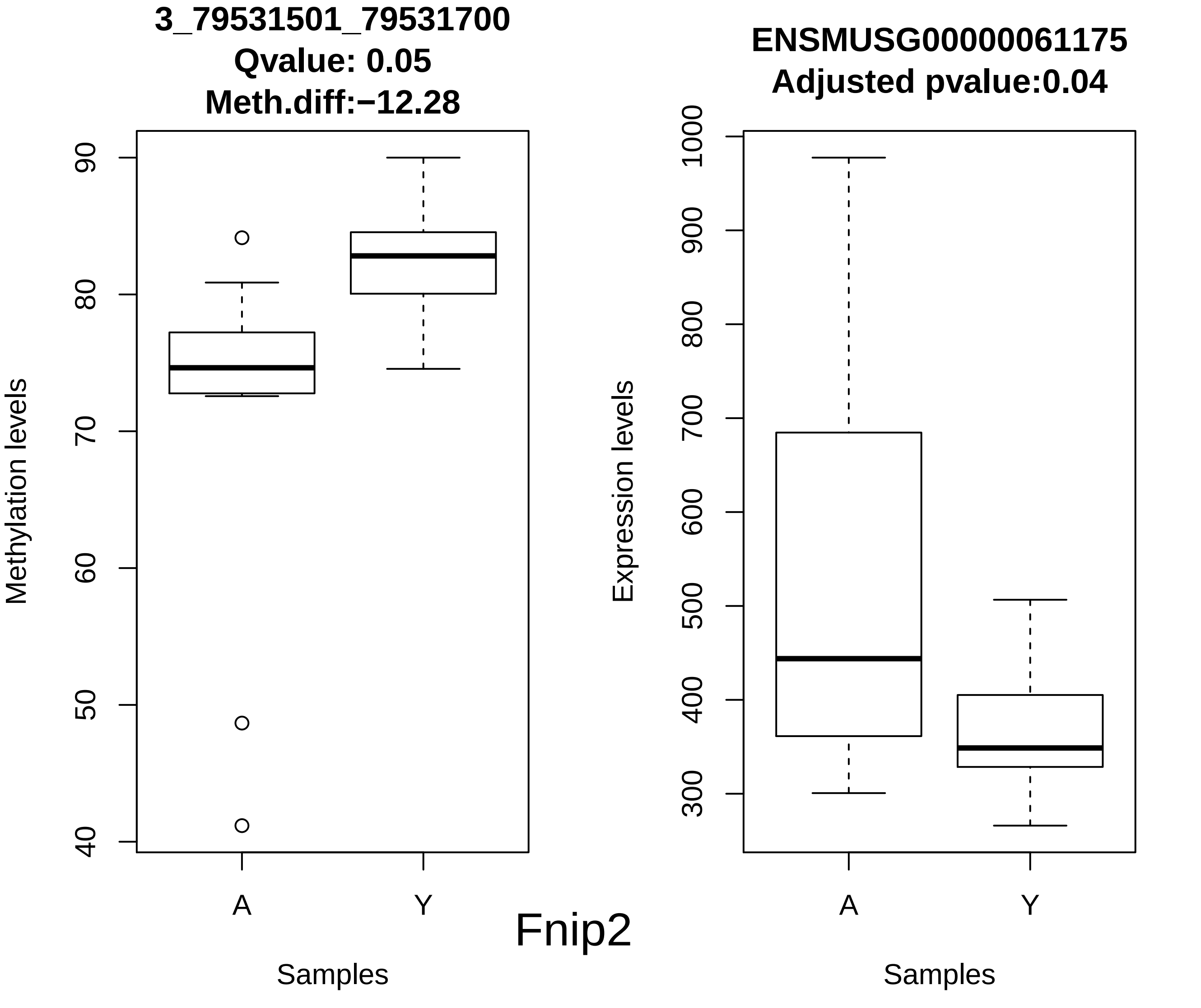
3_79531501_79531700
Qvalue: 0.05
Meth.diff:−12.28
Methylation levels
90
80
70
60
50
40
A
Y
Samples
ENSMUSG00000061175
Adjusted pvalue:0.04
Expression levels
1000
900
800
700
600
500
400
300
A
Y
Samples
Fnip2

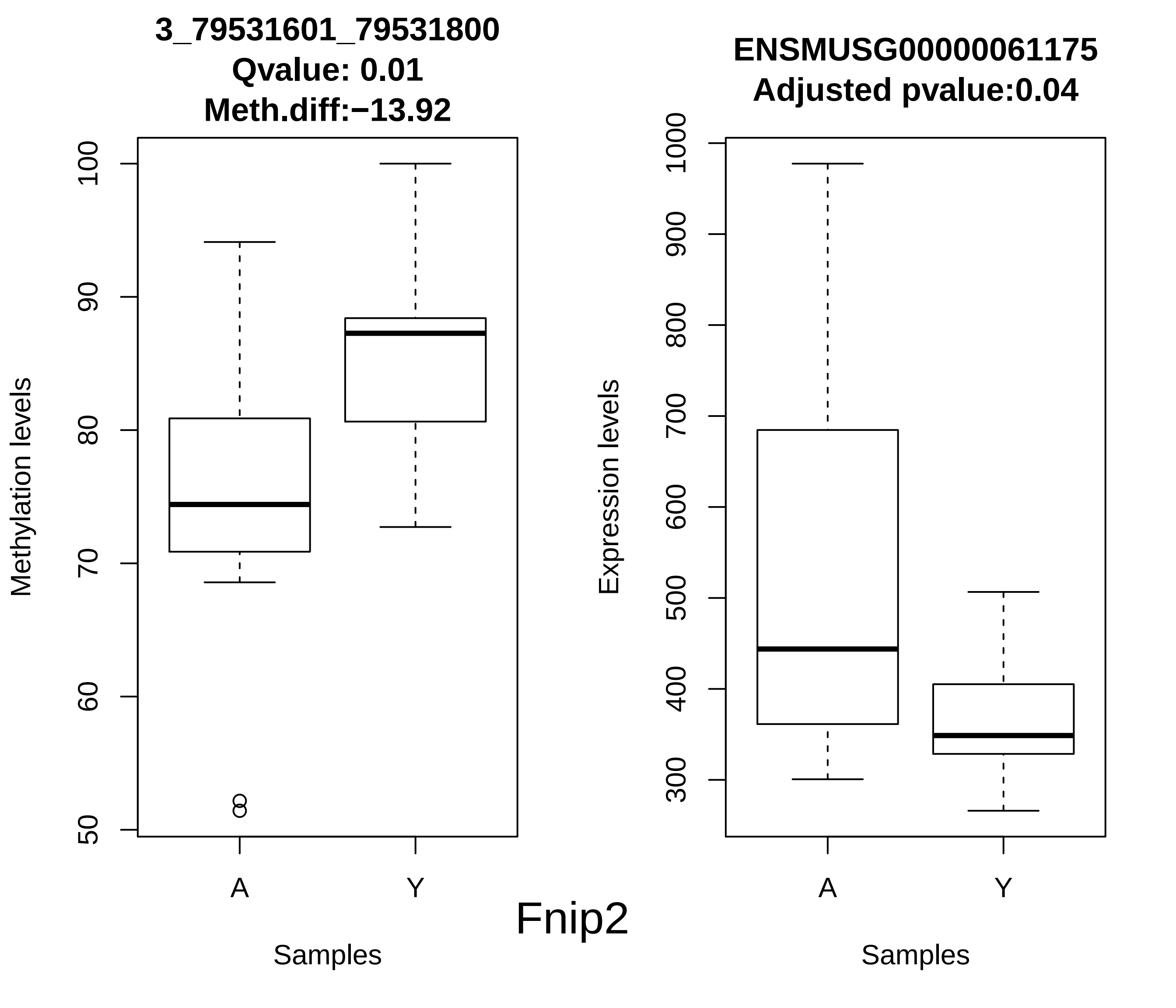
3_79531601_79531800
Qvalue: 0.01
Meth.diff:−13.92
Methylation levels
100
90
80
70
60
50
A
Y
Samples
ENSMUSG00000061175
Adjusted pvalue:0.04
Expression levels
1000
900
800
700
600
500
400
300
A
Y
Samples
Fnip2

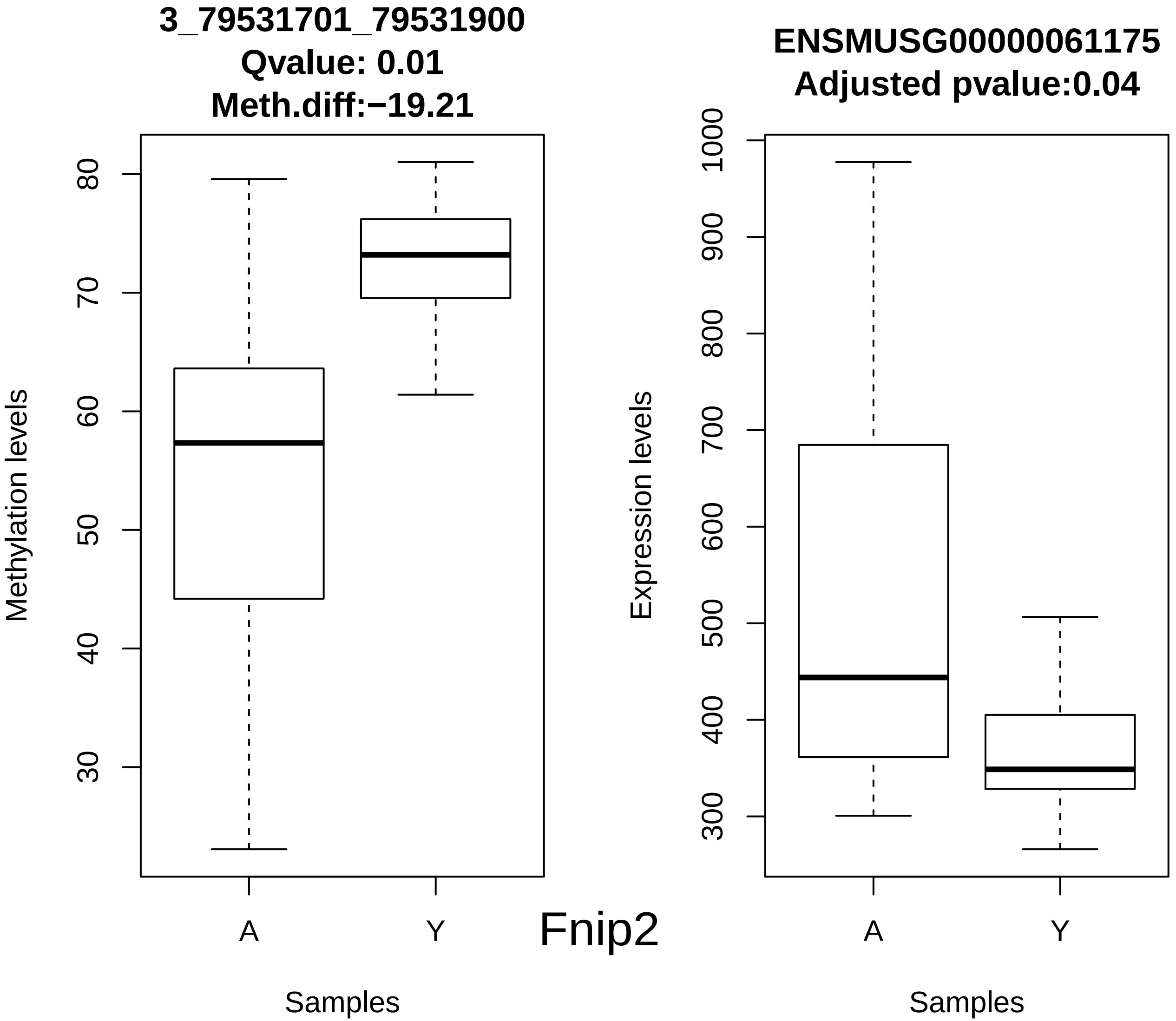
3_79531701_79531900
Qvalue: 0.01
Meth.diff:−19.21
Methylation levels
80
70
60
50
40
30
A
Y
Samples
Fnip2
ENSMUSG00000061175
Adjusted pvalue:0.04
Expression levels
1000
900
800
700
600
500
400
300
A
Y
Samples

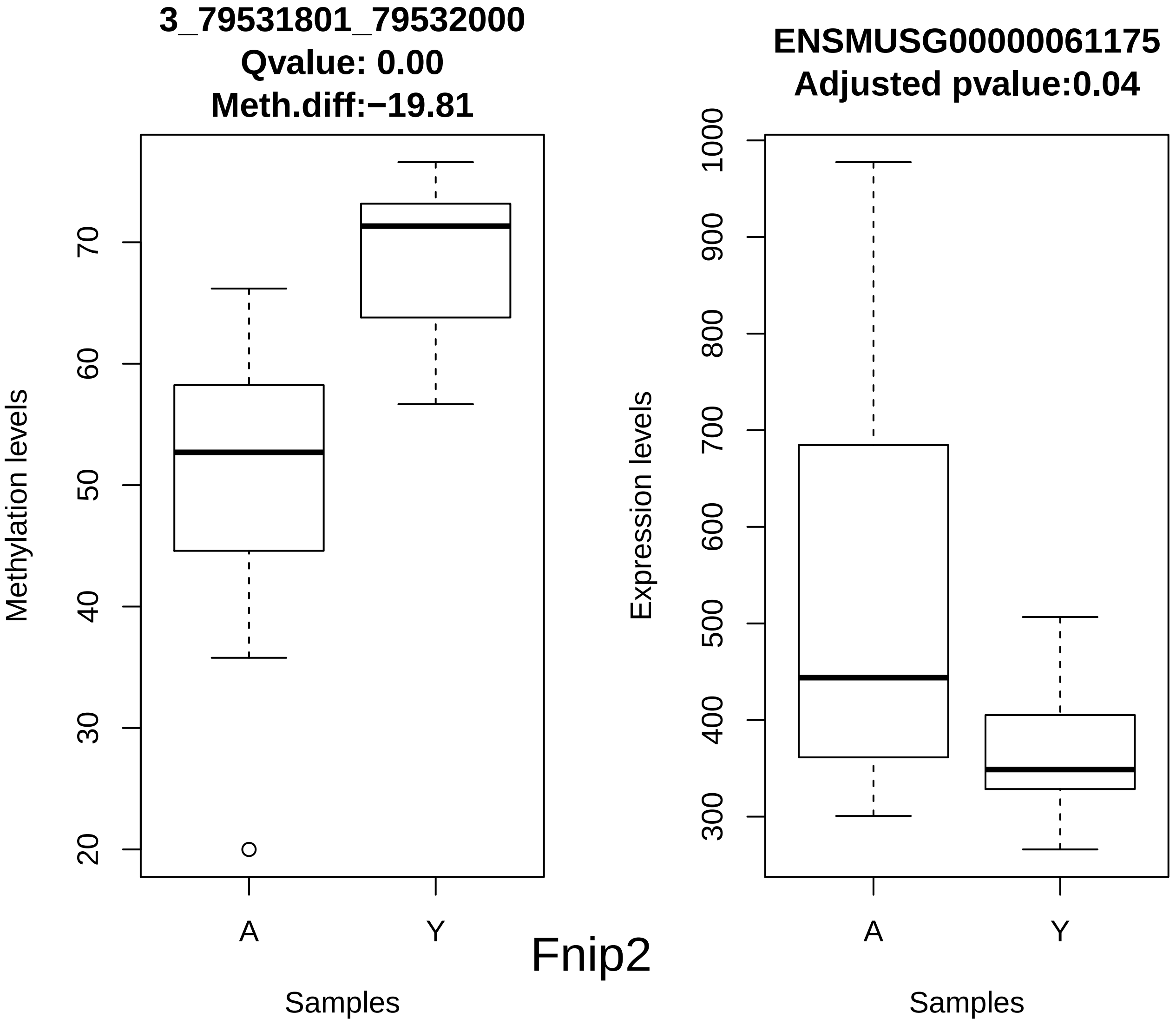
3_79531801_79532000
Qvalue: 0.00
Meth.diff:−19.81
Methylation levels
20
30
40
50
60
70
A
Y
Samples
ENSMUSG00000061175
Adjusted pvalue:0.04
Expression levels
300
400
500
600
700
800
900
1000
A
Y
Samples
Fnip2

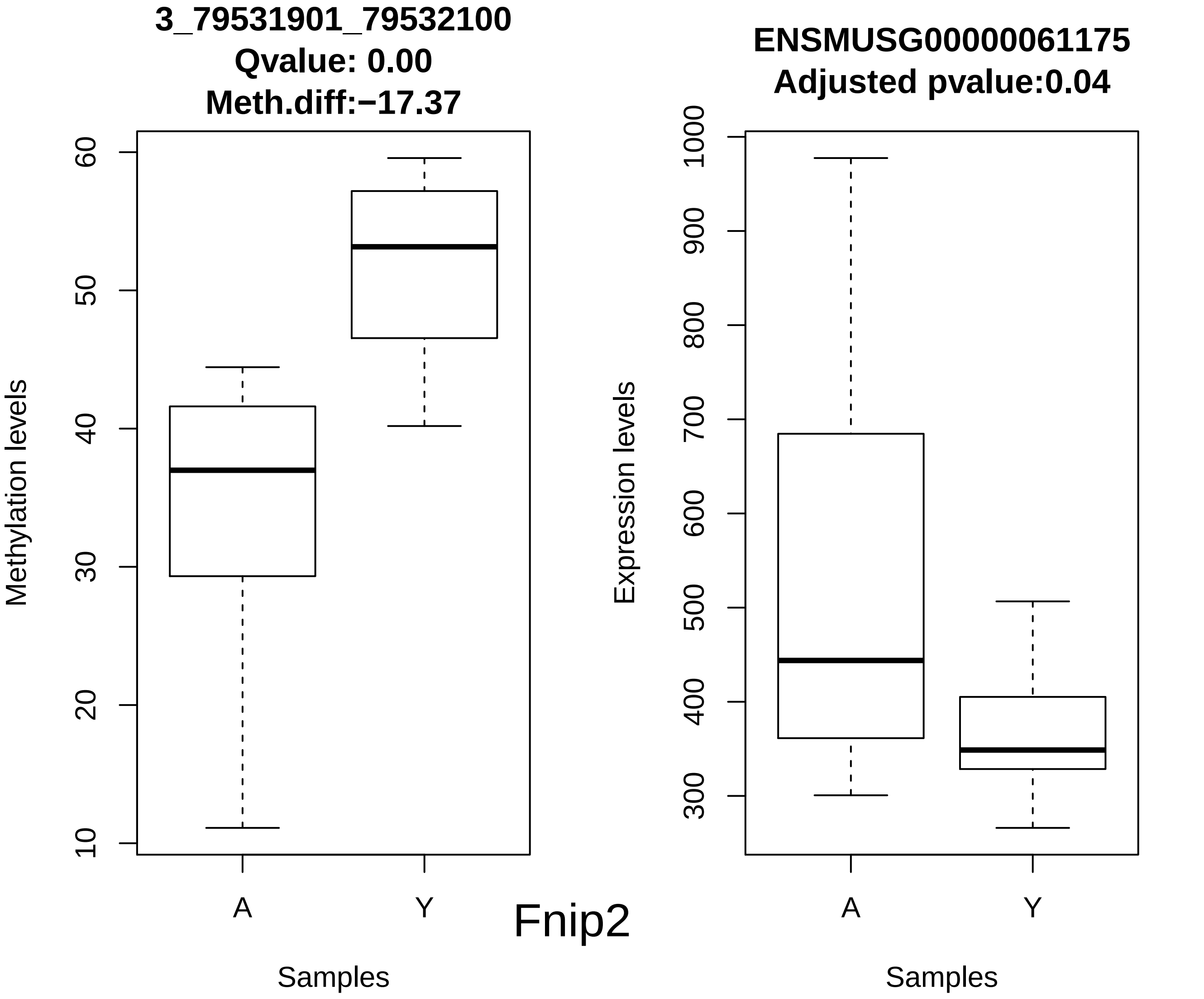

3_79531901_79532100
Qvalue: 0.00
Meth.diff:−17.37
Methylation levels
10
20
30
40
50
60
A
Y
Samples
ENSMUSG00000061175
Adjusted pvalue:0.04
Expression levels
300
400
500
600
700
800
900
1000
A
Y
Samples


Fnip2

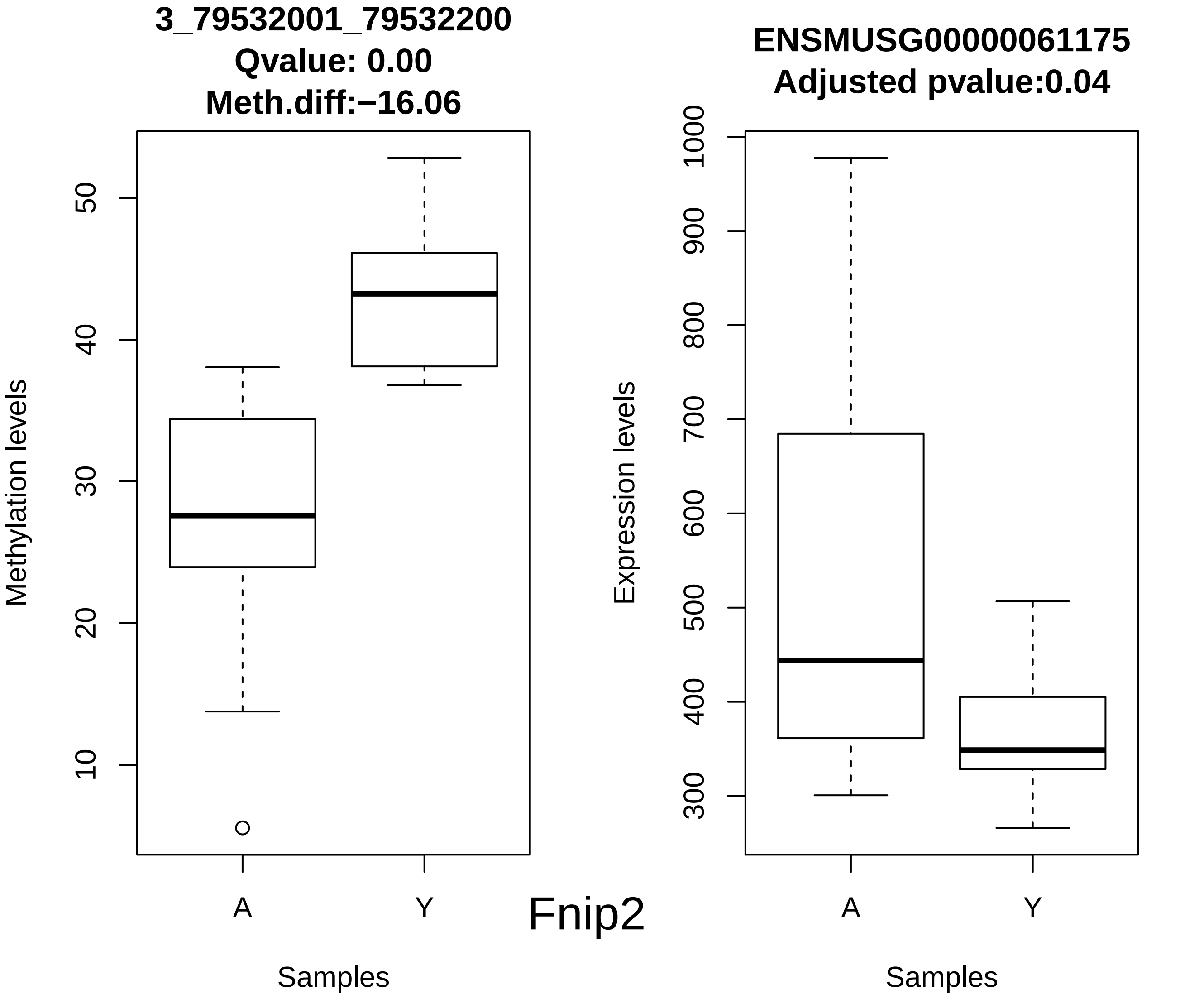
3_79532001_79532200
Qvalue: 0.00
Meth.diff:−16.06
Methylation levels
10
20
30
40
50
A
Y
Samples
Fnip2
ENSMUSG00000061175
Adjusted pvalue:0.04
Expression levels
300
400
500
600
700
800
900
1000
A
Y
Samples

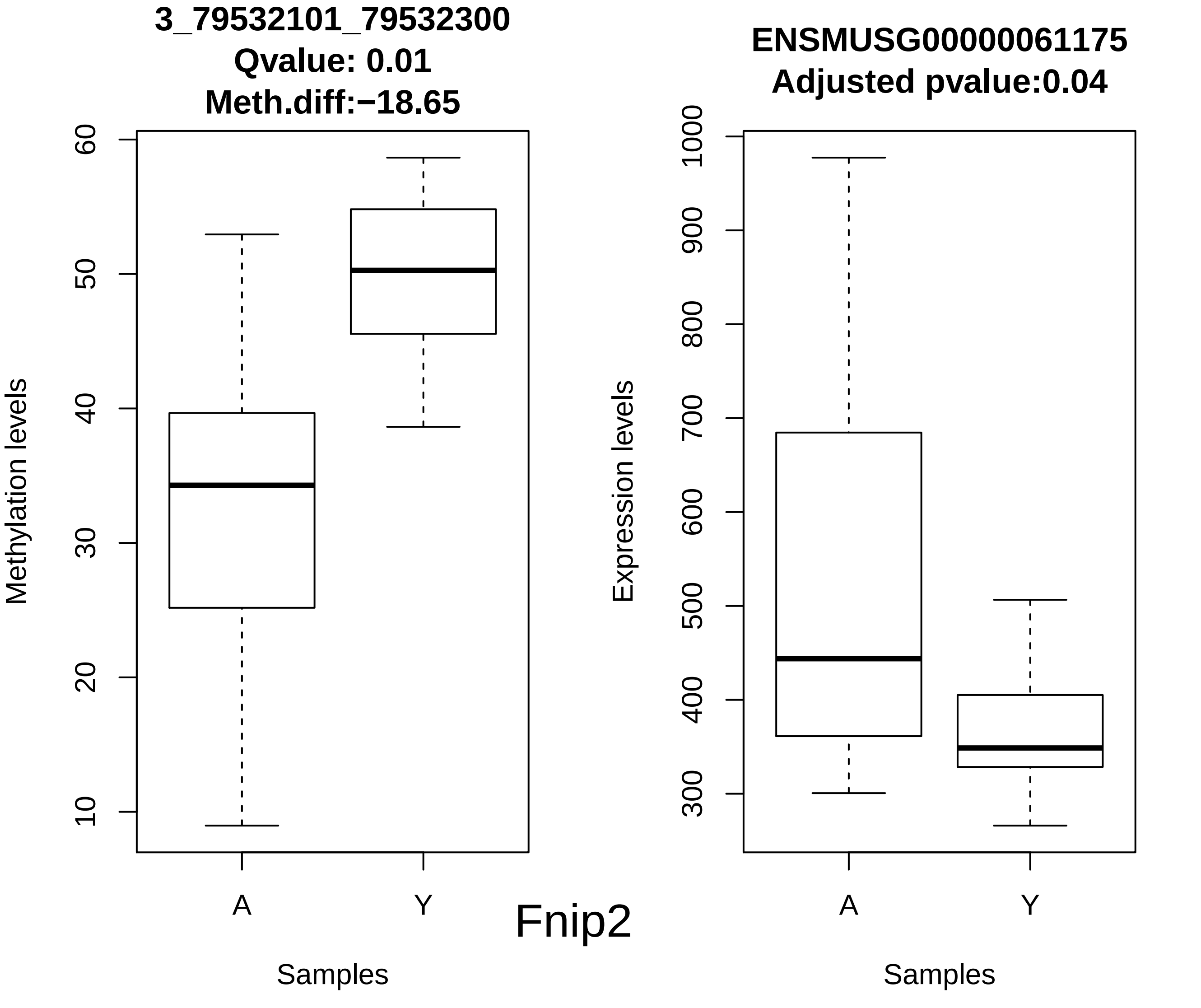
3_79532101_79532300
Qvalue: 0.01
Meth.diff:−18.65
Methylation levels
10
20
30
40
50
60
A
Y
Samples
Fnip2
ENSMUSG00000061175
Adjusted pvalue:0.04
Expression levels
300
400
500
600
700
800
900
1000
A
Y
Samples

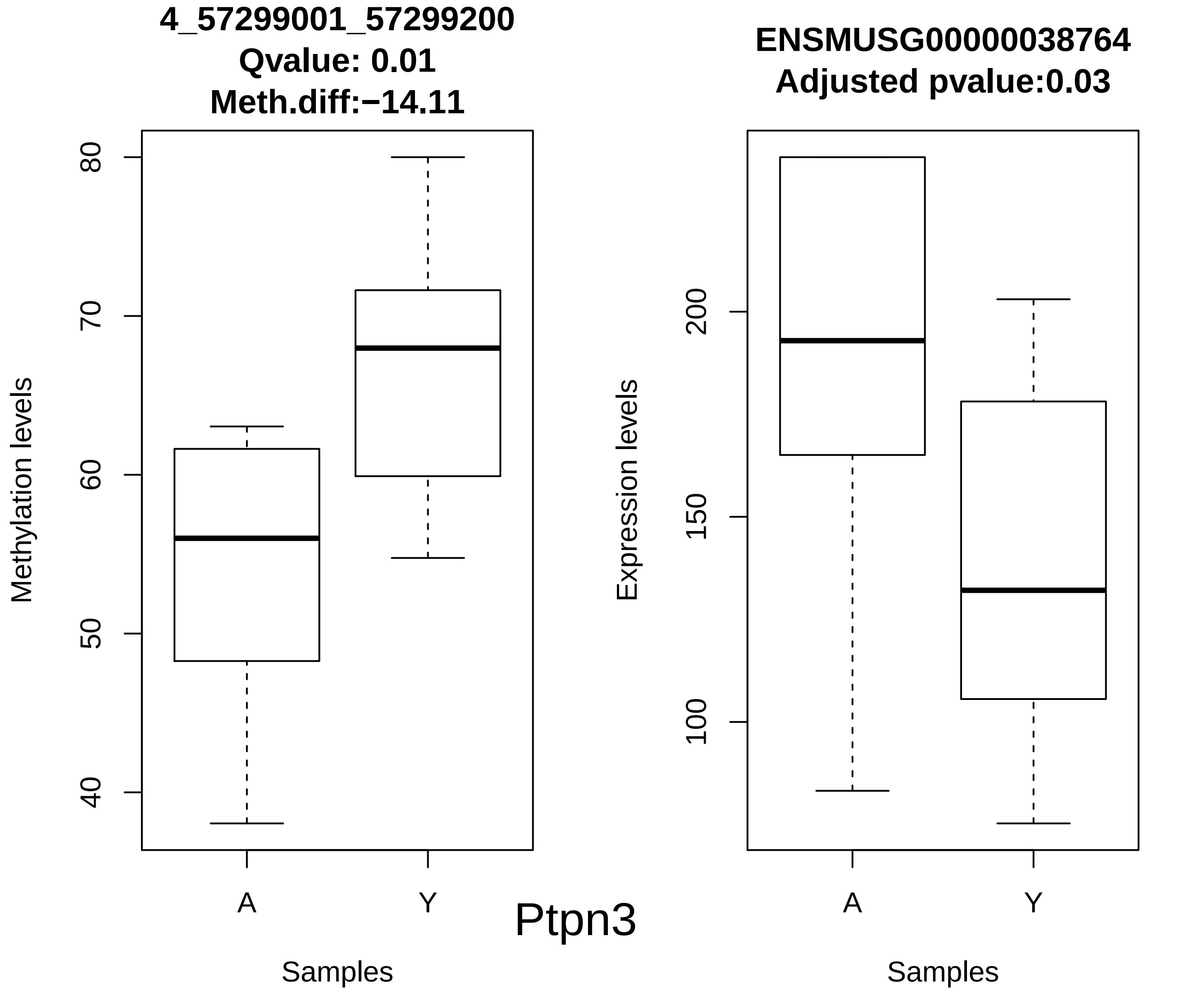
4_57299001_57299200
Qvalue: 0.01
Meth.diff:−14.11
Methylation levels
80
70
60
50
40
A
Y
Samples
Ptpn3
ENSMUSG00000038764
Adjusted pvalue:0.03
Expression levels
200
150
100
A
Y
Samples

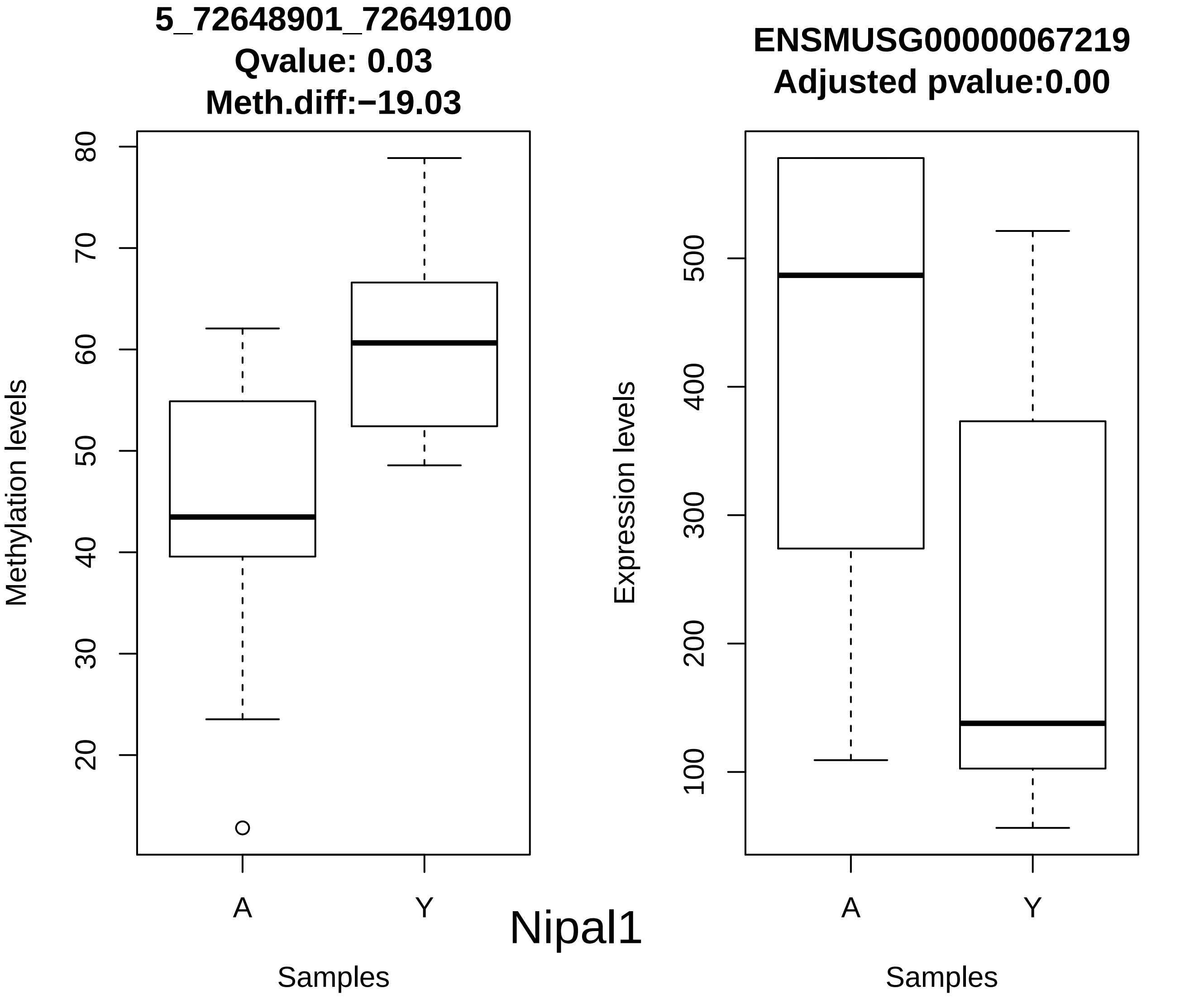
5_72648901_72649100
Qvalue: 0.03
Meth.diff:−19.03
Methylation levels
80
70
60
50
40
30
20
A
Y
Samples
ENSMUSG00000067219
Adjusted pvalue:0.00
Expression levels
500
400
300
200
100
A
Y
Samples
Nipal1

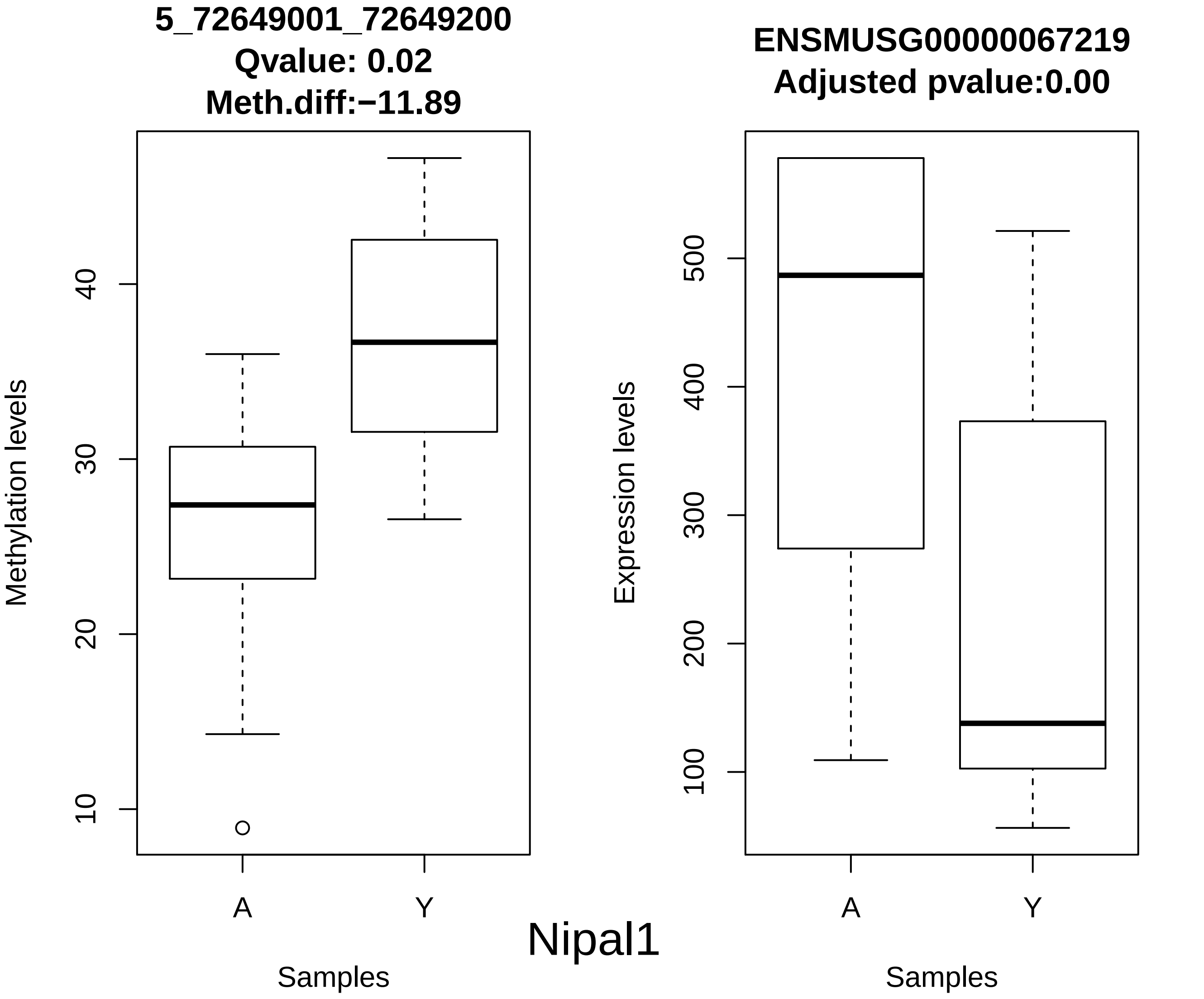
5_72649001_72649200
Qvalue: 0.02
Meth.diff:−11.89
Methylation levels
10
20
30
40
A
Y
Samples
ENSMUSG00000067219
Adjusted pvalue:0.00
Expression levels
100
200
300
400
500
A
Y
Samples
Nipal1

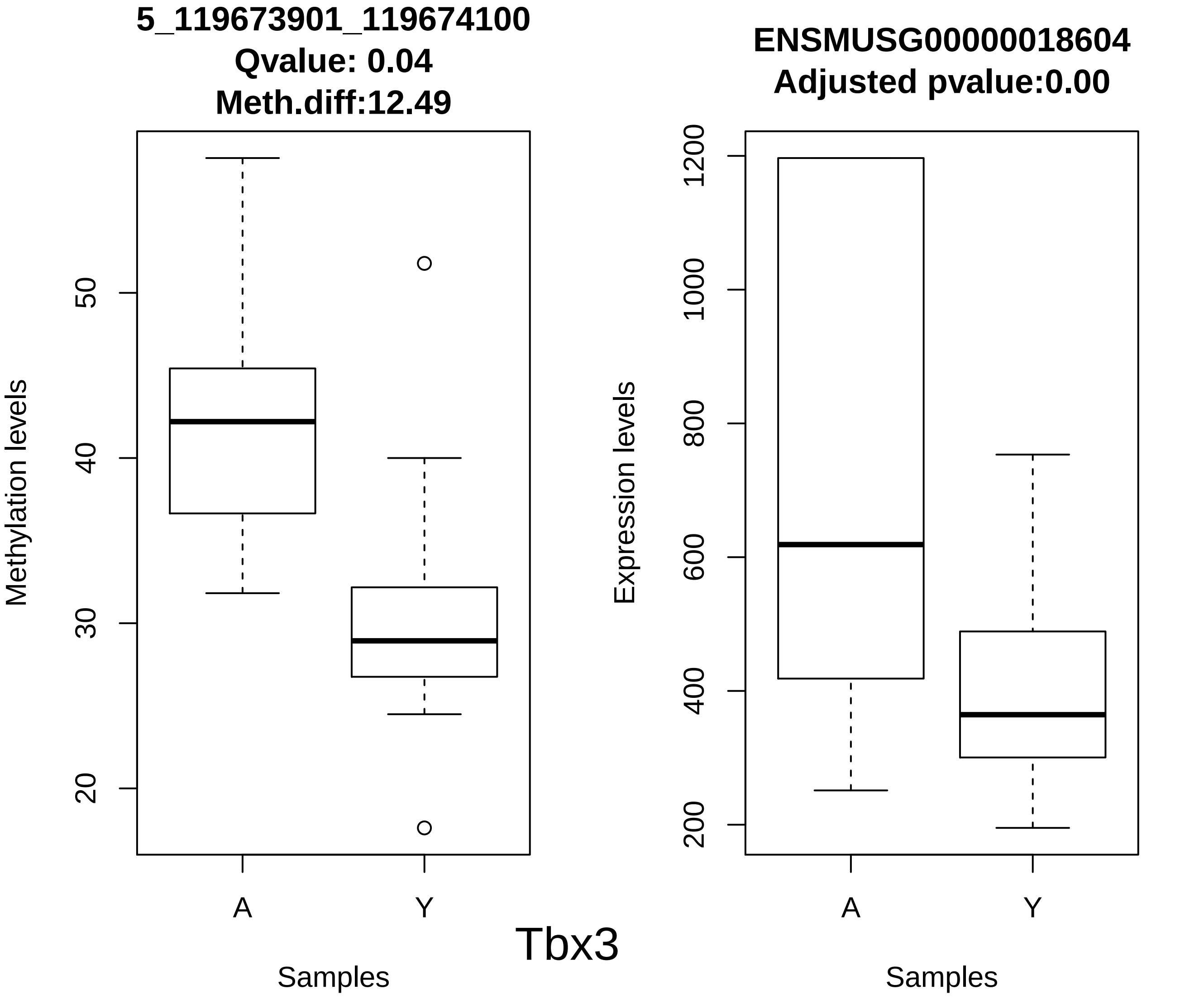

5_119673901_119674100
Qvalue: 0.04
Meth.diff:12.49
Methylation levels
20
30
40
50
A
Y
Samples
ENSMUSG00000018604
Adjusted pvalue:0.00
Expression levels
200
400
600
800
1000
1200
A
Y
Samples
Tbx3

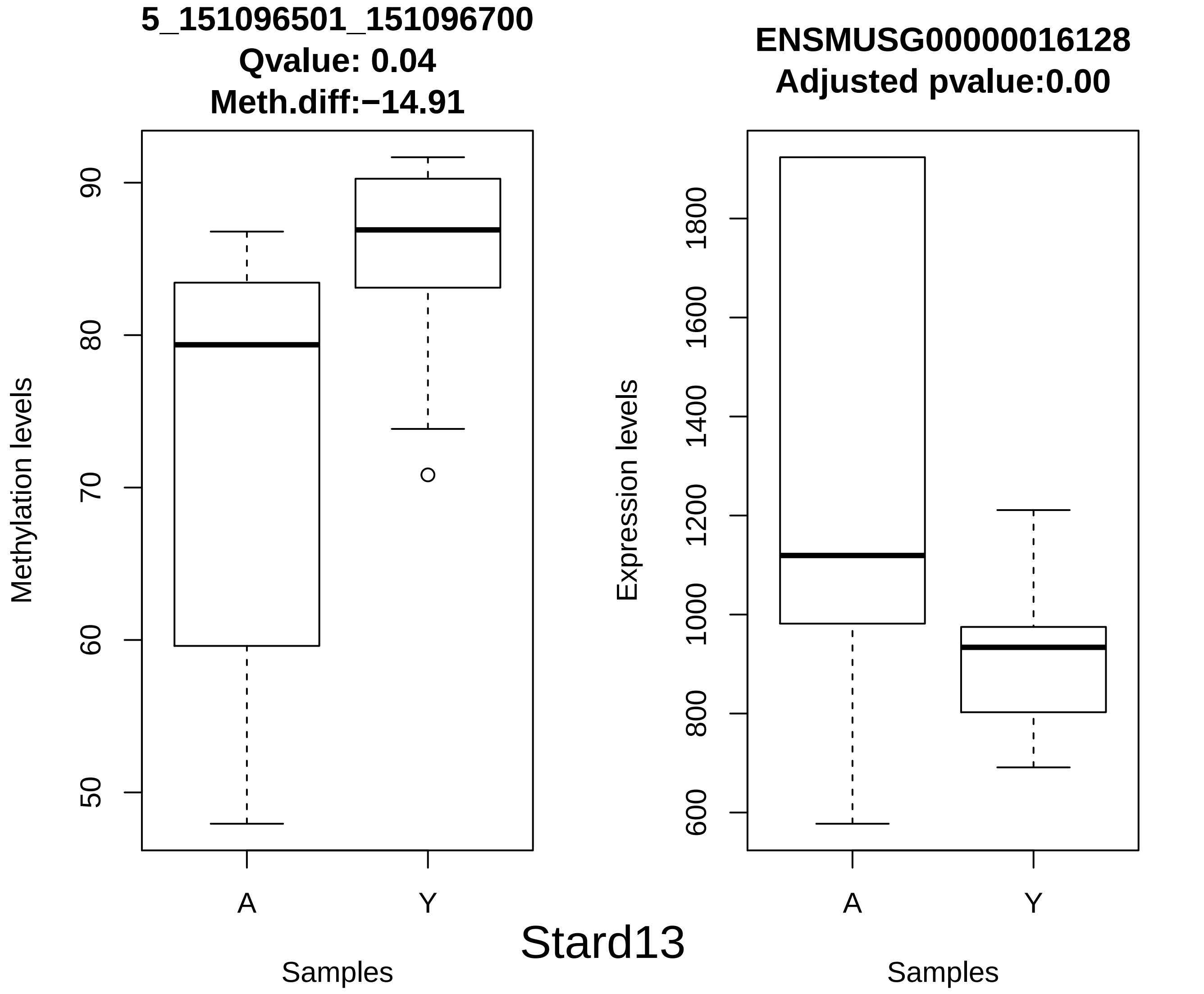
5_151096501_151096700
Qvalue: 0.04
Meth.diff:−14.91
Methylation levels
90
80
70
60
50
A
Y
Samples
ENSMUSG00000016128
Adjusted pvalue:0.00
Expression levels
1800
1600
1400
1200
1000
800
600
A
Y
Samples
Stard13

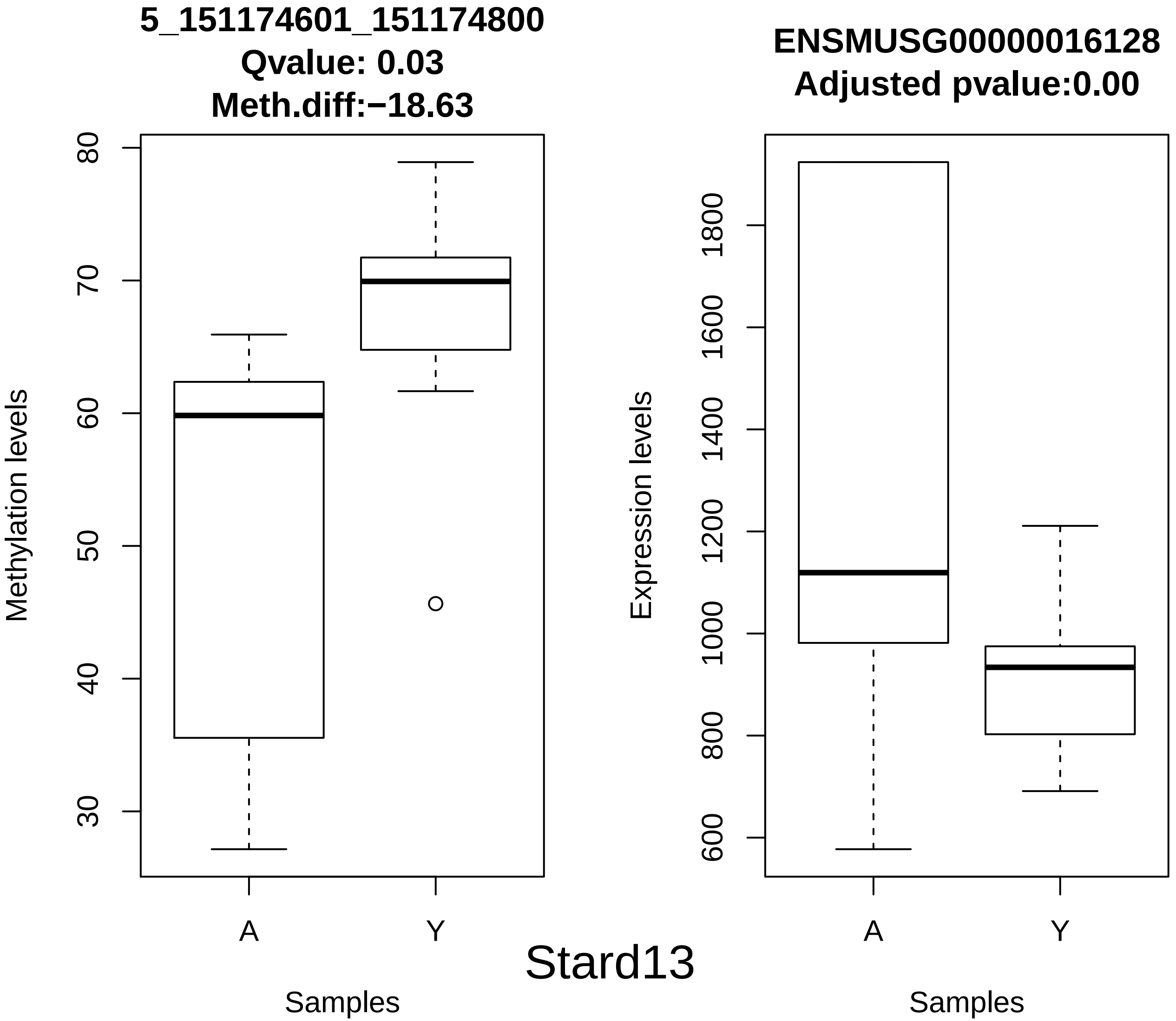
5_151174601_151174800
Qvalue: 0.03
Meth.diff:−18.63
Methylation levels
80
70
60
50
40
30
A
Y
Samples
ENSMUSG00000016128
Adjusted pvalue:0.00
Expression levels
1800
1600
1400
1200
1000
800
600
A
Y
Samples
Stard13

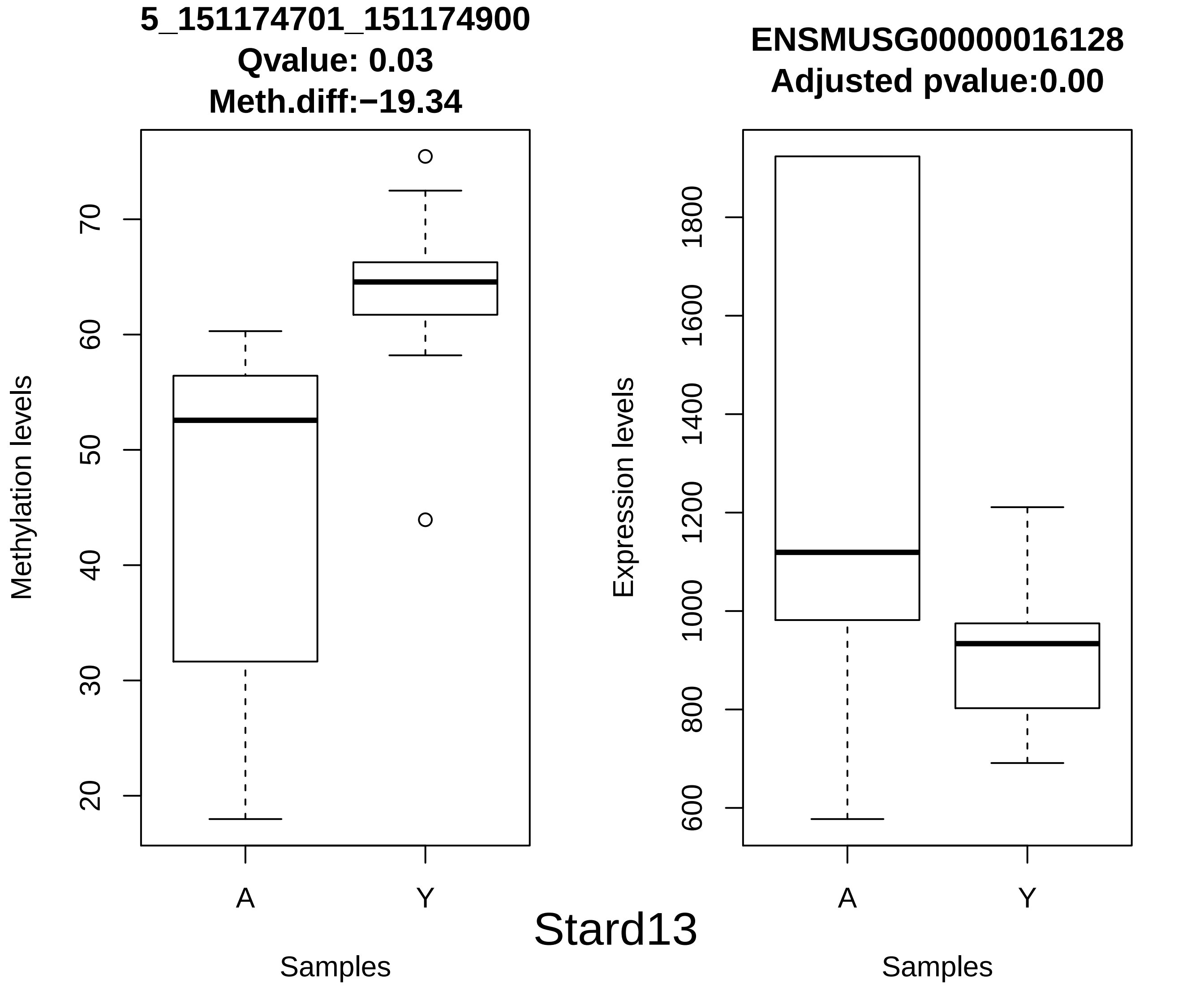
5_151174701_151174900
Qvalue: 0.03
Meth.diff:−19.34
Methylation levels
20
30
40
50
60
70
A
Y
Samples
ENSMUSG00000016128
Adjusted pvalue:0.00
Expression levels
600
800
1000
1200
1400
1600
1800
A
Y
Samples
Stard13

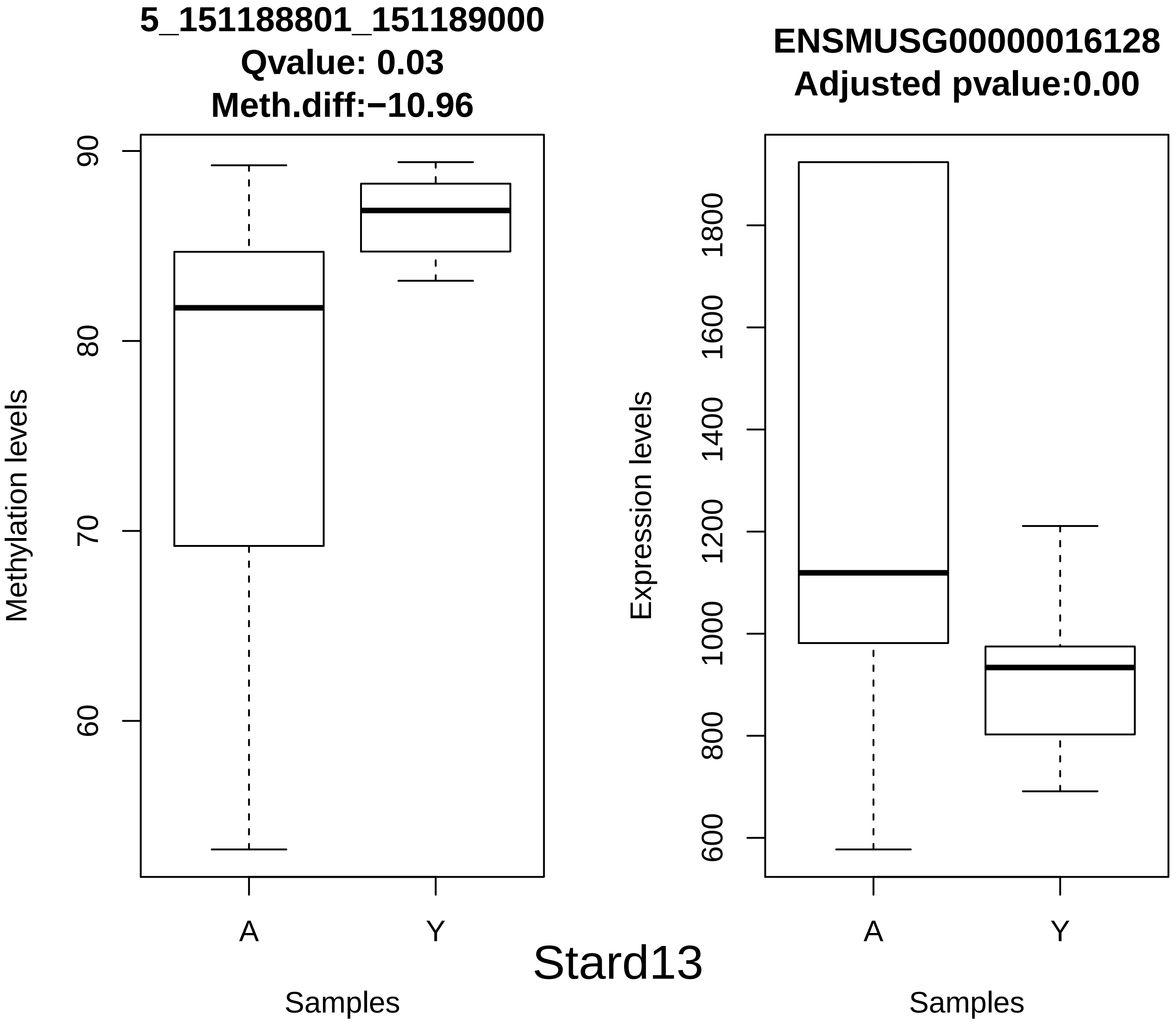

5_151188801_151189000
Qvalue: 0.03
Meth.diff:−10.96
Methylation levels
90
80
70
60
A
Y
Samples
ENSMUSG00000016128
Adjusted pvalue:0.00
Expression levels
1800
1600
1400
1200
1000
800
600
A
Y
Samples
Stard13

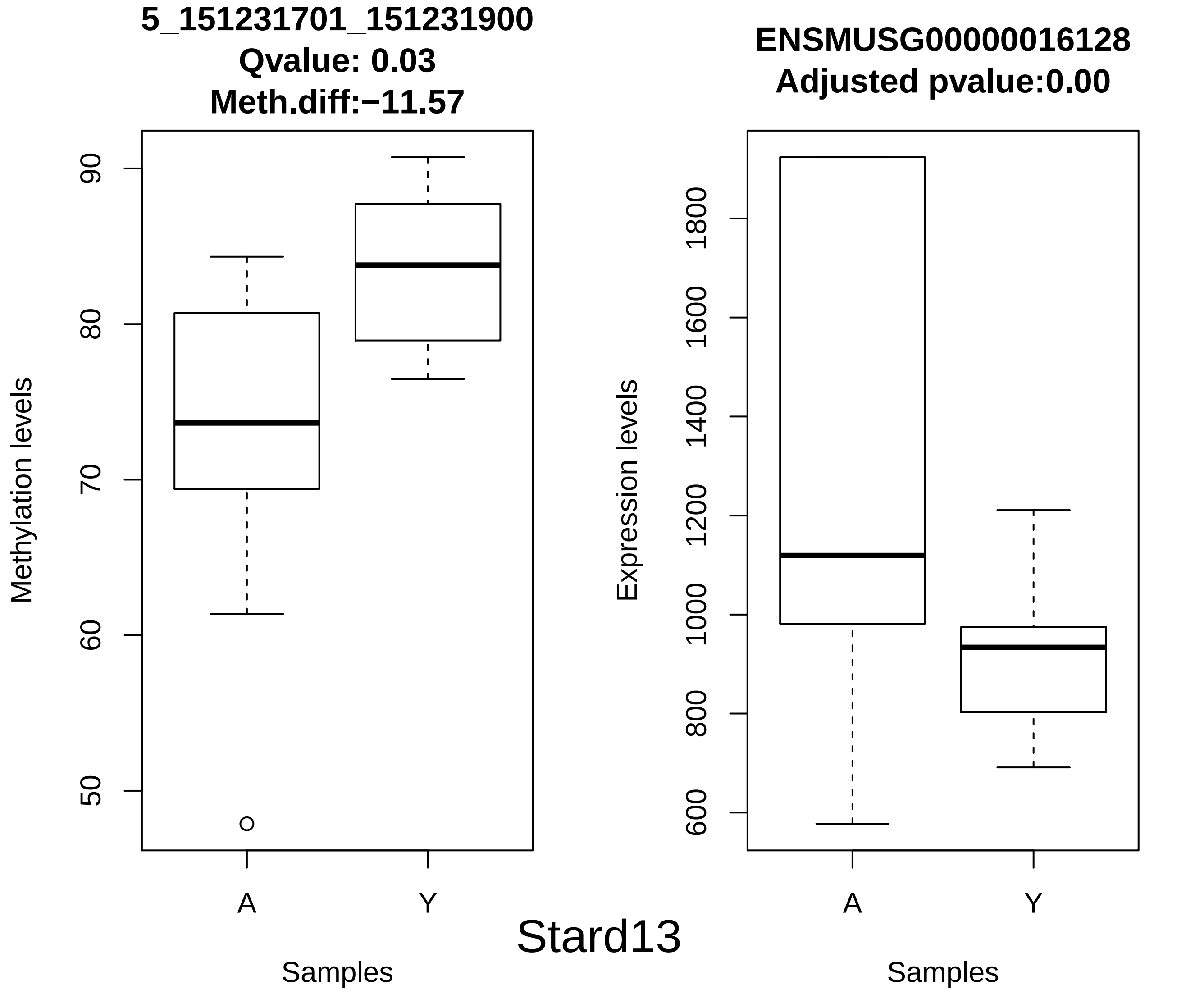
5_151231701_151231900
Qvalue: 0.03
Meth.diff:−11.57
Methylation levels
90
80
70
60
50
A
Y
Samples
ENSMUSG00000016128
Adjusted pvalue:0.00
Expression levels
1800
1600
1400
1200
1000
800
600
A
Y
Samples
Stard13

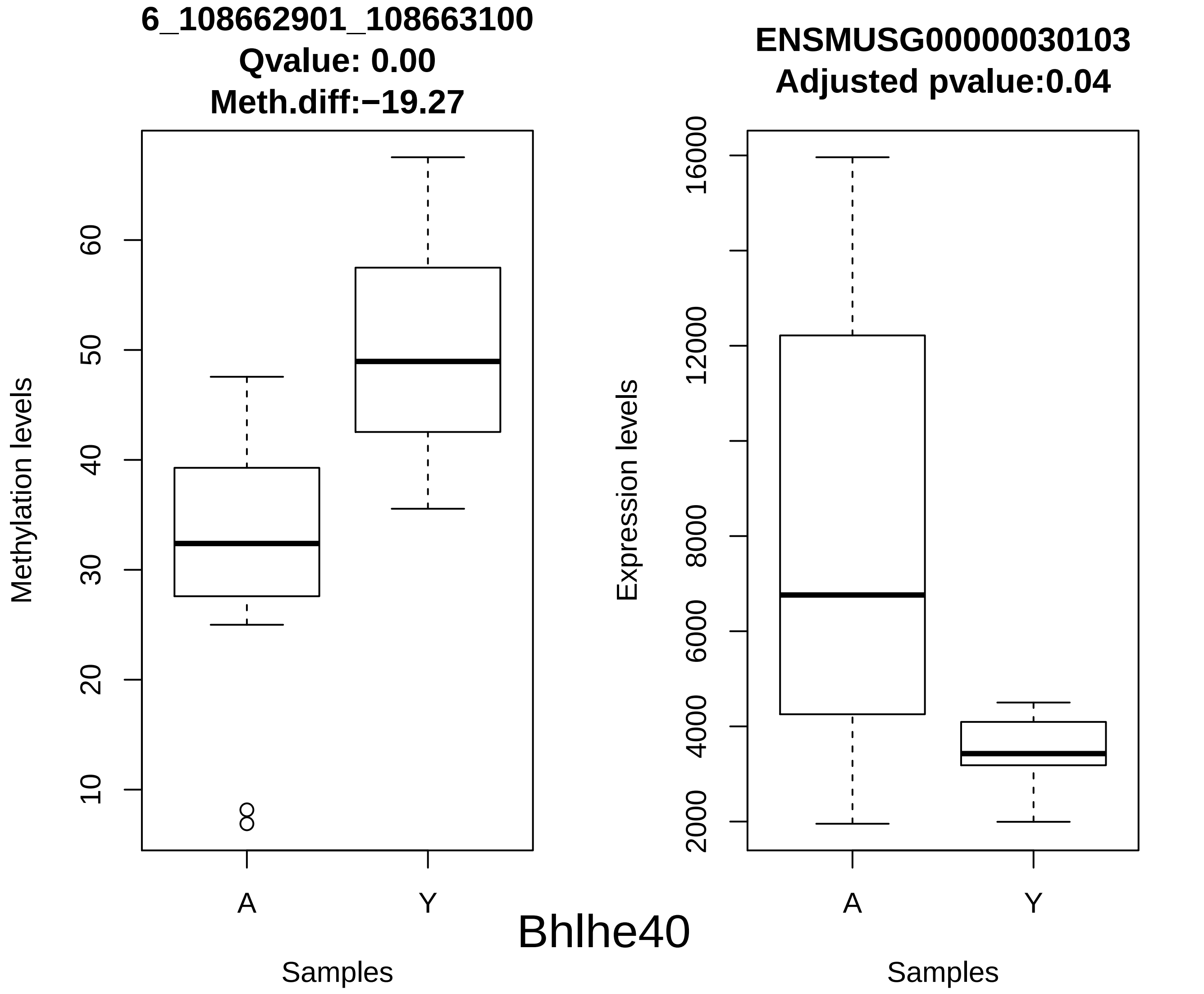

6_108662901_108663100
Qvalue: 0.00
Meth.diff:−19.27
Methylation levels
10
20
30
40
50
60
A
Y
Samples
ENSMUSG00000030103
Adjusted pvalue:0.04
Expression levels
2000
4000
6000
8000
12000
16000
A
Y
Samples
Bhlhe40

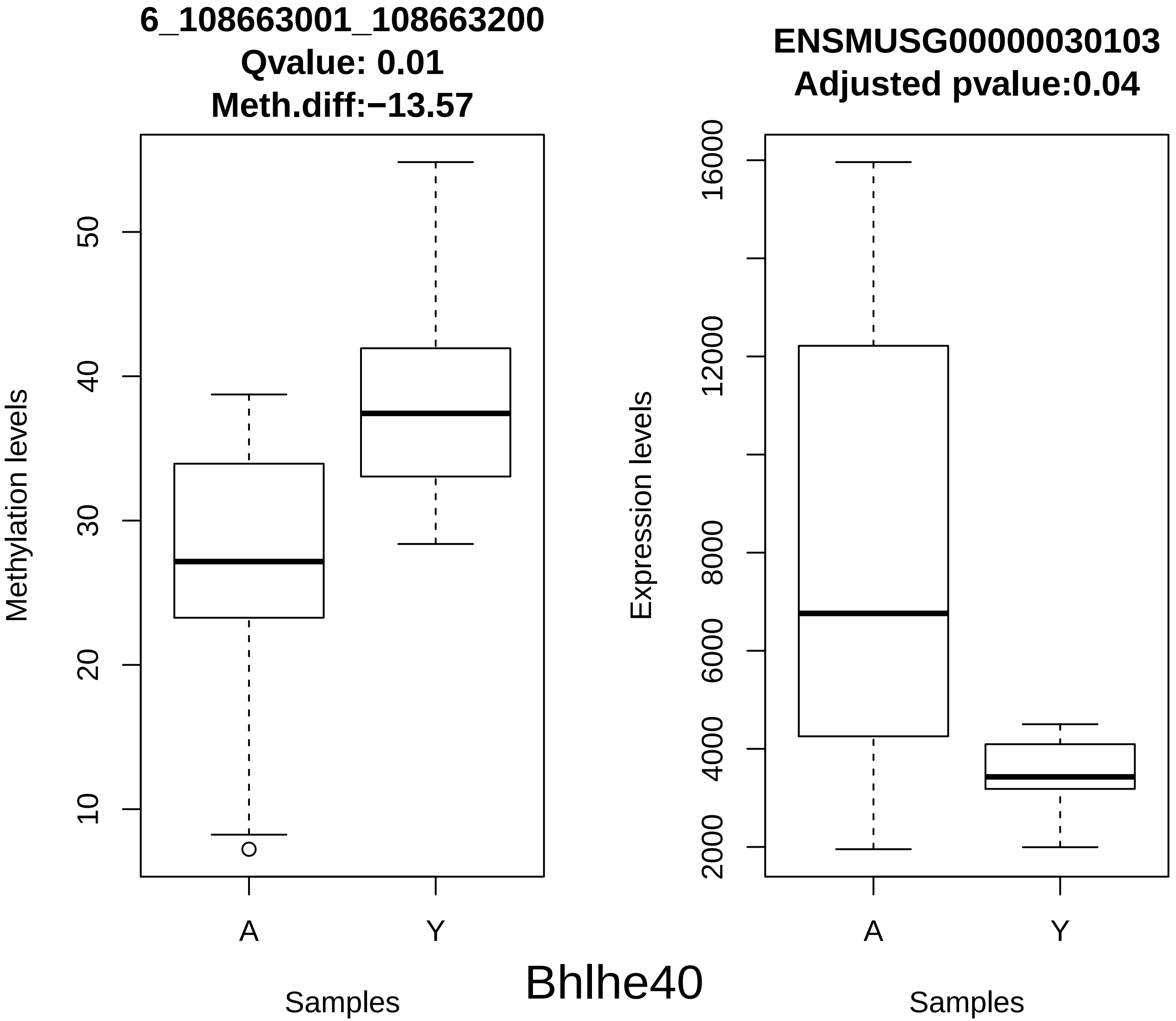
6_108663001_108663200
Qvalue: 0.01
Meth.diff:−13.57
Methylation levels
10
20
30
40
50
A
Y
Samples
ENSMUSG00000030103
Adjusted pvalue:0.04
Expression levels
2000
4000
6000
8000
12000
16000
A
Y
Samples
Bhlhe40

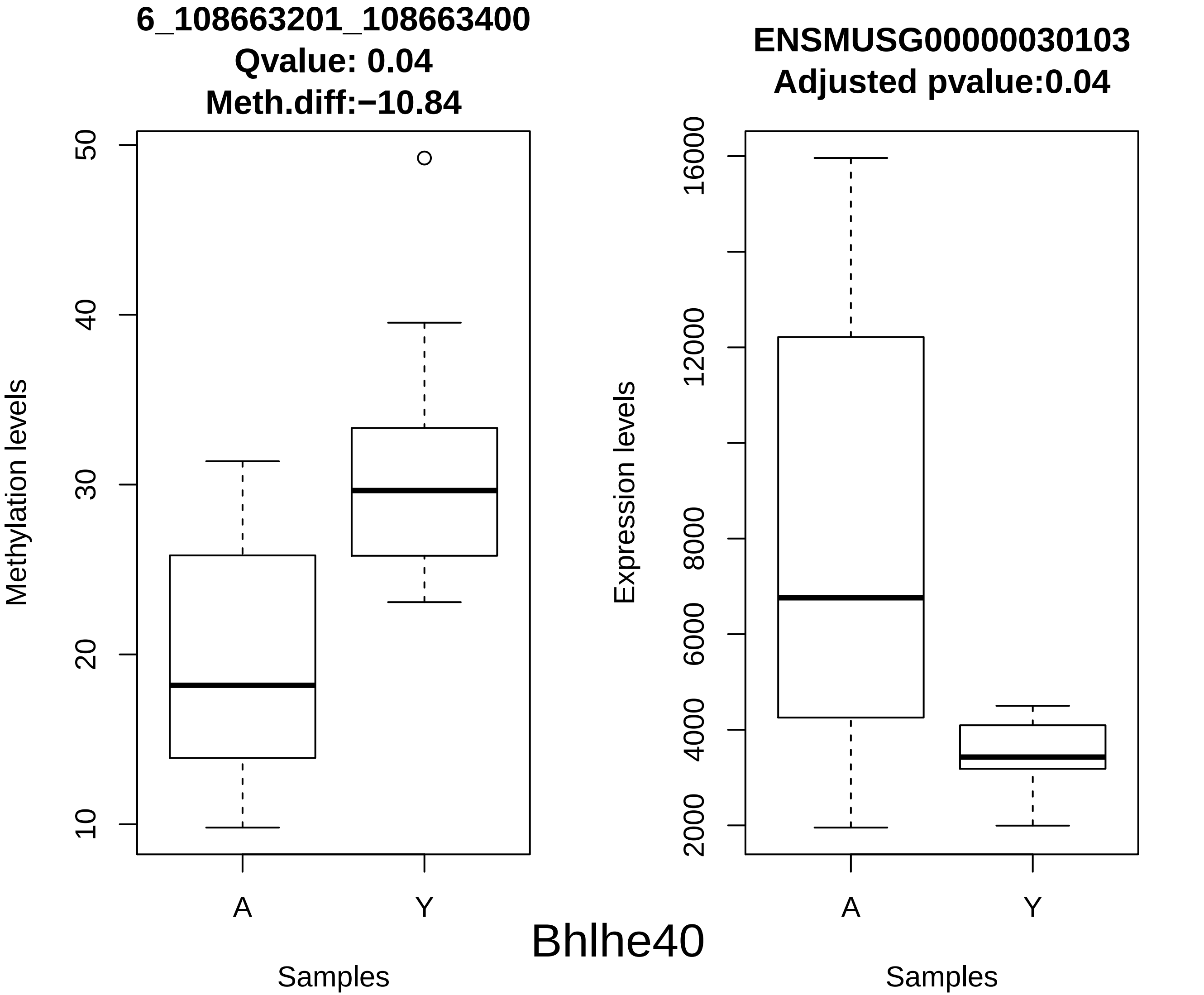
6_108663201_108663400
Qvalue: 0.04
Meth.diff:−10.84
Methylation levels
10
20
30
40
50
A
Y
Samples
ENSMUSG00000030103
Adjusted pvalue:0.04
Expression levels
2000
4000
6000
8000
12000
16000
A
Y
Samples
Bhlhe40

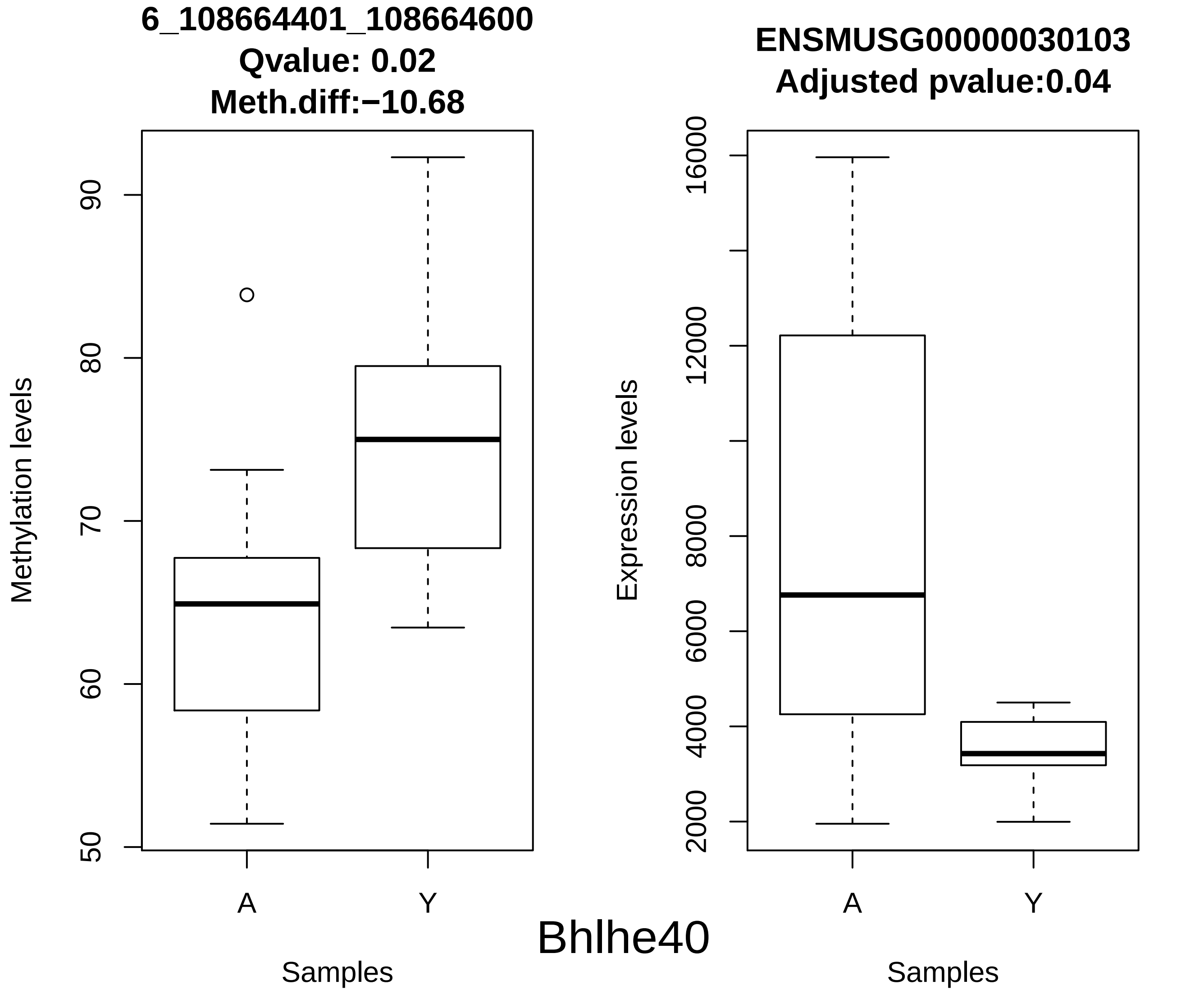
6_108664401_108664600
Qvalue: 0.02
Meth.diff:−10.68
Methylation levels
90
80
70
60
50
A
Y
Samples
ENSMUSG00000030103
Adjusted pvalue:0.04
Expression levels
16000
12000
8000
6000
4000
2000
A
Y
Samples
Bhlhe40

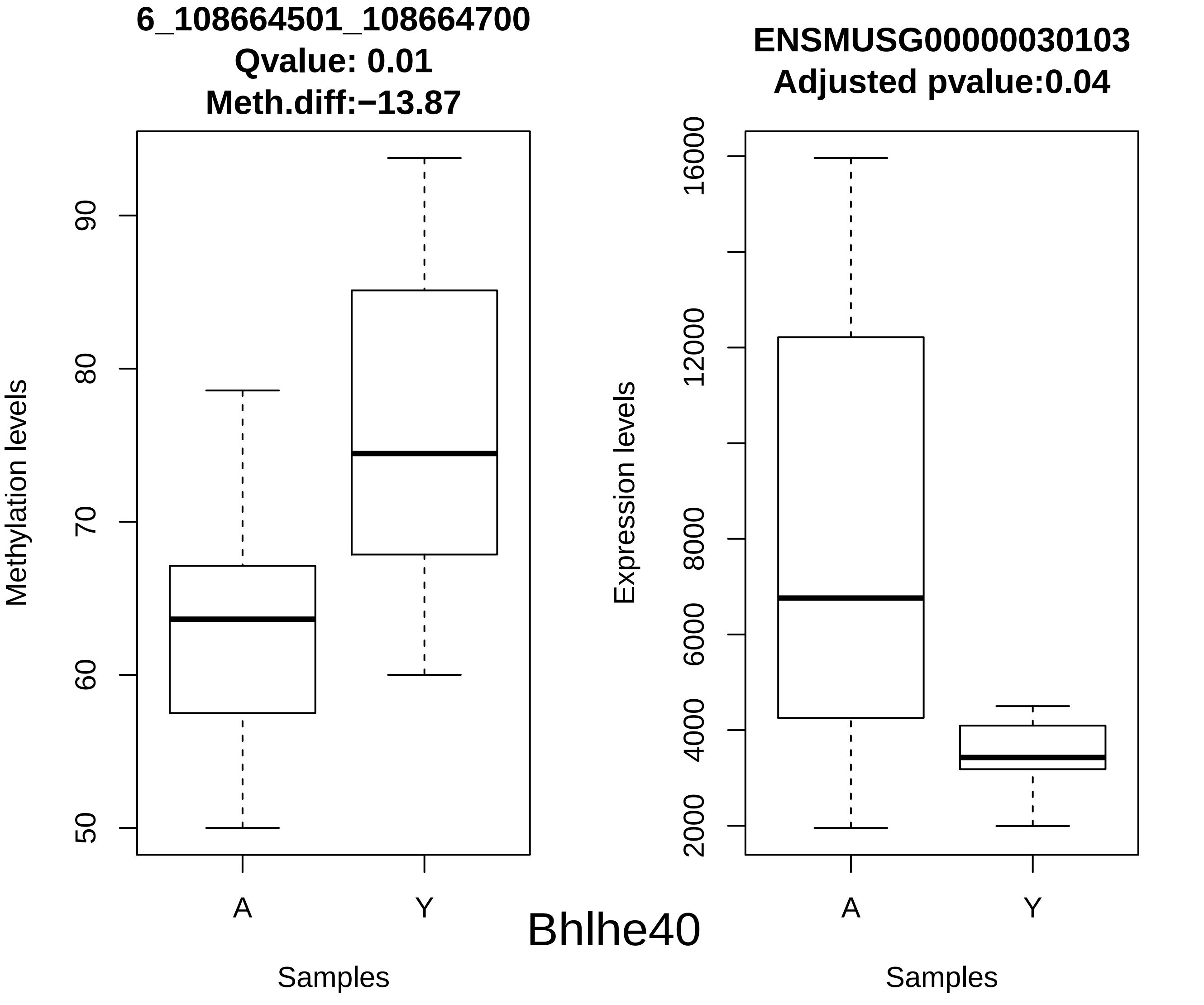
6_108664501_108664700
Qvalue: 0.01
Meth.diff:−13.87
Methylation levels
50
60
70
80
90
A
Y
Samples
ENSMUSG00000030103
Adjusted pvalue:0.04
Expression levels
2000
4000
6000
8000
12000
16000
A
Y
Samples
Bhlhe40

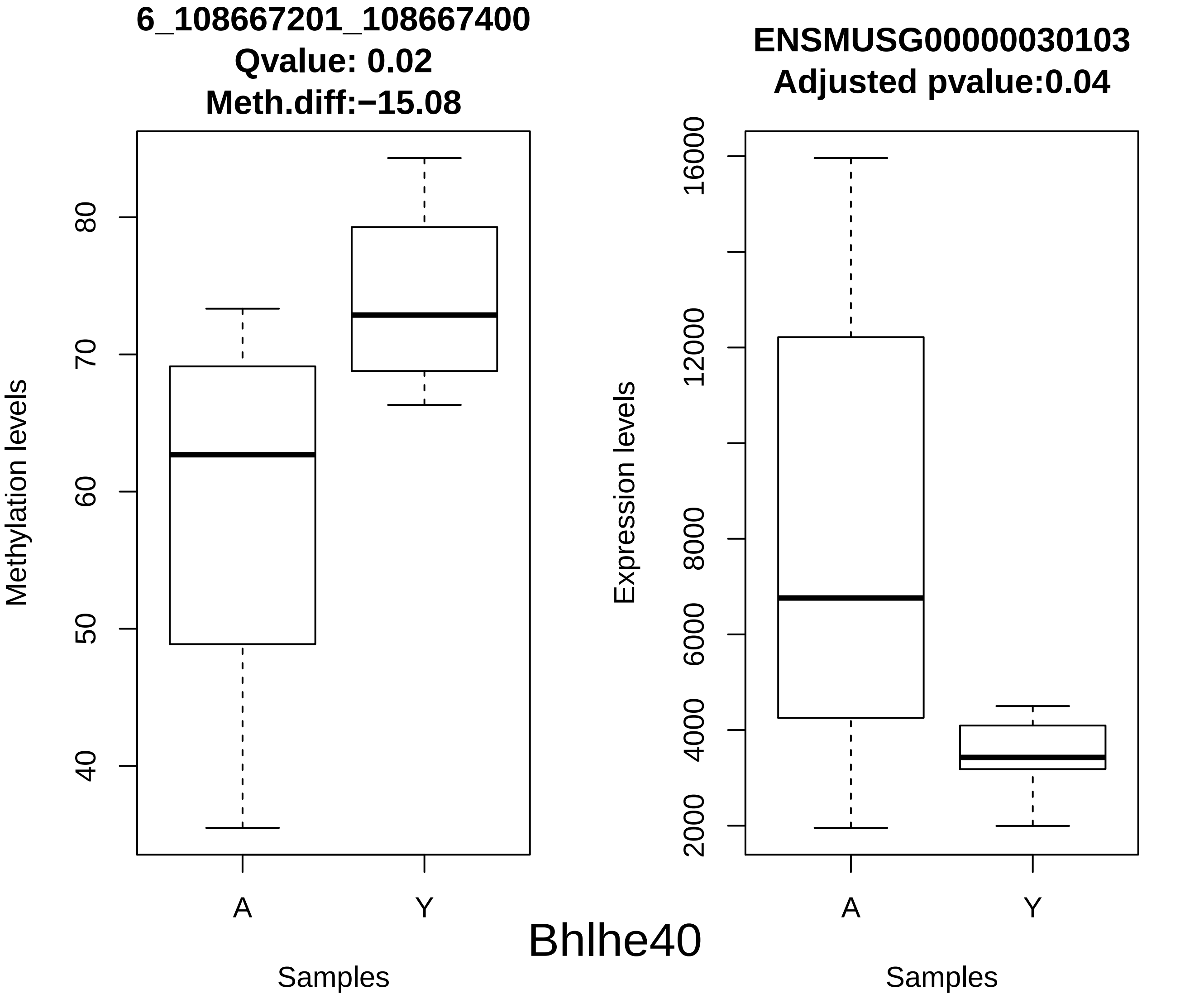
6_108667201_108667400
Qvalue: 0.02
Meth.diff:−15.08
Methylation levels
40
50
60
70
80
A
Y
Samples
ENSMUSG00000030103
Adjusted pvalue:0.04
Expression levels
2000
4000
6000
8000
12000
16000
A
Y
Samples
Bhlhe40

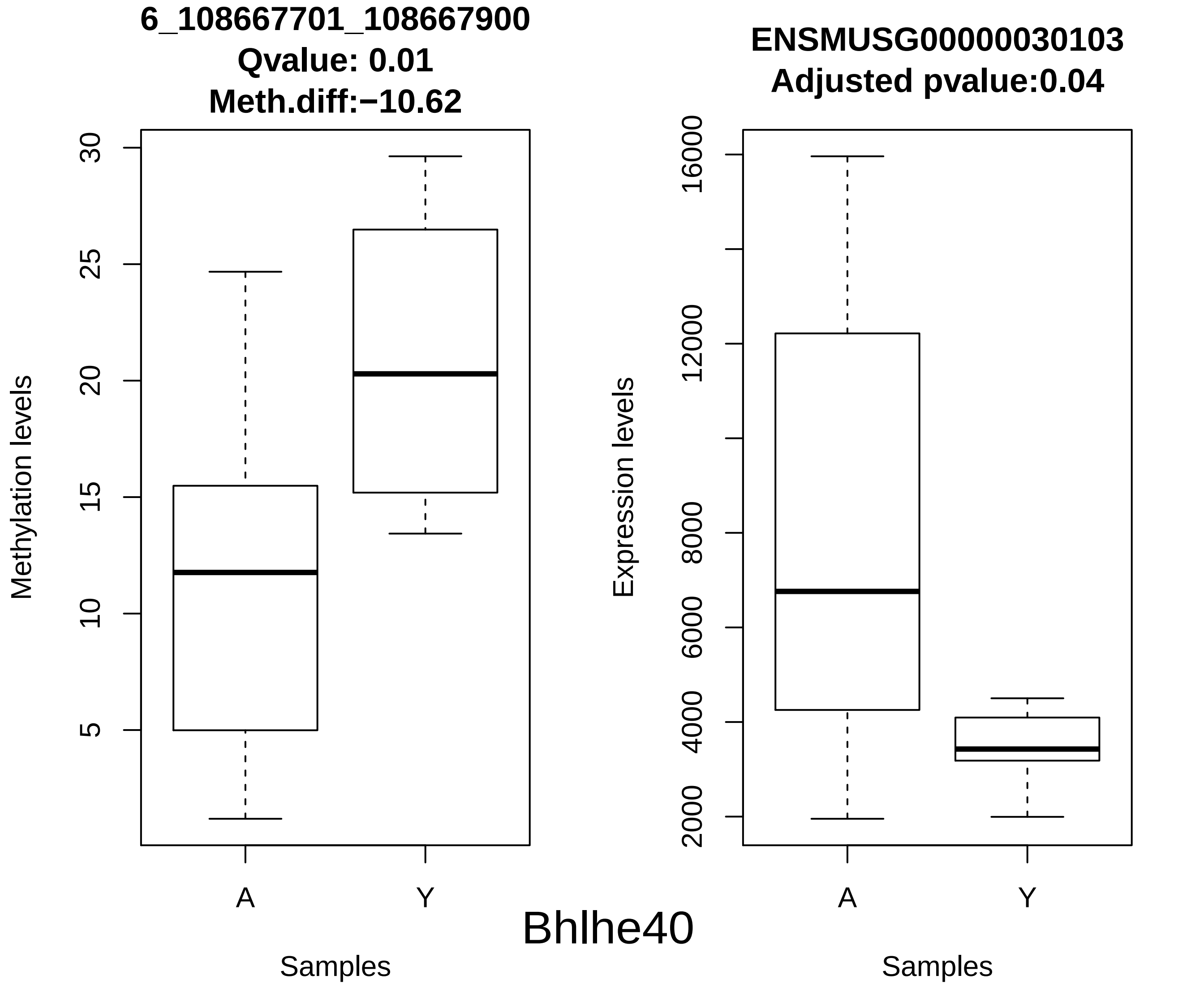
6_108667701_108667900
Qvalue: 0.01
Meth.diff:−10.62
Methylation levels
30
25
20
15
10
5
A
Y
Samples
ENSMUSG00000030103
Adjusted pvalue:0.04
Expression levels
16000
12000
8000
6000
4000
2000
A
Y
Samples
Bhlhe40

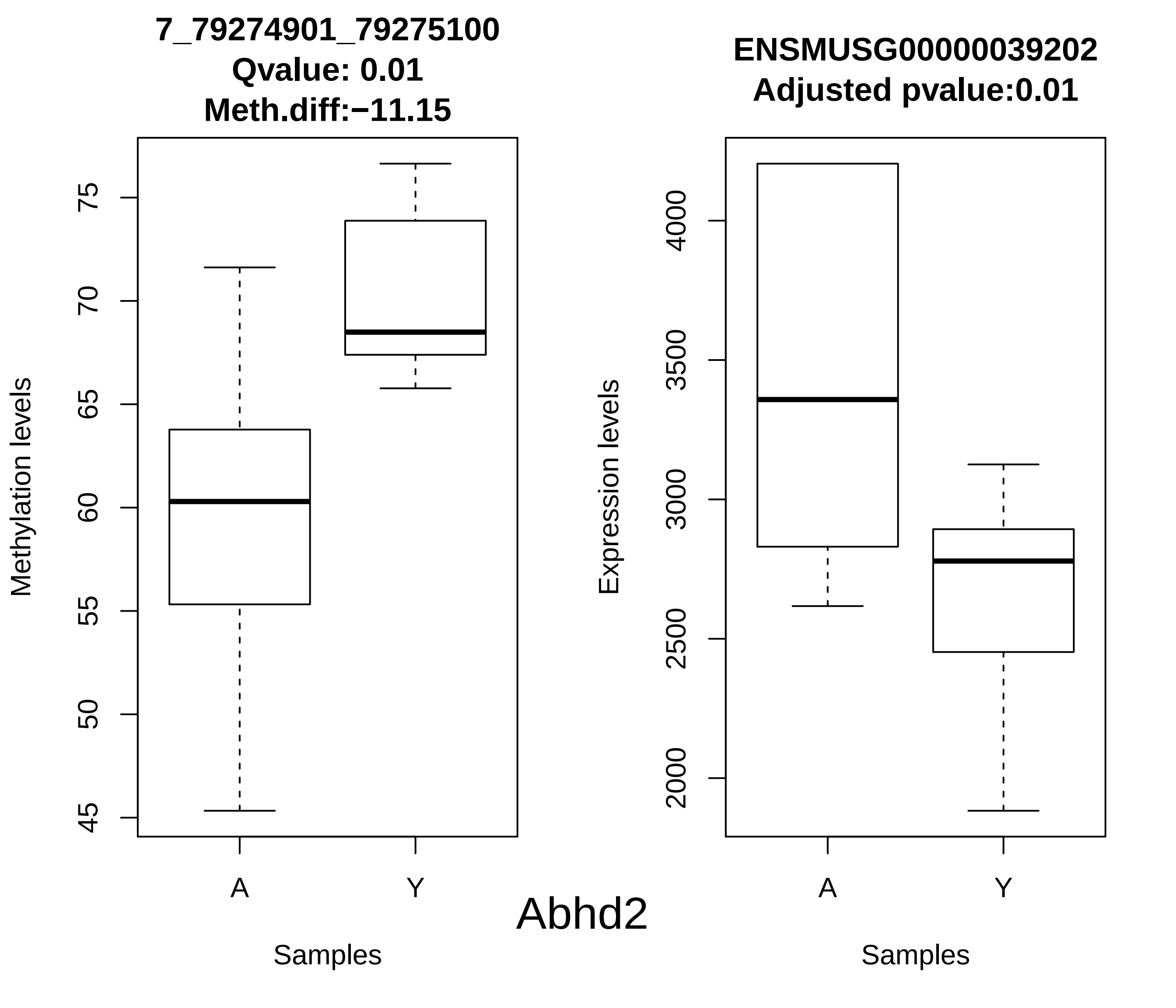
7_79274901_79275100
Qvalue: 0.01
Meth.diff:−11.15
Methylation levels
45
50
55
60
65
70
75
A
Y
Samples
ENSMUSG00000039202
Adjusted pvalue:0.01
Expression levels
2000
2500
3000
3500
4000
A
Y
Samples
Abhd2

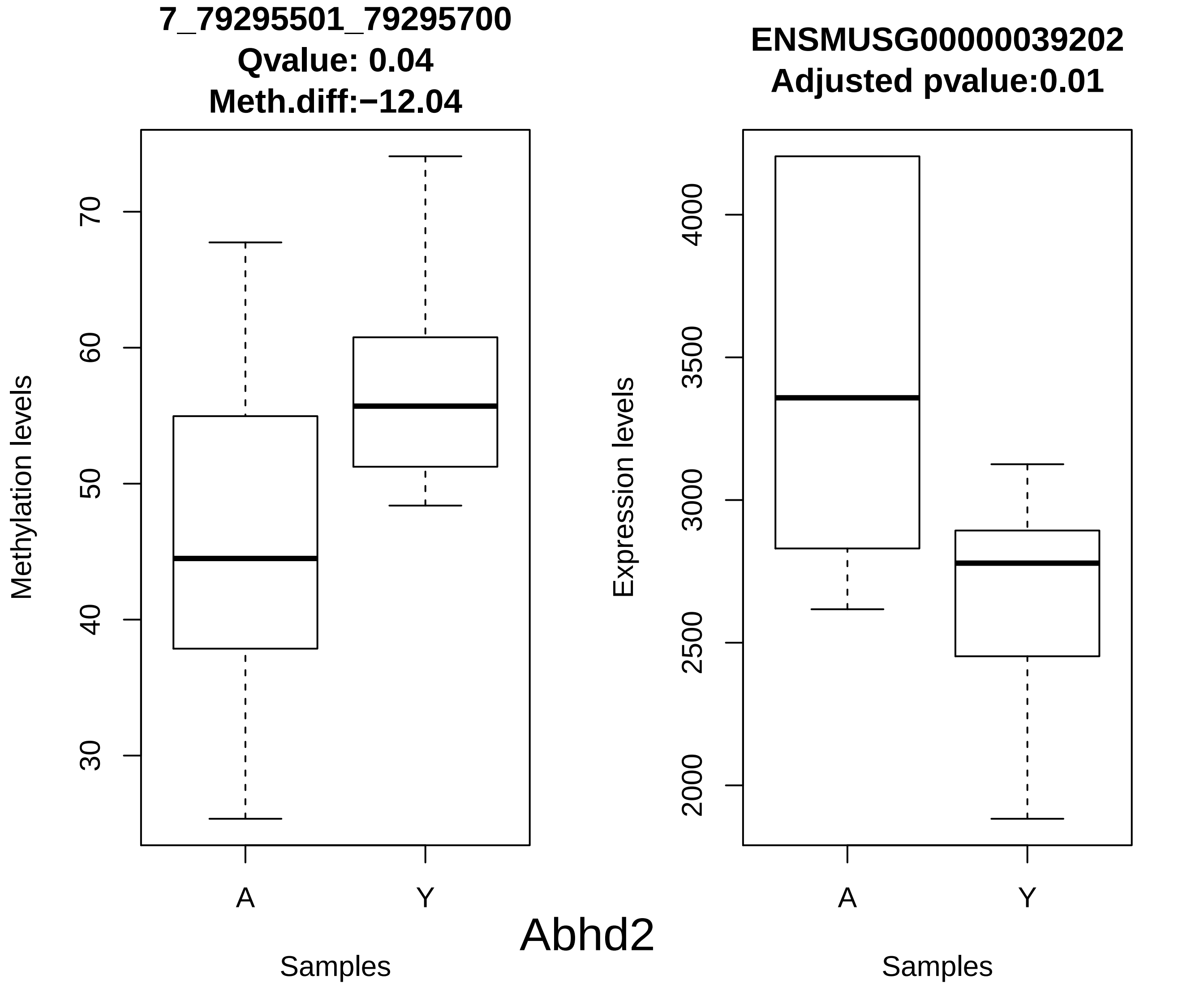
7_79295501_79295700
Qvalue: 0.04
Meth.diff:−12.04
Methylation levels
70
60
50
40
30
A
Y
Samples
ENSMUSG00000039202
Adjusted pvalue:0.01
Expression levels
4000
3500
3000
2500
2000
A
Y
Samples
Abhd2

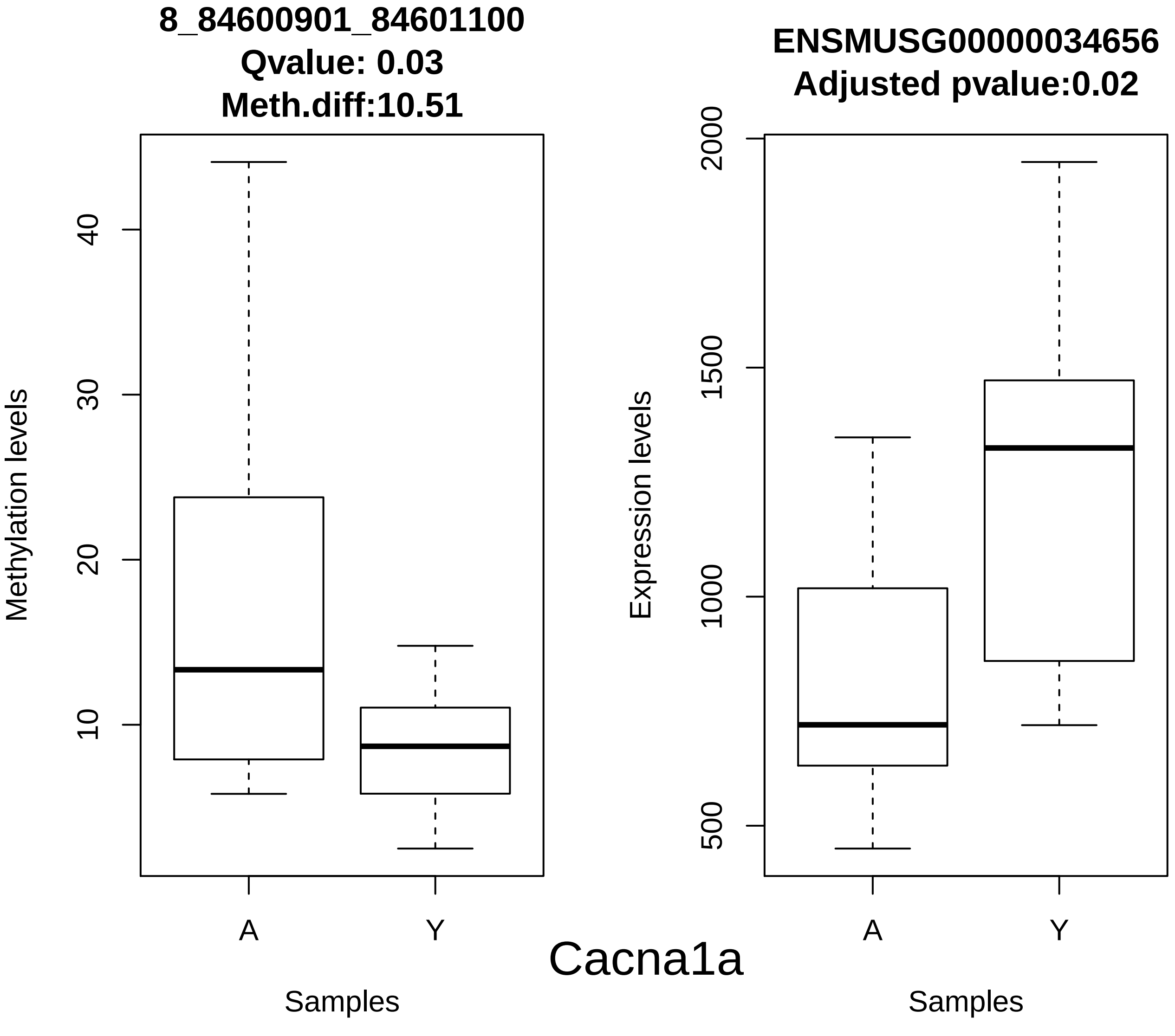
8_84600901_84601100
Qvalue: 0.03
Meth.diff:10.51
Methylation levels
10
20
30
40
A
Y
Samples
ENSMUSG00000034656
Adjusted pvalue:0.02
Expression levels
500
1000
1500
2000
A
Y
Samples
Cacna1a

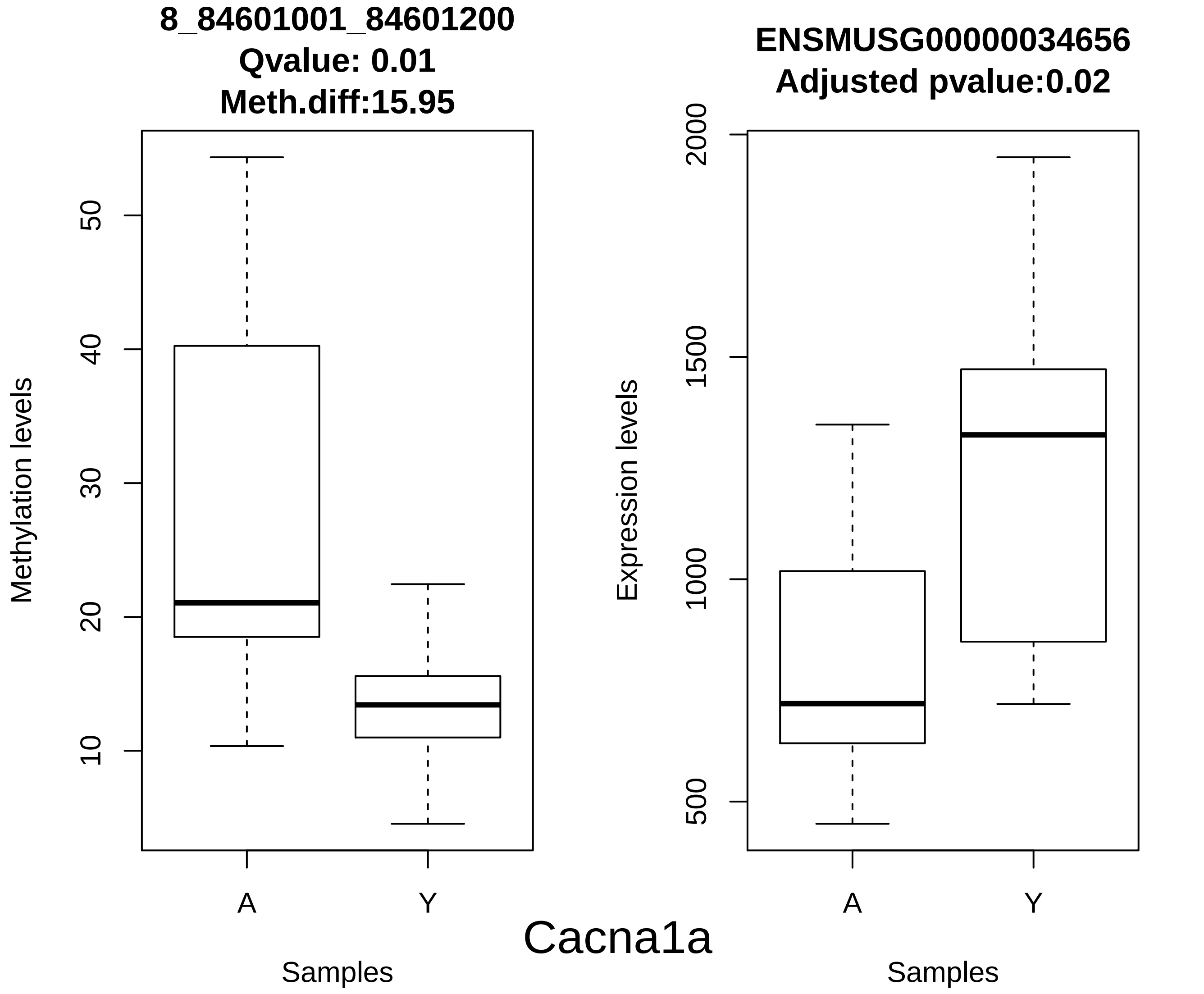
8_84601001_84601200
Qvalue: 0.01
Meth.diff:15.95
Methylation levels
10
20
30
40
50
A
Y
Samples
ENSMUSG00000034656
Adjusted pvalue:0.02
Expression levels
500
1000
1500
2000
A
Y
Samples
Cacna1a

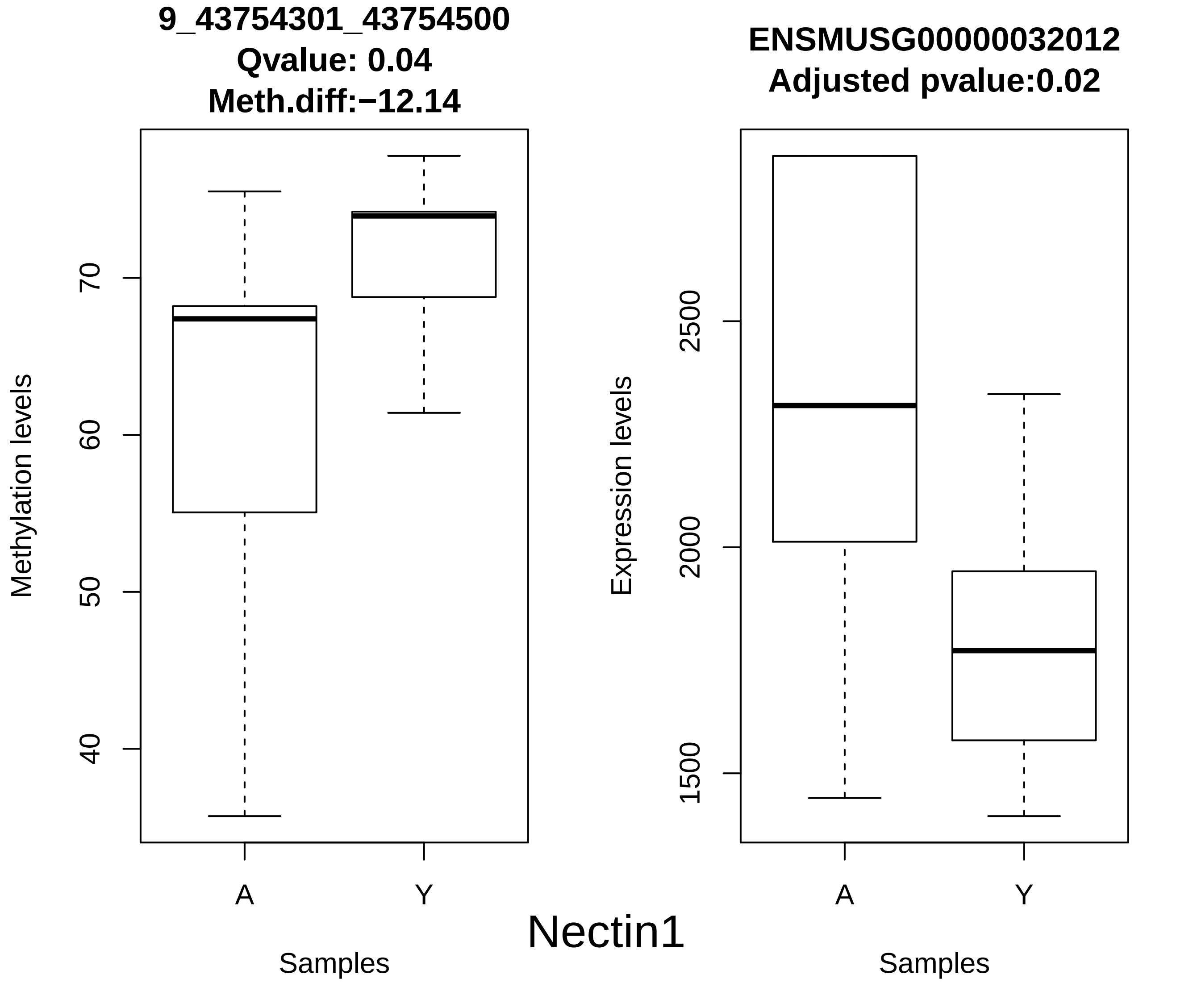
9_43754301_43754500
Qvalue: 0.04
Meth.diff:−12.14
Methylation levels
70
60
50
40
A
Y
Samples
ENSMUSG00000032012
Adjusted pvalue:0.02
Expression levels
2500
2000
1500
A
Y
Samples
Nectin1

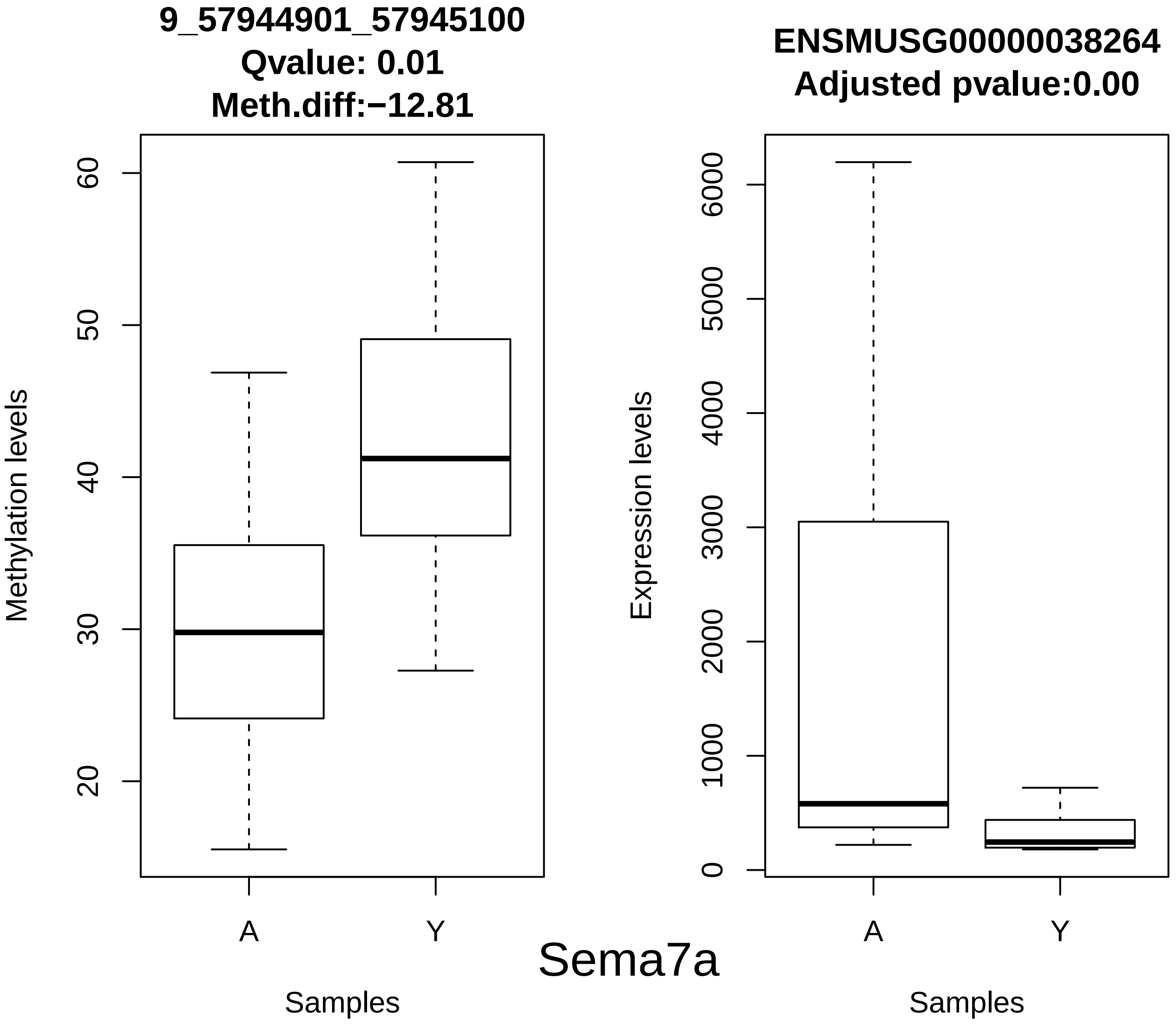
9_57944901_57945100
Qvalue: 0.01
Meth.diff:−12.81
Methylation levels
20
30
40
50
60
A
Y
Samples
ENSMUSG00000038264
Adjusted pvalue:0.00
Expression levels
0
1000
2000
3000
4000
5000
6000
A
Y
Samples
Sema7a

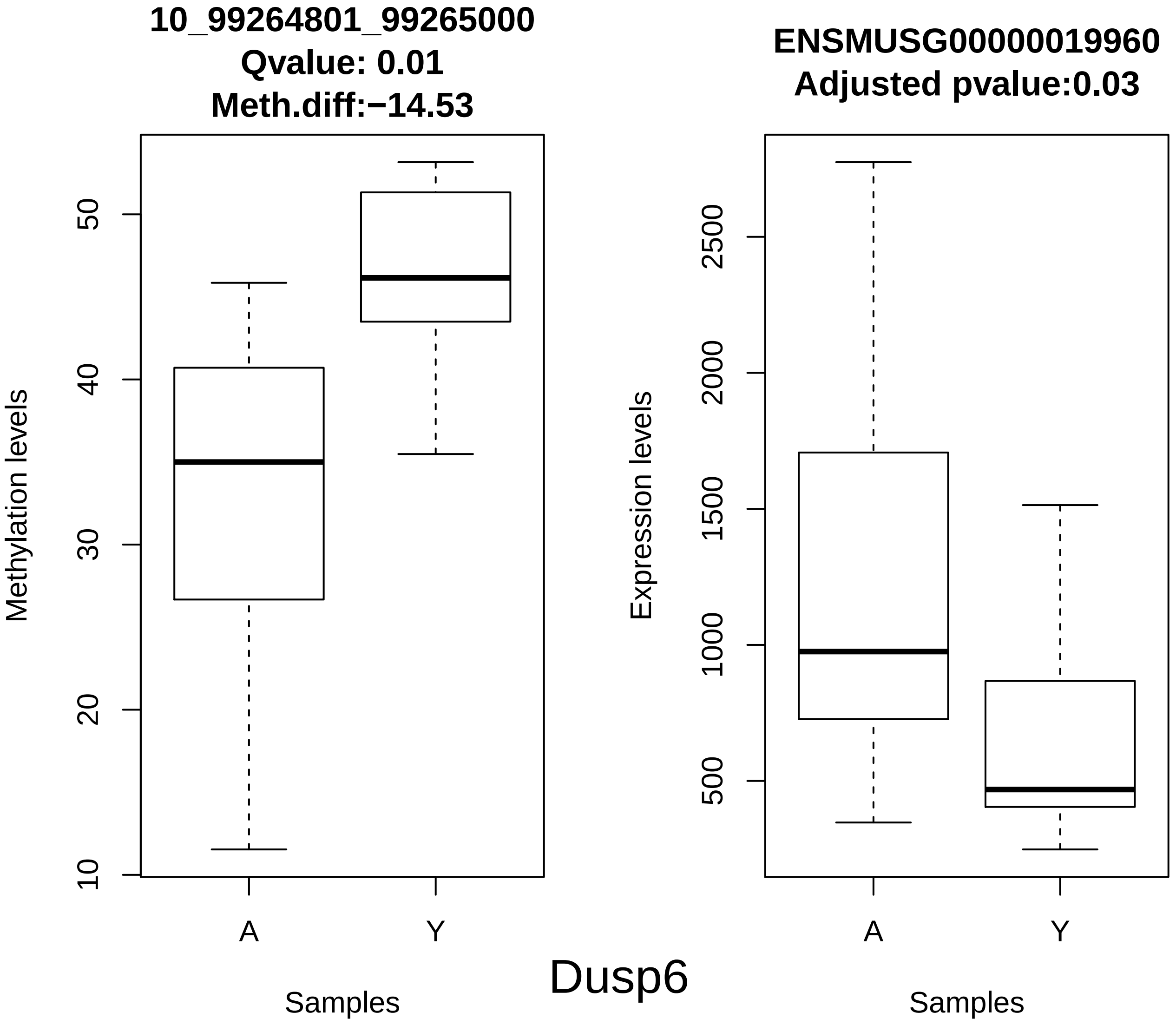
10_99264801_99265000
Qvalue: 0.01
Meth.diff:−14.53
Methylation levels
10
20
30
40
50
A
Y
Samples
ENSMUSG00000019960
Adjusted pvalue:0.03
Expression levels
500
1000
1500
2000
2500
A
Y
Samples
Dusp6

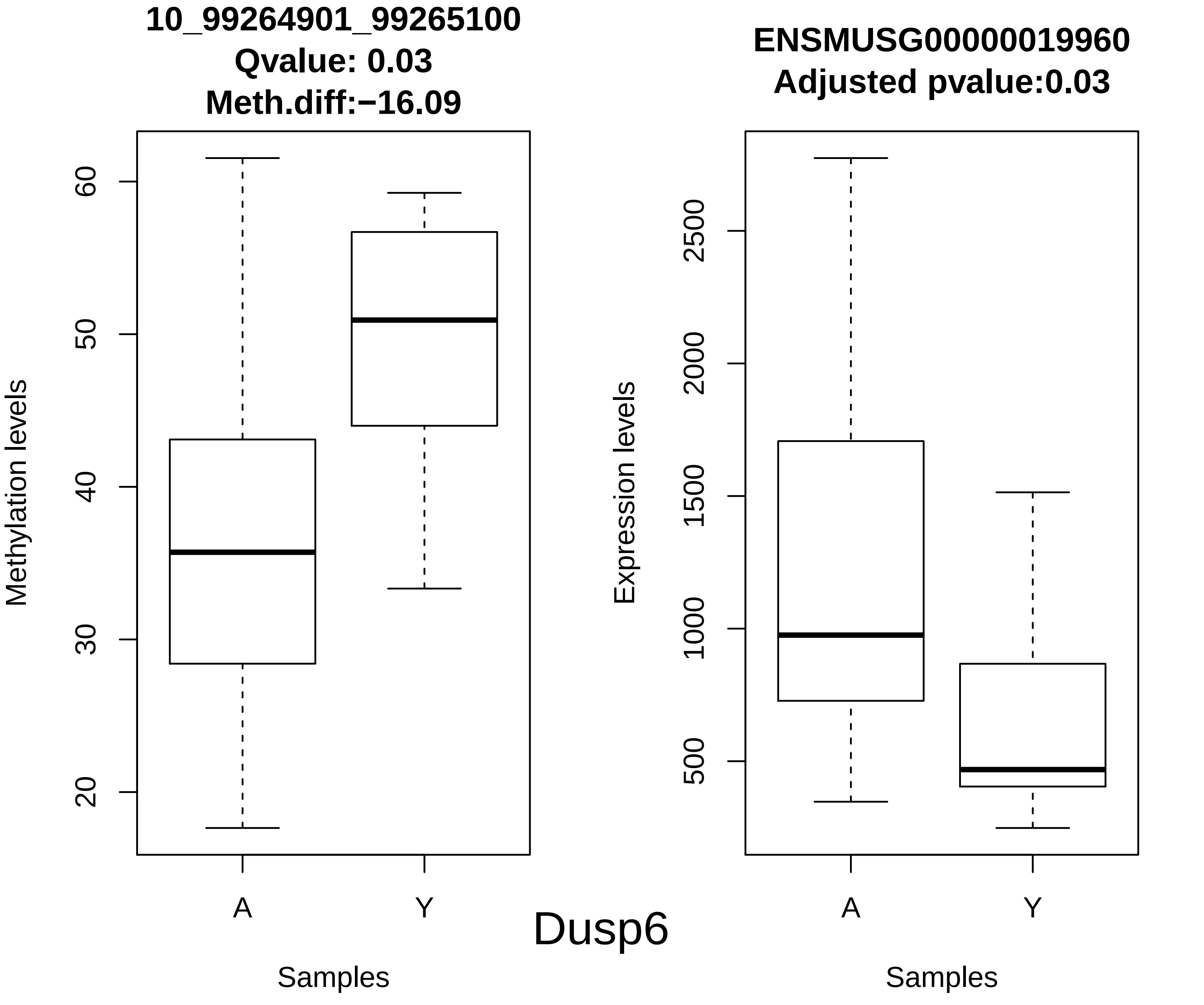
10_99264901_99265100
Qvalue: 0.03
Meth.diff:−16.09
Methylation levels
60
50
40
30
20
A
Y
Samples
ENSMUSG00000019960
Adjusted pvalue:0.03
Expression levels
2500
2000
1500
1000
500
A
Y
Samples
Dusp6

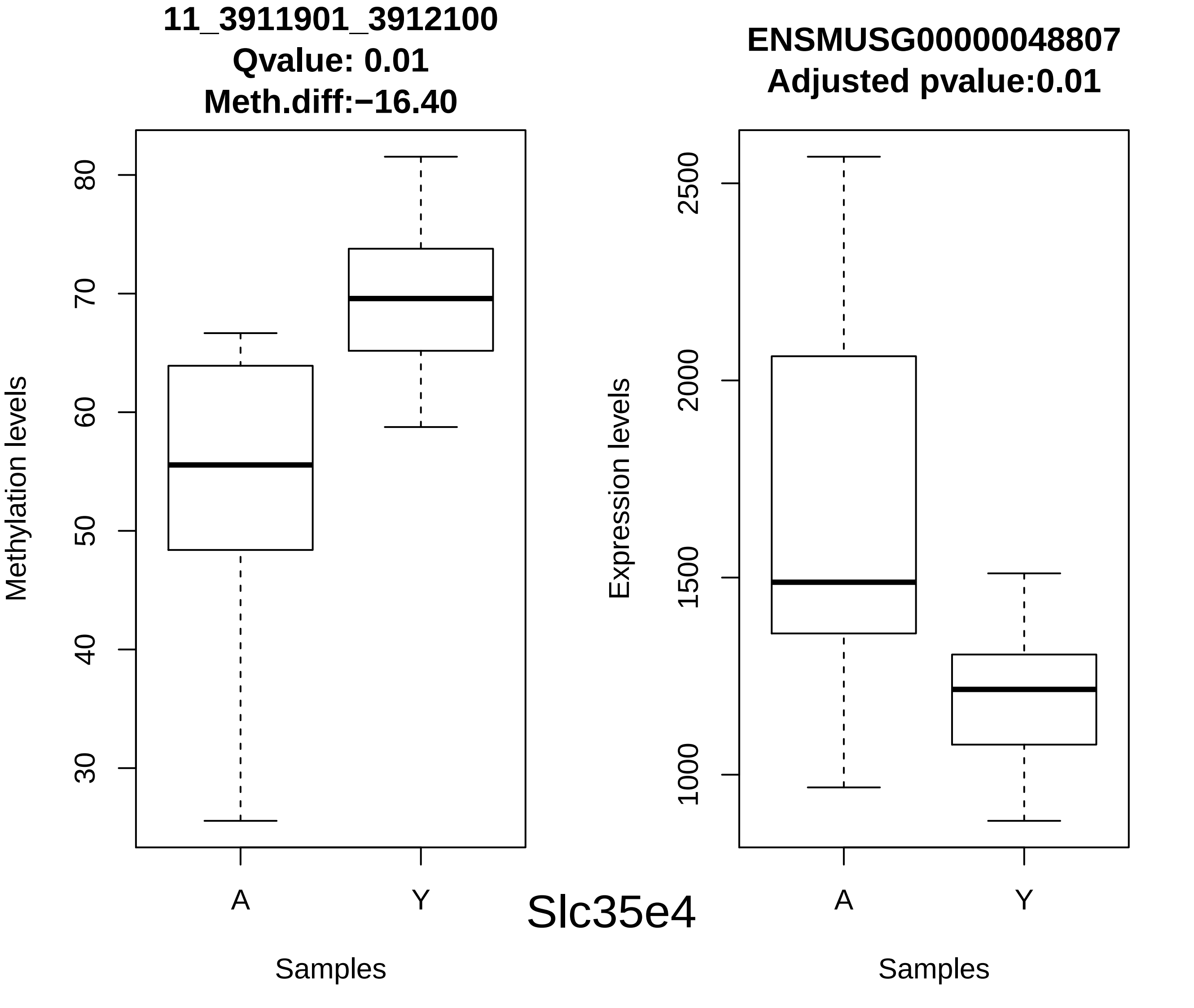
11_3911901_3912100
Qvalue: 0.01
Meth.diff:−16.40
Methylation levels
80
70
60
50
40
30
A
Y
Samples
ENSMUSG00000048807
Adjusted pvalue:0.01
Expression levels
2500
2000
1500
1000
A
Y
Samples
Slc35e4

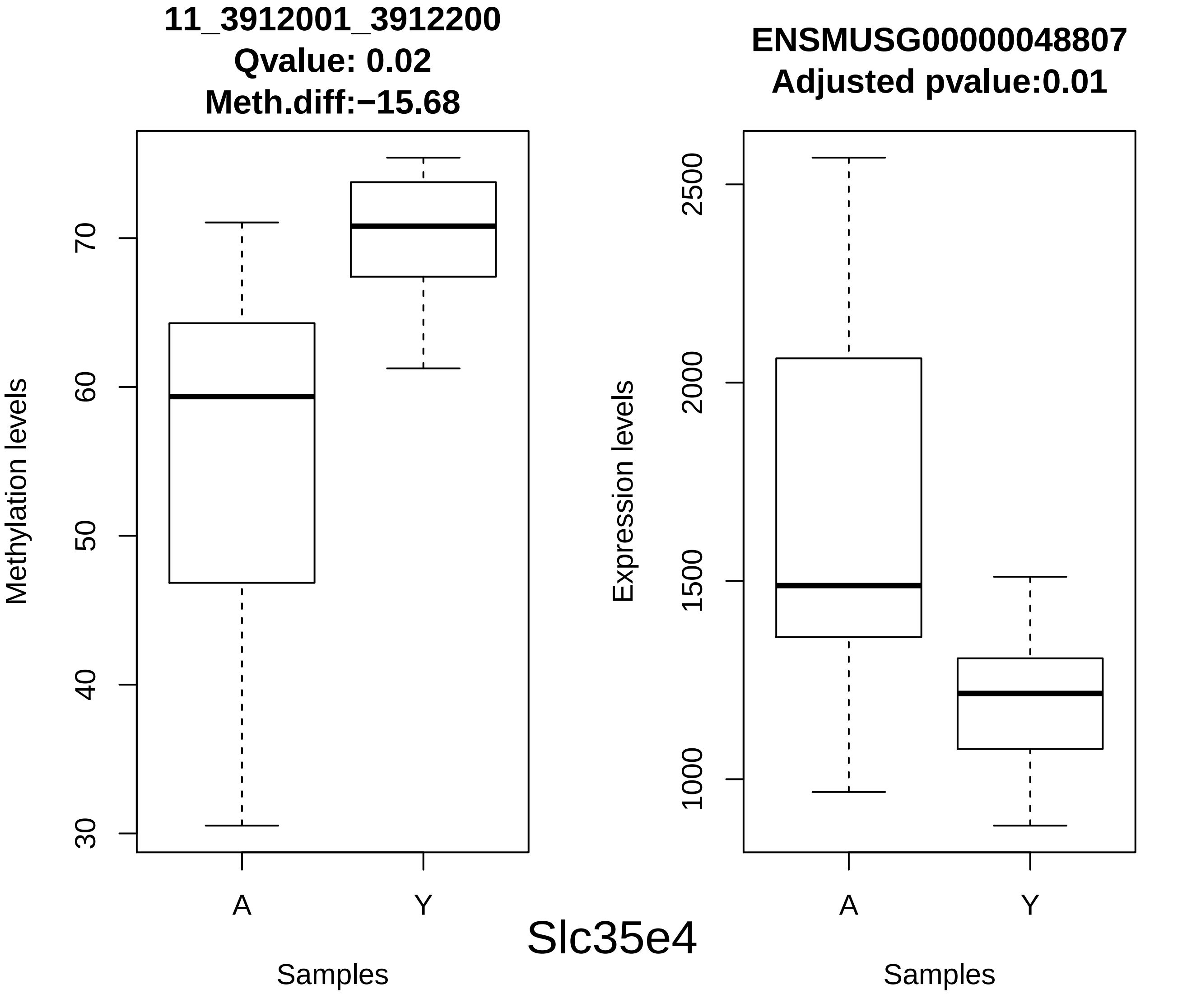
11_3912001_3912200
Qvalue: 0.02
Meth.diff:−15.68
Methylation levels
30
40
50
60
70
A
Y
Samples
ENSMUSG00000048807
Adjusted pvalue:0.01
Expression levels
1000
1500
2000
2500
A
Y
Samples
Slc35e4

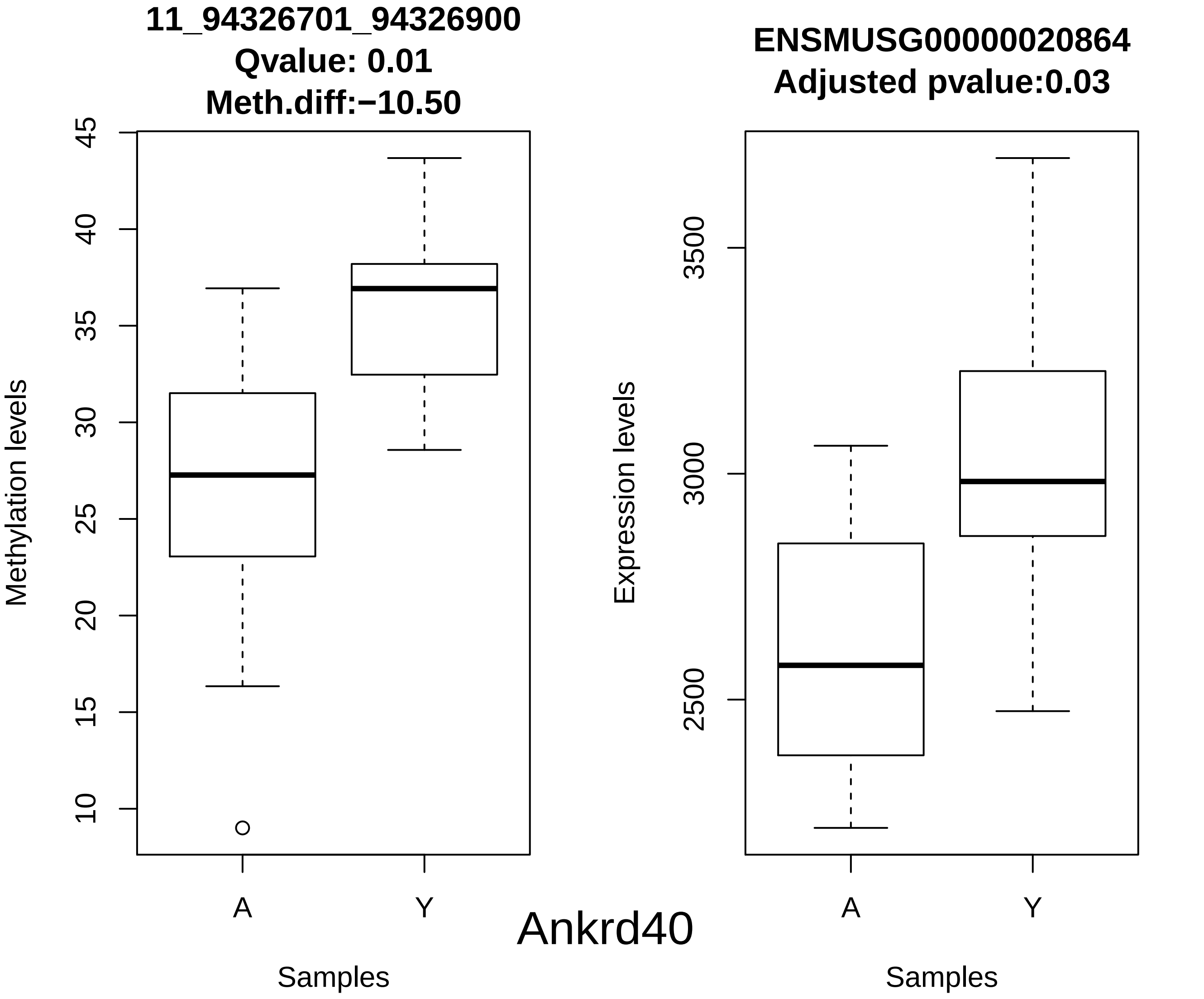
11_94326701_94326900
Qvalue: 0.01
Meth.diff:−10.50
Methylation levels
45
40
35
30
25
20
15
10
A
Y
Samples
ENSMUSG00000020864
Adjusted pvalue:0.03
Expression levels
3500
3000
2500
A
Y
Samples
Ankrd40

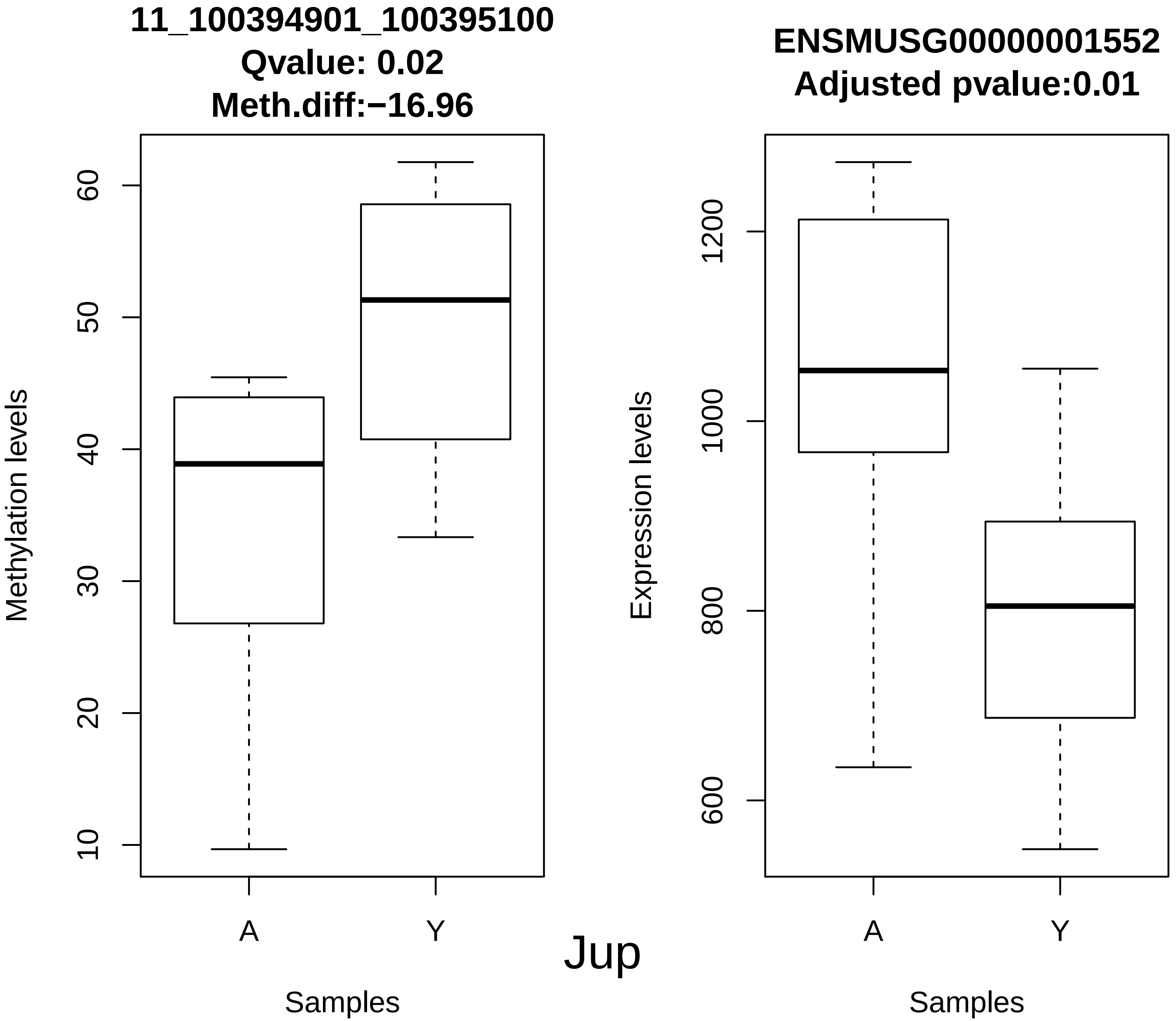

11_100394901_100395100
Qvalue: 0.02
Meth.diff:−16.96
Methylation levels
60
50
40
30
20
10
A
Y
Samples
ENSMUSG00000001552
Adjusted pvalue:0.01
Expression levels
1200
1000
800
600
A
Y
Samples


Jup

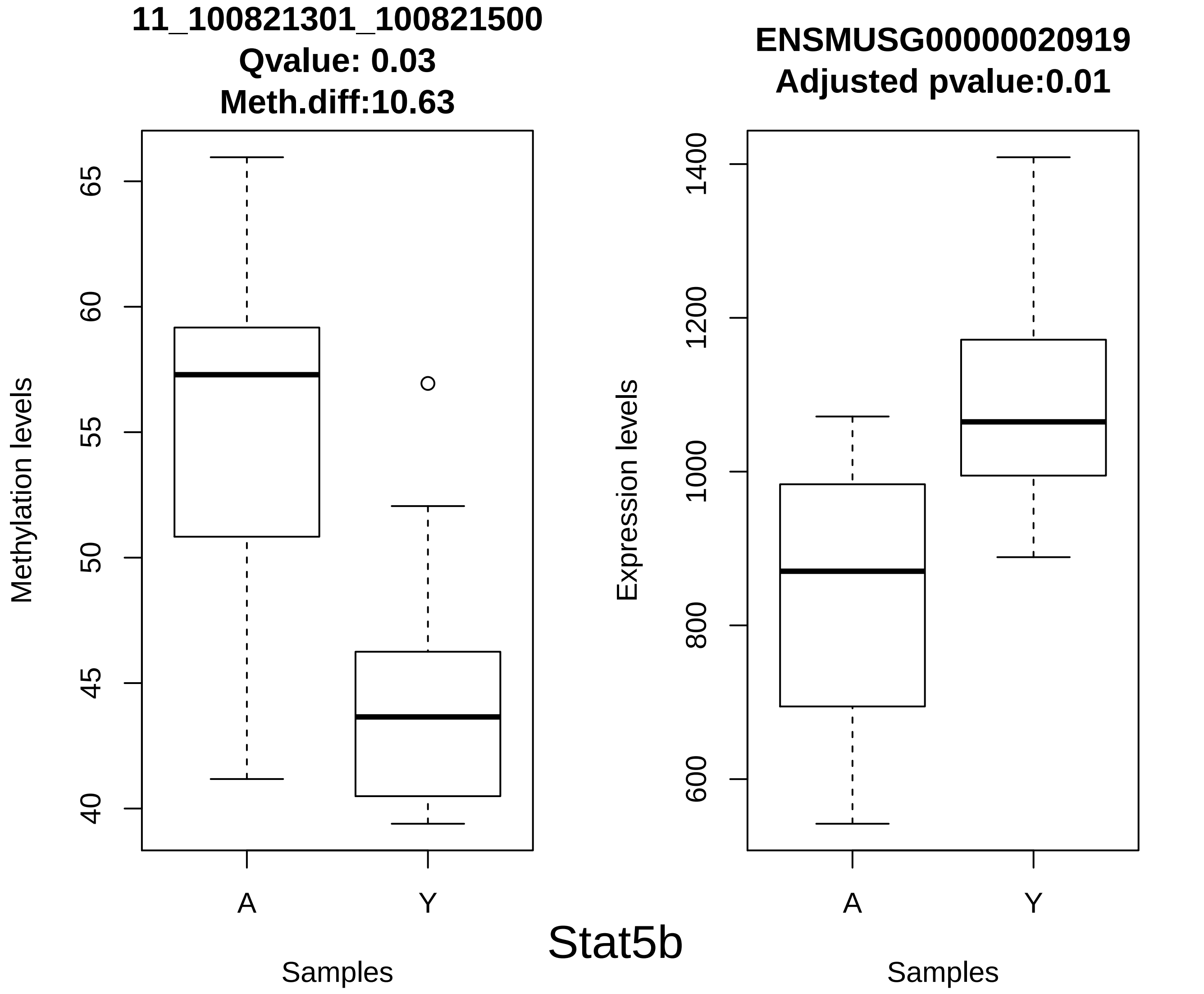
11_100821301_100821500
Qvalue: 0.03
Meth.diff:10.63
Methylation levels
65
60
55
50
45
40
A
Y
Samples
ENSMUSG00000020919
Adjusted pvalue:0.01
Expression levels
1400
1200
1000
800
600
A
Y
Samples
Stat5b

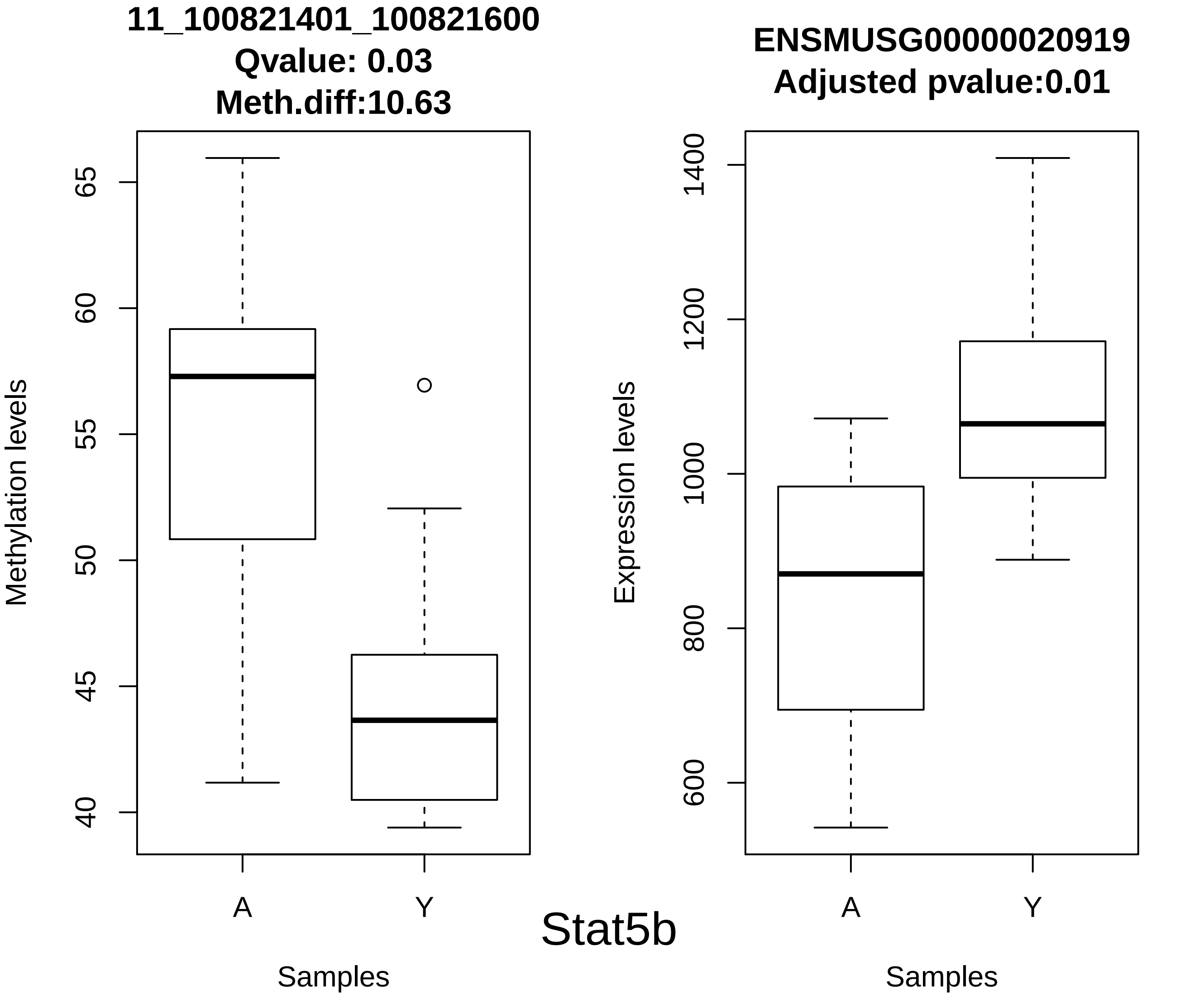
11_100821401_100821600
Qvalue: 0.03
Meth.diff:10.63
Methylation levels
65
60
55
50
45
40
A
Y
Samples
ENSMUSG00000020919
Adjusted pvalue:0.01
Expression levels
1400
1200
1000
800
600
A
Y
Samples

Stat5b

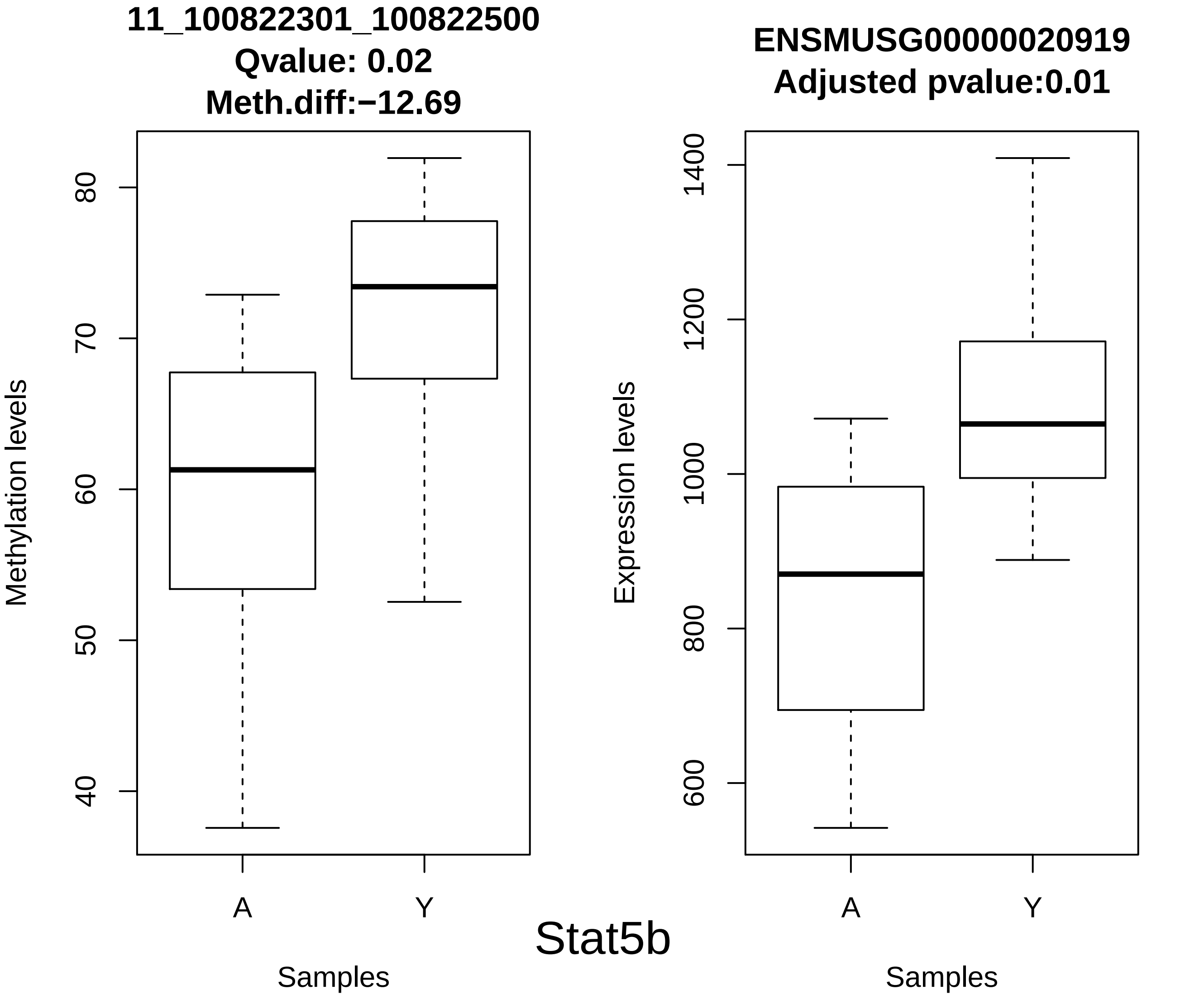
11_100822301_100822500
Qvalue: 0.02
Meth.diff:−12.69
Methylation levels
80
70
60
50
40
A
Y
Samples
ENSMUSG00000020919
Adjusted pvalue:0.01
Expression levels
1400
1200
1000
800
600
A
Y
Samples
Stat5b

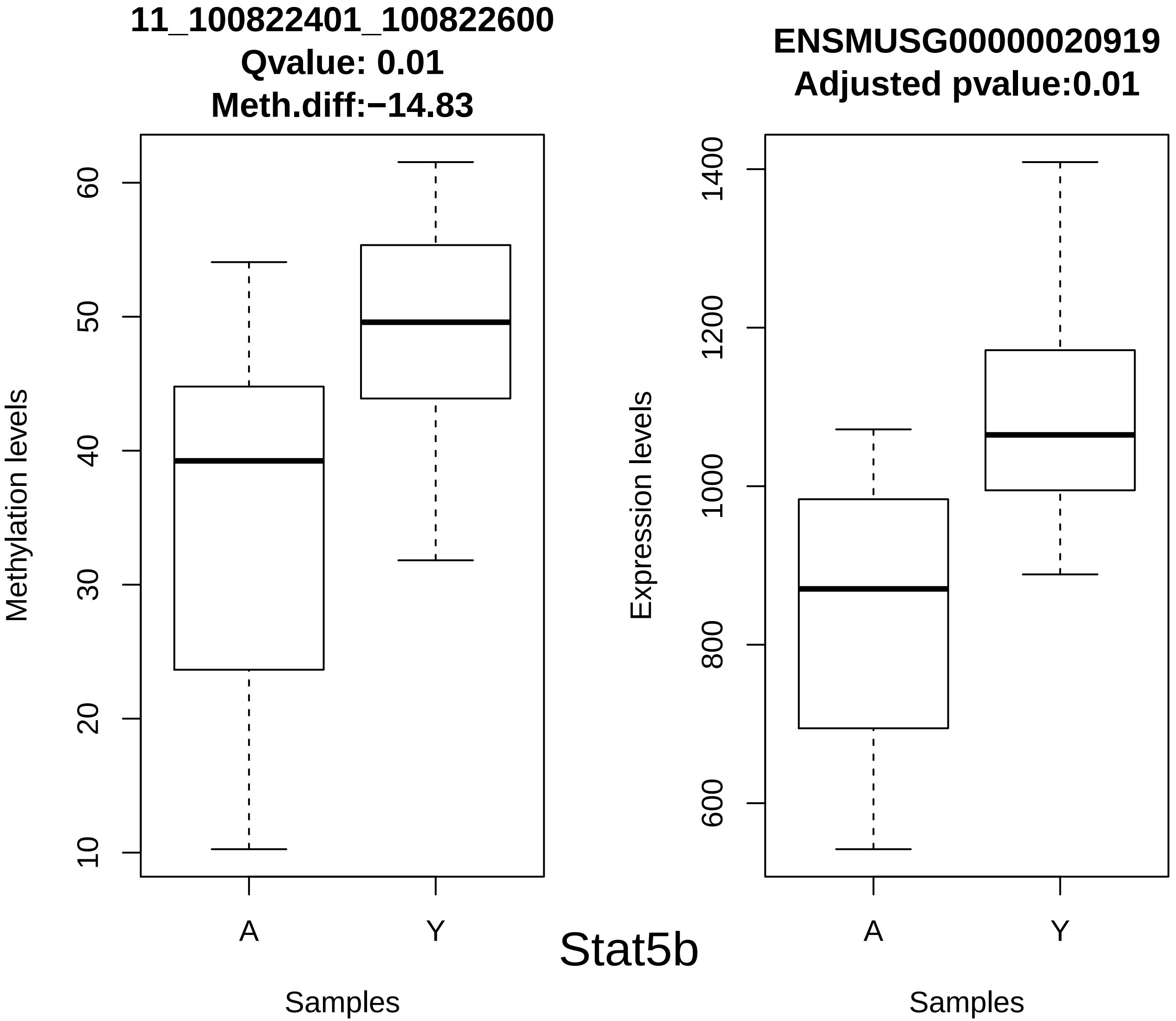
11_100822401_100822600
Qvalue: 0.01
Meth.diff:−14.83
Methylation levels
10
20
30
40
50
60
A
Y
Samples
ENSMUSG00000020919
Adjusted pvalue:0.01
Expression levels
600
800
1000
1200
1400
A
Y
Samples

Stat5b

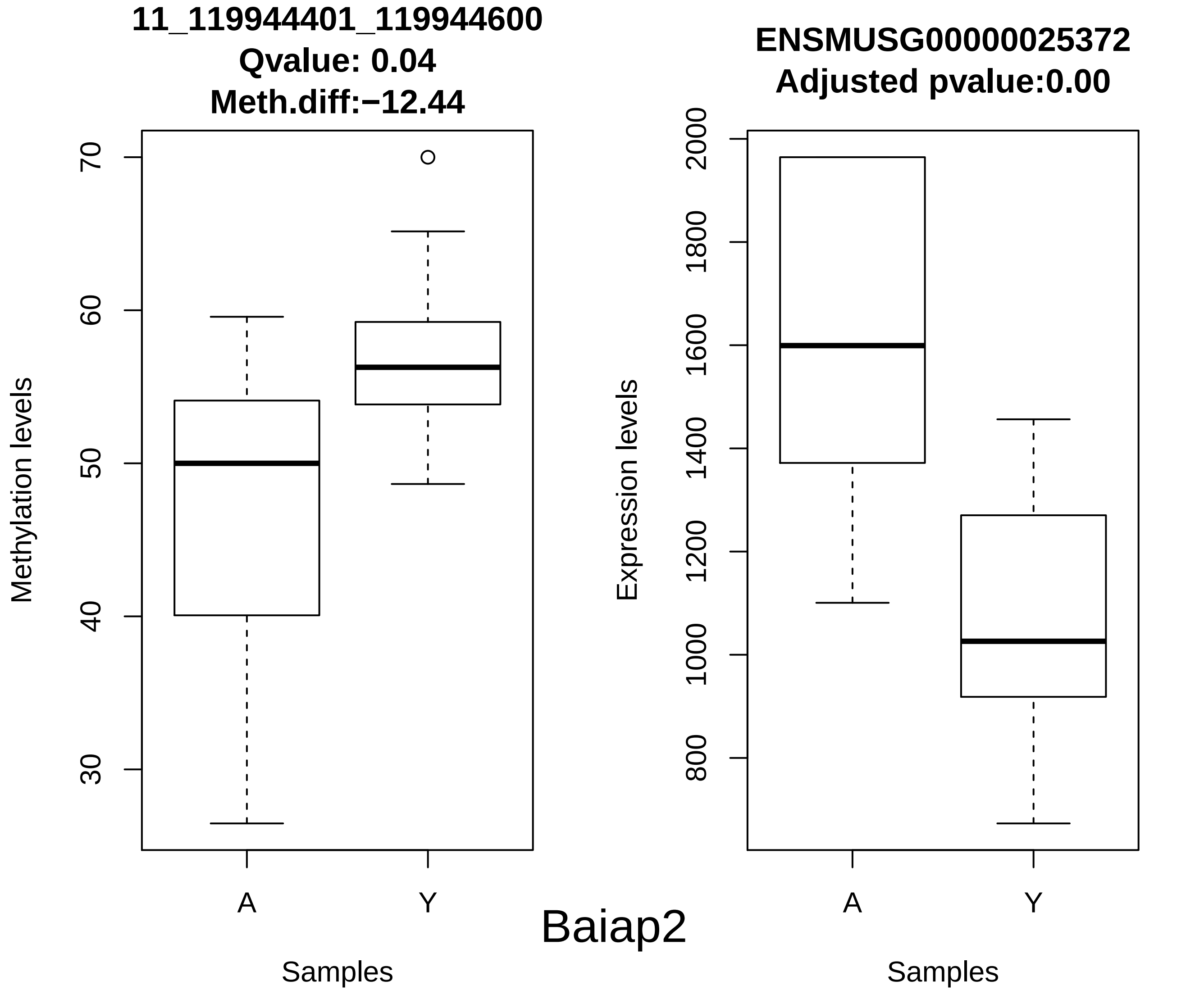
11_119944401_119944600
Qvalue: 0.04
Meth.diff:−12.44
Methylation levels
70
60
50
40
30
A
Y
Samples
ENSMUSG00000025372
Adjusted pvalue:0.00
Expression levels
2000
1800
1600
1400
1200
1000
800
A
Y
Samples
Baiap2

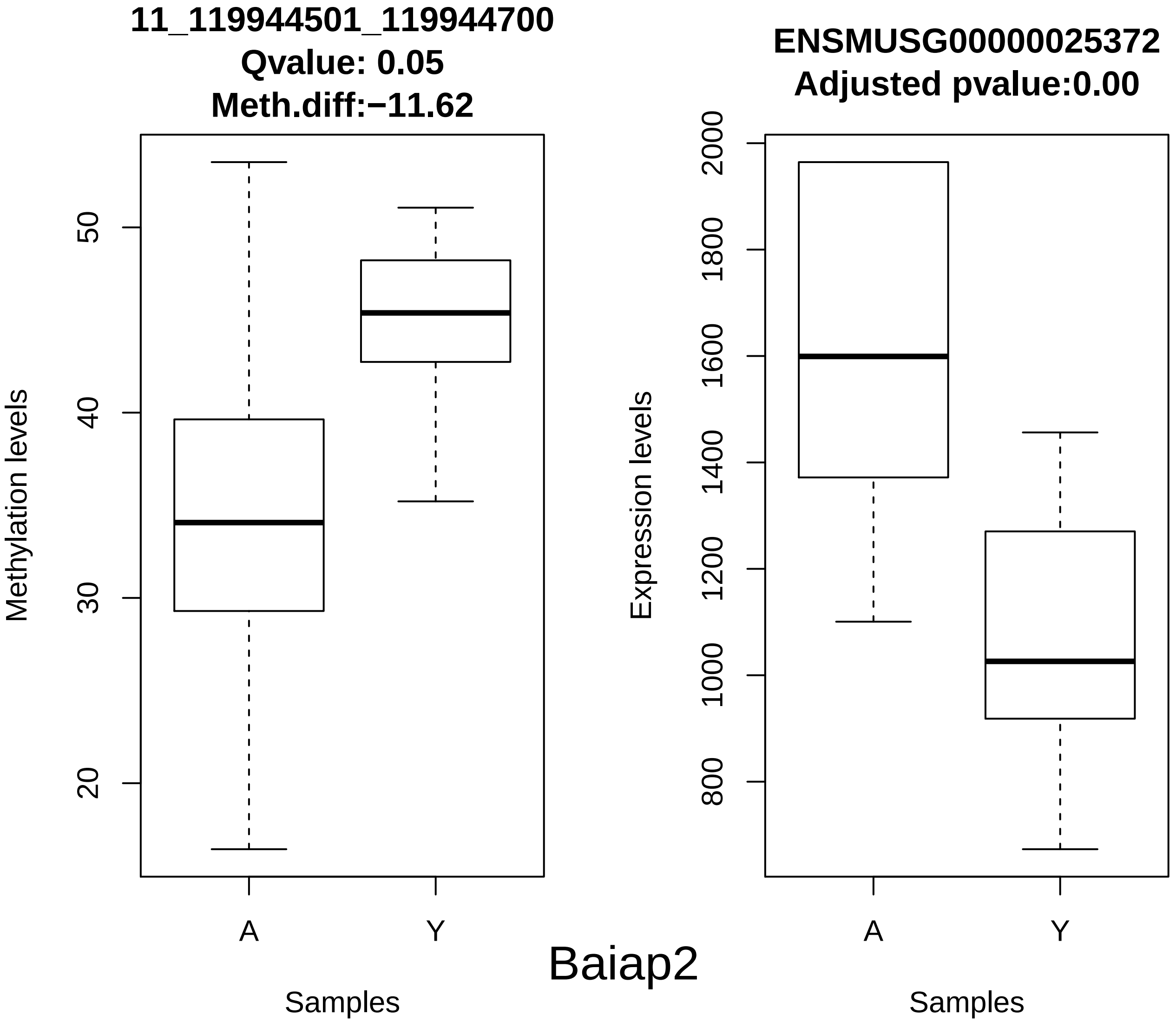

11_119944501_119944700
Qvalue: 0.05
Meth.diff:−11.62
Methylation levels
20
30
40
50
A
Y
Samples
ENSMUSG00000025372
Adjusted pvalue:0.00
Expression levels
800
1000
1200
1400
1600
1800
2000
A
Y
Samples


Baiap2

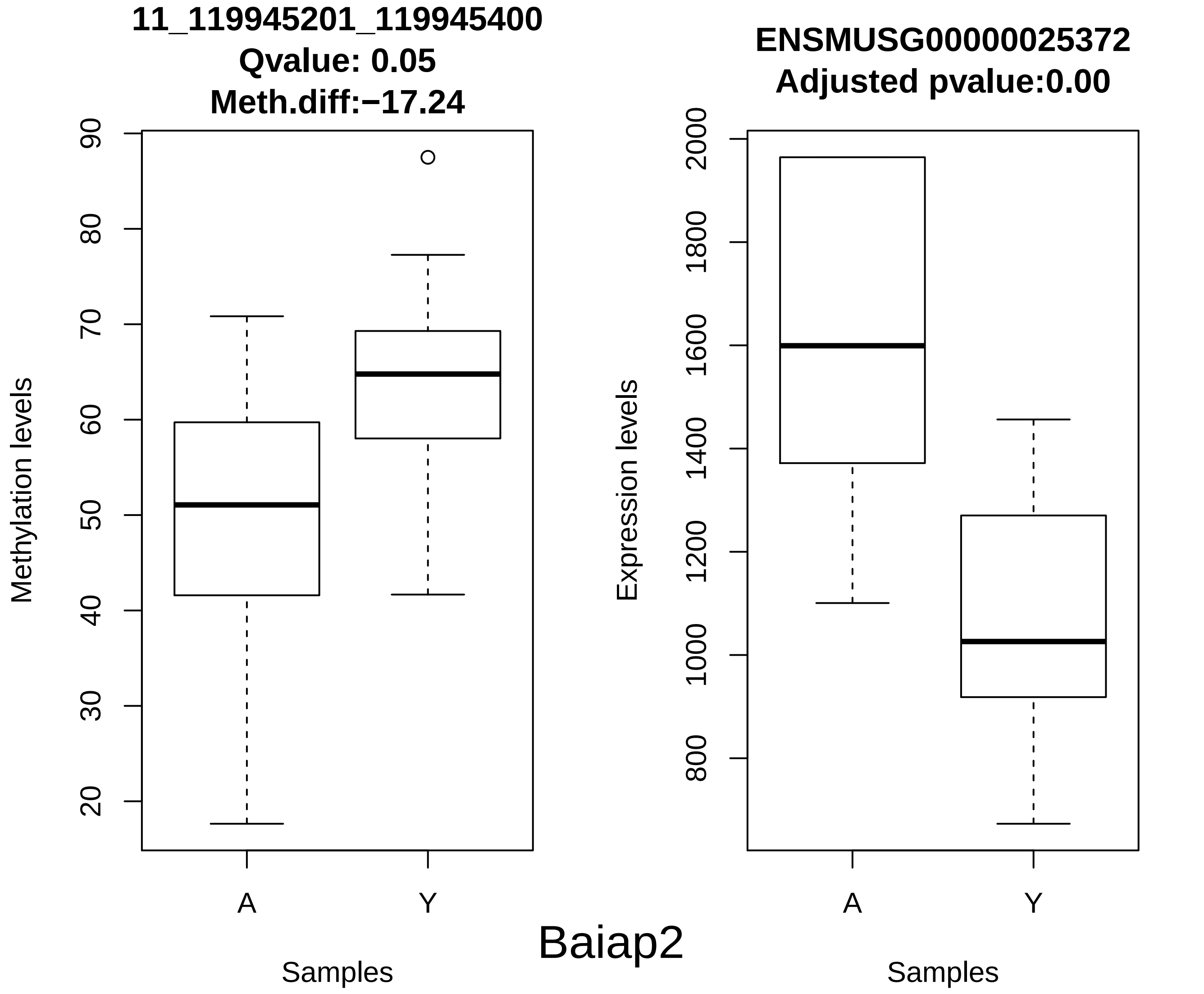
11_119945201_119945400
Qvalue: 0.05
Meth.diff:−17.24
Methylation levels
90
80
70
60
50
40
30
20
A
Y
Samples
ENSMUSG00000025372
Adjusted pvalue:0.00
Expression levels
2000
1800
1600
1400
1200
1000
800
A
Y
Samples
Baiap2

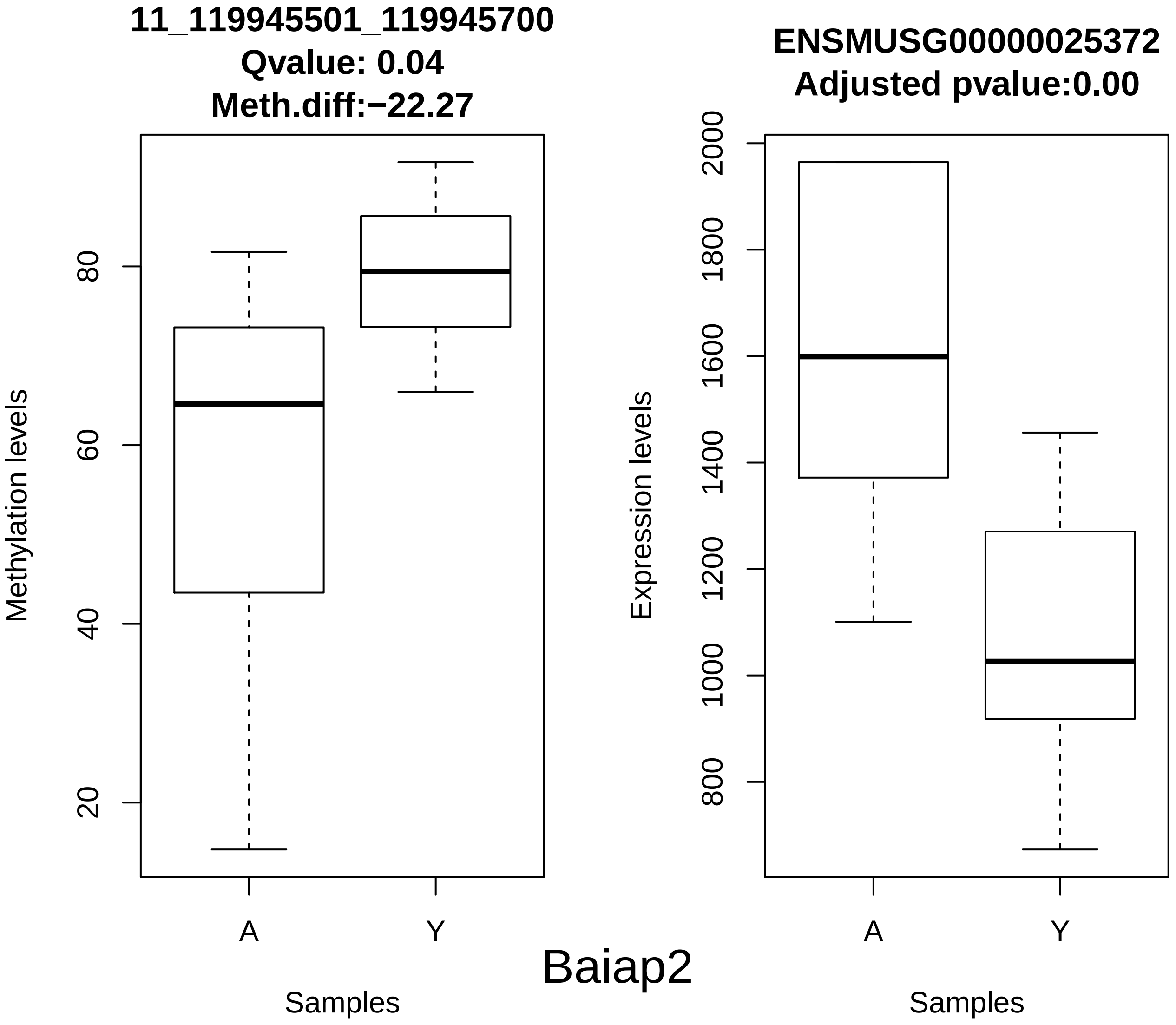
11_119945501_119945700
Qvalue: 0.04
Meth.diff:−22.27
Methylation levels
20
40
60
80
A
Y
Samples
ENSMUSG00000025372
Adjusted pvalue:0.00
Expression levels
800
1000
1200
1400
1600
1800
2000
A
Y
Samples
Baiap2

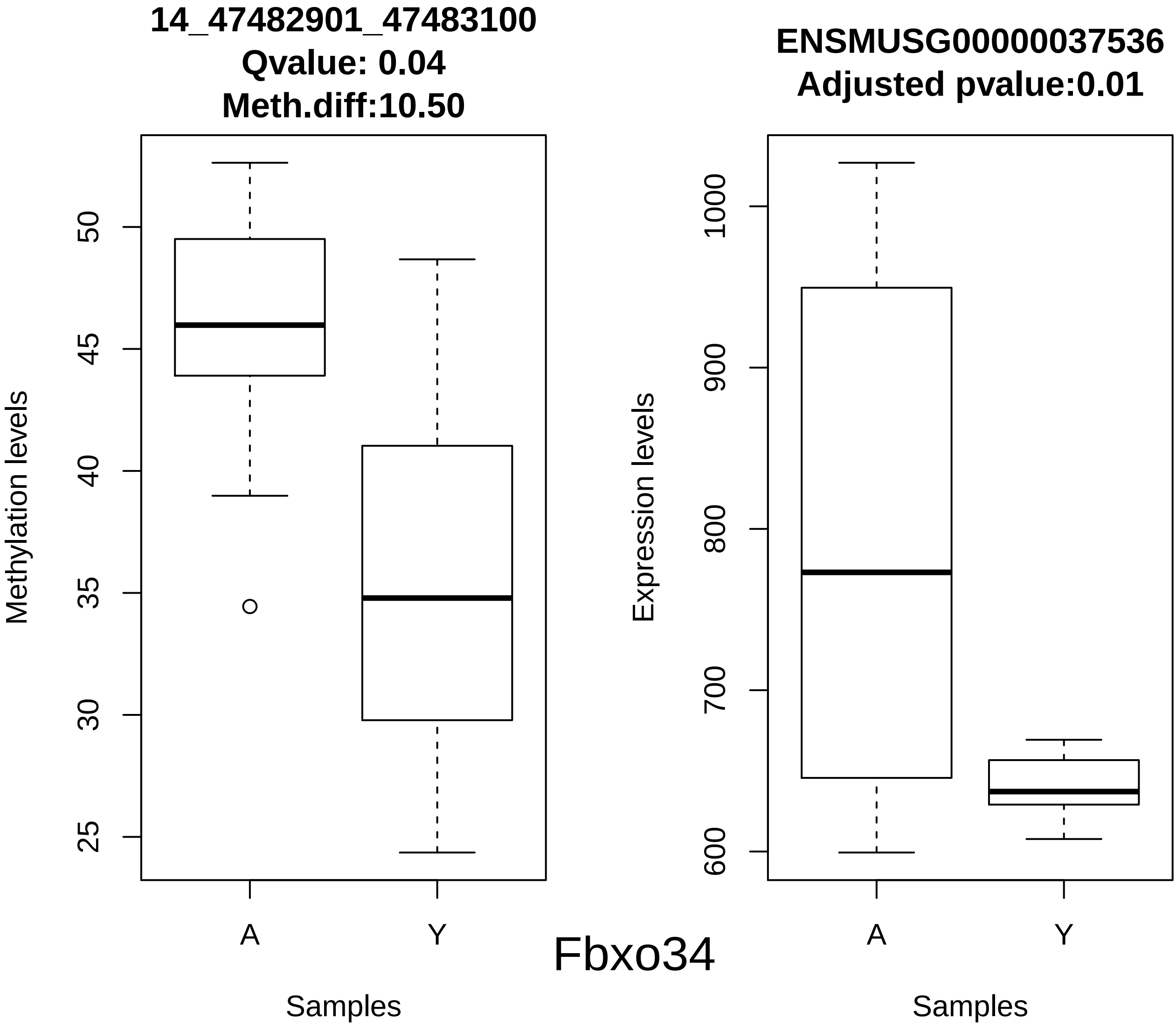
14_47482901_47483100
Qvalue: 0.04
Meth.diff:10.50
Methylation levels
50
45
40
35
30
25
A
Y
Samples
ENSMUSG00000037536
Adjusted pvalue:0.01
Expression levels
1000
900
800
700
600
A
Y
Samples
Fbxo34

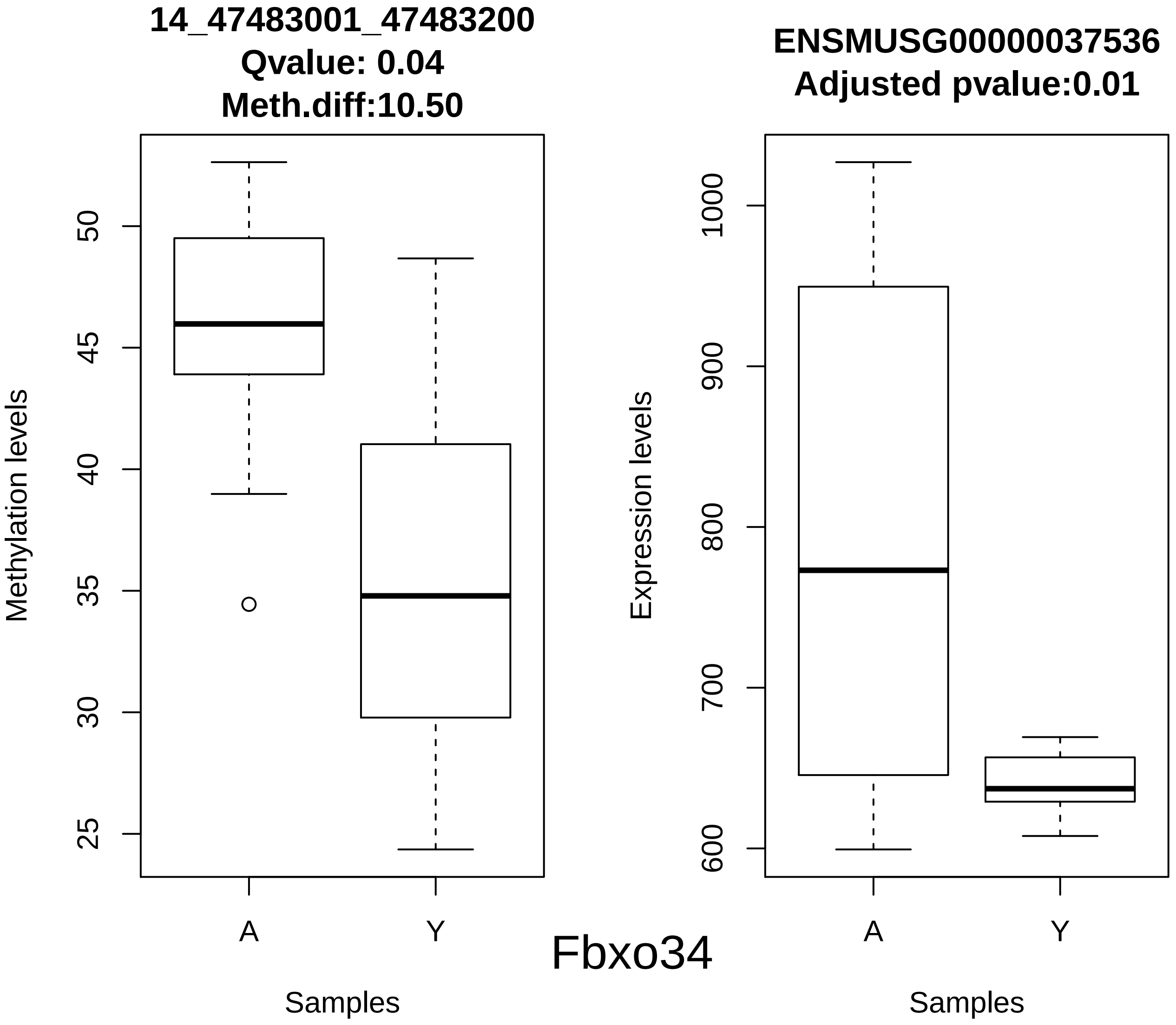
14_47483001_47483200
Qvalue: 0.04
Meth.diff:10.50
Methylation levels
50
45
40
35
30
25
A
Y
Samples
ENSMUSG00000037536
Adjusted pvalue:0.01
Expression levels
1000
900
800
700
600
A
Y
Samples
Fbxo34

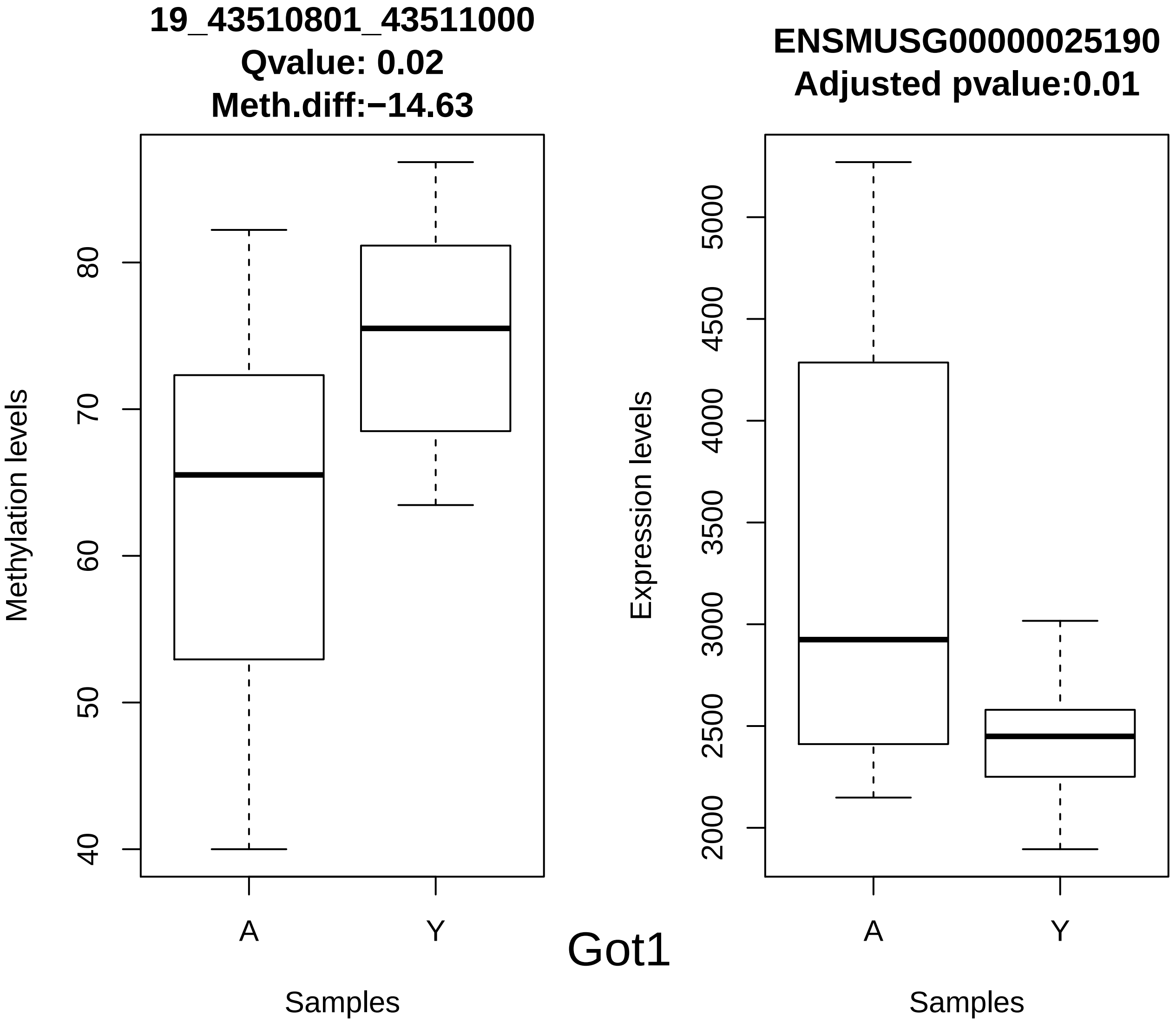
19_43510801_43511000
Qvalue: 0.02
Meth.diff:−14.63
Methylation levels
40
50
60
70
80
A
Y
Samples
ENSMUSG00000025190
Adjusted pvalue:0.01
Expression levels
2000
2500
3000
3500
4000
4500
5000
A
Y
Samples
Got1

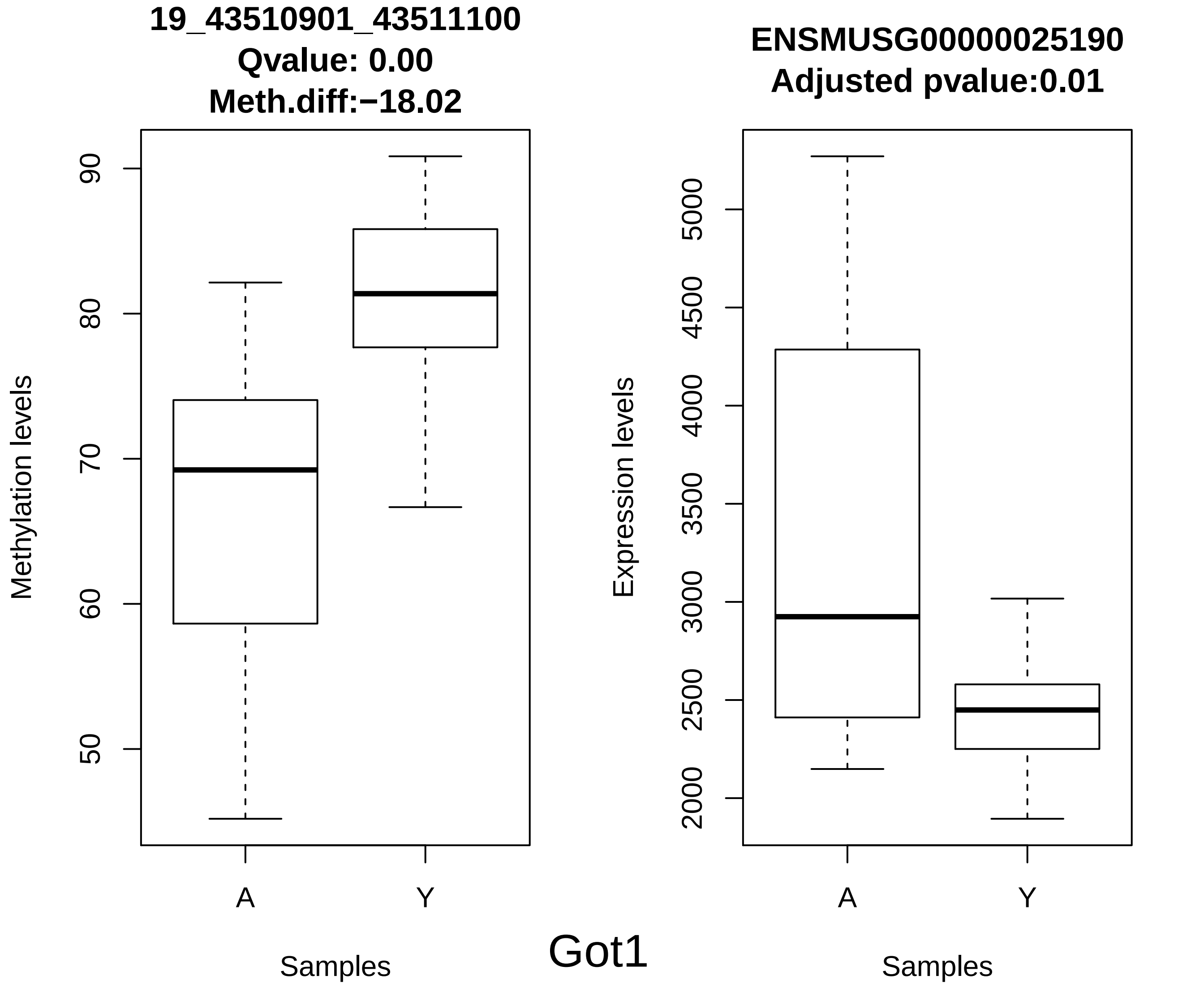
19_43510901_43511100
Qvalue: 0.00
Meth.diff:−18.02
Methylation levels
90
80
70
60
50
A
Y
Samples
ENSMUSG00000025190
Adjusted pvalue:0.01
Expression levels
5000
4500
4000
3500
3000
2500
2000
A
Y
Samples
Got1

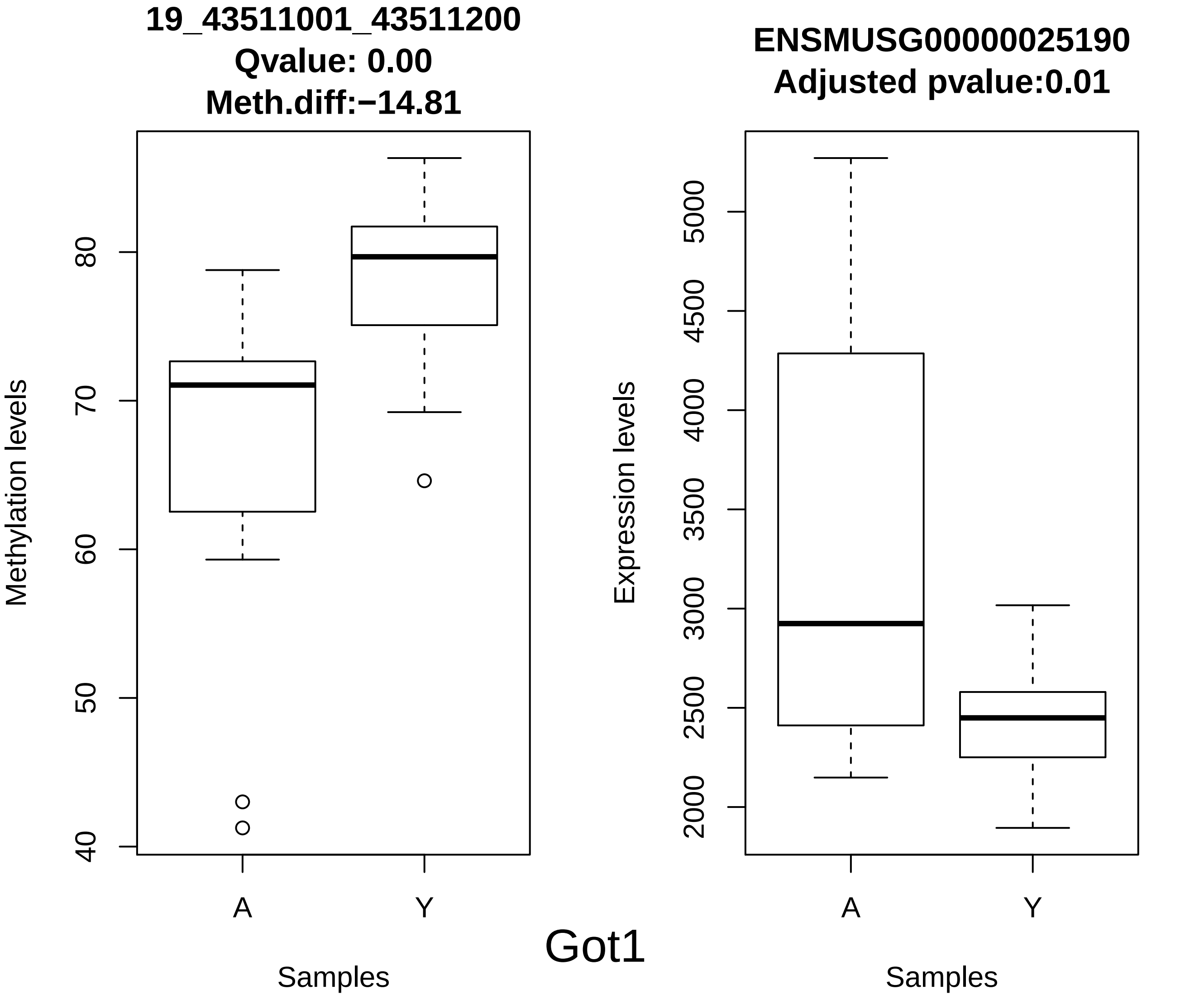

19_43511001_43511200
Qvalue: 0.00
Meth.diff:−14.81
Methylation levels
40
50
60
70
80
A
Y
Samples
ENSMUSG00000025190
Adjusted pvalue:0.01
Expression levels
2000
2500
3000
3500
4000
4500
5000
A
Y
Samples
Got1

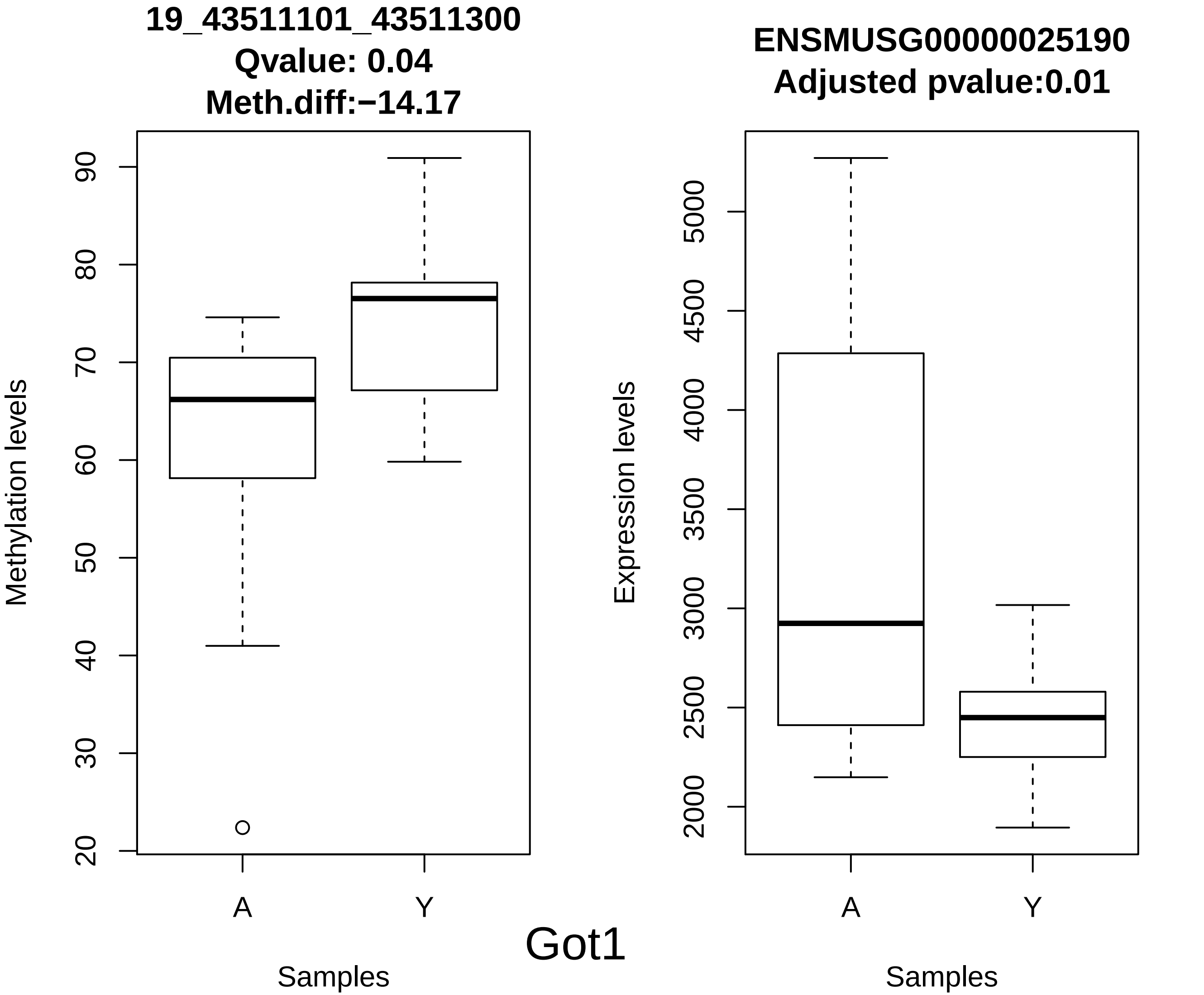

19_43511101_43511300
Qvalue: 0.04
Meth.diff:−14.17
Methylation levels
20
30
40
50
60
70
80
90
A
Y
Samples
ENSMUSG00000025190
Adjusted pvalue:0.01
Expression levels
2000
2500
3000
3500
4000
4500
5000
A
Y
Samples
Got1

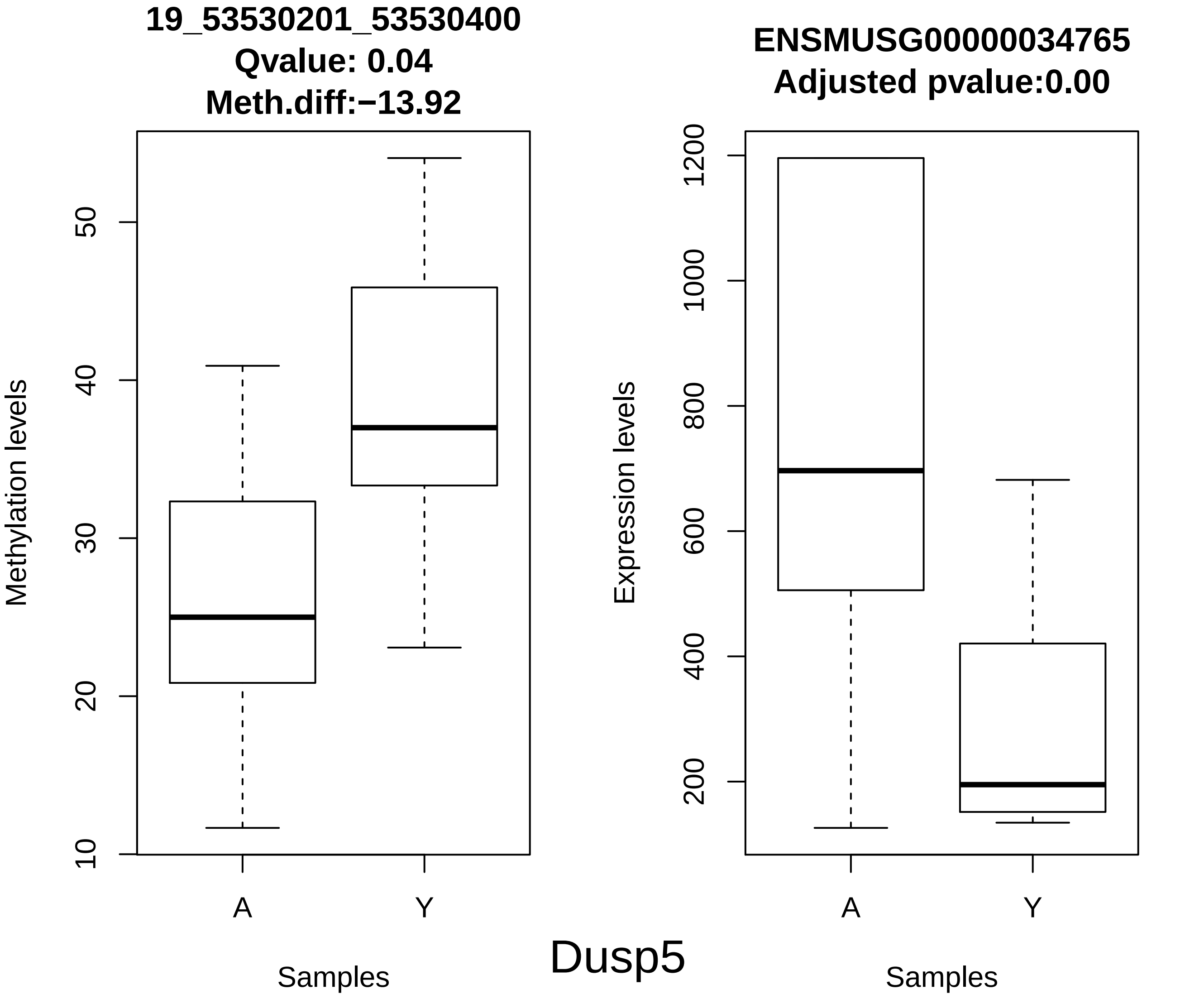
19_53530201_53530400
Qvalue: 0.04
Meth.diff:−13.92
Methylation levels
10
20
30
40
50
A
Y
Samples
ENSMUSG00000034765
Adjusted pvalue:0.00
Expression levels
200
400
600
800
1000
1200
A
Y
Samples

Dusp5

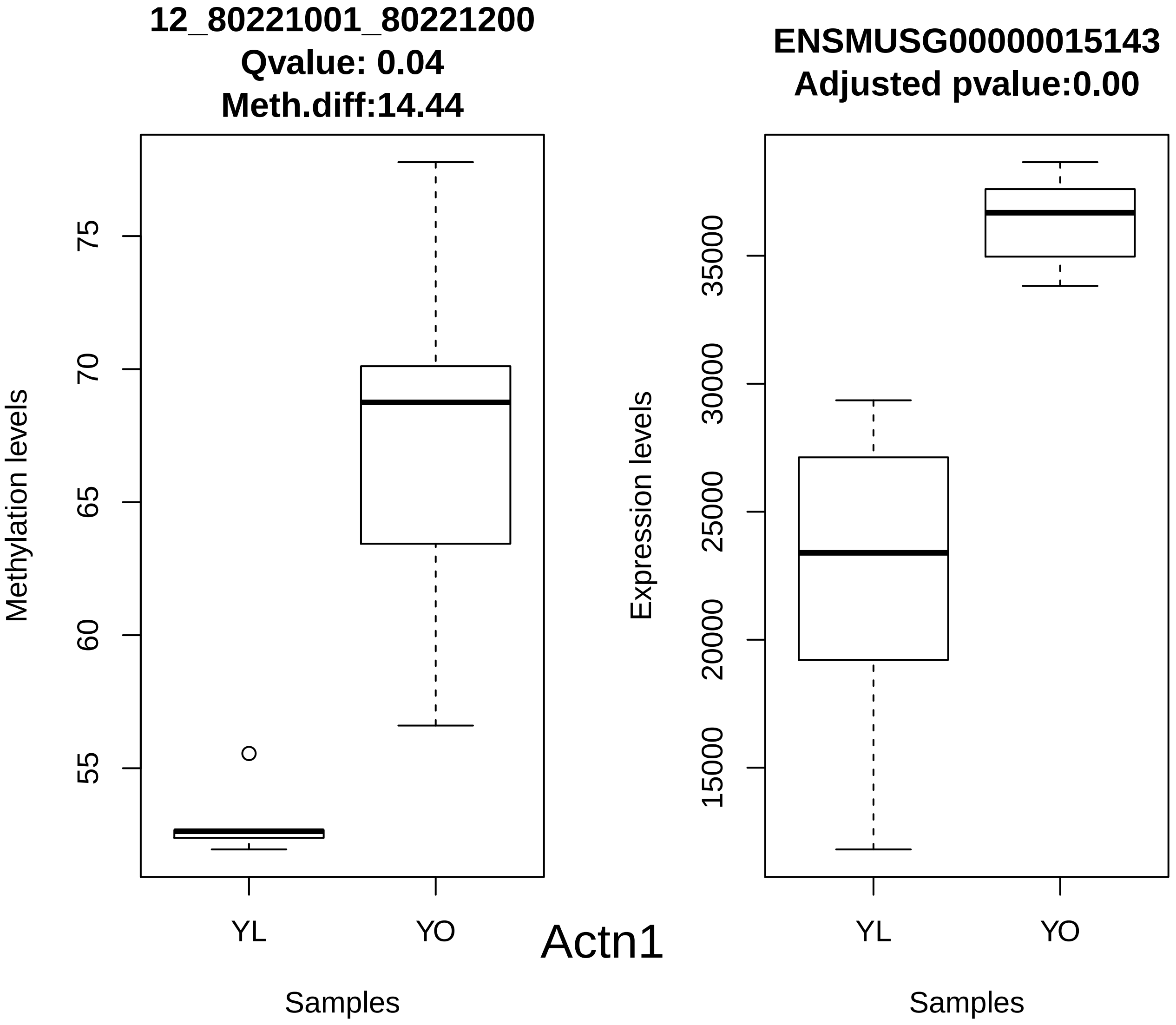
12_80221001_80221200
Qvalue: 0.04
Meth.diff:14.44
Methylation levels
75
70
65
60
55
YL
YO
Samples
ENSMUSG00000015143
Adjusted pvalue:0.00
Expression levels
35000
30000
25000
20000
15000
YL
YO
Samples
Actn1

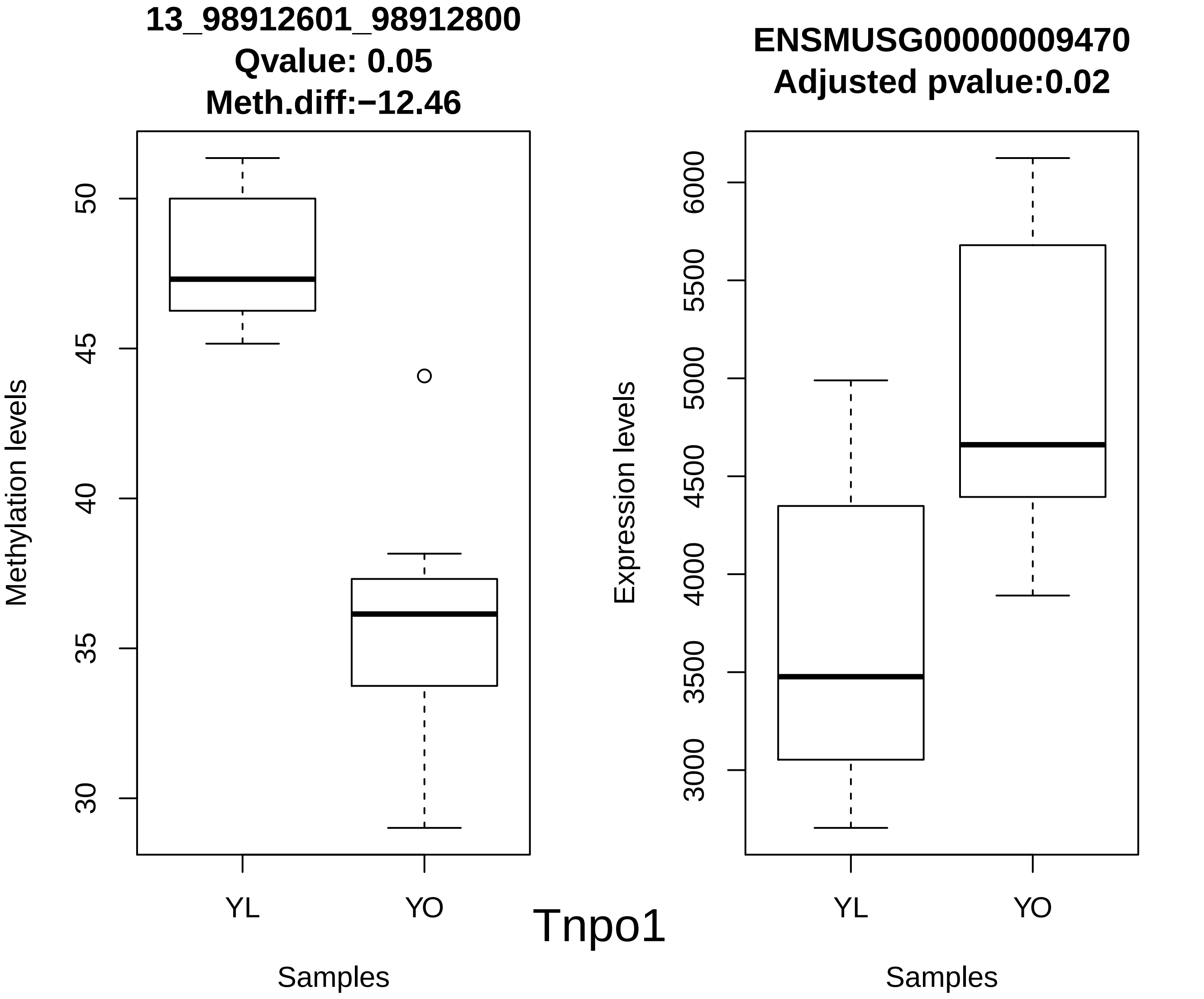
13_98912601_98912800
Qvalue: 0.05
Meth.diff:−12.46
Methylation levels
50
45
40
35
30
YL
YO
Samples
ENSMUSG00000009470
Adjusted pvalue:0.02
Expression levels
6000
5500
5000
4500
4000
3500
3000
YL
YO
Samples
Tnpo1

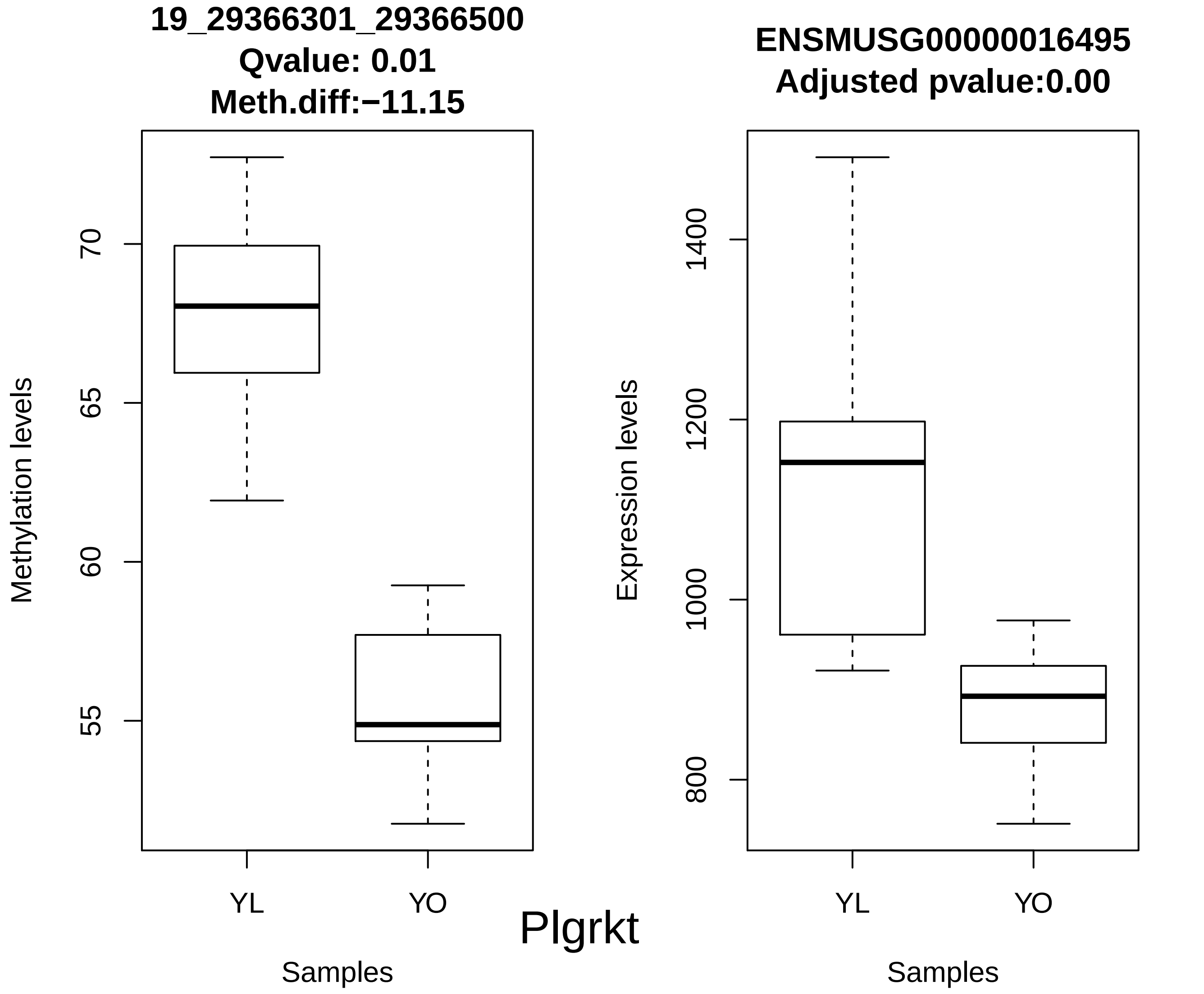

19_29366301_29366500
Qvalue: 0.01
Meth.diff:−11.15
Methylation levels
70
65
60
55
YL
YO
Samples
ENSMUSG00000016495
Adjusted pvalue:0.00
Expression levels
1400
1200
1000
800
YL
YO
Samples
Plgrkt

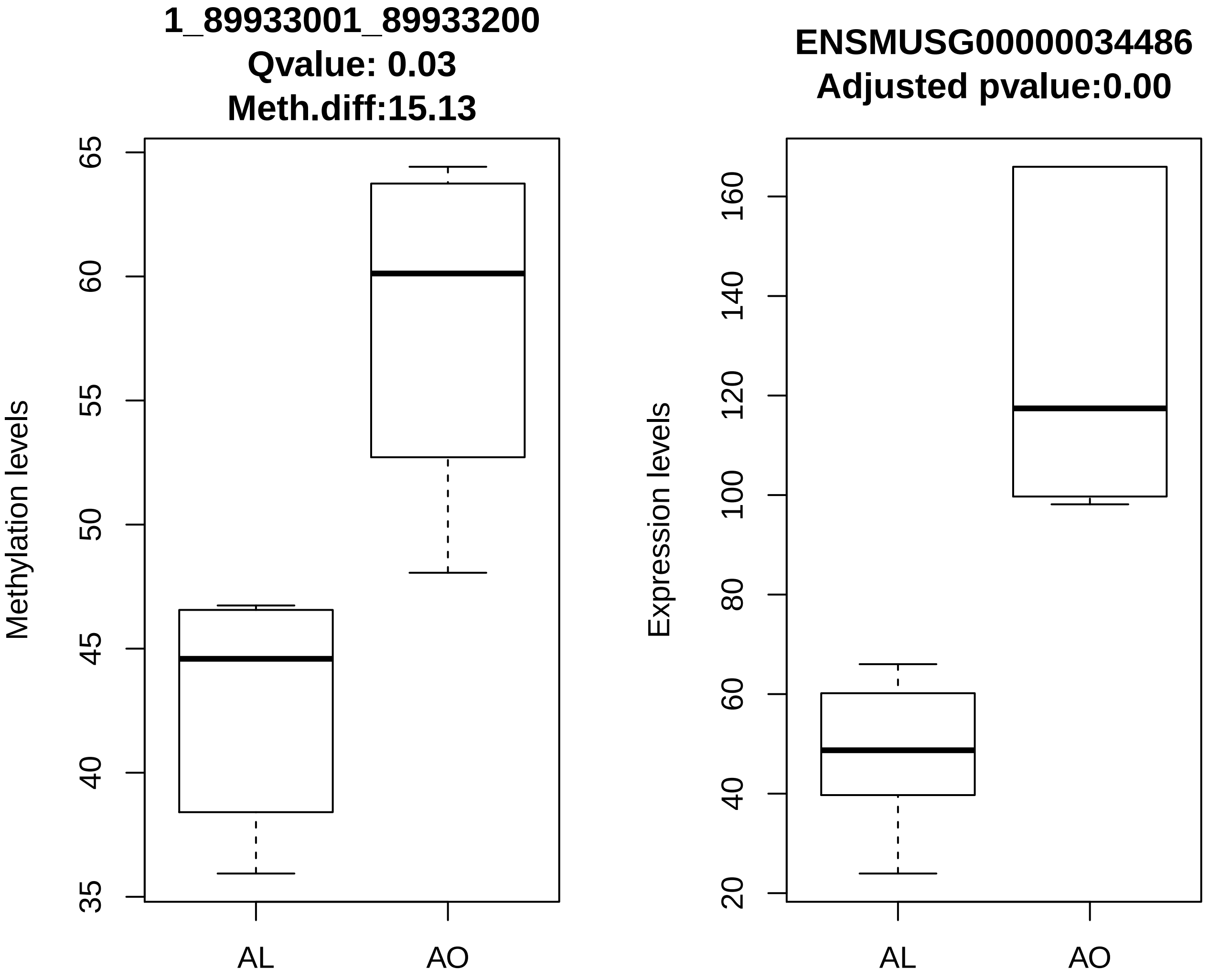
1_89933001_89933200
Qvalue: 0.03
Meth.diff:15.13
Methylation levels
35
40
45
50
55
60
65
AL
AO
ENSMUSG00000034486
Adjusted pvalue:0.00
Expression levels
20
40
60
80
100
120
140
160
AL
AO

Gbx2

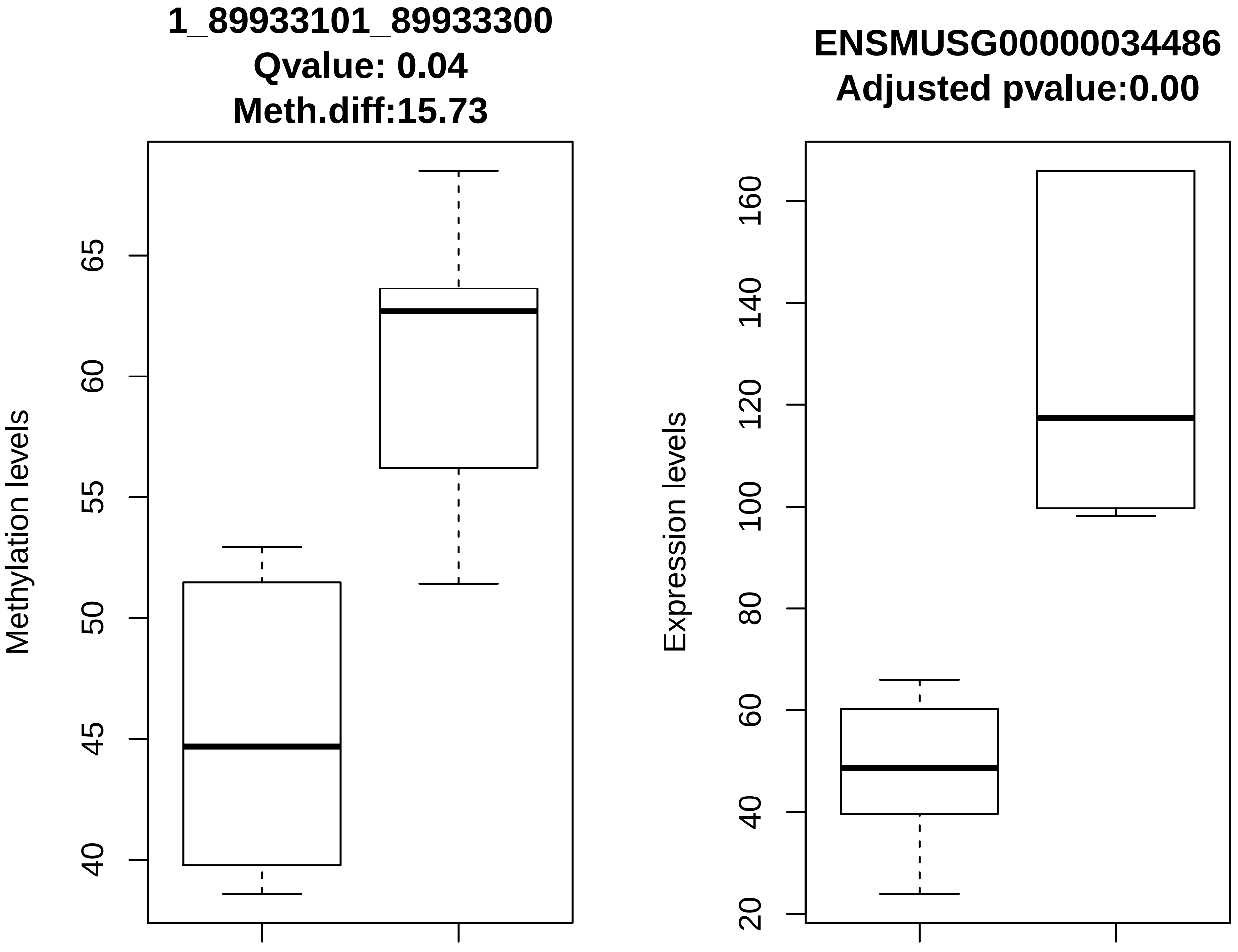
1_89933101_89933300
Qvalue: 0.04
Meth.diff:15.73
Methylation levels
40
45
50
55
60
65
AL
AO
ENSMUSG00000034486
Adjusted pvalue:0.00
Expression levels
20
40
60
80
100
120
140
160
AL
AO

Gbx2

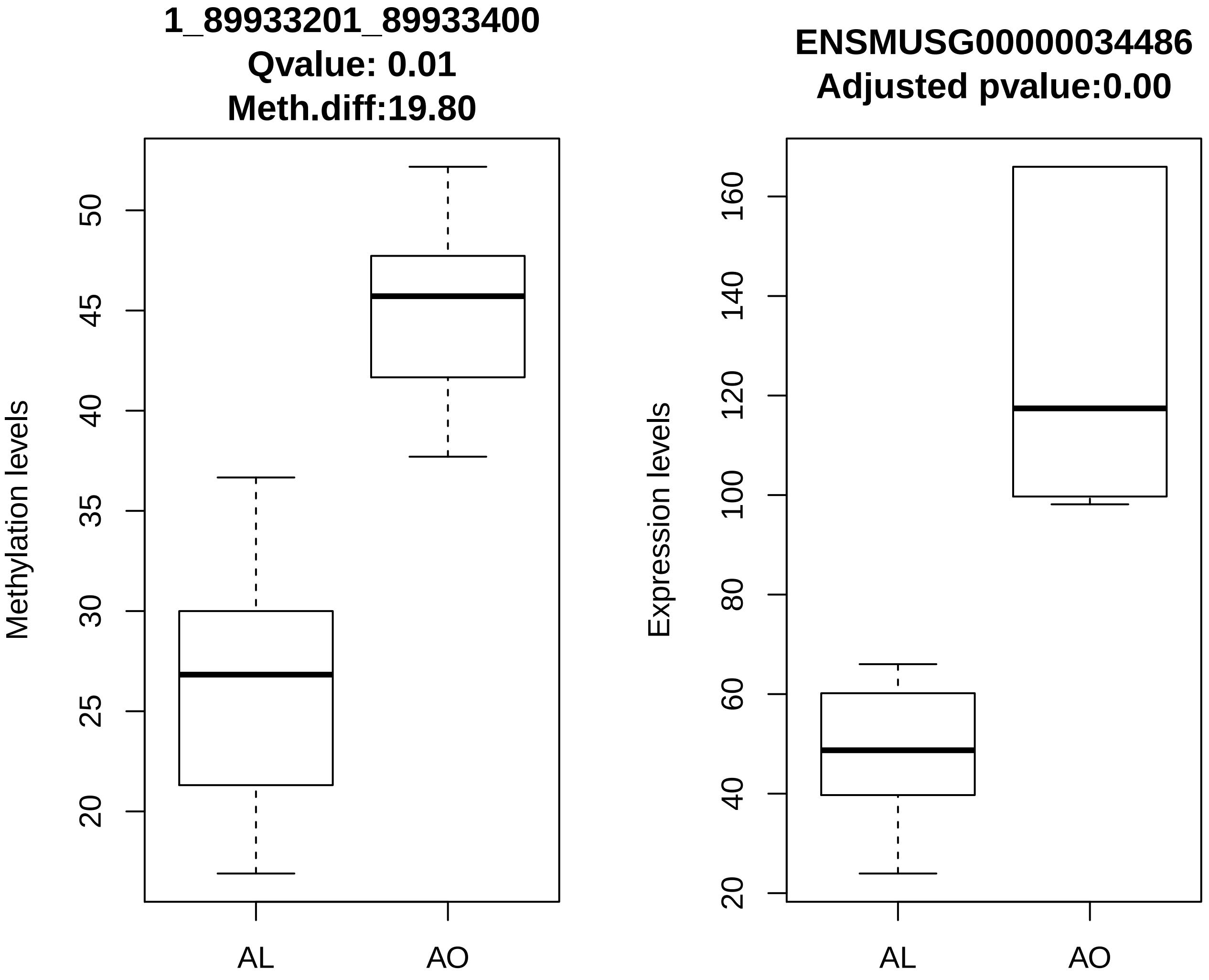
1_89933201_89933400
Qvalue: 0.01
Meth.diff:19.80
Methylation levels
20
25
30
35
40
45
50
AL
AO
ENSMUSG00000034486
Adjusted pvalue:0.00
Expression levels
20
40
60
80
100
120
140
160
AL
AO

Gbx2

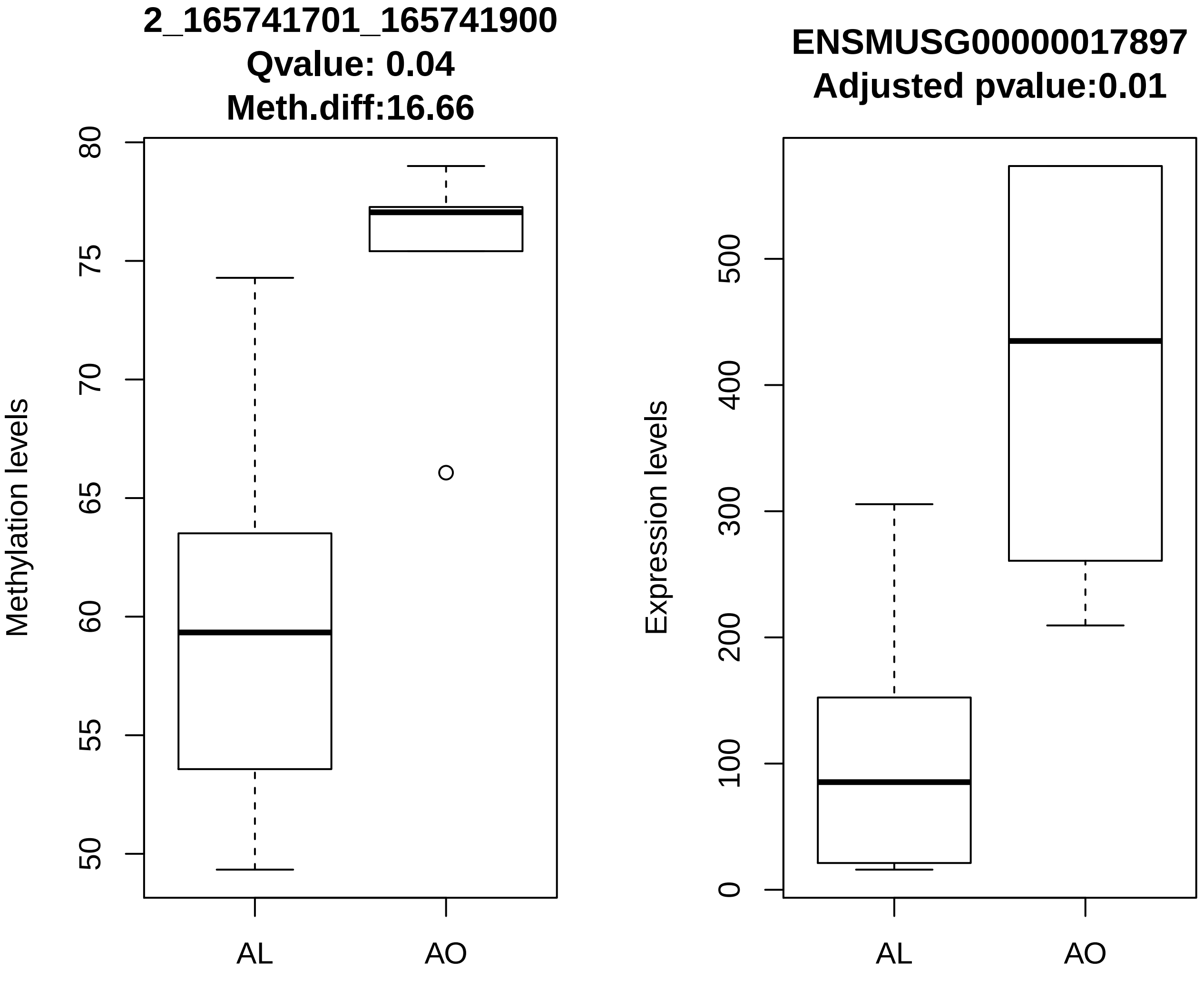
2_165741701_165741900
Qvalue: 0.04
Meth.diff:16.66
Methylation levels
80
75
70
65
60
55
50
AL
AO
ENSMUSG00000017897
Adjusted pvalue:0.01
Expression levels
500
400
300
200
100
0
AL
AO

Eya2

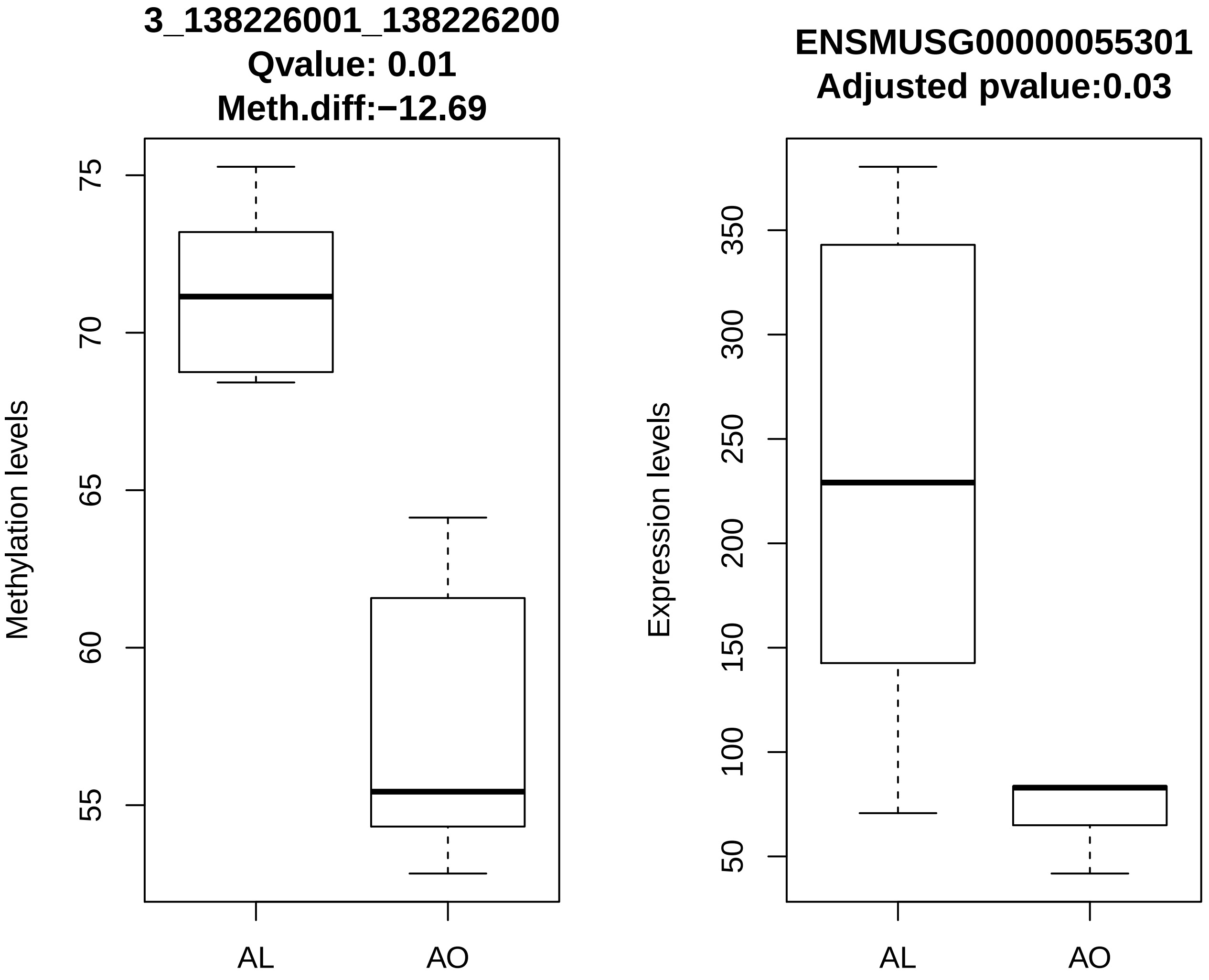
3_138226001_138226200
Qvalue: 0.01
Meth.diff:−12.69
Methylation levels
75
70
65
60
55
AL
AO
ENSMUSG00000055301
Adjusted pvalue:0.03
Expression levels
350
300
250
200
150
100
50
AL
AO

Adh7

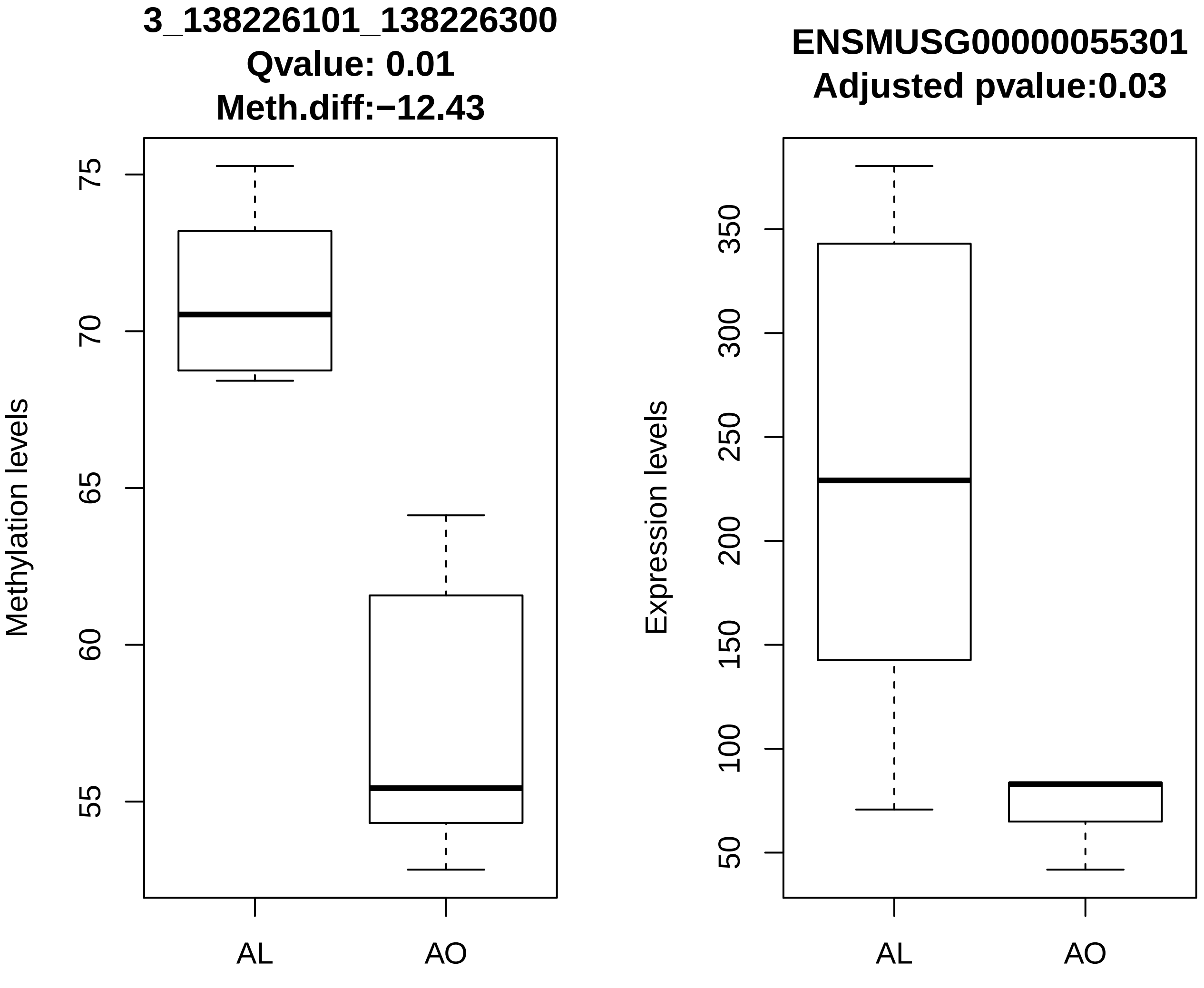
3_138226101_138226300
Qvalue: 0.01
Meth.diff:−12.43
Methylation levels
75
70
65
60
55
AL
AO
ENSMUSG00000055301
Adjusted pvalue:0.03
Expression levels
350
300
250
200
150
100
50
AL
AO

Adh7

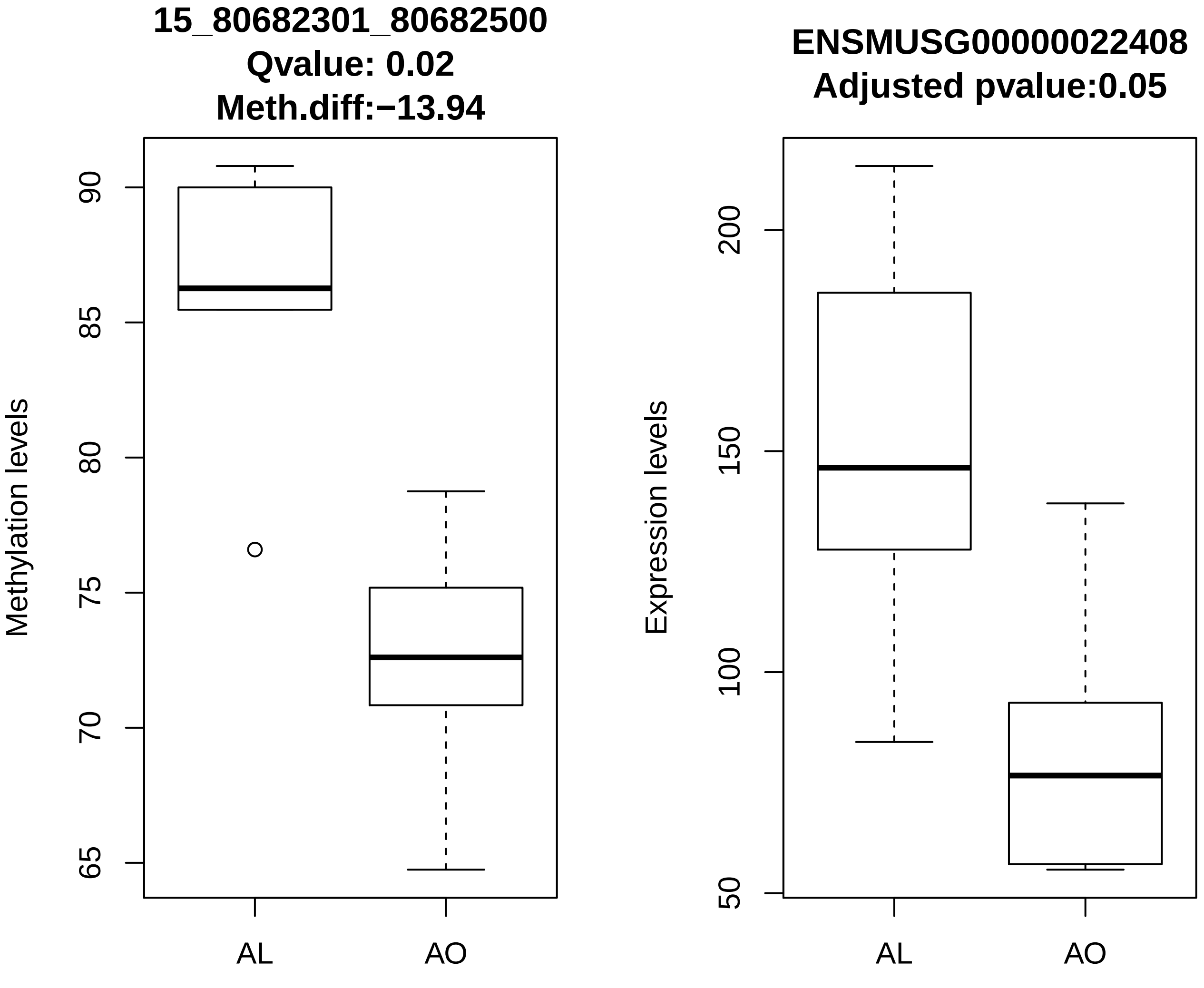
15_80682301_80682500
Qvalue: 0.02
Meth.diff:−13.94
Methylation levels
90
85
80
75
70
65
AL
AO
ENSMUSG00000022408
Adjusted pvalue:0.05
Expression levels
200
150
100
50
AL
AO

Fam83f

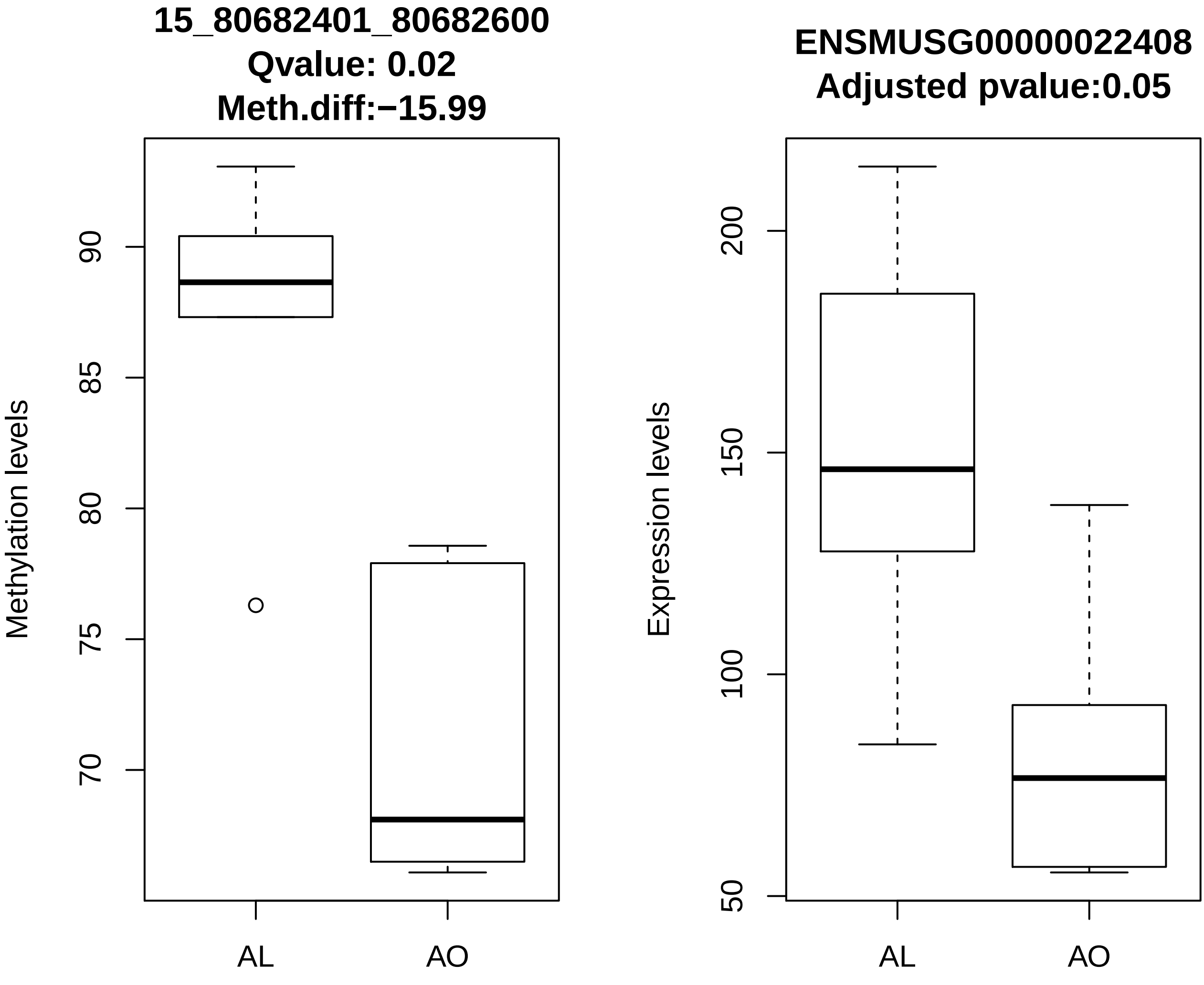
15_80682401_80682600
Qvalue: 0.02
Meth.diff:−15.99
Methylation levels
90
85
80
75
70
AL
AO
ENSMUSG00000022408
Adjusted pvalue:0.05
Expression levels
200
150
100
50
AL
AO

Fam83f

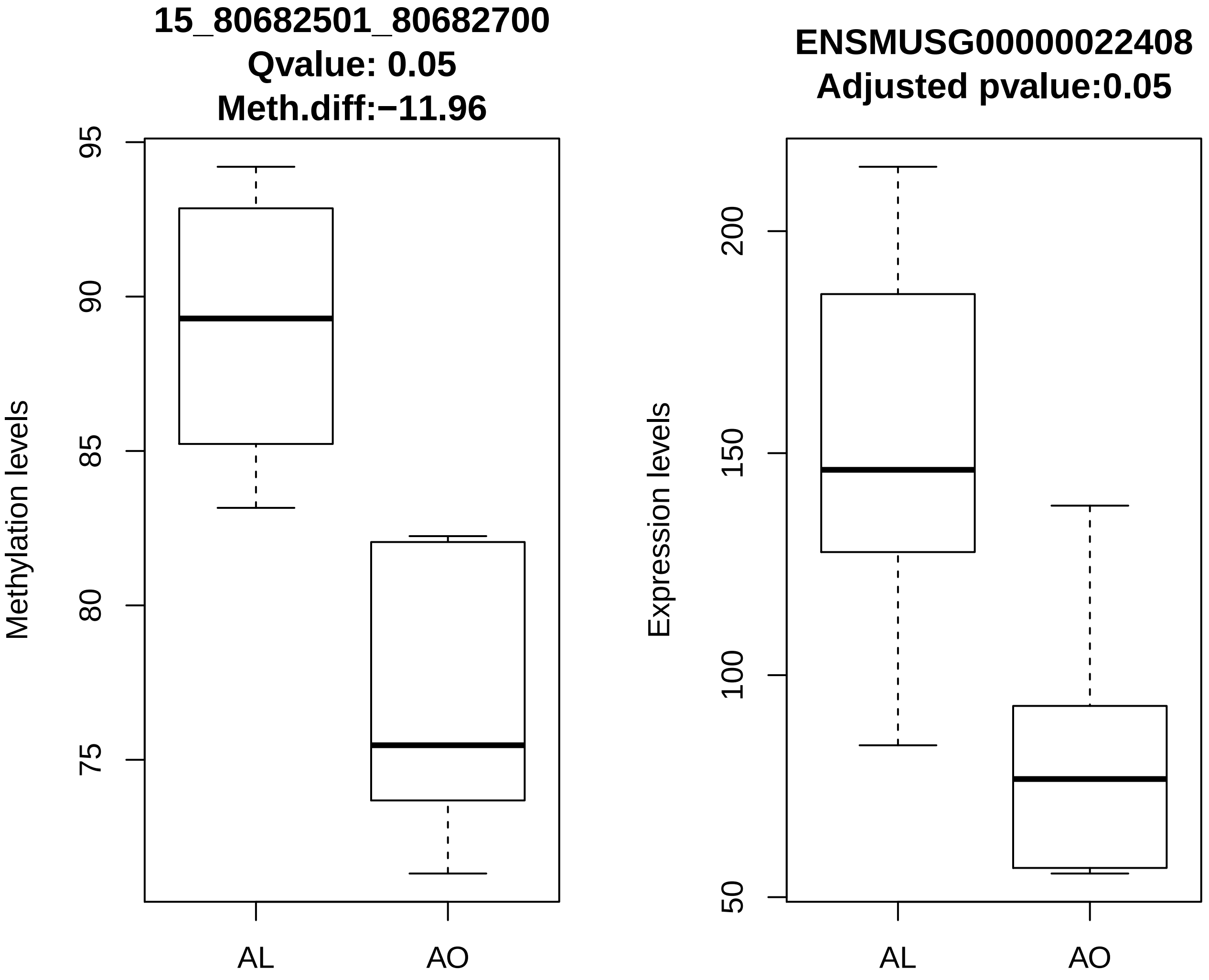
15_80682501_80682700
Qvalue: 0.05
Meth.diff:−11.96
Methylation levels
95
90
85
80
75
AL
AO
ENSMUSG00000022408
Adjusted pvalue:0.05
Expression levels
200
150
100
50
AL
AO

Fam83f

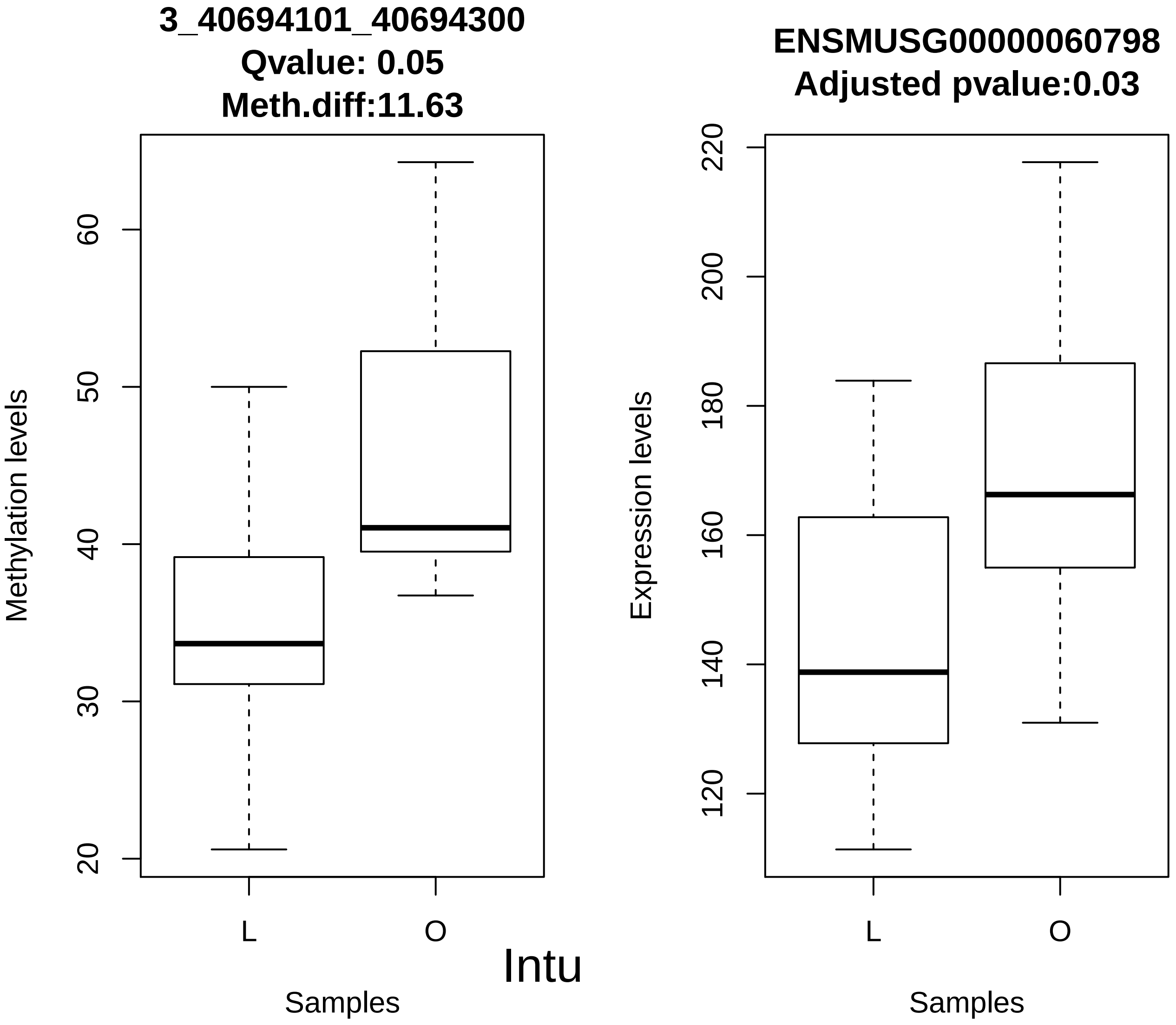
3_40694101_40694300
Qvalue: 0.05
Meth.diff:11.63
Methylation levels
20
30
40
50
60
L
O
Samples
ENSMUSG00000060798
Adjusted pvalue:0.03
Expression levels
120
140
160
180
200
220
L
O
Samples
Intu

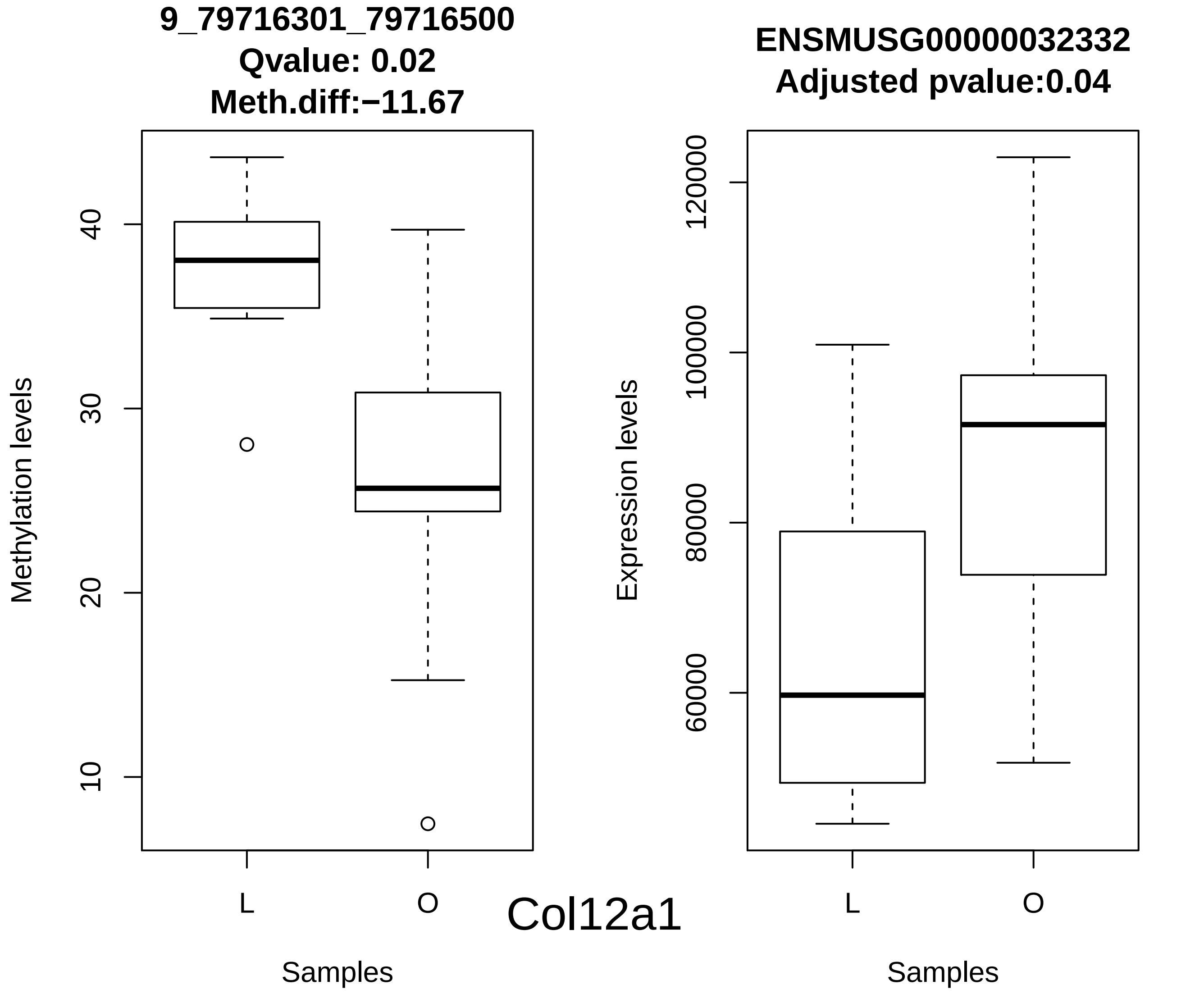
9_79716301_79716500
Qvalue: 0.02
Meth.diff:−11.67
Methylation levels
10
20
30
40
L
O
Samples
Col12a1
ENSMUSG00000032332
Adjusted pvalue:0.04
Expression levels
60000
80000
100000
120000
L
O
Samples

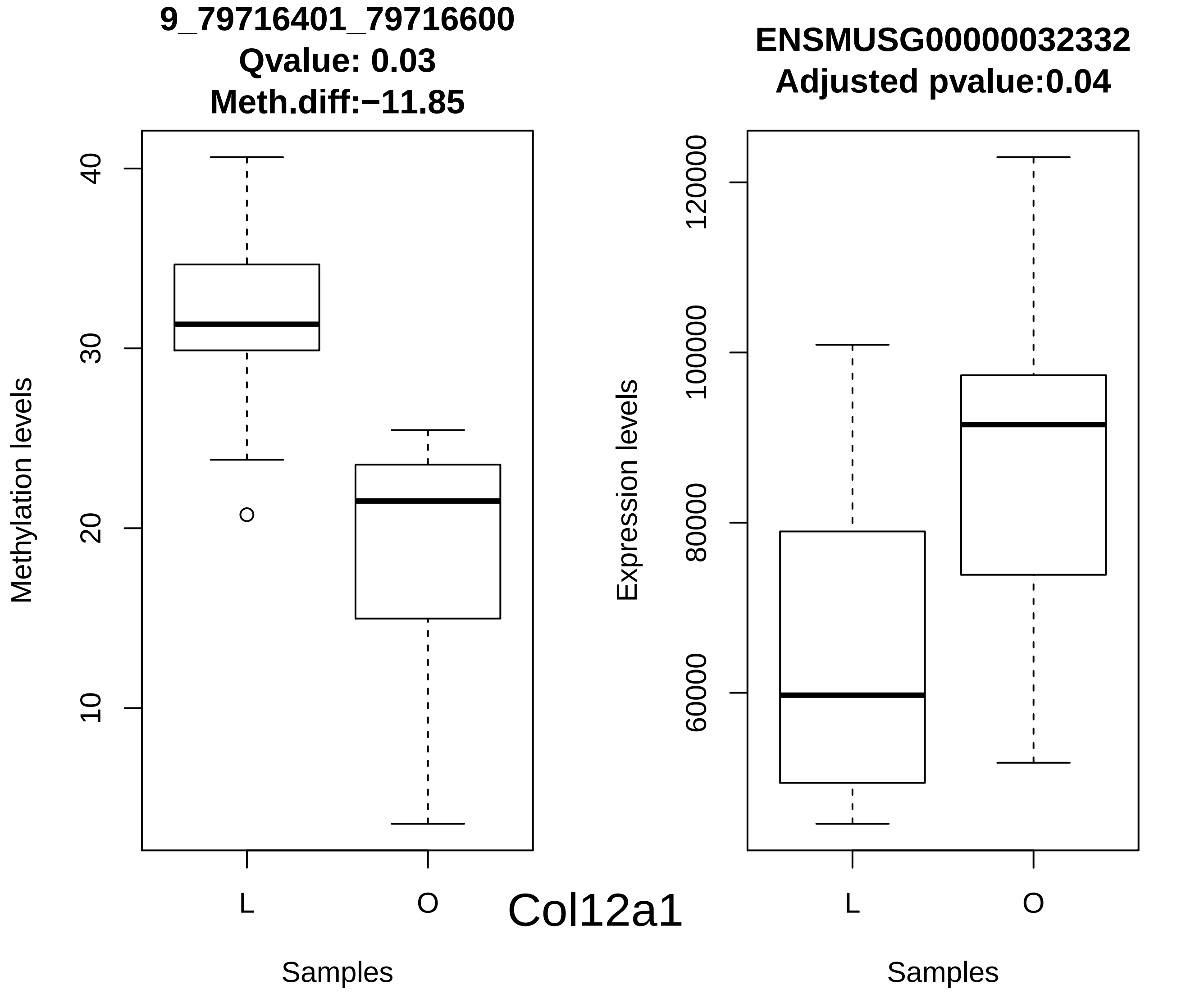
9_79716401_79716600
Qvalue: 0.03
Meth.diff:−11.85
ENSMUSG00000032332
Adjusted pvalue:0.04
Methylation levels
Expression levels
10
20
30
40
60000
80000
100000
120000
L
O
L
O
Col12a1
Samples
Samples

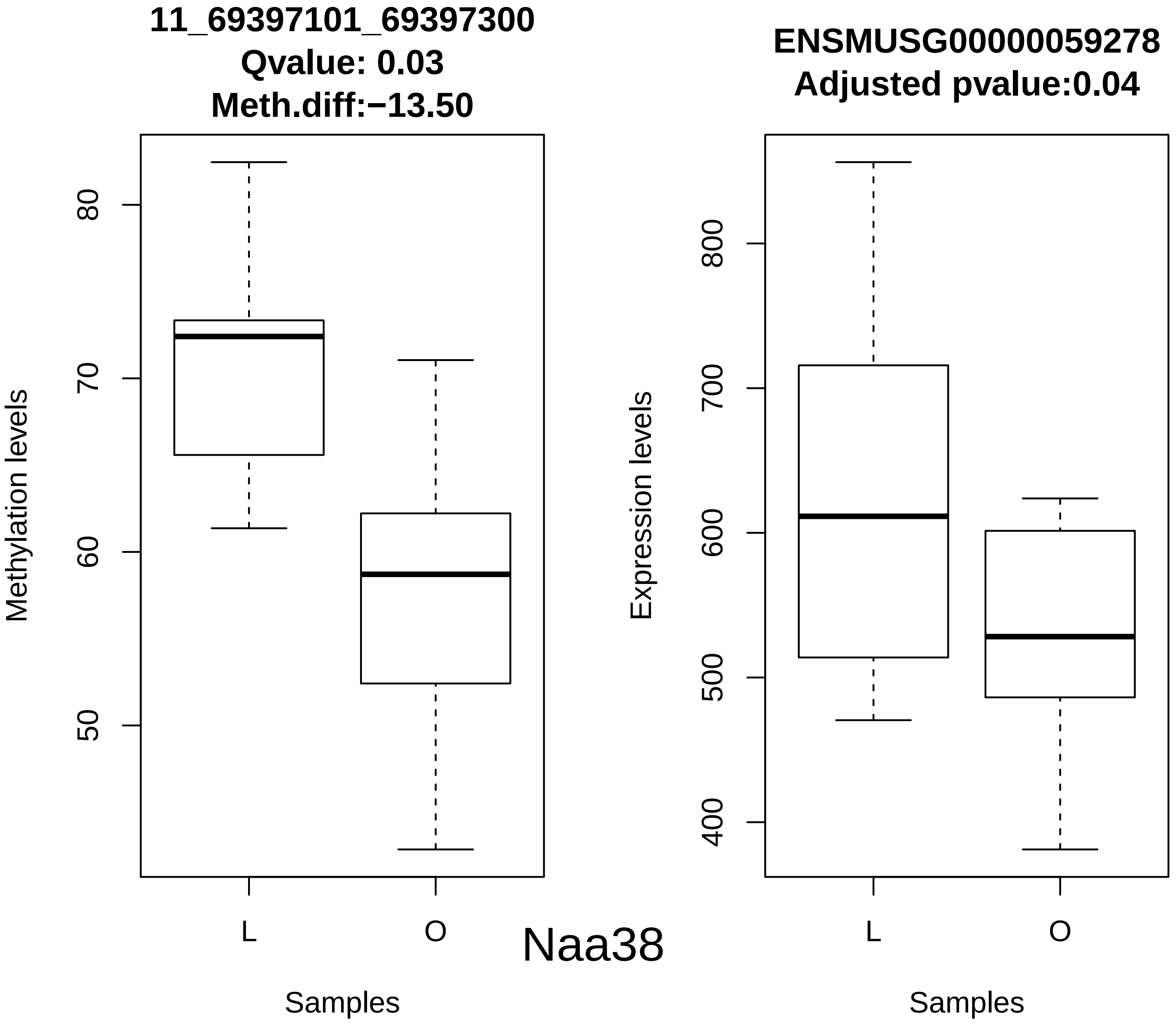
11_69397101_69397300
Qvalue: 0.03
Meth.diff:−13.50
Methylation levels
80
70
60
50
L
O
Samples
Naa38
ENSMUSG00000059278
Adjusted pvalue:0.04
Expression levels
800
700
600
500
400
L
O
Samples

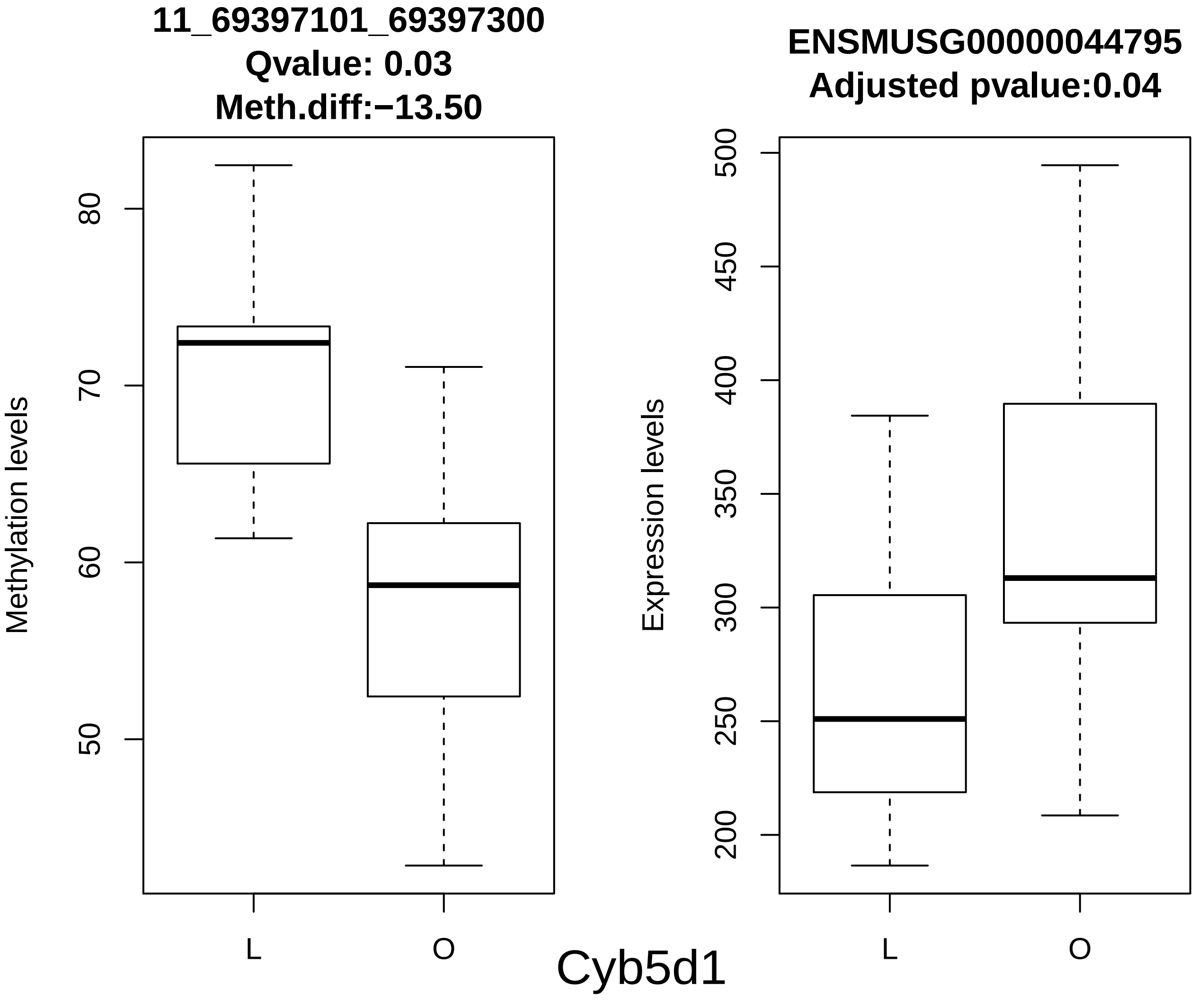
11_69397101_69397300
Qvalue: 0.03
Meth.diff:−13.50
Methylation levels
80
70
60
50
L
O
Samples
ENSMUSG00000044795
Adjusted pvalue:0.04
Expression levels
500
450
400
350
300
250
200
L
O
Samples
Cyb5d1

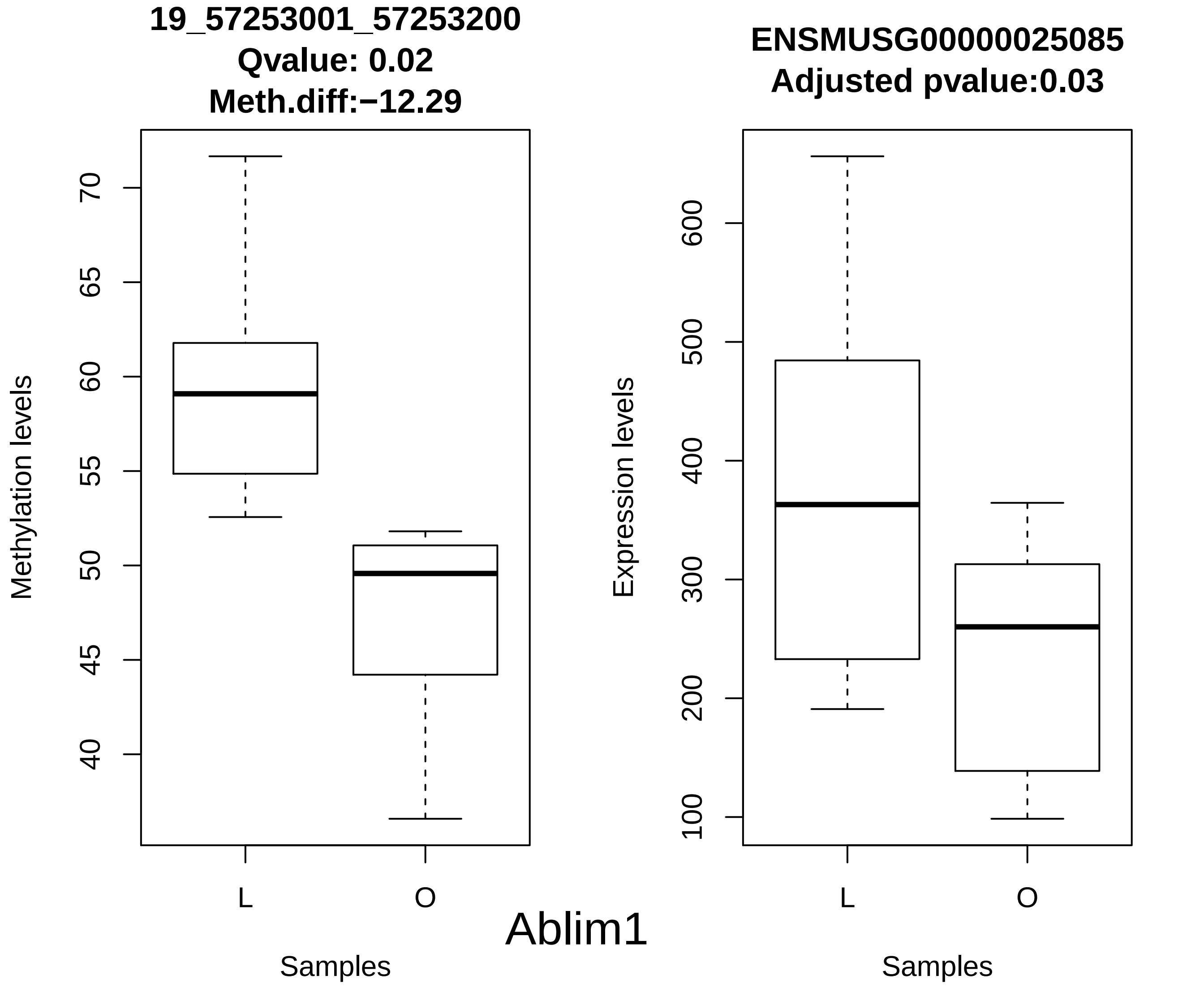
19_57253001_57253200
Qvalue: 0.02
Meth.diff:−12.29
Methylation levels
70
65
60
55
50
45
40
L
O
Samples
ENSMUSG00000025085
Adjusted pvalue:0.03
Expression levels
600
500
400
300
200
100
L
O
Samples
Ablim1
